# PI3K regulates TAZ/YAP and mTORC1 axes that can be synergistically targeted

**DOI:** 10.1101/2025.01.21.634138

**Authors:** Keith C. Garcia, Ali A. Khan, Krishnendu Ghosh, Souradip Sinha, Nicholas Scalora, Gillian DeWane, Colleen Fullenkamp, Nicole Merritt, Yuliia Drebot, Samuel Yu, Mariah Leidinger, Michael D. Henry, Patrick Breheny, Michael S. Chimenti, Munir R. Tanas

## Abstract

**Purpose:** Sarcomas are a heterogeneous group of cancers with few shared therapeutic targets. PI3K signaling is activated in various subsets of sarcomas, representing a shared oncogenic signaling pathway. Oncogenic PI3K signaling has been challenging to target therapeutically. An integrated view of PI3K and Hippo pathway signaling is examined to determine if this could be leveraged therapeutically.

**Experimental design:** A tissue microarray containing sarcomas of various histological types was evaluated for PTEN loss and correlated with levels of activated TAZ and YAP. PI3K and Hippo pathways were dissected in sarcoma cell lines. The role of TAZ and YAP were evaluated in a PI3K-driven mouse model. The efficacy of mTORC1 inhibition and TEAD inhibition were evaluated in sarcoma cell lines and *in vivo*.

**Results:** PI3K signaling is frequently activated in sarcomas due to PTEN loss (in 30-60%), representing a common therapeutic target. TAZ and YAP are transcriptional co-activators regulated by PI3K and drive a transcriptome necessary for tumor growth in a PI3K-driven sarcoma mouse model. Combination therapy using IK-930 (TEAD inhibitor) and everolimus (mTORC1 inhibitor) synergistically diminished proliferation and anchorage independent growth of PI3K-activated sarcoma cell lines at low, physiologically achievable doses. Furthermore, this combination therapy showed a synergistic effect *in vivo*, reducing tumor proliferation and size.

**Conclusions:** TAZ and YAP are transcriptional co-activators downstream of PI3K signaling, a pathway that has lacked a well-defined oncogenic transcription factor. This PI3K-TAZ/YAP axis exists in parallel to the known PI3K-Akt-mTORC1 axis allowing for synergistic combination therapy targeting the TAZ/YAP-TEAD interaction and mTORC1 in sarcomas.

## Introduction

Soft tissue sarcomas (STS) are a heterogenous group of cancers that consist of over 50 histological subtypes (1) making these cancers challenging to both diagnose and treat. Additional therapeutic targets active in different histological types of sarcoma are needed. Phosphotidylinositol-3 kinase (PI3K) signaling has been implicated in the pathogenesis of several different sarcoma types (2–4), but as a group, sarcomas have not been identified as a cancer type in which PI3K signaling plays a central role or can be successfully targeted. This is because in contrast to other cancers such as breast cancer or ovarian cancer, there is a scarcity of mutations in *PIK3CA, AKT1, AKT2, or AKT3* (5). Thus targeting PI3K signaling represents a potentially underdeveloped therapeutic approach in sarcomas.

There are three classes of PI3K enzymes of which the Class I enzymes are the best characterized (6). Class I PI3Ks are heterodimers consisting of a catalytic subunit (p110) and a regulatory subunit (p85). When activated by growth factor receptor tyrosine kinases, PI3K enzymes will phosphorylate P(4,5)P_2_ (PIP_2_) to generate P(3,4,5)P_3_ (PIP_3)_ at the cell membrane that results in a docking site for PDK1 and Akt via their individual pleckstrin homology (PH) domains. Upon phosphorylation of Akt by PDK1, Akt phosphorylates a number of different substrates including GSK-3β, p21, Bad, FOXO1, Mdm2, and Tsc2 that coordinately drives proliferation, inhibits apoptosis, and drives growth (7–14). By inactivating the Tsc2/Tsc1 complex, mTOR is activated as part of the mTORC1 complex also composed of the Raptor and GβL proteins. The mTORC1 complex subsequently phosphorylates and activates an S6K1-S6 axis and 4E-BP1, together which stimulate cap-dependent translation of proteins (15–18). A key negative regulator of the PI3K pathway is the phosphatase and tensin homolog (PTEN) tumor suppressor that dephosphorylates PIP_3_ converting it back to PIP_2_ at the membrane, thereby dampening PI3K signaling (19–21).

One underappreciated phenotype of PI3K signaling is the regulation of cell size (22), a phenotype that overlaps with that of the Hippo pathway (23). Indeed, several connections have been made between the PI3K signaling pathway and the Hippo pathway as it pertains to amino acid regulation (24–27), but also at a signal transduction level (28–33).

The Hippo pathway is a highly conserved pathway involved in regulating tissue/organ and cell size (34–39). This pathway consists of a core serine/threonine kinase cascade that consists of the STE20-like protein kinases 1 and 2 (MST1/2) and the <u>la</u>rge tumor <u>s</u>uppressor kinases 1 and 2 (LATS1/2) (34,35,37–42). The downstream effectors of the Hippo pathway are the transcriptional co-activators TAZ (gene name is *WWTR1*; <u>WW</u> domain-containing transcription regulator <u>1</u>) and YAP (gene name is *YAP1*; <u>Y</u>es-<u>a</u>ssociated <u>p</u>rotein <u>1</u>). TAZ and YAP have been implicated as oncoproteins in a number of different cancers including breast, colon, liver, lung, pancreas, thyroid cancers(43). More recently, TAZ and YAP have been shown to be frequently activated in sarcomas (44–56). Various external cues such as cell confluence or detachment can activate the core signaling cascade in the Hippo pathway leading to LATS1/2 mediated phosphorylation of TAZ and YAP at multiple serine residues that promote their accumulation into the cytoplasm where they are subsequently ubiquitinated and degraded by proteasome (38,39,57,58). Inactivation of the core serine/threonine kinase cassette results in the nuclear accumulation of TAZ and YAP where they bind to the TEA domain (TEAD) family of transcription factors to stimulate target gene expression (59,60). Upstream signaling pathways such as RAS-MEK-ERK and Wnt signaling (61,62) have been implicated to activate TAZ and YAP in various cancer types (63,64).

One notable absence in the known PI3K signaling cascade and an enigma in the field is the absence of downstream oncogenic transcription factors that could transduce signals from the cell membrane into the nucleus (FOXO1 is a transcription factor downstream of PI3K but has a tumor suppressive function) (65). Herein, we show using *in vitro* and *in vivo* approaches that TAZ and YAP are oncogenic transcriptional co-activators and effectors downstream of PI3K signaling, and that this finding can be leveraged to better therapeutically target PI3K signaling in sarcomas.

## Results

### Phosphotidylinositol-3 kinase is activated in sarcomas

To evaluate the mutational landscape of the PI3K signaling pathway, we analyzed publicly available datasets (The Cancer Genome Atlas) (66–68) for genetic alterations involving the pathway’s core components such as *PIK3CA* (p110α subunit)*, PIK3R1* (p85 subunit), *AKT1-3,* and *PDPK1* (PDK1). We observed that 5% or less of sarcomas exhibit gain-of-function genetic alterations for each of these members of the PI3K signaling pathway (**Figure 1A**). Scattered mutations in PTEN have been identified in various cancers, including the loss of function R130Q mutation in cancers including undifferentiated pleomorphic sarcoma (69–71) (**Figure S1A**). Further analysis of data from the TCGA showed alterations of *PTEN* such as deletions and truncations (**Figure 1B**). To determine if PTEN was lost at the protein level in clinical samples, we performed immunohistochemistry on a tissue microarray composed of 144 untreated sarcomas encompassing 18 histological types (**Figure 1C-D**). H-scores were calculated (intensity x % positive cells) to quantitate PTEN expression. Sarcomas demonstrated a spectrum of PTEN loss including those with complete loss of expression (H-score of 0) and samples with very low expression (H-score of ≤ 50). Approximately 1/3 (32%) of sarcomas showed a complete loss of PTEN expression consistent with the 20-30% loss that has been reported in the literature (2). However, approximately 2/3 of sarcomas (63%) of sarcomas showed complete loss to very low expression of PTEN (H-score of ≤ 50) across different histological types, suggesting that PI3K may be activated in a higher percentage of sarcomas than previously appreciated.

**Figure 1:**
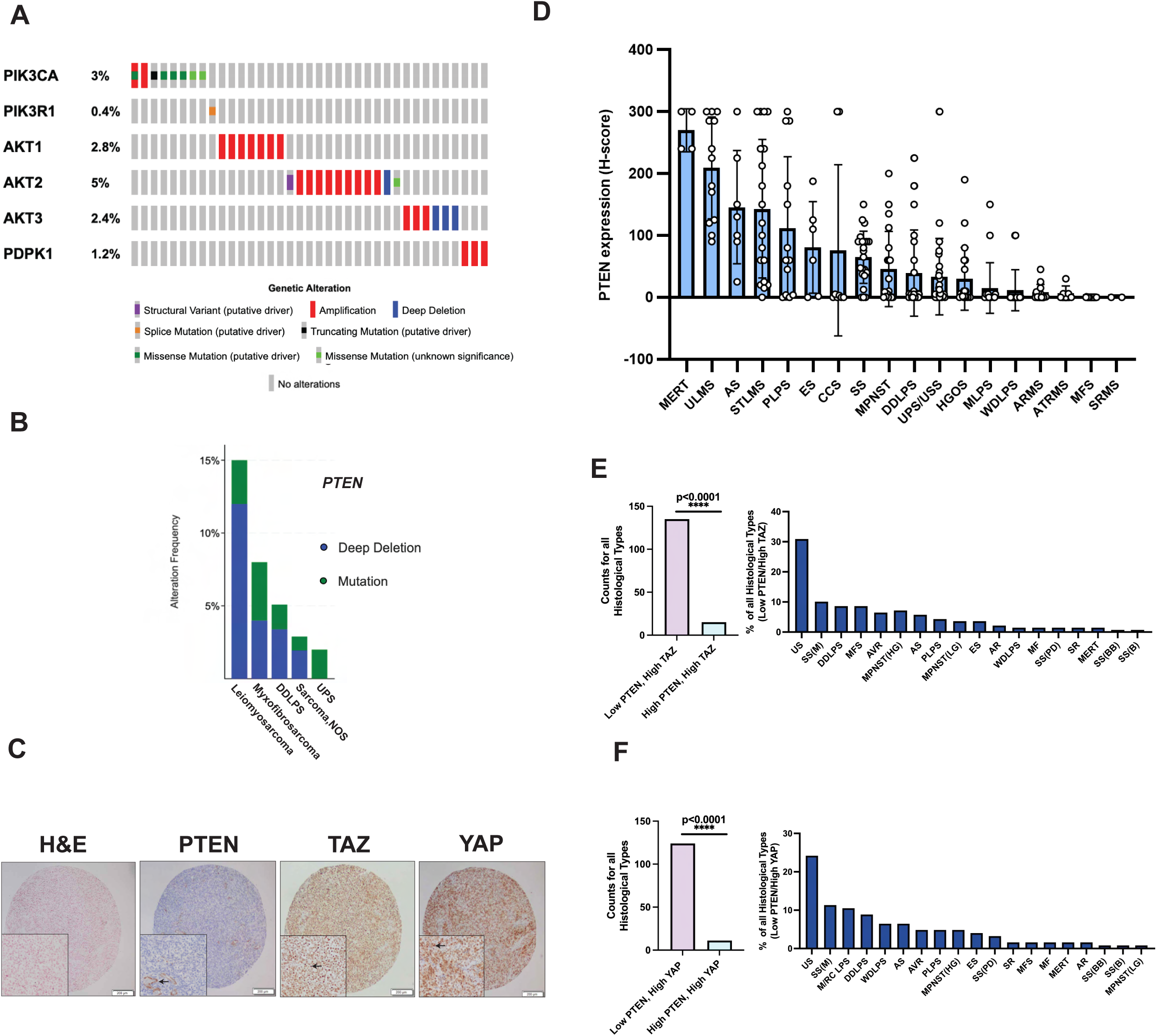
PI3K is widely activated in sarcomas. **(A)** The Cancer Genome Atlas (TCGA) data for PI3K signaling alterations in sarcomas (n=253) demonstrating the incidence of mutations in the core kinases of the PI3K signaling pathway. Data from TCGA was analyzed using the cBioportal online software. (**B**) TCGA data demonstrating the frequency of genetic alterations of *PTEN* in sarcomas (n=253). Data from TCGA was analyzed using the cBioportal online software. **C**) H&E and immunohistochemistry performed for PTEN, TAZ, and YAP on adult-type rhabdomyosarcoma from tissue microarray. Inset for PTEN shows vascular positive control. Insets for activated (nuclear) TAZ and YAP. (**D**) H-scores from clinical tissue microarray panel of various sarcoma histological subtypes. MERT– malignant extrarenal rhabdoid tumor; ULMS–uterine leiomyosarcoma; AS–angiosarcoma; STLMS–soft tissue leiomyosarcoma; PLPS–pleomorphic liposarcoma; ES–epithelioid sarcoma; CCS–clear cell sarcoma of soft tissue; SS–synovial sarcoma; MPNST–malignant peripheral nerve sheath tumor; DDLPS–dedifferentiated liposarcoma; UPS/USS–undifferentiated pleomorphic sarcoma/undifferentiated spindle cell sarcoma; HGOS–high grade osteosarcoma; MLPS–myxoid liposarcoma; WDLPS–well differentiated liposarcoma; ARMS–alveolar rhabdomyosarcoma; ATRMS–adult-type rhabdomyosarcoma; MFS–myxofibrosarcoma; SRMS– sclerosing rhabdomyosarcoma. Histology, H&E staining, and immunohistochemistry (IHC) for PTEN, TAZ, and YAP (nuclear staining) in a sample of ARMS (bottom four panels). (**E)** Bar graph (left) showing sarcomas exhibiting Low PTEN/High TAZ expression profile (n=135) relative to High PTEN/High TAZ expression profile (n=35). Bar graph (fight) showing composition of sarcomas exhibiting Low PTEN/High TAZ profile according to histological type. (**F**) Bar graph (left) showing sarcomas exhibiting Low PTEN/High YAP expression profile (n=124) relative to High PTEN/High YAP expression profile (n=11). Bar graph (right) showing composition of sarcomas exhibiting Low PTEN/High TAZ profile according to histological type. Two-tailed p-values for IHC counts were evaluated using the chi-square test with one degree of freedom. For all panels, ** p<0.01, ***p<0.001.

To determine if PTEN expression and thus PI3K activation correlates with TAZ and YAP expression and activation (nuclear localization), H-scores for TAZ and YAP from a previously published data set (46) were plotted as a function of H-score for PTEN. The resulting scatter plot for TAZ (**Figure S1B**) was divided into 4 quadrants/categories: TAZ High/PTEN Low, TAZ High/PTEN High, TAZ Low/PTEN Low, and TAZ Low/PTEN High. The scatter plot for YAP was similarly divided (**Figure S1C**). The cut-off for low expression of TAZ, YAP, and PTEN was established at an H-score of ≤ 150 (the median H score possible); H-scores of > 150 were considered high. Tumors with high TAZ expression/activation were more likely to demonstrate low PTEN expression than high PTEN expression (p<0.0001) (**Figure 1E**). Similarly, tumors with high YAP expression/activation were more likely to demonstrate low PTEN expression than high PTEN expression (p<0.0001) (**Figure 1F**). The sarcoma histological type with the highest percentage of low PTEN/high TAZ tumors and low PTEN/high YAP tumors was undifferentiated sarcoma (composed of undifferentiated pleomorphic sarcoma and undifferentiated spindle cell sarcoma). Other histological types notable for having a relatively high percentage of low PTEN/high TAZ/YAP tumors include monophasic synovial sarcomas, well-differentiated/dedifferentiated liposarcoma, myxoid/round cell liposarcoma, malignant peripheral nerve sheath tumor, myxofibrosarcoma, and alveolar rhabdomyosarcoma (**Figure 1E, F**).

We subsequently assayed 8 sarcoma cell lines (**Figure S1D**) to examine hyperactive PI3K signaling by western blot after serum starvation. Phosphorylated AKT (S473) levels were elevated for SJCRH30 (alveolar rhabdomyosarcoma) and RD (embryonal rhabdomyosarcoma) cell lines compared to the primary skeletal muscle cell control (**Figure S1D**). Phosphorylated AKT (S473) was also elevated for the A204 (malignant extrarenal rhabdoid tumor) and SW872 (liposarcoma) cell lines compared to primary cell controls that were available (primary smooth muscle and skeletal muscle cells). Consistent with findings from the above mentioned clinical samples (**Figure 1C, D**), 3 of the 4 cell lines (75%) mentioned above demonstrated elevated phospho-Akt levels and were simultaneously characterized by reduced PTEN expression. To determine whether these cell lines were dependent on PI3K for cell survival and thus driven by PI3K, clonogenic outgrowth assays were performed on the SJCRH30 (PTEN lost) and A204 (PTEN intact) sarcoma cell lines. Treatment with the pan class I PI3K inhibitors, GDC-0941 (Pictilisib) and LY294002 (**Figure S1E-H**), resulted in a significant decrease in cell survival in a concentration dependent manner. Even so, inhibition did not entirely prevent clonogenic outgrowth even at higher concentrations, providing an impetus to better target PI3K signaling in these cell lines and sarcomas overall.

### TAZ/YAP are transcriptional end effectors regulated by PI3K

Observations from analysis of the above clinical samples (**Figure 1E, F)** led to the hypothesis that PTEN loss leads to activation PI3K signaling, which subsequently activates TAZ and YAP in sarcomas. To determine if TAZ and YAP were oncoproteins in the PI3K-activated SJCRH30 and A204 cell lines (**Figure S1D**), we performed shRNA mediated knockdown of TAZ or YAP in these cell lines which resulted in reduced anchorage independent growth (**Figure S1I, J**). These results demonstrated that SJCRH30 and A204 were two PI3K-activated sarcoma cell lines in which TAZ and YAP functioned as oncoproteins. To determine if TAZ and YAP are regulated by PI3K signaling, SJCRH30 and A204 cells were treated with 30 µM of the pan class I PI3K inhibitor, LY294002 (LY) over a 12 hour time course. For both the SJCRH30 and A204 cell lines, we saw significant decreases in total TAZ levels beginning at 1 hour of treatment (**Figure 2A**). These observations are consistent with the observable half-life of TAZ (∼2-6 hours) depending on the cellular context (72). Total YAP protein expression levels remained essentially unchanged in both SJCHR30 and A204 cells (**Figure 2A**) with 30 µM of LY.

**Figure 2:**
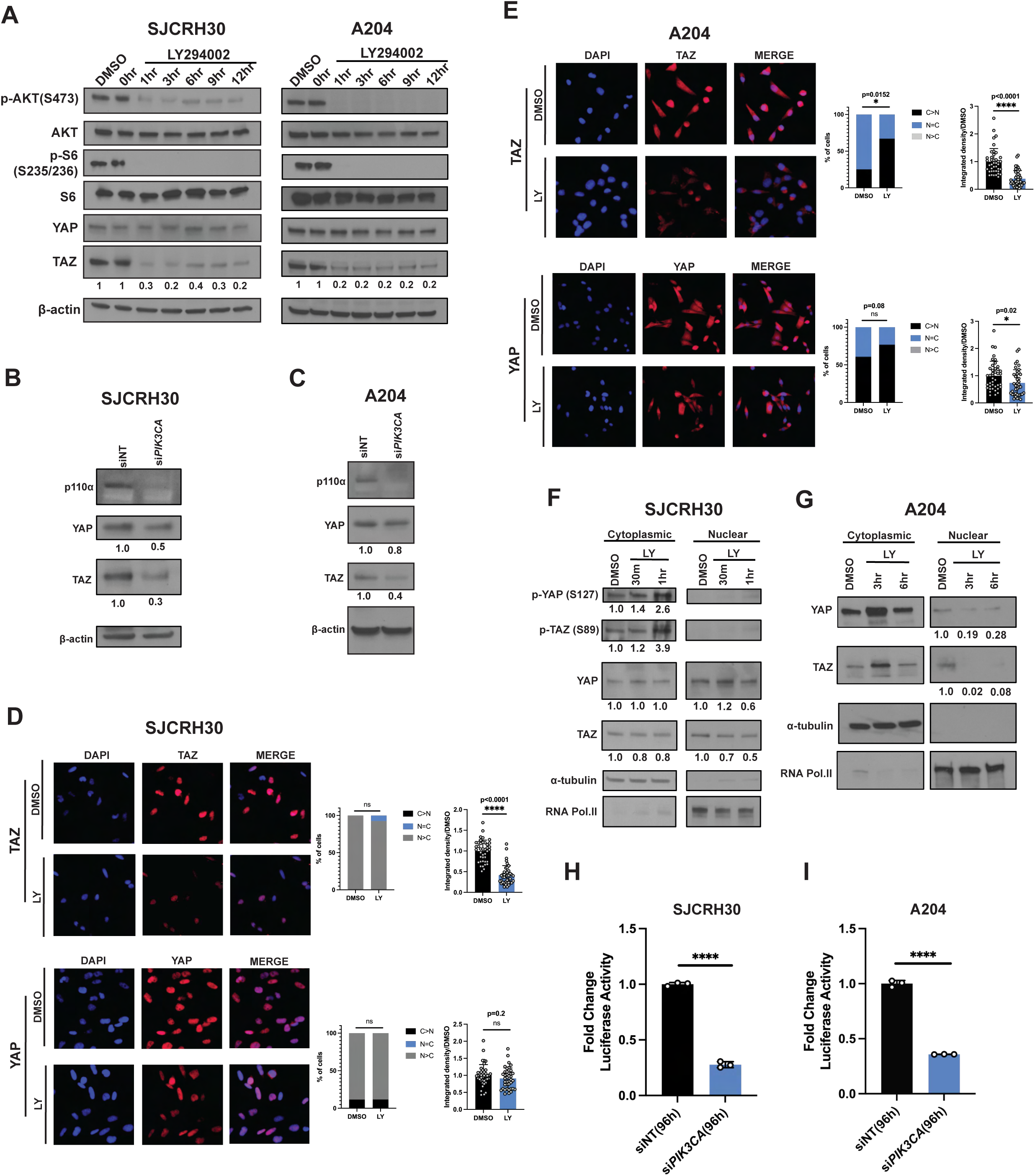
PI3K inhibition alters the stability and localization of TAZ/YAP in SJCRH30 and A204 cells. **(A)** Western blot of SJCRH30 and A204 cells that were treated with 30 µM LY294002 for the indicated timepoints. Total YAP and TAZ protein levels were normalized to β-actin and compared relative to vehicle control (DMSO). (**B, C**) Western blot analysis of TAZ/YAP upon siRNA (pooled) mediated knock down of *PIK3CA* (p110a) in SJCRH30 (**B**) and A204 (**C**) cells. Quantitation for TAZ and YAP protein levels were normalized to β-actin and compared to siNT (non-targeting) control. β-actin was used as the loading control. (**D, E**) Immunofluorescence analysis of SJCRH30 (**D**) or A204 (**E**) cells that were treated with 30 µM LY2094002 for 3hrs and 6 hrs, respectively. % of cells localized in the cytosol vs. nucleus (left graph) and integrated density of fluorescent signal (right graph) was analyzed for TAZ (bottom) and YAP. C>N (cytosolic>nuclear), N=C (nuclear=cytosolic), and N>C (nuclear>cytosolic). % localization and integrated density analysis were compared relative to DMSO treated cells (see Methods section for details). (**F, G**) Nuclear and cytoplasmic fractionation of SJCRH30 cells (**F**) and A204 cells (**G**) treated with vehicle (DMSO) or 60 µM LY294002. p-TAZ and p-YAP quantitation is normalized to cytoplasmic total TAZ and YAP, respectively, and compared relative to DMSO treated cells. Cytoplasmic and nuclear total TAZ/YAP protein levels are normalized to 𝛼-tubulin and RNA polymerase II, respectively, and compared relative to DMSO controls. 𝛼-tubulin is the cytoplasmic loading control and RNA polymerase II is the nuclear loading control. Luciferase activity measured upon siRNA (pooled) mediated knockdown of *PIK3CA* for 96 hours in (**H**) SJCRH30 cells and (**I**) A204 cells stably expressing a TEAD luciferase reporter. Fold change was analyzed for si*PIK3CA* relative to siNT (non-targeting) control. Statistical analysis was performed using student’s unpaired two-tailed *t*-test. Each experiment was repeated at least twice. Error bars were used to define one standard deviation. For all panels, ****p<0.0001, *** p<0.001, ** p<0.01, *p<0.05, ns (not statistically significant). Each experiment was repeated at least twice.

To further determine if PI3K regulates TAZ/YAP in sarcoma cells, we performed siRNA mediated knockdown of *PIK3CA*, the catalytic subunit of the PI3K heterodimer, that resulted in a significant decrease in total TAZ protein levels, and to a lesser extent, YAP, in both SJCRH30 and A204 cell lines (**Figure 2B, C**). YAP is hypothesized to be more stable than TAZ due to additional serine residues involved in its protein stability (39,57,58). To test the hypothesis that PI3K signaling was affecting the localization of YAP rather than total levels of YAP, we performed immunofluorescence (IF) analysis on SJCHR30 cells treated with 60 µM LY294002 (LY) (**Figure 2D**). Although the ratio of nuclear:cytoplasmic TAZ did not change after treatment with LY, total TAZ levels decreased after treatment with LY by integrated density (p<0.0001) (**Figure 2D**). No significant change in YAP localization was identified after treatment with 60µM LY. When A204 cells were treated with LY, localization of TAZ was shifted from the nucleus into the cytoplasm (**Figure 2E**). A similar trend towards cytoplasmic localization was noted for YAP, however this was not statistically significant. Total TAZ and YAP expression, as calculated by integrated density, was decreased in A204 cells after treatment with LY (**Figure 2E**).

Although a shift in the nuclear to cytoplasmic localization of YAP was not identified by IF in the SJCRH30 cells, nuclear and cytoplasmic fractionation demonstrated a decrease in total YAP in the nuclear compartment after treatment with 60 µM LY for 1 hour accompanied by a greater than 2-fold increase in phospho-YAP S127 in the cytoplasmic compartment (**Figure 2F**). In addition, nuclear and cytoplasmic fractionation showed that total TAZ protein levels decreased between 30 minutes and one hour, and a greater than 3-fold increase in phosphorylated TAZ (S89) was observed in the cytoplasmic fraction (**Figure 2F**). Consistent with the above findings, PI3K inhibition in A204 cells revealed that TAZ and YAP expression decreased in the nuclear fraction after treatment with LY relative to DMSO control (**Figure 2G**).

The above studies have shown that PI3K regulates the expression and localization of TAZ and YAP. To determine whether this inhibition of PI3K signaling was functionally significant and impacted the transcriptional activity of TAZ/YAP, SJCRH30 and A204 cells stably expressing an 8xTEAD luciferase reporter was subsequently transfected with siRNA targeting *PIK3CA* for 96 hours using a dual luciferase reporter assay with *Renilla* luciferase control (see Methods section for details). siRNA mediated knockdown of *PIK3CA* in the two cell lines caused a signification reduction in TEAD reporter activity; a 3.6 fold reduction (p < 0.0001) was noted for SJCRH30, and a 2.9 fold reduction (p < 0.0001) was seen in the A204 cell line (**Figure 2H, I**). These findings demonstrate that inhibition of PI3K signaling leads to reduced transcriptional activity of TAZ/YAP in sarcoma cells. Collectively, the above data indicates TAZ/YAP are transcriptional end effectors regulated by PI3K in sarcoma cell lines.

### PI3K inhibition results in LATS-mediated regulation of TAZ and YAP in sarcoma cells

To determine how PI3K may be regulating the expression and activity of TAZ and YAP, we examined the role of core enzymes downstream of PI3K. Work from others have suggested that PDK1 and AKT are key regulators of TAZ and YAP (31,33,73). To determine if PDK1 was a regulator of TAZ/YAP in these two cell lines, we performed siRNA-mediated silencing of *PDK1* and found that it did not affect total TAZ or YAP expression levels (**Figure S2A**). To determine if AKT is a potential upstream regulator of TAZ and YAP in SJCRH30 and A204 cells we performed siRNA mediated knockdown of *AKT1* (**Figure S2B**) and treatment with MK2206, an allosteric pan-AKT inhibitor, (**Figure S2C, D**). Pharmacological and genetic inhibition of AKT had no effect on the expression of TAZ and YAP protein levels in both cell lines (**Figure S2B-D**). The above data indicate PDK1 and AKT are not regulators of TAZ and YAP expression in these PI3K-activated sarcoma cell lines.

There is some evidence suggesting that GSK3β may regulate TAZ by phosphorylating serine residues 58 and 62 that are part of the N-terminal phosphodegron of TAZ (74) (**Figure S3A**). To determine if GSK3β plays a role in TAZ stability subsequent to PI3K inhibition, SJCRH30 and A204 cells were engineered to stably express wild-type Flag-TAZ or the Flag-TAZ S58/62A GSK3β resistant mutant (**Figure S3A, B**). When treated with LY294002, Flag-TAZ and Flag-TAZ S58/62A both decreased over the course of one hour in SJCRH30 cells suggesting that TAZ is regulated by PI3K in a GSK3β−independent manner (**Figure S3C, D**). Similarly, Flag-TAZ and Flag-TAZ S58/62A in A204 cells decreased over time when treated with LY294002 (**Figure S3E, F**). Taken together, the above findings indicate that PI3K is regulating TAZ and YAP by a mechanism that does not include PDK1, AKT, or GSK3β.

To gain further insight into how TAZ and YAP are regulated by PI3K, we evaluated whole cell lysates of SJCRH30 after treatment of LY by western blot (**Figure 3A**). Consistent with previous observations (**Figure 2F**), 30 minutes to one hour post treatment of LY demonstrated elevated phosphorylation levels of TAZ (S89) and YAP (S127), target sites for the LATS1/2 kinases, relative to DMSO (vehicle) treated cells (**Figure 3A**). In addition, phospho-LATS1 (S909) was elevated after treatment with LY, consistent with activation of LATS1 (75,76). Quantitative immunoblot analysis showed that total TAZ protein levels decreased approximately 2-fold relative to vehicle control around three hours post treatment (**Figure 3A**). No considerable change was observed for total YAP protein levels in whole cell lysates, (**Figure 3A**), although this does not take into account the shift in nuclear and cytoplasmic localization seen in **Figure 2F**. To address potential off target effects with LY294002, SJCRH30 and A204 cells were treated with a second pan class I PI3K inhibitor, GDC-0941 (10uM) which displayed similar results (**Figure S4A, B**). To test the hypothesis that PI3K inhibition results in LATS1/2 mediated phosphorylation of TAZ and YAP, we generated and expressed a LATS1/2-insensitive form of TAZ, Flag-TAZ4SA, in both SJCRH30 and A204 cells (**Figure 3B, C and S4C, D**)(77). Treatment of LY294002 in A204 Flag-TAZ cells exhibited an observable decrease in Flag-TAZ protein levels (**Figure 3B**). In contrast, LY294002 did not diminish expression of Flag-TAZ4SA in A204 cells (**Figure 3C**), consistent with the hypothesis that PI3K dependent regulation of TAZ is mediated via LATS1/2. Similar findings were observed in the SJCRH30 Flag-TAZ and Flag-TAZ4SA cell lines when treated with LY (**Figure S4C, D**). Finally, we sought to determine if depletion of TAZ levels upon PI3K inhibition was due to proteasomal degradation secondary to phosphorylation by LATS1/2. A204 cells simultaneously treated with LY294002 and 10µM of MG-132 showed that MG-132 rescued proteasomal degradation of TAZ promoted by LY294002 (**Figure 3D**). Similar results were found in the SJCRH30 cell line (**Figure 3E**). The above findings support a working model that inhibition of PI3K signaling in sarcoma cells results in LATS1/2-mediated phosphorylation of TAZ and YAP to promote their destabilization and proteasomal degradation (**Figure 3F**).

**Figure 3:**
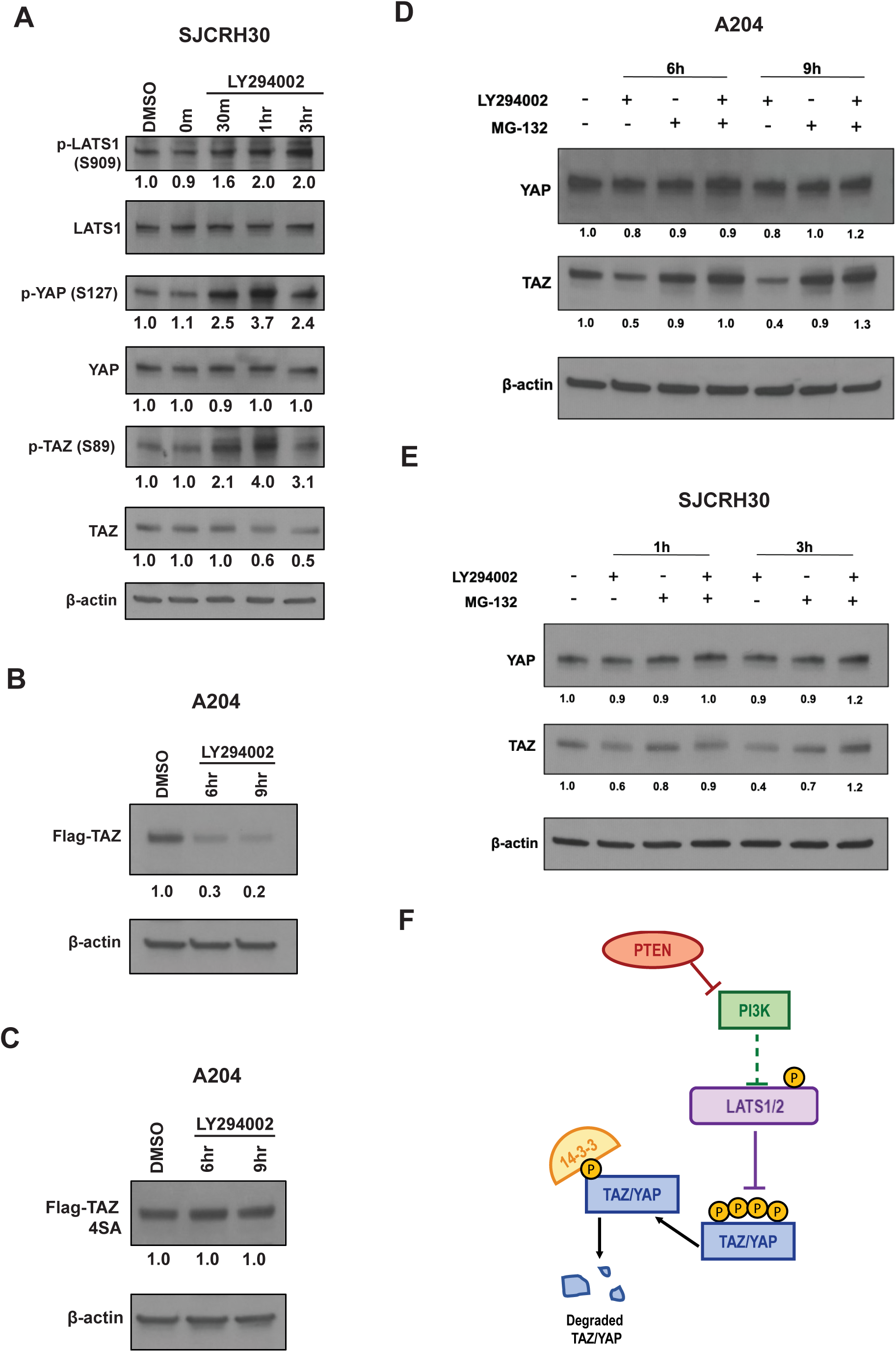
PI3K inhibition promotes LATS-mediated degradation of TAZ/YAP in SJCRH30 and A204 cells. (**A**) Western blot of SJCRH30 cells that were treated with 60 µM LY294002 for the indicated timepoints. p-YAP (S127), and p-TAZ (S89) were quantitated relative to total YAP and TAZ, respectively, and compared to vehicle (DMSO) treated cells. Total YAP and TAZ protein levels were normalized to β-actin and compared relative to vehicle control (DMSO). (**B**) Flag-TAZ or (**C**) Flag-TAZ 4SA were treated with 60 µM LY294002 for 6 and 9 hours. Flag-TAZ 4SA LATS-insensitive mutants are resistant to degradation when treated with LY294002. Quantitation of Flag-TAZ and TAZ 4SA protein levels were normalized to β-actin and compared to vehicle (DMSO) control. (**D**) A204 cells were treated with or without LY294002 (60 µM) or MG132 (10 µM) as indicated for 6 and 9 hours. Quantitation of YAP and TAZ protein levels were normalized to β-actin and compared to vehicle (DMSO) control. (**E**) SJCRH30 cells were treated with or without LY294002 or MG132 as indicated for 1 and 3 hours. Quantitation of YAP and TAZ protein levels were normalized to β-actin and compared to vehicle (DMSO) control. (**F**) Working model illustrating crosstalk between PI3K and LATS1/2 in TAZ/YAP regulation. Each experiment was repeated at least twice.

### Taz and Yap are central oncoproteins in a *Trp53^fl/fl^Pten^fl/fl^* sarcoma mouse model

We previously showed that conditional knock out of *Trp53* and *Pten* by adenoviral mediated Cre recombinase (Ad CMV-Cre) expression injected intramuscularly resulted in the formation of predominantly undifferentiated pleomorphic sarcoma at the injection site (78). Tumor lysates in undifferentiated pleomorphic sarcomas derived from *Trp53^fl/fl^Pten^fl/fl^* mice showed activation of PI3K signaling as demonstrated by elevated phospho-Akt (Ser473) and phospho-S6 (Ser235/236) relative to skeletal muscle controls (**Figure 4A**). Consistent with the above *in vitro* studies, total TAZ and YAP in these tumor lysates were elevated relative to skeletal muscle controls. To determine whether TAZ and YAP were activated in this tumors, we performed immunohistochemistry on tumors from *Trp53^fl/fl^Pten^fl/fl^* mice. In agreement with the above findings by western blot, immunohistochemistry for TAZ and YAP demonstrated diffuse and strong expression and nuclear localization of the two transcriptional co-activators in essentially all of the tumors (**Figure 4B**).

**Figure 4:**
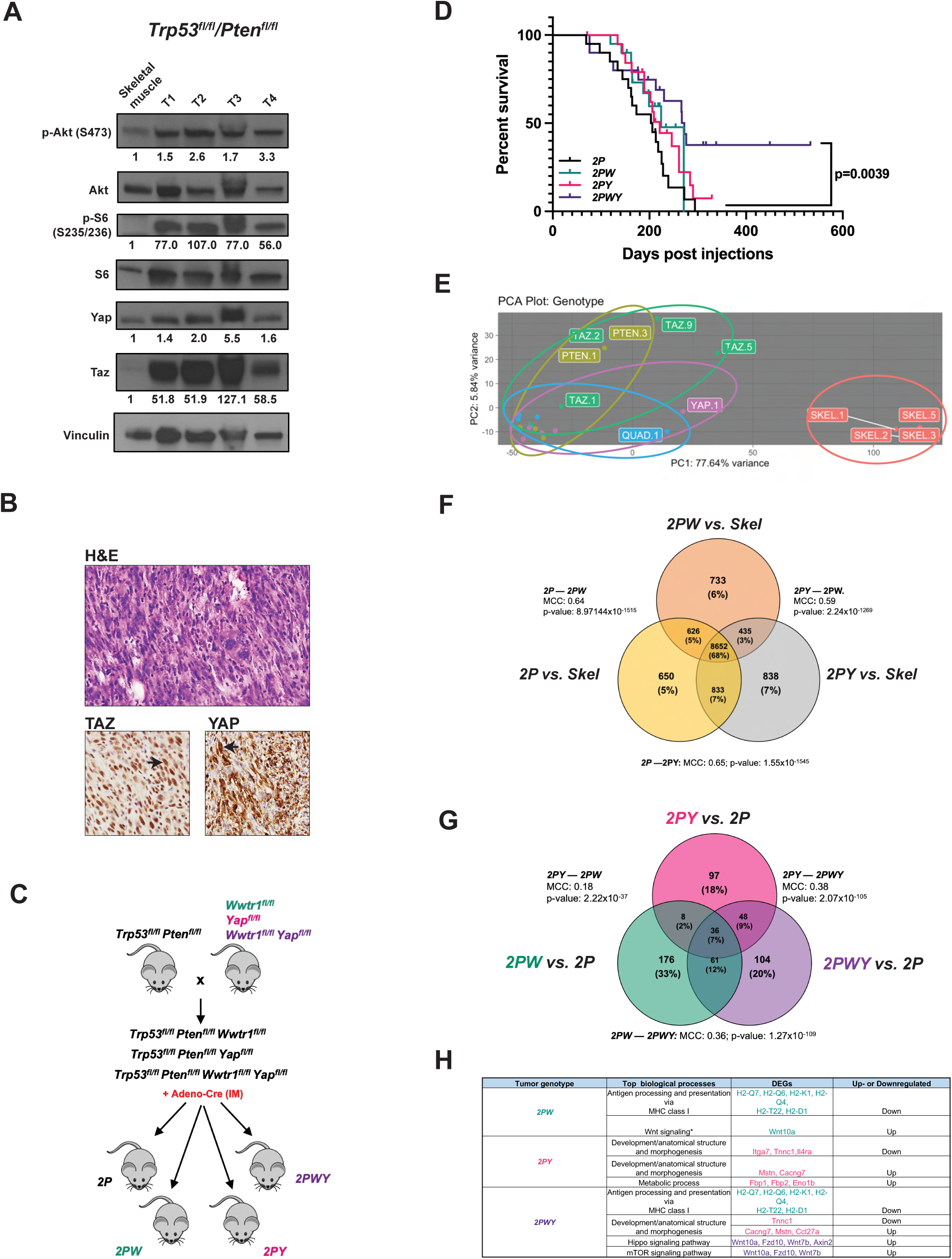
TAZ and YAP are critical oncoproteins in a PI3K activated mouse model of sarcoma. **(A)** Western blot analysis of four representative tumors from *Trp53^fl/fl^Pten^fl/fl^* mice evaluating activation of PI3K-AKT-mTOR signaling and TAZ/YAP expression relative to normal murine skeletal muscle. Vinculin was used as the loading control. (**B**) H&E section from a *Trp53^fl/fl^Pten^fl/fl^* primary tumor exhibiting atypical spindle cells and high mitotic figures (top panel). Immunohistochemistry for TAZ and YAP demonstrating nuclear positivity indicative of activation (bottom panels). (**C**) Schematic for mouse crosses; *Trp53^fl/fl^Pten^fl/fl^* mice were crossed with *Wwtr1^fl/fl^*(Taz^fl/fl^), *Yap1^fl/fl^*, or *Wwtr1^fl/fl^Yap^fl/fl^* to generate *2P*, *2PW*, *2PY*, and *2PWY* mice. AdenoCre virus was injected intramuscularly into the left quadriceps. (**D**) Survival curves for each mouse cohort. (**E**) Principal component analysis of RNA expression after variance-stabilizing transformation for 5 representative samples from murine skeletal muscle and *2P*, *2PW*, *2PY*, and *2PWY* tumors. (**F**) Number of differentially expressed genes in *2P*, *2PW*, and *2PY* tumors normalized to murine skeletal muscle. MCC=Matthew’s correlation coefficient. p values calculated via hypergeometric analysis. (**G**) Number of differentially expressed genes in *2PW*, *2PY*, and *2PWY* tumors normalized to *2P* tumors. MCC=Matthew’s correlation coefficient. p values calculated via hypergeometric analysis. (**H**) List of top biological processes and their respective differentially expressed genes (DEGs) for *2PW*, *2PY*, and *2PWY* tumors; * although Wnt10a is differentially expressed in *2PW* mice, Wnt signaling as a biological process, overall, is not enriched. Statistical analysis for Kaplan-Meier survival analysis was performed with the log-rank (Mantel-Cox) test. The above experiment with *2P*, *2PW*, *2PY*, and *2PWY* mice was performed twice.

To determine if TAZ and YAP were functionally important to the formation of PI3K-driven sarcomas *in vivo, Tr<u>p</u>53^fl/fl^<u>P</u>ten^fl/fl^* mice (hereafter referred to as *2P*) were crossed with: *Wwtr1^fl/fl^* mice to generate *Tr<u>p</u>53^fl/fl^<u>P</u>ten^fl/fl^Wwtr1^fl/fl^* mice (*2PW)*, *Yap1^fl/fl^* mice to generate *Tr<u>p</u>53^fl/fl^<u>P</u>ten^fl/fl^Yap1^fl/fl^ mice* (*2PY*), and *Wwtr1^fl/fl^ Yap1^fl/fl^* mice to generate (*2PWY*) (**Figure 4C**). Mice (n=20) in each arm of the study were injected intramuscularly with Ad CMV-Cre recombinase with Ad CMV-eGFP controls as described in Materials and Methods. Subsequent to intramuscular delivery of Ad CMV-Cre, survival analysis for each cohort was performed once tumors reached 1.5 cm in the greatest dimension at the site of injection (according to animal use protocol; see Materials and Methods for details). No significant differences in histological types of sarcomas generated were seen between the different groups, which were comprised predominantly of undifferentiated pleomorphic sarcoma/undifferentiated spindle cell sarcoma as previously described in the *Trp53^fl/fl^Pten^fl/fl^* mouse model (78) (see **Table S1**). Survival analysis showed that relative to *2P* mice, conditional knock-out of either *Wwtr1^fl/fl^* (*2PW*, p=0.17) or *Yap1* (*2PY*, p=.17) alone does not improve overall survival whereas loss of both significantly extends the overall survival (*2PWY*, p=0.0039) (**Figure 4D**). This indicates that TAZ and YAP have distinct, complementary functions in this PI3K activated, sarcoma-prone mouse model.

To determine whether differences in survival was due to differences in time to tumor formation, we compared the time to tumor formation among the four different phenotypes, and saw that 2*PY* mice showed an increased latency to tumor formation compared to *2P* mice (176 days vs. 126 days; p-value of p=0.007) (**Figure S5A**). *2PWY* mice showed a trends towards increase in the latency of tumor formation, however this did not reach statistical significance (p-value of p=0.11). Finally, we compared the fraction of mice that developed tumors in the span of the experiment, 533 days, and found that while tumors developed in all mice of the *2P*, *2PY*, and *2PW* genotypes, 6 of the 20 mice (30%) with the *2PWY* genotype did not develop tumors at the injection site by the end of the study. To determine whether there were differences in the growth rate of the tumors, we evaluated the time to tripling of the volume of the tumors, and found that tumors from the *2PW*, *2PY*, and *2PWY* genotypes showed no difference compared to the *2P* controls (**Figure S5B**). Taken together, TAZ and YAP when coordinately inactivated *in vivo* play a larger role in tumor initiation than proliferation.

To determine if the above phenotype *in vivo* seen with inactivation of TAZ and YAP was due to changes in transcription, RNA was isolated from 5 tumors (undifferentiated pleomorphic sarcoma) from each genotype and total RNA-Seq was performed. By PCA analysis, samples from the groups tended to cluster together and were distinct relative to normal skeletal muscle tissue (**Figure 4E** and **Figure S6A-D**). Beginning with *2P* (designated PTEN), ellipses representing tumor samples from each arm of the study demonstrated a clock-wise rotation corresponding to sequential loss of Taz (*2PW*), Yap (*2PY*), or both (*2PWY;* designated QUAD). (**Figure 4E**). This arrangement suggests a step-wise change in the overall transcriptome with inactivation of TAZ, YAP, or both. The transcriptomes of 2PW and 2PYoverlapped significantly with 2P (<u>M</u>atthews <u>c</u>orrelation <u>c</u>oefficient or MCC of 0.64 and 0.65 respectively, p < 0.0001 for both comparisons) **(Figure 4F)**. These findings show that TAZ and YAP drive a significant portion of the PI3K transcriptome.

Differentially expressed genes (DEGs) in the conditional knock-out of both *Wwtr1* and *Yap1* (*2PWY*) overlapped with DEGs seen in the *2PW* and *2PY* transcriptomes. However, conditional knock-out of Taz and Yap in *2PWY* mice did not simply have an additive effect on gene expression, the *2PWY* transcriptome was also composed of a subset of genes that were unique and likely represented a set of genes coordinately regulated by both Taz and Yap (**Figure 4G**). By pathway analysis, conditional knock-out of *Wwtr1* (*2PW*) showed inactivation of genes important for antigen processing and presentation and Wnt signaling (**Figure 4H**). Conditional knock-out of *Yap1* (*2PY*) showed differential expression of a series of genes important for development (**Figure 4H**). The *2PWY* transcriptome demonstrated DEGs identified by pathway analysis in the *2PW* and *2PY* transcriptomes. However, consistent with the above Venn diagram analysis showing DEGs unique to the *2PWY* transcriptome (**Figure 4G**), pathway analysis also showed differential expression of genes involved in Hippo signaling (*Wnt10a, Fzd10*, *Wnt7b*, *Axin2*) and mTOR biological process (*Wnt10a*, *Fzd10*, and *Wnt7b*) unique to the *2PWY* transcriptome (**Figure 4H**). Well-defined transcriptional targets of TAZ and YAP, *Ccn1* (Cyr61) and *Ccn2* (Ctgf), were downregulated in *2PWY* mice vs *2P* mice (log2 fold change of −0.193 for *Ccn2*, and −0.059 for *Ccn1*), although this did not reach statistical significance (**Table S2**). Collectively, the above *in vivo* data suggest that TAZ and YAP are key oncogenic transcriptional effectors in a PI3K activated mouse model of sarcoma.

### mTORC1 inhibition has a limited therapeutic effect *in vitro* and *in vivo*

Taken together, the above *in vitro* and *in vivo* data suggest that the PI3K-TAZ/YAP axis represents another key signaling axis in addition to the PI3K-Akt-mTORC1 axis. To begin assessing the relative efficacy of targeting the two axes, we evaluated the efficacy of targeting the PI3K-Akt-mTORC1 axis utilizing everolimus (a rapamycin derivative) *in vivo* in the PI3K-driven *Trp53^fl/fl^/Pten^fl/fl^* sarcoma mouse model mentioned above (**Figure 4**). Everolimus (5mgs/kg) or vehicle (DMSO) was administered orally on a weekly basis as previously described (79). Tumor dimensions were measured until an endpoint of 2 cm was met. Kaplan-Meier analysis indicated an initial survival benefit to single agent therapy with everolimus with the maximum difference between the survival curves seen at 220 days with Everolimus showing a survival fraction of 0.79 and vehicle with a survival fraction of 0.44. However, this difference in survival decreased over the course of the experiment, with no change in overall survival at the culmination of the study (p=0.2336) **(Figure 5A)**. This is consistent with clinical data showing that mTORC1 inhibition alone is not effective in reducing tumor burden/survival *in vivo* as has been documented in various contexts (80–82).

**Figure 5:**
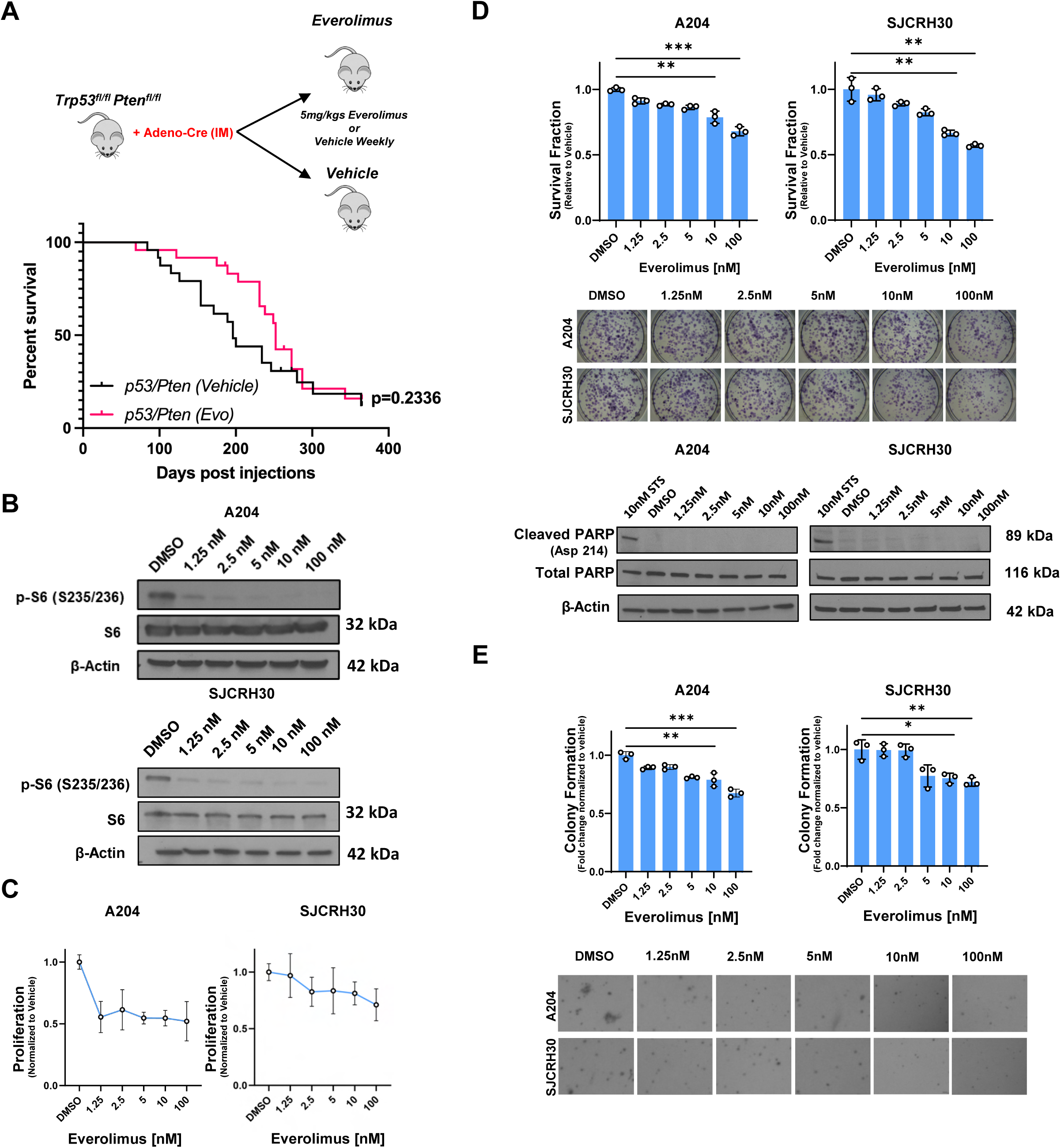
mTORC1 inhibition shows a modest effect on proliferation and growth *in vivo* and *in vitro*. **A)** Everolimus (5mgs/kgs) or vehicle (DMSO) was given weekly by oral gave in *2P* mice (n=16/group). Kaplan-Meier survival analysis shown. **B**) Western blot evaluating p-S6 (S235/236) as a function of everolimus concentration in both A204 and SJCRH30 cell lines. **C)** Drug response curve after treatment with everolimus for 72 hours in the A204 and SJCRH30 cell lines. **D)** Clonigenic outgrowth assay and evaluation of PARP-cleavage (staurosporine is the positive control) after treatment with everolimus for 72 hours in the A204 and SJCRH30 cell lines. **E)** Soft agar assay after treatment with everolimus. For soft agar and clonogenic assays, statistical significance was evaluated using an unpaired two-tailed *t*-test. All in vitro assays were repeated at least twice. Approximately equal numbers of male and female mice were used for the above mouse study. Statistical analysis for Kaplan-Meier survival analysis was performed with the log-rank (Mantel-Cox) test. For all panels, ****p<0.0001, ***p<0.001, **p<0.01, *p<0.05.

To dissect the therapeutic limitations of everolimus observed *in vivo*, we treated the SJRCH30 and A204 cell lines with everolimus and assessed various hallmarks of cancer *in vitro*. Everolimus was shown to decrease phosphorylation of the S6 ribosomal subunit at a wide range of concentrations spanning from 1 nM to 100 nM (10 nM is a concentration often utilized clinically)(83) **(Figure 5B).** Reduction in S6 phosphorylation was observed at sub-10 nM concentrations of everolimus in both the SJCRH30 and A204 cell lines **(Figure 5B)**. In contrast, this inhibitory effect resulted in only a modest decrease in proliferation, even at higher concentrations of 100 nM (10-fold higher than the clinically relevant dose) **(Figure 5C)**. To determine whether this decrease in proliferation could be due to cytotoxic effects of everolimus, we performed clonogenic outgrowth assays over a range of concentrations. Clonogenic assays showed survival fractions that were marginally reduced at physiologically relevant doses of everolimus and only reduced by 25-40% at the highest concentration of 100 nM **(Figure 5D).** No change in cleaved PARP was seen via western blot over this range of concentrations of everolimus. All together, the above findings demonstrate low cytotoxicity and an absence of apoptosis with treatment with everolimus. Similarly, effects of mTORC1 inhibition on anchorage independent growth were modest and not seen until higher concentrations **(Figure 5E)**.

### TEAD inhibition alone has modest efficacy *in vitro* and predominantly affects anchorage independent growth

We then assessed whether it may be more efficacious to target the PI3K-TAZ/YAP axis. To determine whether IK-930, a recently described TEAD inhibitor (84), disrupts the interaction of YAP with TEAD1/TEAD transcription factors, A204 and SJCRH30 were treated with 1 µM IK-930 for 24 hours. To evaluate the effect of TEAD inhibition on well-defined transcriptional targets of YAP and TAZ, qRT-PCR was done including the clinically relevant concentration of 1 µM IK-930 in the SJCRH30 and A204 cell lines **(Figure 6A)**. Expression of *CCN2* (CTGF) and *CCN1* (CYR61) was decreased approximately 5 fold upon treatment with 1 µM, a clinically relevant dose, of IK-930 in both cell lines **(Figure 6A)**. To confirm the ability of IK-930 to inhibit TEAD-dependent transcription, A204 and SJCRH30 cell lines were transduced with a TEAD Luciferase (pLV-8XTEADrep) reporter construct plasmid. Luciferase activity was measured in the SJCRH30 and A204 cell lines after 72 hours of treatment with various concentrations of IK-930 **(Figure 6B)**. A dose dependent reduction in TEAD reporter activity was seen in both A204 and SJCRH30 with IK-930 treatment (**Figure 6B).**

**Figure 6:**
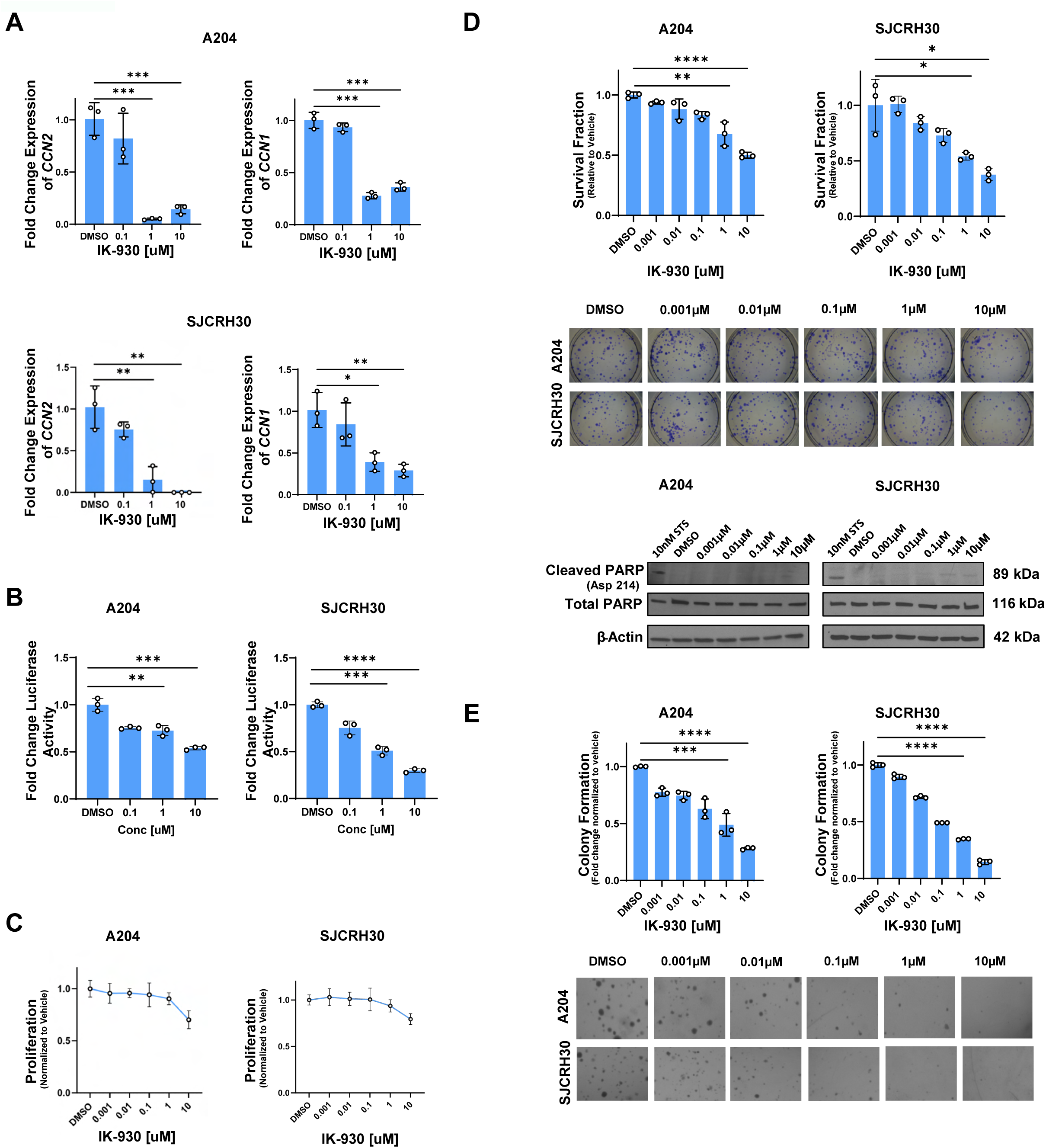
YAP/TAZ inhibition significantly reduces growth-related phenotype. **A)** Quantitative RT-PCR (qRT-PCR) showed reduced *CCN1* (CYR61) and *CCN2* (CTGF) expression upon treatment with IK-930. **B)** Dual luciferase reporter assay after treatment with IK-930. **(C)** Drug response curve after treatment with IK-930 for 72 hours. **D)** Clonogenic outgrowth assay and evaluation of PARP-cleavage (staurosporine is the positive control) after treatment with varying concentrations of IK-930 after 72 hours. **E)** Soft agar assay after treatment with IK-930. For quantitative RT-PCR and luciferase reporter assays, standard deviation was calculated from fold change values for each triplicate. For all assays, statistical significance was evaluated using an unpaired two-tailed *t*-test. All in vitro assays were repeated at least twice. For all panels, ****p<0.0001, ***p<0.001, **p<0.01, *p<0.05.

To examine the effect of IK-930 on sarcoma cell proliferation, treatment of SJCRH30 and A204 cells with various concentrations of IK-930 resulted in modest reductions in proliferation with only the higher concentrations of IK-930 (1 µM, 10 µM) showing a slight effect **(Figure 6C).** Similar effects were observed when performing a clonogenic assay for IK-930 with no significant effects on the survival fraction until concentrations reached higher concentrations between 1 and 10 µM **(Figure 6D)**. To investigate the effect of TEAD inhibition on apoptosis, western blot was performed on A204 and SJCRH30 cells to examine PARP cleavage (**Figure 6D**). No increase in cleaved PARP was identified with increasing concentrations of IK-930 suggesting the limited cytotoxicity seen in higher concentrations of IK-930 may be due to other mechanisms of cell death (e.g. necrosis, ferroptosis) **(Figure 6D)**. YAP and TAZ have been implicated in anchorage independent growth in various cancer types (85–89). To test the effect of TEAD inhibition on anchorage independent growth, we performed soft agar assays for both cell lines **(Figure 6E)**. In contrast to proliferation, IK-930 demonstrated reduction in anchorage independent growth at lower concentrations including the 1-10 nM range **(Figure 6E).** All together, TEAD inhibition had the greatest effect on anchorage independent growth, with more modest effects on proliferation and cytotoxicity.

### mTORC1 and TEAD inhibitors show synergistic activity in reducing proliferation and soft agar growth

The above data showed that inhibition of mTORC1 (everolimus) predominantly affected proliferation while inhibition of the TAZ/YAP (with TEAD inhibitor IK-930) predominantly affected anchorage independent growth; both drugs exhibited limited cytotoxic effect. This led to the hypothesis that targeting both the PI3K-Akt-mTORC1 and PI3K-TAZ/YAP axes might be more effective from a therapeutic standpoint. To test this hypothesis, we treated the SJCRH30 and A204 cell lines with varying concentrations of everolimus and IK-930 in combination. A range of concentrations centered around their IC-50s were used and the online tool, SynergyFinder (90), was used to identify synergistic concentrations of the two drugs. Using the Bliss quantitative model within the SynergyFinder algorithm we found that the effect of combination therapy on proliferation was greater than that of either single agent (90,91) **(Figure 7A).** The synergy score was higher for A204 (47.3) as compared to SJCRH30 cell line (14.7), but the contours of the synergy maps showed similar patterns with a primary peak at lower concentrations of IK-930 (≤10 nM) and everolimus (≤5 nM) **(Figure 7A).** For A204, 10 nM of IK-930 and 2.5 nM of everolimus showed a synergistic reduction in proliferation by approximately 43% in comparison with vehicle. Whereas for SJCRH30, 1 nM of IK-930 and 5 nM of Everolimus showed a synergistic reduction in proliferation by 37% in comparison with vehicle. In contrast, at the above concentrations, single agent IK-930 showed a 0-4% reduction in proliferation, while single agent everolimus demonstrated a 20% reduction in proliferation.

**Figure 7:**
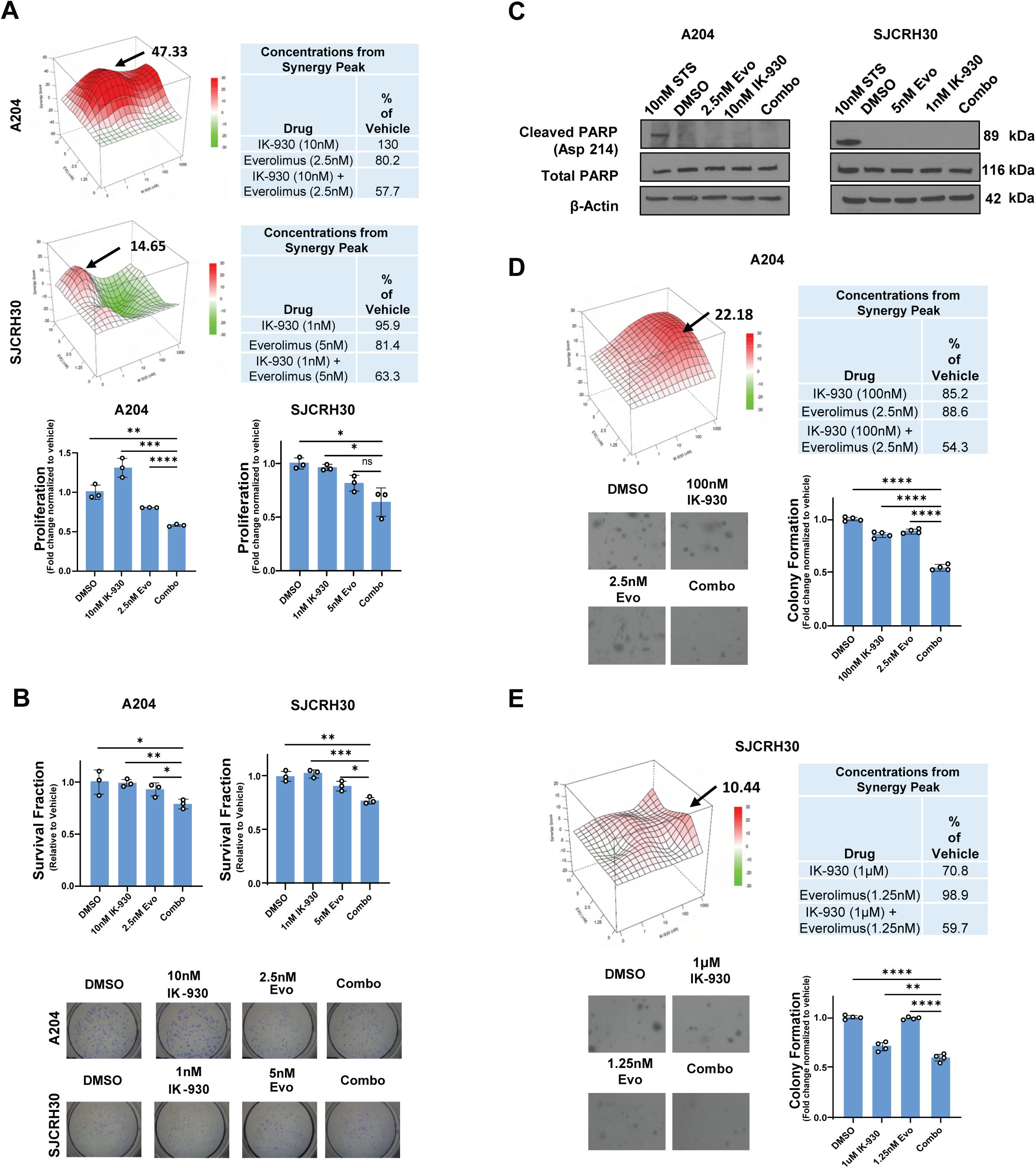
Pharmacological inhibition of YAP/TAZ and mTORC1 works synergistically *in vitro*. **A)** SJCRH30 and A204 cells were treated with combinational drug matrix of IK-930 and everolimus at indicated doses for 72 hours and cell viability evaluated by MTT-style assay. Synergy scores and 3D surface plots of cell viability in both cell lines were quantified and analyzed with the Bliss model using SynergyFinder. Arrows demonstrate concentrations of IK-930 and everolimus further evaluated in table form and graphically. **B)** Clonogenic assay using the synergistic concentrations in (**A**) showed statistically significant reduction in clonogenic outgrowth. **C)** Western blot evaluated cleaved PARP after monotherapy and combination therapy for the synergistic concentration in (**B**). **D, E)** Synergy scores and 3D surface plots of combination studies in soft agar in the A204 cell line (**D**) and SJCRH30 cell line (**E**) were also quantified and analyzed with the Bliss model using SynergyFinder. Arrows demonstrate concentrations of IK-930 and everolimus further evaluated in table form. All *in vitro* assays were repeated at least twice. Staurosporine was used as a positive control. For soft agar assay, statistical significance was evaluated using an unpaired two-tailed t-test. For all panels, ****p<0.0001, ***p<0.001, **p<0.01, *p<0.05.

These synergistic concentrations in **Figure 7A** were then used to study the effect on clonogenic outgrowth in SJCRH30 and A204 cell lines **(Figure 7B)**. Statistically significant reductions in clonogenic outgrowth compared to vehicle control, 23% for SJCRH30 (p=0.043) and 21% for A204 (p=0.0027), were seen using these low, physiologically relevant concentrations. To determine if this effect on clonogenic outgrowth was due to apoptosis, western blot analysis was used to evaluate changes in cleaved PARP. PARP cleavage was not identified with the combination of everolimus and IK-930 or the single agent controls (**Figure 7C**). All together, the above data suggests the combination therapy does cause some cell death in an apoptosis-independent manner such as necrosis or ferroptosis (92,93) at lower, more clinically relevant concentrations than everolimus alone **(Figure 5D**) or IK-930 alone (**Figure 6D**). To investigate the effect of combination therapy on anchorage independent growth, soft agar assays were performed utilizing varying concentrations of both IK-930 and everolimus in agarose.

Synergistic activity of TEAD inhibition and mTORC1 inhibition at 100 nM and 2.5 nM, respectively, was seen represented by a peak with a synergy score of 22.2 in the A204 cell line **(Figure 7D)**. The combination therapy reduced anchorage independent growth by 46%, while single agent therapy with everolimus or IK-930 at those concentrations only reduced anchorage independent growth by approximately 15% in A204 cell line **(Figure 7D)**. For SJCRH30, the synergy peak was seen at 1µM of IK-930 and 1.25 nM of everolimus with a synergy score of 10.4 **(Figure 7E).** The combination therapy reduced anchorage independent growth by 40%, while single agent IK-930 and everolimus reduced anchorage independent growth by around 30% and 2%, respectively, in the SJCRH30 cell line **(Figure 7E).**

While everolimus alone predominantly affected proliferation (**Figure 5**), and TEAD inhibition alone predominantly affected anchorage independent growth (**Figure 6**), the above data show that the combination therapy synergistically affected proliferation (**Figure 7A**) and anchorage independent growth (**Figure 7D**), while causing some apoptosis-independent cell death at more physiologically relevant doses than the single agent approaches (**Figure 7B**).

### Combination therapy with TEAD and mTORC1 inhibitors reduces tumor volume *in vivo*

To determine if this combination therapy would be effective *in vivo*, we further evaluated the effect of combination therapy in a mouse xenograft model. 5×10^6^ SJCRH30 cells were subcutaneously injected in *Prkdc^scid^Il2rg^tm1Wjl^/SzJ* (NSG) mice. Once tumors were palpable (5 mm^3^) mice were divided into 4 groups: vehicle, IK-930 alone, everolimus alone, and IK-930 combined with everolimus. IK-930 (75mgs/kgs) was administered orally every day while everolimus (5mgs/kgs) was administered orally on a weekly basis (79). Tumor volume was measured every other day until tumors in the vehicle arm averaged 2 cm. Significantly, in the combination therapy arm, every treatment of everolimus (weekly administration of everolimus denoted by arrows) resulted in a reduction of tumor growth **(Figure 8A)**. At the end of the study, tumor volume was measured *in situ* in the mice showing an almost two-fold reduction in tumor volume for combination therapy-treated mice in comparison to vehicle and single agent therapy controls (p < 0.01)**(Figure 8B)**. Histological evaluation of resected tumors showed no change in percent necrosis (**Figure 8C**). Immunohistochemistry showed no change in cleaved caspase 3 activity (**Figure 8D**). However, an approximately 20% reduction in Ki-67, a proliferation marker, was seen in the combination therapy cohort in comparison with vehicle (p=0.001) and single agent treated mice (p=0.0002 for Everolimus and p=0.002 for IK-930) **(Figure 8E).** Collectively, these results support the working model that targeting both the PI3K-Akt-mTORC1 axis and PI3K-TAZ/YAP axes can be leveraged to more completely inhibit oncogenic PI3K signaling in sarcomas and potentially other cancers **(Figure 8F)**.

**Figure 8:**
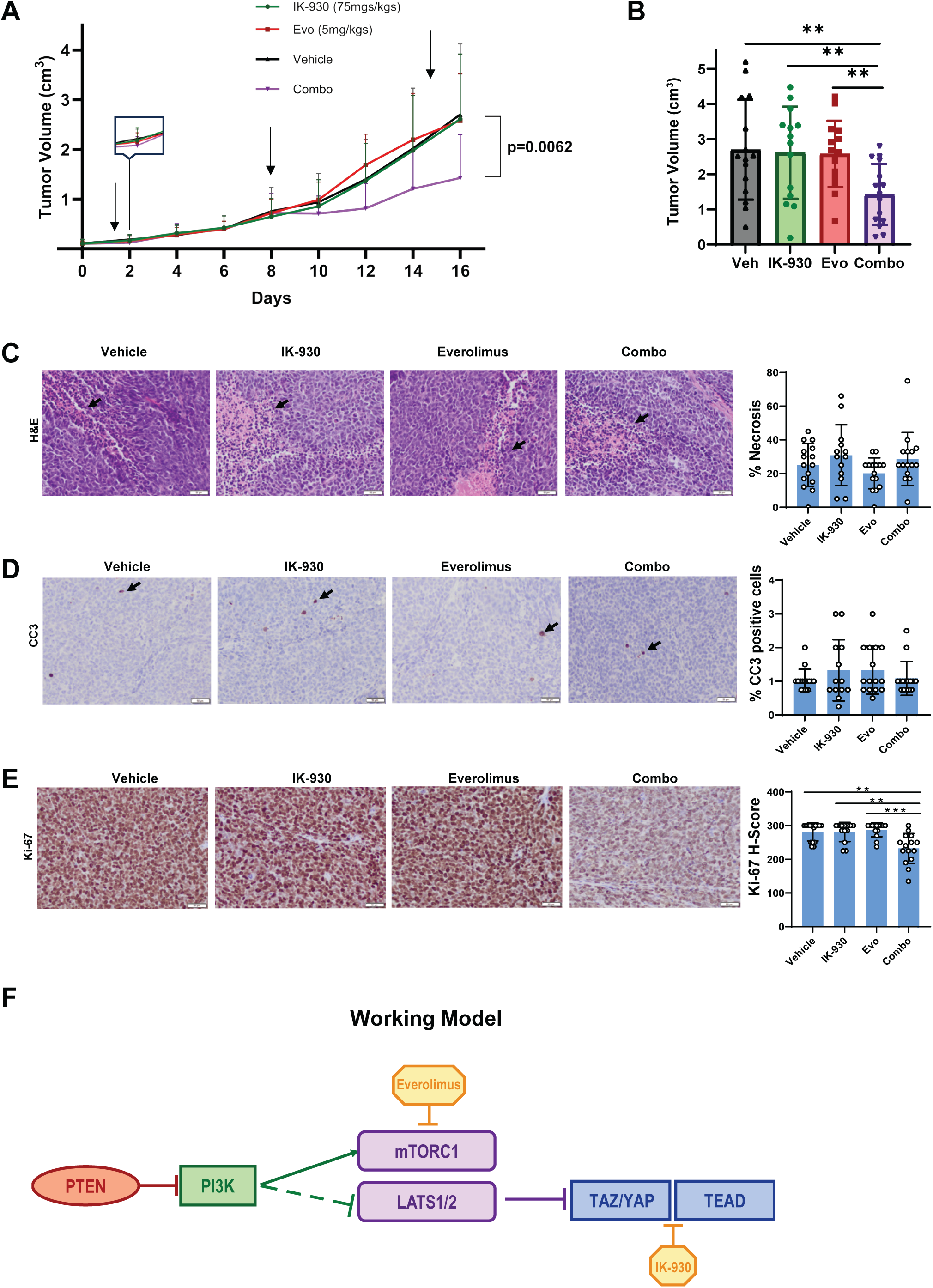
Combination therapy targeting YAP/TAZ and mTORC1 inhibits tumor growth *in vivo*. **A)** Tumor growth curve SJCRH30 cells injected subcutaneously into NSG mice. Mice were divided into 4 groups and given IK-930 (75mgs/kgs-daily), everolimus (Evo) (5mg/kgs-weekly), DMSO (Vehicle) and both IK-930 and everolimus in combination (Combo) by oral gavage (n=16/group). Arrows denote weekly administration of everolimus. **B)** Tumor volume at the end of study in in (**A**) for all four treatment groups. **C)** H&E slides showing morphology and necrosis (arrows demonstrate necrosis); graphically represented to the right**. D)** Immunohistochemistry for cleaved caspase 3 (arrows demonstrate positive cells); graphically represented to the right**. E)** Immunohistochemistry for Ki-67. H-score for Ki-67 staining shown; graphically represented to the right. **F)** Working model showing mTORC1 and YAP/TAZ as two axes downstream of PI3K signaling, providing the rationale for combination therapy targeting mTORC1 and the YAP/TAZ-TEAD interface. Scale bar for photomicrographs is 50 µm (200X). Approximately equal numbers of male and female mice were used for the Xenograft study. For tumor volume and histological analysis, statistical significance was evaluated using an unpaired two-tailed *t*-test. For all panels, ****p<0.0001, ***p<0.001, **p<0.01, *p<0.05.

## Discussion

### Phosphotidylinositol-3 kinase is frequently activated in sarcomas

Our studies show that despite a low mutational burden for some of the core PI3K signaling enzymes, PI3K signaling could be activated at some level in up to 60% of sarcomas due to absent or very low PTEN expression, representing an underutilized, common therapeutic target active in various sarcomas. Loss of expression of PTEN varied across different histological types but is prevalent in key sarcoma histological types including undifferentiated pleomorphic sarcoma/undifferentiated spindle cell sarcoma, the most common histological type of sarcoma (1). This observation is validated by the observation that the *Trp53*^fl/fl^*Pten*^fl/fl^ sarcoma mouse model generates predominantly undifferentiated pleomorphic sarcomas (78). The loss of PTEN expression seen at the protein level in clinical samples (30-60%) (**Figure 1 C, D)** exceeds the frequency of mutations-deletions seen in *PTEN* at the genomic level (5-10%) (**Figure 1B**), suggesting the epigenetic mechanisms may be playing a role in the regulation of PTEN. Previous reports have shown that YAP suppresses PTEN expression by inducing expression of miR-29 in epithelial cell systems and carcinomas (28). Additional studies are needed to elucidate the mechanisms leading to the loss of expression of PTEN in sarcomas.

### Targeting mTORC1 demonstrates limited efficacy in PI3K-driven sarcomas

PI3K signaling is activated in a number of cancers including breast and ovarian cancer due to mutations in *PIK3CA*, *AKT*1/2/3, and loss of *PTEN* (94–97). Several approaches have been used to target these cancers driven by PI3K signaling including: targeting of PI3K, inhibition of AKT, combination approaches targeting PI3K and AKT, inhibition of mTORC1 (98,99), and combination approaches targeting PI3K and mTORC1 (100). Most current approaches targeting PI3K signaling have utilized rapamycin derivatives (e.g. sirolimus, everolimus) while acknowledging limitations due to various feedback mechanisms (24,101–104). To address these limitations, mTORC1 inhibition has been pursued in sarcomas for clinical trials in combination with other therapeutic agents (105,106). Malignant perivascular epithelioid cell tumor (PEComa) is currently the only sarcoma in which mTORC1 inhibition is approved (107). Our studies largely recapitulate these observations (**Figure 5**). Although everolimus showed a modest decrease in proliferation there was limited cytoxicity *in vitro.* Initial gains in overall survival *in vivo* diminished over time as tumors became resistant to rapamycin-based therapy. All together, our findings and previous studies demonstrate limitations in treating PI3K-driven cancers including sarcomas with mTORC1 inhibition alone. Additional therapeutic approaches are needed to target PI3K signaling in these cancers including targeting of the PI3K-TAZ/YAP axis.

### PI3K regulates YAP/TAZ driven transcription via a LATS1/2 dependent mechanism

One persistent question in the study of PI3K signaling is how signals from the cells surface are transduced into the nucleus. Unlike other oncogenic signal transduction pathways, a *bona fide* oncogenic transcription factor downstream of PI3K has not been elucidated. In these studies, we find that TAZ and YAP represent an additional arm of the PI3K signaling axis and represent oncogenic transcription factors/transcriptional coactivators identified downstream of PI3K signaling. Our observations support a working model that a PI3K-TAZ/YAP axis exists in parallel to a PI3K-Akt-mTORC1 axis in sarcomas and potentially other cancers. Our work shows that PI3K regulates TAZ/YAP in via a LATS1/2 dependent mechanism in sarcomas, which may differ from mechanisms implicating PDK1, AKT1, or GSK-3β derived from studies in epithelial cells (31,33,73,74). In gastric carcinomas, phosphorylated (inactivated) PTEN disrupted the MOB1-LATS1/2 interaction resulting in YAP nuclear localization and activation (108). Future studies are warranted to identify the intermediate steps by which PI3K signaling inactivates LATS1/2 to activate TAZ/YAP in sarcomas.

### TAZ and YAP are central oncoproteins in PI3K driven oncogenesis and coordinately mediate the PI3K transcriptome

TAZ and YAP are well known drivers of initiation, progression, and metastasis (109). However, to our knowledge, analysis of the role and relative contributions of Taz and Yap in genetically engineered mouse models of sarcomas have not been explored. It is increasingly appreciated that TAZ and YAP do not completely phenocopy each other and may functionally compensate for one another (110,111). We showed for the first time in a PI3K-driven sarcoma mouse model, that inactivation of both Taz and Yap are needed to inhibit tumor initiation and have an effect on overall survival. This suggests that TAZ and YAP have complimentary roles in PI3K-driven sarcomagenesis, and is consistent with the transcriptomic data shown above, which showed that TAZ and YAP have overlapping yet distinct transcriptomes and coordinately regulate a unique set of genes in the PI3K transcriptome (**Figure 4F** and **4G**). Collectively, our findings and those of others indicate that Taz and Yap play distinct and complementary roles in sarcoma development and mediate the oncogenic PI3K transcriptome. Future studies are needed to further elucidate how the PI3K-TAZ/YAP transcriptome drives sarcoma formation.

### mTORC1 and YAP/TAZ constitute effector arms of PI3K signaling providing a rationale for combination therapy

The above findings show that YAP/TAZ are downstream oncogenic transcriptional co-activators of PI3K signaling thus providing a rationale for combination therapy targeting the TEAD transcription factors along with mTORC1 inhibitors. We show that while IK-930 as a single agent was effective in decreasing anchorage independent growth, and that everolimus inhibited proliferation *in vitro*, they were both ineffective as single agent therapies *in vivo*. Consistent with our *in vitro* studies, combining the two drugs *in vivo* showed a synergistic decrease in proliferative index and tumor volume that could have relevance to the treatment of sarcomas and potentially other PI3K-dependent cancers. The synergistic effect, both *in vitro* and *in vivo,* between YAP/TAZ-TEAD and mTORC1 inhibition can potentially be explained by the hypothesis this approach inhibits both the transcriptional (YAP/TAZ) and translational (mTORC1) apparatus downstream of PI3K signaling (85,112,113).

### Combination therapy may enhance the efficacy of TEAD inhibitors

Although TEAD inhibitors may be effective as a single agent in some settings, concerns regarding their efficacy as a monotherapy exist. Indeed our studies show that as a single agent, a TEAD inhibitor was insufficient to inhibit sarcoma growth in a TAZ/YAP-dependent xenograft mouse model. It was only with combination targeting of a parallel signaling axis that the TEAD inhibitor was effective. Rational combination of a TEAD inhibitor with additional agents targeting a relevant pathway is emerging as a viable therapeutic strategy (114–118). Combination therapy with TEAD inhibitors may also be a reasonable approach to mitigating the risk of secondary resistance developing to TEAD inhibitors. For these reasons, additional investigation into the basic mechanisms driving or complementing TAZ and YAP activation in different cancers is needed to more effectively utilize TEAD inhibitors in TAZ/YAP dependent cancers.

## Materials and Methods

### Cell culture

SJCRH30 (human alveolar rhabdomyosarcoma) and A204 (human malignant extrarenal rhabdoid tumor) cells were obtained from American Type Culture Collection (ATCC, Manassas, VA, USA). SJCRH30 and A204 cells were cultured in RPMI and McCoy’s 5A media (Gibco), respectively. Media was supplemented with (according to ATCC recommendations) 10% fetal bovine serum (Invitrogen-Life Technologies), 1 mM sodium pyruvate, and 50 µg/mL penicillin/streptomycin. All cells were cultured at 37 °C and 5% CO_2_. For PI3K inhibition studies, SJCRH30 and A204 cells were treated with 30 µM/60 µM LY294002 (TOCRIS, catalog# 1130), 10 µM GDC-0941 (Selleckchem, catalog# S1065). SJCRH30 and A204 cells were treated with 10 nM-10 µM of the AKT inhibitor MK-2206 2HCl (Selleckchem, catalog# S1078). To evaluate proteasomal degradation, SJCRH30 and A204 cells were treated with 10 µM of MG-132 (Sigma-Aldrich, catalog# C2211) alone or in combination with 60 µM LY294002. To evaluate PARP cleavage by western blot, Staurosporine (10nM) was used as a positive control for apoptosis in A204 and SJCRH30 cells.

### Histopathology

Sarcoma samples were retrieved from the University of Iowa Department of Pathology with previous approval from the Institutional Review Board. The tissue microarray was constructed by arraying 1.0 mm cores taken from formalin fixed paraffin embedded tissue and assembled using a MTA-1 tissue arrayer from Beecher Instruments (Sun Prairie, WI) as previously described(46). Sarcomas were classified according to World Health Organization criteria (1).

### Expression constructs

Double Flag TAZ and TAZ4SA were derived as previously described in (47). TAZ S58/62A mutations were introduced to Double Flag TAZ by site directed mutagenesis using the QuikChange II Site-directed mutagenesis kit (Agilent #200521) and the following primers:

TAZ S58A Primer – Forward

5’ – ctttctttaaggagcctgatgcgggctcgcactc – 3’

TAZ S58A Primer – Reverse

5’ – gagtgcgagcccgcatcaggctccttaaagaaag – 3’

TAZ S62A Primer – Forward

5’ – cgggctcgcacgcgcgccagtcc – 3’

TAZ S62A Primer – Reverse

5’ – ggactggcgcgcgtgcgagcccg – 3’

### Transfection and retroviral transduction

Retroviral transduction with pBabeNeo constructs was performed by transfecting PhoenixA retroviral packaging cells with 10 µg of plasmid DNA. Transfection was done using Lipofectamine Reagent and Plus Reagent (Invitrogen-Life Technologies) according to manufacturer’s instructions. Supernatant was collected at 48 and 72 hours after transfection, filtered with 0.45 µm filters and supplemented with 8 µg/mL polybrene (EMD Millipore, Burlington, MA, USA). Serial transductions (48 and 72-hour supernatants) were applied to either SJCRH30 or A204 cells for 8 hours each. Pooled stable lines were generated by selecting with G418 for two weeks.

### siRNA knockdown

A204 and SJCRH30 cells were grown to 60-70% confluence in 6cm tissue culture plates. Each plate received 5.5uL of 20uM On-TARGETplus SMARTPOOL (Dharmacon) siRNA (AKT1, cat# L-003000-00-0005, PIK3CA, cat# L-003018-00-0005, PDK1, cat# L-003017-00-0005) that was forward transfected with Lipofectamine® RNAiMAX (cat# 13778075) according to manufacturer’s protocol. Target cells treated with RNAi-Lipofectamine® RNAiMAX duplexes were incubated for 12, 24, or 48hrs before harvesting for western blot analysis.

### RNA interference-mediated silencing

The following pLKO.1-puromycin constructs were obtained from Sigma-Aldrich. Empty vector construct (SHC001) scrambled negative control (SHC002), and TAZ knock-down constructs TRCN0000319224 (shTAZ#3), TRCN0000370007 (shTAZ#5). YAP knock-down constructs were obtained from Addgene #42540 (shYAP#1), #42541 (shYAP1#2). Constructs were transfected into LentiX (HEK293T) cells, along with pcMVΔ8.12 and pVSVG packaging plasmids, using Lipofectamine Reagent and Plus Reagent (Invitrogen) according to manufacturer’s instructions. Supernatants were collected at 48 hours, filtered using 0.45 µm filters, and 40% polyethylene glycol (PEG) 8000 was added to a final concentration of 12%, and stored at 4 °C overnight. The following day, 48-hour supernatants were centrifuged, and viral pellets were resuspended in 0.45 µm filtered 72-hour media and 8 µg/mL polybrene (EMD Millipore, Burlington, MA, USA) added to either SJCRH30 or A204 overnight. Pooled stable lines were generated by selection in puromycin.

### Western blot

Cell pellets were lysed in radioimmunoprecipitation assay (RIPA) buffer, containing cOmplete™ Protease Inhibitor Cocktail (EDTA-free) (Roche) and PhosSTOP™ Phosphatase Inhibitor Cocktail (Roche) according to the manufacturer’s instructions. Total protein concentration was measured using Pierce BCA™ Protein Assay Kit (ThermoFisher Scientific, Waltham, MA, USA). Between 20 µg and 100 µg of total protein was loaded onto a gradient (4-15%) polyacrylamide gel (BioRad, Hercules, CA, USA). Proteins were then transferred to a polyvinylidene difluoride (PVDF) membrane and probed with antibodies described below. Each experiment was repeated at least twice. ImageJ (NIH) was used to quantitate protein densitometry.

### Nuclear and cytoplasmic fractionation

Nuclear and cytoplasmic fractionation was performed with the Nuclear Extract Kit (Active Motif, 40010). Cells collected from two 10cm cell culture plates that were centrifuged and resuspended in 500 µL 1X Hypotonic buffer and incubated for 5 min on ice. 25 µL of detergent was added to resuspended cells which were subsequently vortexed for 10s. After centrifugation, the supernatant cytoplasmic fraction was immediately processed for western blot analysis (see above). Remaining nuclear pellet was resuspended in 50-100 µL complete lysis buffer followed by pestle-motor vortexing for maximal nuclear protein extraction. After centrifugation, the supernatant nuclear fraction was immediately processed for western blot analysis.

### Antibodies for Western Blot and Histopathology

Anti-FLAG antibody (mouse monoclonal clone M2 (catalog # F3165) utilized for western blot (1:1000) was obtained from Sigma-Aldrich (St. Louis, MO, USA). Anti-β-actin antibody (AC-15; catalog #A544) utilized for western blot (1:10,000) was obtained from Sigma-Aldrich (St. Louis, MO, USA). Anti-alpha tubulin antibody (clone DM1A, catalogue #T9026) was utilized for western blot (1:5000) was obtained from Sigma-Aldrich (St. Louis, MO, USA). Anti-TAZ (catalog# HPA007415) utilized for western blot (1:2000) was obtained from Sigma-Aldrich (St. Louis, MO, USA). YAP (D8H1X XP; catalog #14074) (1:1000), GAPDH (D16H11;catalog#5174) XP (1:5000), Phospho-S6 Ribosomal Protein (Ser235/236)(D57.2.2E;catalog#4858)(1:2000), S6 Ribosomal Protein (5G10;catalog#2217)(1:2000), Phospho-Akt (Ser473) (D9E; catalog# 4060) XP (1:500), Akt (pan) (11E7;catalog# 4685) (1:500), PTEN (D4.3;catalog# 9188) XP (1:500), Phospho-YAP/TAZ(Ser127)(catalog#4911)(1:1000),Phospho-PRAS40(Thr246) (C77D7;catalog#2997) (1:2000), PRAS40 (D23C7;catalog# 2691)(1:2000), PI3 Kinase p110α (C73F8;catalog#4249)(1:1000), and Vinculin (E1E9V;catalog#13901)(1:5000) were obtained from Cell Signaling (Danvers, MA, USA) and used for western blot analysis. PDPK1(E-3) (catalog #sc-17765) (1:1000) used for western blots was obtained from Santa Cruz Biotechnology. RNA Pol II (4H8;catalog# 39097) (1:2000) was obtained from Active Motif.

The following antibodies were utilized for immunohistochemistry on formalin fixed paraffin embedded tissue. Anti phospho-AKT Ser473 (rabbit monoclonal D9E; Cell Signaling Technology catalog #4060 (Danvers, MA, USA). Anti-PTEN (rabbit monoclonal D4.3 XP; catalog# 9188) was obtained from Cell Signaling Technology. Anti-TAZ (catalog #BD560235). Anti-YAP (catalog #sc-15407). Dilutions utilized for immunohistochemistry are as follows phospho-AKT (1:50), PTEN (1:50), anti-TAZ (1:500), anti-YAP (1:100). Horseradish peroxidase-conjugated secondary antibodies were obtained from Dako. Horseradish peroxidase-conjugated secondary antibodies used for western blots were obtained from Bethyl Laboratories (catalog# A120-101P; A90-116P) and used at 1:5,000 or 1:10,000.

### Soft agar assay

The base layer of 0.5% agarose was plated into 6-well plates (2 mLs/well) and allowed to solidify for 1 hour. For SJCRH30 cells, 2 x 10^3^ cells/2 mL in 0.35% agarose was added to form the top layer. Plates were left in the hood and allowed to solidify at room temperature for 3 hours. Colonies were allowed to grow for 2-3 weeks at 37 °C and 5% CO_2_ before imaging. Each experiment was repeated at least twice.

### Clonogenic assay

100-500 cells were seeded in a 6 well plate format. The following day cells were treated with varying concentrations of LY294002, GDC-0941, or DMSO for 72 hours. After treatment, drug containing media was removed and replaced with fresh media and outgrowth was between 2-3 weeks. Colonies were then washed with 1X PBS twice and cells were fixed and stained with 0.5% crystal violet for 30 minutes. Plates were rinsed, dried, and colonies were subsequently counted. Survival fraction= (# of colonies formed post-treatment)/(# of cells seeded).

### Mouse studies

All animal procedures were performed with approval from the University of Iowa Institutional Animal Care and Use Committee.

#### Experiment 1 (Figure 4)

Both the *Pten* and *Trp53* floxed alleles were bred to homozygosity as previously described in (78). Offspring were genotyped by PCR for the presence of the *Trp53*^fl/fl^ and *Pten*^fl/fl^ alleles using gene-specific primers. *Wwtr1*^fl/fl^*Yap1*^fl/fl^ mice (119) were generously provided by Dr. Eric Olson (University of Texas Southwestern Medical Center, Dallas, TX) and crossed with *Trp53*^fl/fl^*Pten*^fl/fl^ mice to generate *Trp53*^fl/fl^*Pten*^fl/fl^*Wwtr1*^fl/fl^, *Trp53*^fl/fl^*Pten*^fl/fl^*Yap1*^fl/fl^, and *Trp53*^fl/fl^*Pten*^fl/fl^*Wwtr1*^fl/fl^*Yap1*^fl/fl^ mice. 8-12 week old mice were used for the study, with approximately 50% female and 50% male mice.

#### Experiment 2 (Figure 5)

For the everolimus study, 8-12 week old *Trp53*^fl/fl^*Pten*^fl/fl^ mice were injected with adeno-Cre virus as described above and separated into treatment and vehicle control arms composed of 16 mice, with approximately equal numbers of female and male mice used. 100 µl of 5mg/kgs Everolimus or vehicle (DMSO) was delivered by oral gavage on a weekly basis (79).

#### Experiment 3 (Figure 8)

For the combination drug study, *Prkdc^scid^Il2rg^tm1Wjl^/SzJ* (NSG) (RRID:IMSR_JAX:005557) were obtained from The Jackson Laboratory (Bar Harbor, ME). NSG mice ranging in age from 8-10 weeks old were utilized in the SJCRH30 xenograft/combination therapy study. 5×10^6^ RH30 cells in 100 µL 1X PBS were injected subcutaneously IK-930 (75 mg/kg daily) and everolimus (5 mg/kg weekly) were administered by oral gavage to the respective treatment arms of their study.

### Adenovirus injection

For the experiment evaluating sarcoma formation and overall survival in *Trp53*^fl/fl^*Pten*^fl/fl^, *Trp53*^fl/fl^*Pten*^fl/fl^*Wwtr1*^fl/fl^, *Trp53*^fl/fl^*Pten*^fl/fl^*Yap1*^fl/fl^, and *Trp53*^fl/fl^*Pten*^fl/fl^*Wwtr1*^fl/fl^*Yap1*^fl/fl^ mice, 20 mice were included for each genotype and injected with Ad5 CMV-Cre recombinase virus, while 4 mice (1 from each genotype) were injected with Ad5 CMV-eGFP virus for a total of 84 mice examined. No tumors developed in Adeno CMV-eGFP mice control mice. The experiment was repeated. *2P*, *2PY*, *2PW* and *2PWY* mice were first anesthetized in a chamber using 3% isoflurane in order to facilitate accurate injections. Ad5 CMV-eGFP (control animals) or Ad5 CMV-Cre recombinase (experimental animals) obtained from The University of Iowa Viral Vector Core was injected intramuscularly into the posterior left quadriceps. Injections were performed using 1×10^9^ pfu in a total of 20 μL injection volume (viral stocks were diluted to final concentration using 1x phosphate-buffered saline). Successful injections were confirmed either by manual stabilization of the quadriceps muscle and palpation of the femur with the needle tip prior to injection in the intramuscular group.

### Mouse examinations

Once tumors were palpable, they were monitored/measured every 2-3 days until completion of the study. Tumor volume was estimated using the equation V = 0.5 x L x W^2^ (V=volume, L=length, W=width). Euthanasia and dissections were performed when mice reached pre-determined end points, including 20% loss of starting body weight, primary tumor size 1.5 cm (Experiment 1 – see above Mouse studies section) or 2.0 cm (Experiments 2, 3 – see above Mouse studies section) at the site of injection, decreased mobility, lethargy/lack of grooming, or other gross morbidity. Control mice were also euthanized at the completion of the study. Primary tumors and other appropriate tissues were collected and incubated in 10% neutral buffered formalin and embedded in paraffin prior to sectioning.

### RNA-Seq

Total RNA was extracted using Trizol Reagent (Ambion-Life Technologies). Total RNA was isolated using the PureLink RNA mini kit (Invitrogen-ThermoFisher Scientific). On-column DNase (Invitrogen) treatment was performed according to manufacturer’s instructions. Transcription profiling using RNA-Seq was performed by the University of Iowa Genomics Division using manufacturer recommended protocols. Initially, 500 ng of DNase I-treated total RNA. The enriched total RNA pool was then fragmented, converted to cDNA and ligated to sequencing adaptors containing indexes using the Illumina TruSeq Stranded Total RNA w/ RiboZero sample preparation kit (Cat. No. RS-122-2201, Illumina, Inc, San Diego, CA). The molar concentrations of the indexed libraries were measured using the 2100 Agilent Bioanalyzer (Agilent Technologies, Santa Clara, CA) and combined equally into pools for sequencing. The concentration of the pools was measured using the Illumina Library Quantification Kit (KAPA Biosystems, Wilmington, MA) and sequenced on the Nova Seq 6000 genome sequencer using 150 bp paired-end SBS chemistry.

Barcoded samples were pooled and sequenced using an Illumina NovaSeq 6000 in the Iowa Institute of Human Genetics (IIHG) Genomics Core Facility. Paired-end reads were demultiplexed and converted from the native Illumina BCL format to FASTQ format using a custom python workflow wrapper to Illumina’s ‘bcl2fastq’ conversion utility. FASTQ data were processed with nf-core/rnaseq (v3.10), a best-practices pipeline available at the open-source ‘nf-core’ project (120) (https://nf-co.re/, Nextflow version 22.10) (121) running on the Argon HPC resource at the University of Iowa. The pipeline was invoked with command:

> nextflow run nf-core/rnaseq -r 3.10 -profile argon --igenomes_base /Users/mchiment/igenomes/references --input samplesheet_NOSKEL4.csv --outdir nf_NOSKEL4 --genome GRCm38 -- -- save_merged_fastq FALSE --skip_markduplicates TRUE --skip_preseq TRUE --skip_dupradar TRUE -- skip_stringtie TRUE.

Reads from the samples were aligned against the mouse reference genome ‘GRCm38’ using the STAR aligner (122) and quantified with ‘salmon’ (123). Samtools (124) was used in conjunction with Qualimap (125) (126) and MultiQC {10.1093/bioinformatics/btw354} to inspect alignment results. Length-normalized gene-level counts from the STAR/salmon pipeline were used for differential gene expression analysis with DESeq2 (127). Bioconductor package ‘PCAExplorer’ (128) was used for exploratory analysis. The DE gene lists were analyzed using AdvaitaBio’s iPathwayGuide (https://www.advaitabio.com/ipathwayguide) (129) (130) (131) (132). This software analysis tool implements the ‘Impact Analysis’ approach that takes into consideration the direction and type of all signals on a pathway. The raw FASTQ files and associated metadata have been made available for download at GEO accession: GSE274982.

### Immunofluorescence

For both SJCRH30, integrated density of nuclear signal for YAP or TAZ (ImageJ software) was taken for 10 cells from four different fields. The integrated density for LY294002 treated cells were compared relative to DMSO for final graphical presentation (GraphPad Prism). Since TAZ or YAP signal was more diffuse in both the nuclear and cytoplasmic compartments in the A204 cell line, the integrated density was taken for the entire cell. For both SJCRH30 and A204 cell lines, the fluorescent signal in approximately 100 cells for DMSO and LY204002 treated conditions were counted as predominantly nuclear (N>C), cytoplasmic (C>N), or both (N=C). Percentages for the respective counts were calculated relative to the total number of cells counted.

### Mouse Studies

NOD scid gamma (NSG) mice were obtained from The Jackson Laboratory (Bar Harbor, ME, USA) and carry the strain NOD.Cg-Prkdcscid Il2rgtm1Wjl/SzJ (RRID:IMSR_JAX:005557). All animal work was approved by the University of Iowa Institutional Animal Use and Care Committee. SJCRH30 cells containing (5 × 10^6^ cells/100 µL PBS) were injected subcutaneously into the NSG mice. The mice were 8-10 weeks of age at the time of injection.

### Luciferase reporter assay

SJCRH30 and A204 cells were transduced with TEAD Luciferase (pLV-8xTEADrep-Luc2-pgk-Rluc-2A-Neo) reporter construct plasmid (Ikena Oncology, Boston, MA) to generate stable cell lines. A Dual luciferase reporter assay (Promega, Madison, WI, USA) detecting both firefly luciferase and *Renilla* luciferase activity was performed in a six-well plate (biological triplicates) with a seeding density of 5×10^5^/well. Three days after the drug treatment, the cells were collected and lysed, and extracts were assayed in technical triplicate for firefly and *Renilla* luciferase activity using the Dual Luciferase Reporter Assay System (Promega, Madison, WI, USA) and a Synergy H1 Hybrid Multi-Mode Microplate Reader (Biotek, Winooski, VT, USA). Each experiment was repeated at least twice. Due to extremely high luminescence values of *Renilla* control for A204 cells, the firefly luciferase activity values were considered alone after being normalized with the untreated group (DMSO).

### Proliferation assay

1000 cells/well from the SJCRH30 and A204 cell lines were plated in a 96 well plate. Drugs were added and proliferation was measured with the Dojindo Assay Cell Counting Kit 8 (Dojindo Molecular Technologies, Rockville, MD) according to the manufacturer’s instructions. Absorbance was measured using the BioTek: Synergy H1 Hybrid Reader (Biotek). Each experiment was repeated at least twice.

### Statistics

For soft agar and clonogenic assays, statistical significance was evaluated student’s unpaired two-tailed *t*-test. For Kaplan-Meier curves significance was determined by Log-rank (Mantel-Cox) test. For tumor initiation, volume tripling, and weight, statistical signification was determined using the unpaired two-tailed *t-*test with Welch’s correction to account for different sample sizes between groups. For immunofluorescence assays, statistical significance was evaluated using the student’s unpaired two-tailed *t-*test. To assess enrichment on RNA-Seq data sets, hypergeometric analyses and Matthews correlation coefficient (MCC) calculations were carried out in R, version 4.4.1. The upper tail of the hypergeometric distribution was calculated using the phyper() function. Each experiment was repeated at least twice. Error bars were used to define one standard deviation. For all panels, ****p<0.0001, *** p<0.001, ** p<0.01, *p<0.05.

## Acknowledgements

This work was supported by a University of Iowa Sarcoma Multidisciplinary Oncology Group pilot award (M.R.T), by a grant from the Veterans Health Administration Merit Review Program 1 I01 BX003644-01 (M.R.T), by grants from the National Institutes of Health, National Cancer Institute 1 R01 CA237031-01A1 (M.R.T) and 1 R01 CA237031-01A1S1 (K.C.G., M.R.T), by an award from the University of Iowa Stead Family Scholars Program (M.R.T), by a Center for Biocatalysis and Bioprocessing Predoctoral Fellowship (A.A.K), and by an NCI Core Grant P30 CA086862 (University of Iowa Holden Comprehensive Cancer Center). We would like to acknowledge Dr. Eric Olson (UT Southwestern, Dallas, TX) for contributing *Wwtr1*^fl/fl^*Yap1*^fl/fl^ mice. We would also like to acknowledge Ikena Oncology (Boston, MA) for providing IK-930.

## Author contributions

Conceptualization, M.R.T.; Methodology, M.S.C., P.B., and M.R.T.; Software, M.S.C., P.B.; Formal analysis, M.S.C., P.B., and M.R.T.; Investigation, K.C.G., A.A.K., K.G., S.S., N.S., G.D., C.F., N.M., Y.D., S.Y., M.L., P.B., M.S.C., and M.R.T.; Resources, M.D.H., P.B., and M.R.T.; Writing-Original Draft, K.C.G., A.A.K., and M.R.T.; Writing-Review & Editing, K.C.G., A.A.K., and M.R.T.; Supervision, M.R.T.; Funding Acquisition, K.C.G., A.A.K., and M.R.T.

### Data and Code Availability

The accession number for the RNA-Seq data reported in this paper for mouse tumors is GEO: GSE274982.

---

**Figure S1:**
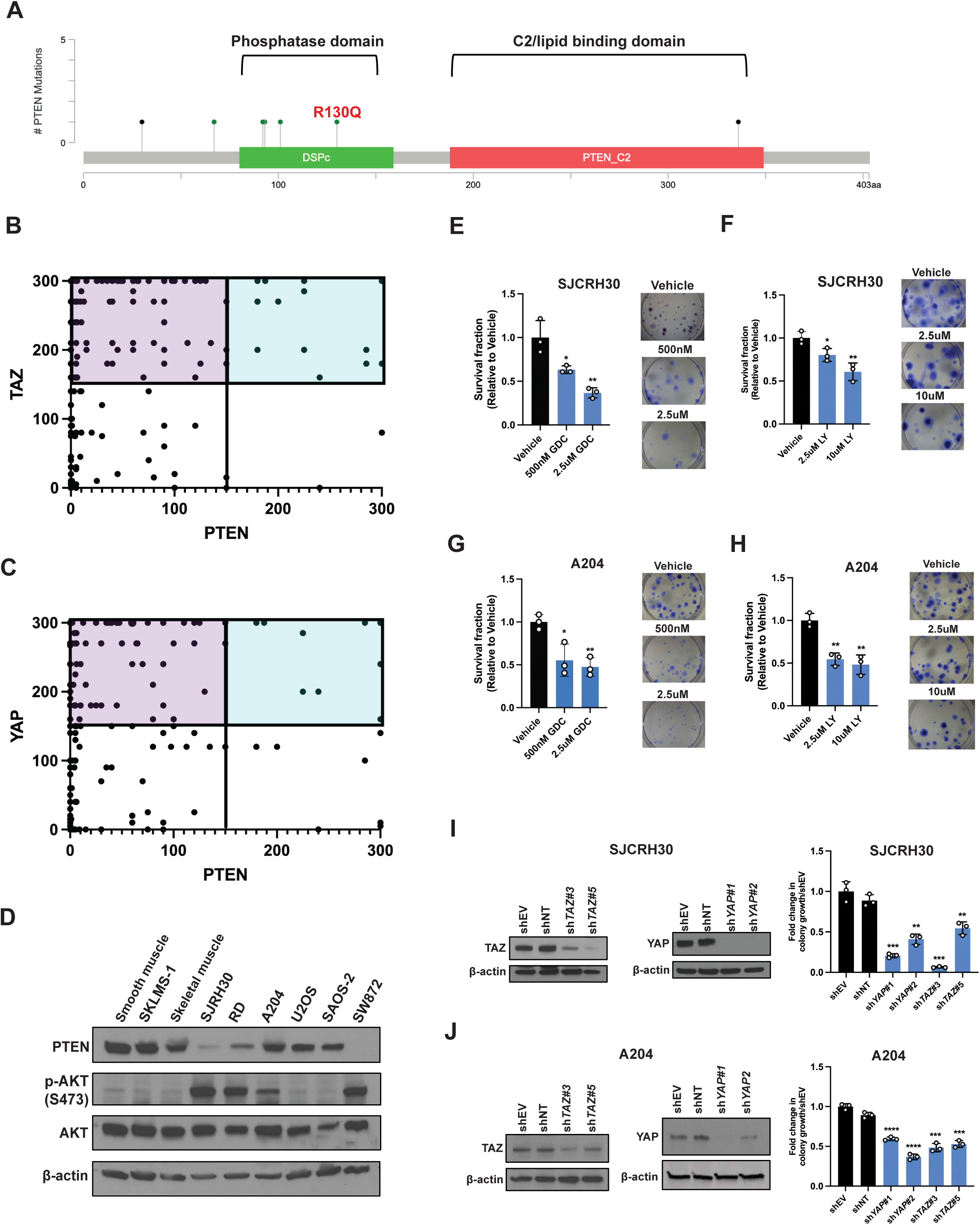
Initial assessment of PI3K activation in clinical samples and sarcoma cell lines. **(A)** PTEN point mutations within the phosphatase and lipid binding domains in sarcomas (n=253). R130Q (red) missense mutation in UPS known to lead to defective phosphatase activity. Data from TCGA was analyzed using the cBioportal online software. (**B**) Scatter plot showing H-scores for TAZ plotted vs. H-scores for PTEN. Tumors that demonstrated activated TAZ (high TAZ) are located within the upper two quadrants (upper left quadrant – low PTEN/high TAZ; upper right quadrant – high PTEN/high TAZ). (**C**) Scatter plot showing H-scores for YAP plotted vs. H scores for PTEN. Tumors that demonstrated activated YAP (high YAP) are located within the upper two quadrants (upper left quadrant – low PTEN/high YAP; upper right quadrant – high PTEN/high YAP). (**D**) Western blot analysis of sarcoma cell lines and primary cell cultures. Cells were temporarily serum starved for 4hrs. Where appropriate, PI3K activation (p-AKT) and PTEN expression are compared to normal cellular controls such as smooth muscle cells, skeletal muscle cells. (**E**-**F**) Clonogenic outgrowth analysis for SJCRH30 cells with PI3K class I inhibitors GDC-0941 (**E**) and LY294002 (**F**) after treatment for 72 hours. (**G**-**H**) Clonogenic outgrowth analysis for A204 cells with PI3K class I inhibitors GDC-0941 (**G**) and LY294002 (**H**) after treatment for 72 hours. For clonogenic assays, statistical significance was evaluated using student’s unpaired two-tailed *t*-test compared to DMSO. Each experiment was repeated at least twice. (**I**) Western blot and graph showing decrease of anchorage independent growth in soft agar in SJCRH30 cells after knock-down of TAZ or YAP with two different shRNAs. (**J**) Western blot and graph showing decrease of anchorage independent growth in soft agar in A204 cells after knock-down of TAZ or YAP with two different shRNAs. For soft agar assays, statistical significance was evaluated using student’s unpaired two-tailed *t*-test. Each experiment was repeated at least twice. For all panels, ****p<0.0001, *** p<0.001, ** p<0.01, *p<0.05.

**Figure S2:**
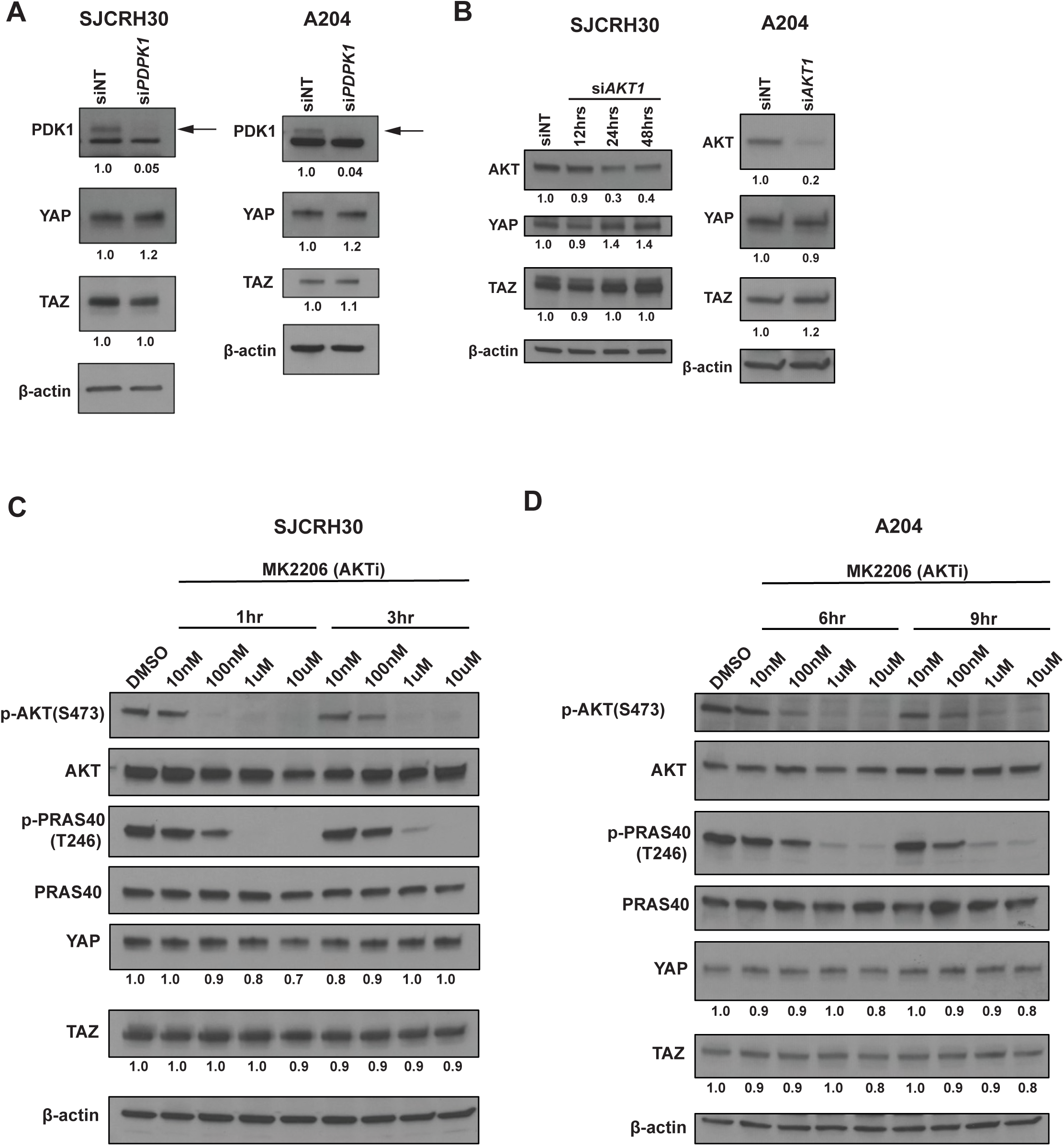
PI3K regulates TAZ and YAP independent of PDK1 and AKT1 in sarcomas. **(A)** siRNA (pooled) mediated knockdown of PDK1 for 48 hours in SJCRH30 and A204 cells; western blot for PDK1, YAP, and TAZ. **B**) siRNA (pooled) mediated knockdown or AKT1; western blot for AKT, YAP, and TAZ. (**C, D**) Pharmacological inhibition of AKT with MK2206 in the SJCRH30 (**C**) and A204 (**D**) cell lines for the indicated timepoints. TAZ and YAP quantitation normalized to β-actin and compared to DMSO (vehicle) control. Experiments were repeated at least twice.

**Figure S3:**
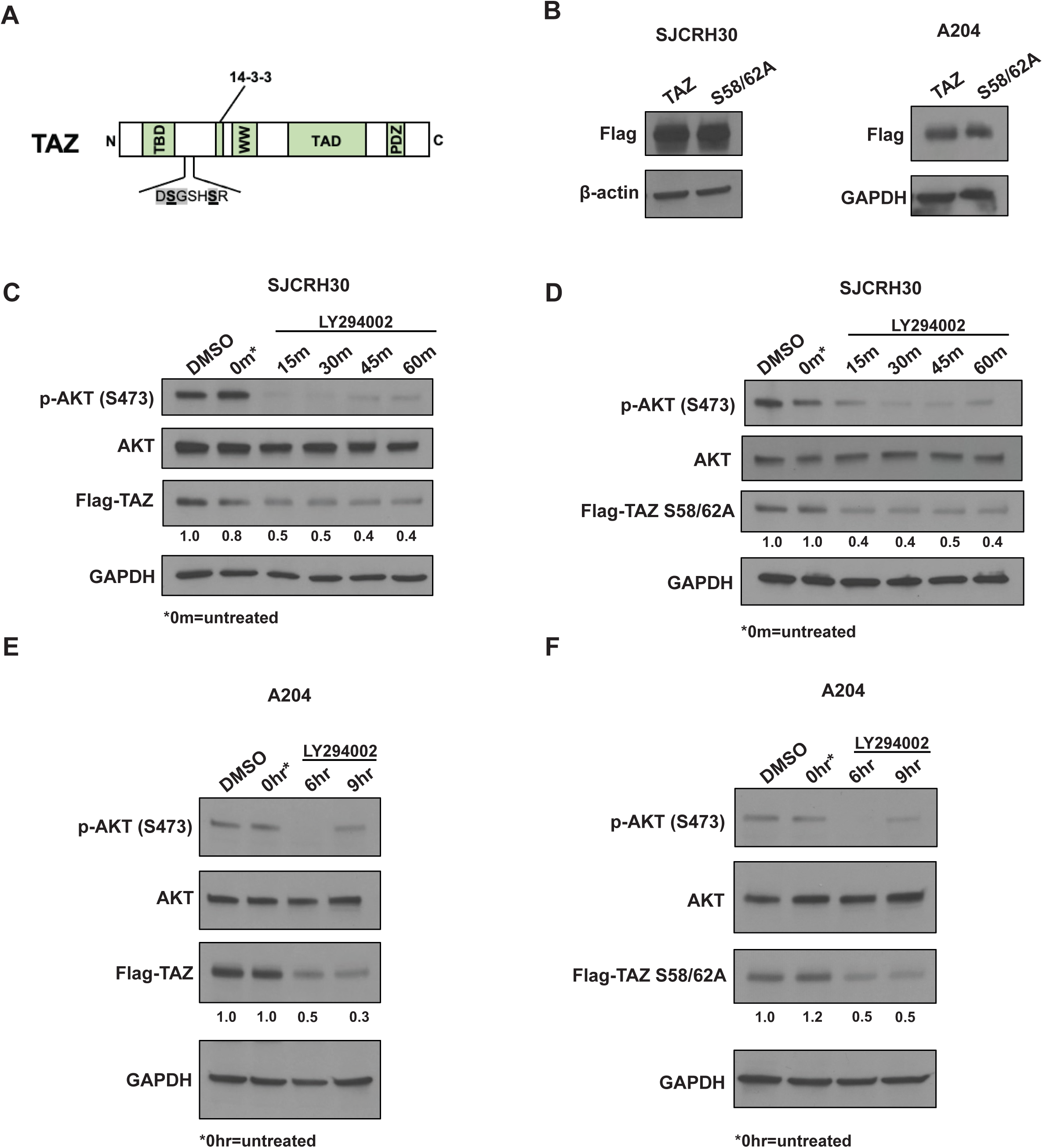
Regulation of TAZ is GSK3β independent in SJCRH30 and A204 cells. (**A**) TAZ N- terminal phosphodegron (DSGXXS motif) for SCFβTrCP dependent degradation. Target serines 58 and 62 are highlighted in bold and underlined. TEAD Binding Domain (TBD), 14-3-3 binding motif (14-3-3), WW domain (WW), Transactivating domain (TAD), PDZ binding motif (PDZ). (**B**) Western blots for SJCRH30 and A204 stable lines expressing Flag-TAZ and Flag-TAZ S58/62A showing equivalent levels of expression of the wild-type and mutated protein in both cell lines. β-actin and GAPDH were used as loading controls. (**C, D**) SJCRH30 cells stably expressing Flag-TAZ (**C**) or GSK3β resistant Flag-TAZ S58/62A (**D**) were treated with 30 µM LY294002. (**E, F**) A204 cells stably expressing wild-type Flag-TAZ (**E**) or GSK3β resistant Flag-TAZ S58/62A (**F**) were treated with 30 µM LY294002. Experiments were repeated at least twice. Flag-TAZ and Flag-TAZ S58/62A protein levels were quantitated and normalized to β-actin or GAPDH and compared to vehicle (DMSO) controls.

**Figure S4:**
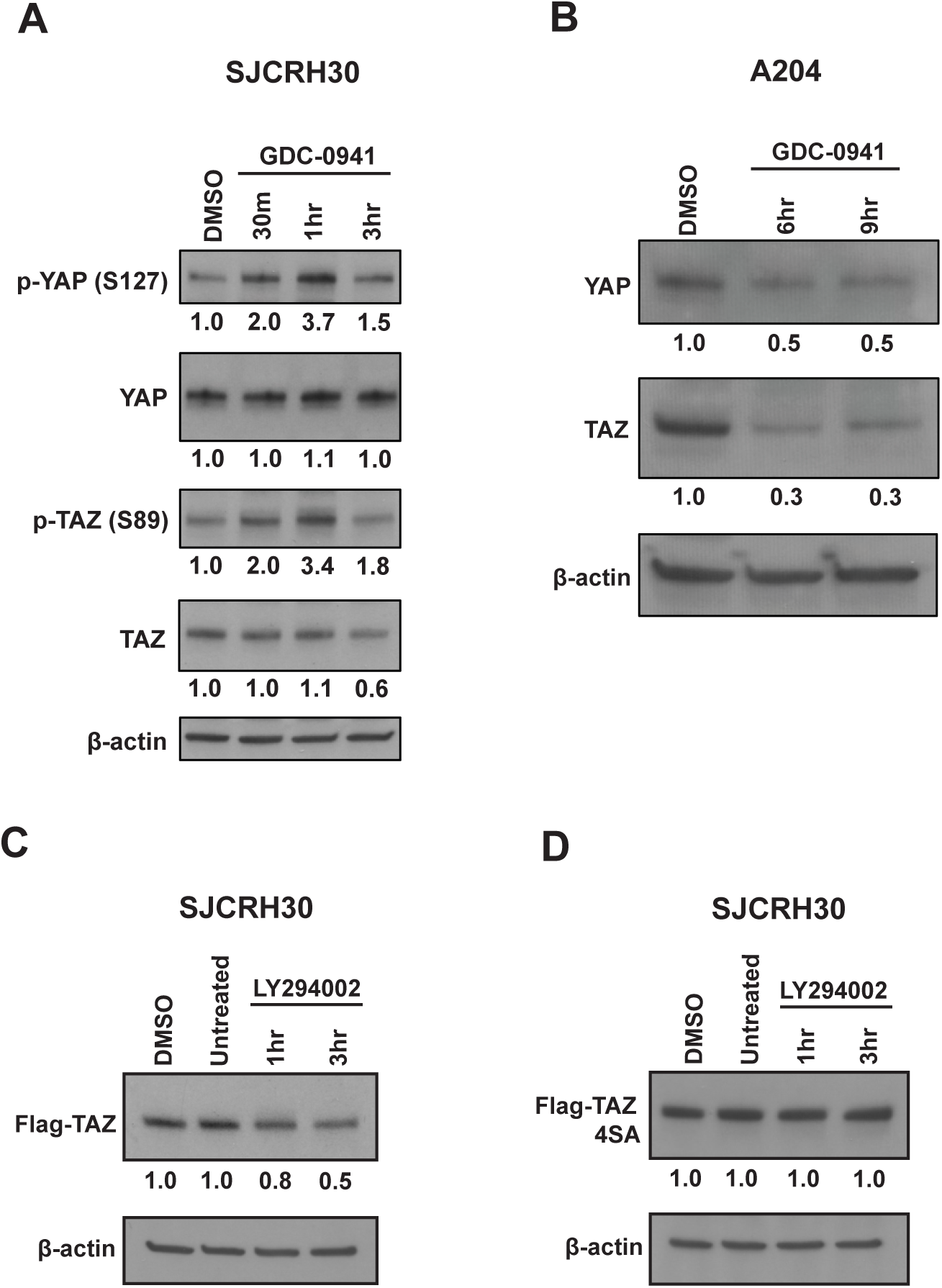
PI3K inhibition promotes LATS-mediated degradation of TAZ/YAP in SJCRH30 and A204 cells. (**A**) SJCRH30 cell line treated with 10 µM GDC-0941 for the indicated timepoints; p-YAP (S127), and p-TAZ (S89) were quantitated relative to total YAP and TAZ, respectively, and compared to vehicle (DMSO) treated cells. (**B**) A204 cell line treated with 10 µM GDC-0941 for the indicated time points. Western blot for total YAP and TAZ. (**C,D**) SJCRH30 cells expressing Flag-TAZ (**C**) or Flag-TAZ 4SA (**D**) were treated with 60 µM LY294002 for the indicated time points.

**Figure S5:**
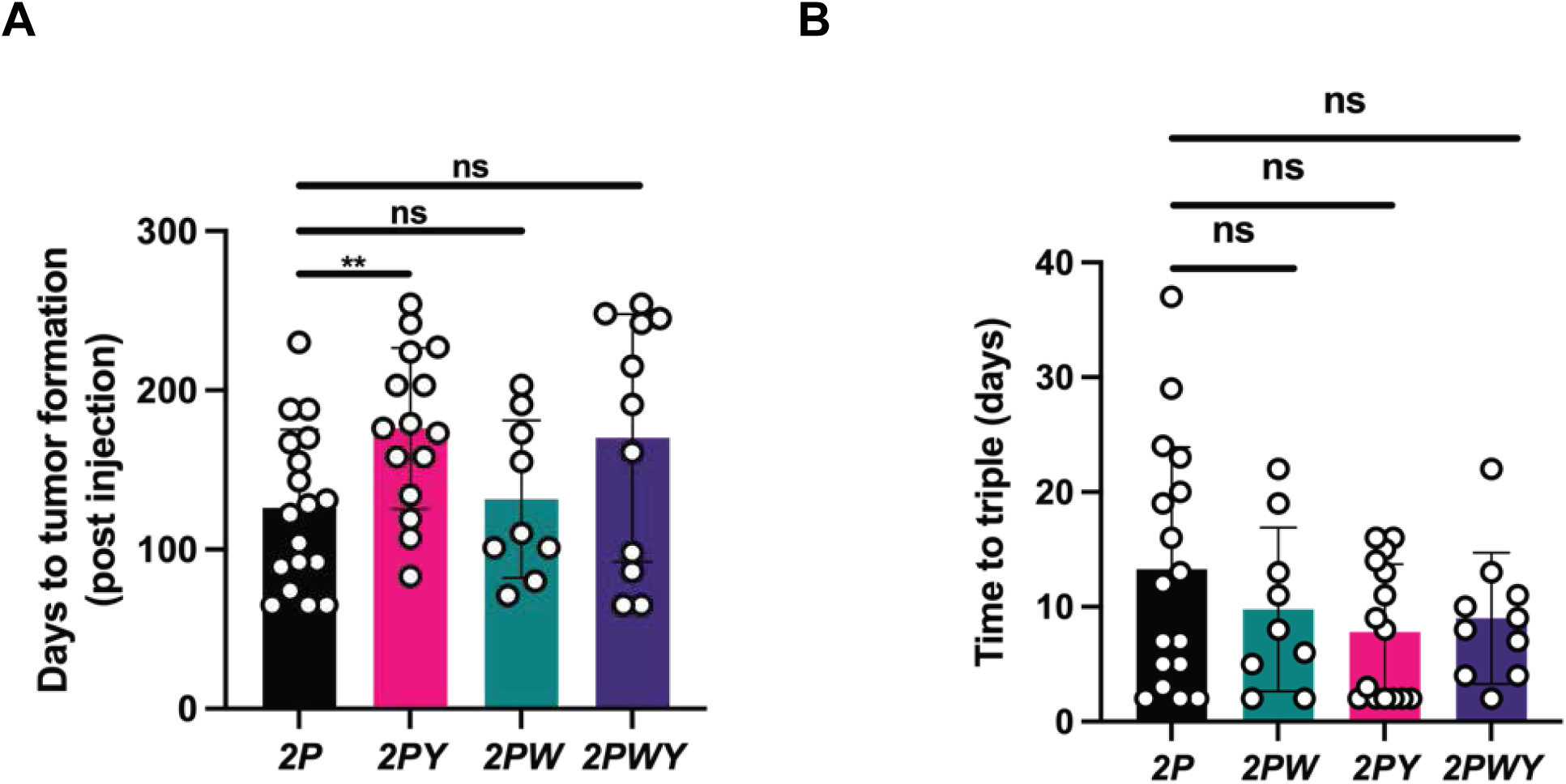
Metrics of tumor growth in *2P*, *2PW*, *2PY*, and *2PWY* mice. **(A**) Days to tumor formation. (**B**) Days to tripling of tumor volume.

**Figure S6:**
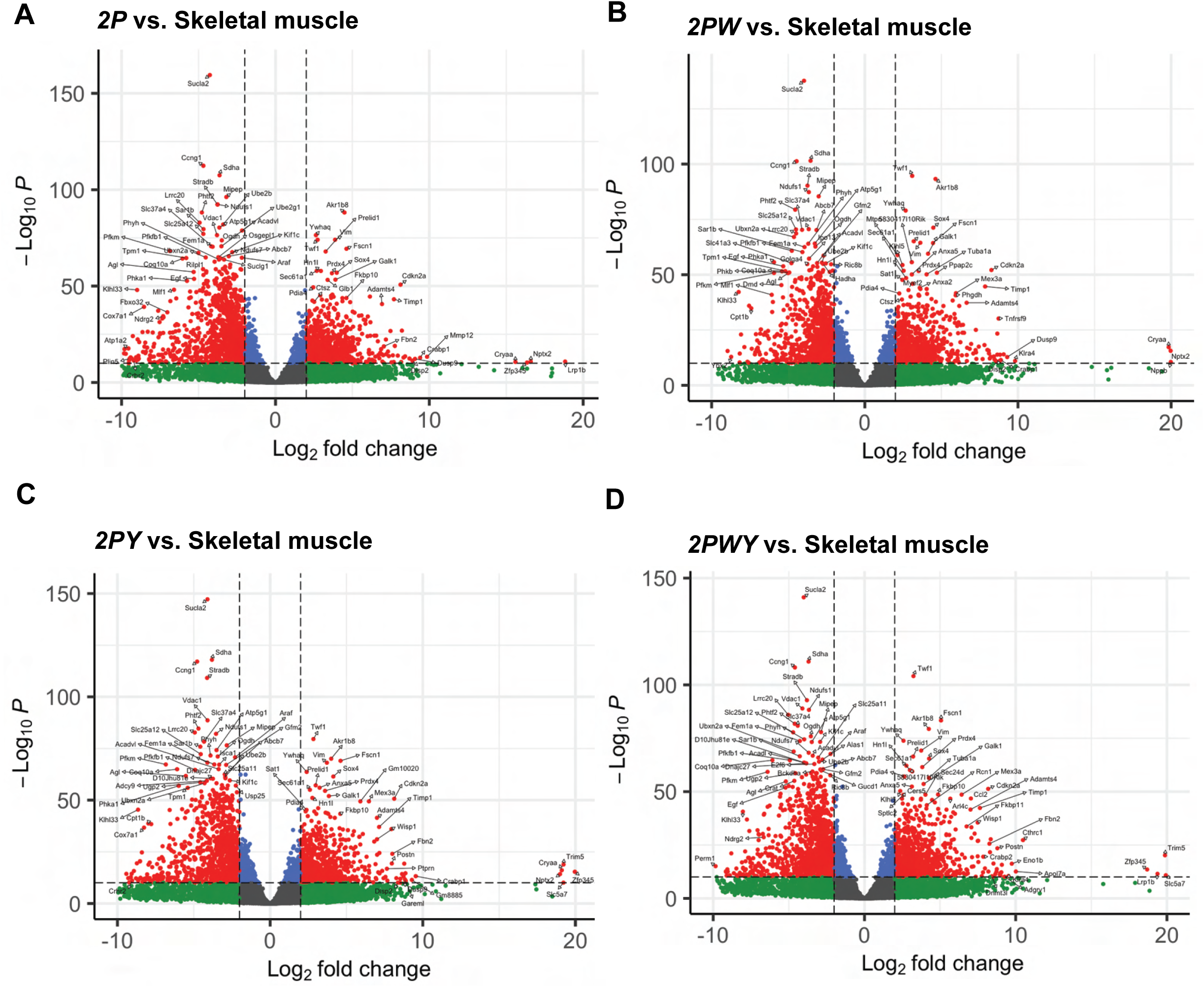
Total RNA-seq analysis of *2P*, *2PW*, *2PY*, and *2PWY* tumors. (**A-D**) Volcano plots from total RNA-seq data for *2P* (**A**)*, 2PW* (**B**), *2PY* (**C**), and *2PWY* (**D**) tumors normalized to murine skeletal muscle.

**Figure S7:**
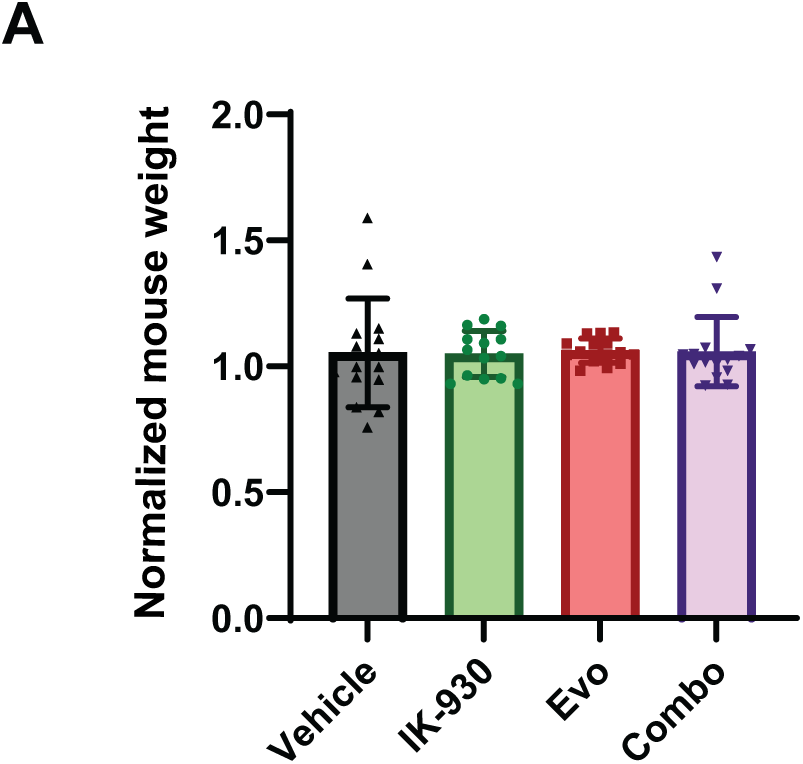
Effects of everolimus/IK-930 combination therapy on mouse weight. (**A**) Normalized weight (post-treatment weight normalized to pre-treatment weight) of SJCRH30 xenografted mice in different treatment arms of the study.

**Table S1:**
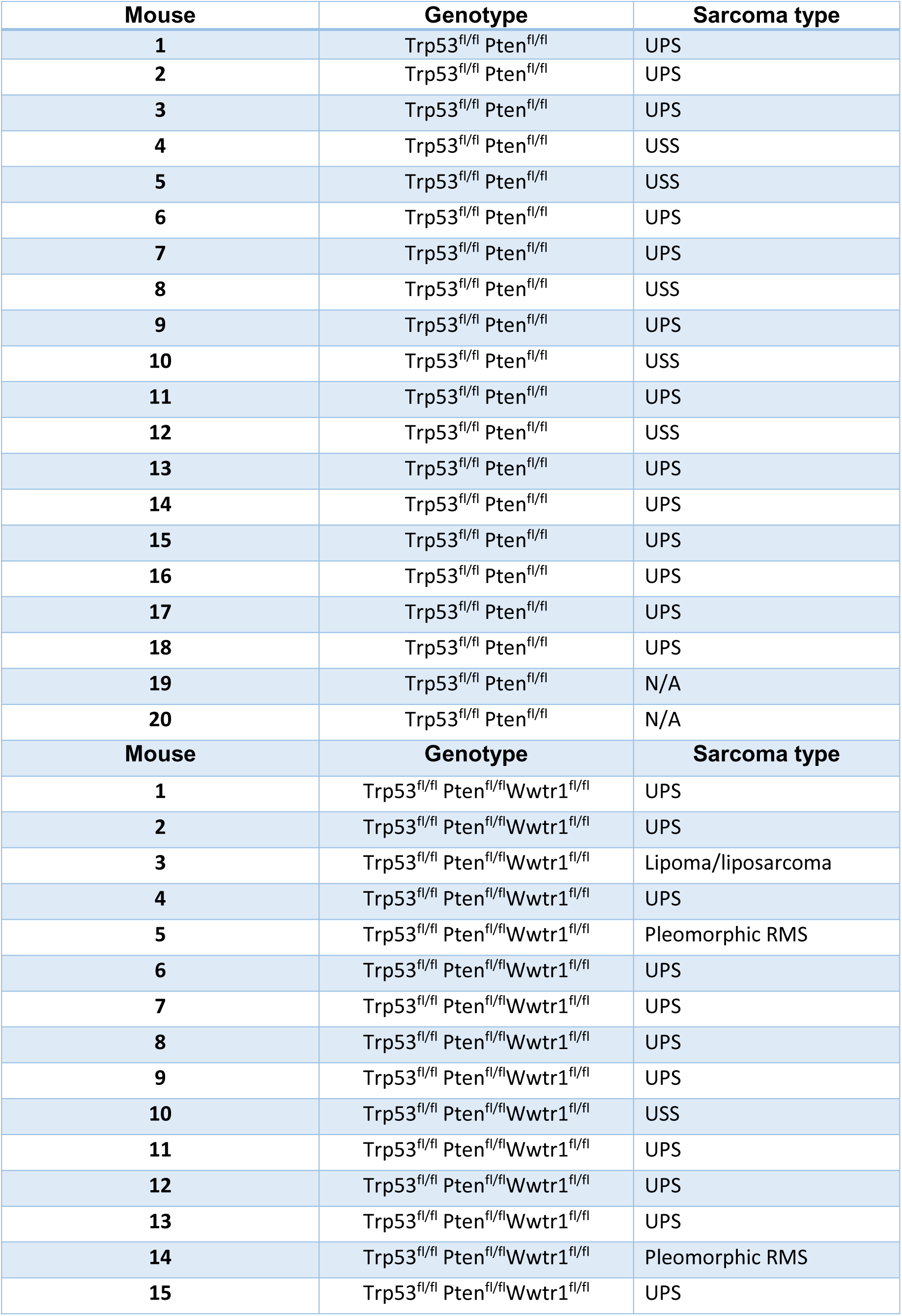
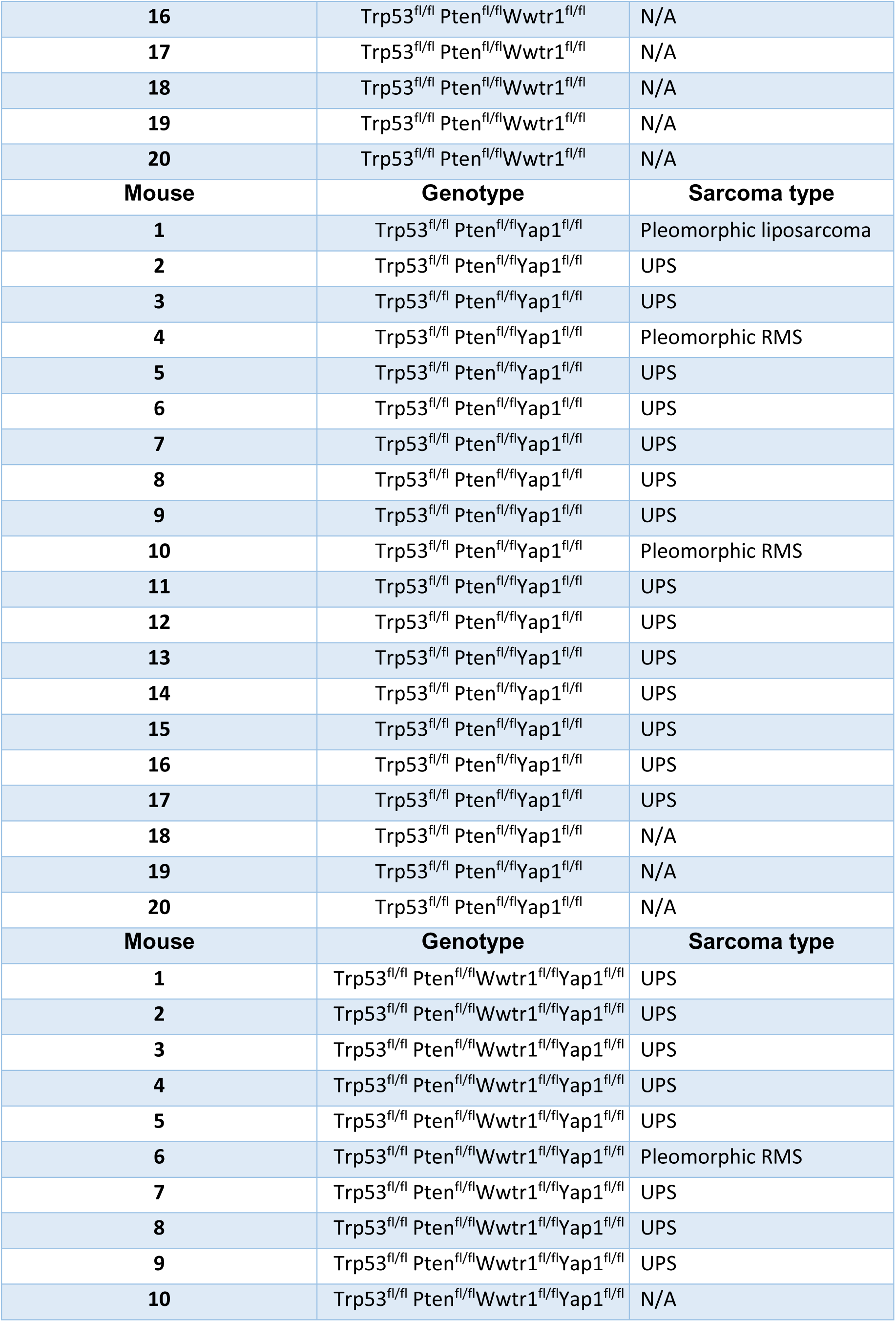
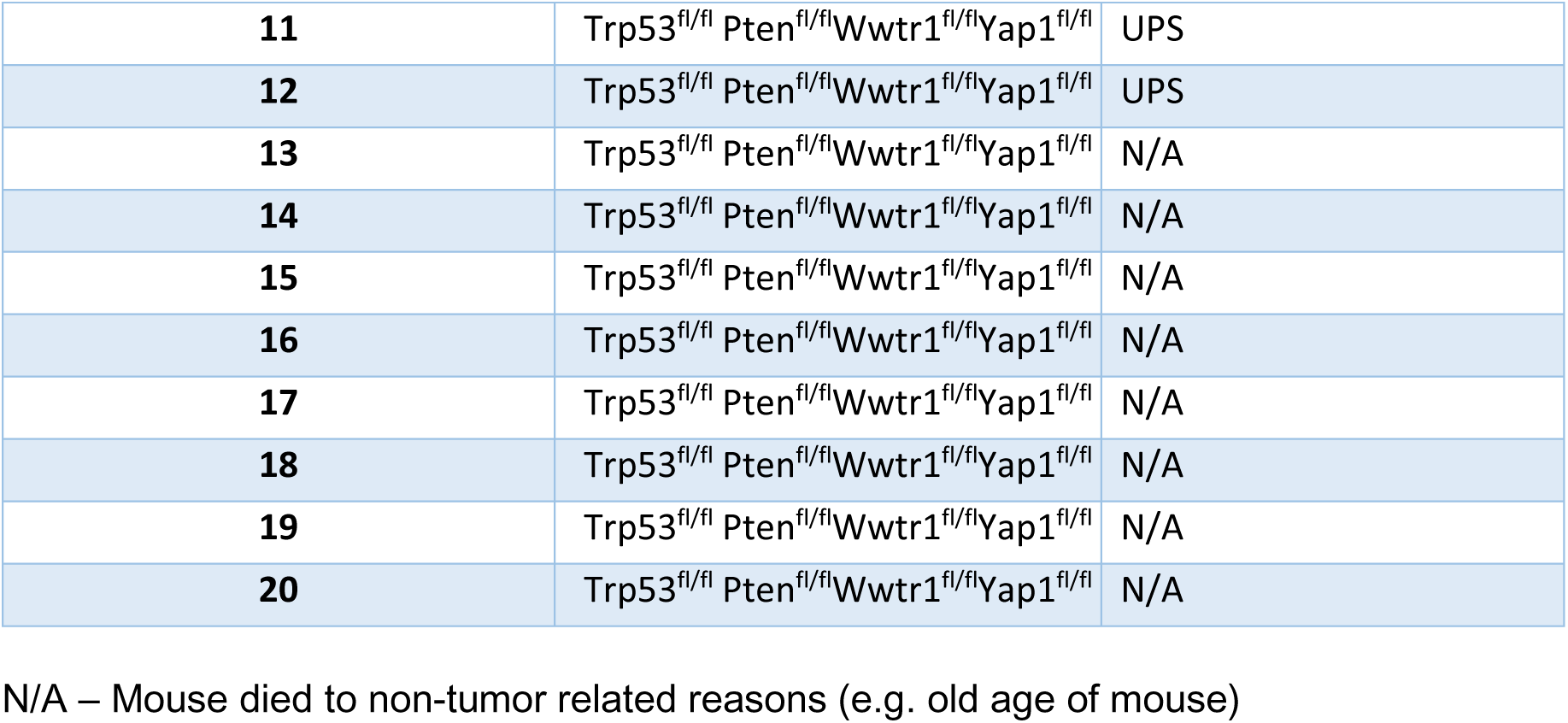
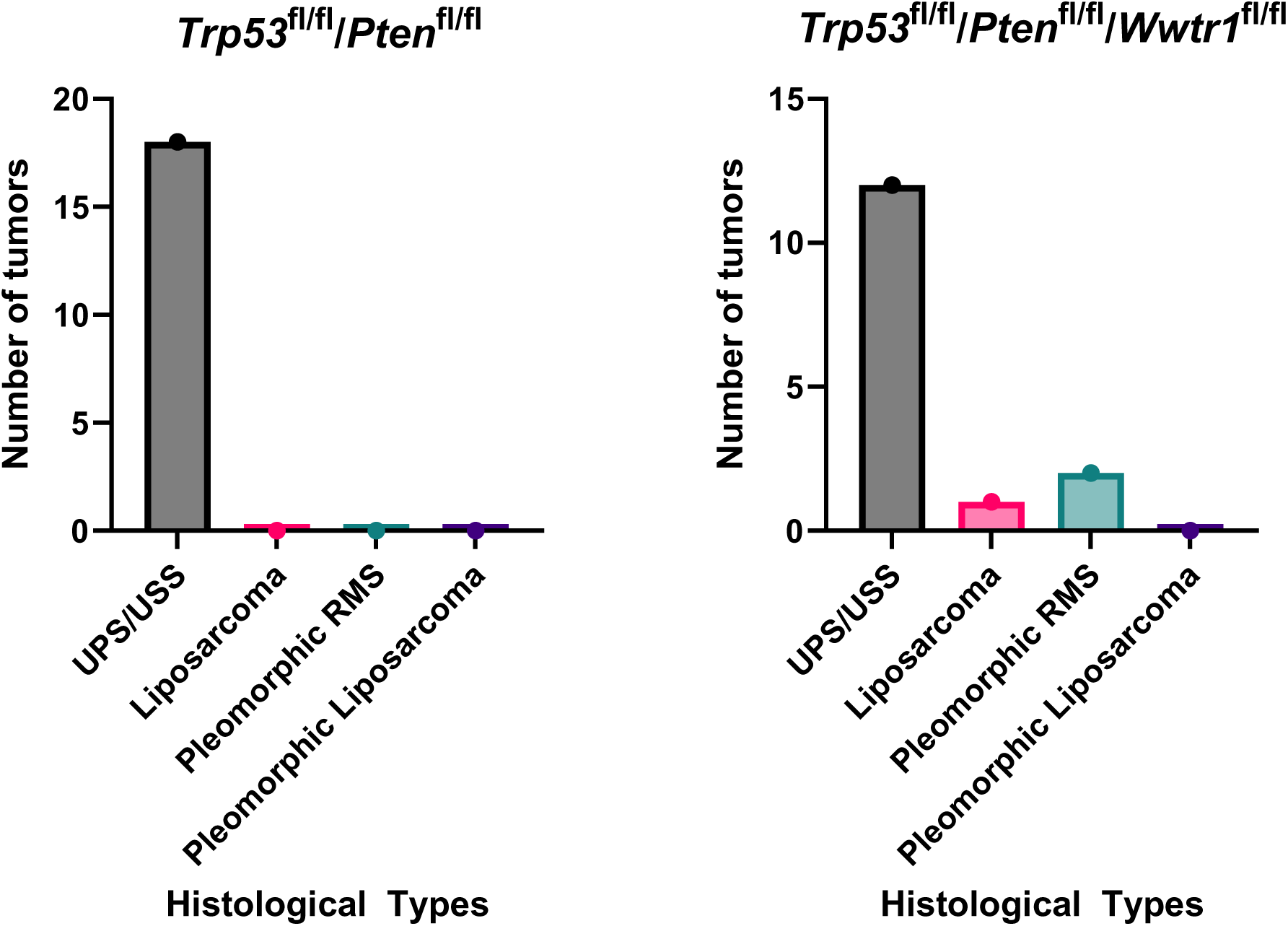
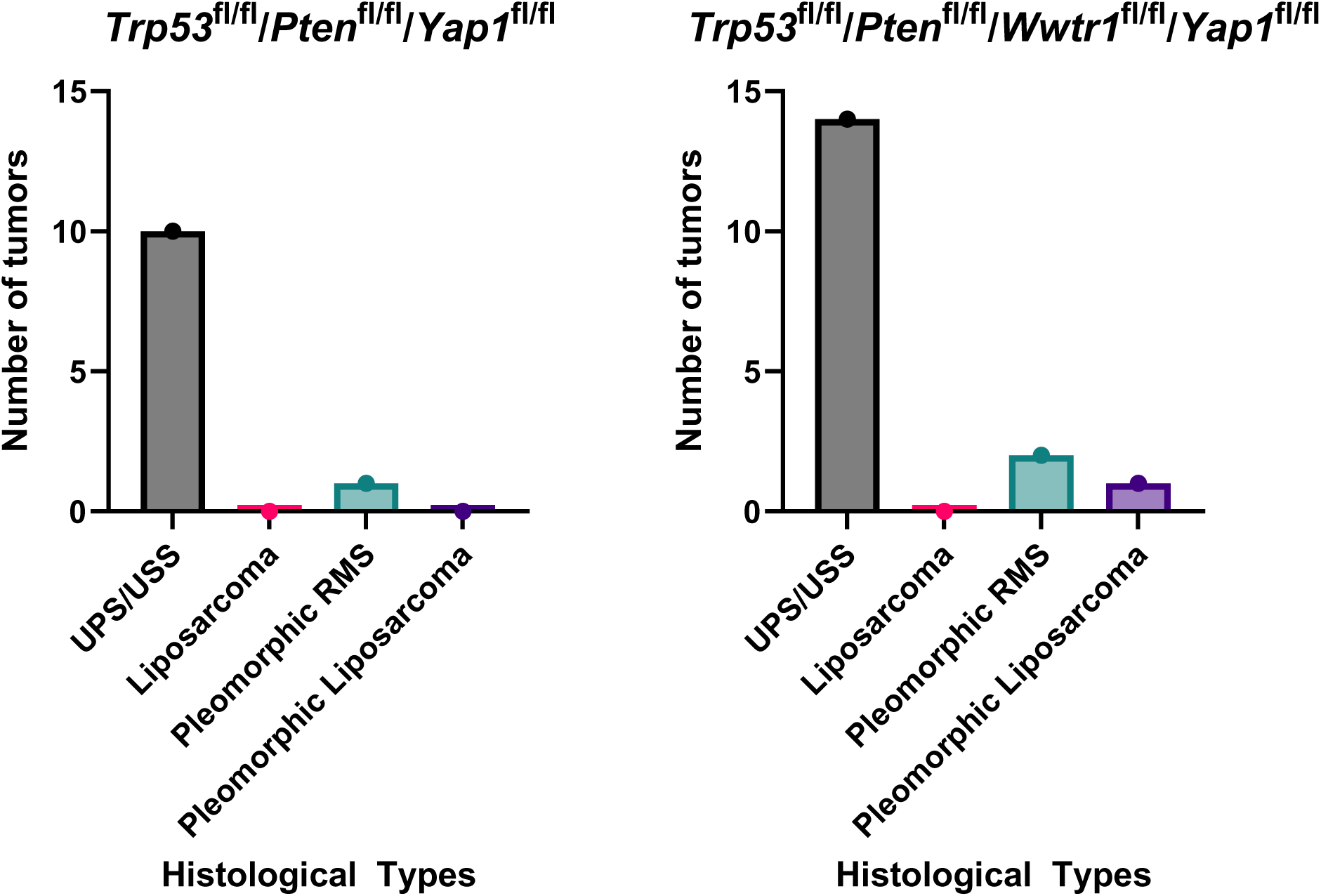
Histological classification of mouse tumors from *2P*, *2PW*, *2PY*, and *2PWY* mice.

**Table S2:**
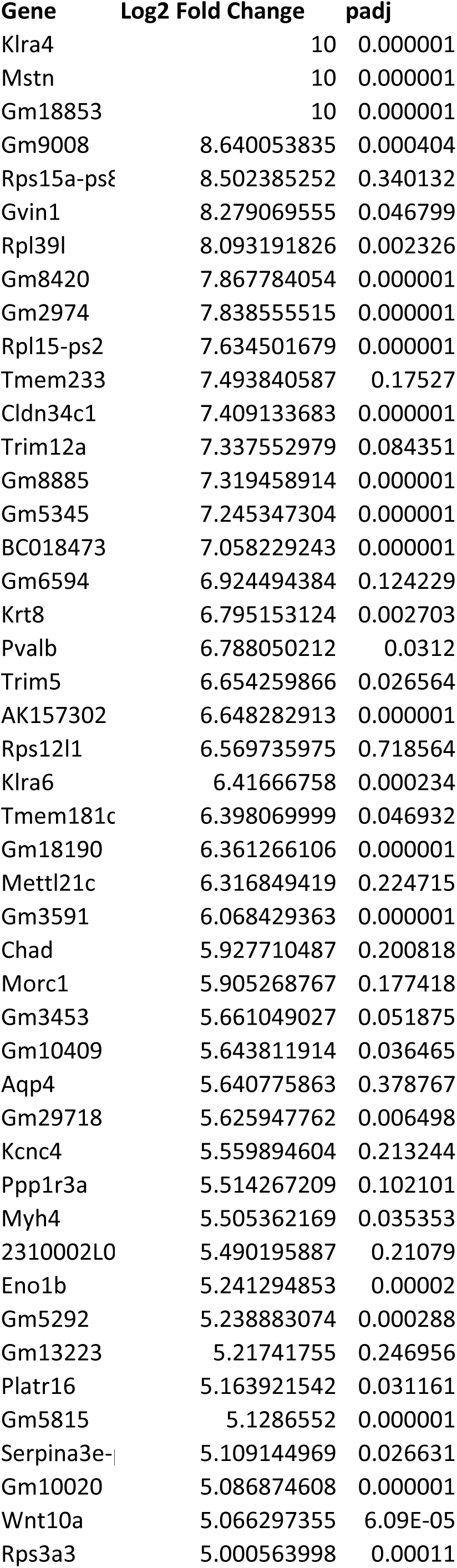
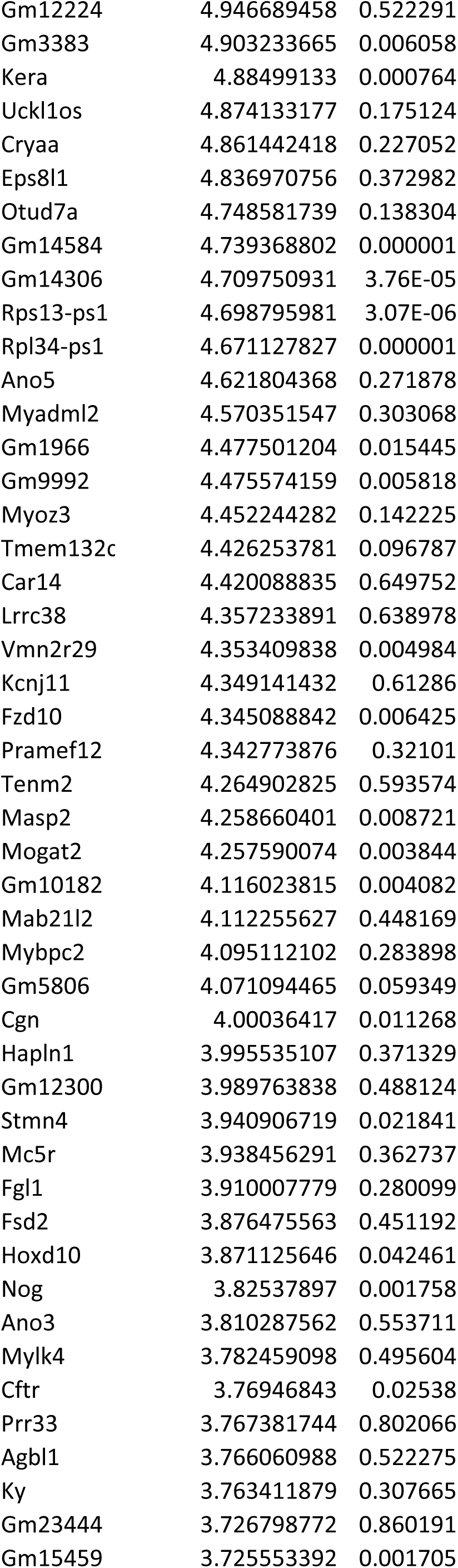
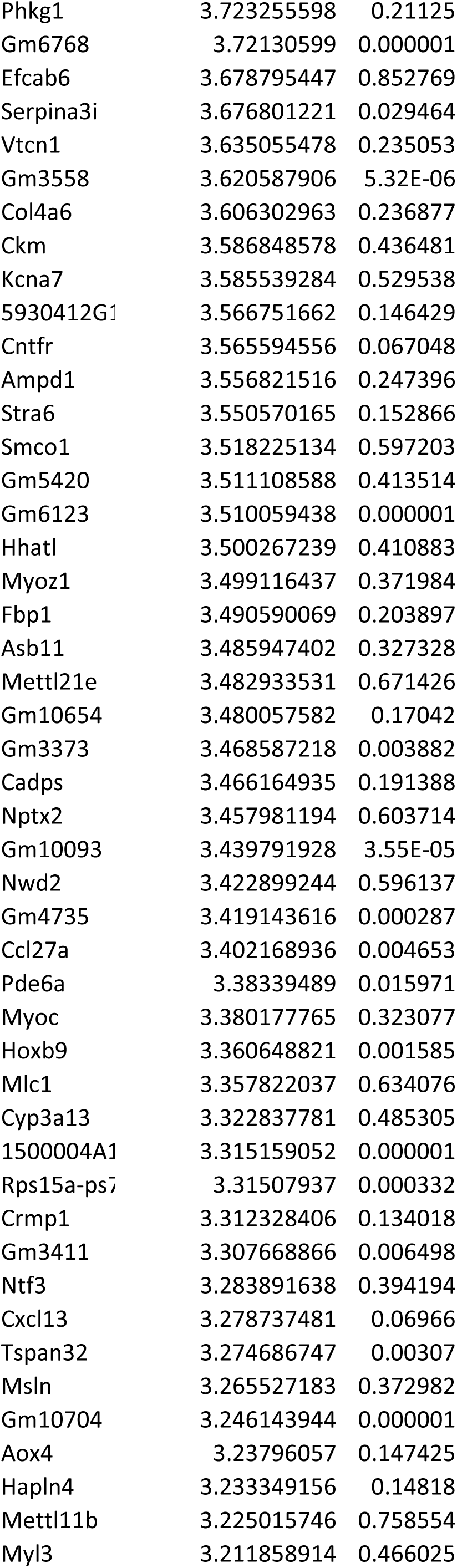
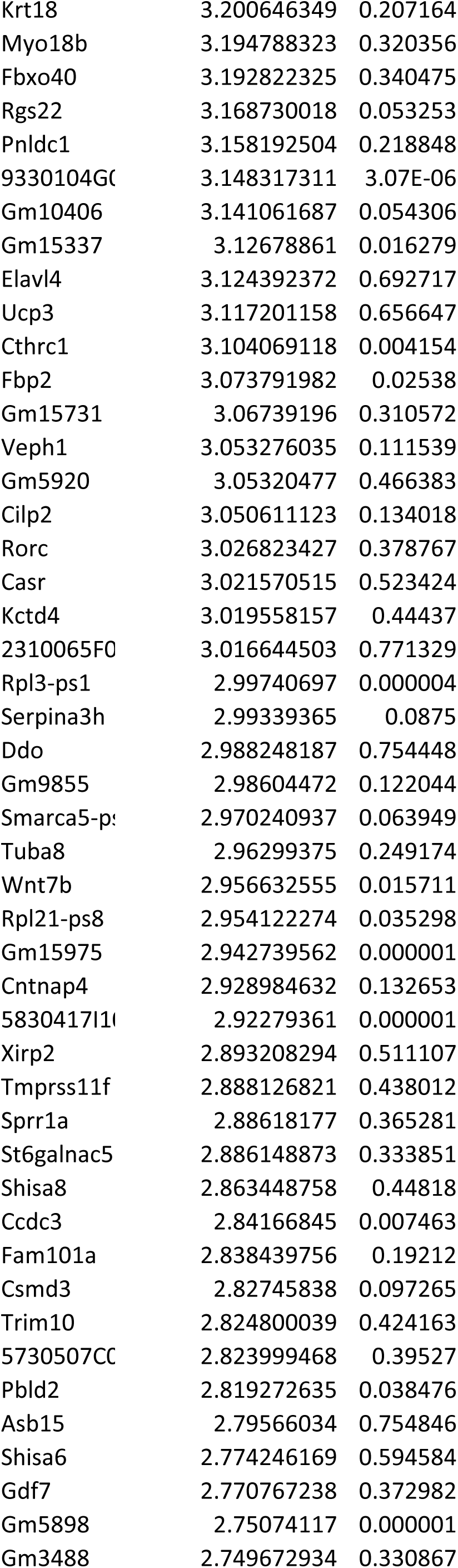
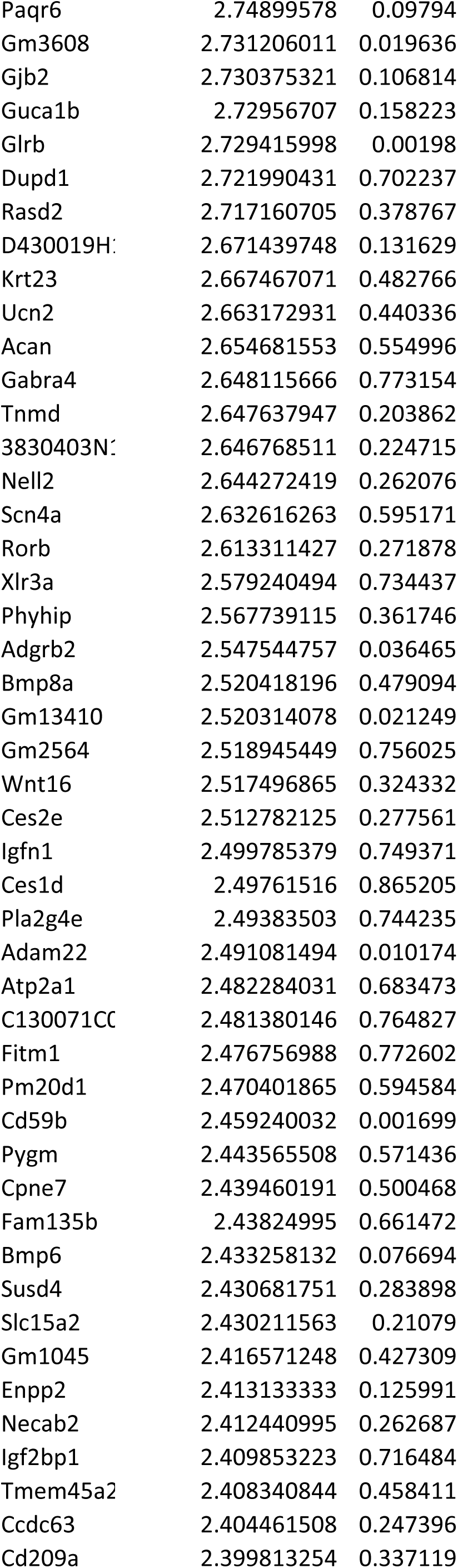
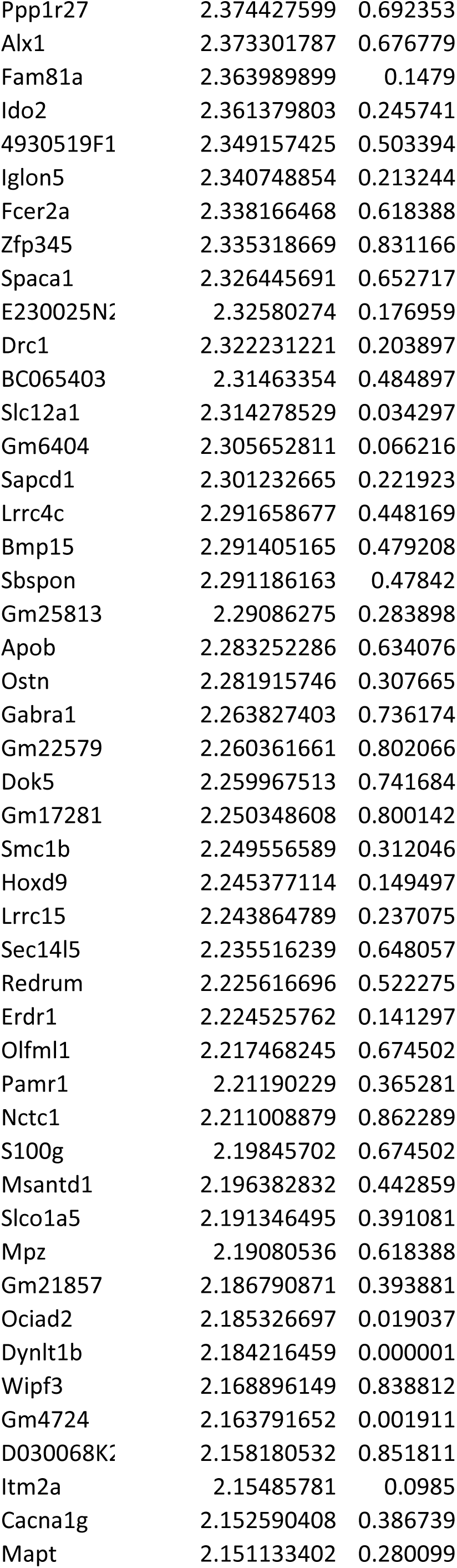
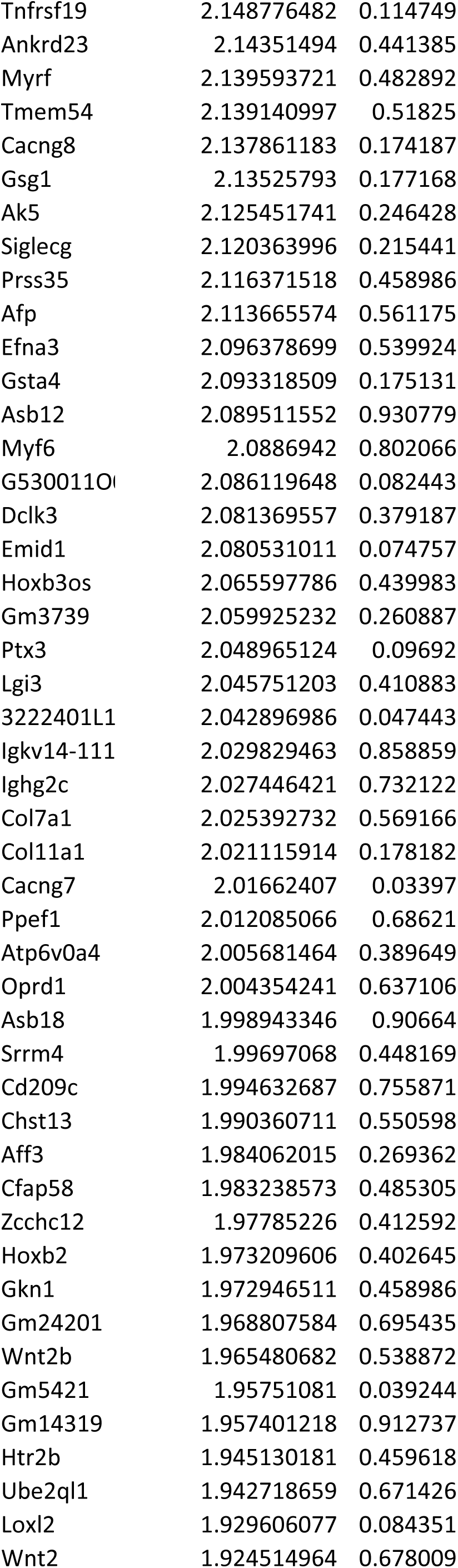
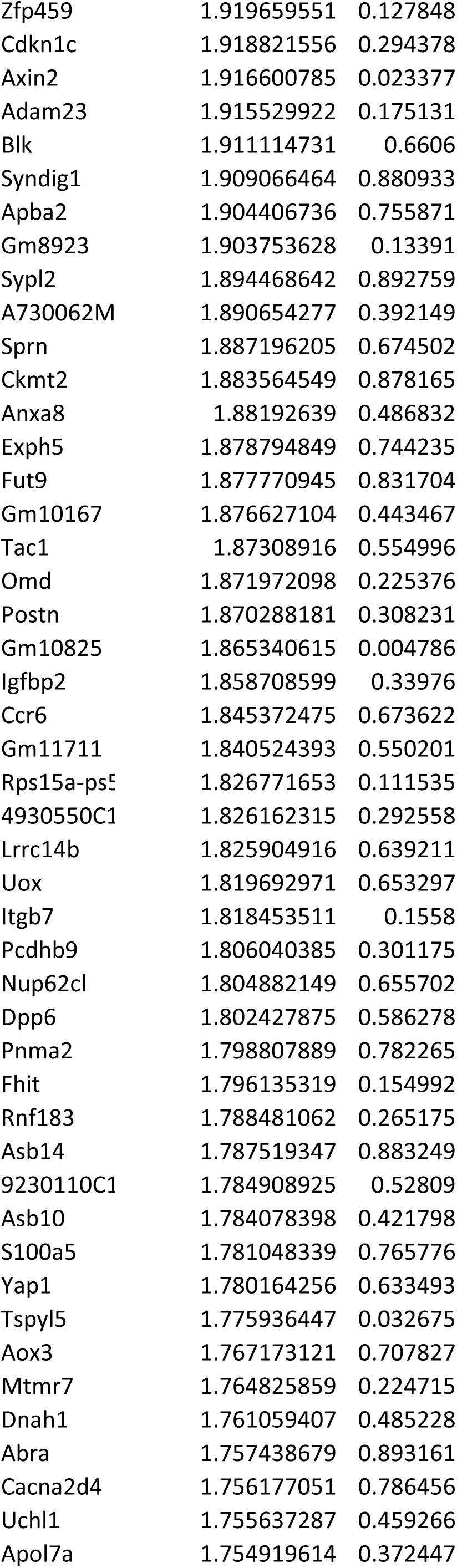
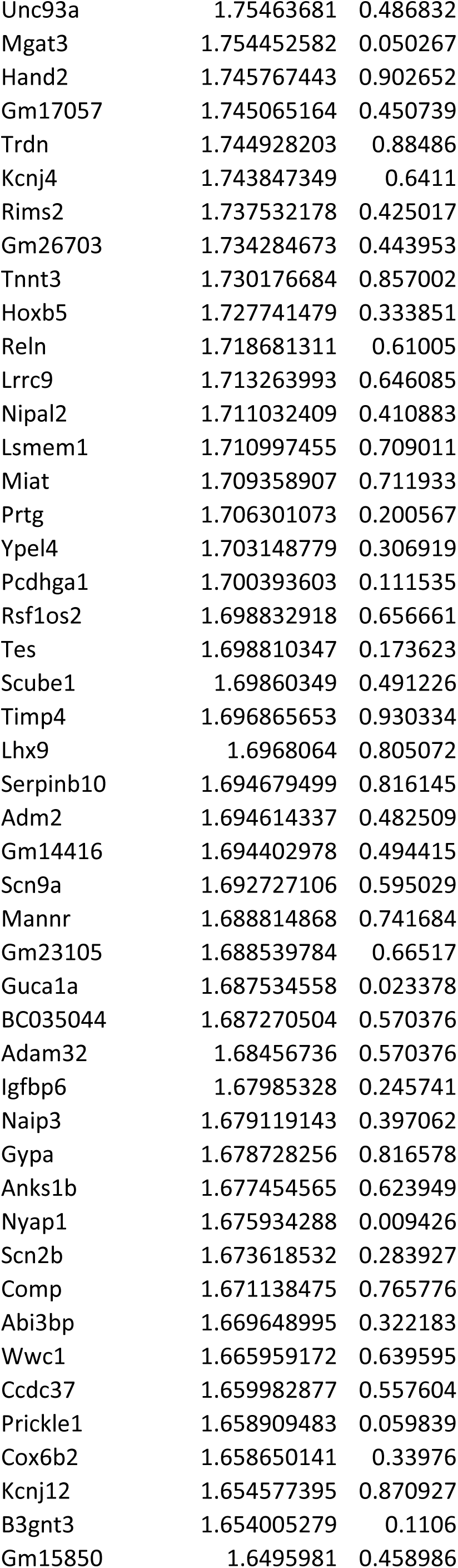
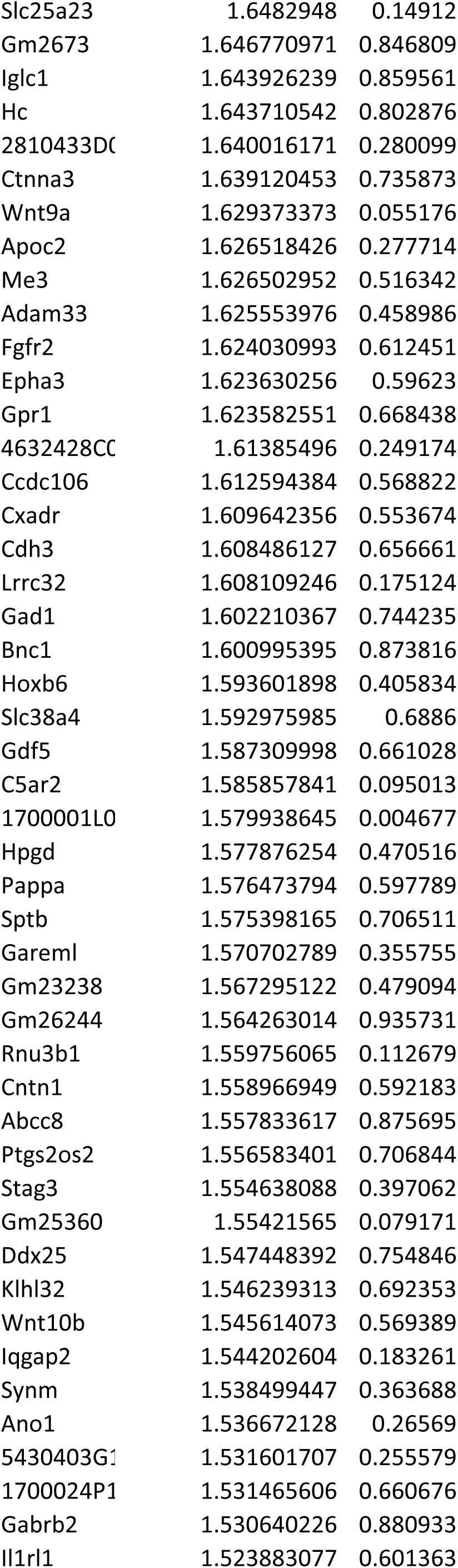
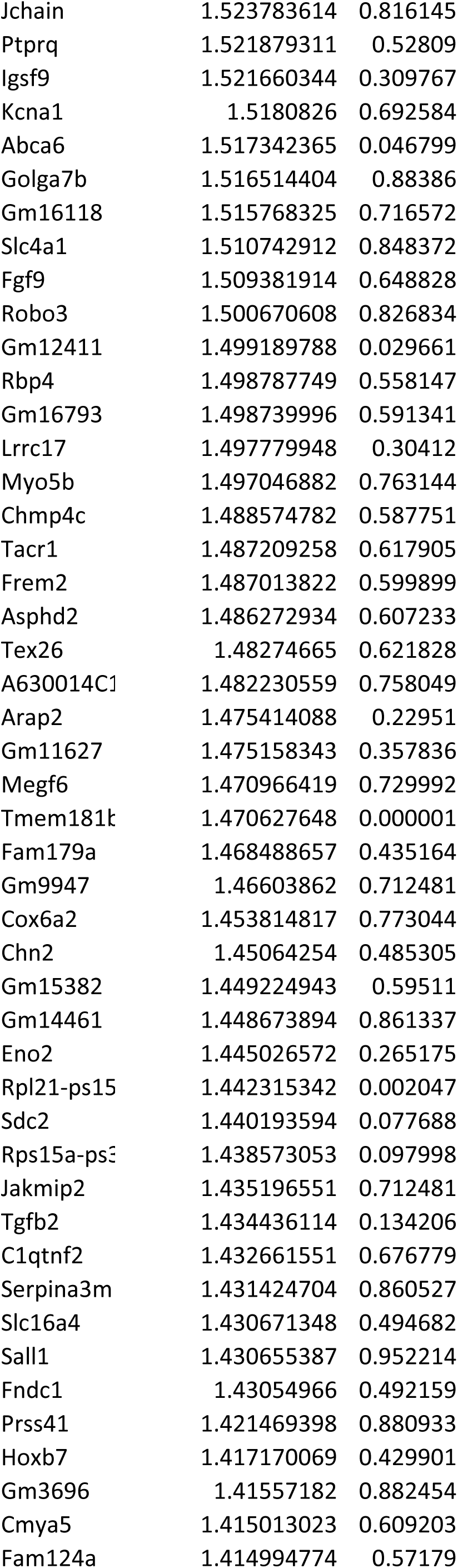
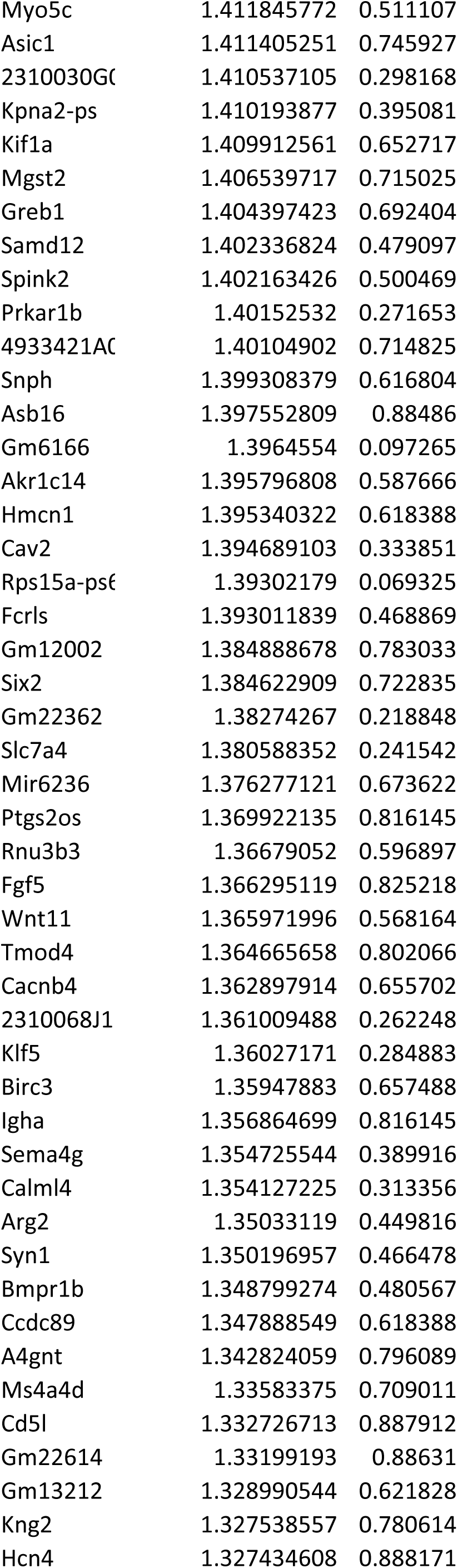
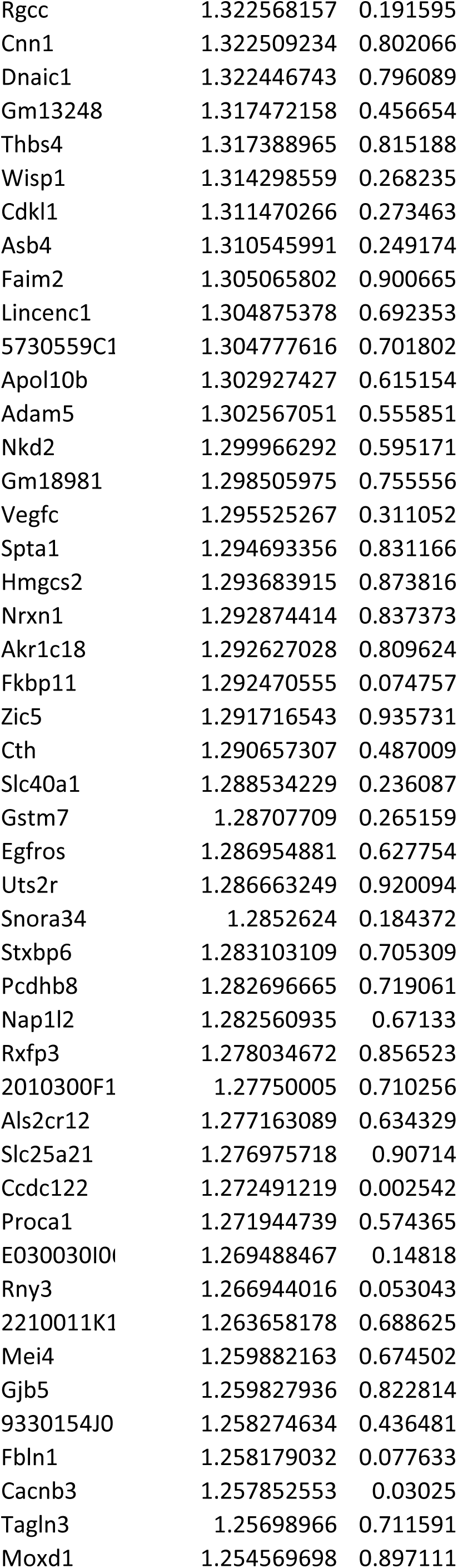
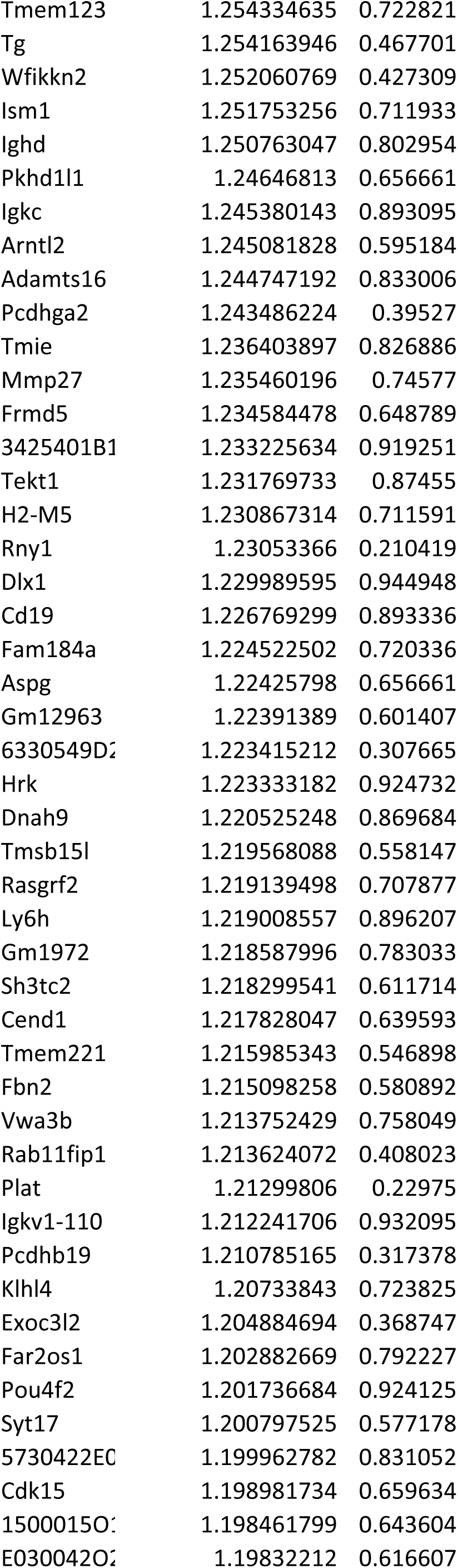
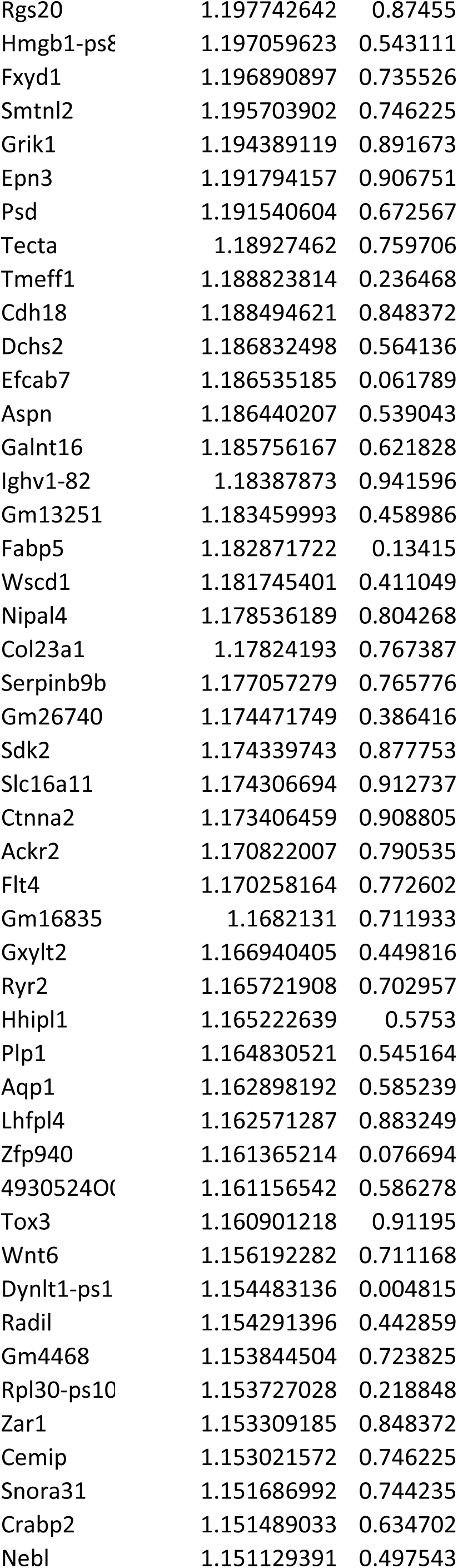
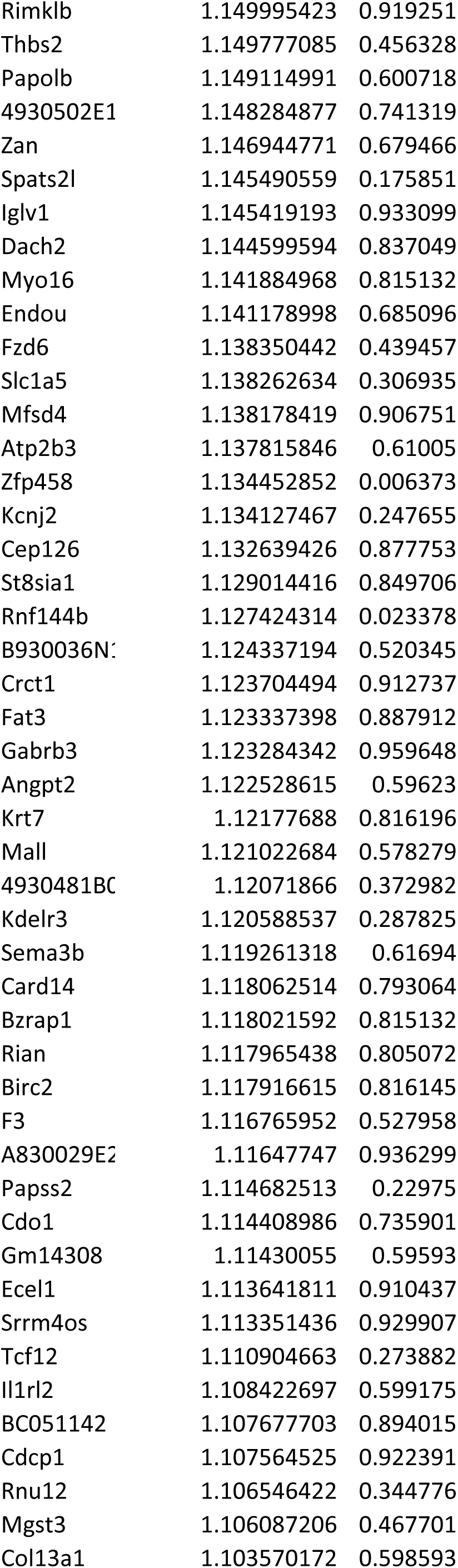
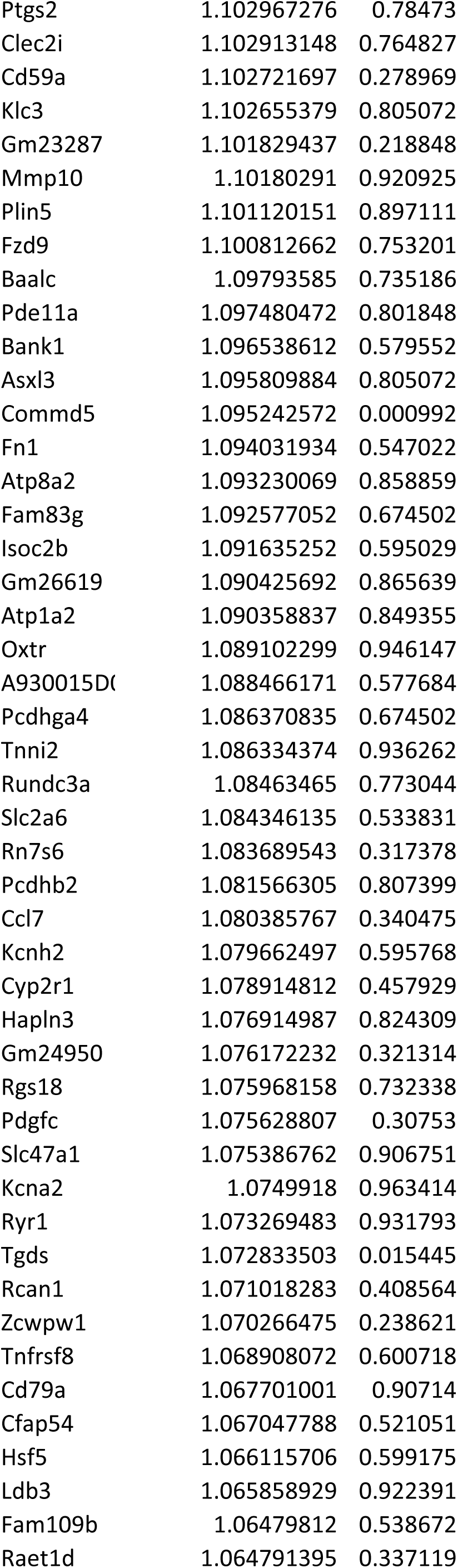
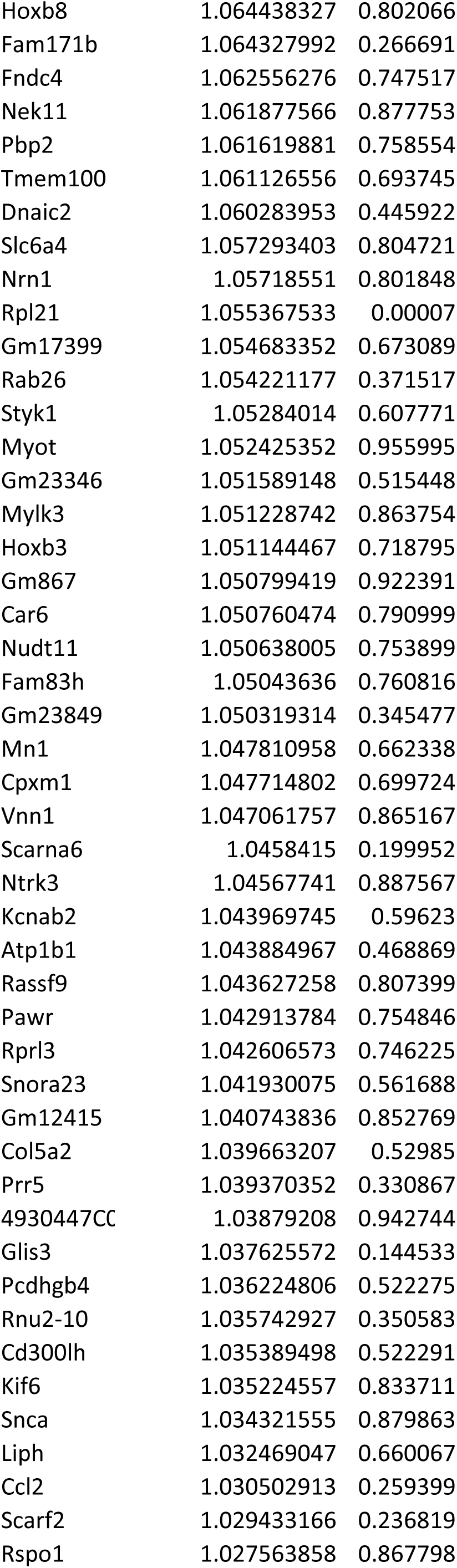
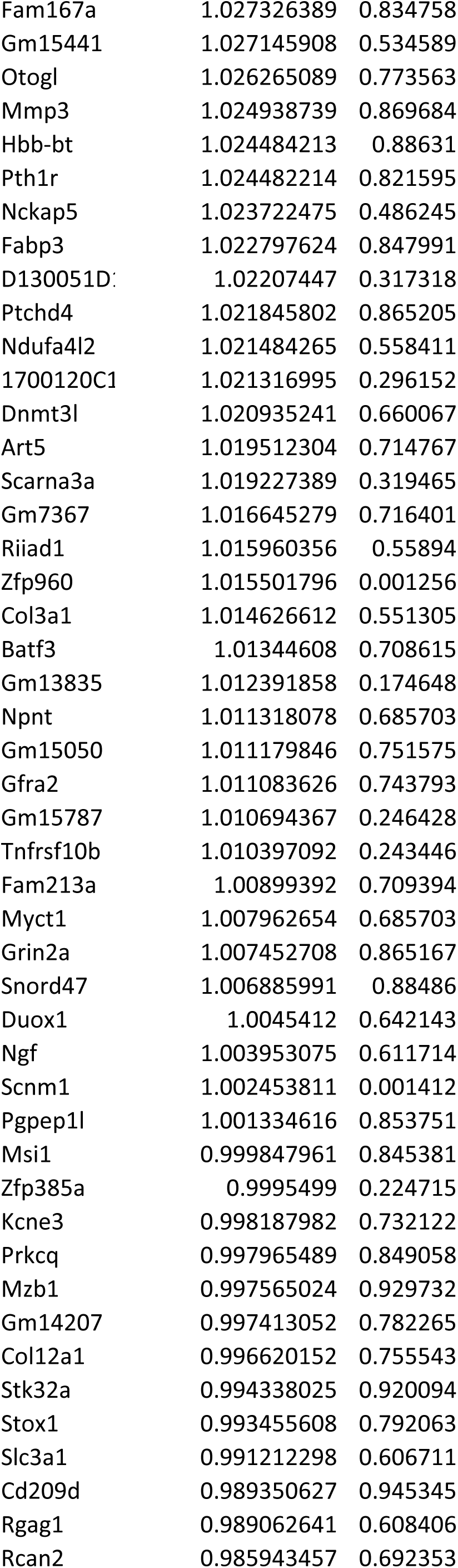
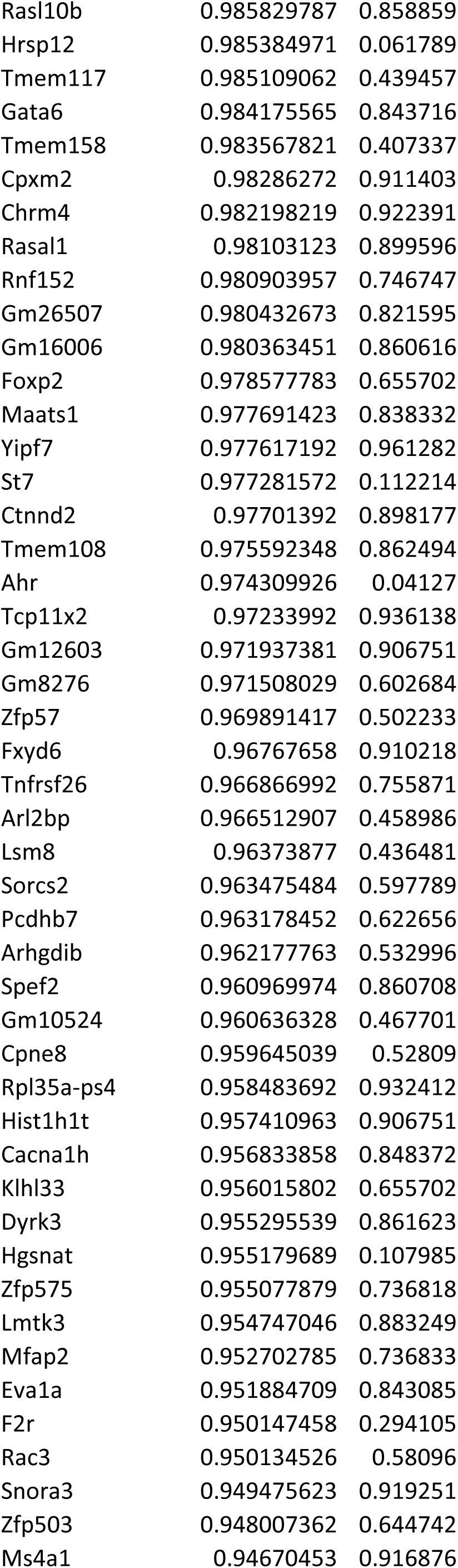
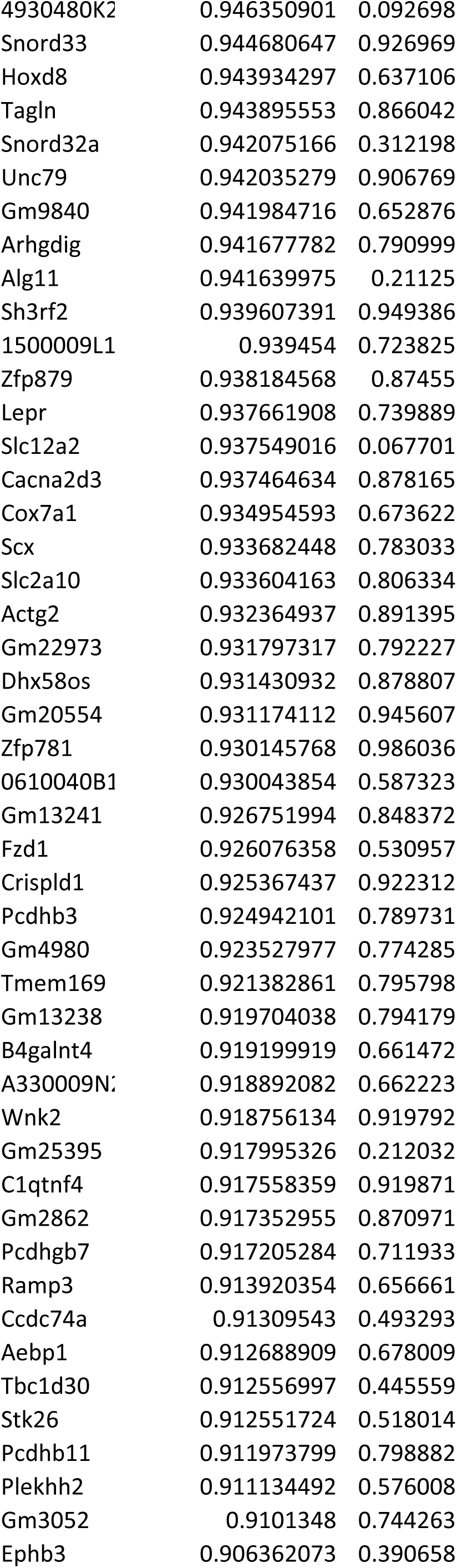
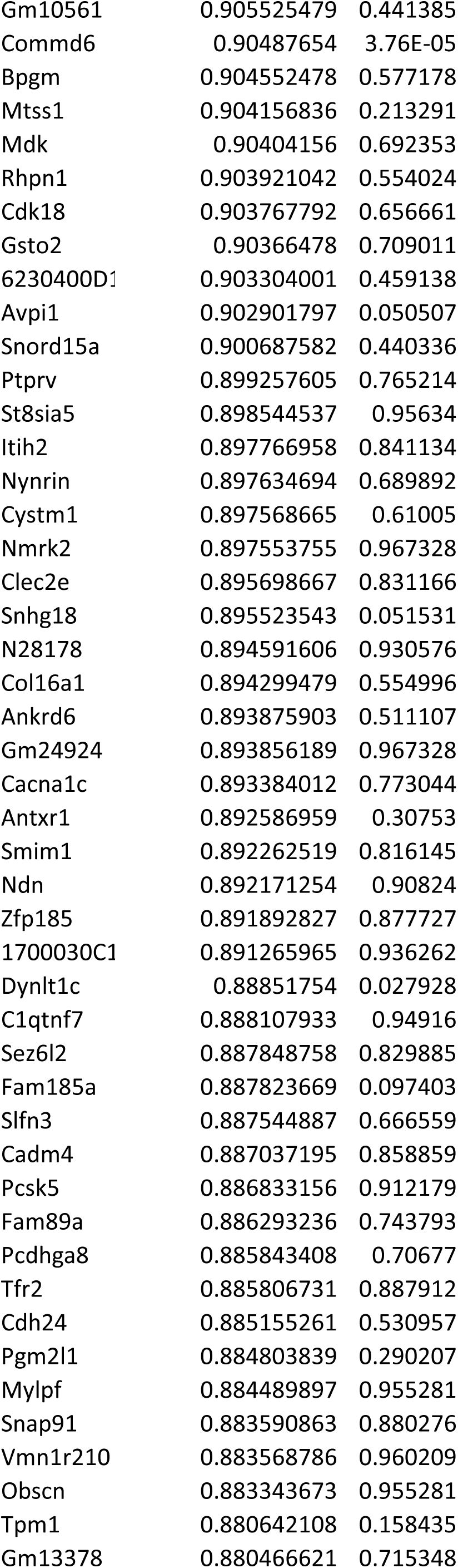
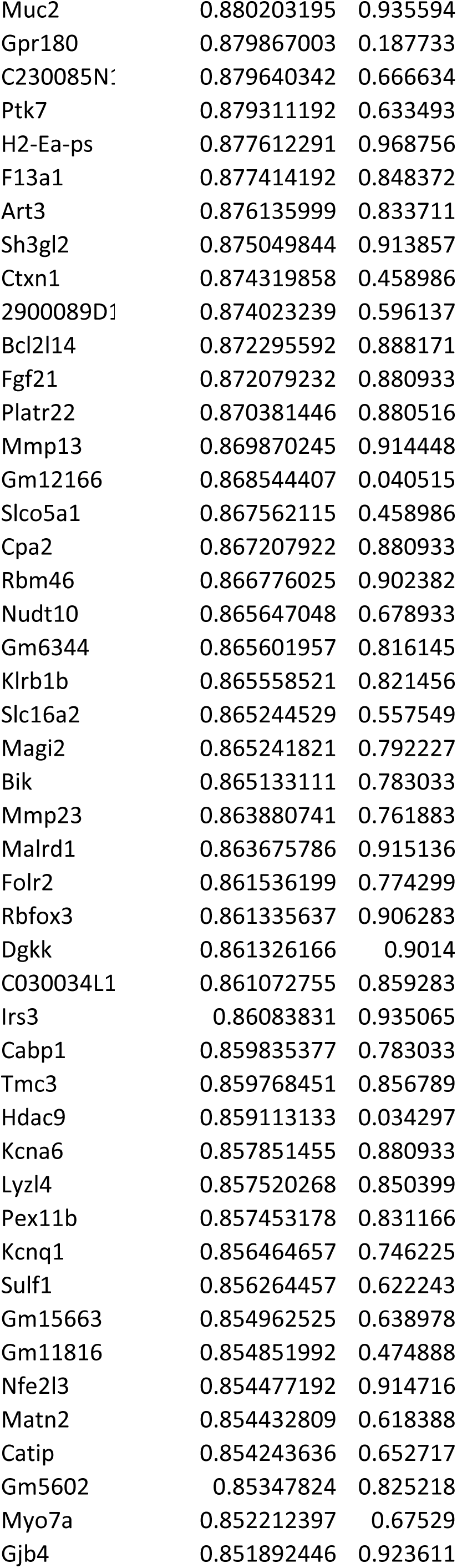
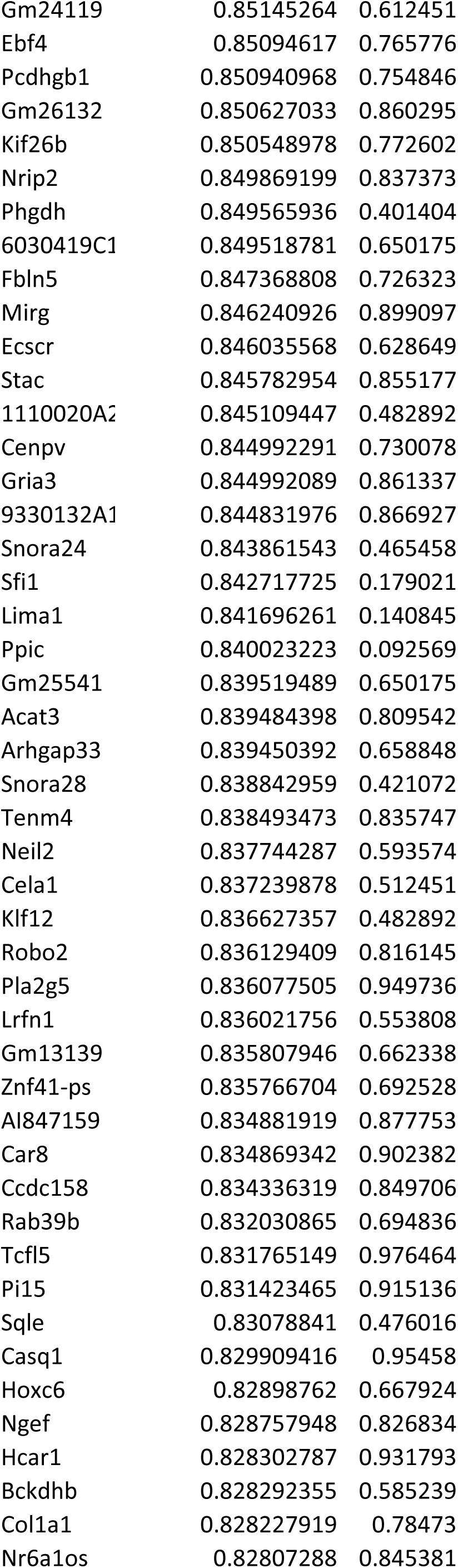
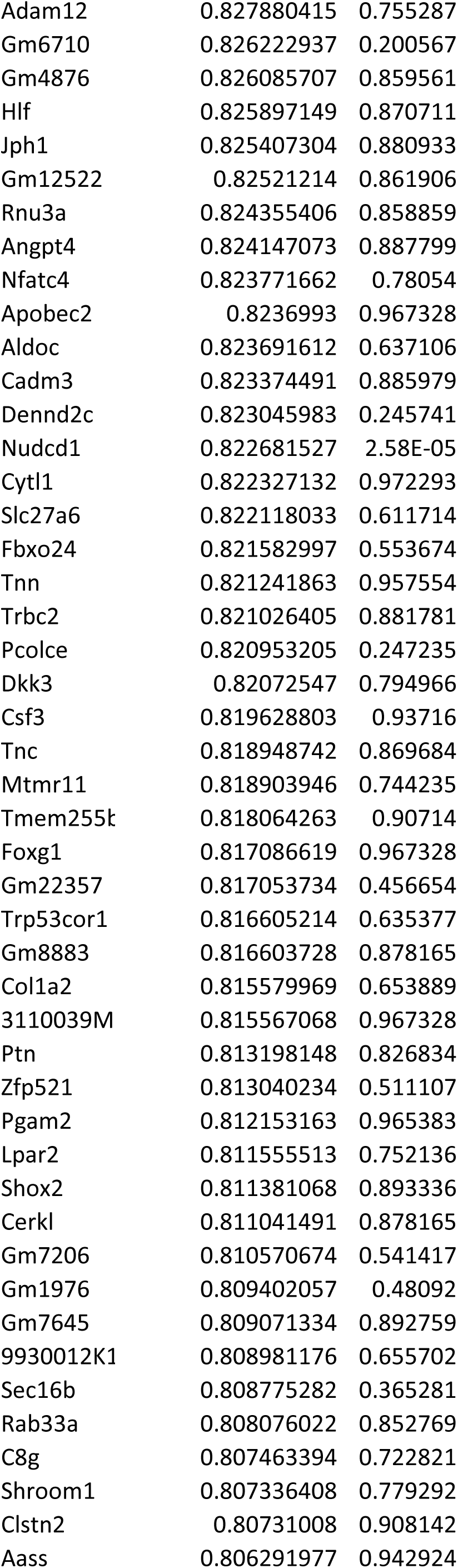
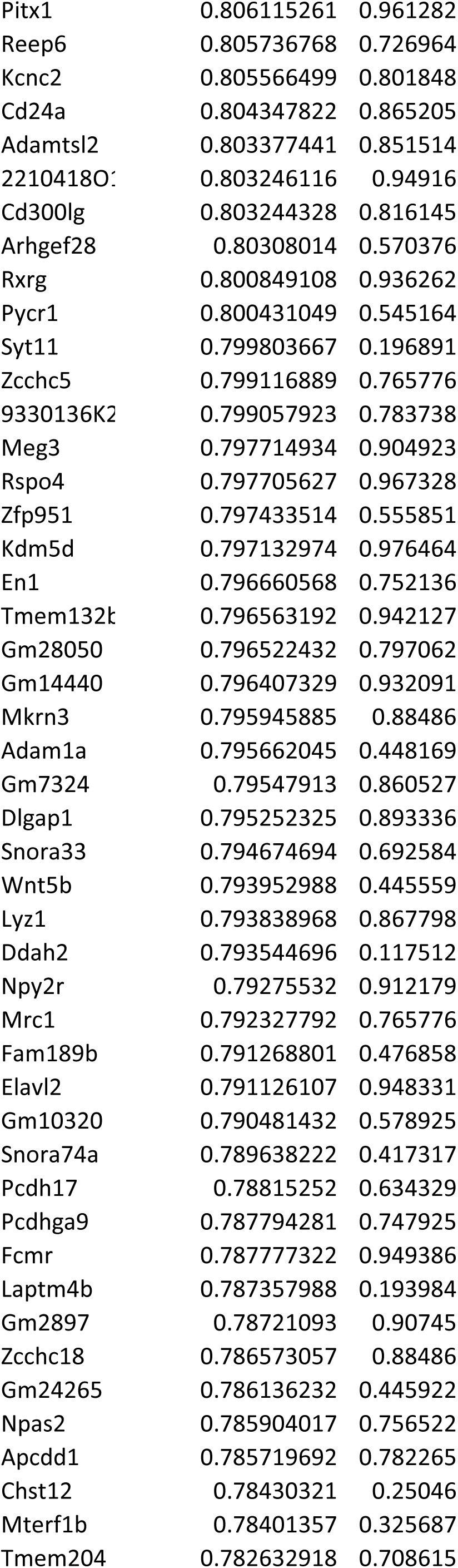
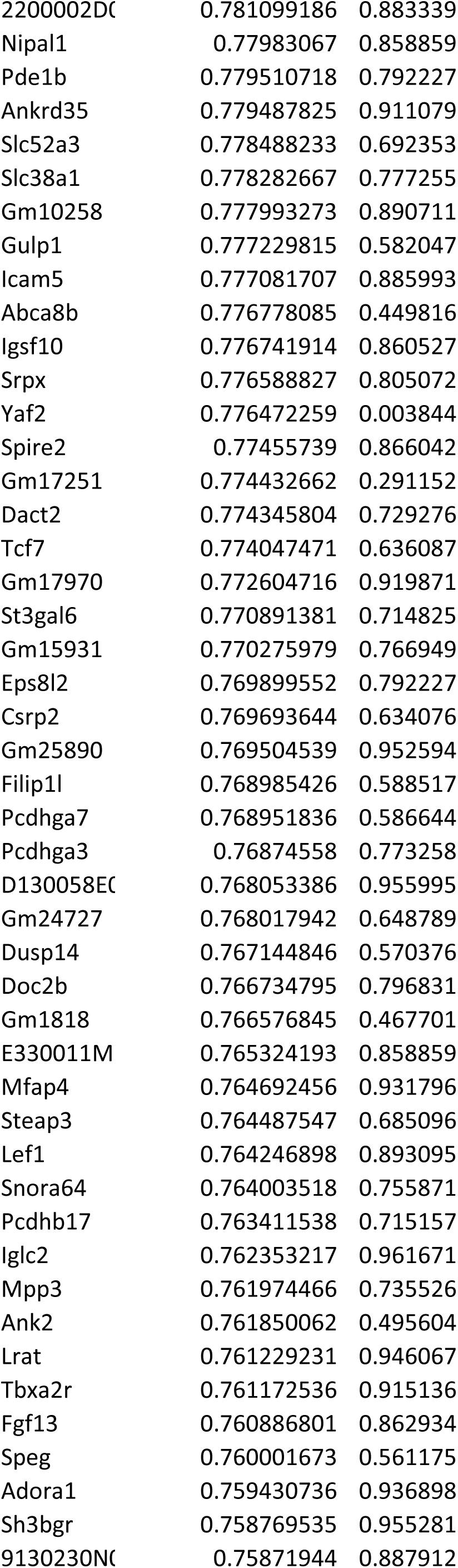
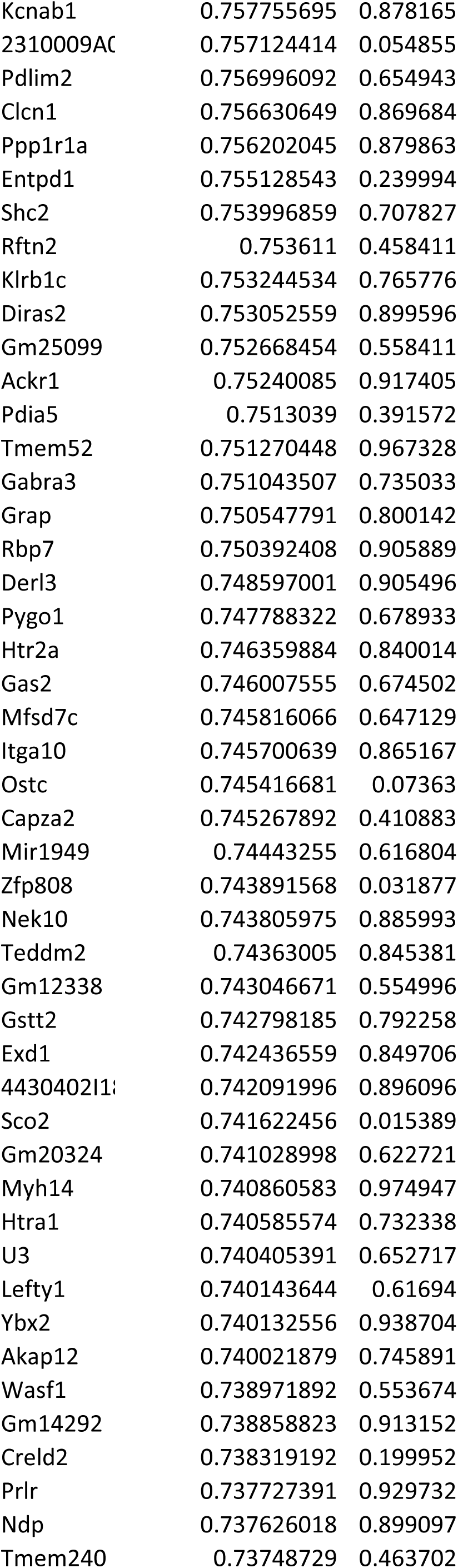
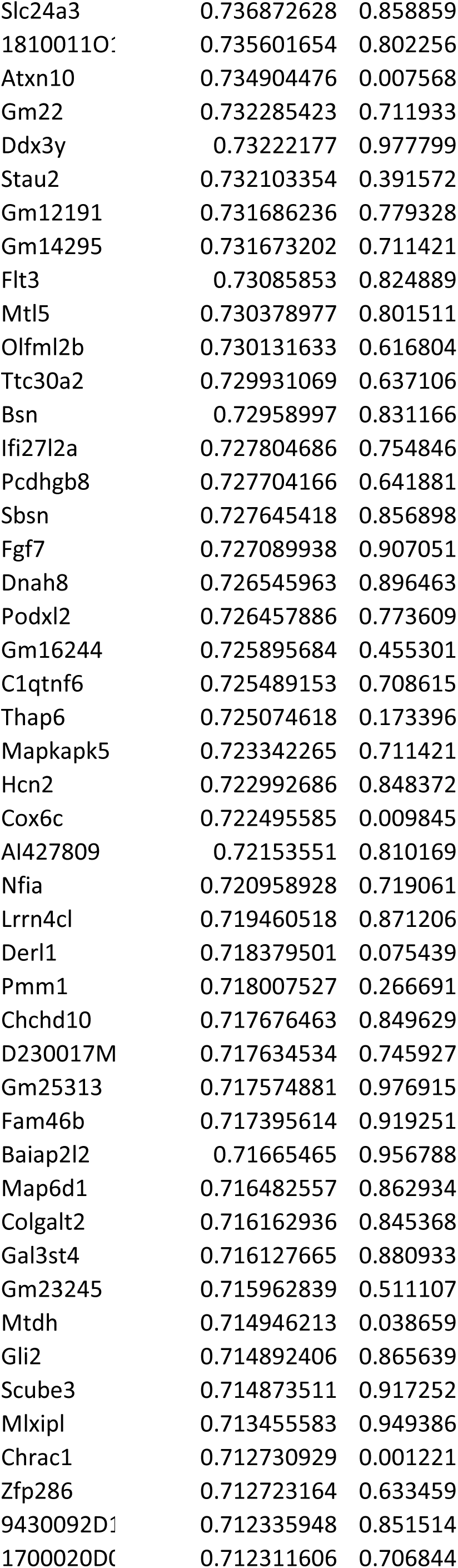
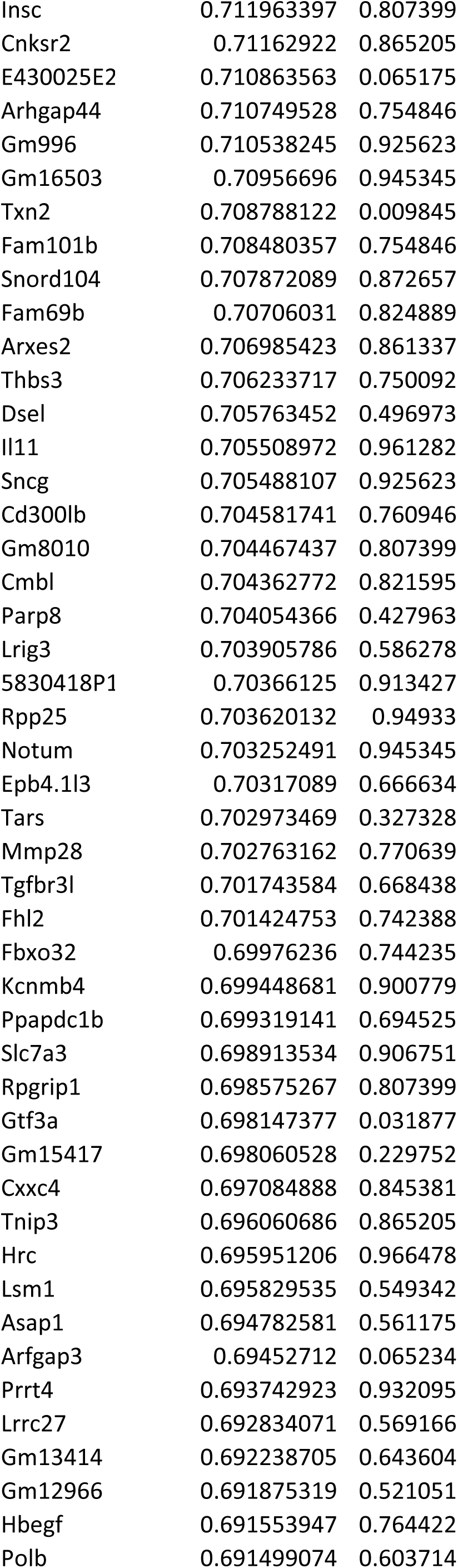
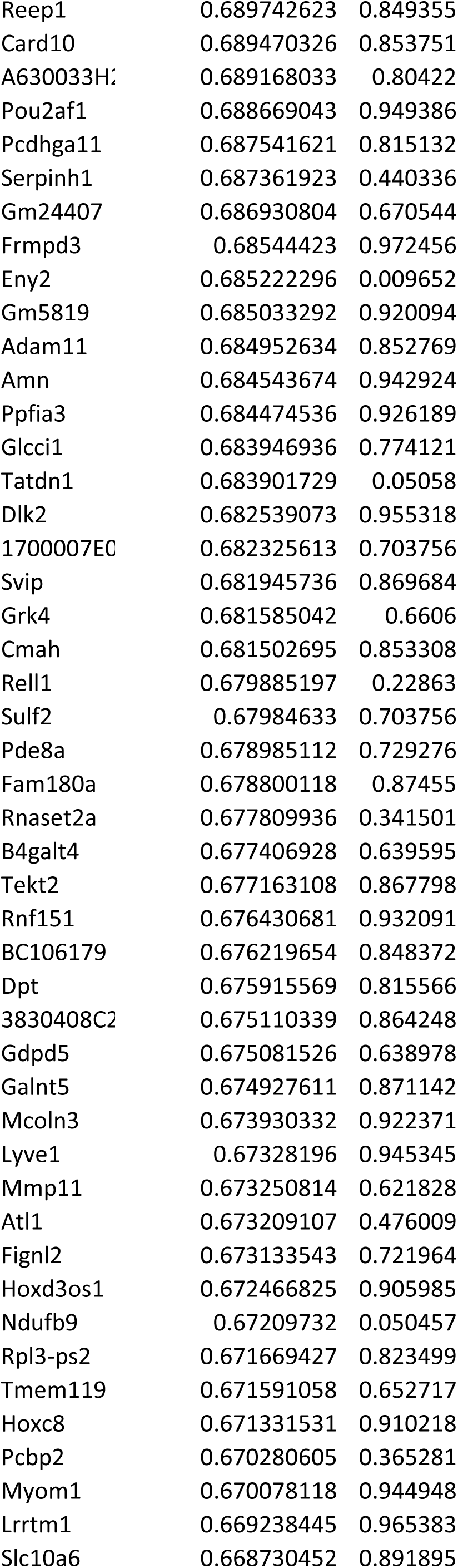
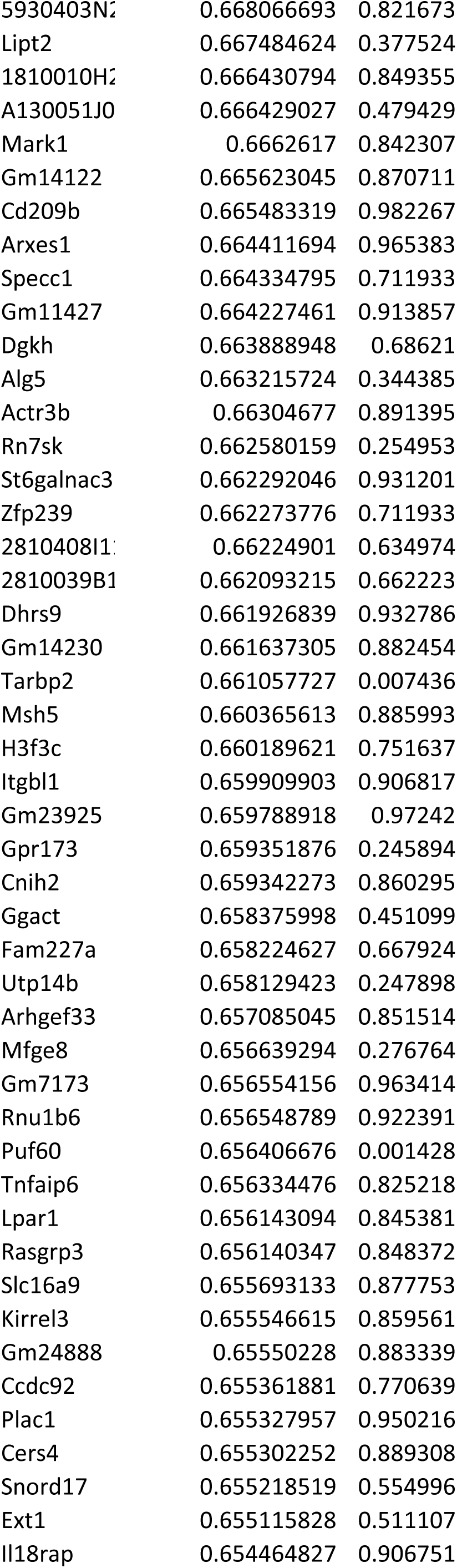
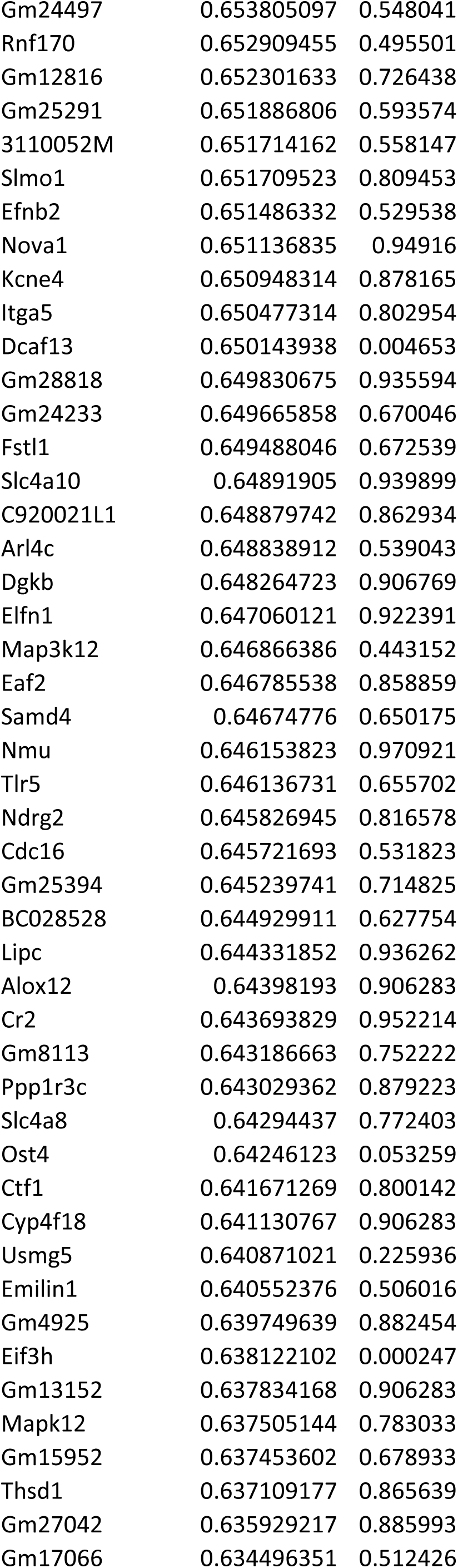
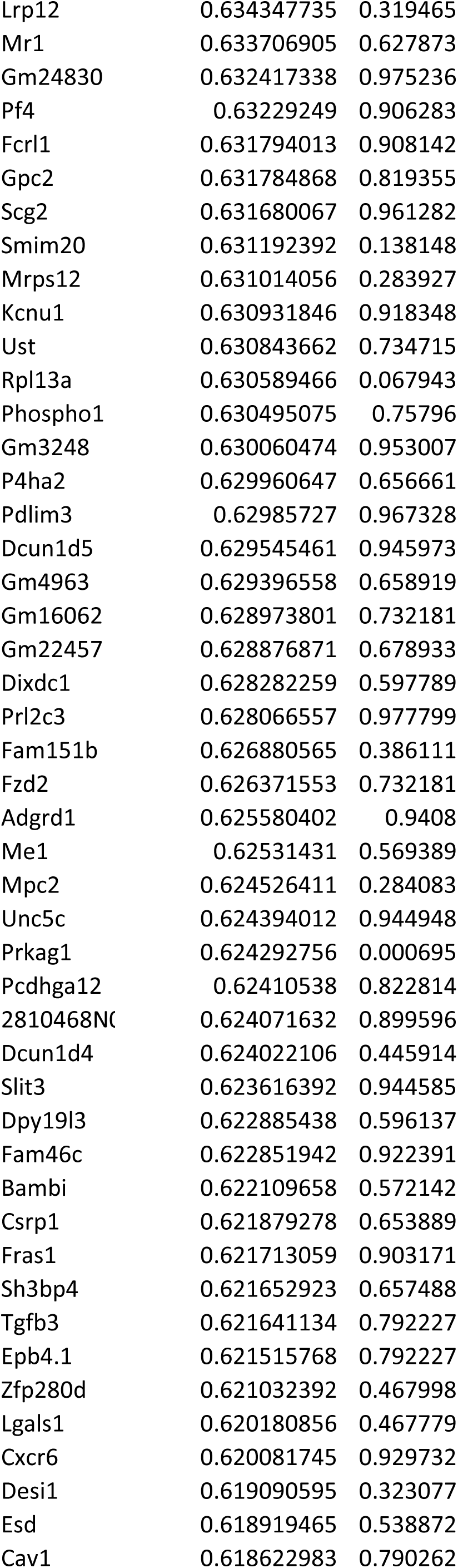
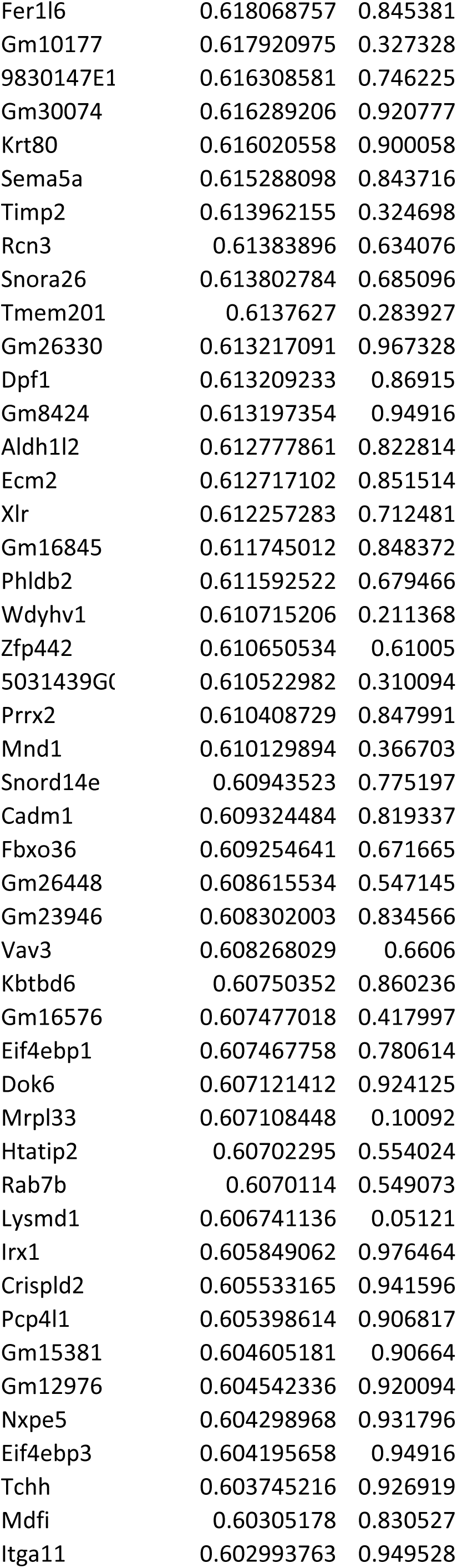
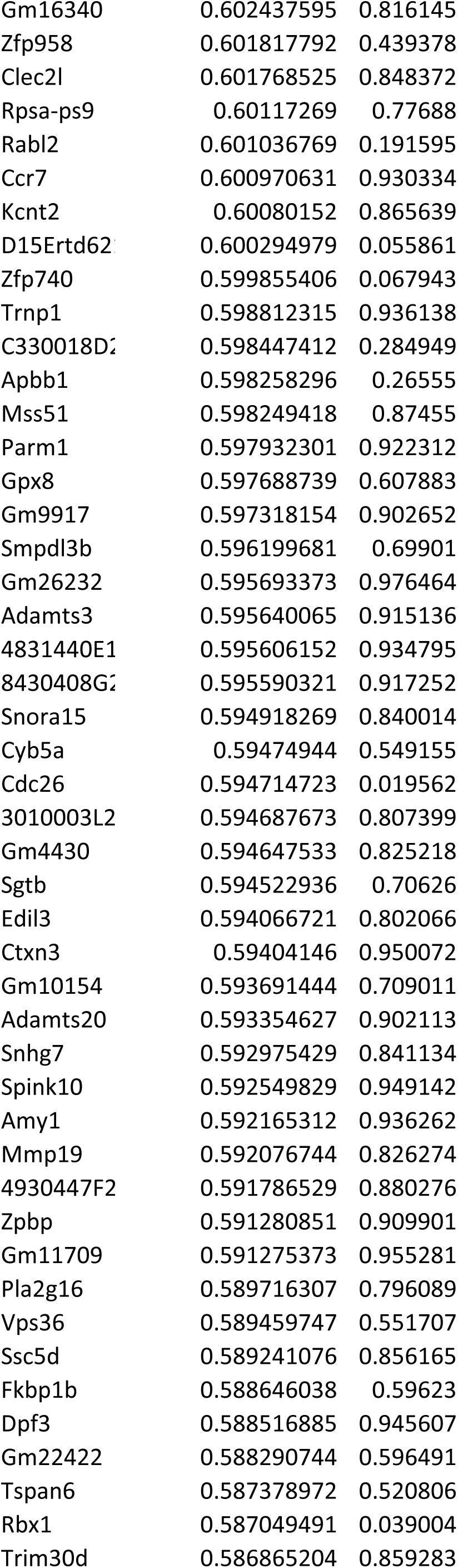
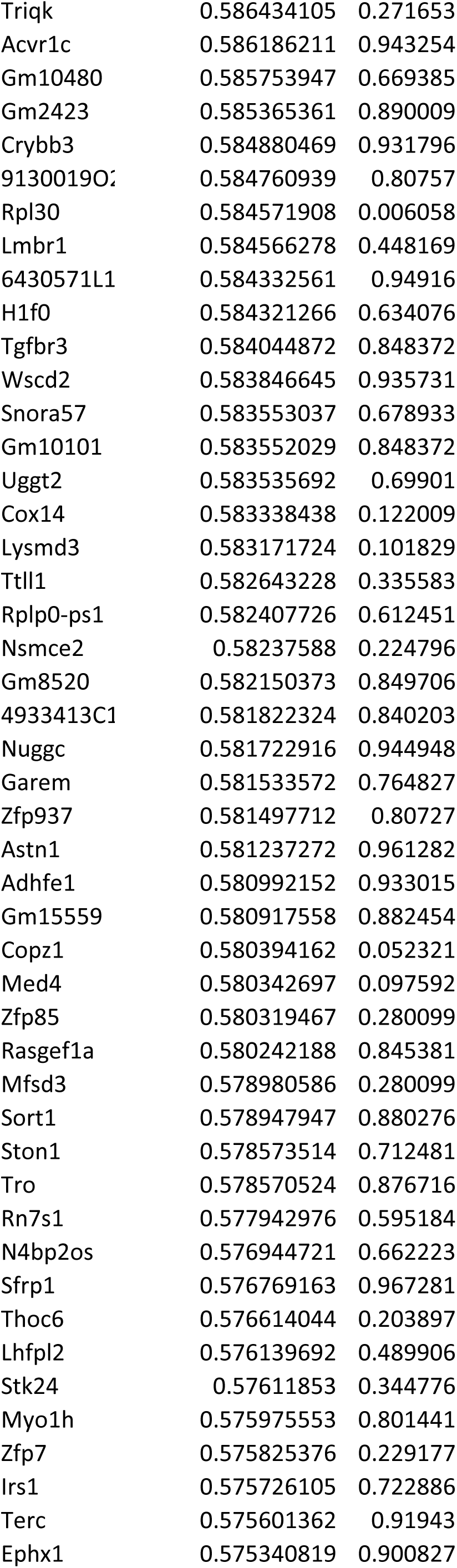
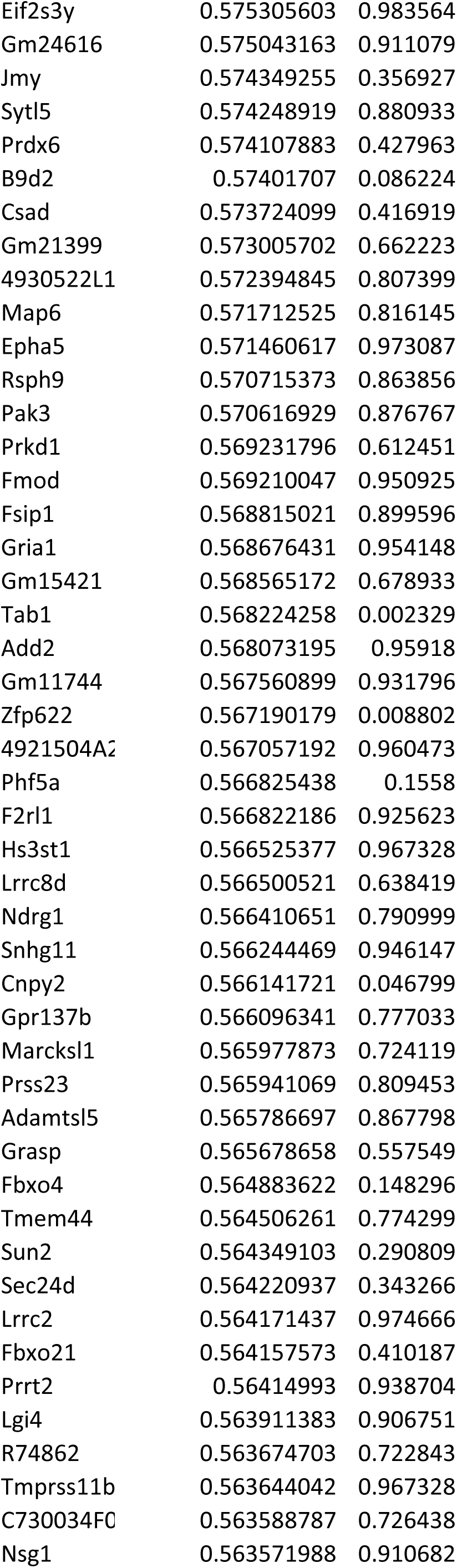
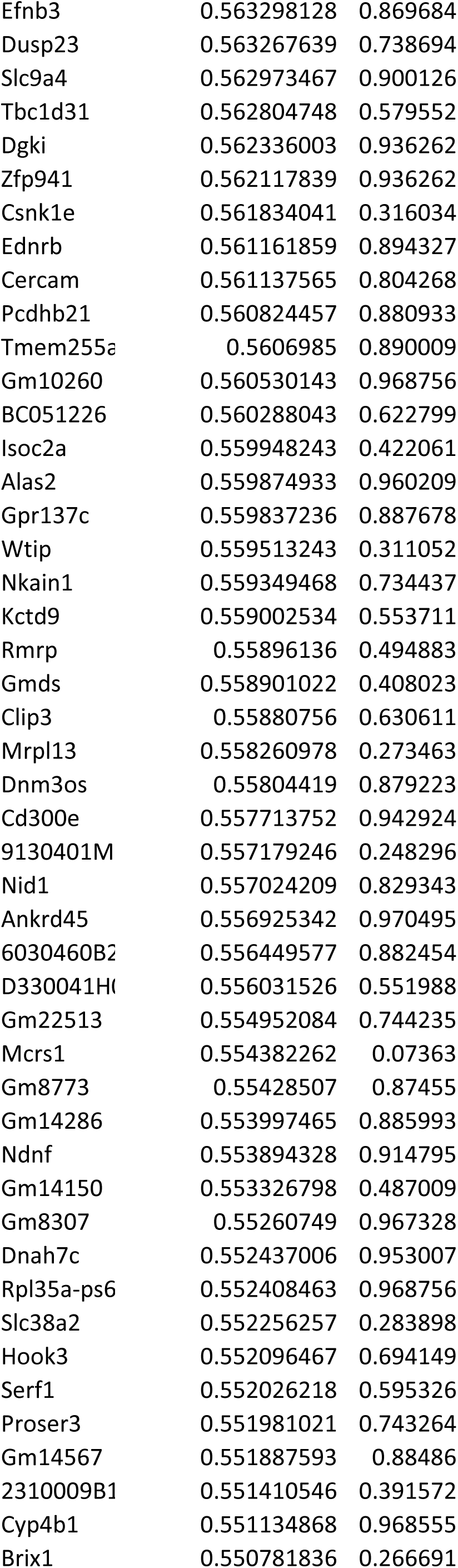
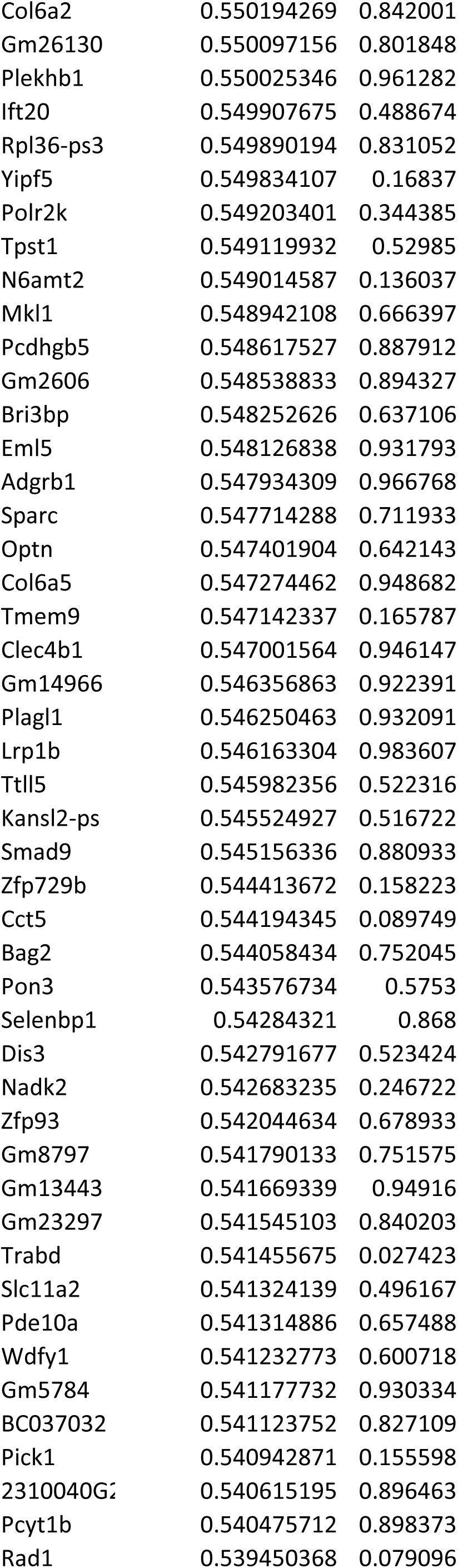
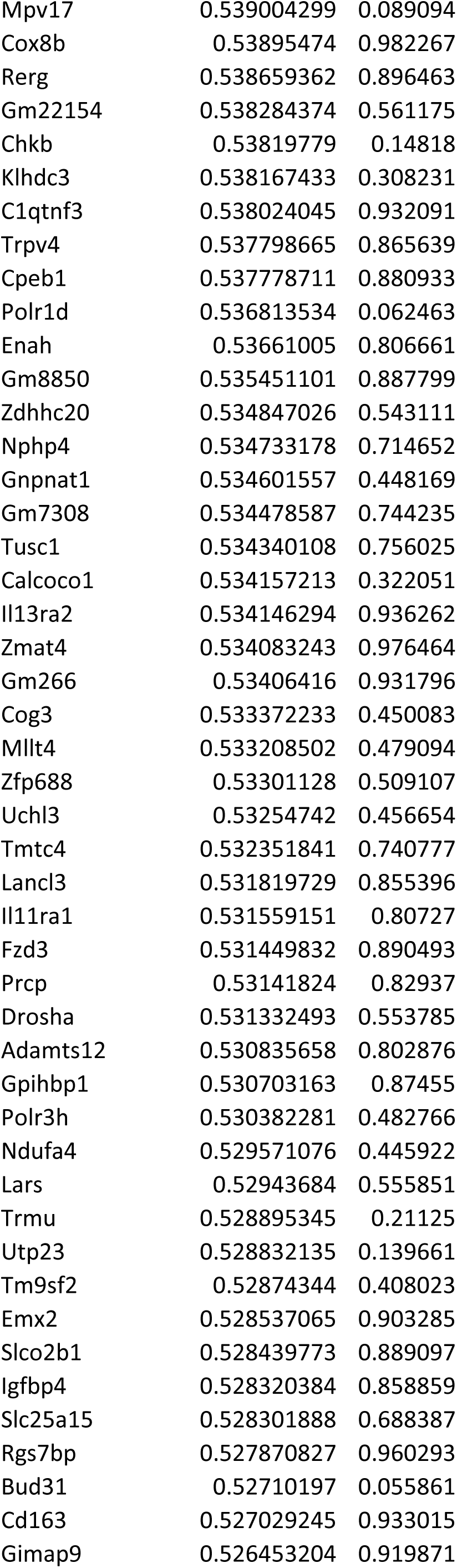
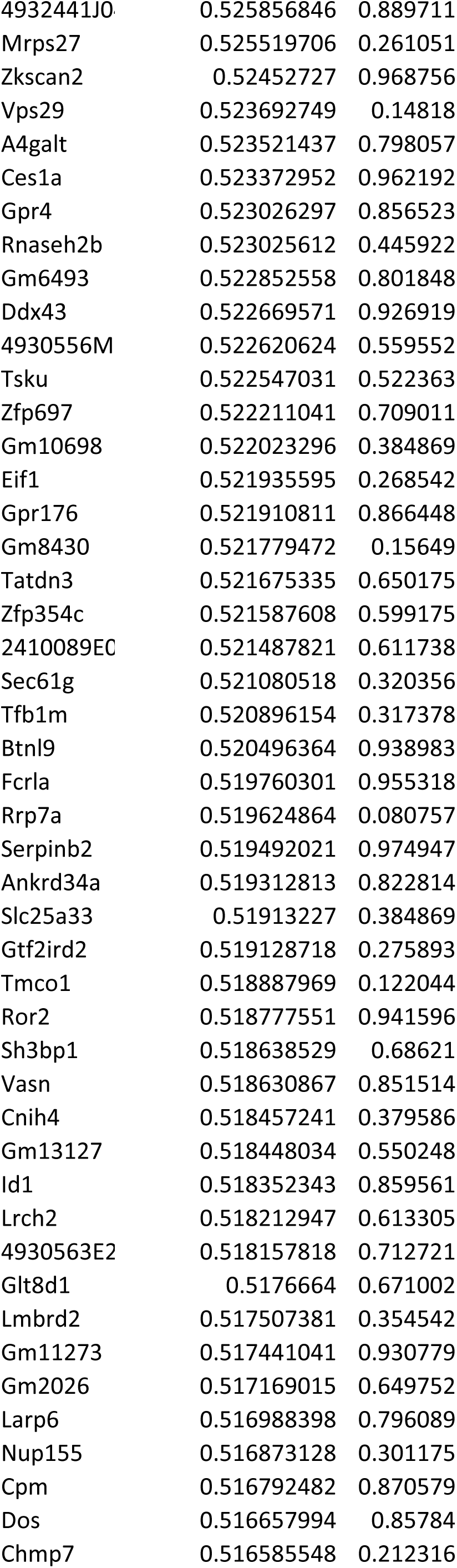
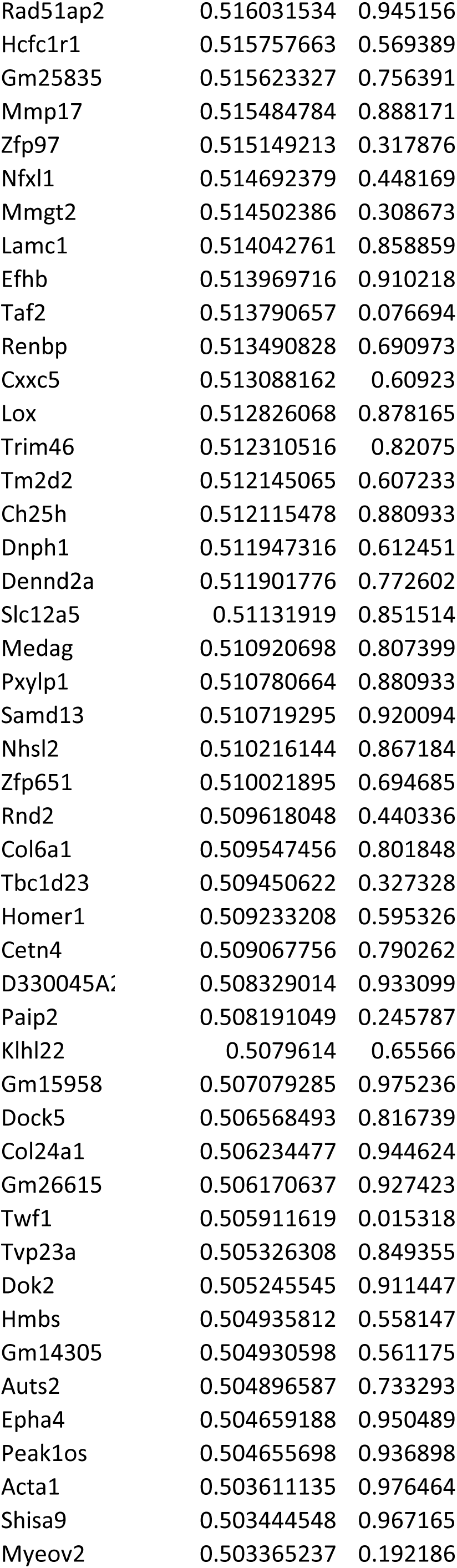
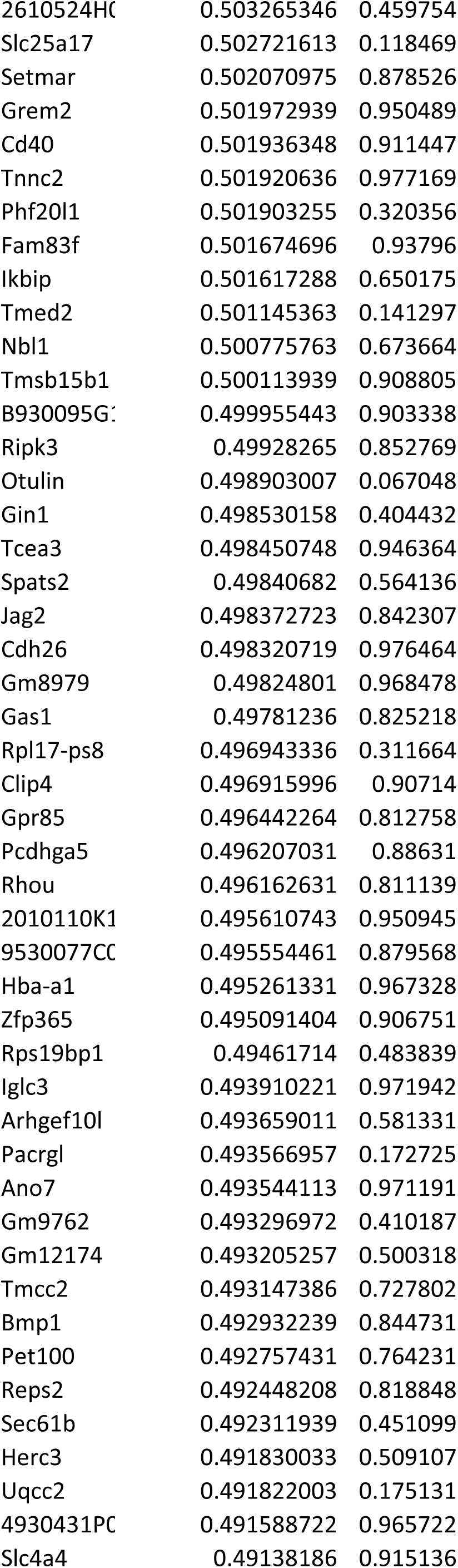
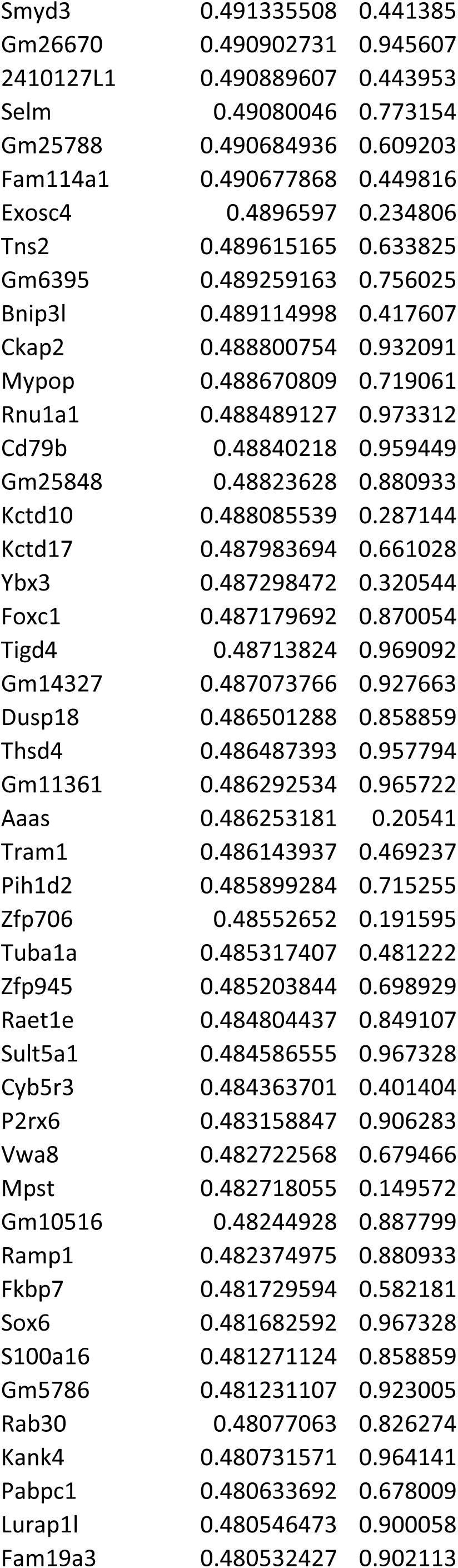
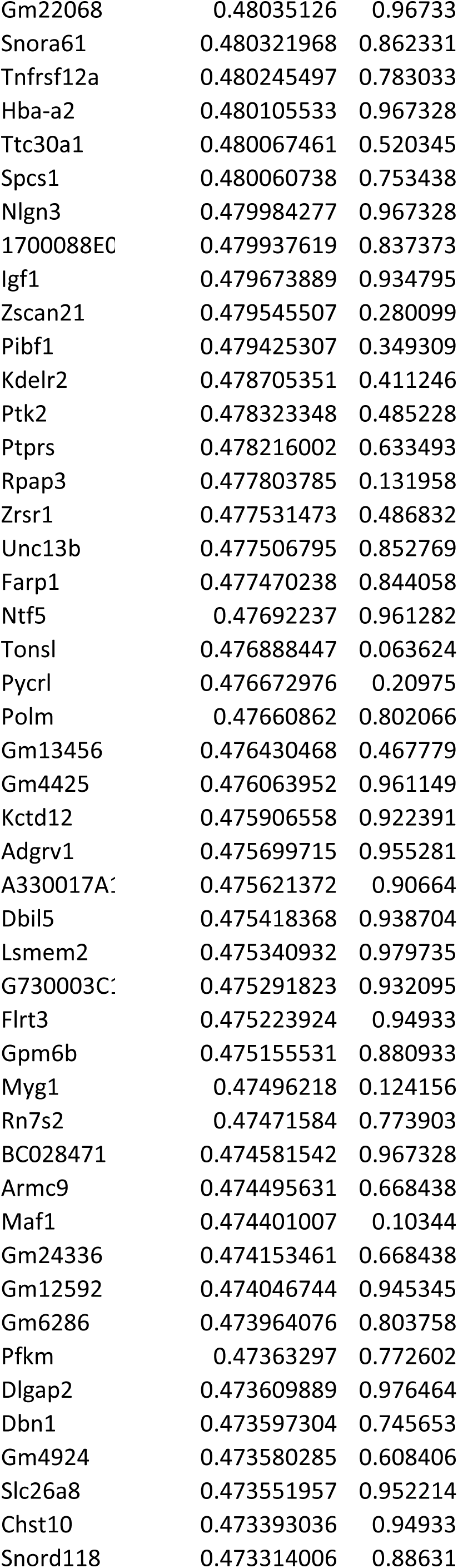
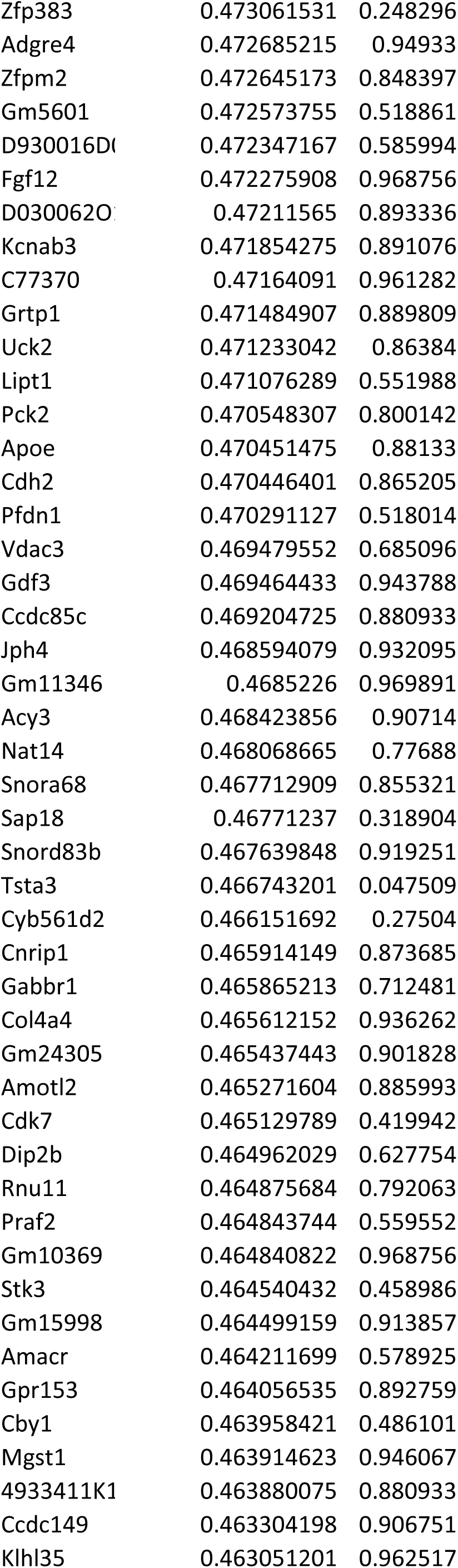
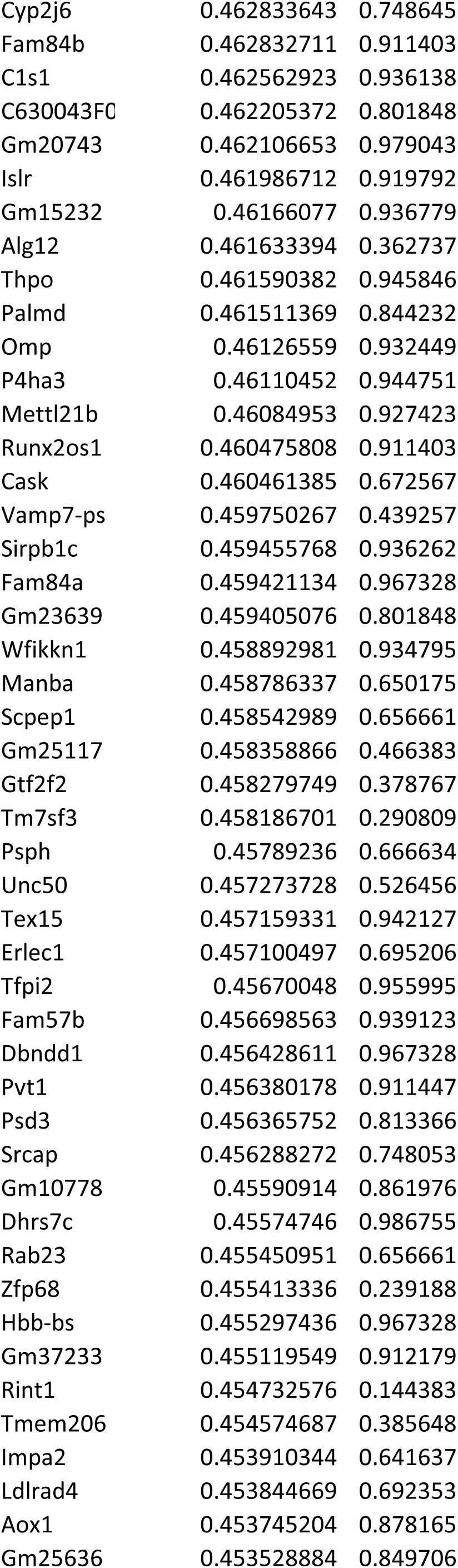
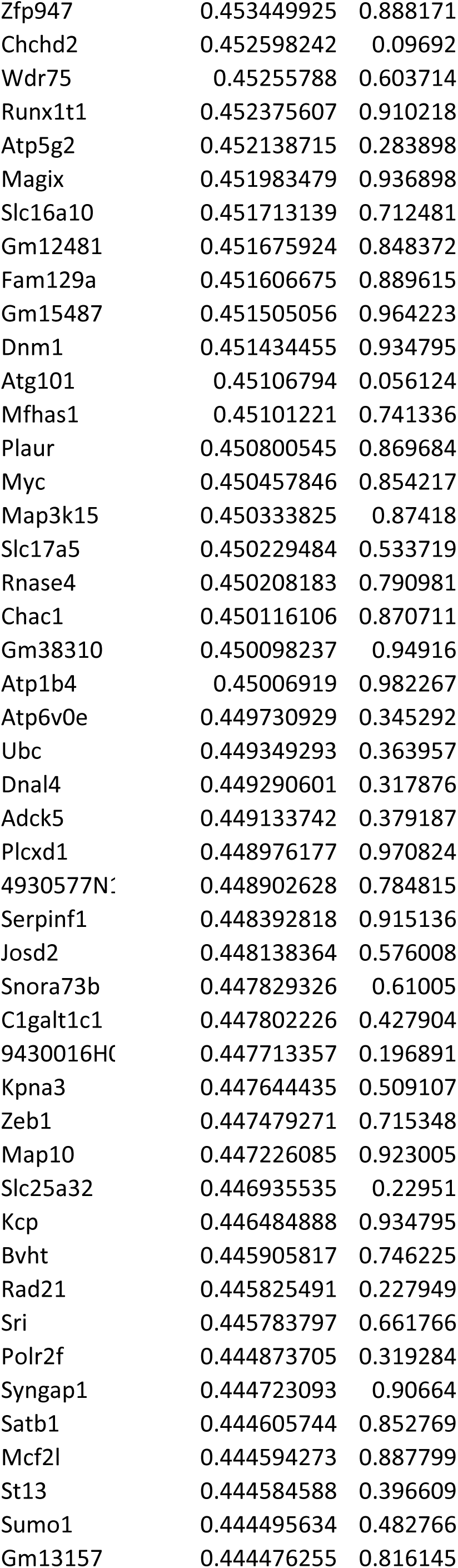
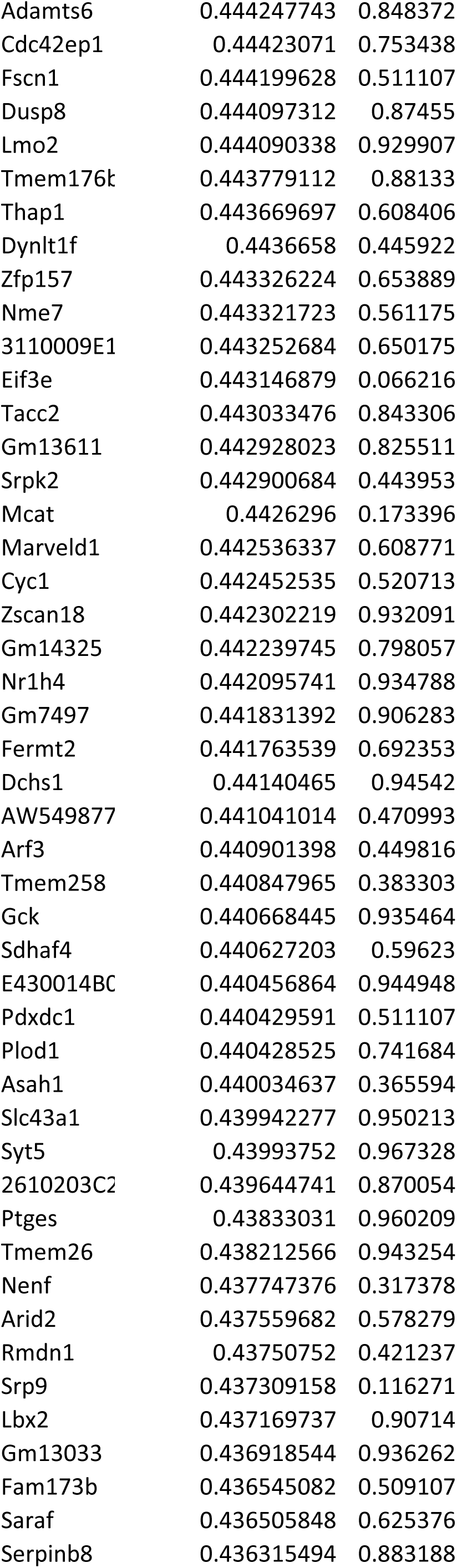
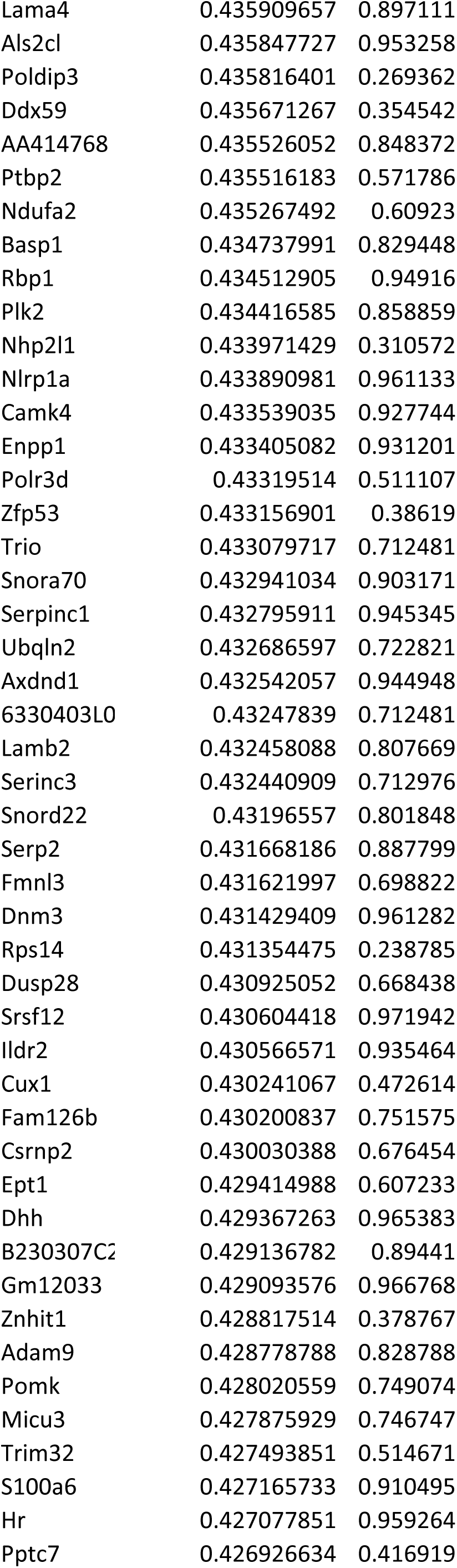
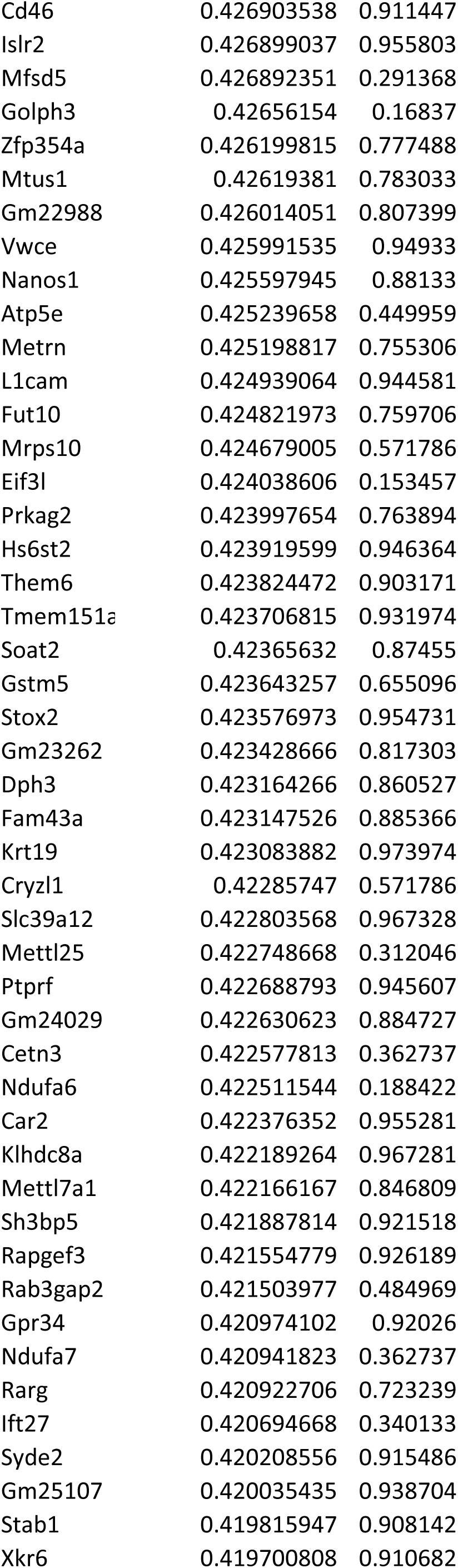
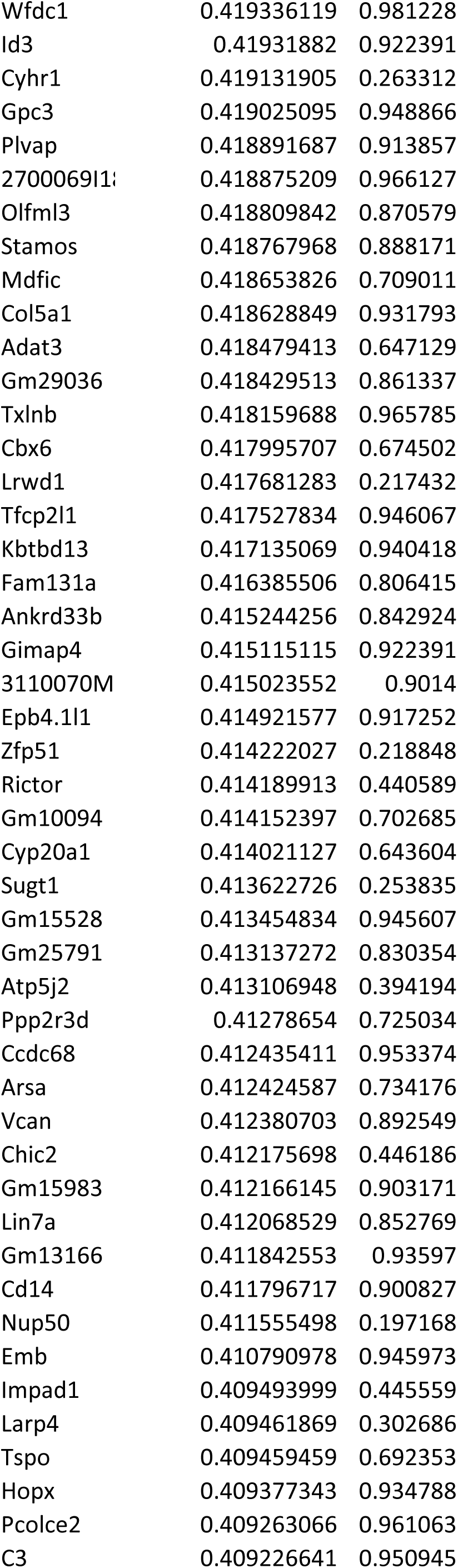
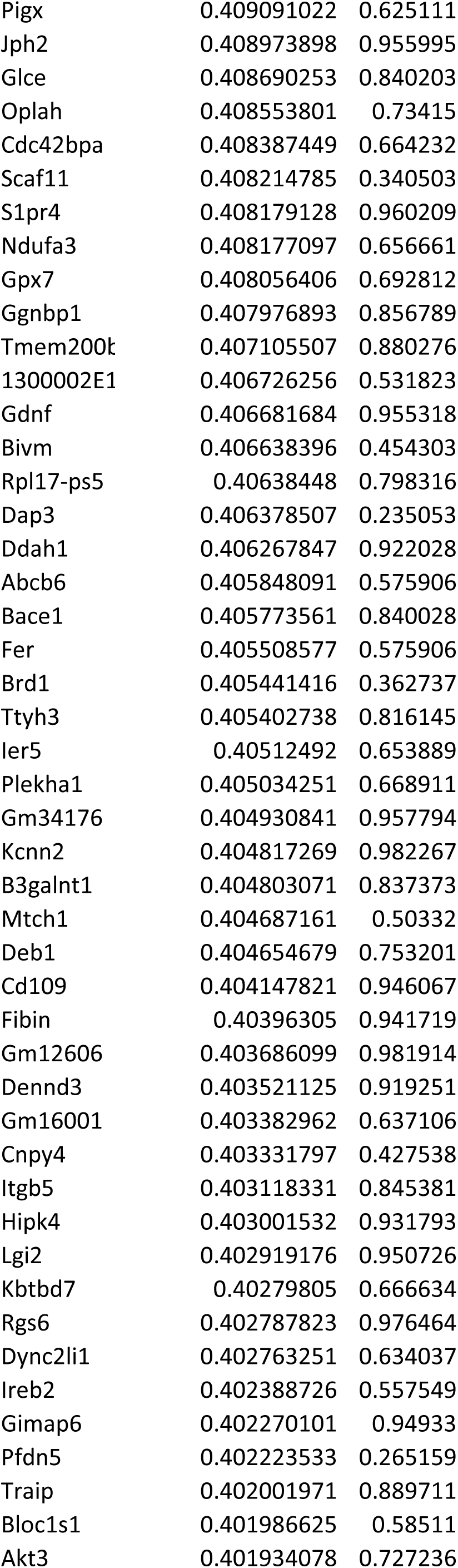
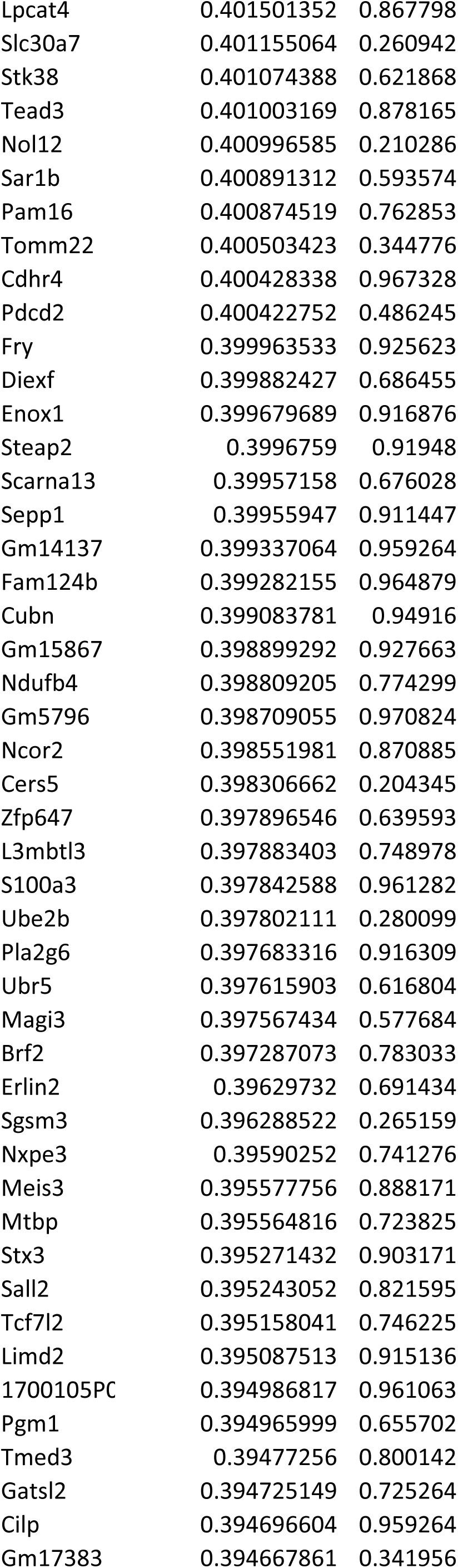
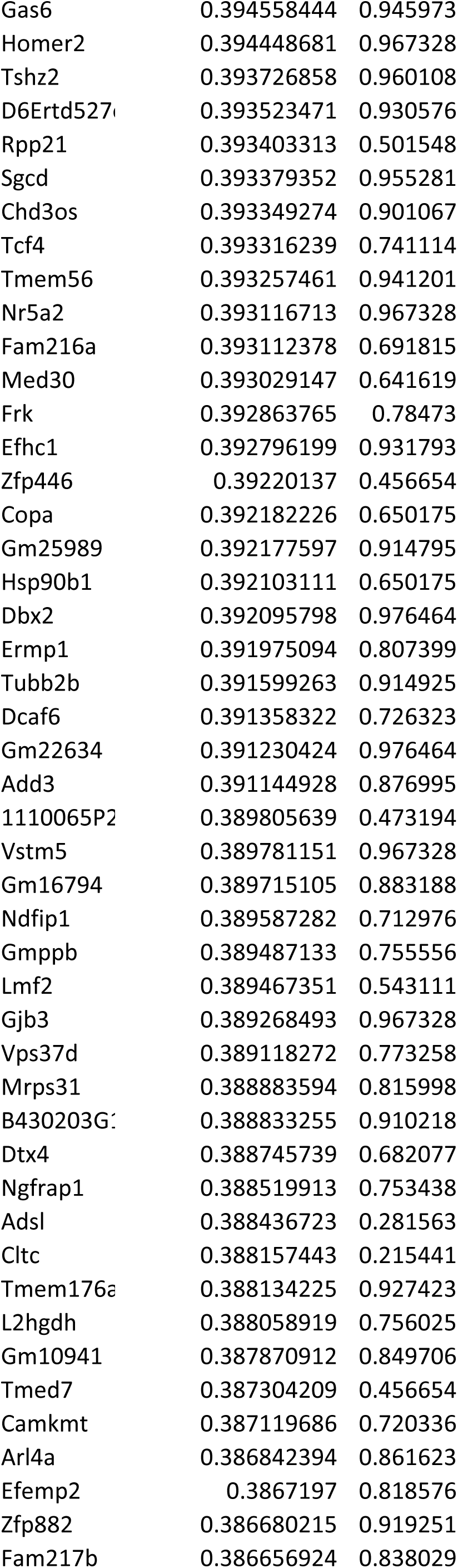
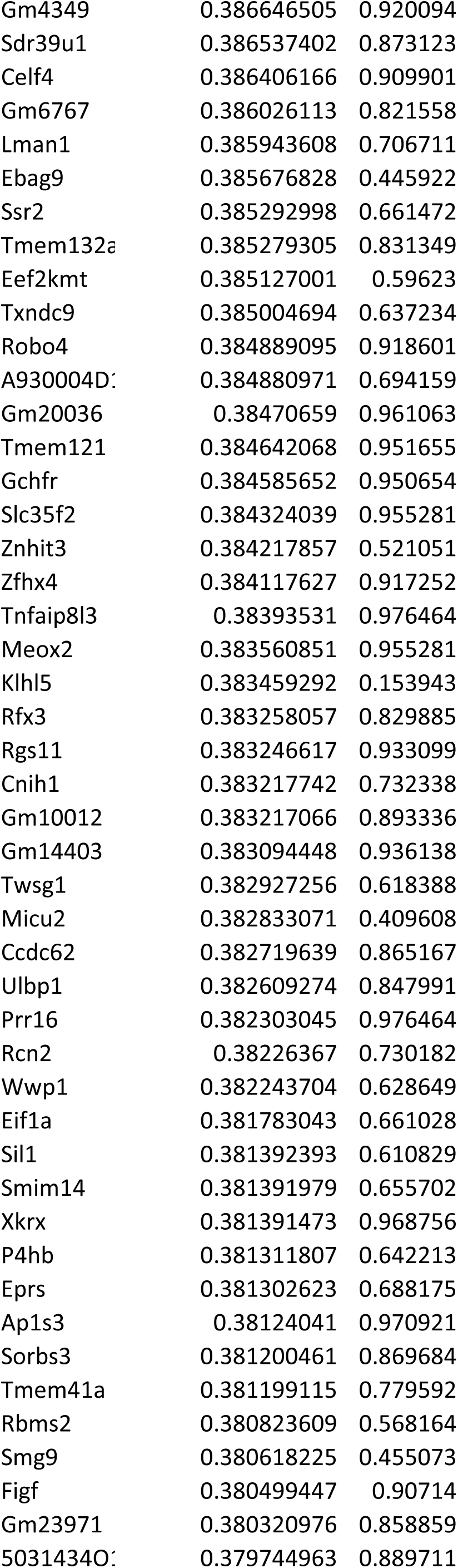
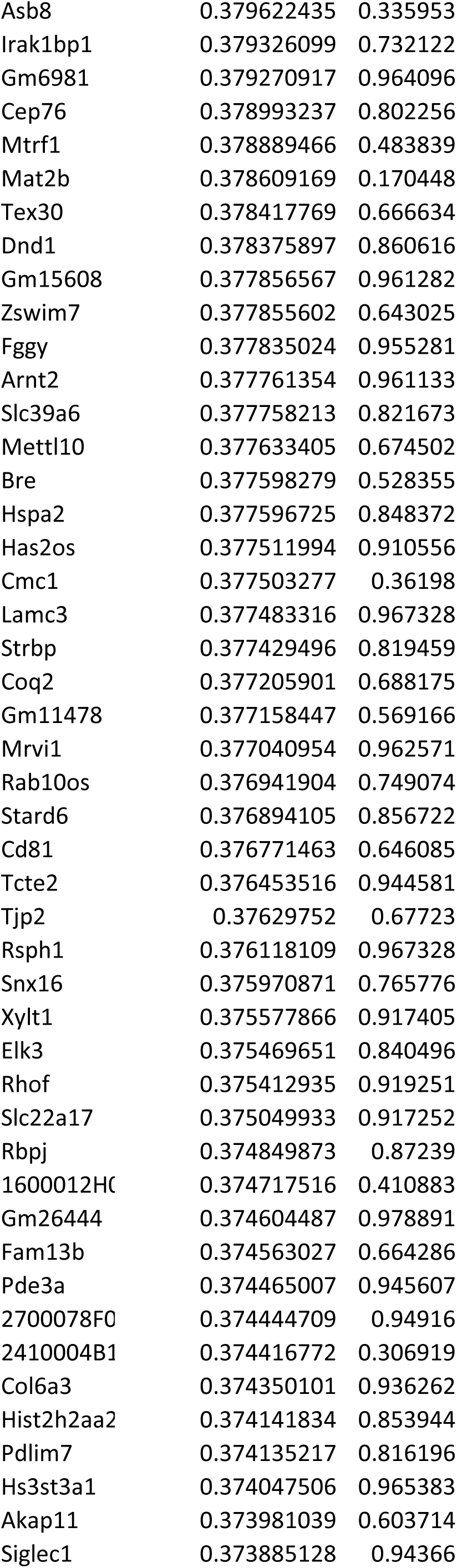
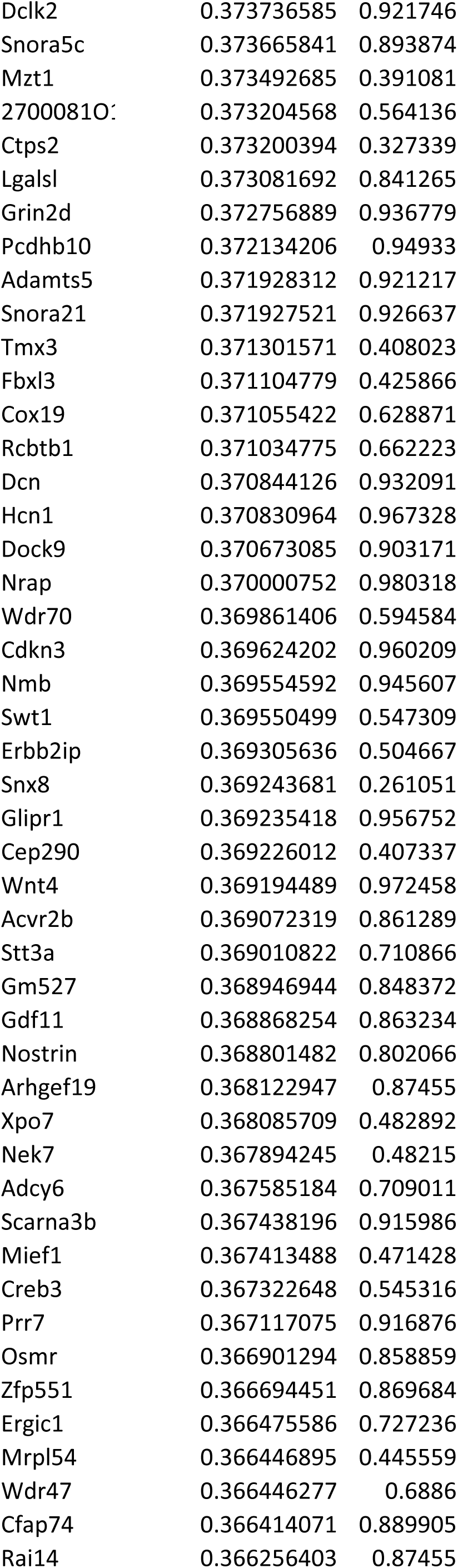
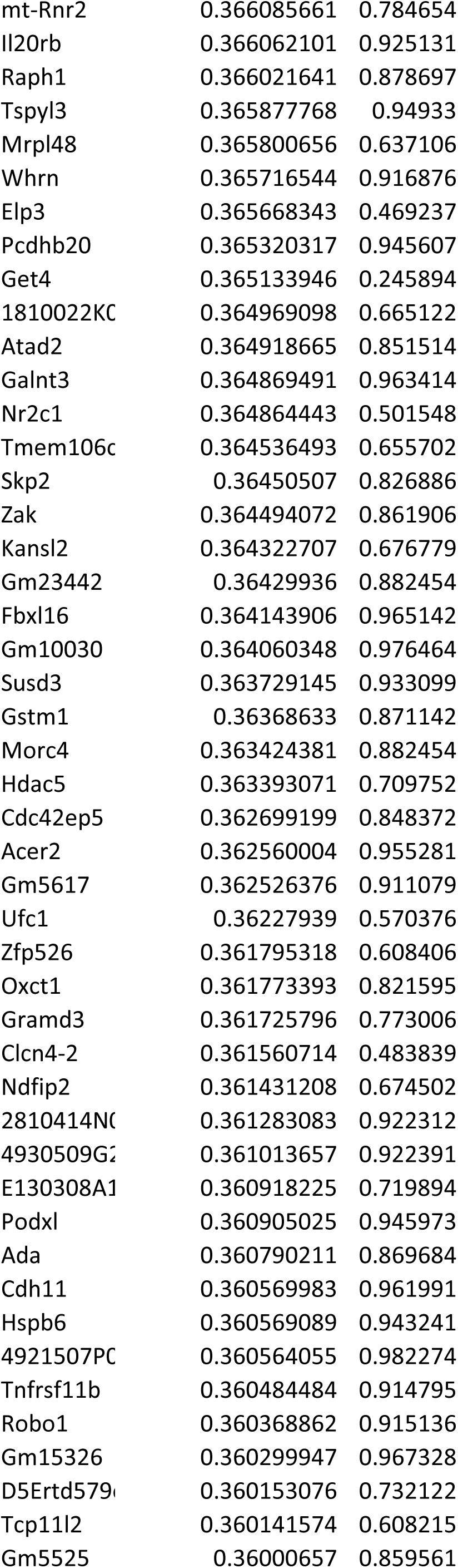
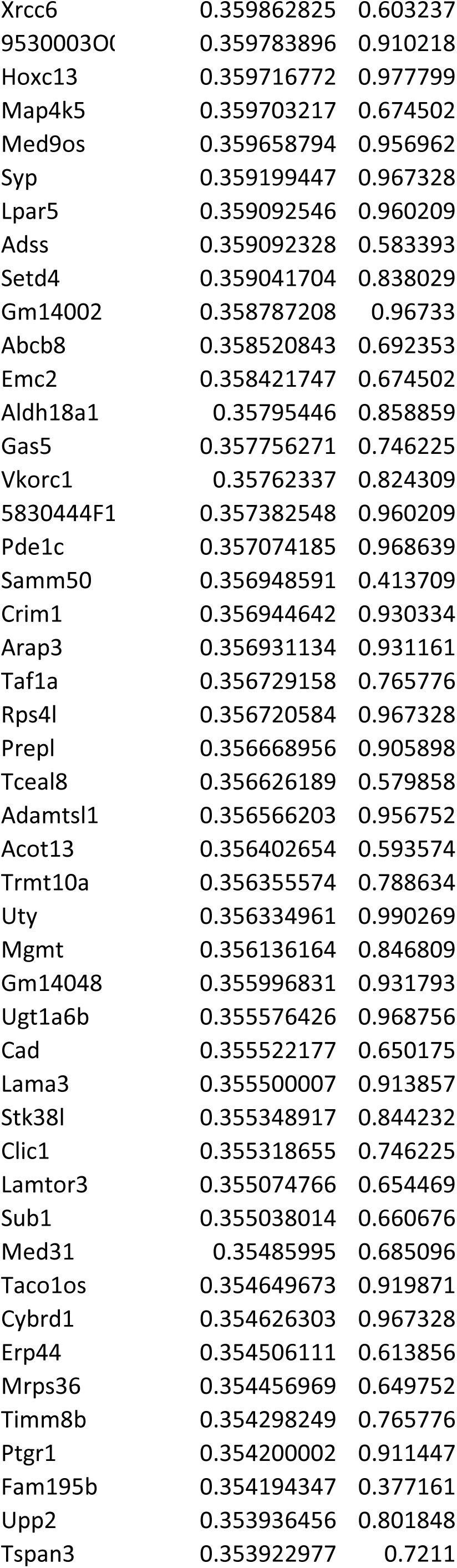
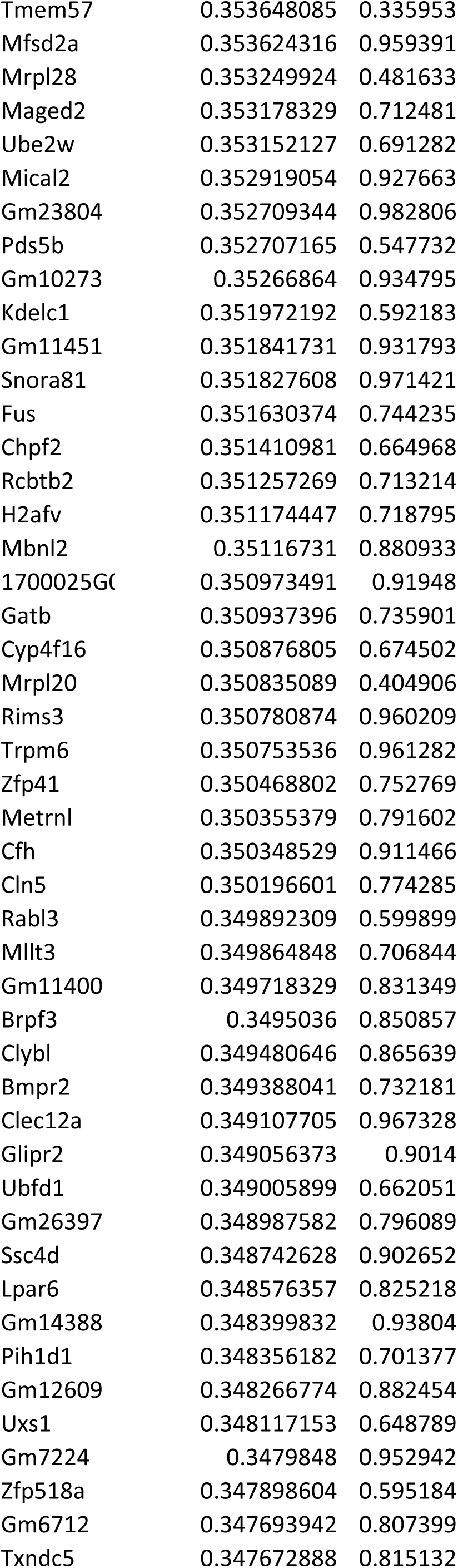
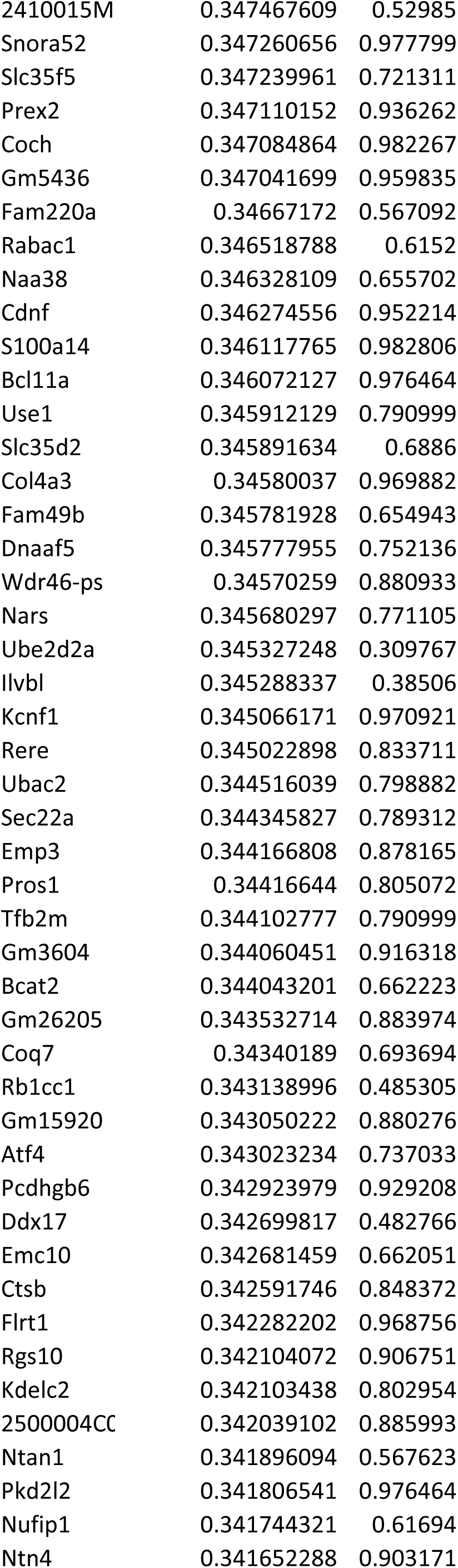
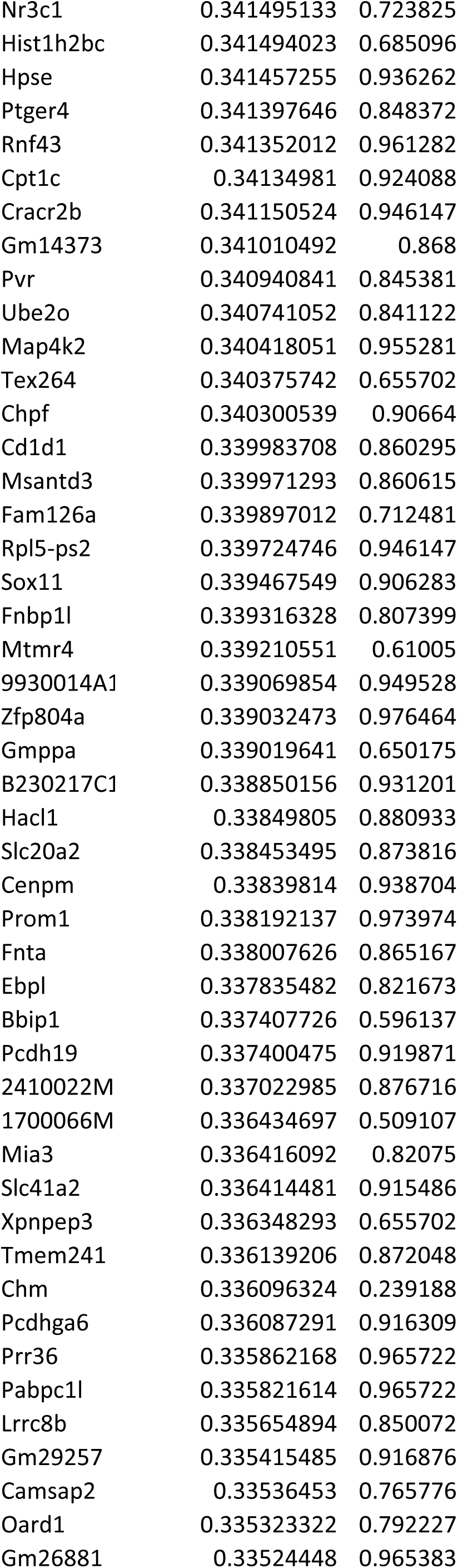
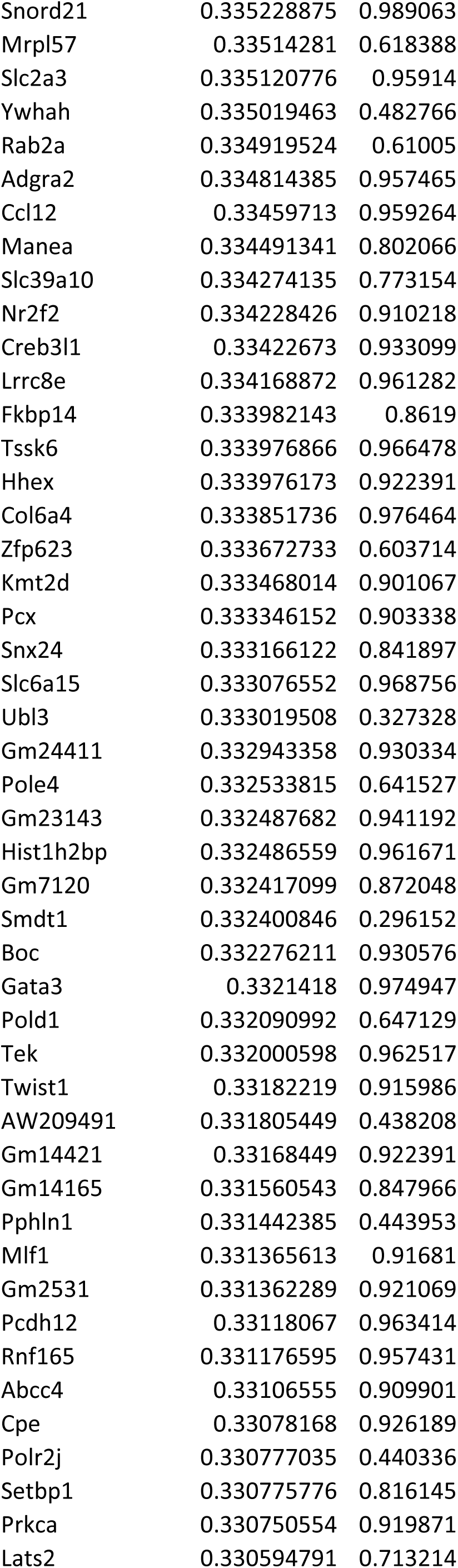
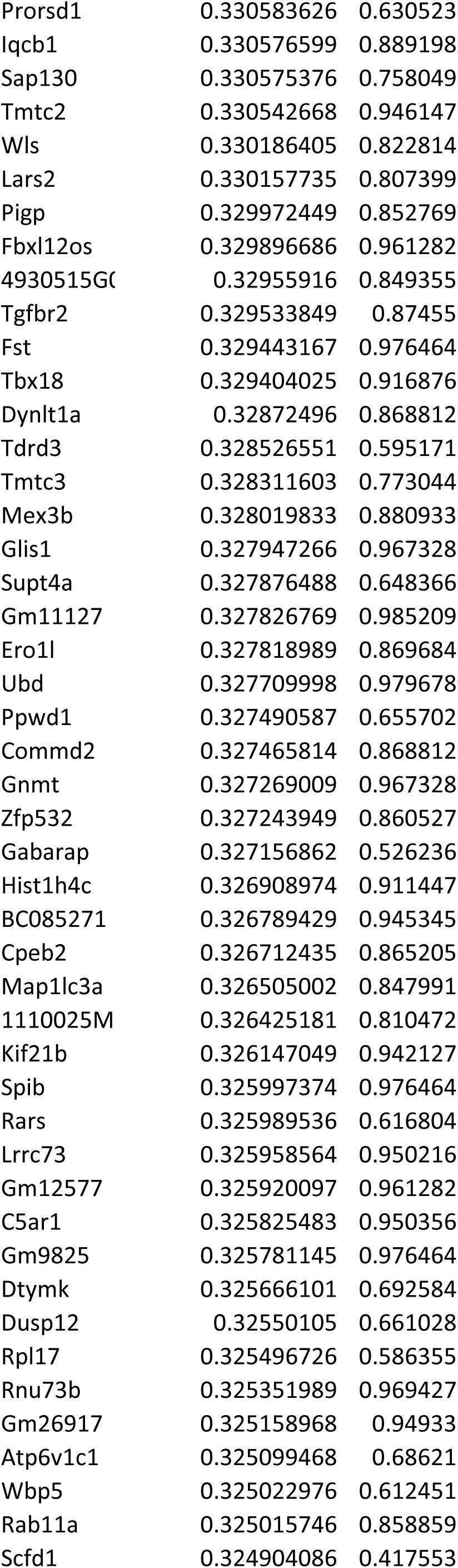
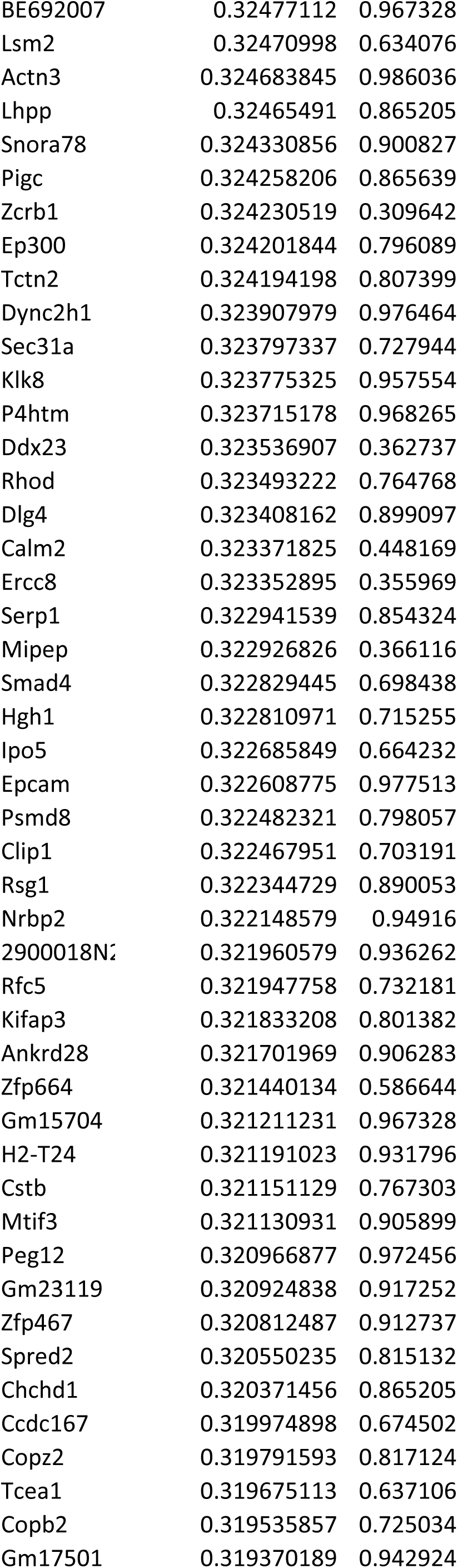
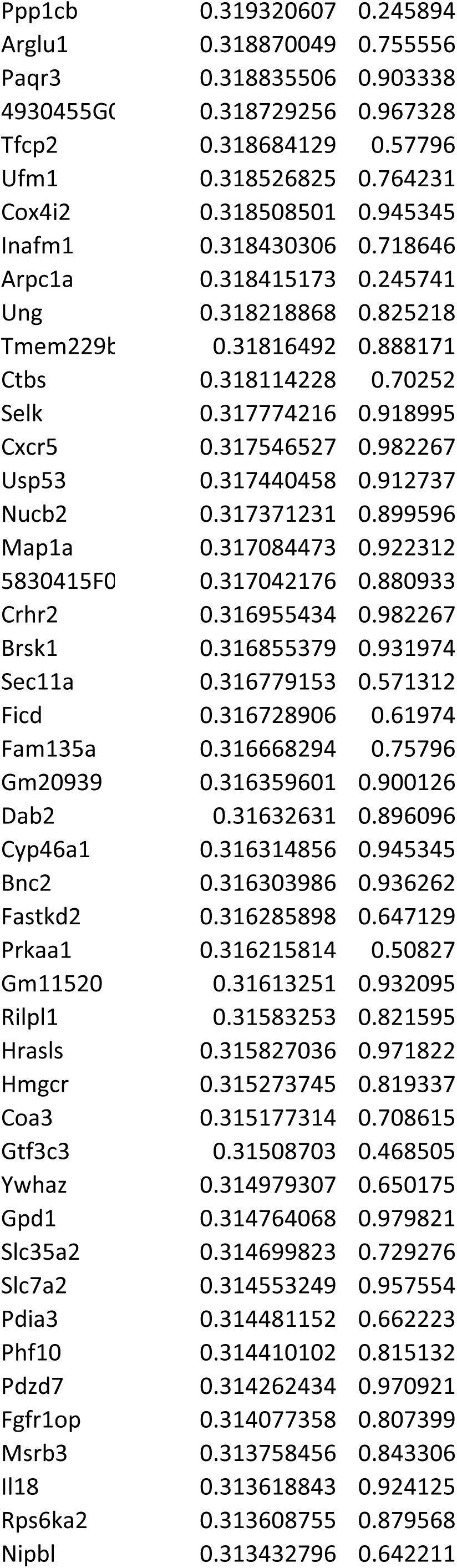
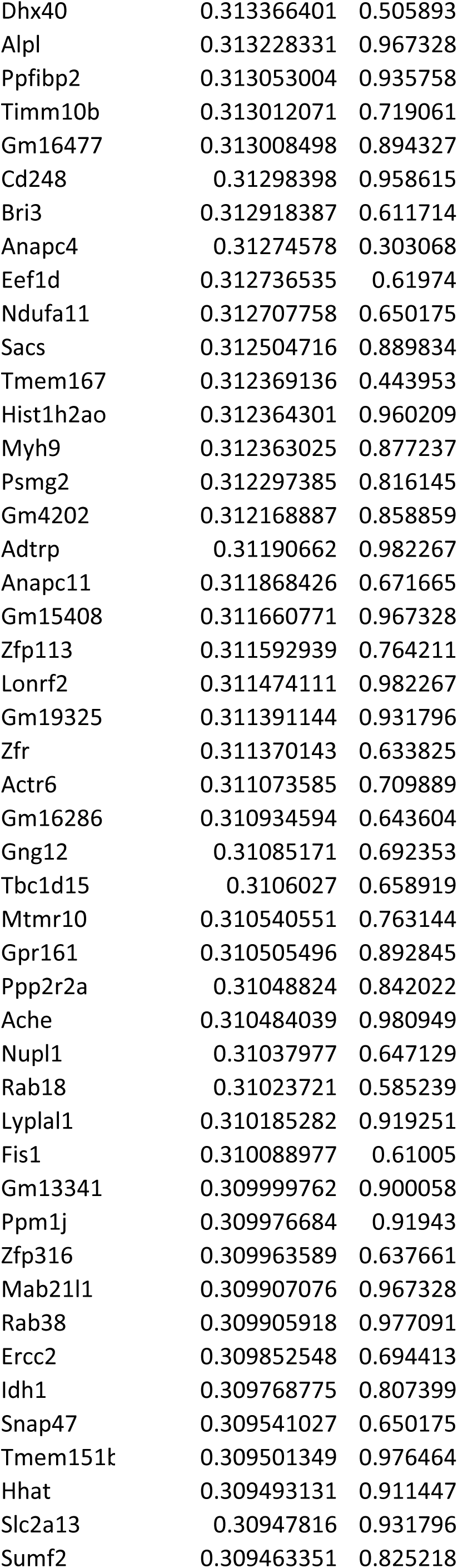
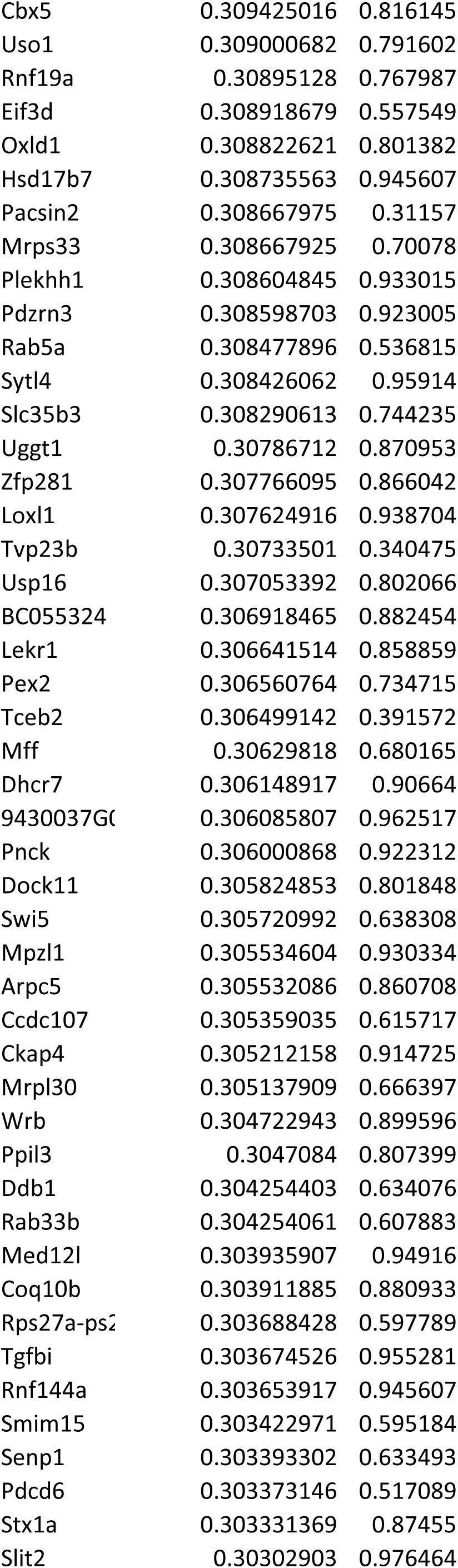
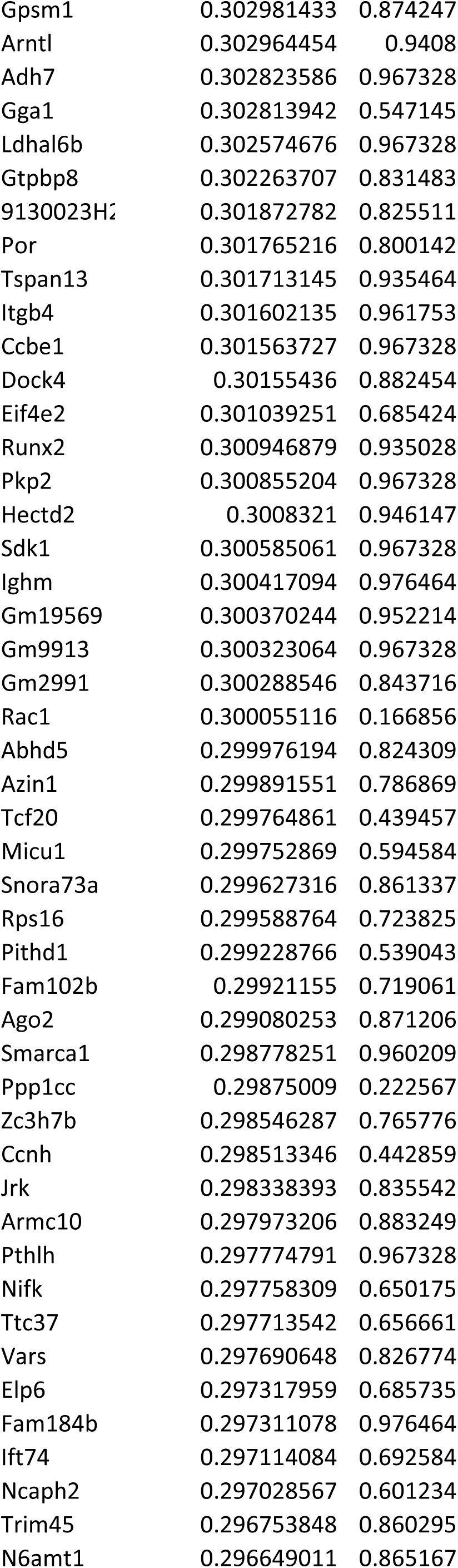
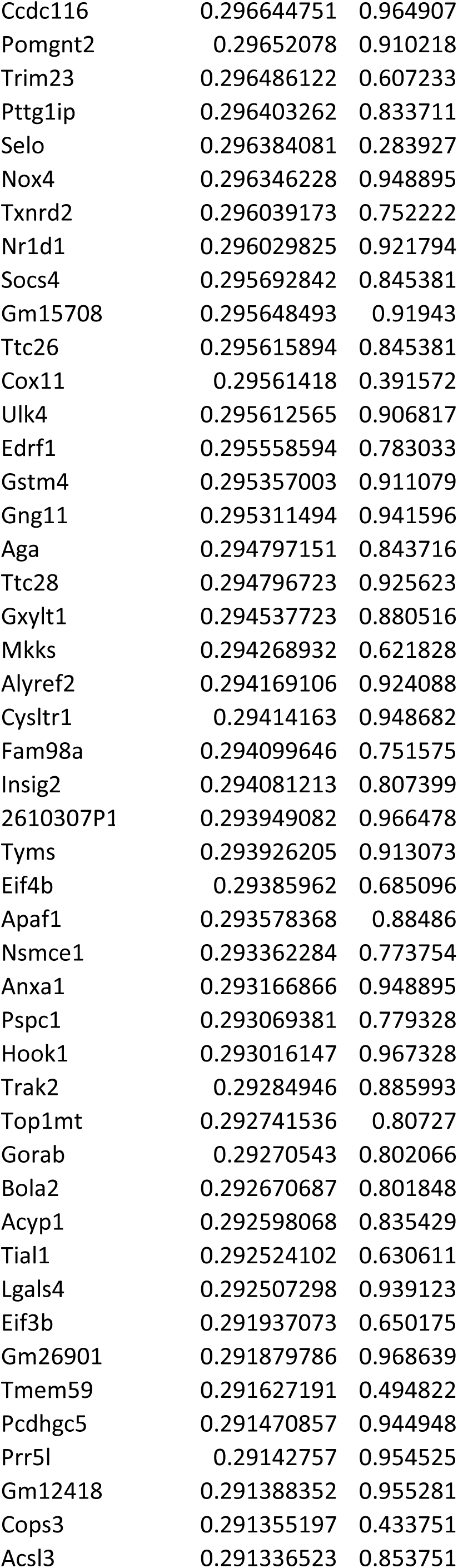
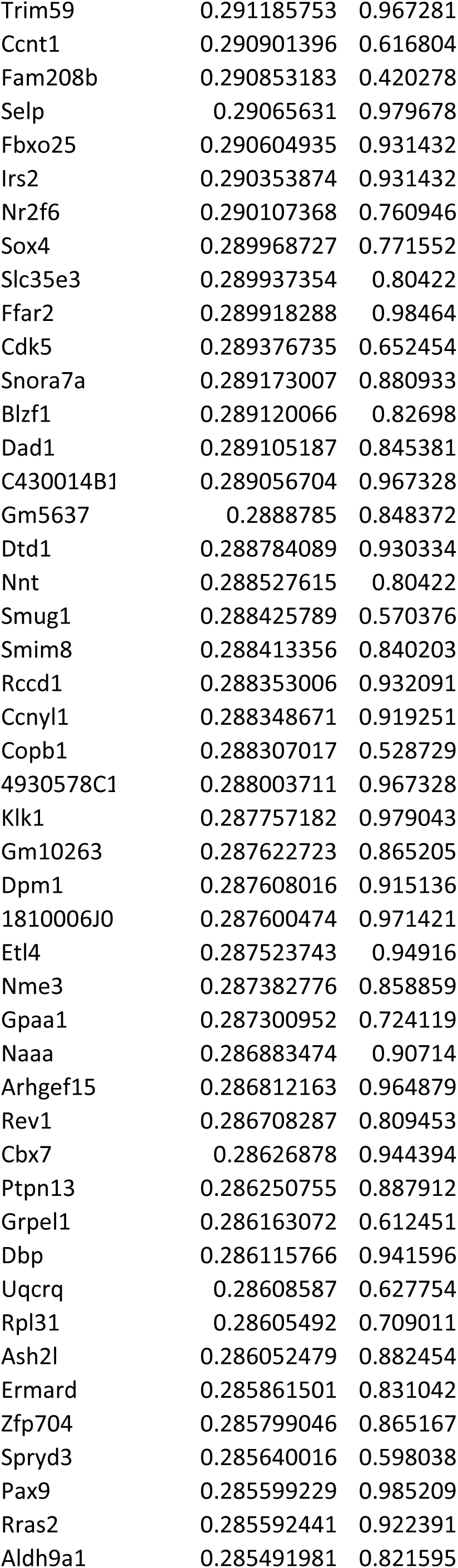
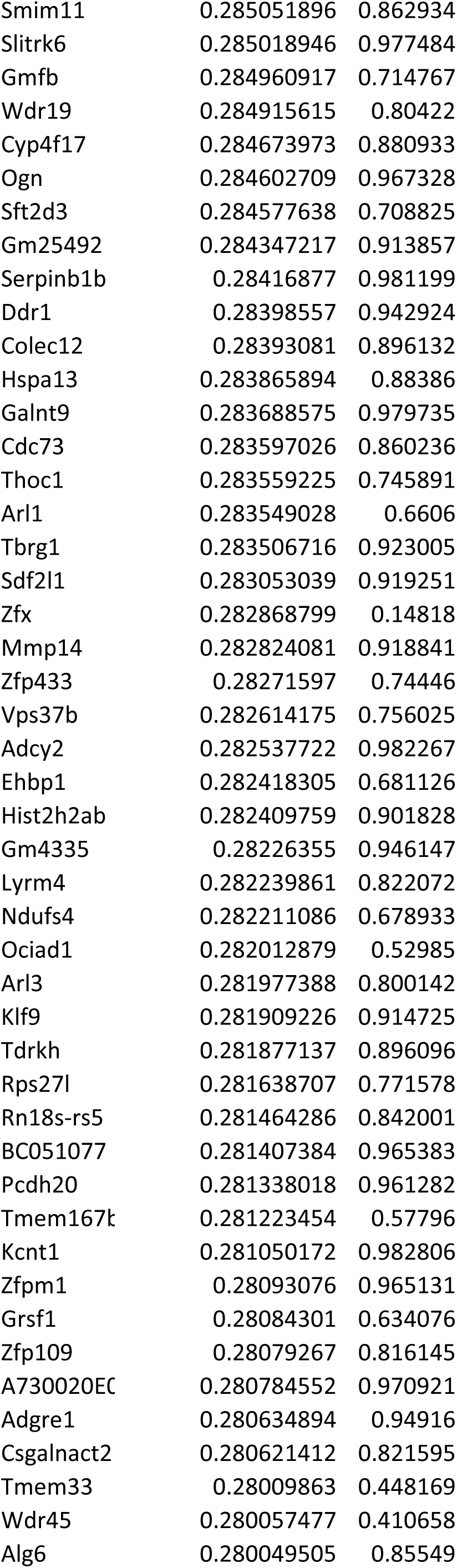
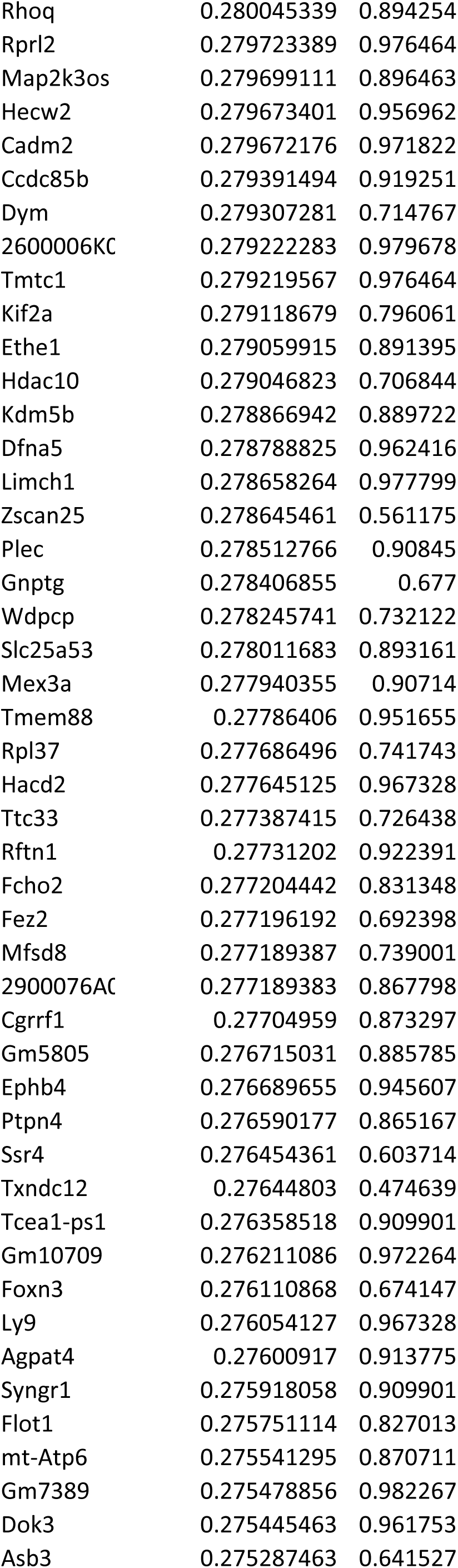
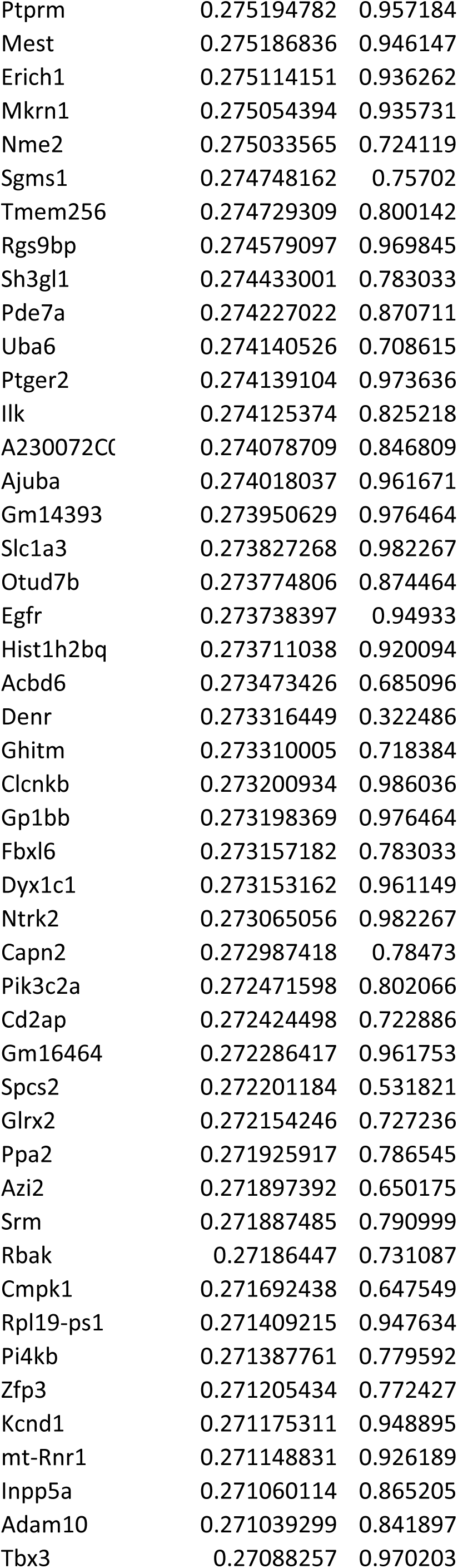
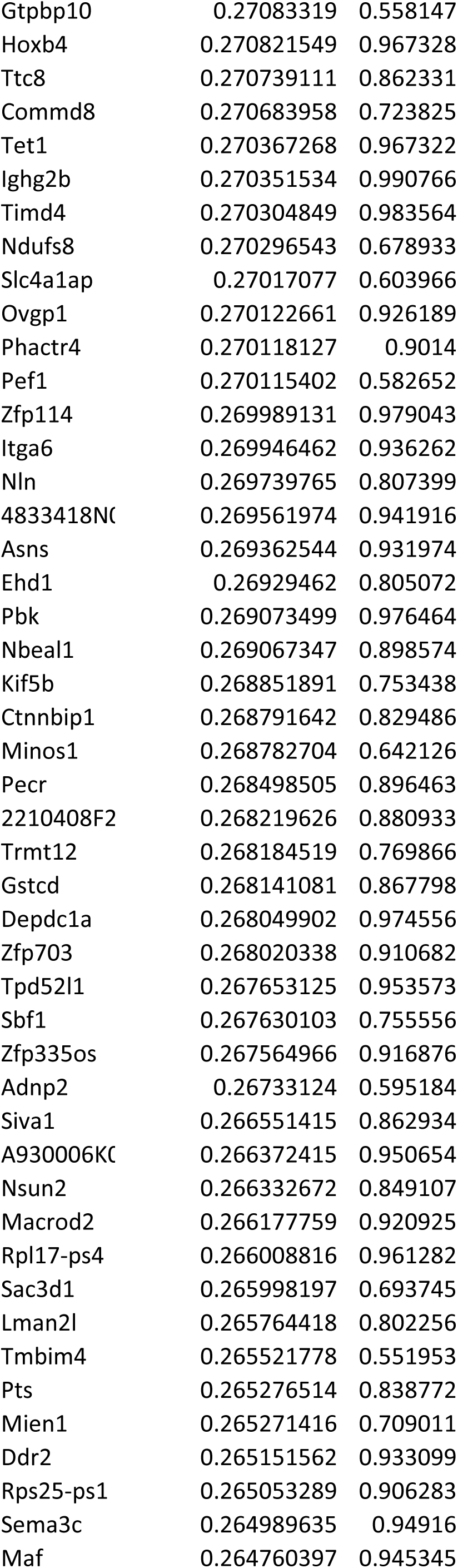
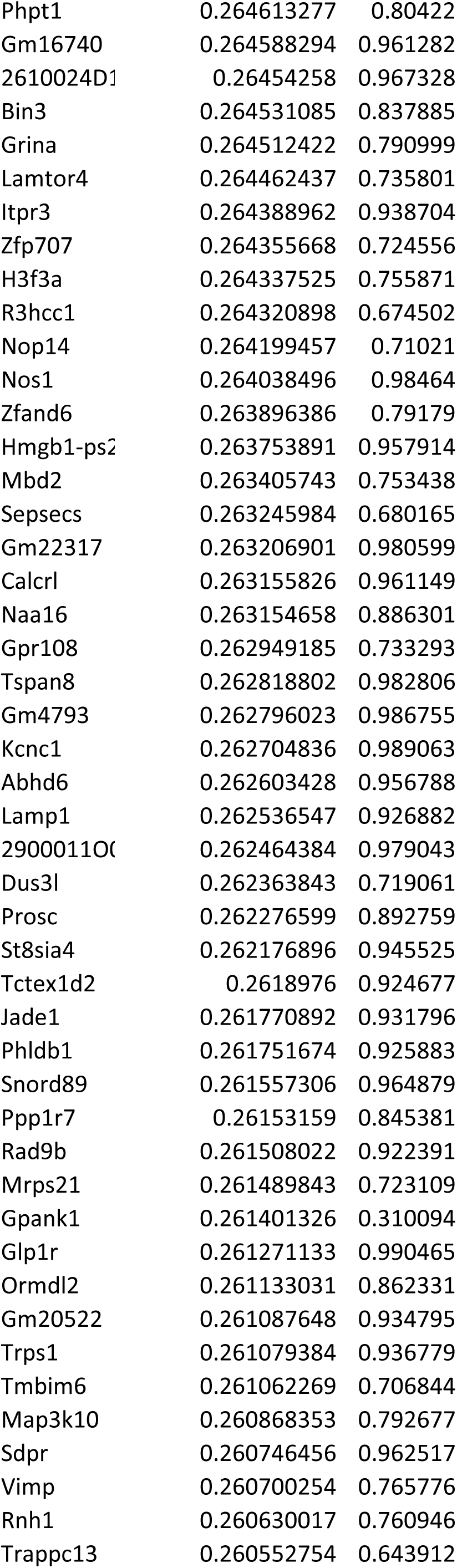
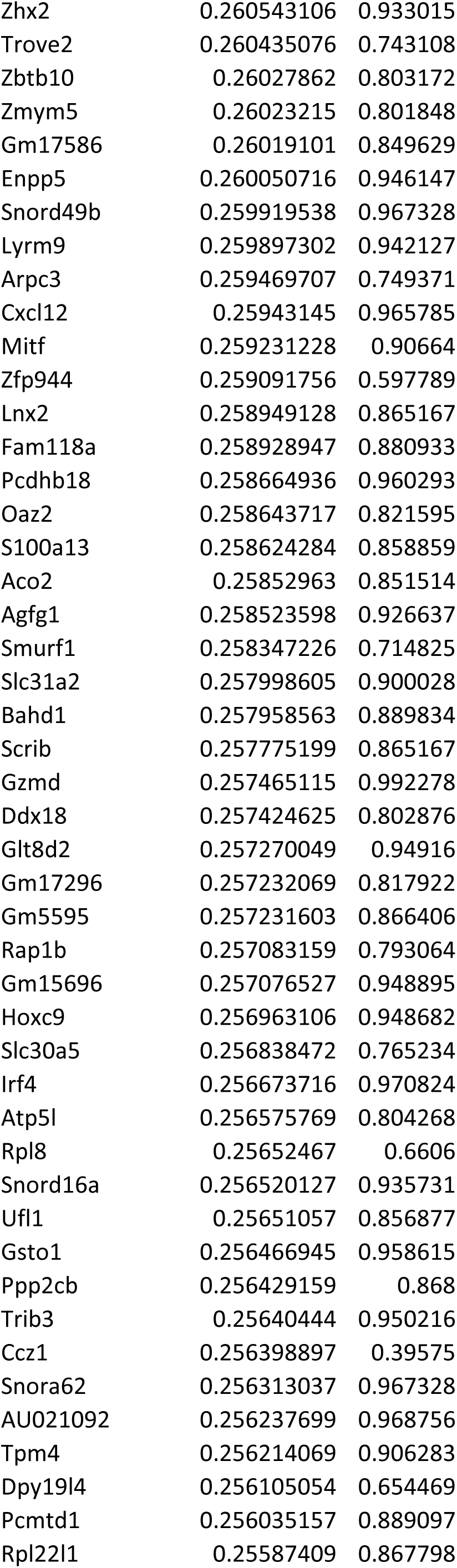
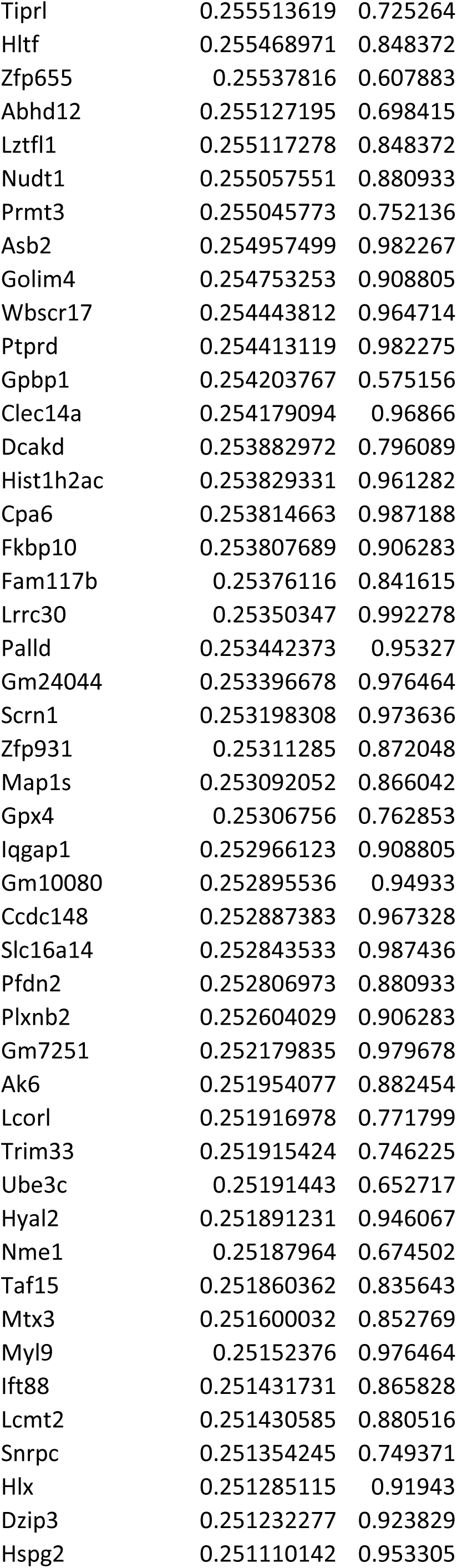
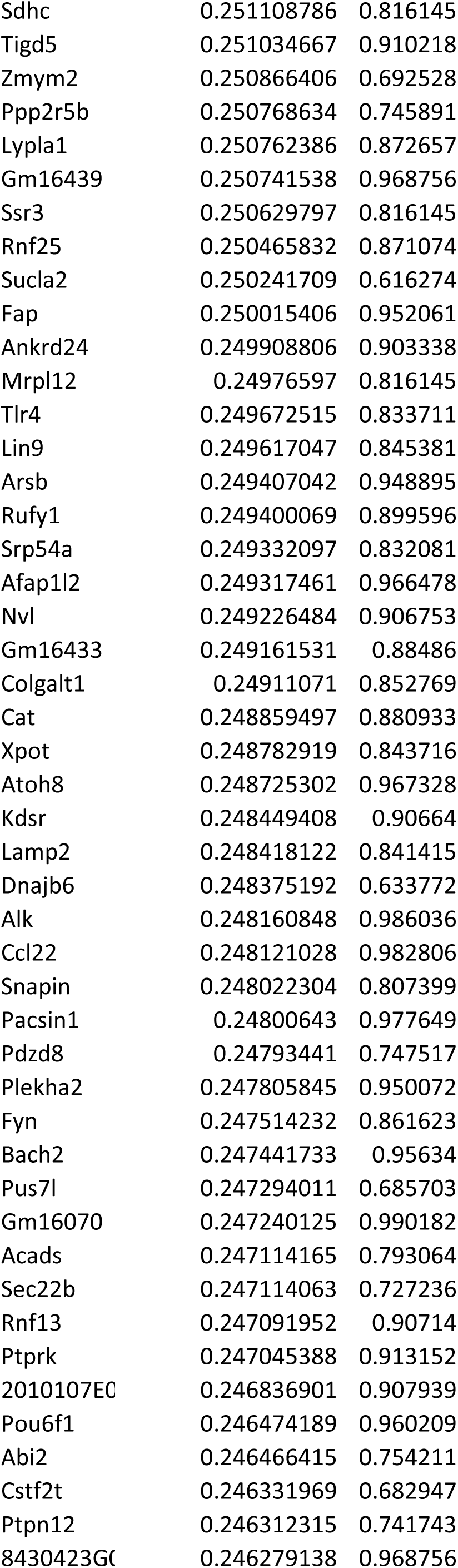
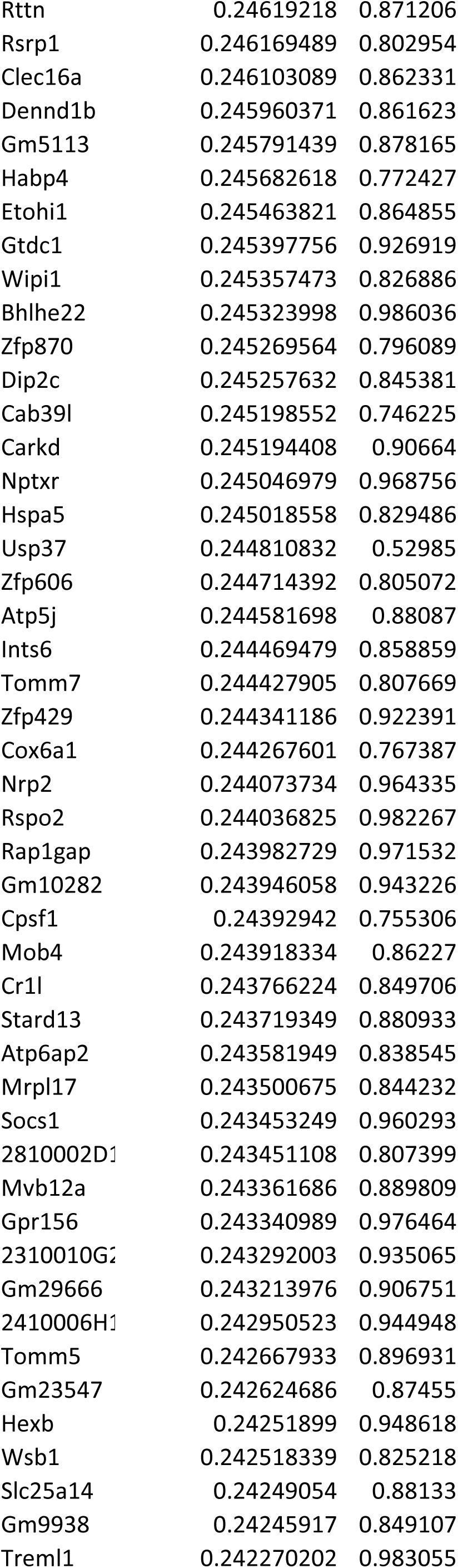
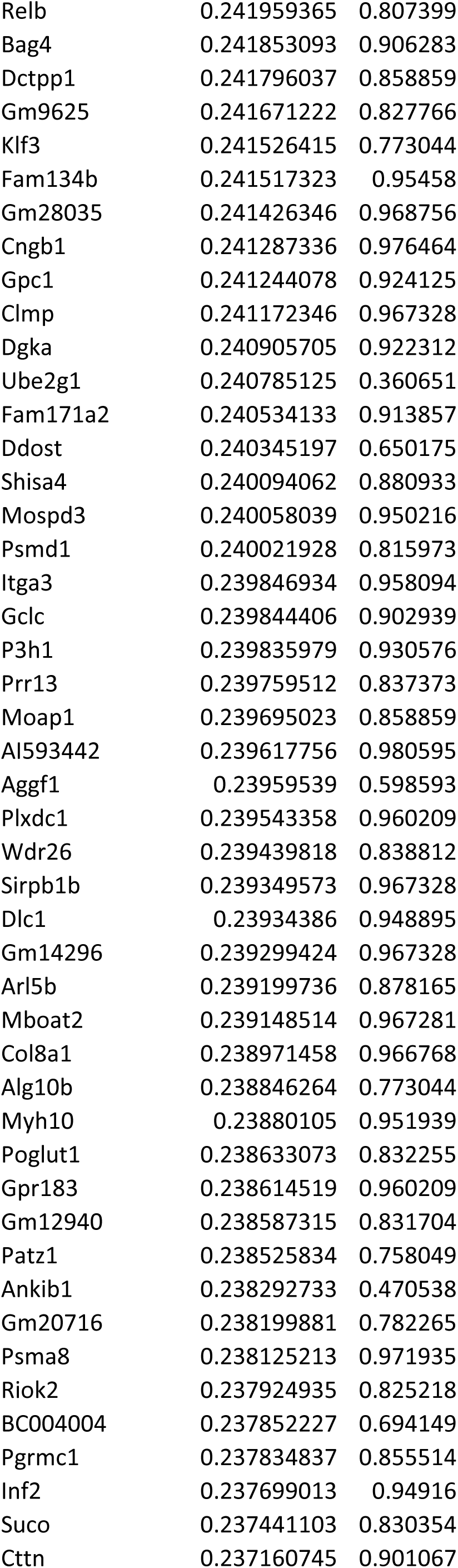
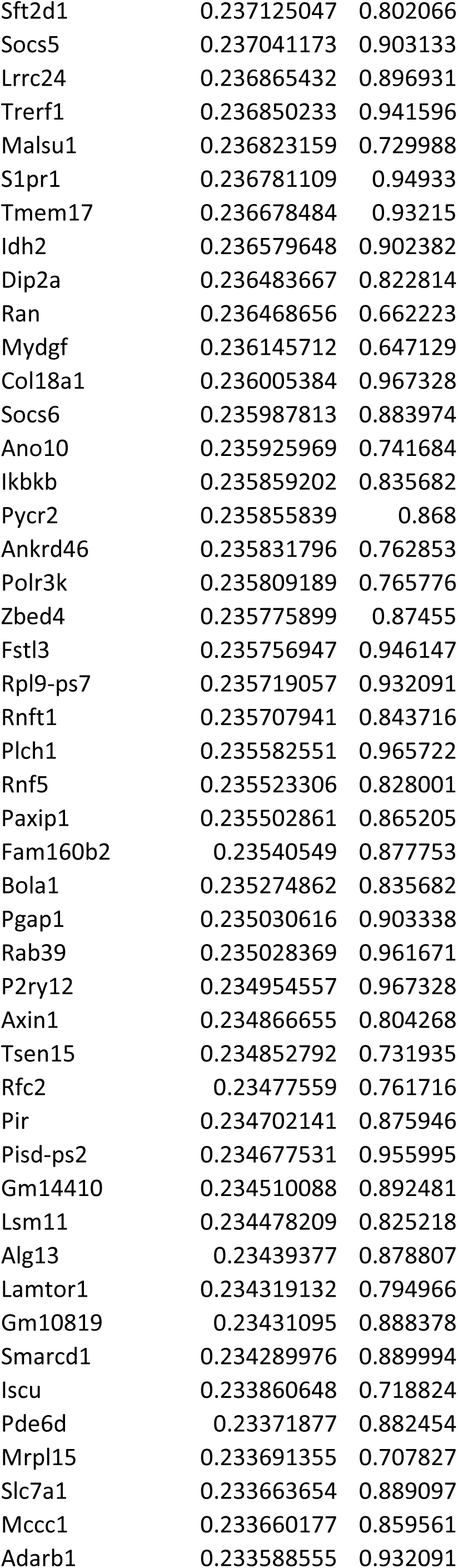
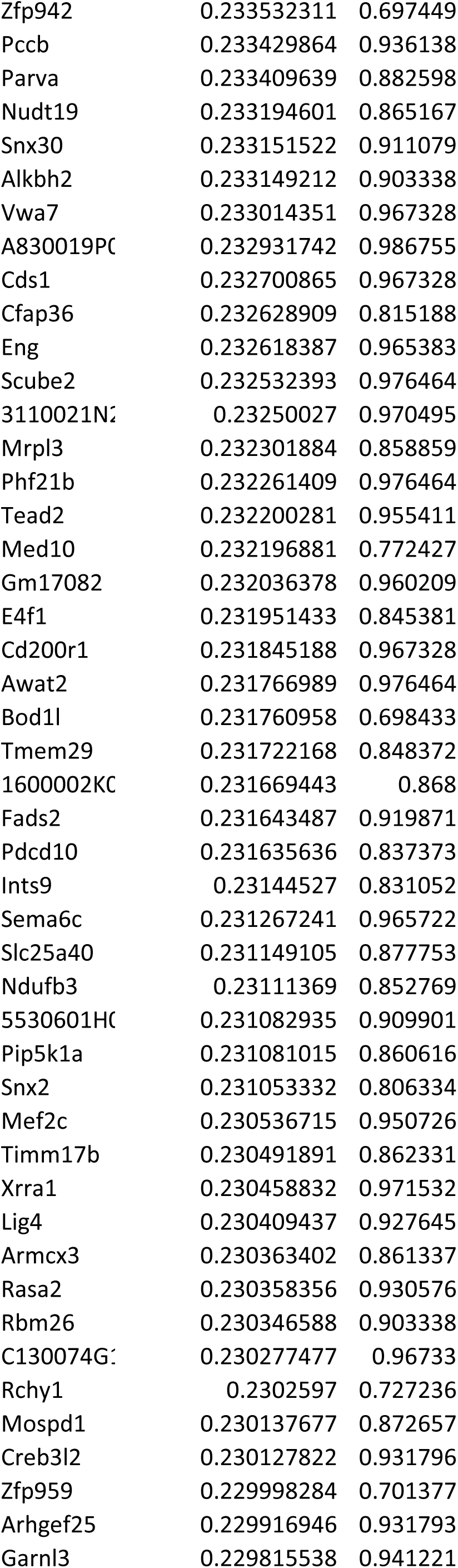
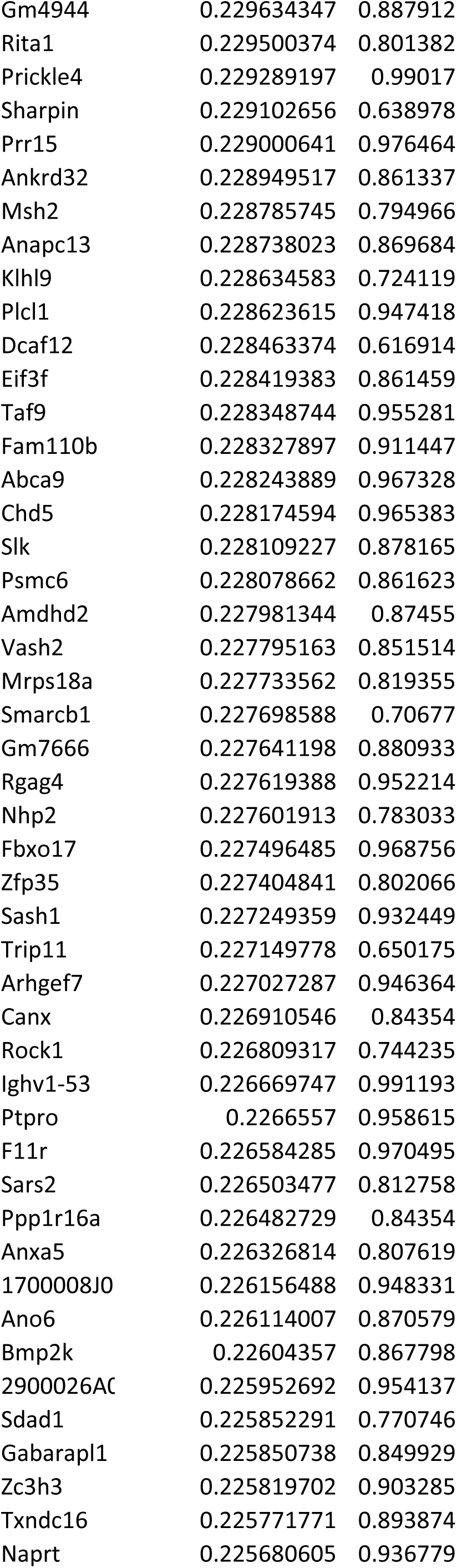
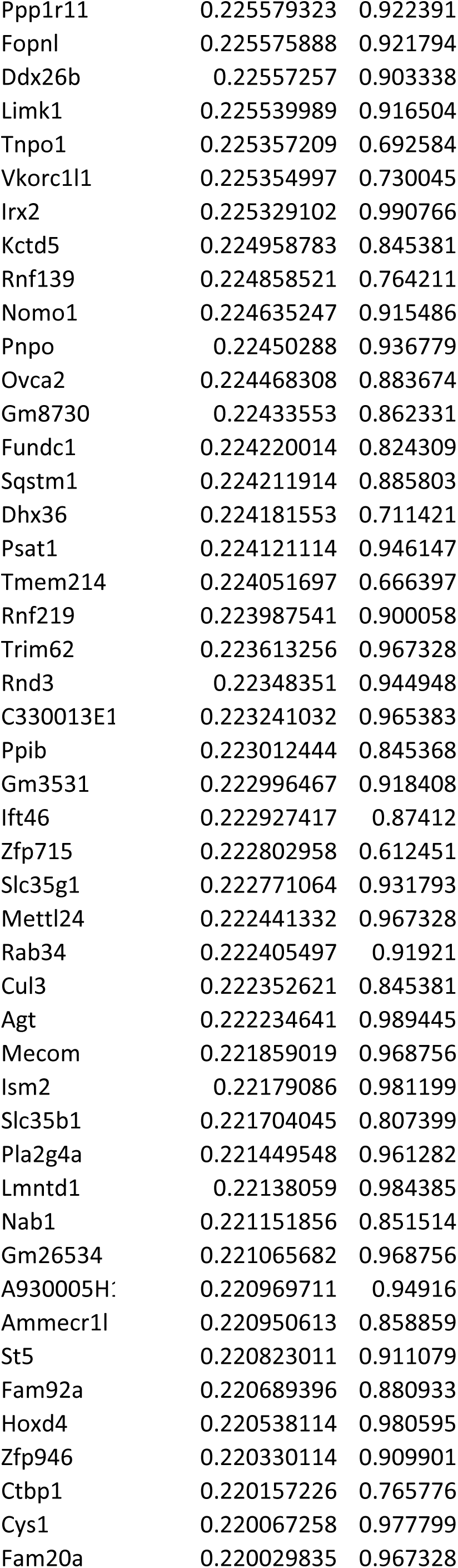
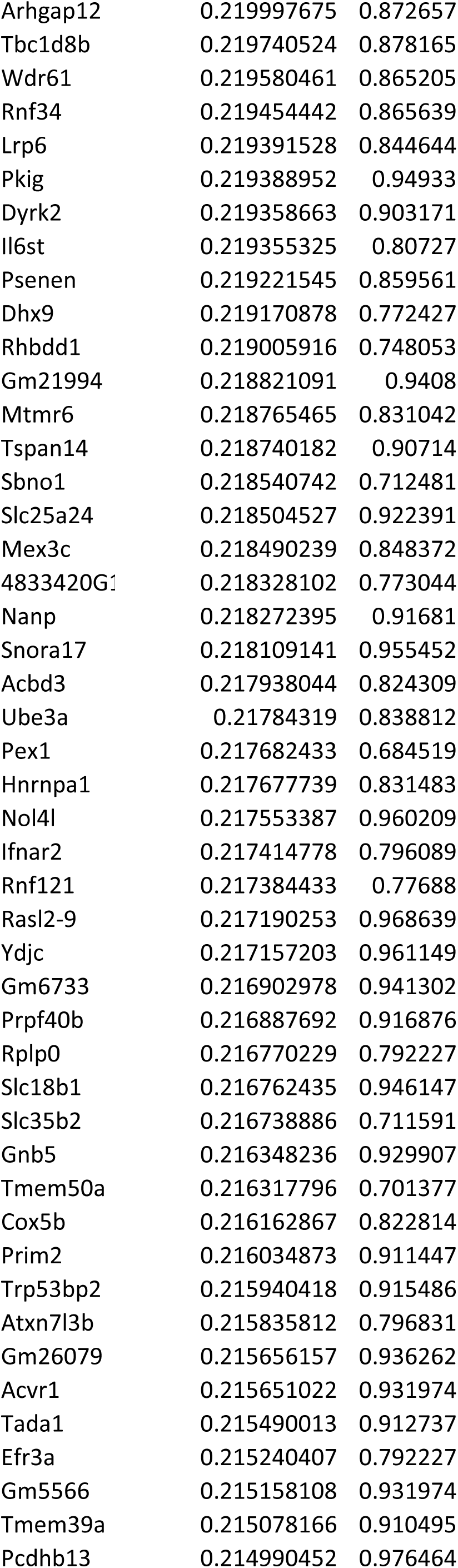
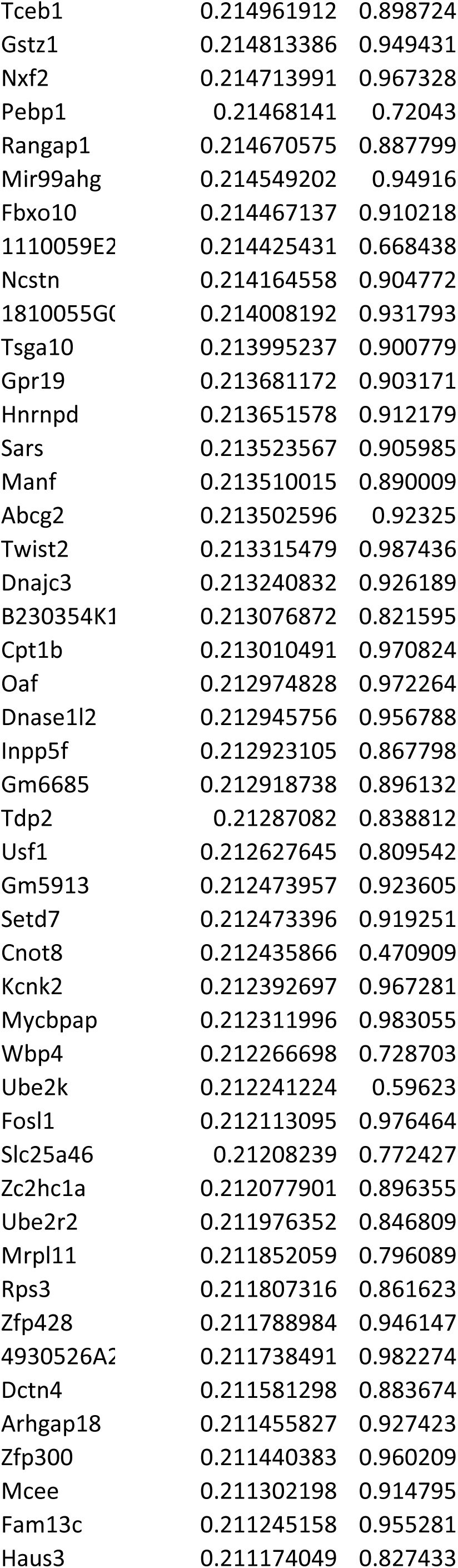
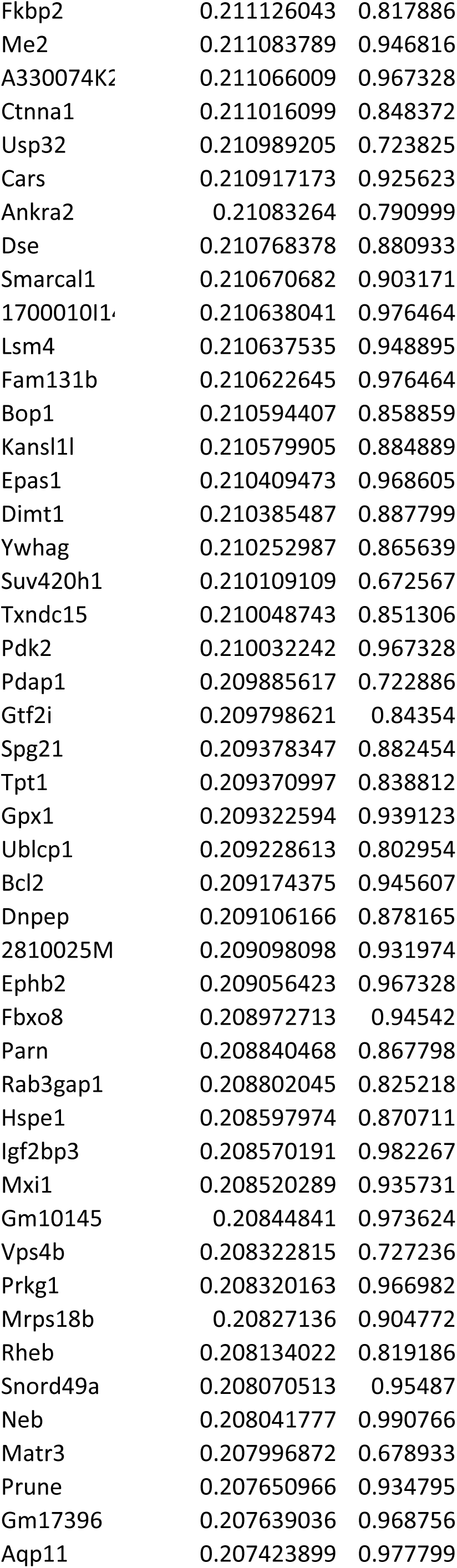
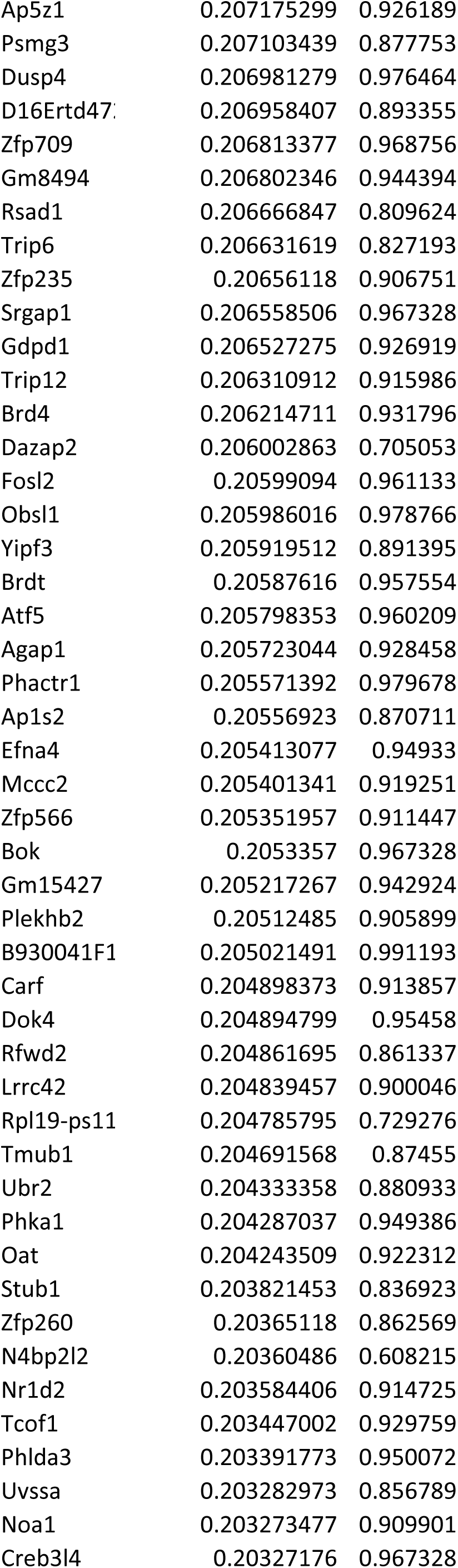
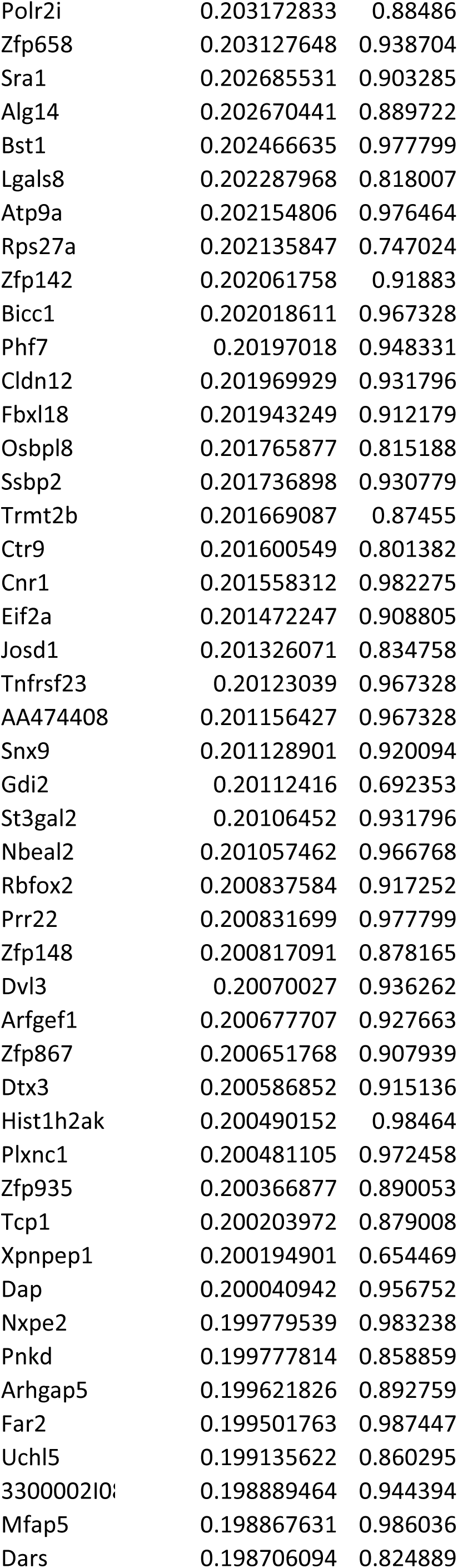
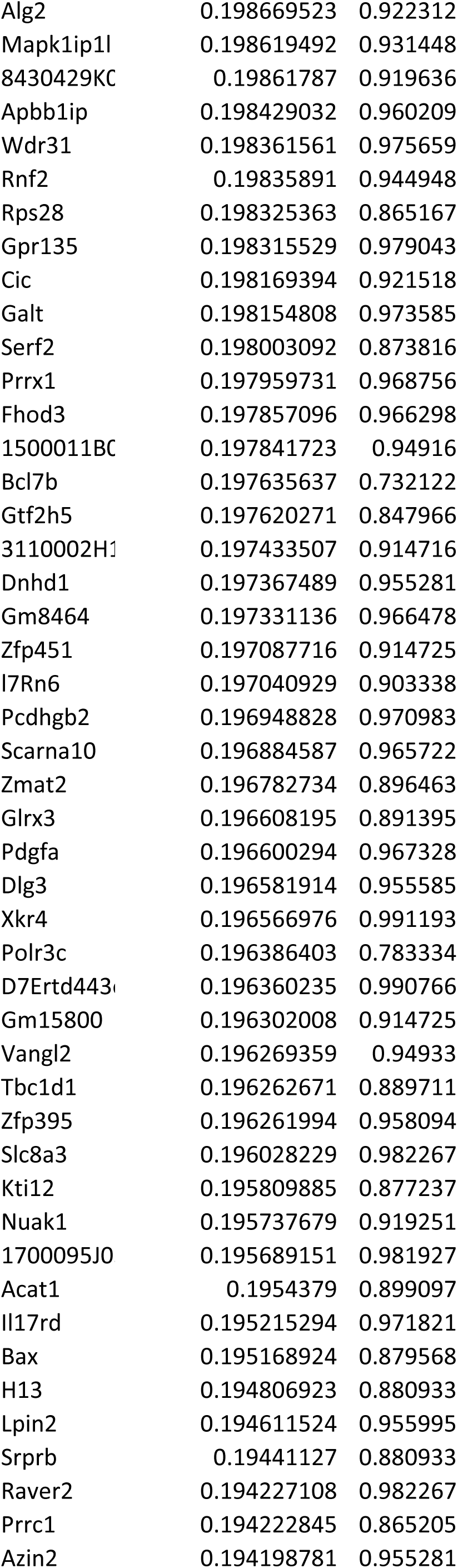
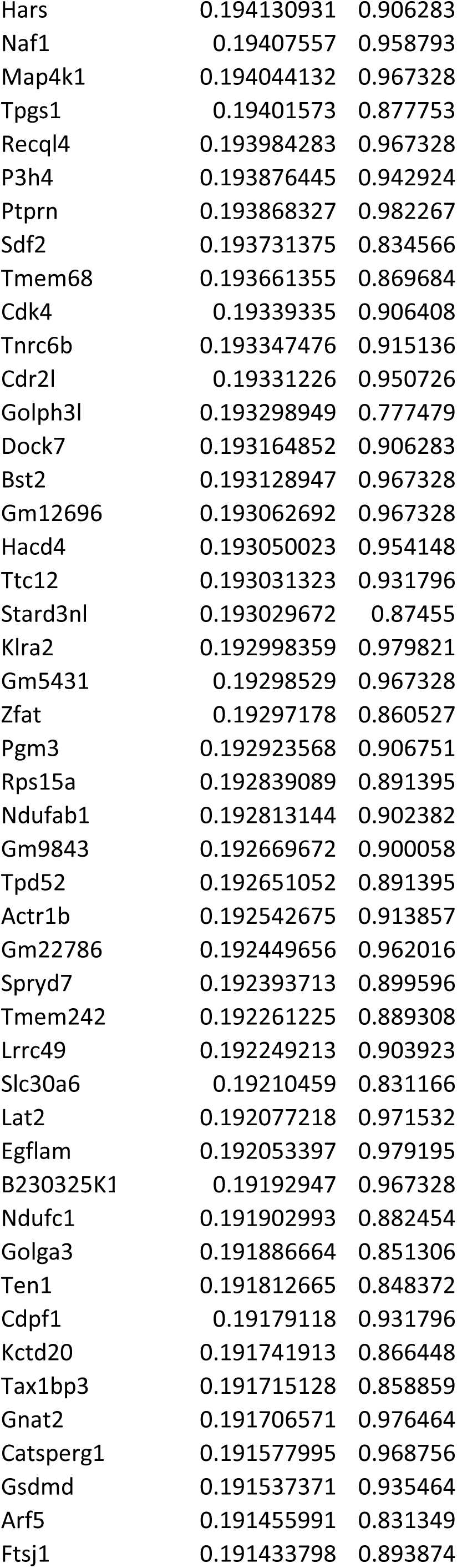
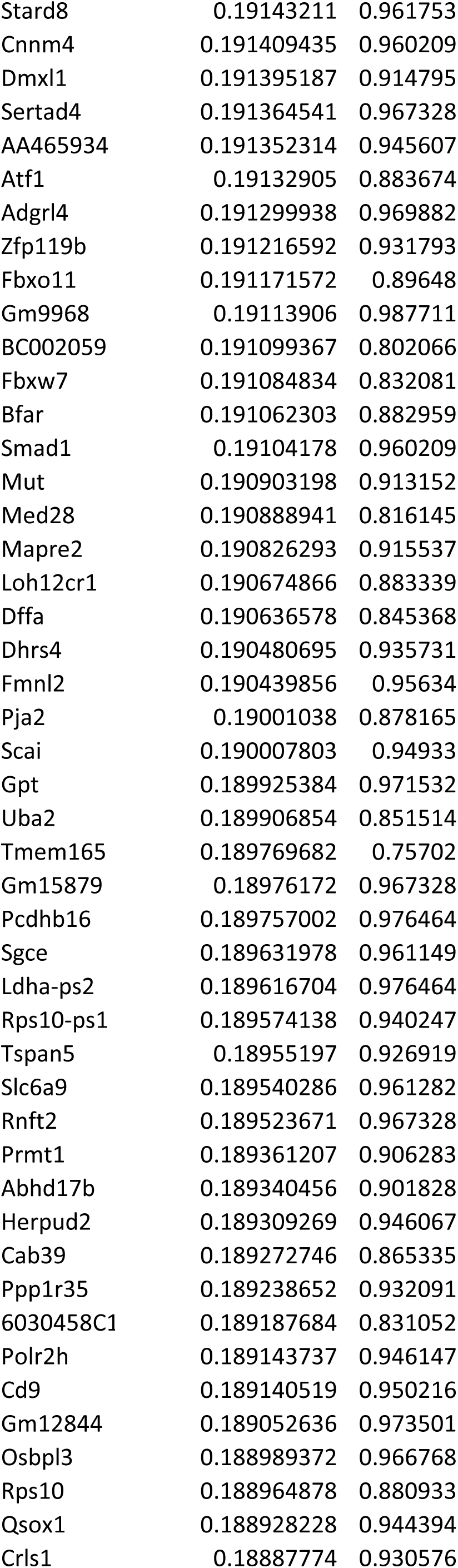
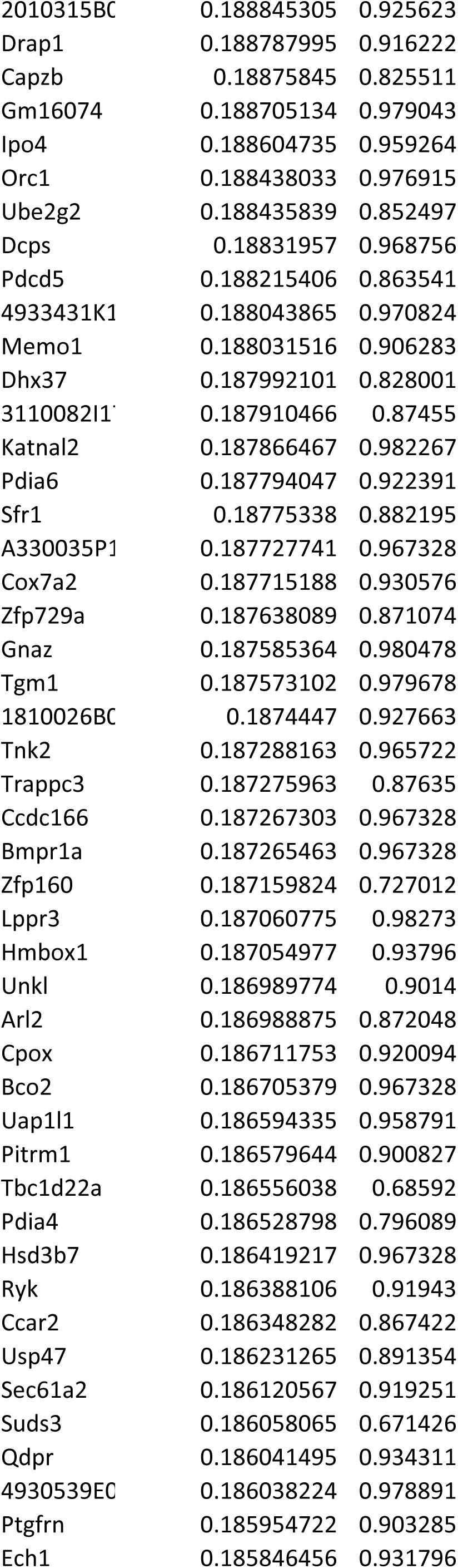
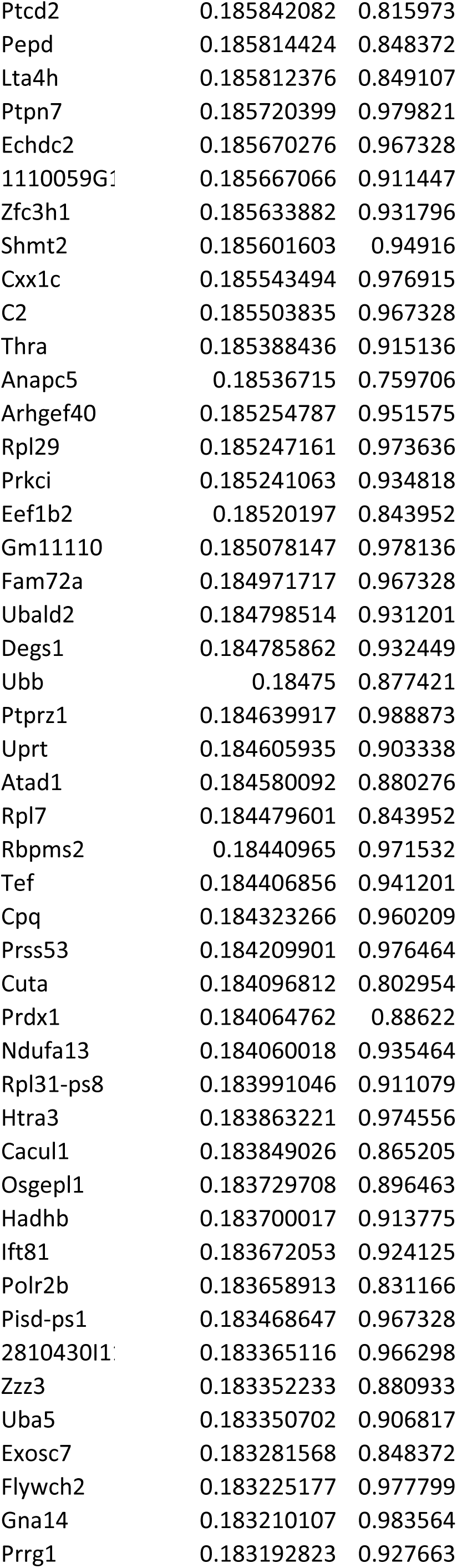
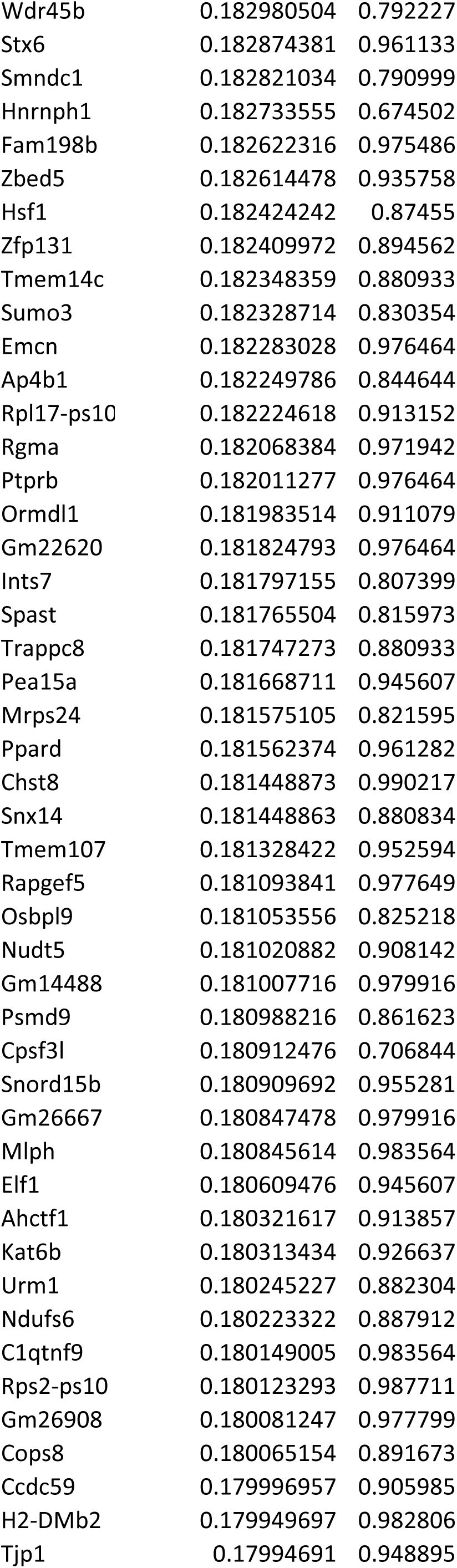
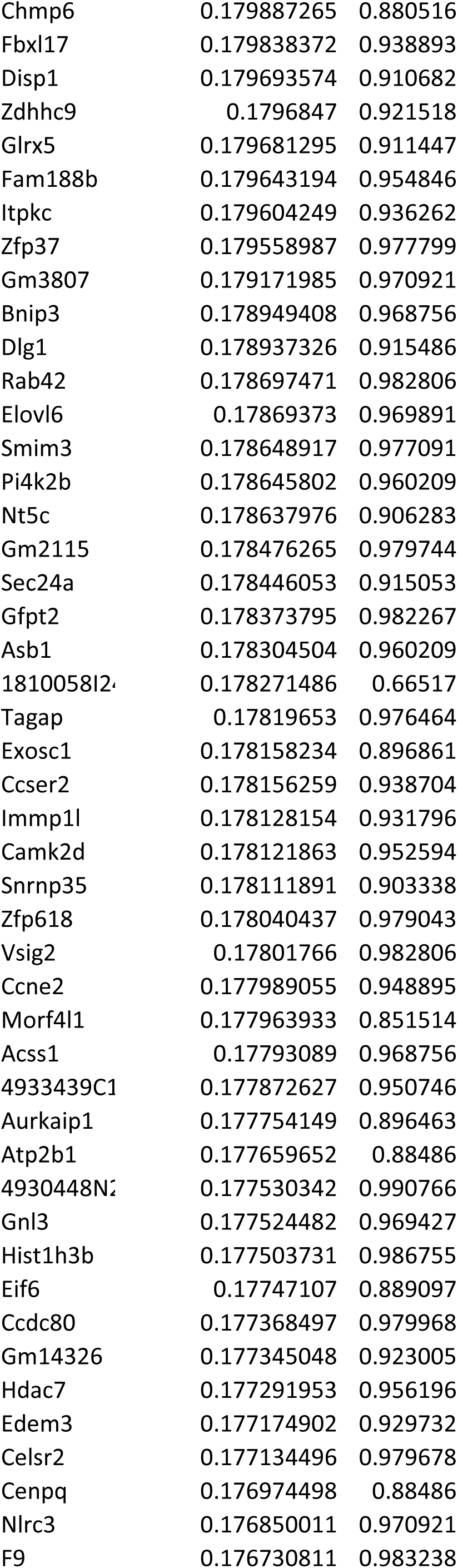
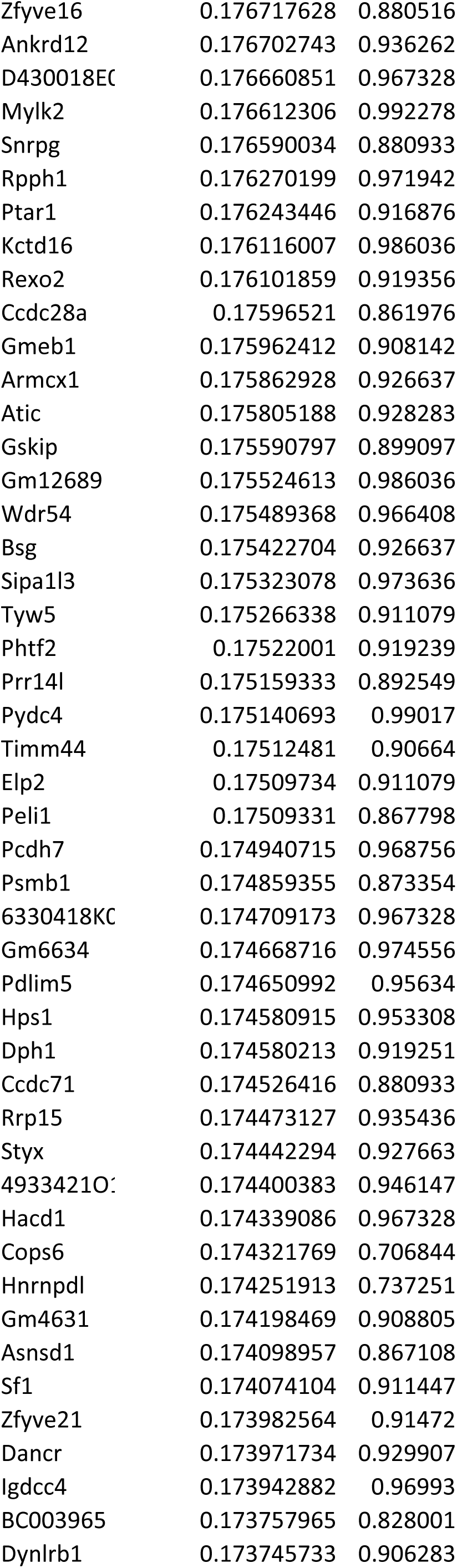
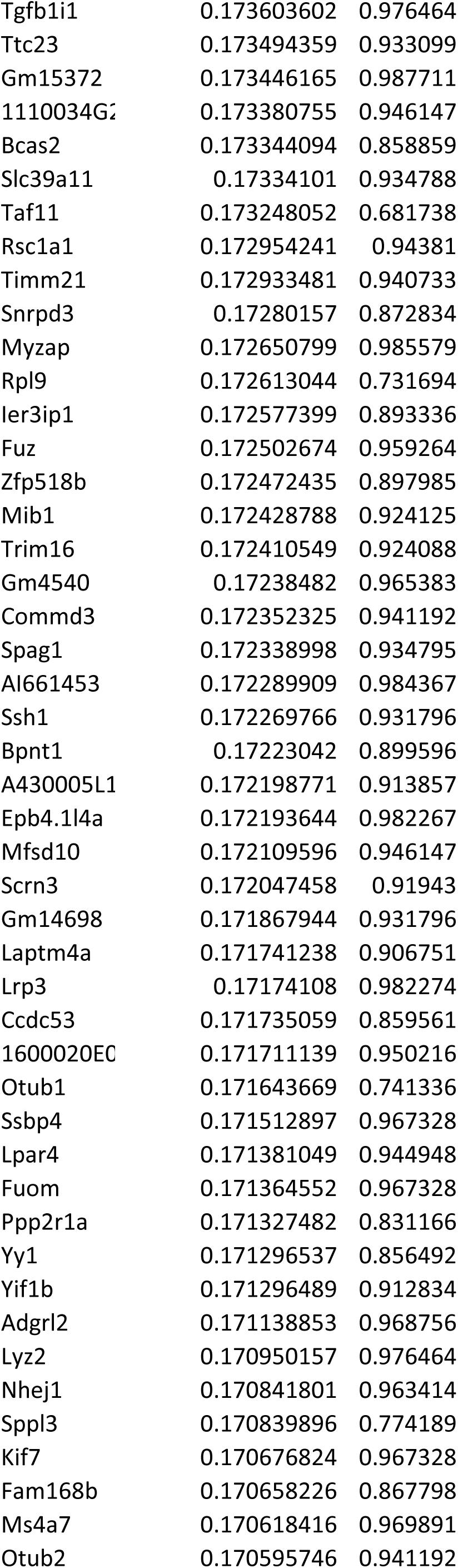
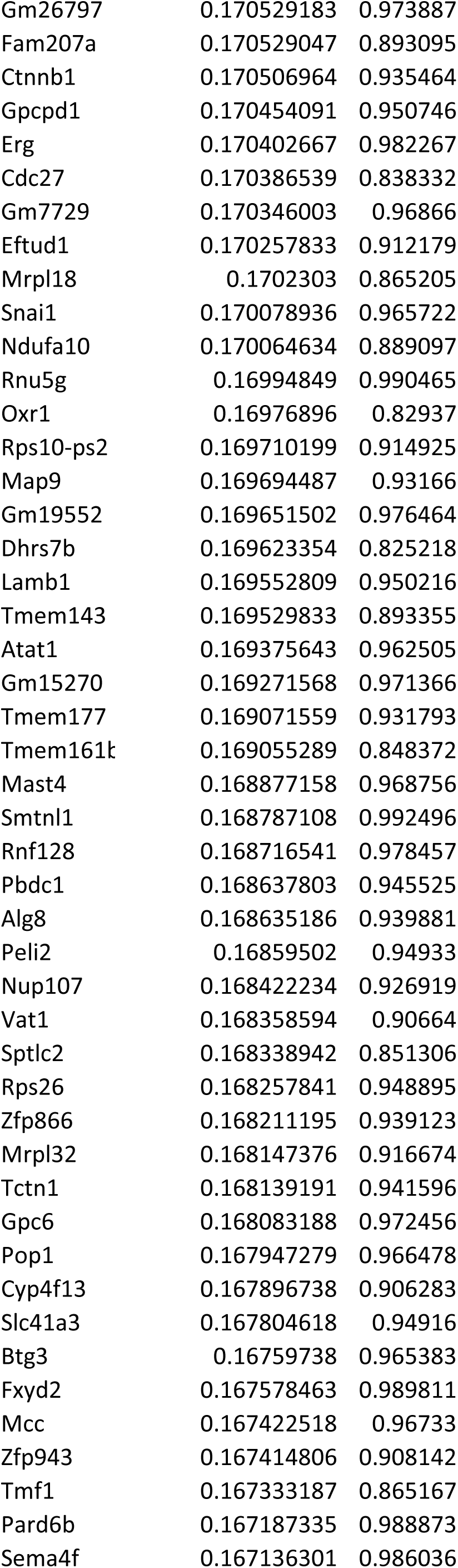
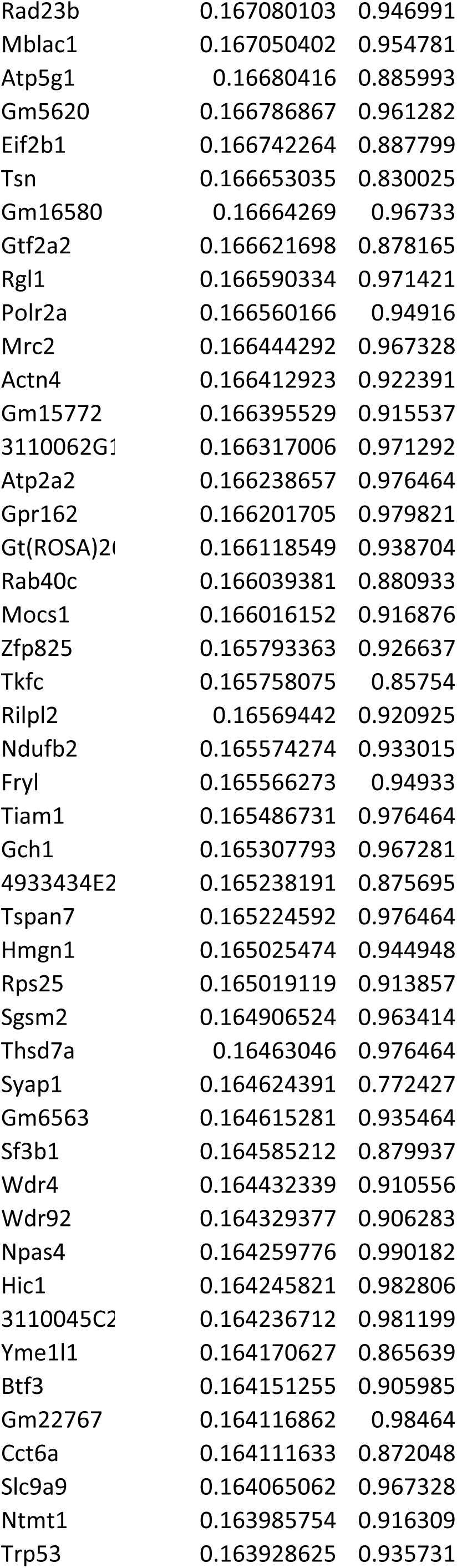
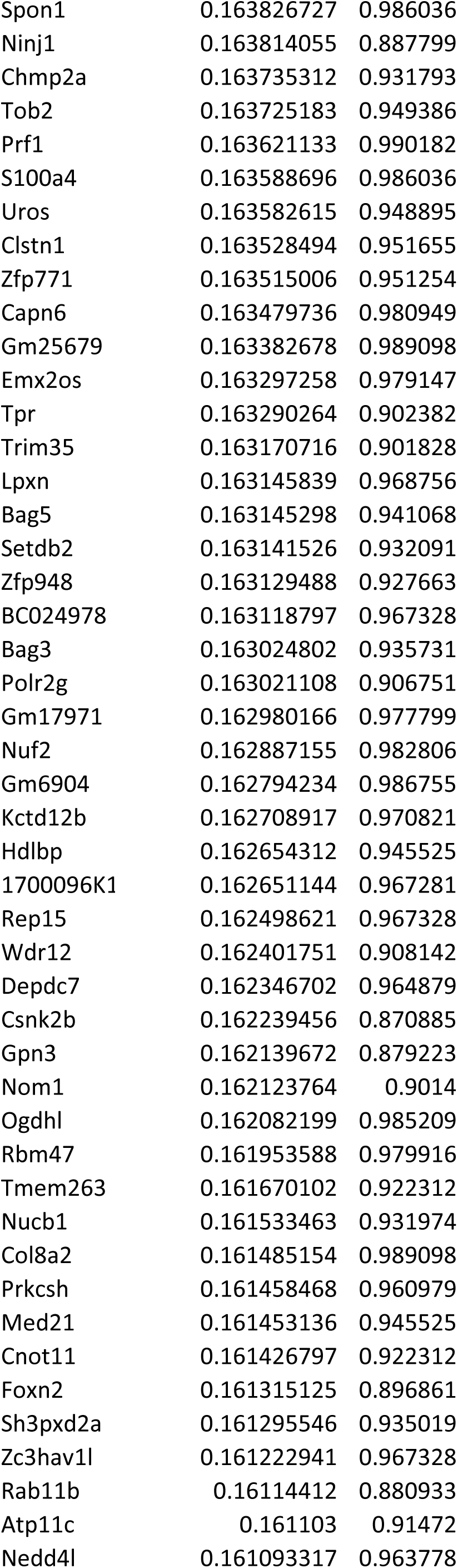
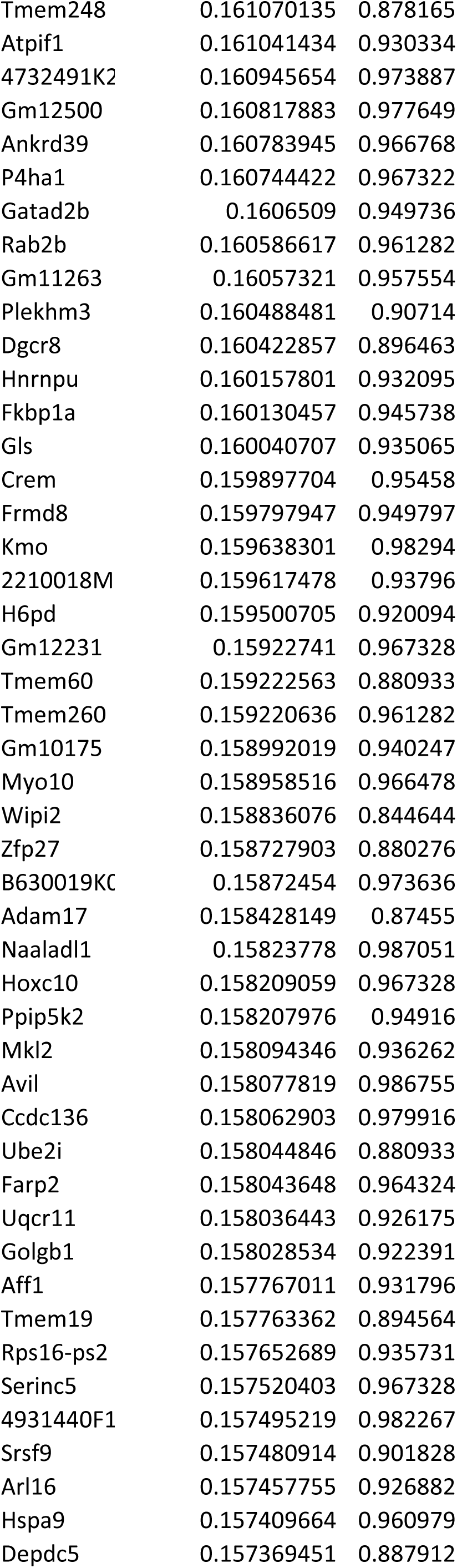
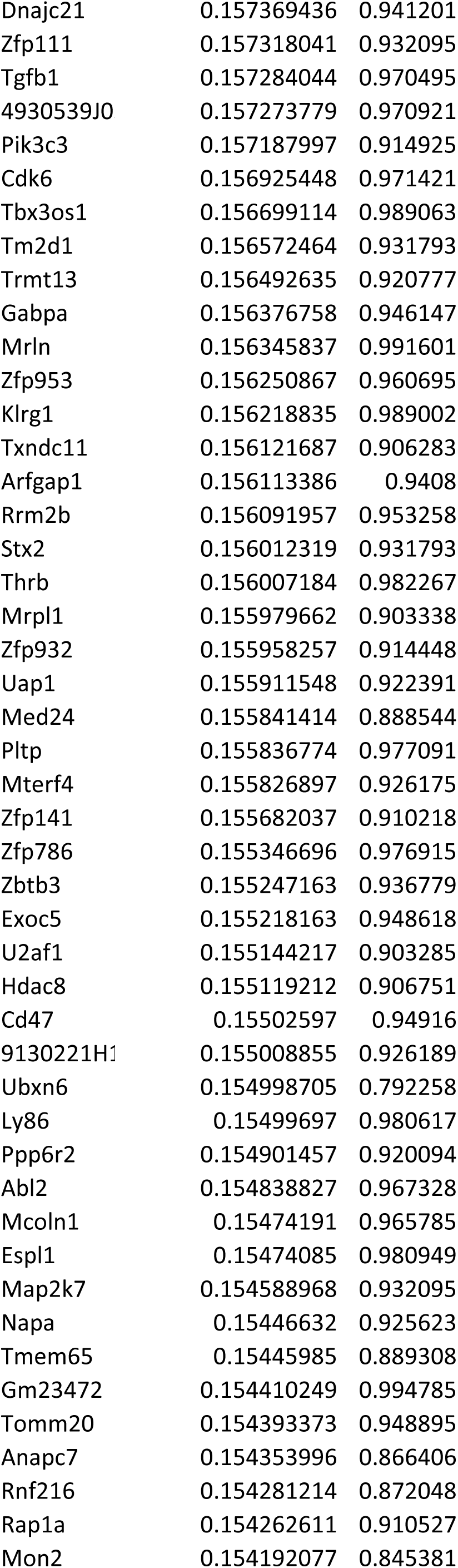
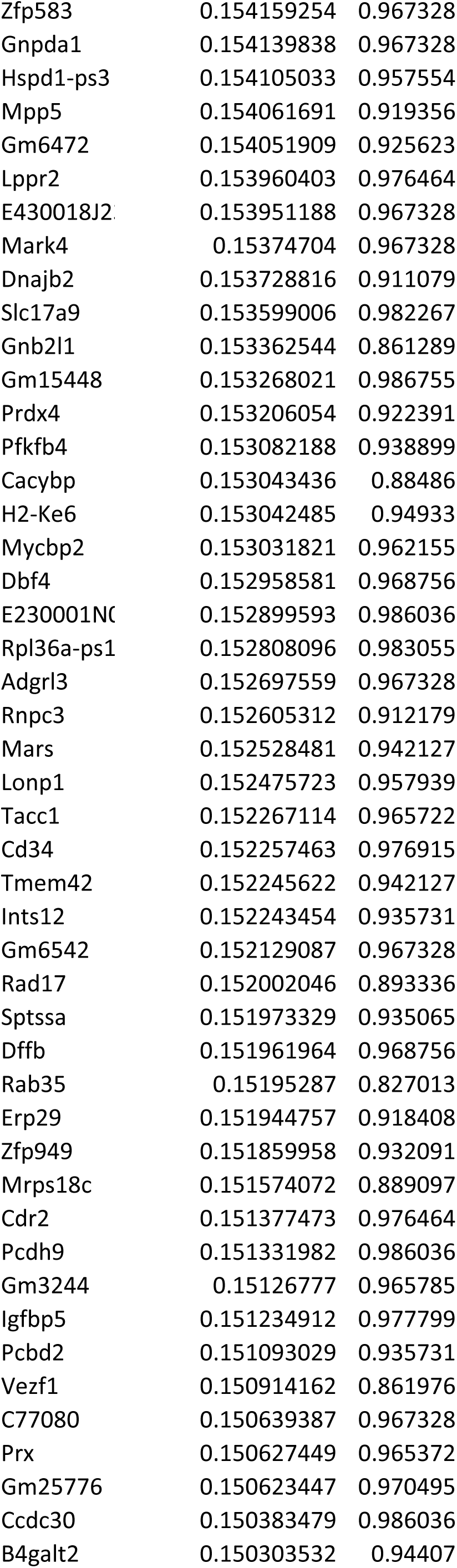
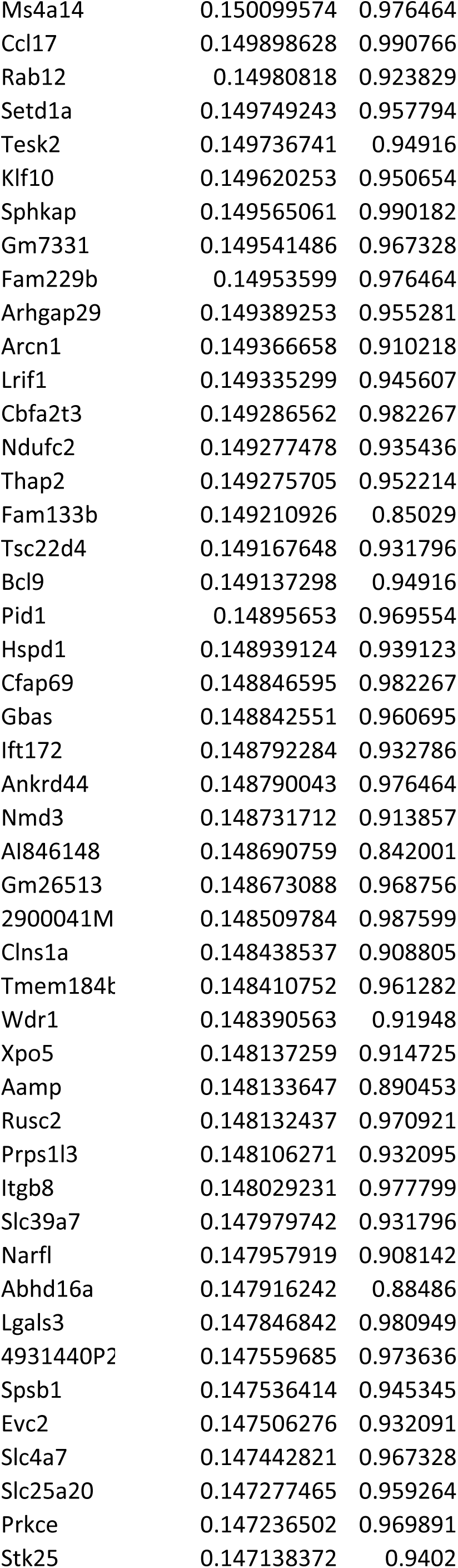
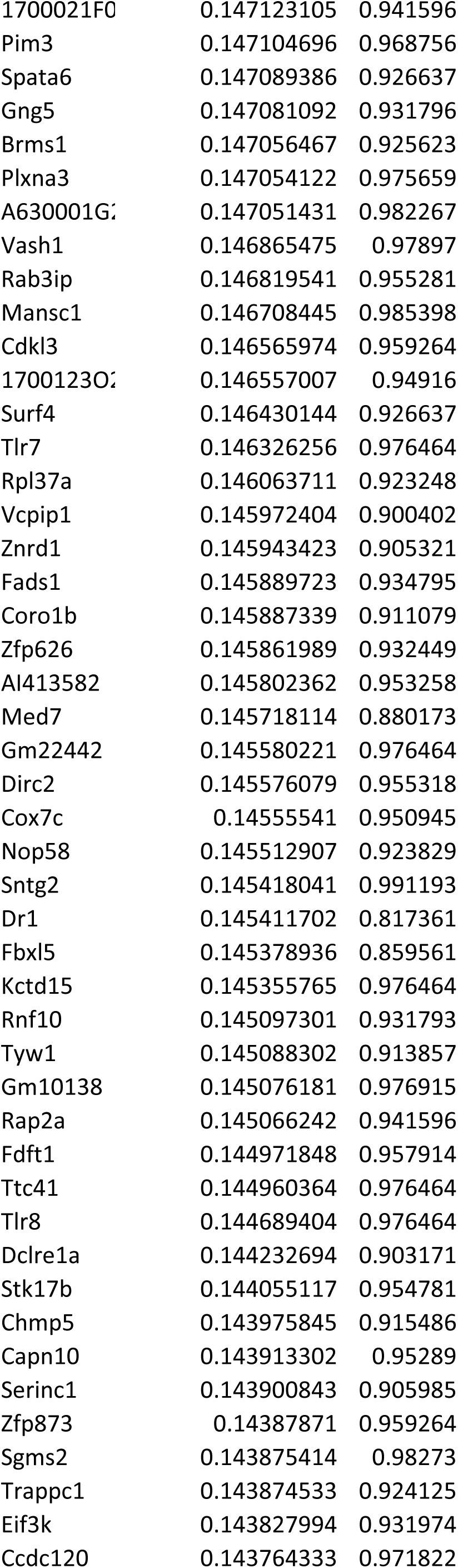
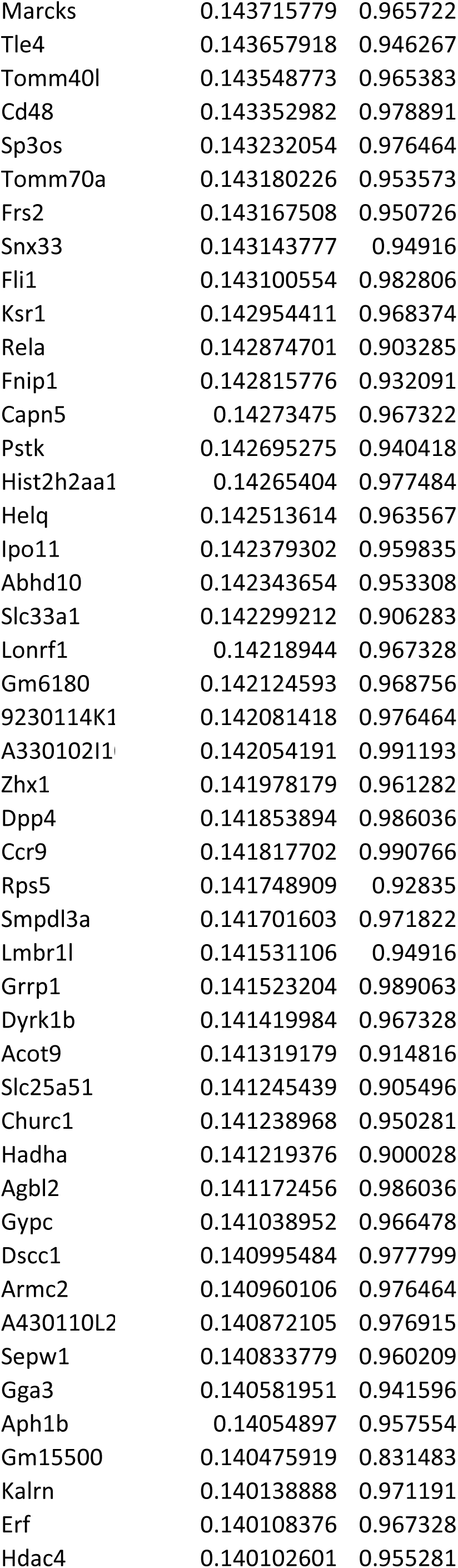
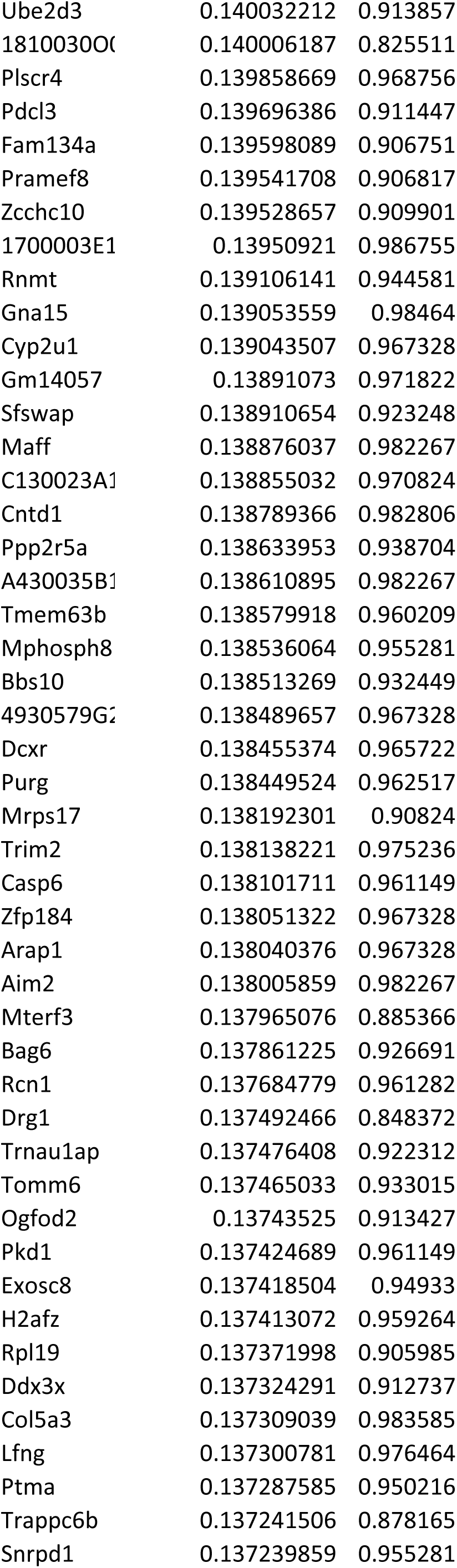
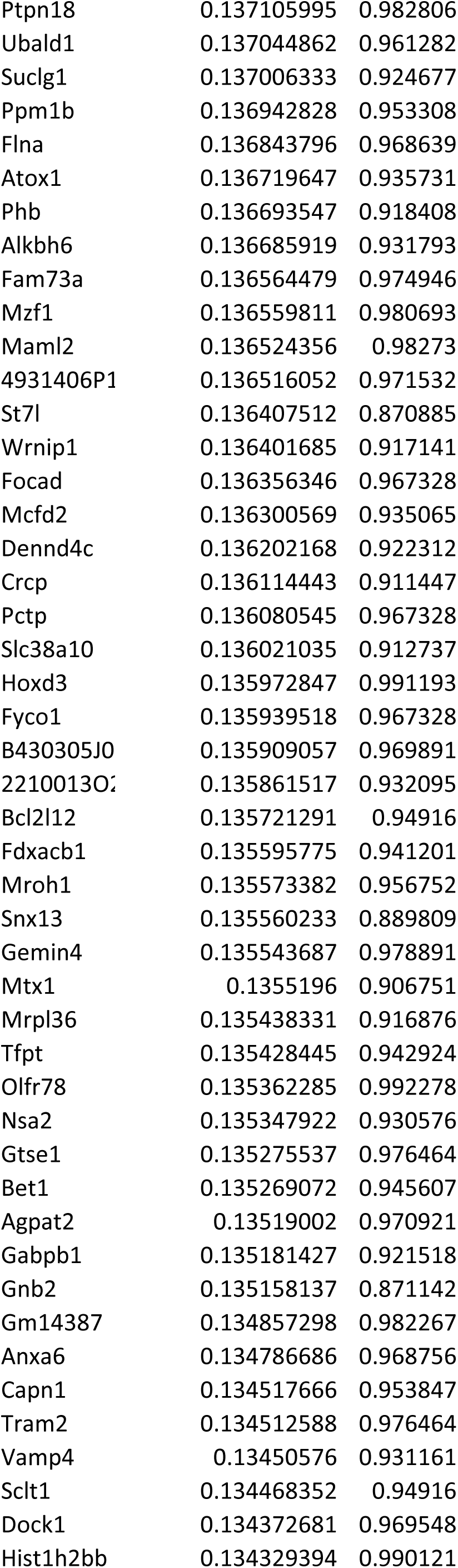
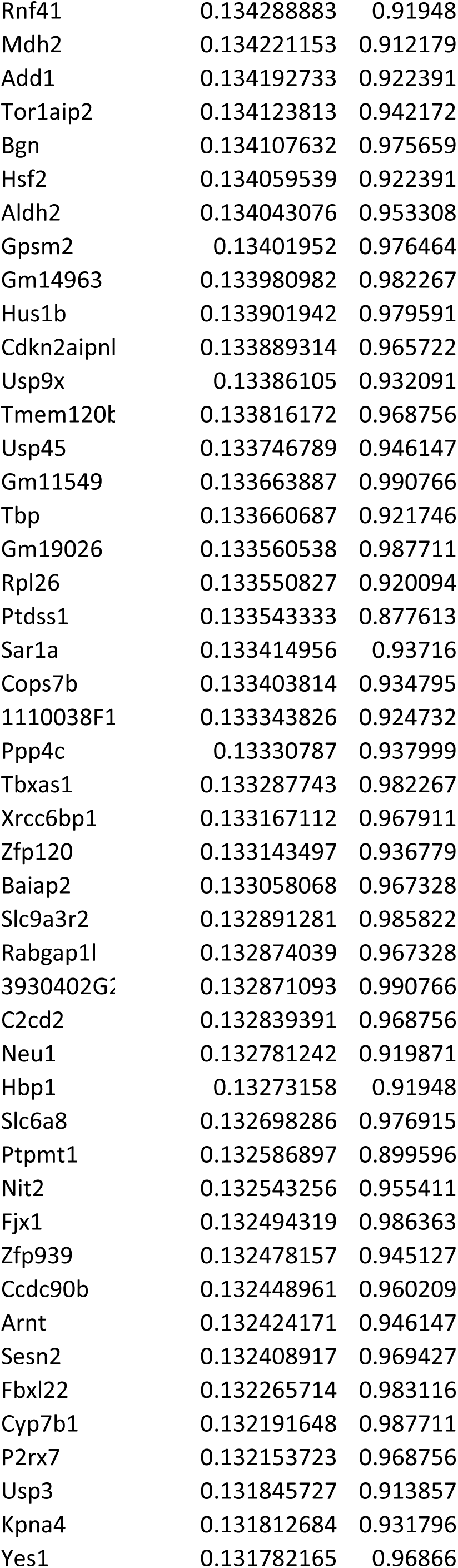
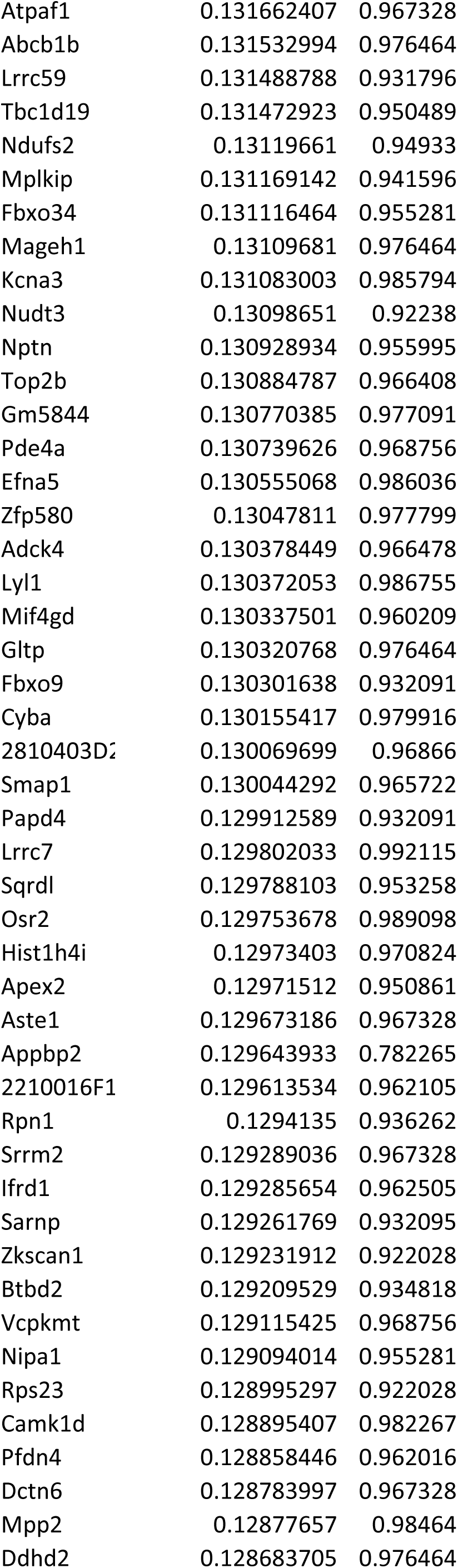
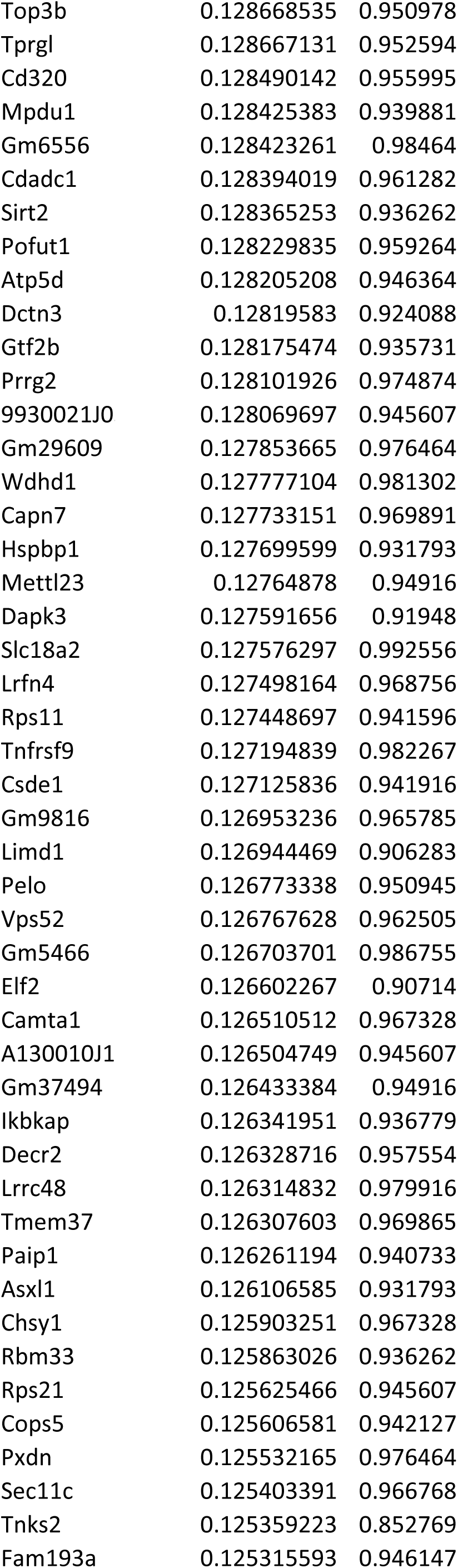
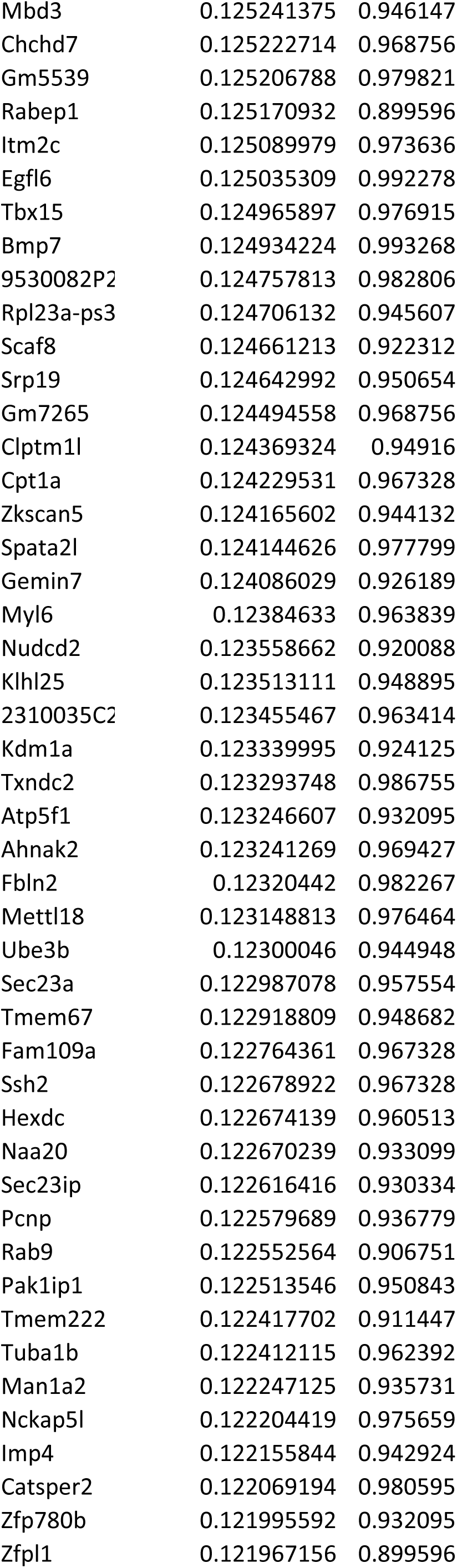
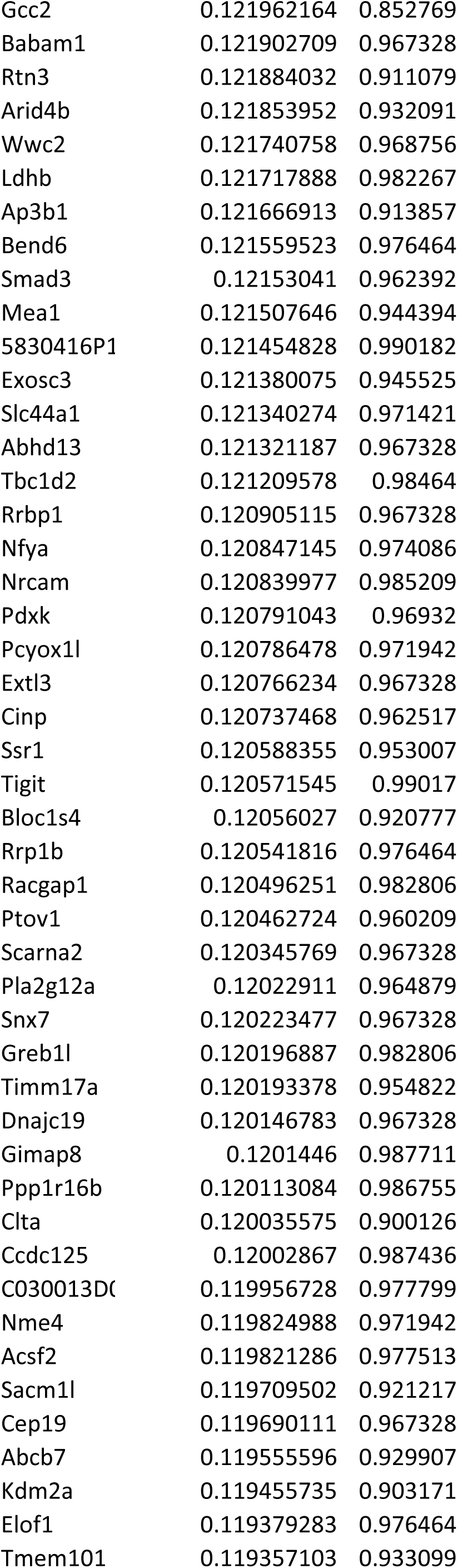
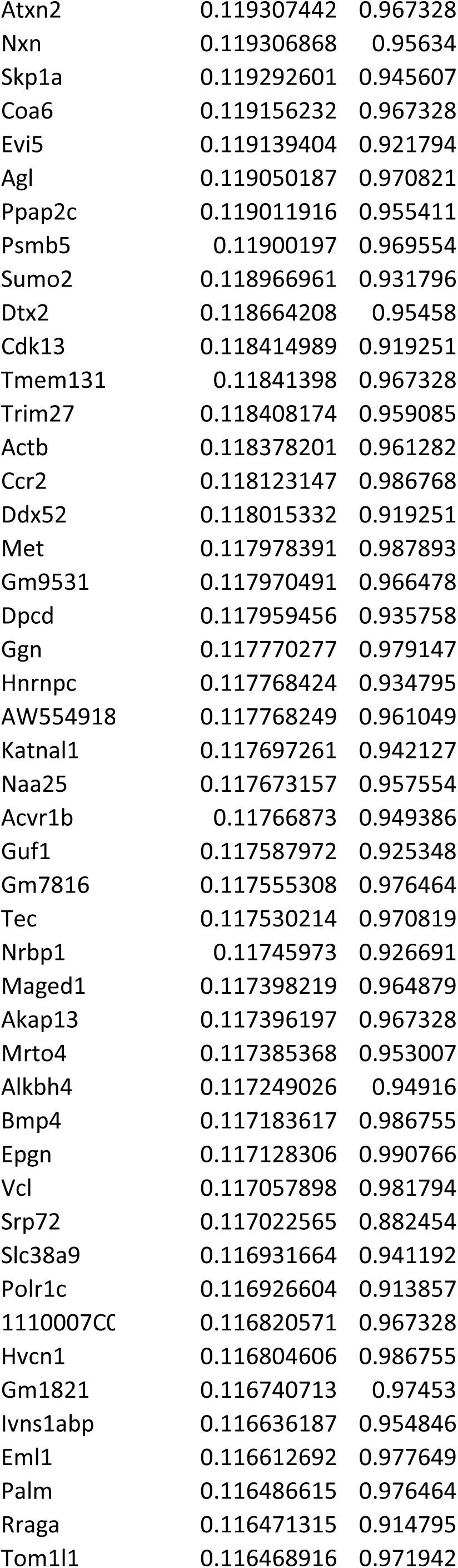
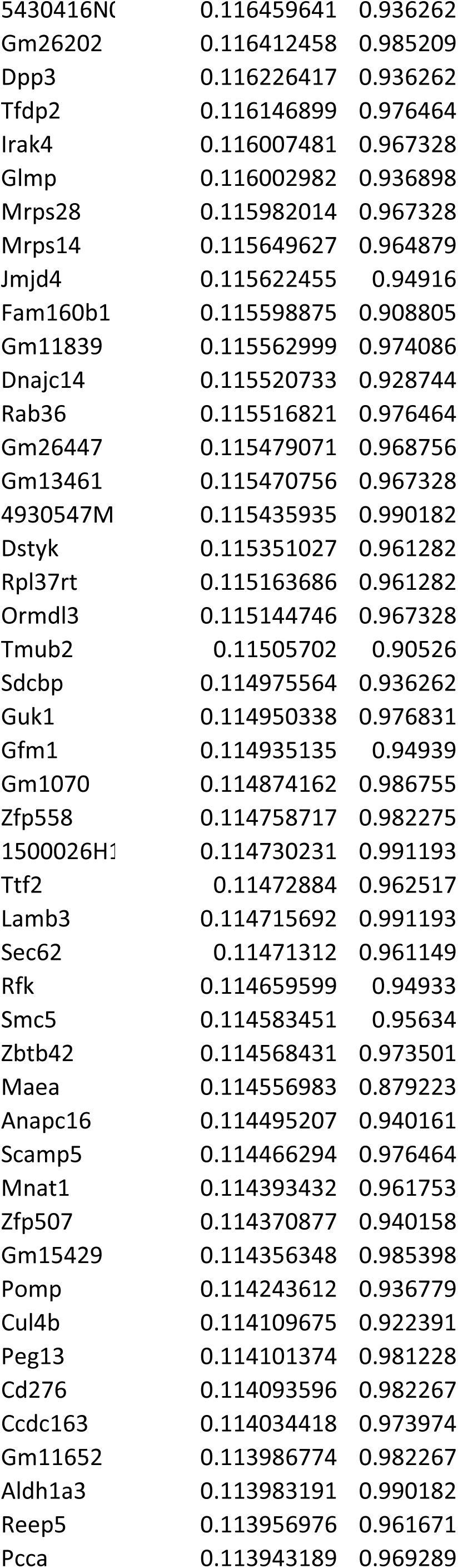
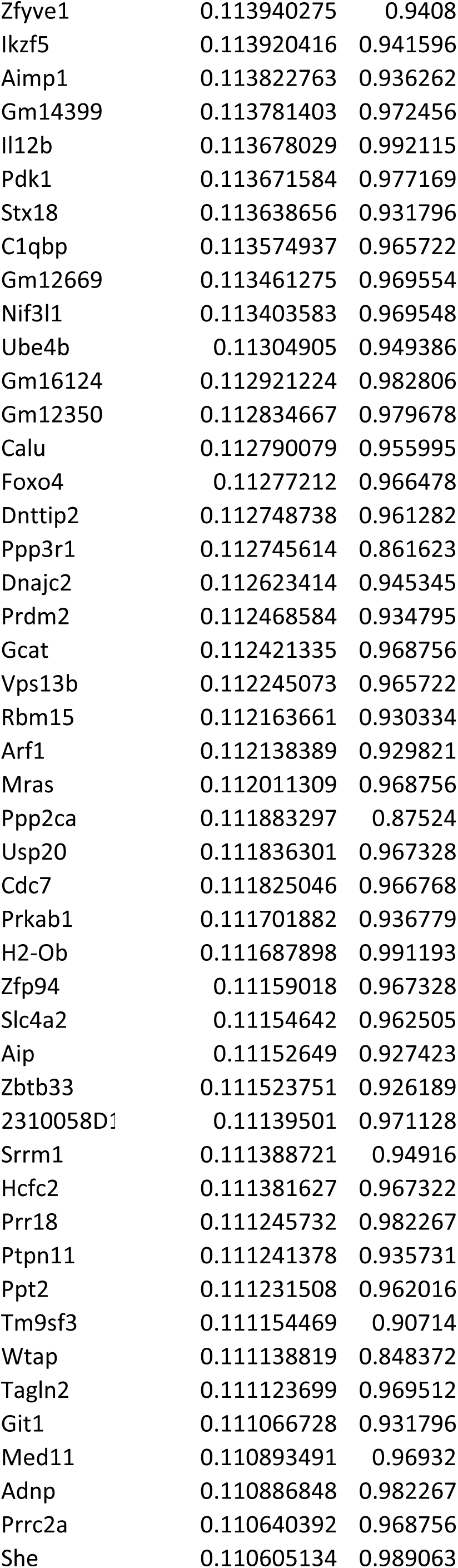
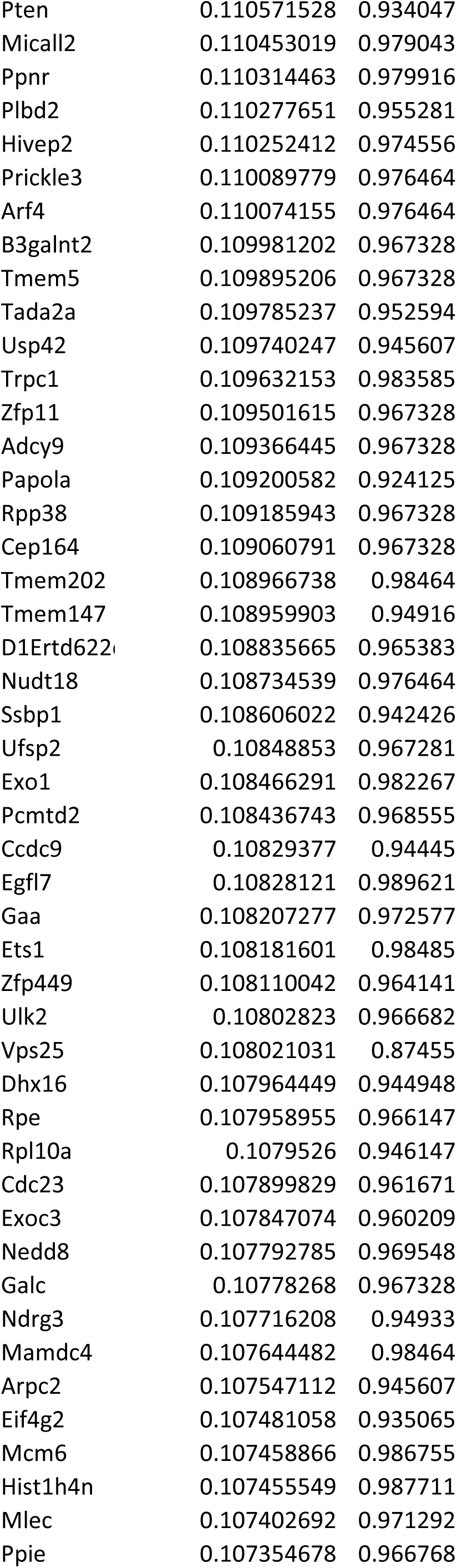
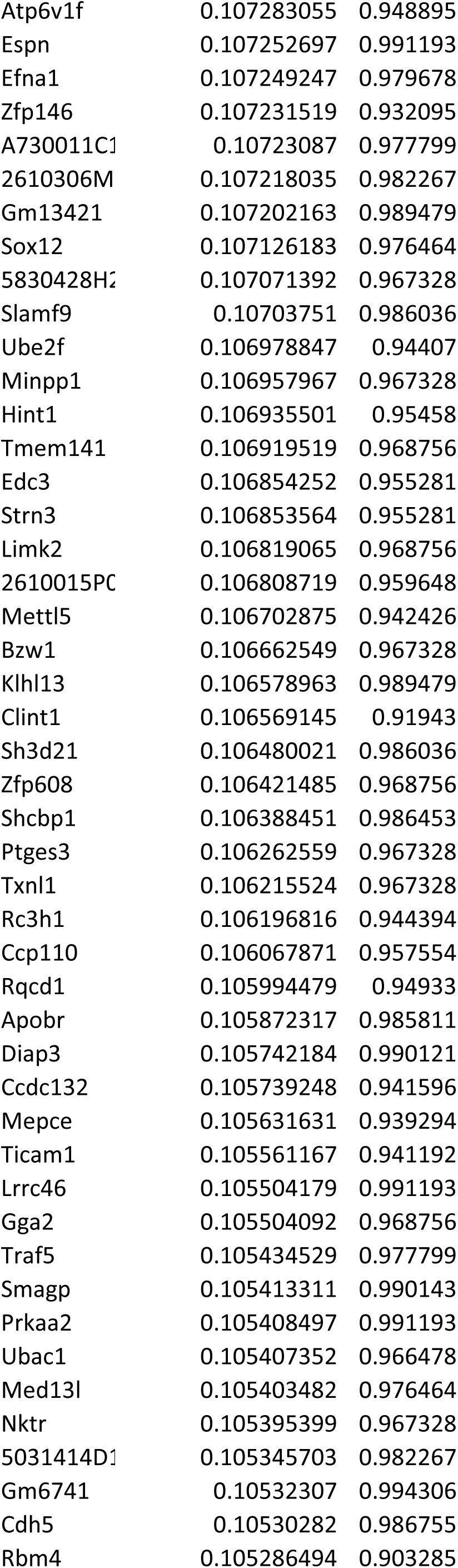
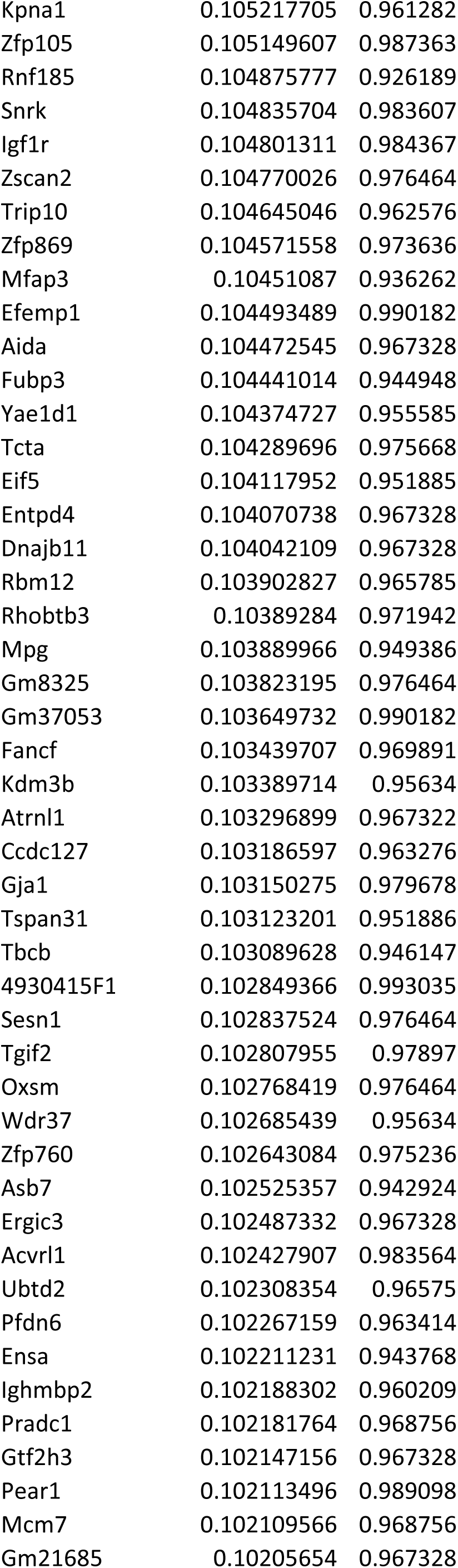
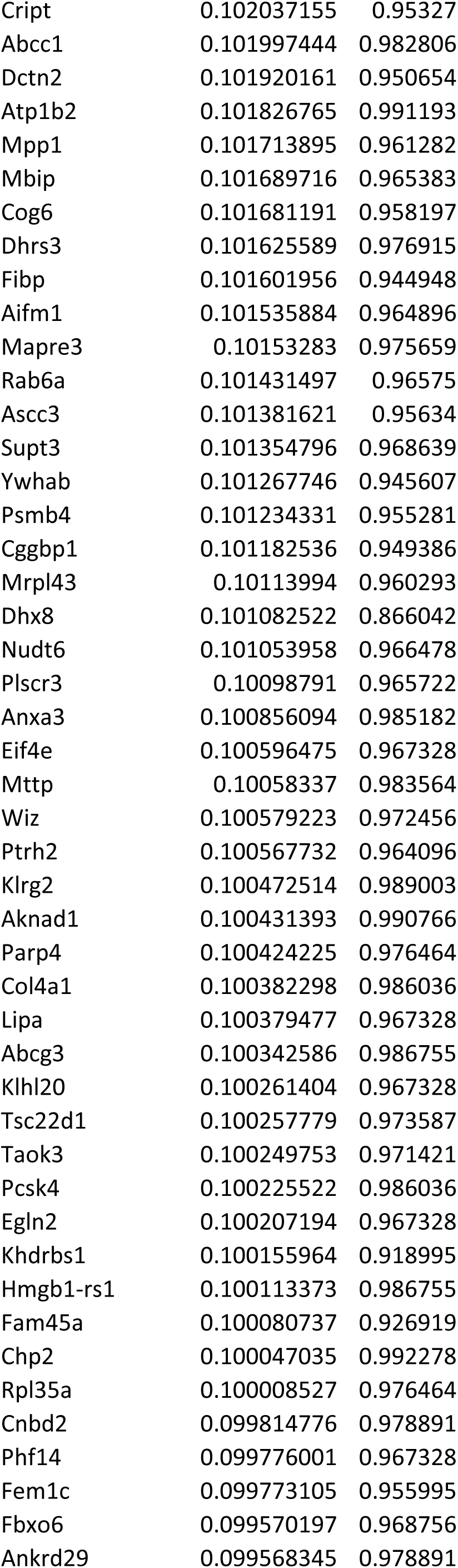
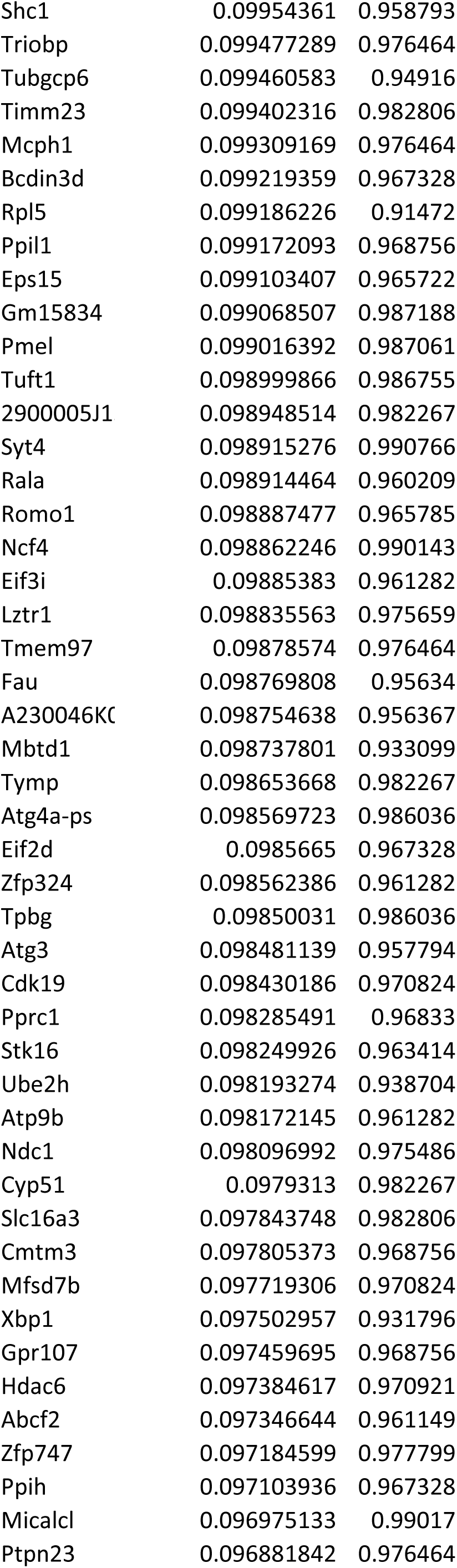
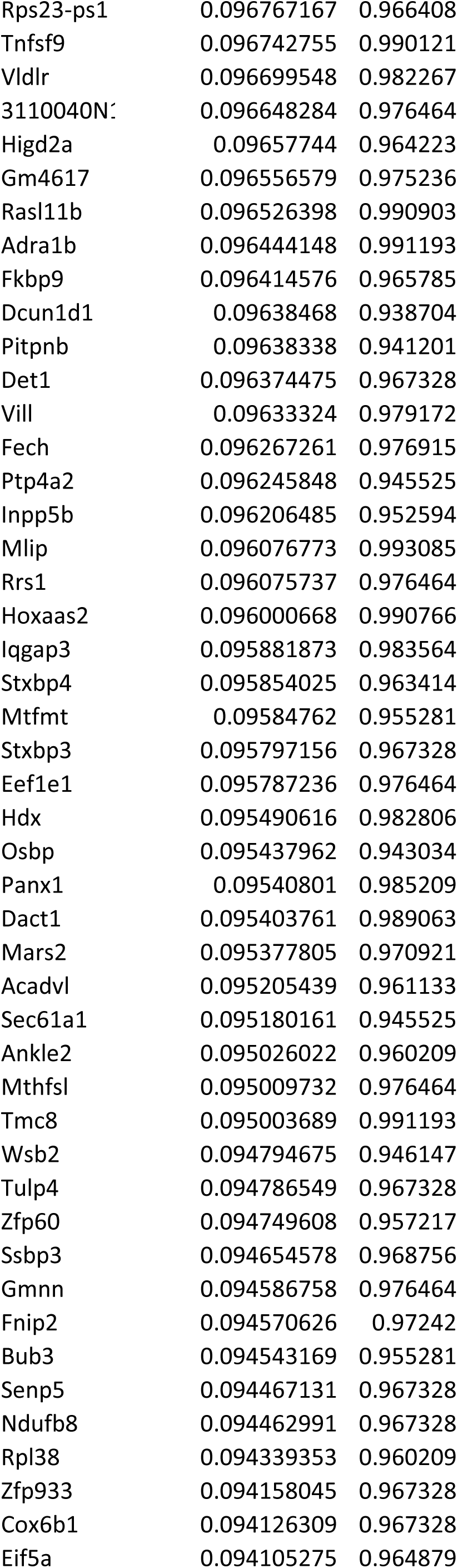
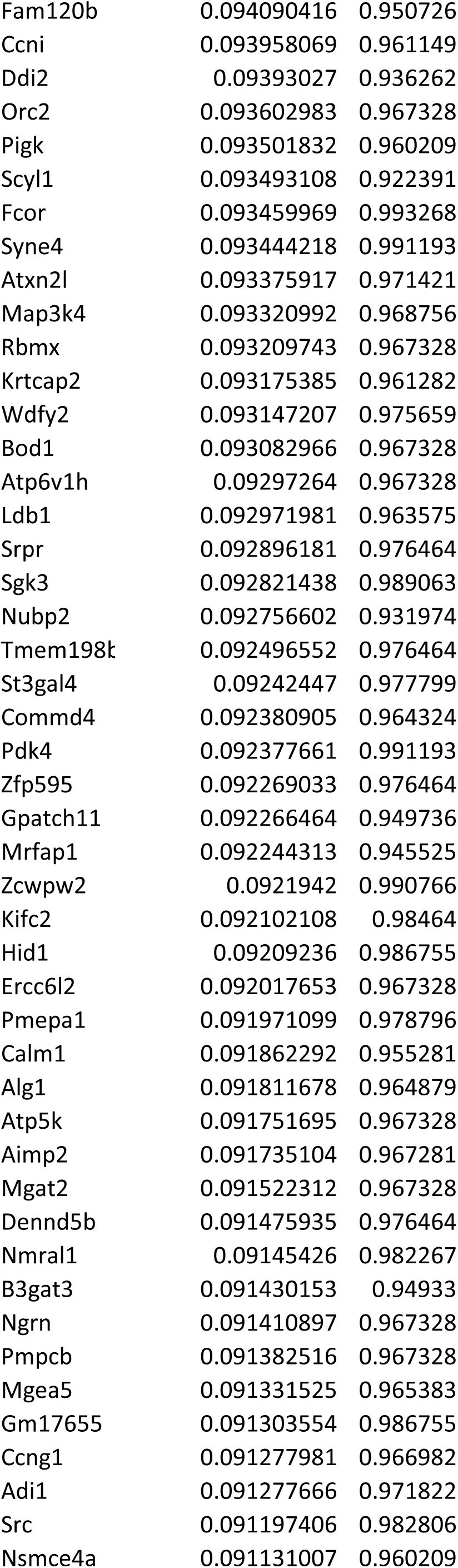
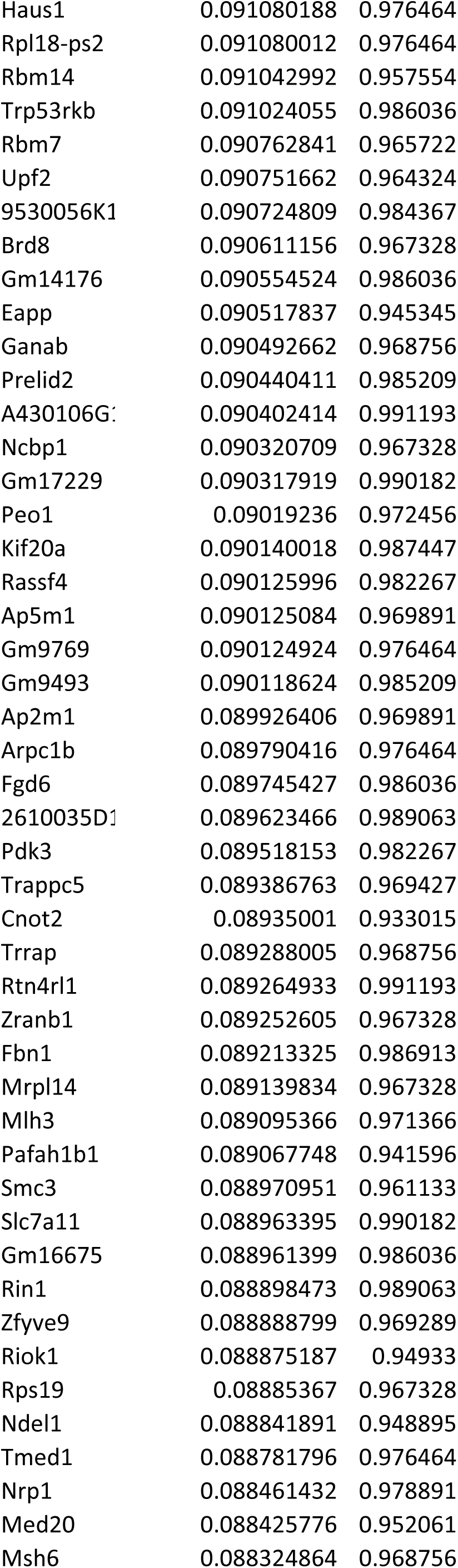
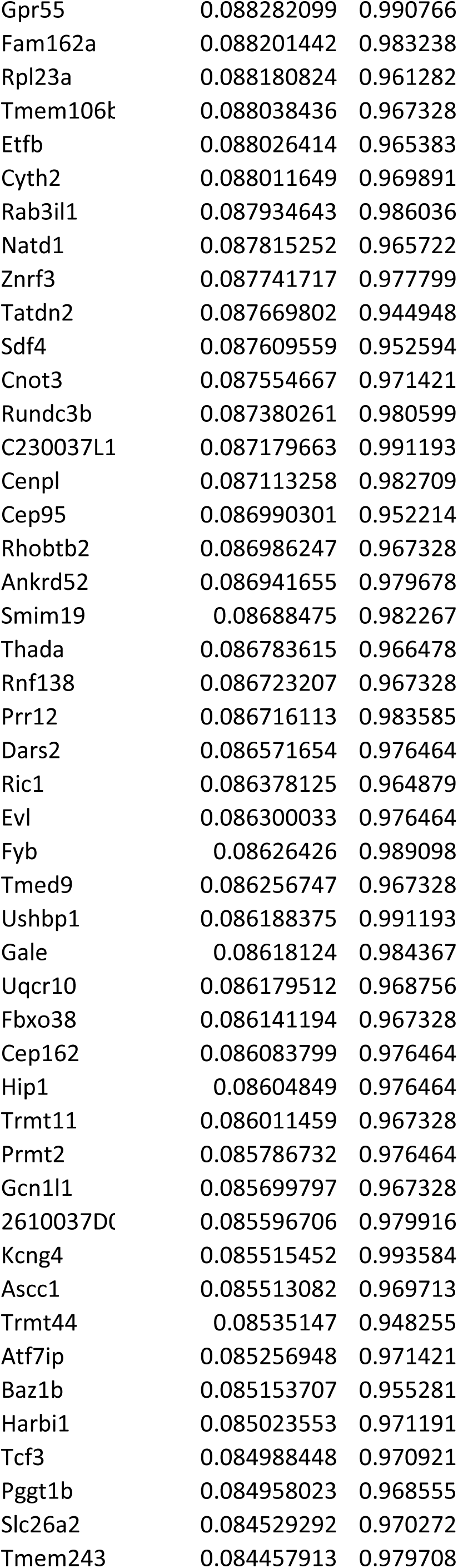
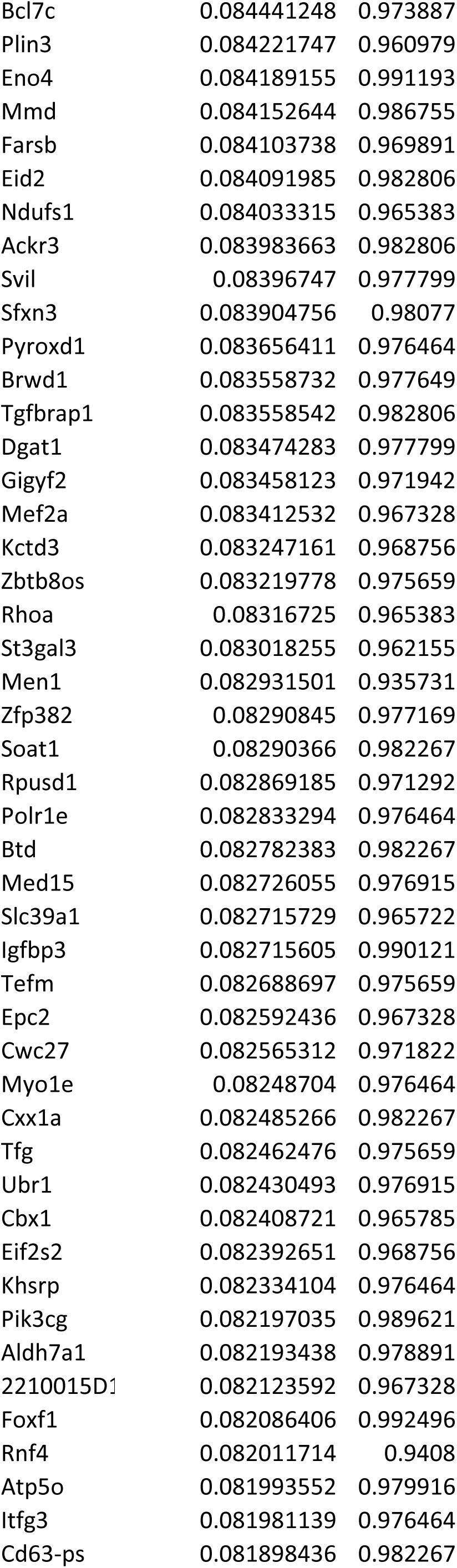
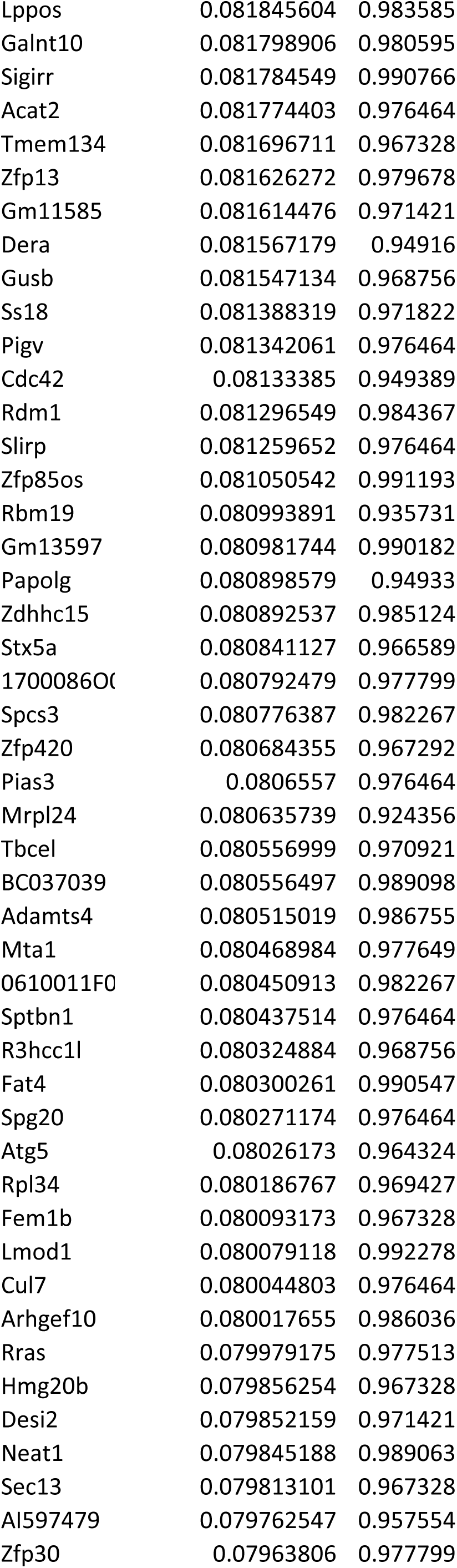
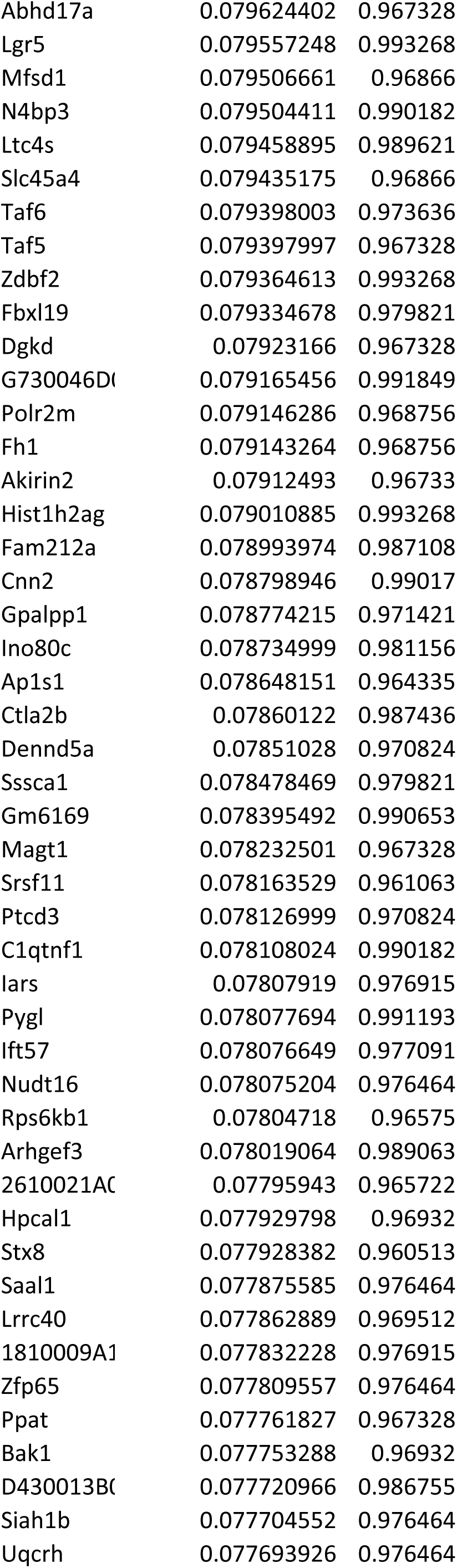
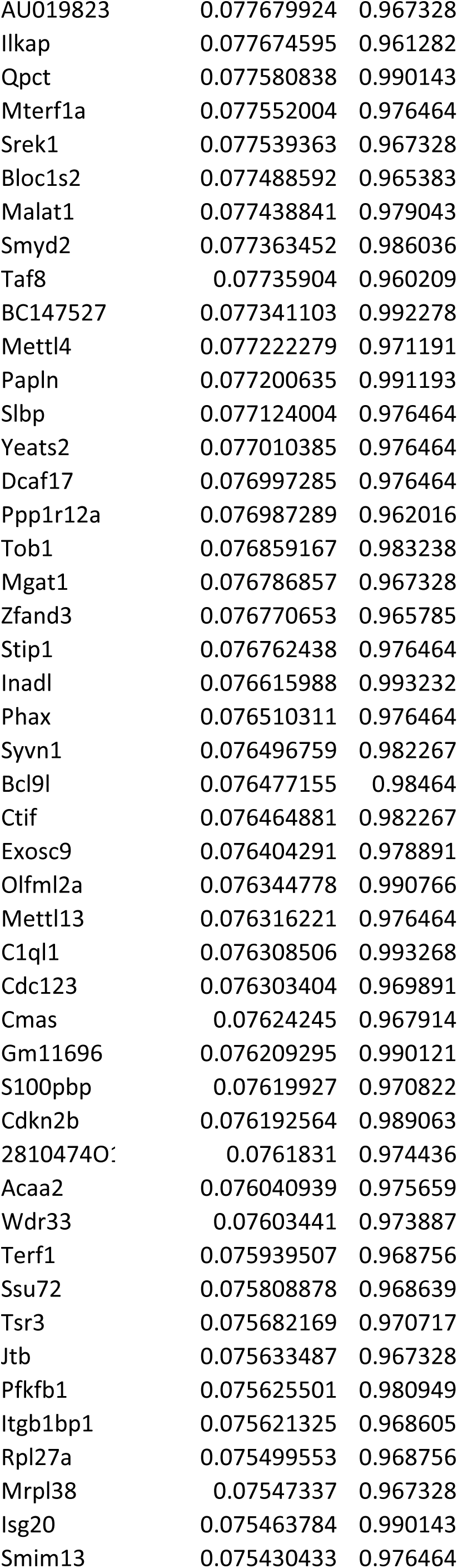
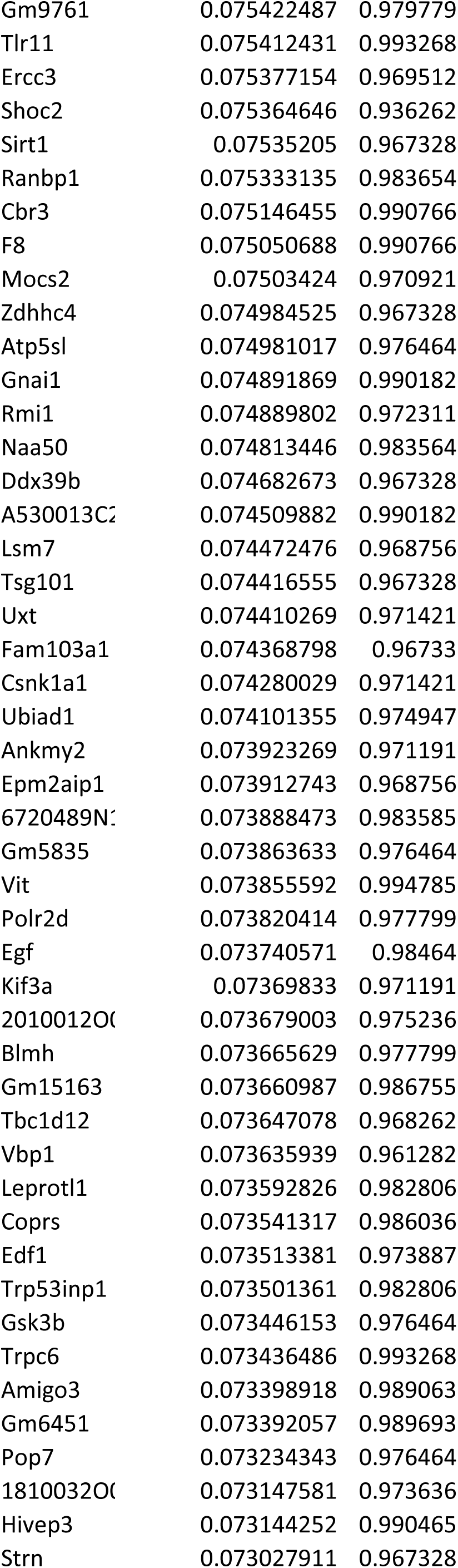
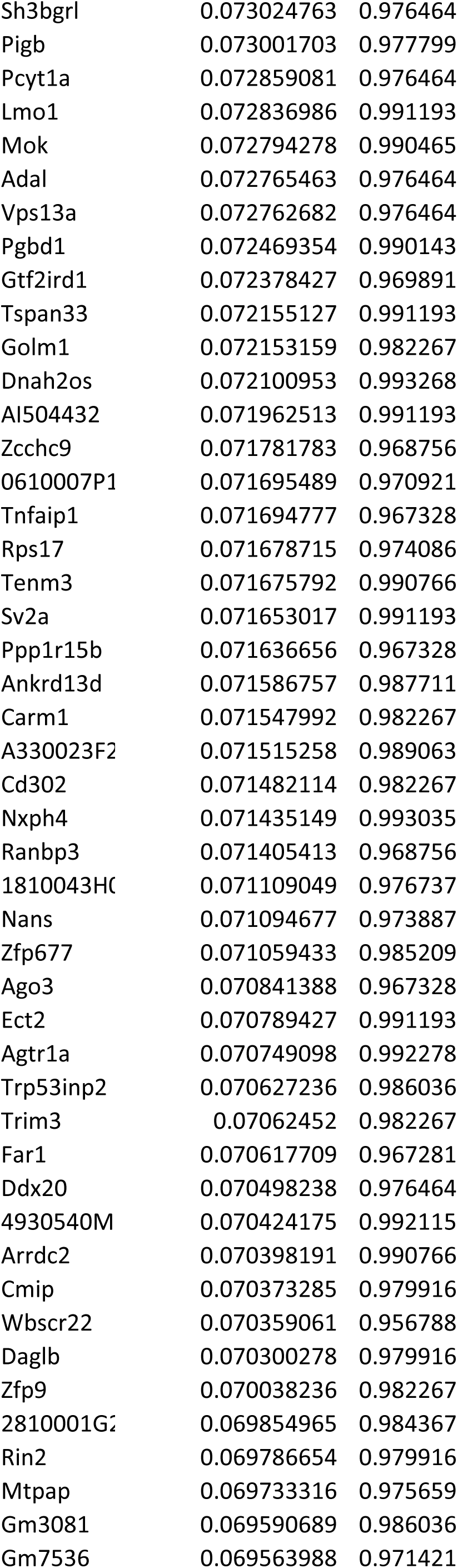
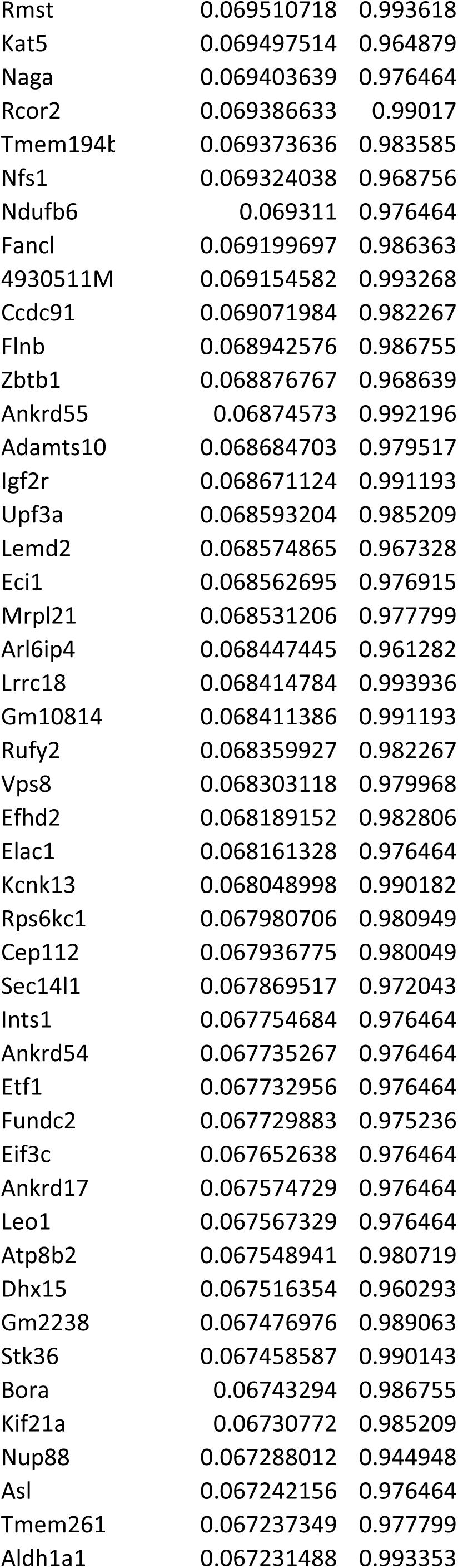
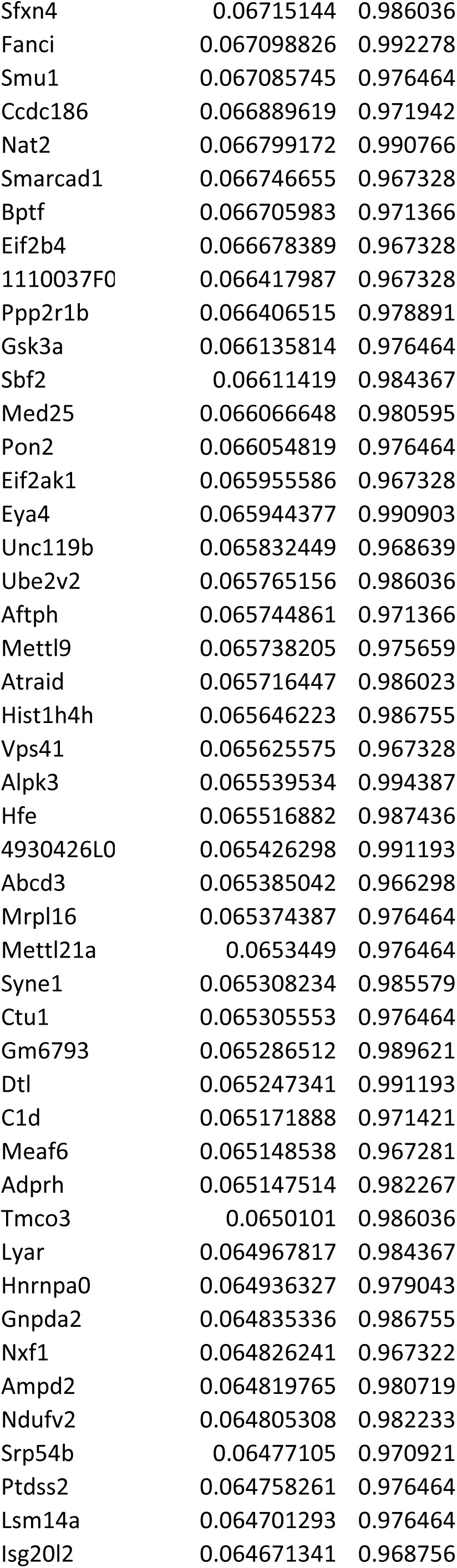
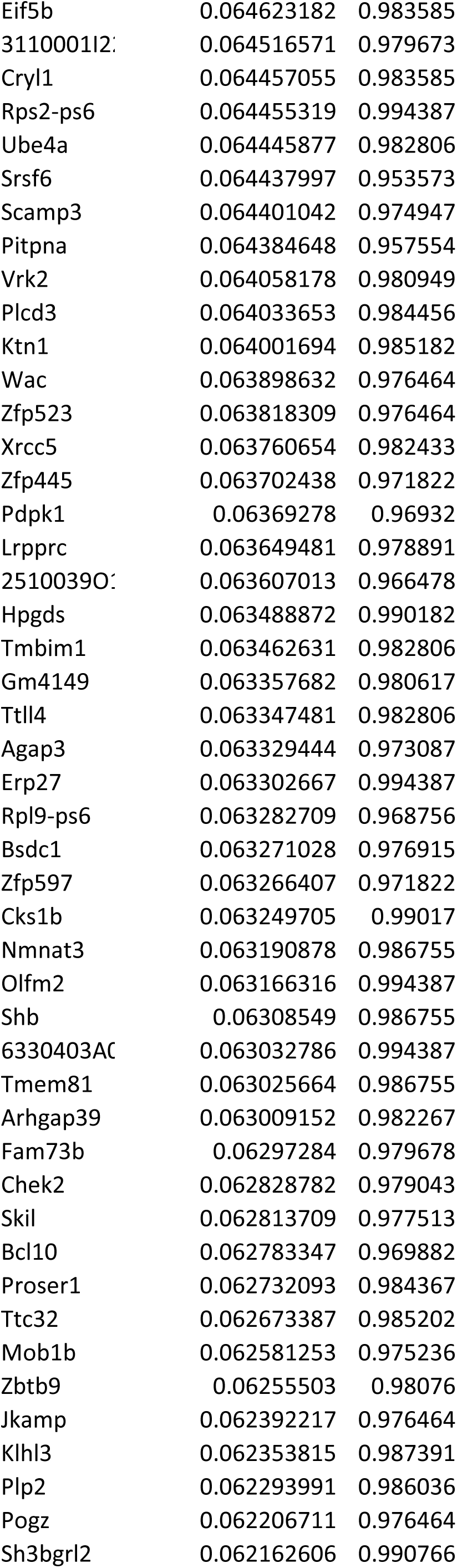
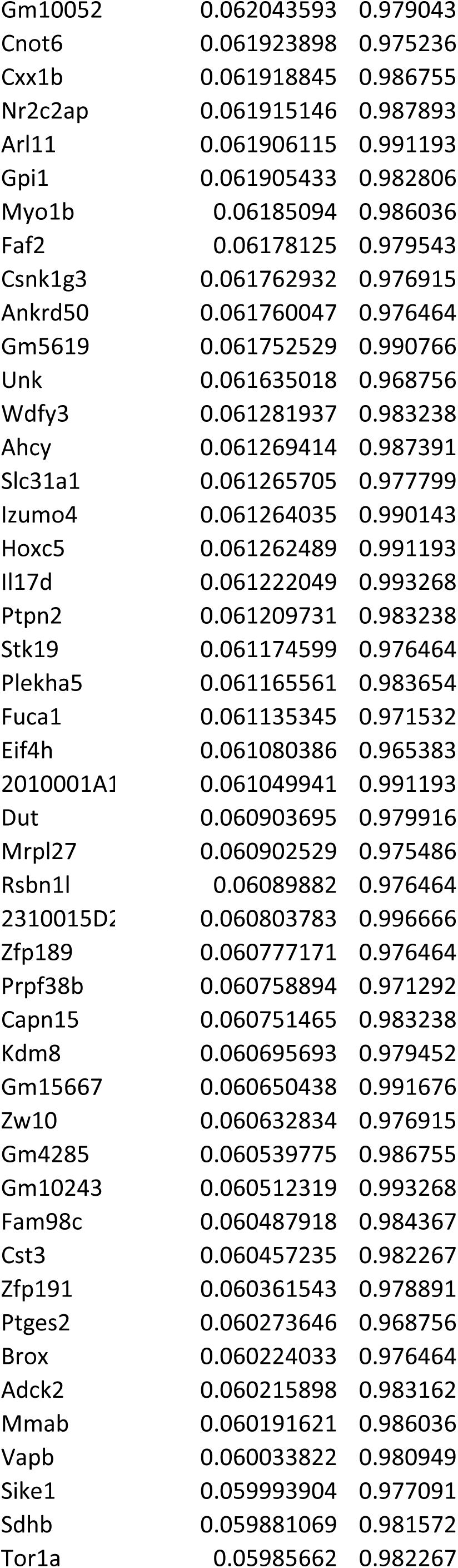
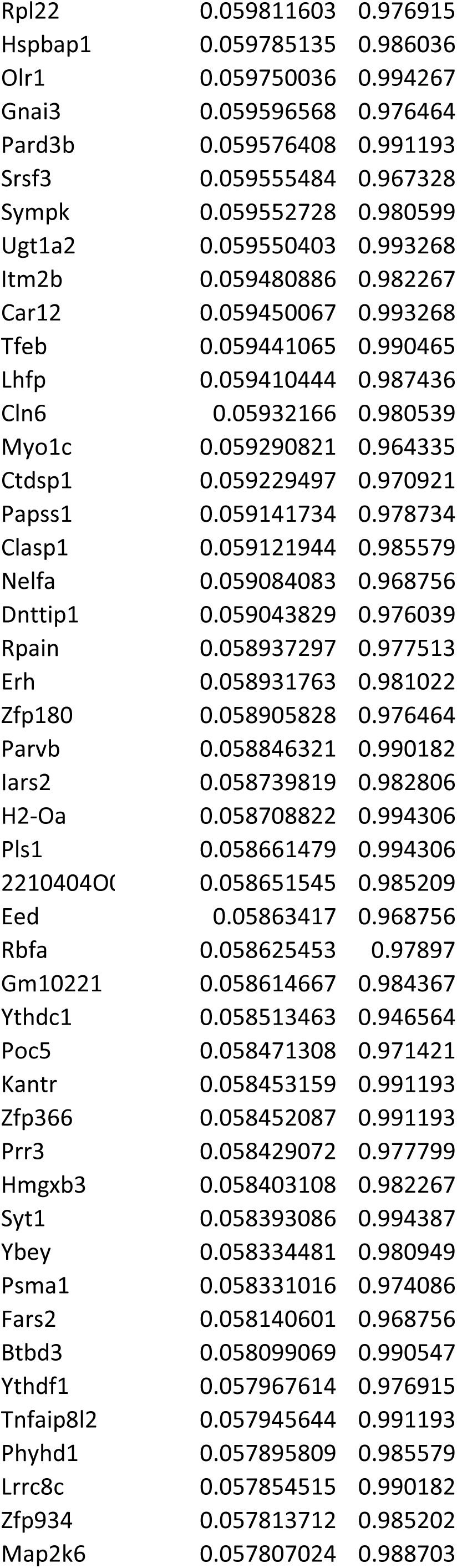
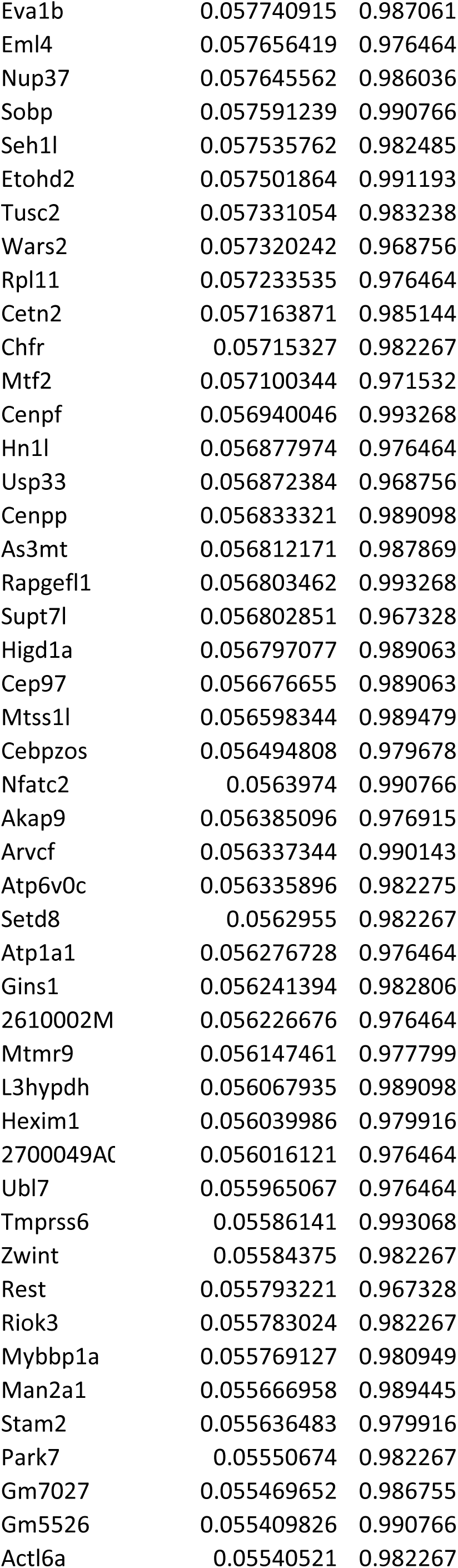
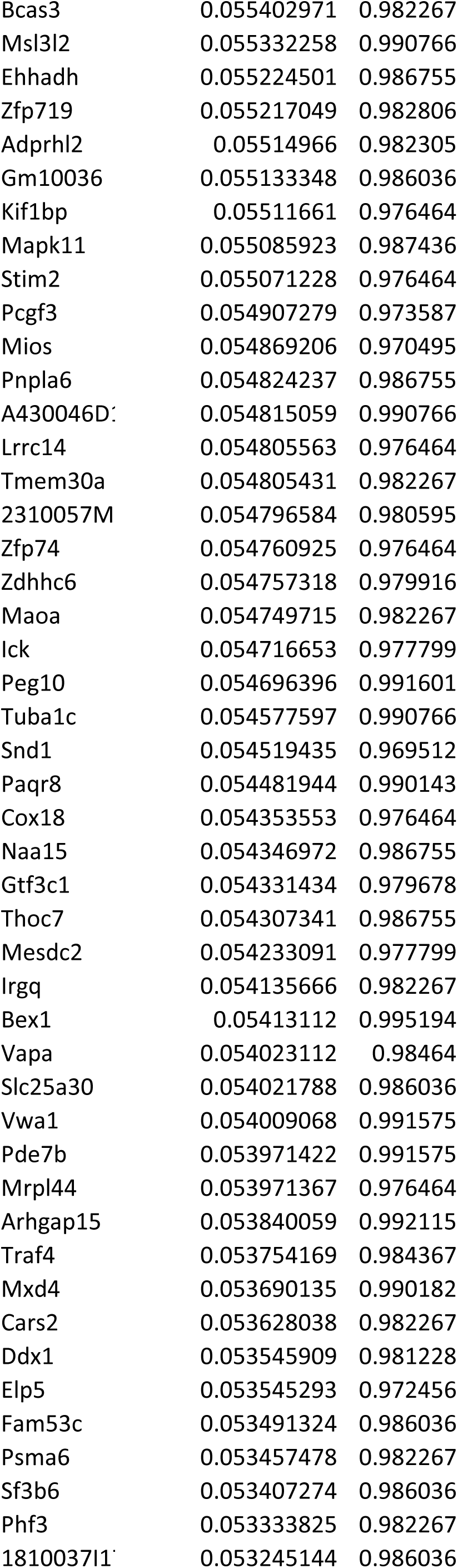
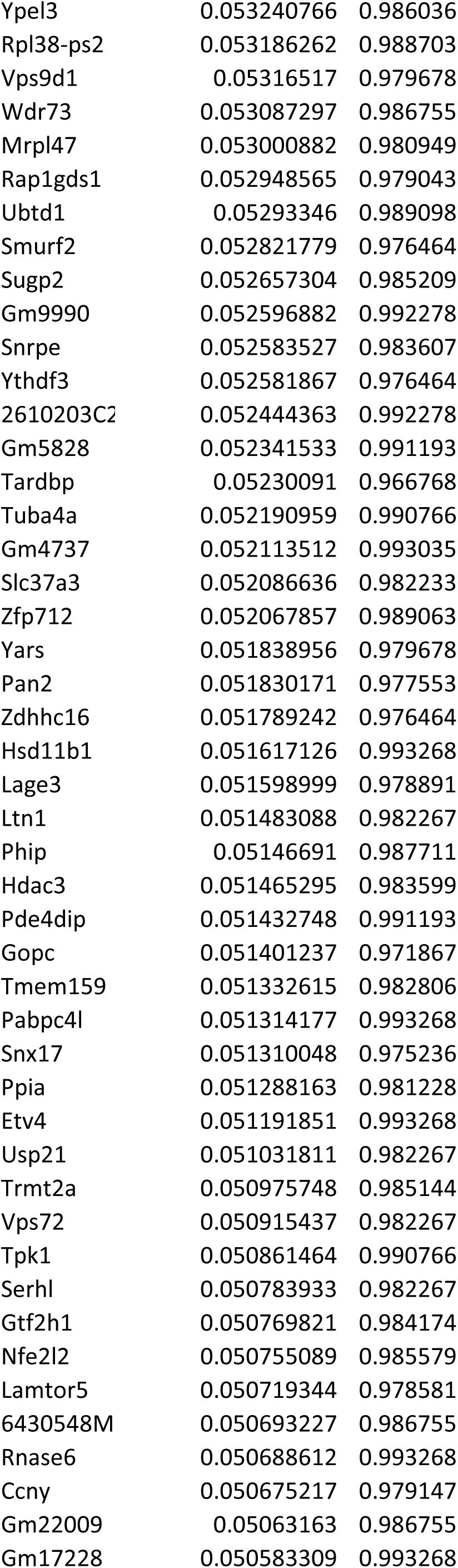
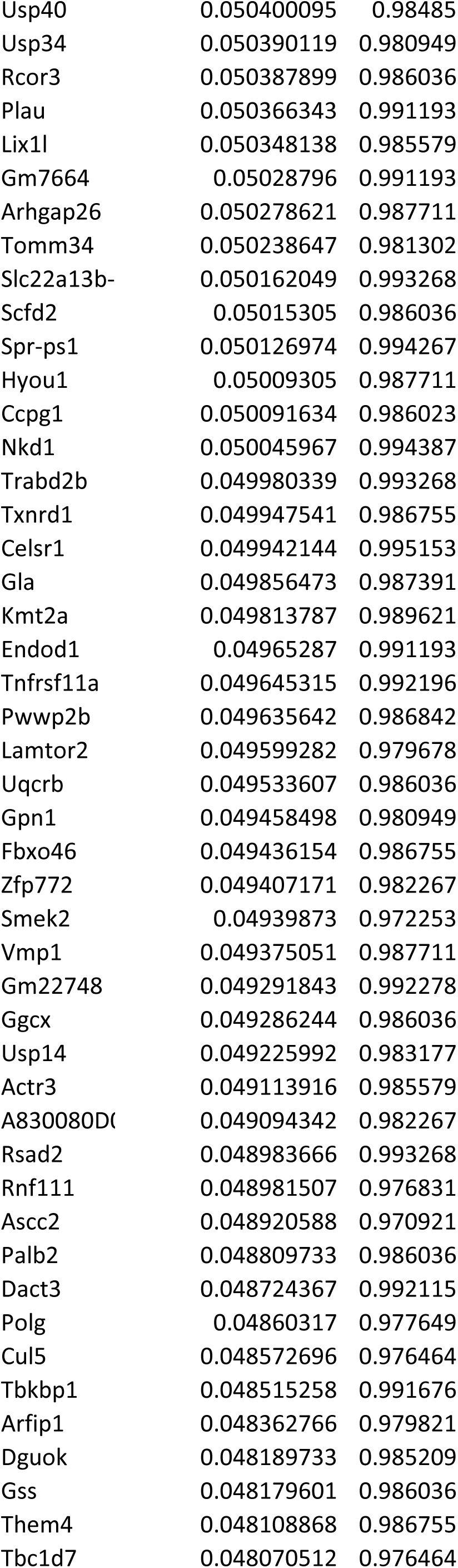
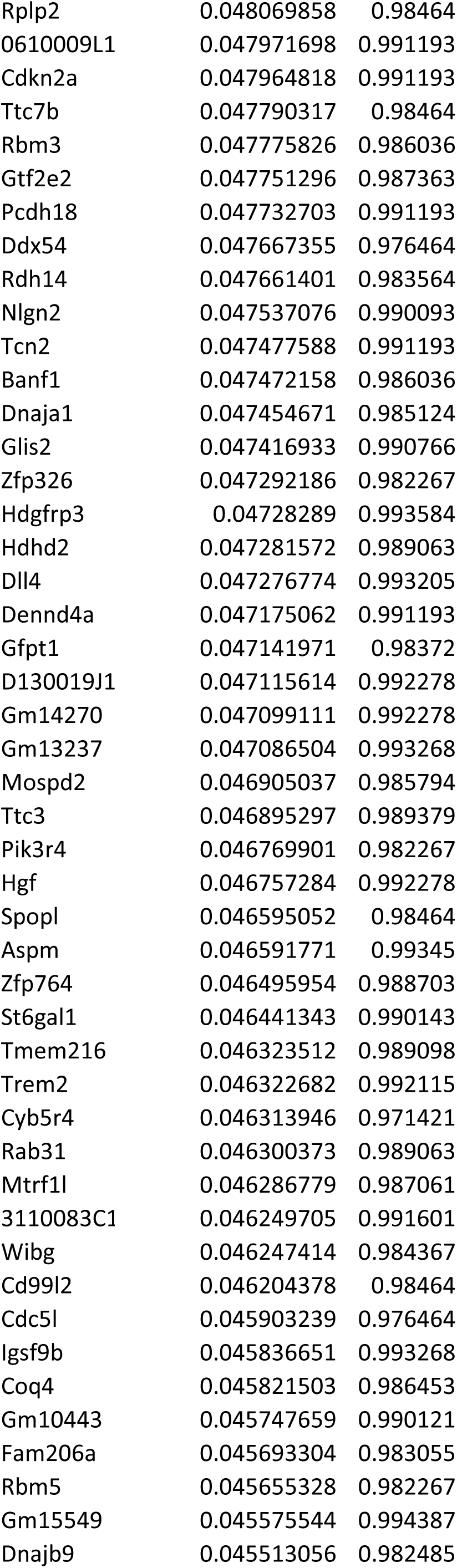
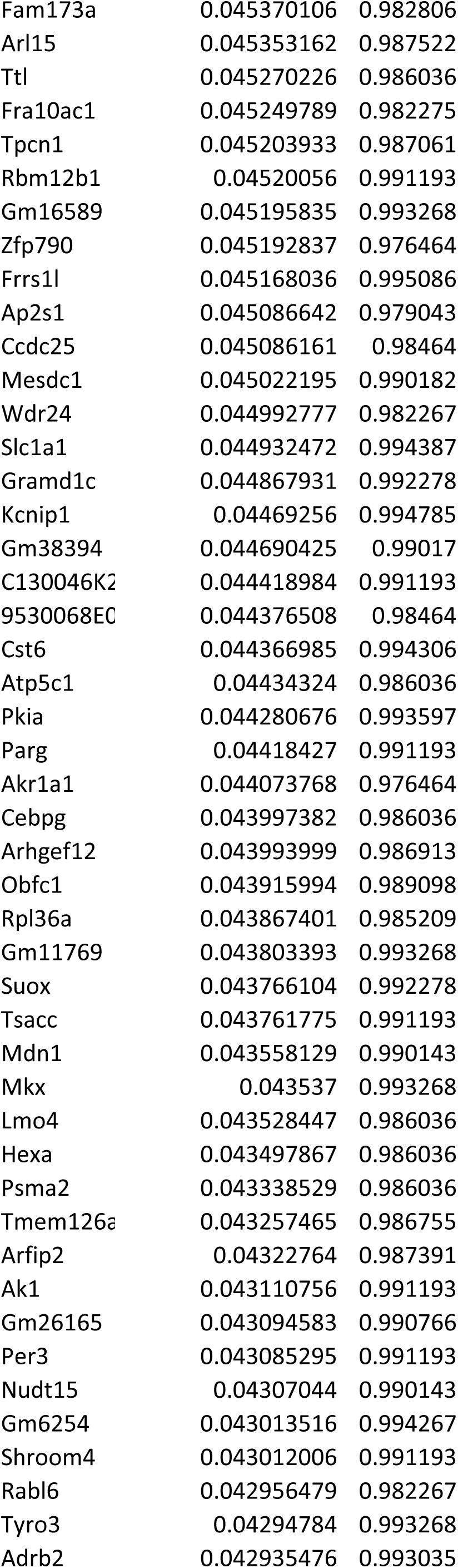
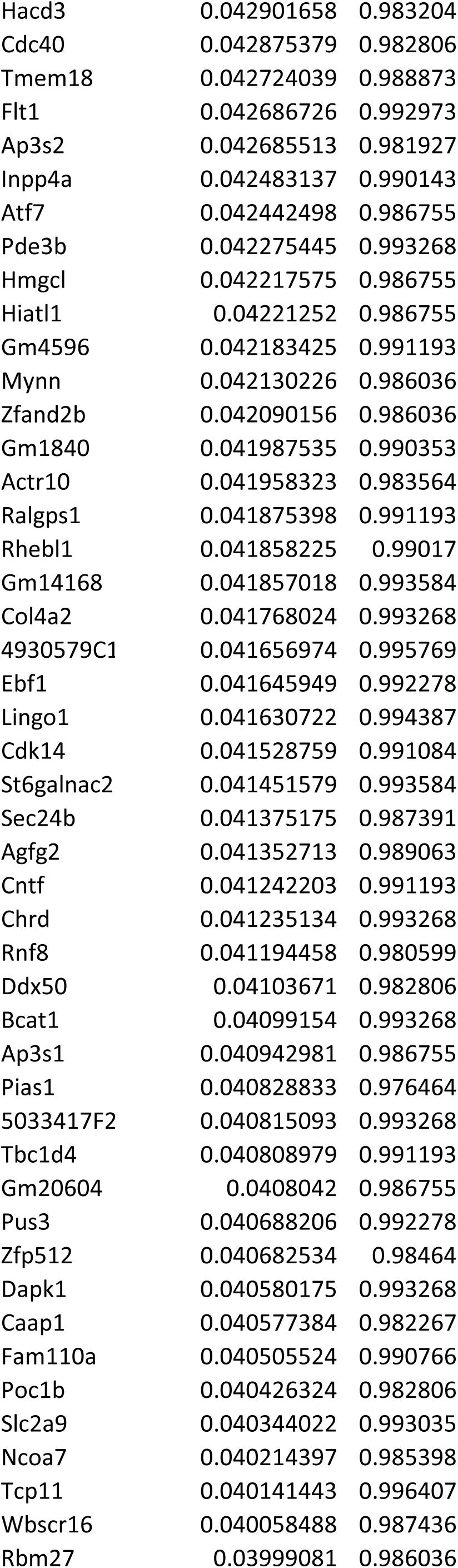
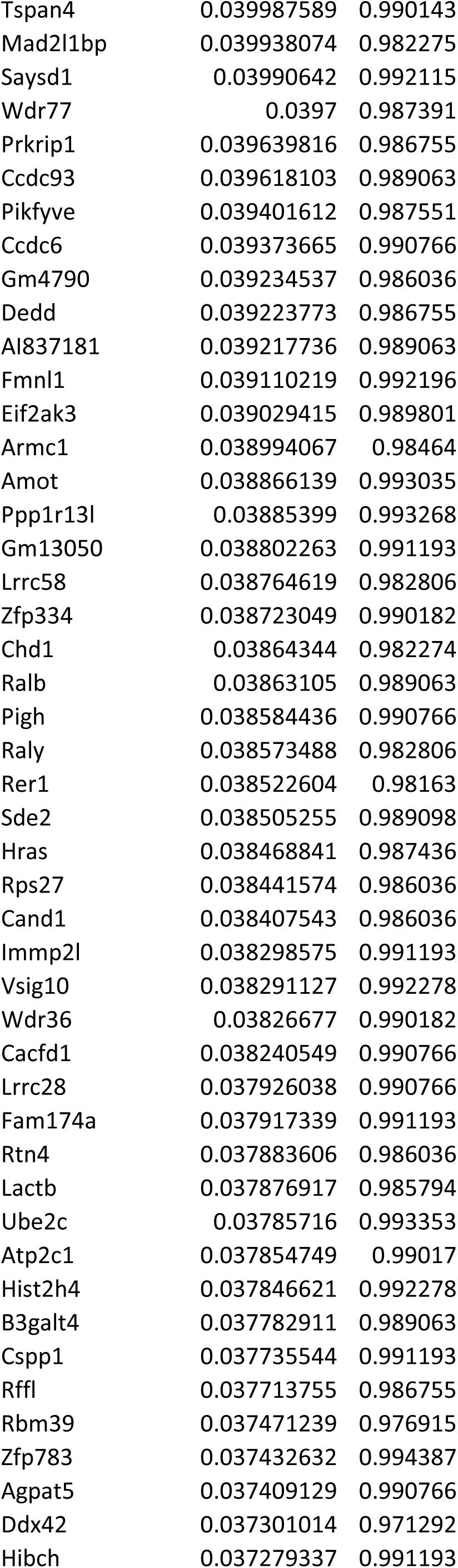
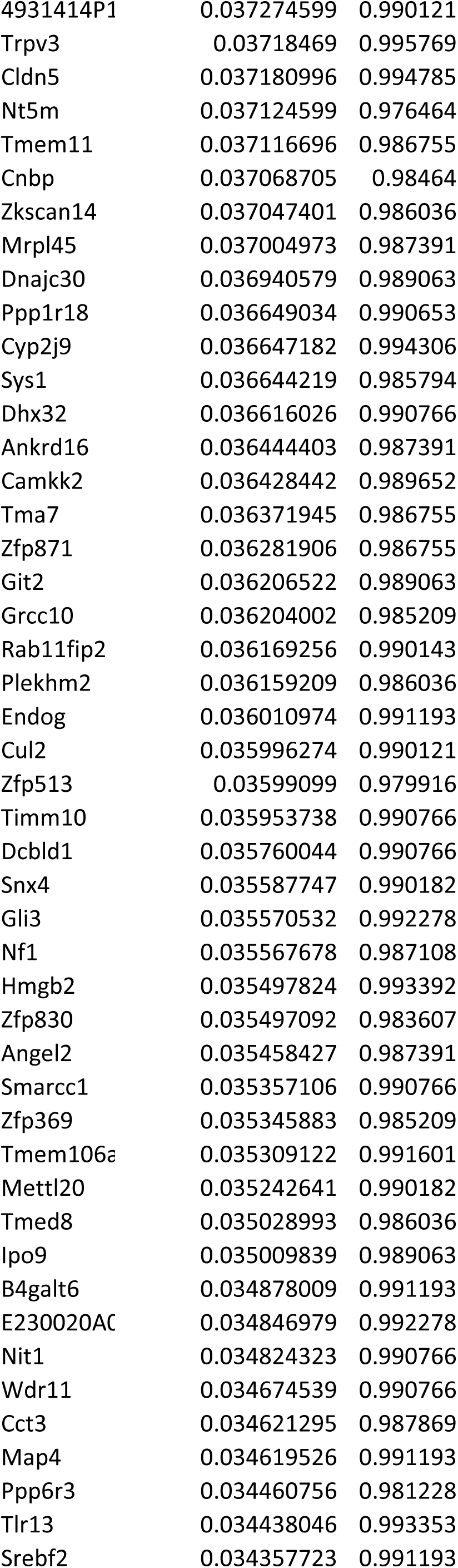
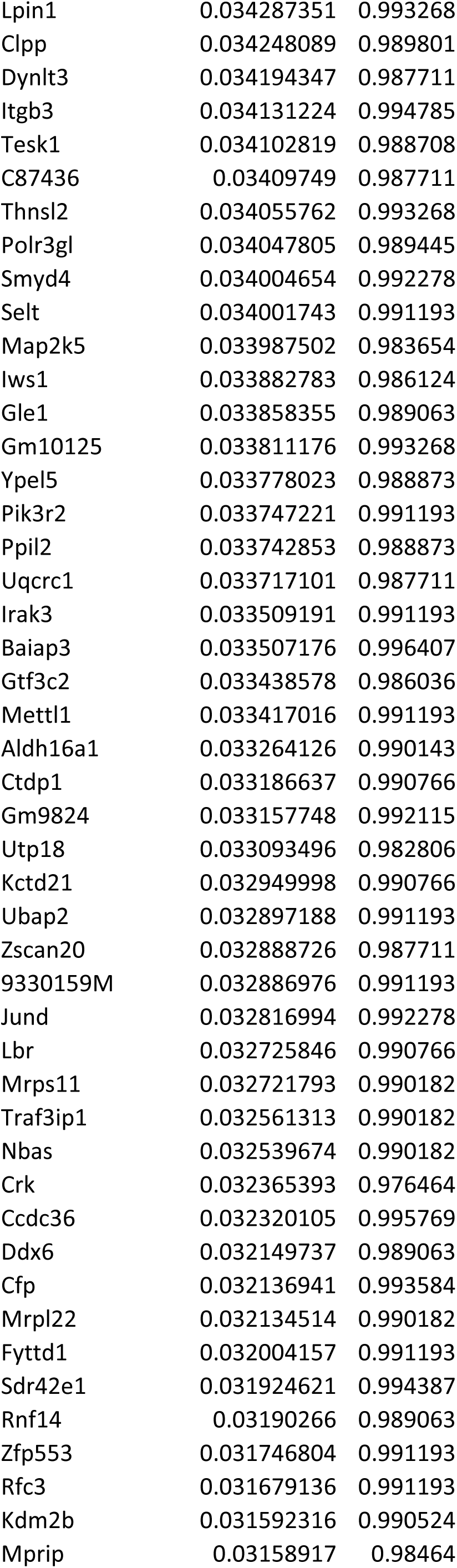
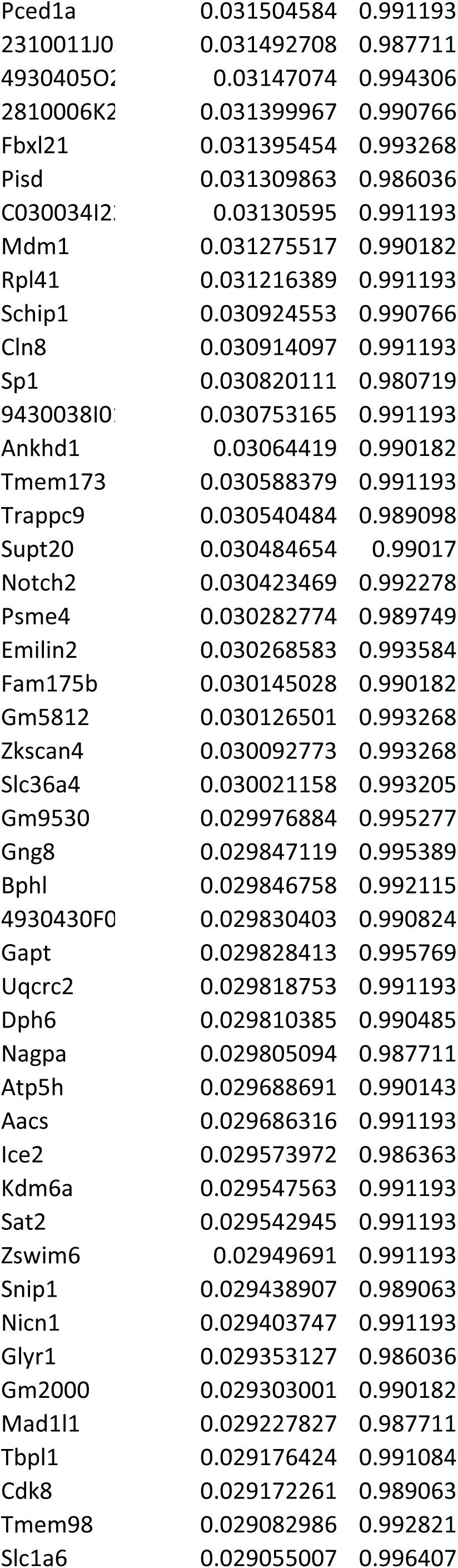
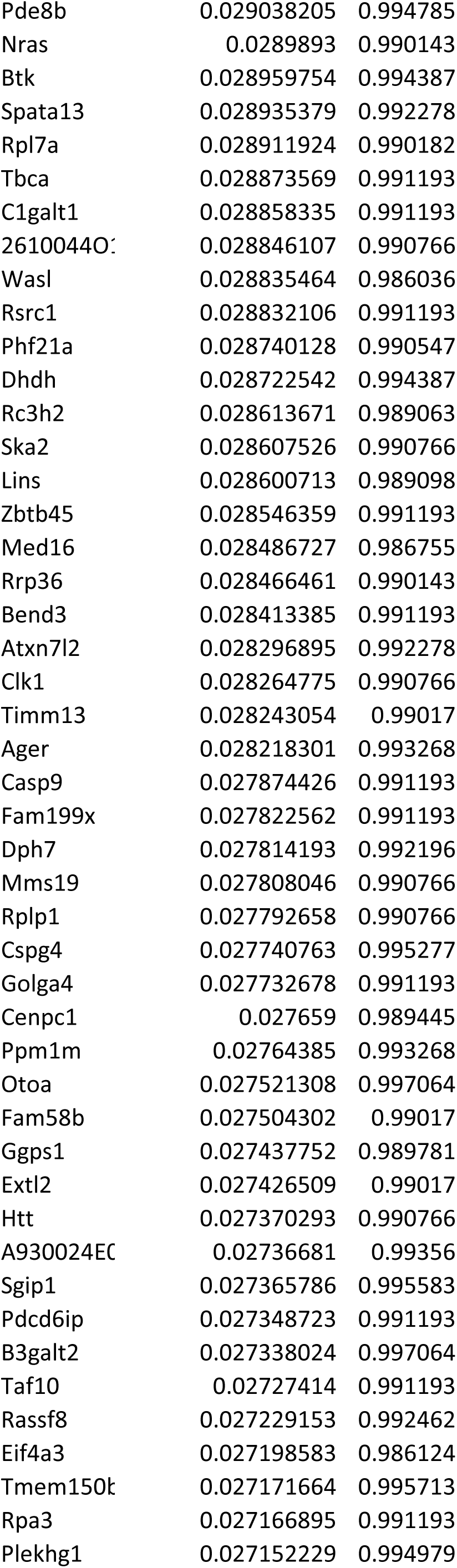
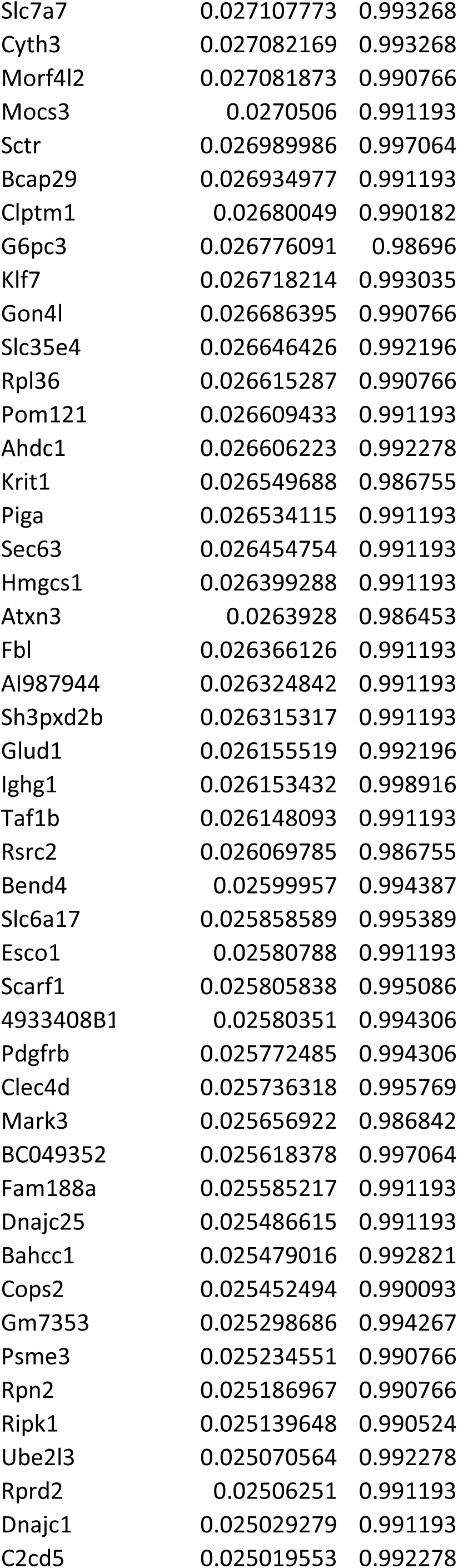
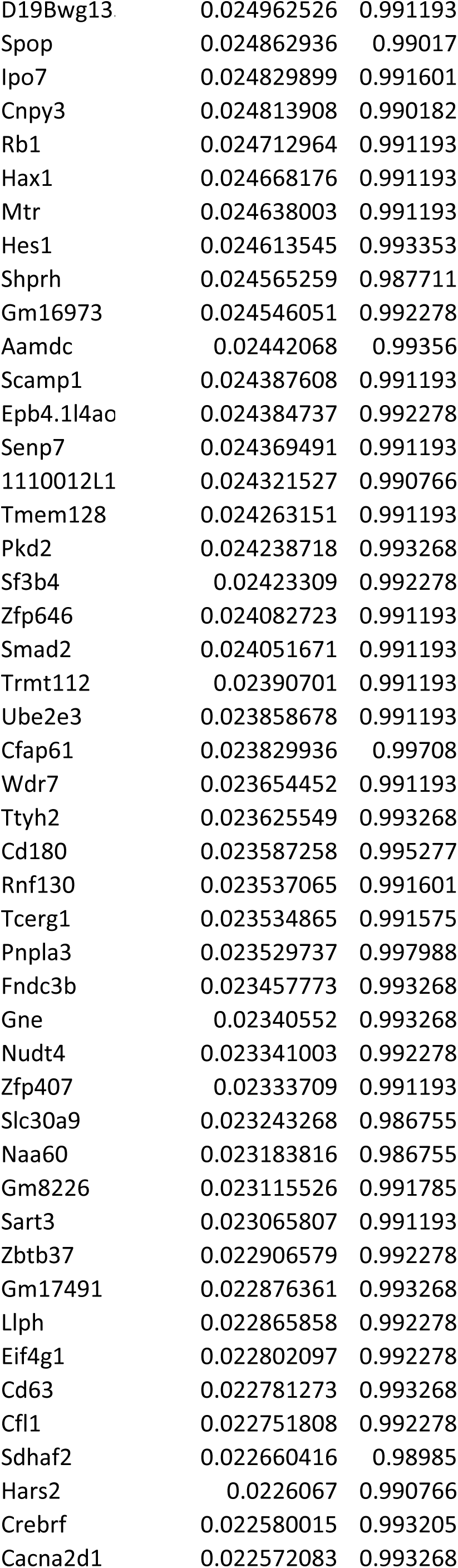
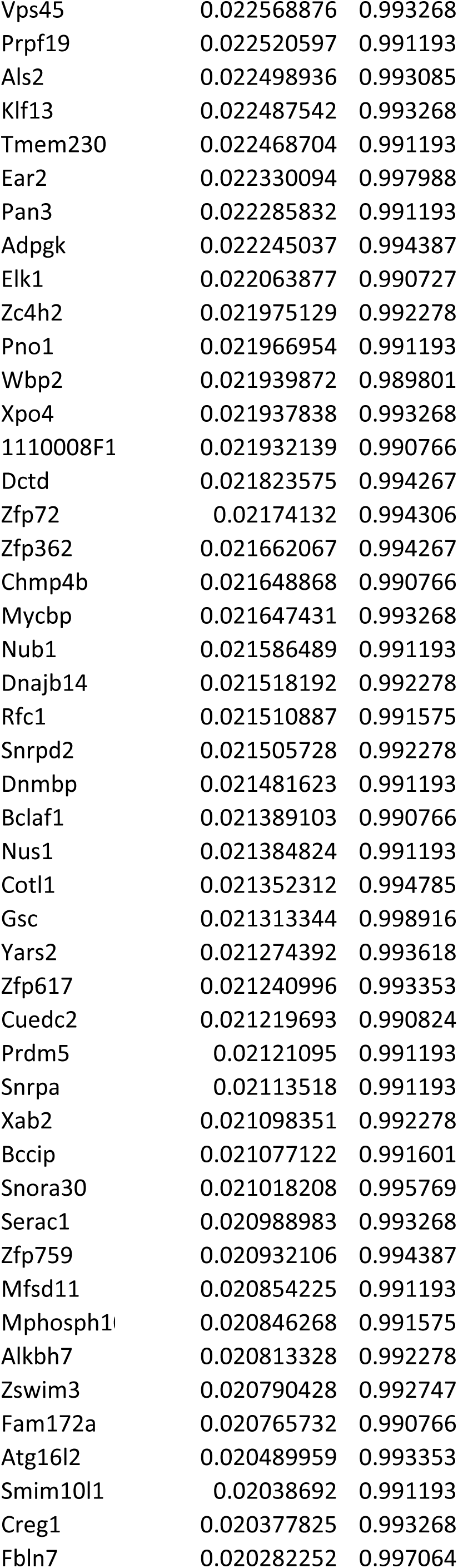
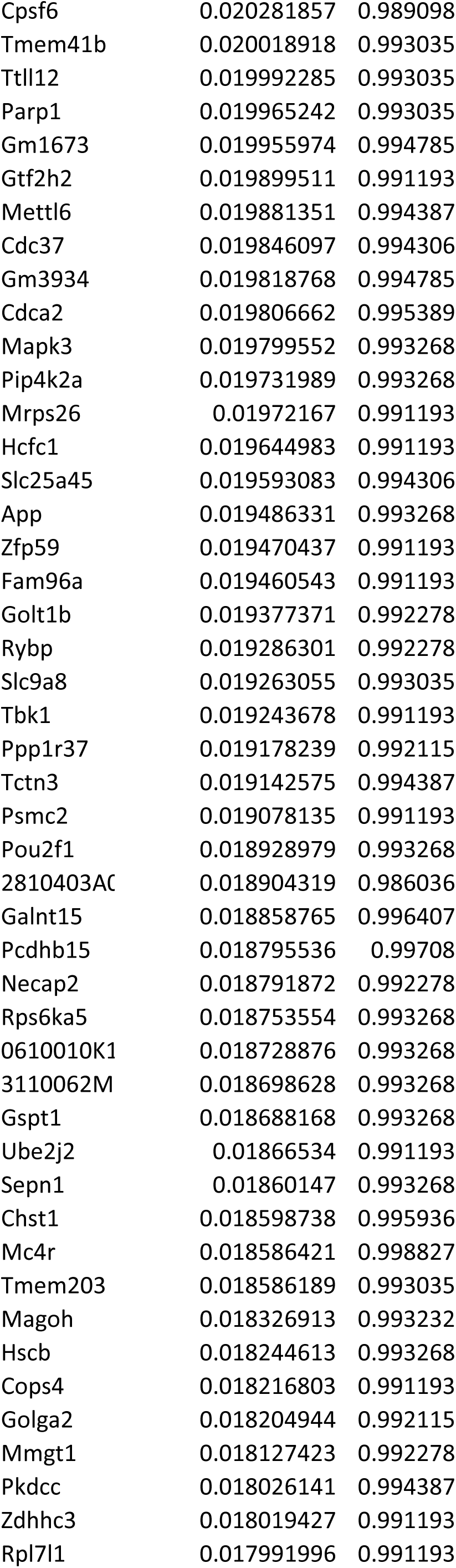
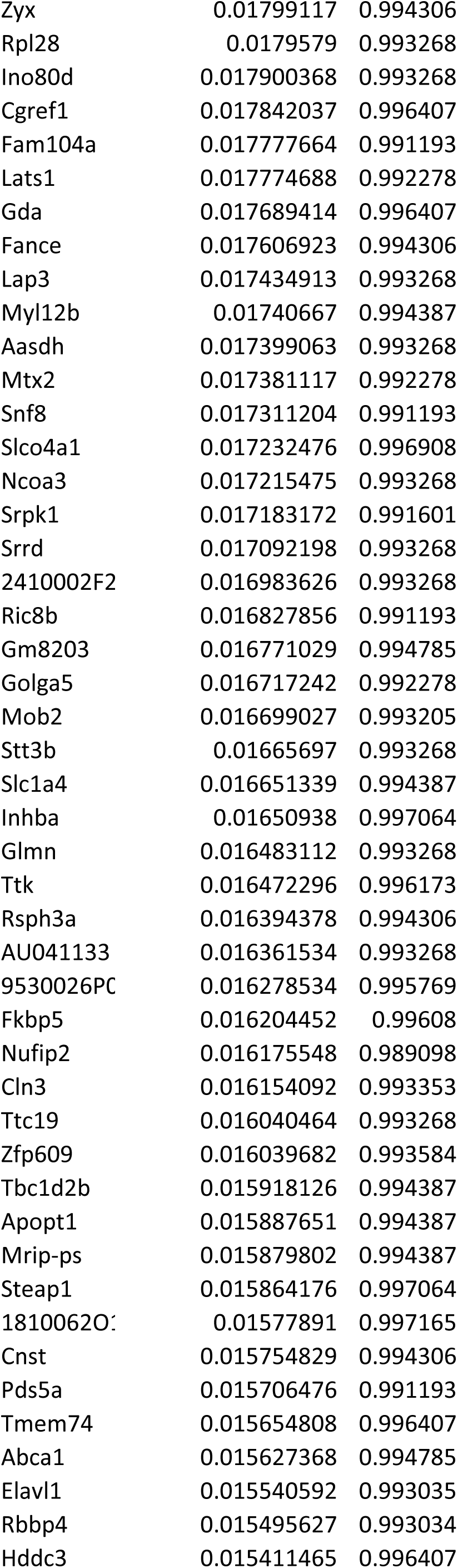
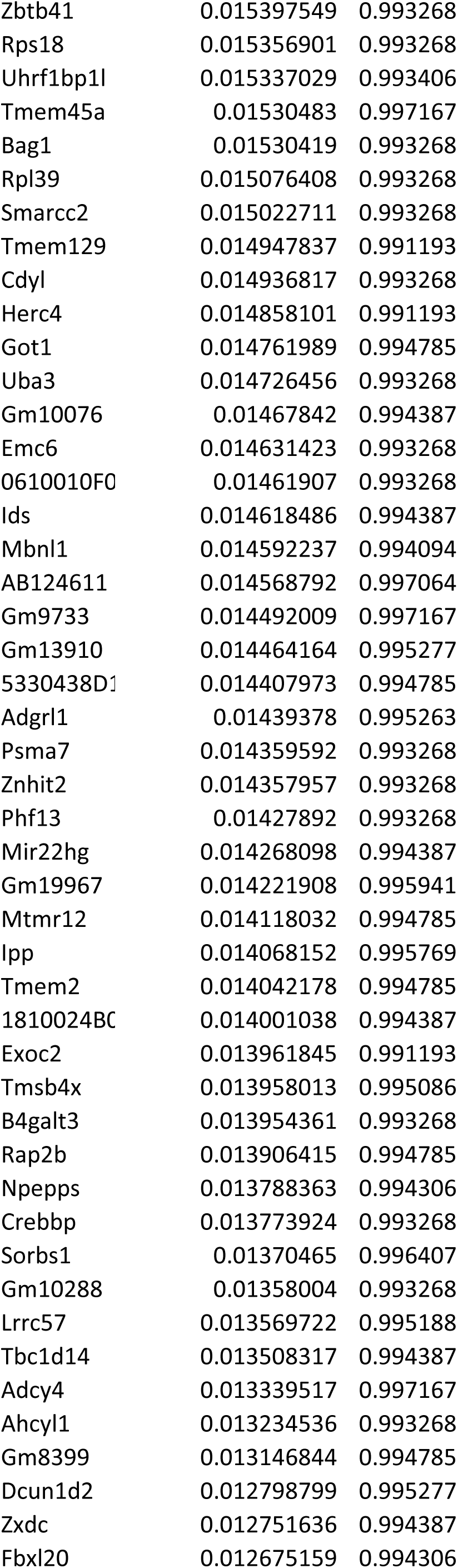
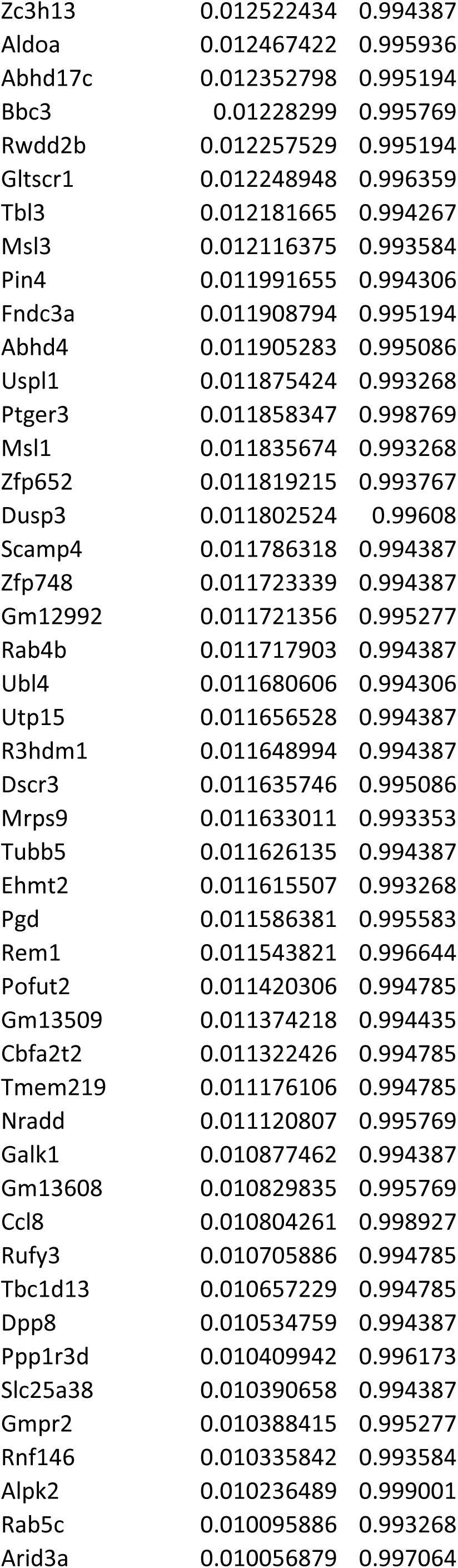
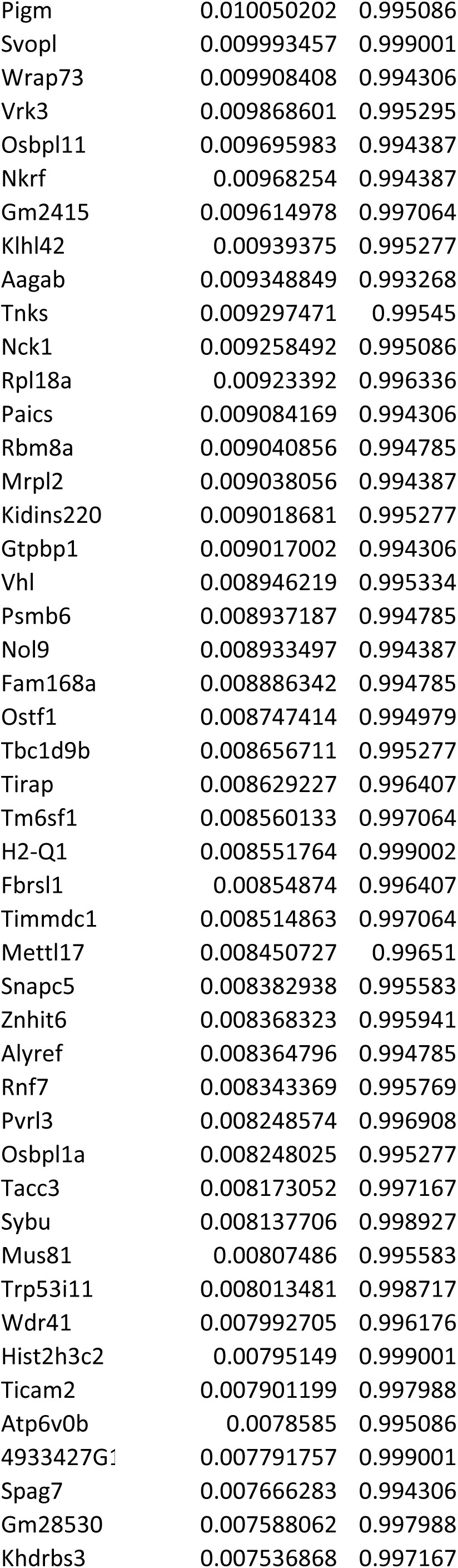
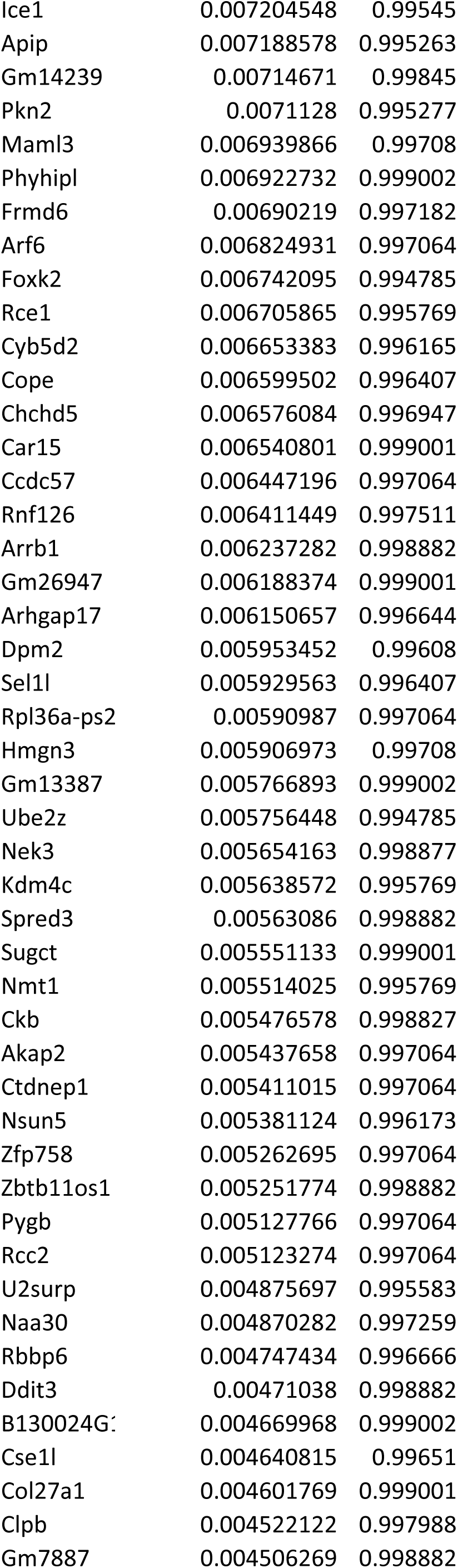
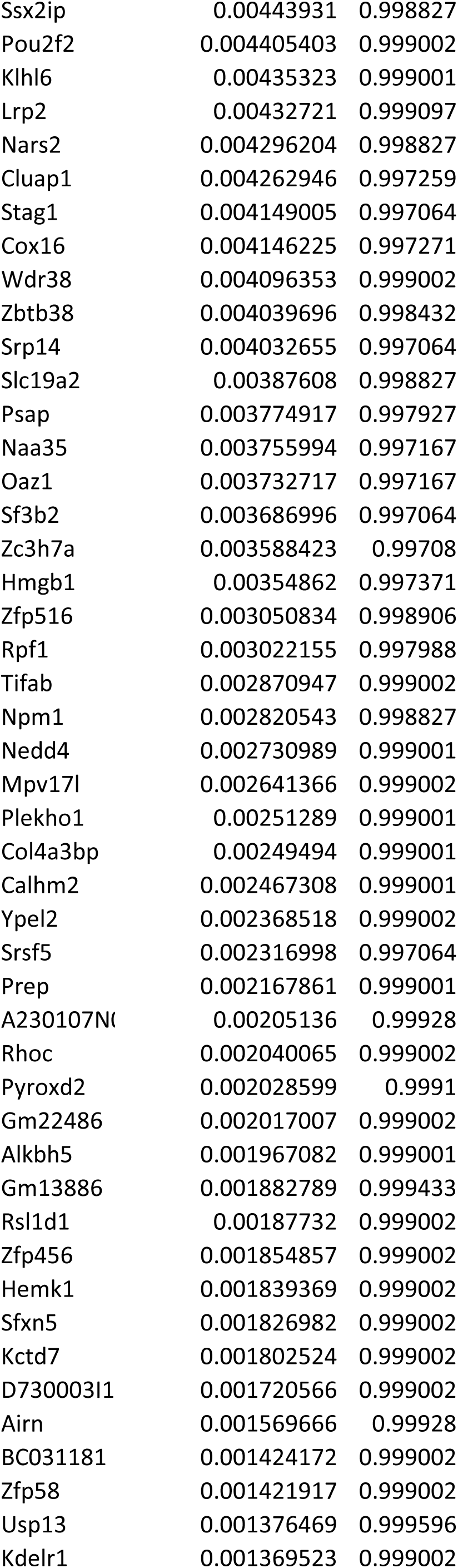
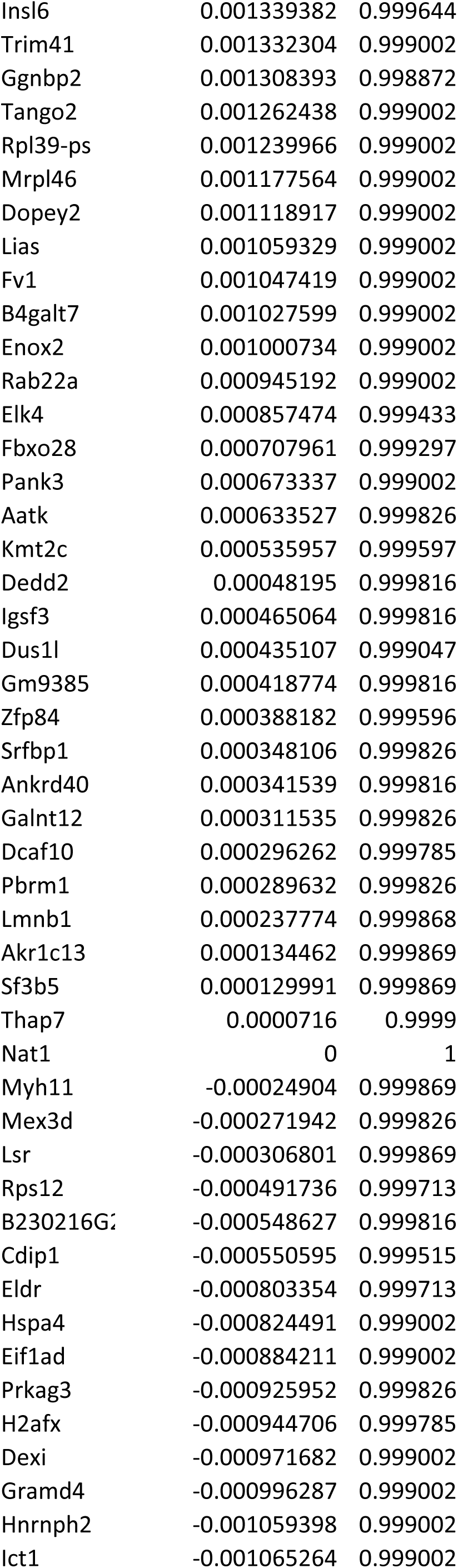
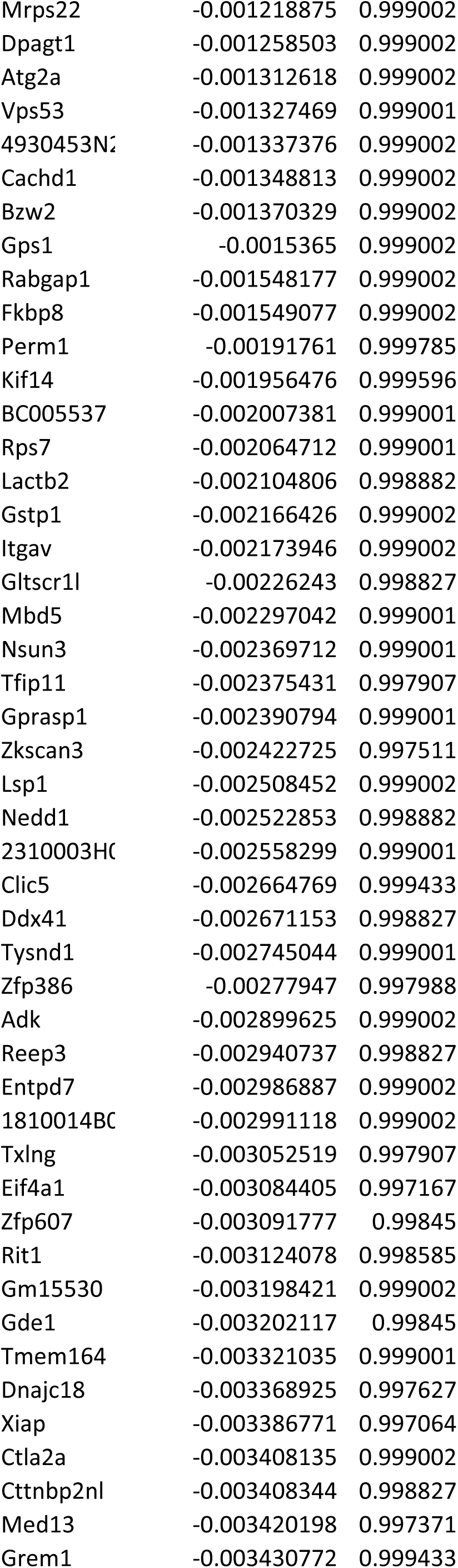
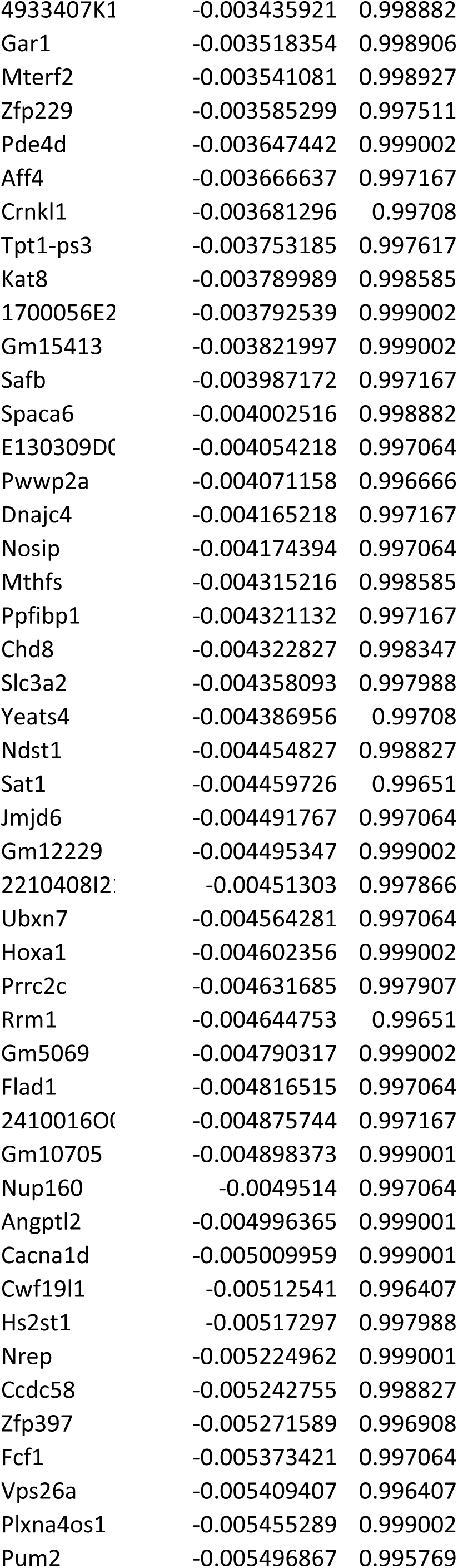
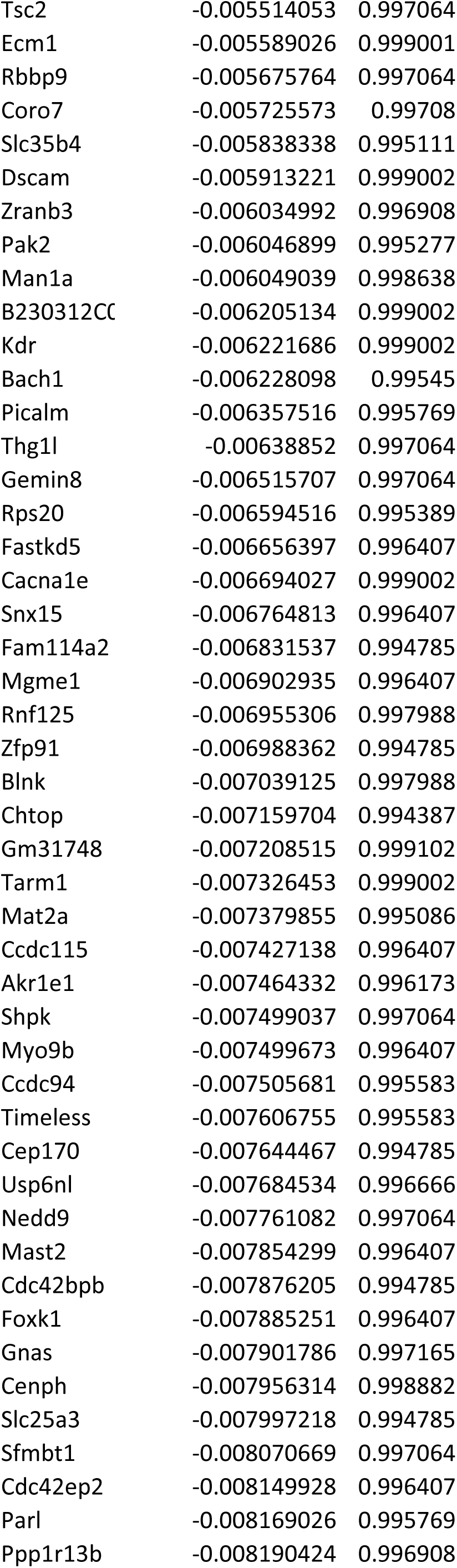
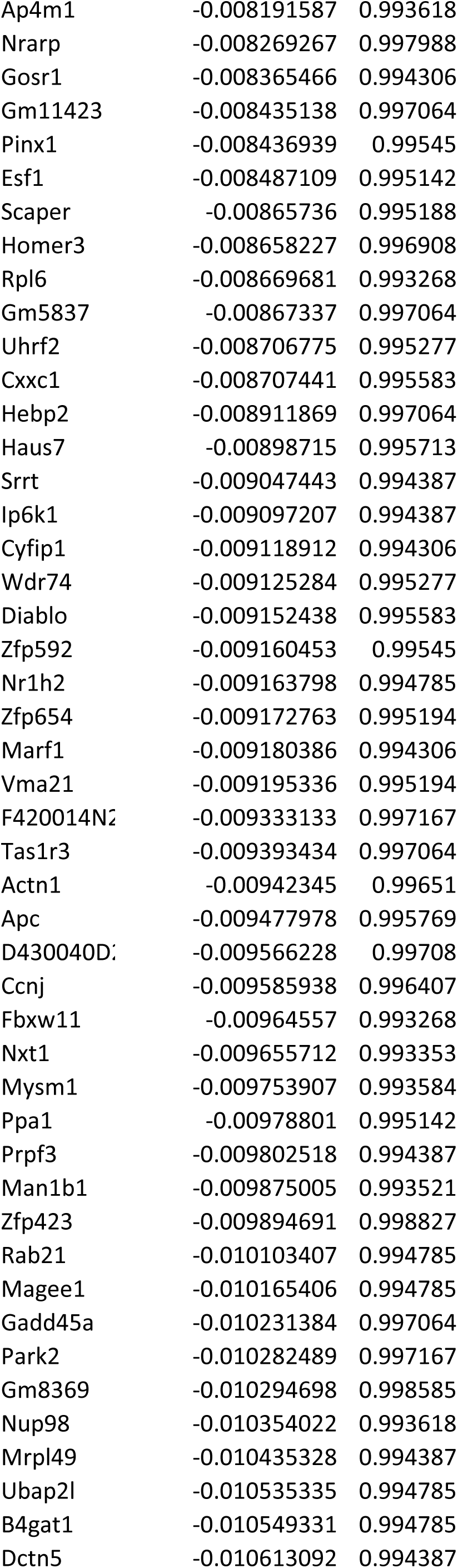
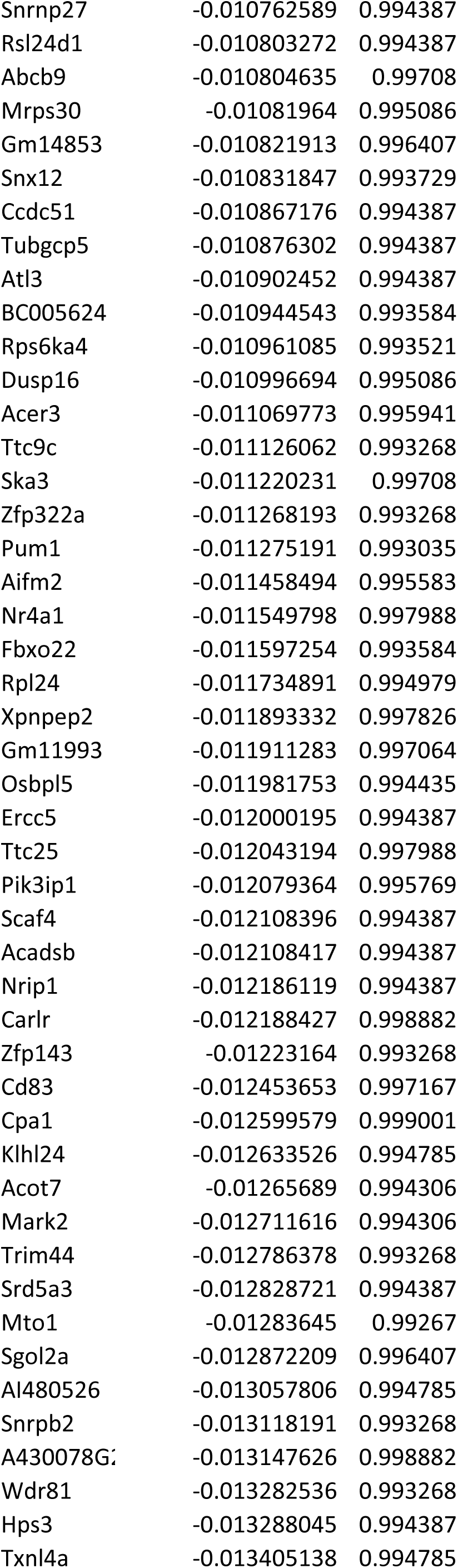
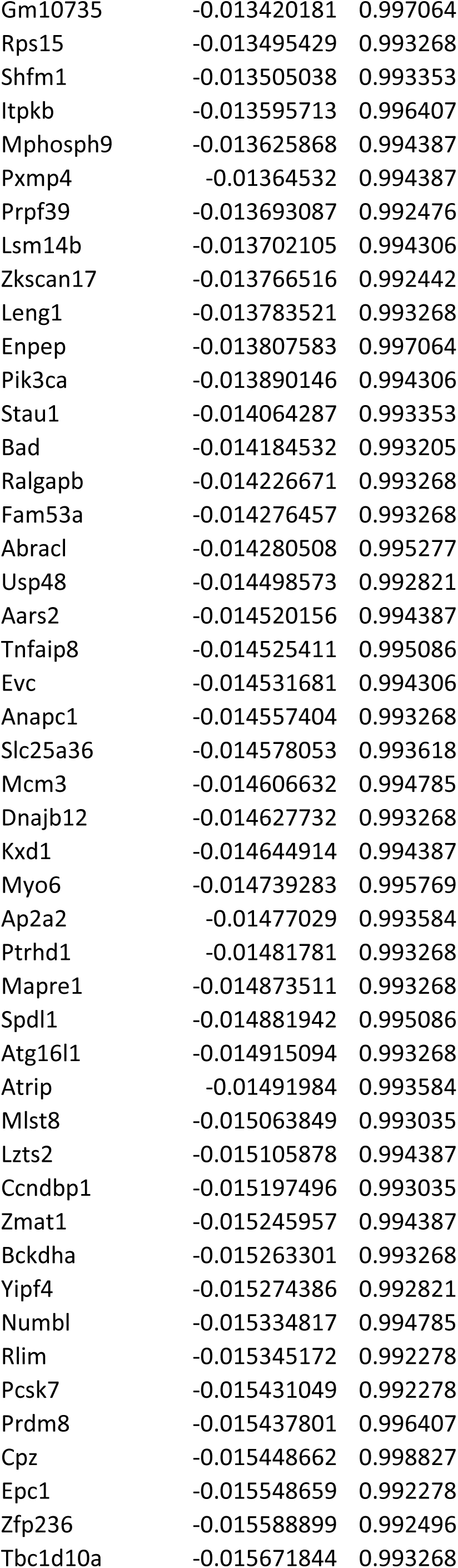
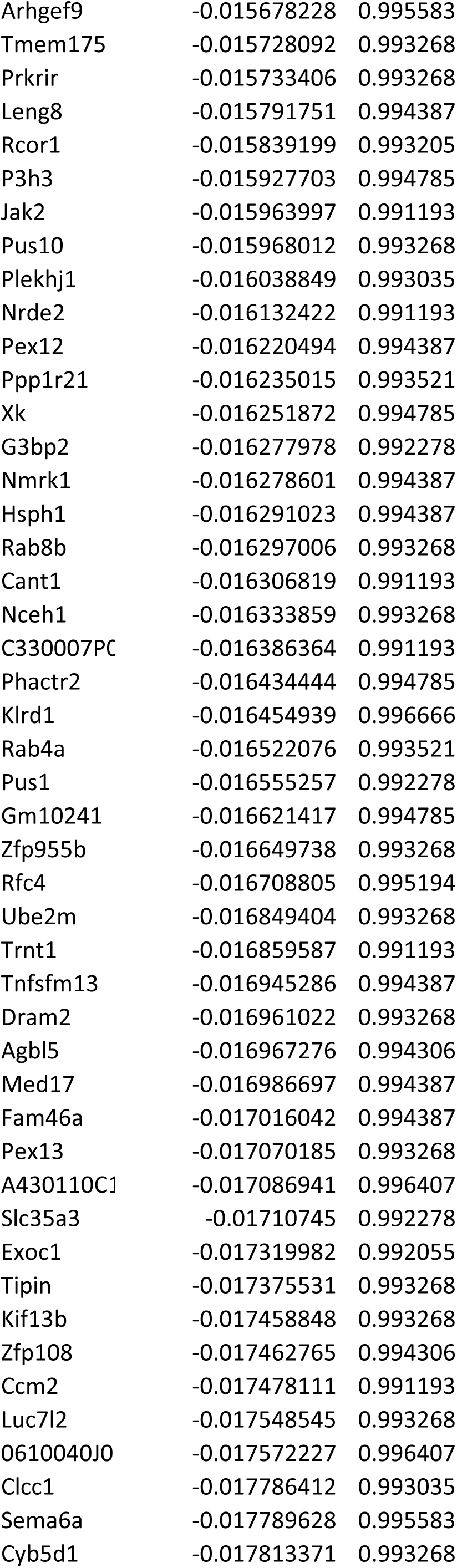
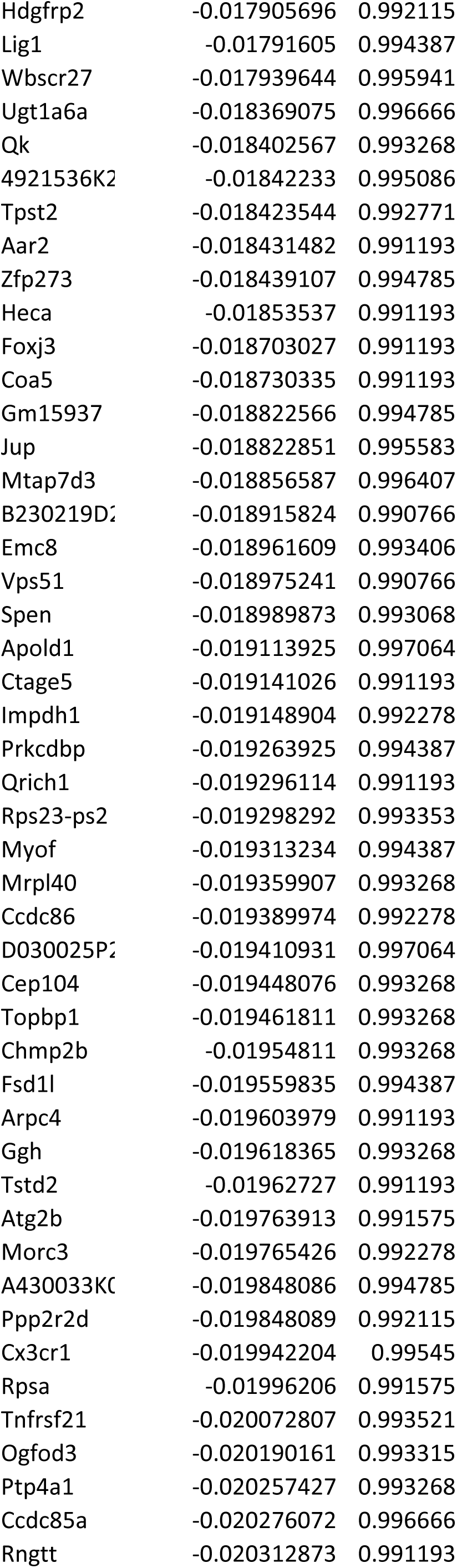
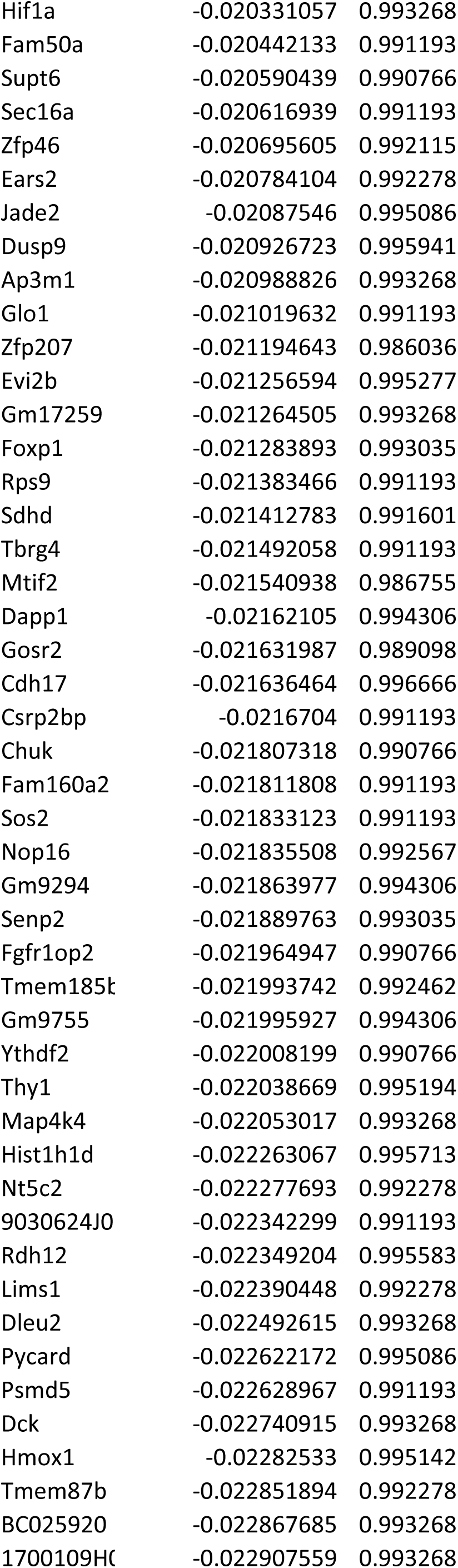
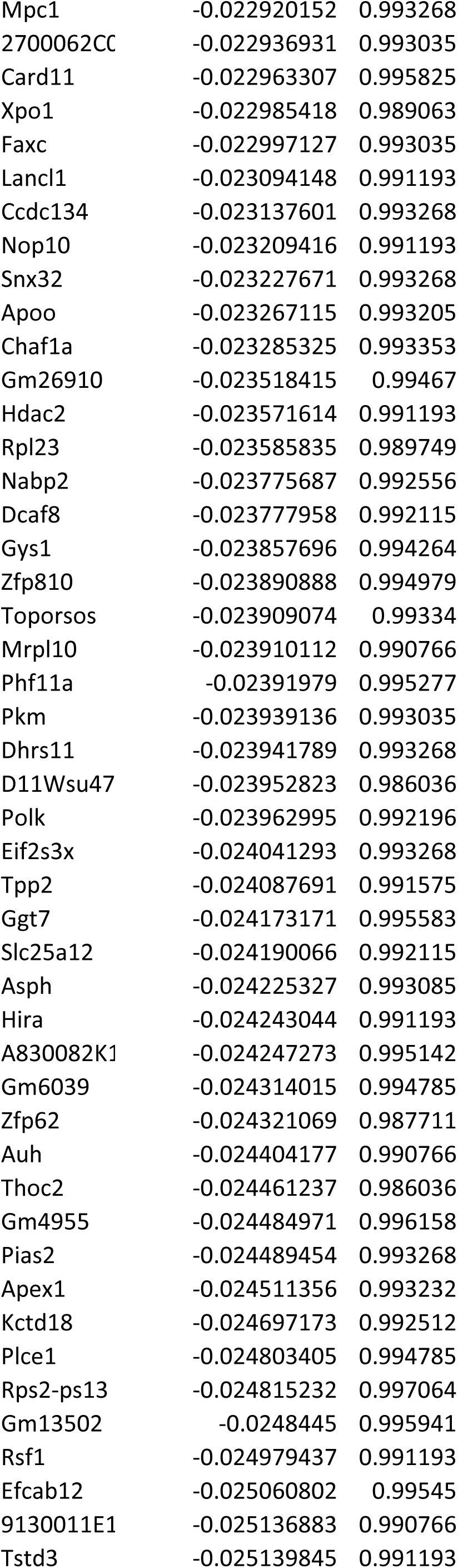
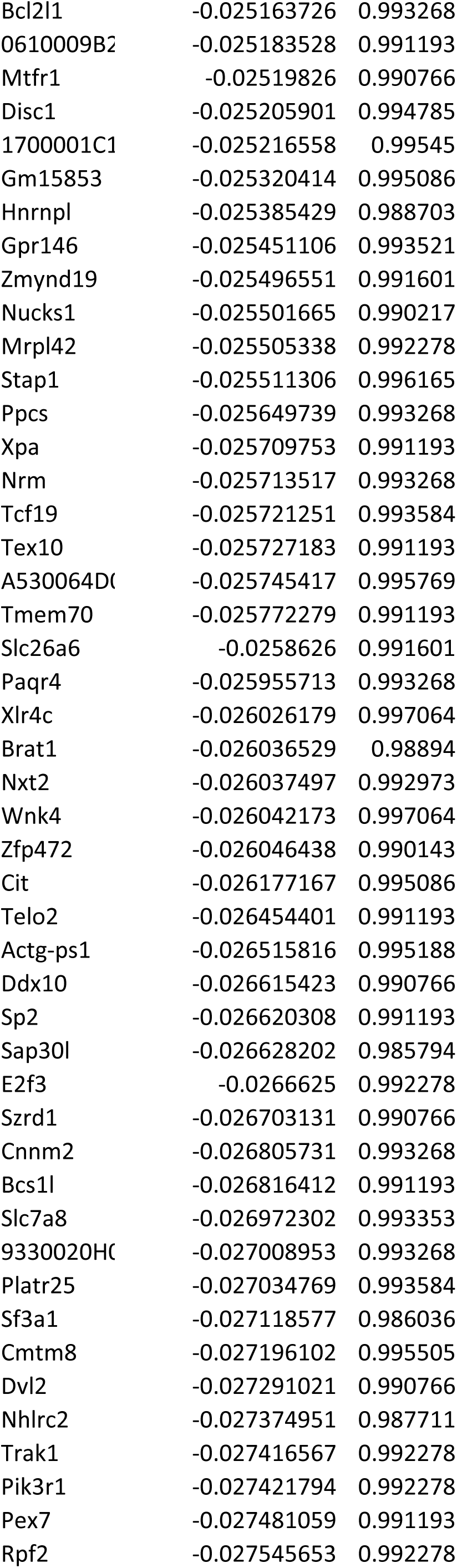
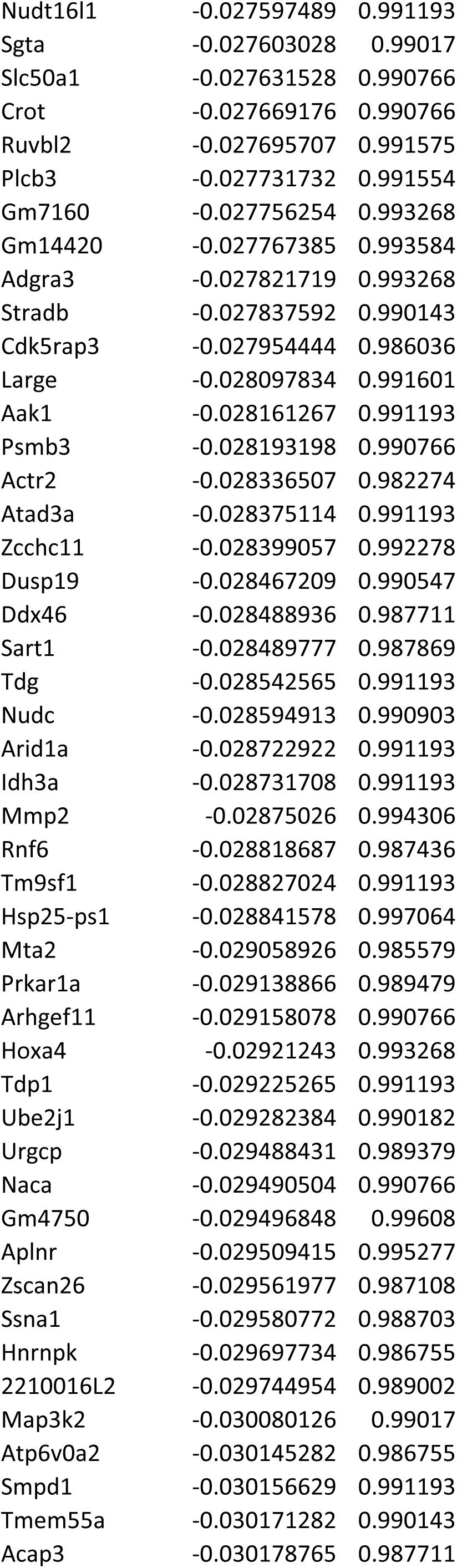
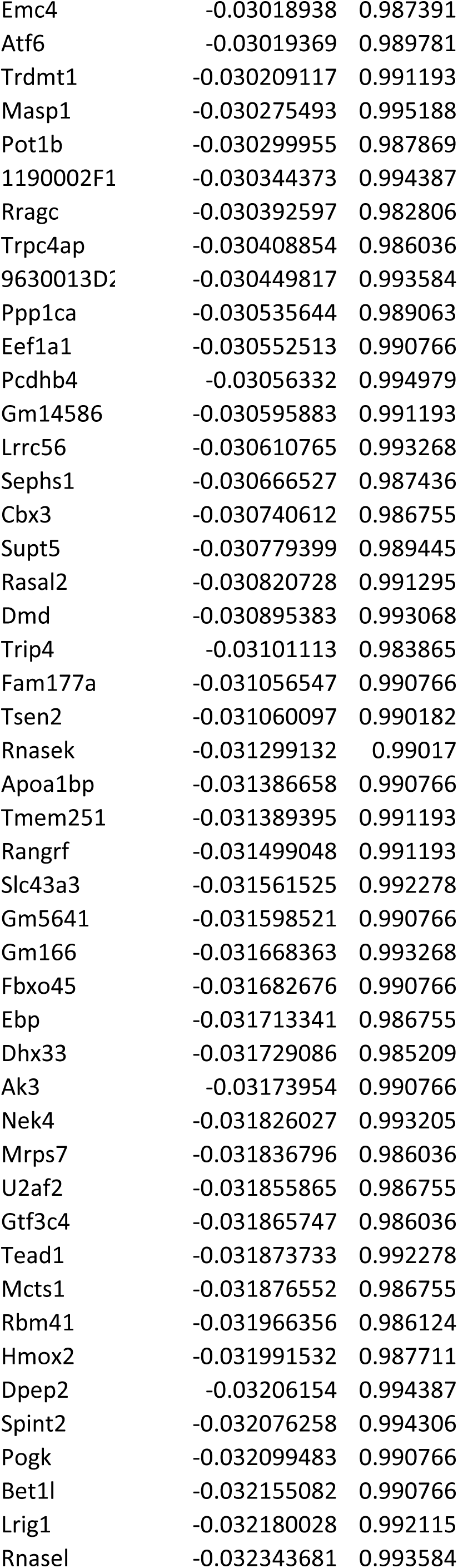
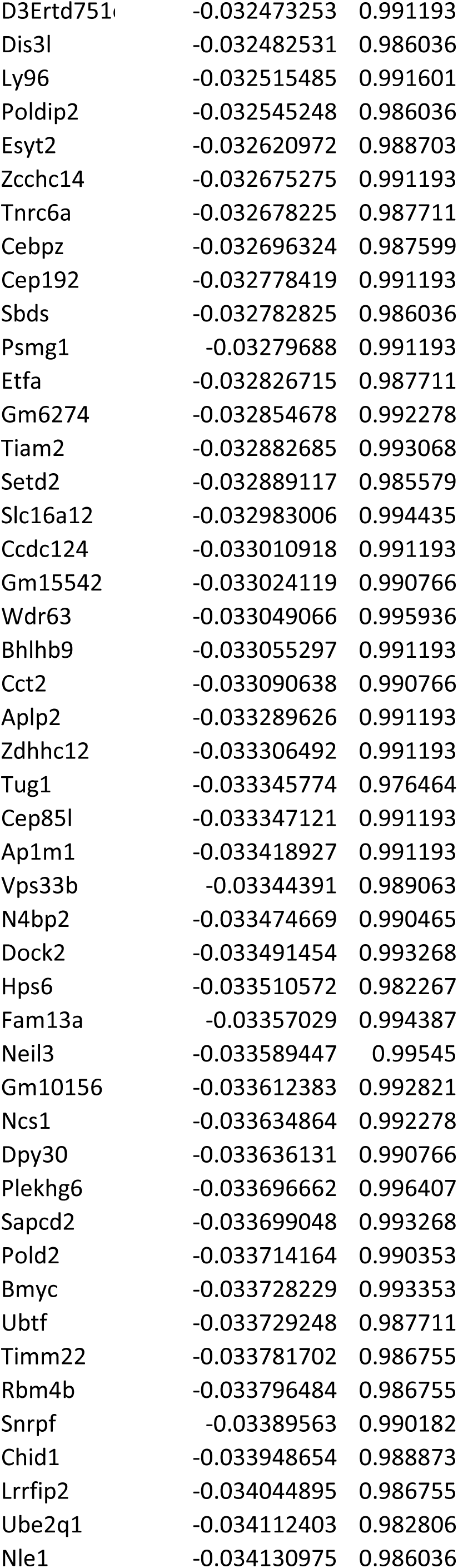
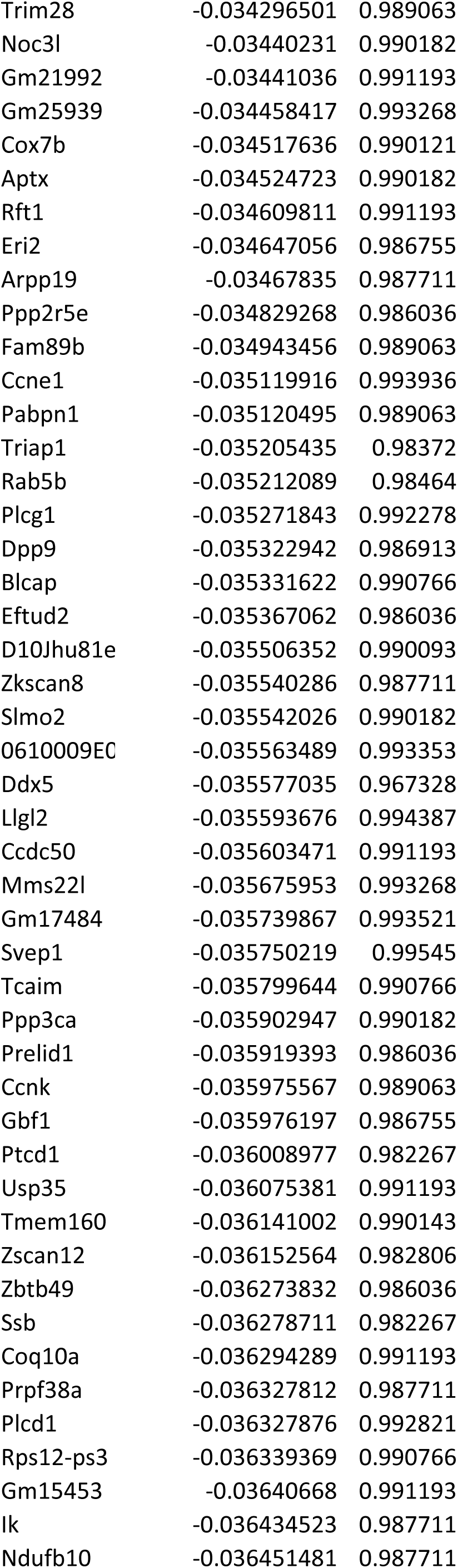
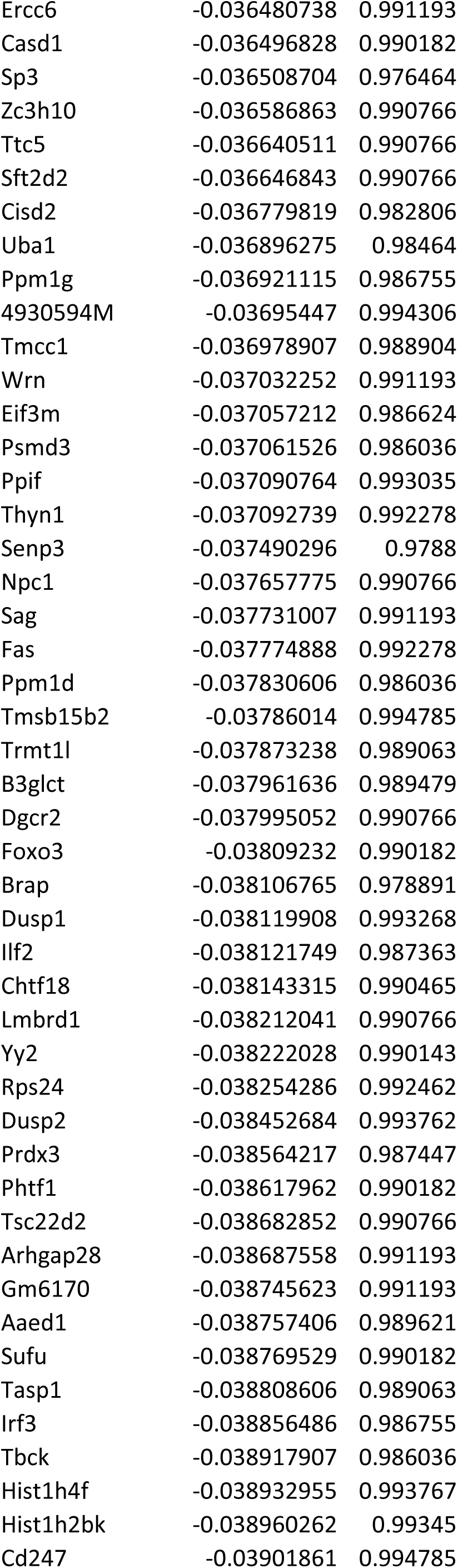
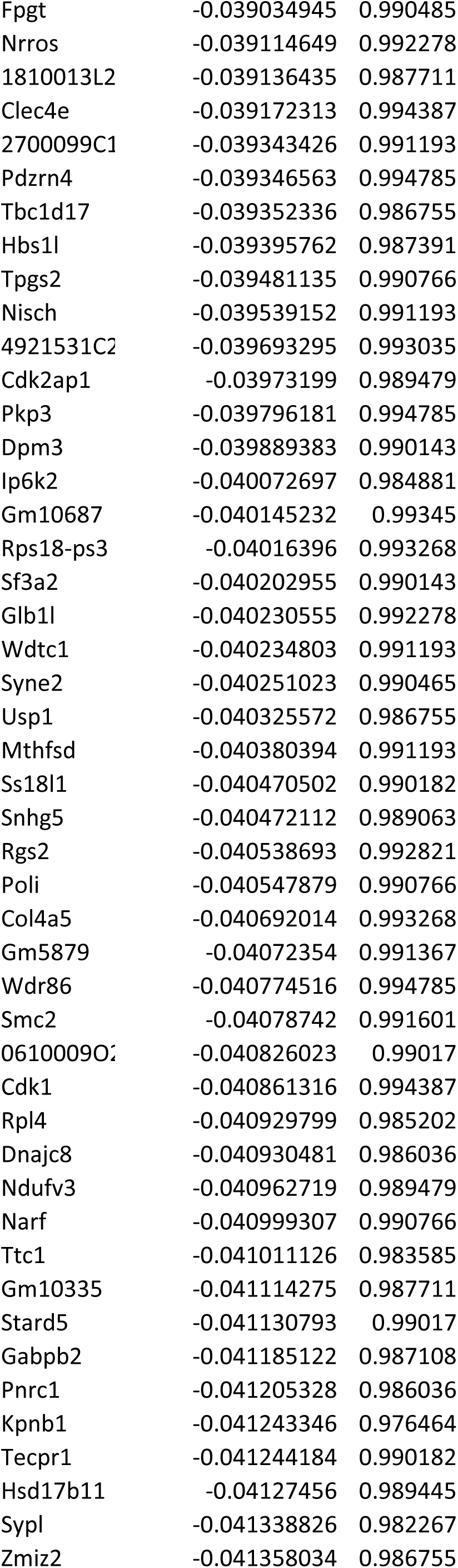
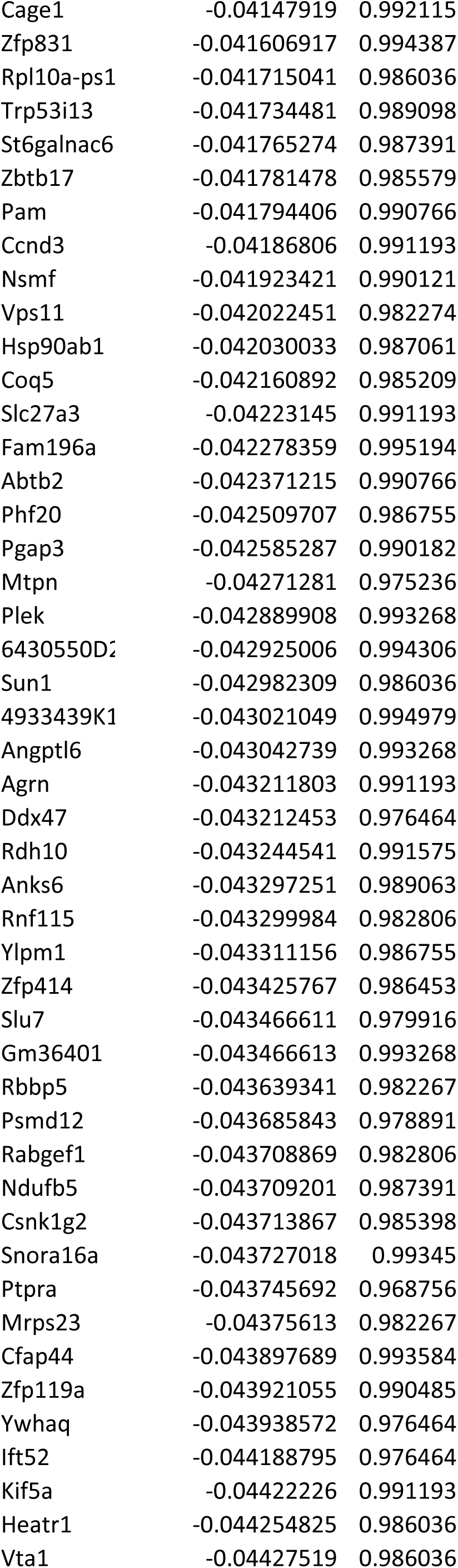
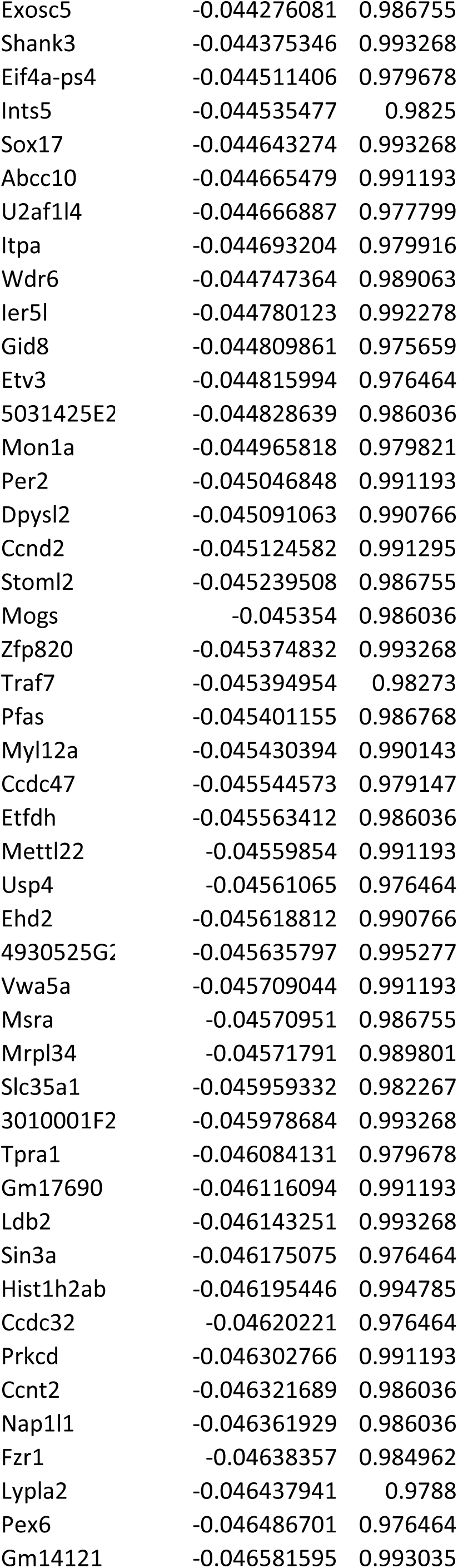
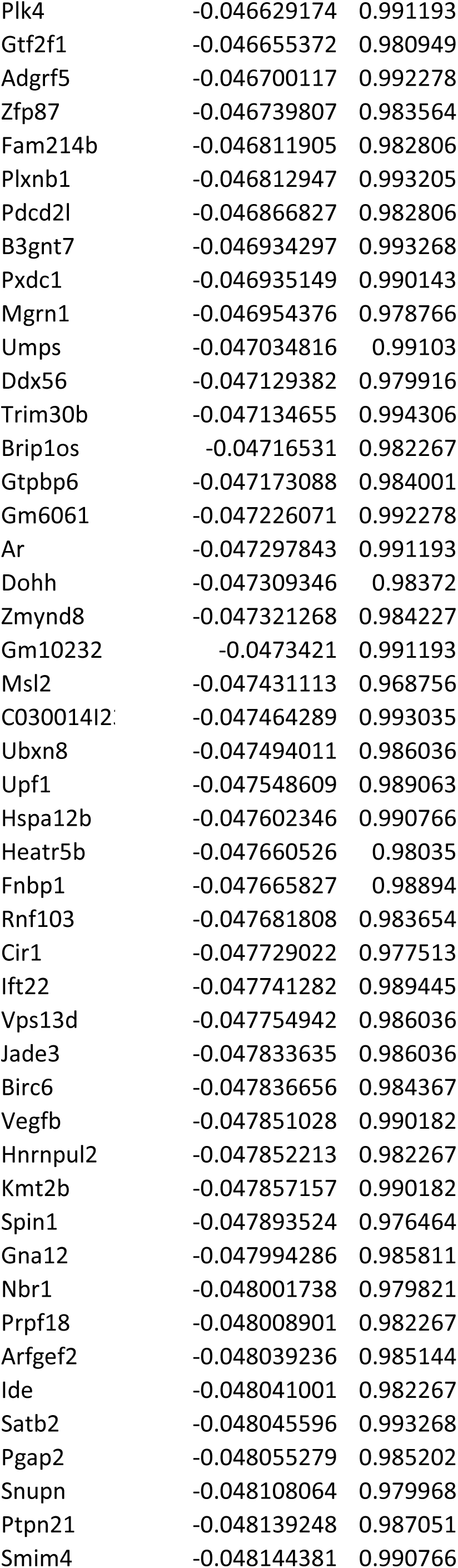
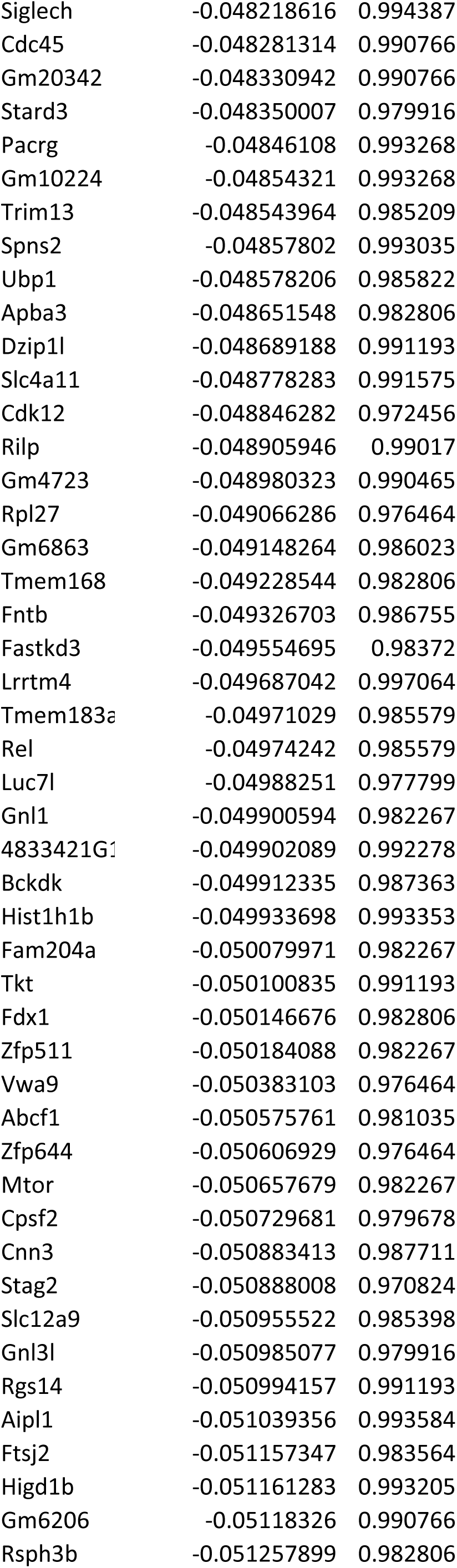
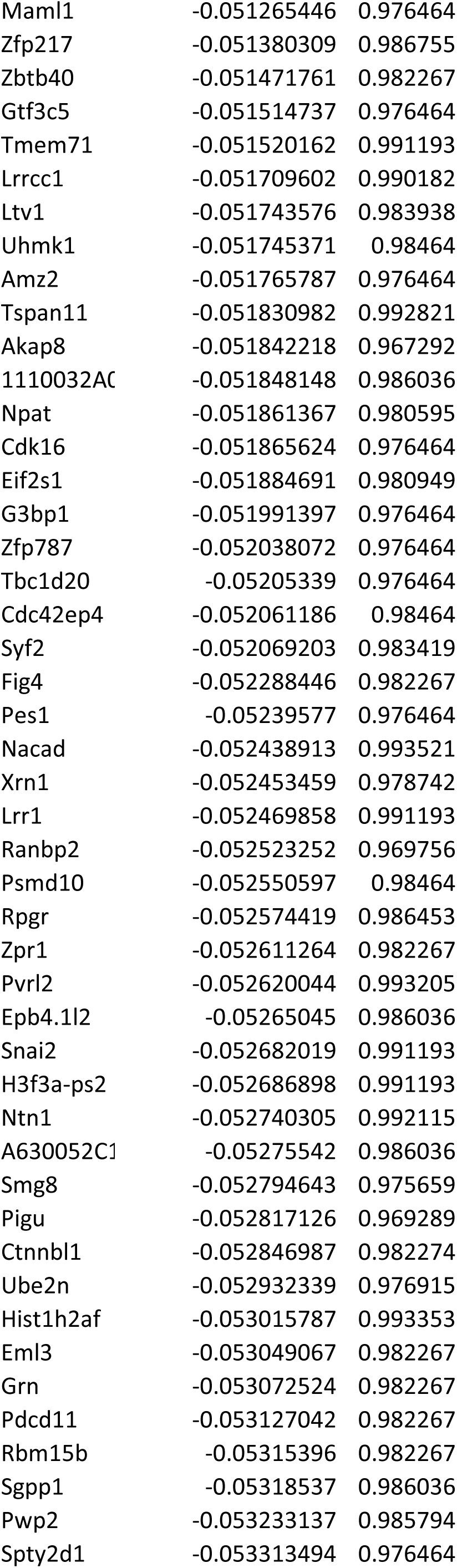
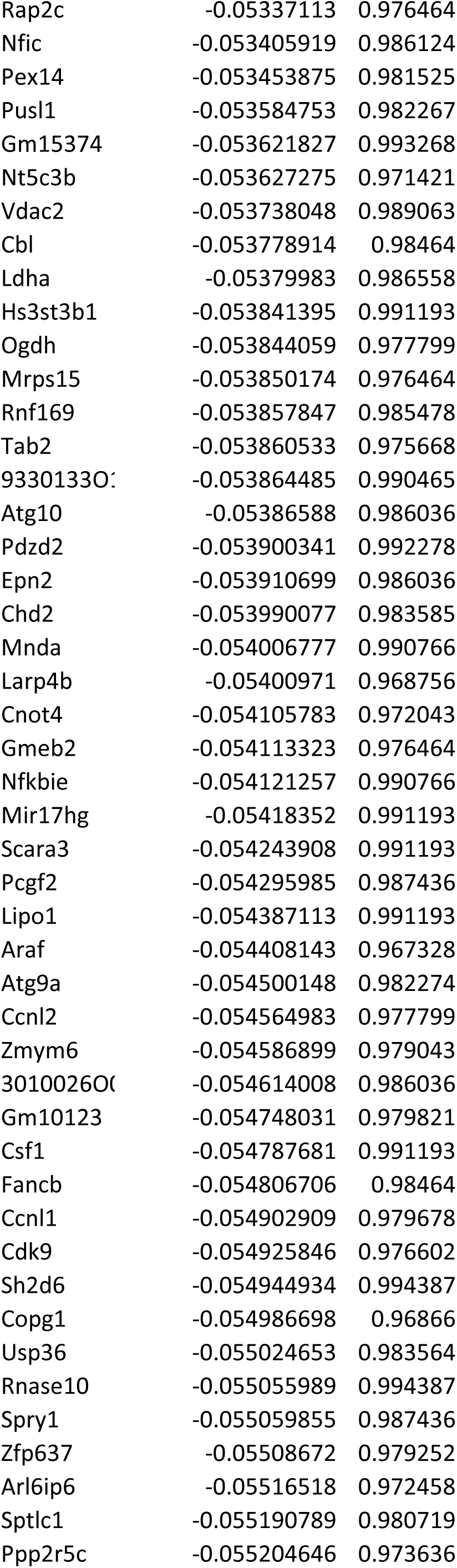
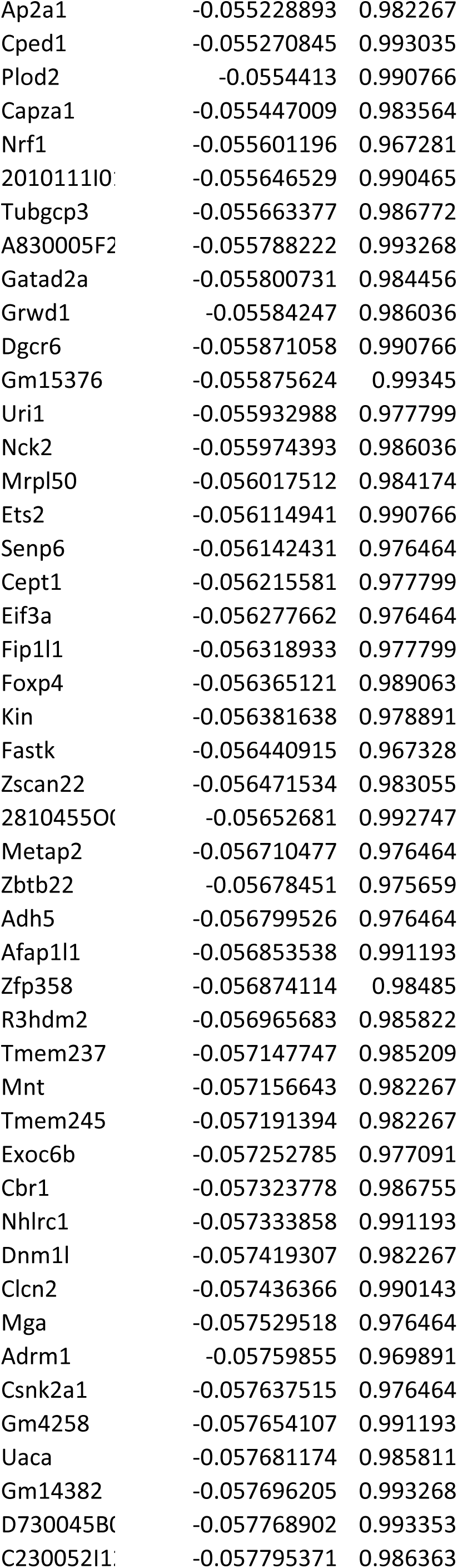
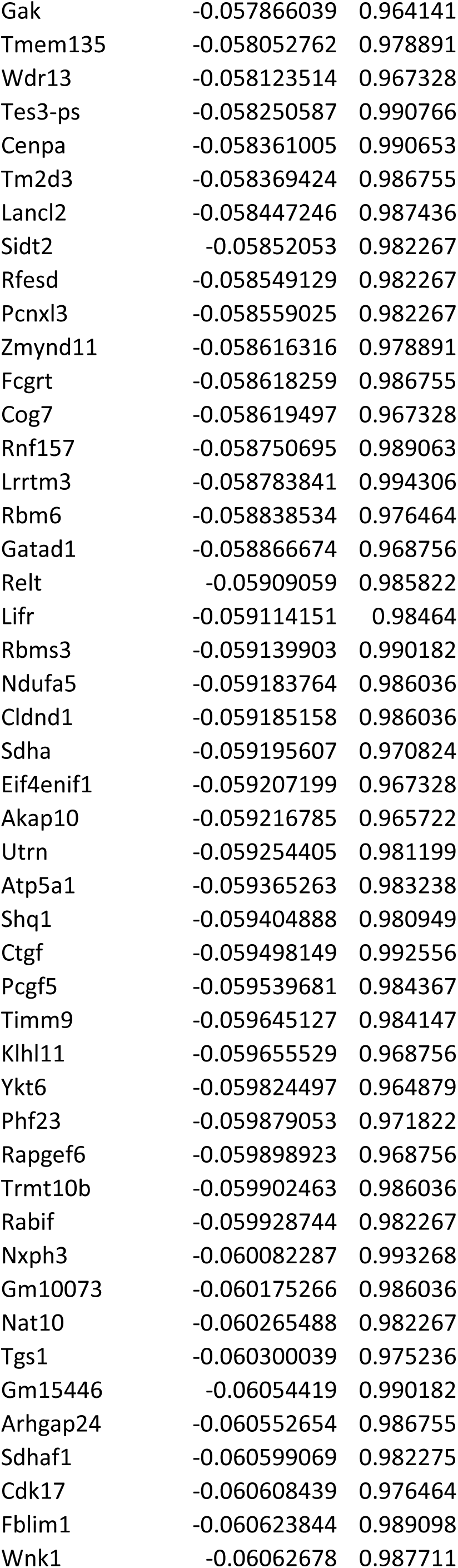
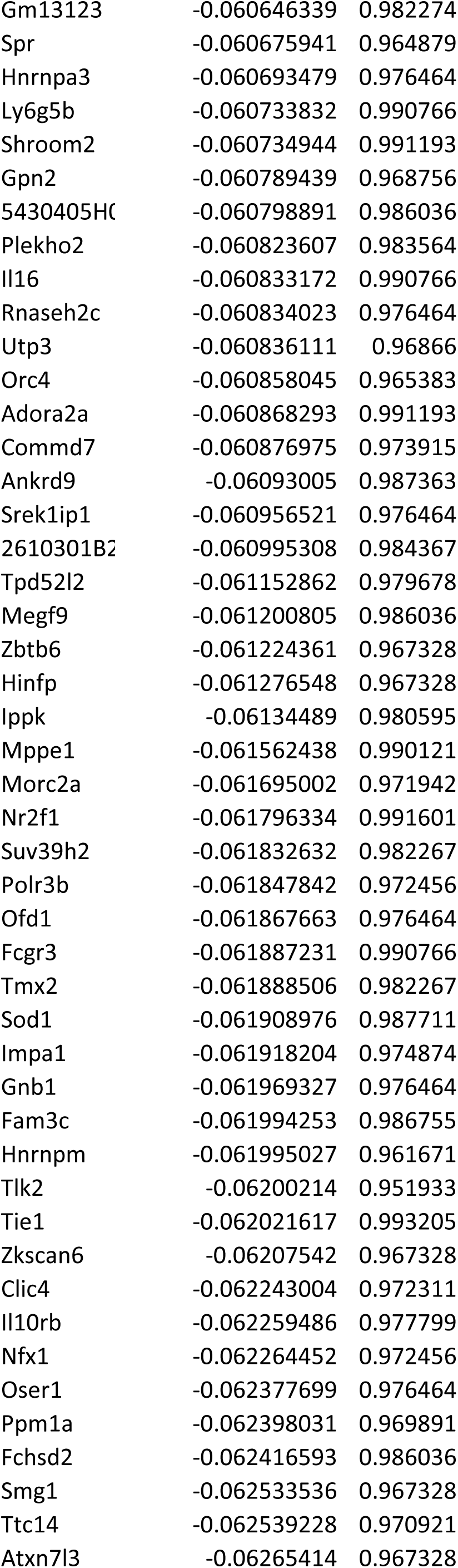
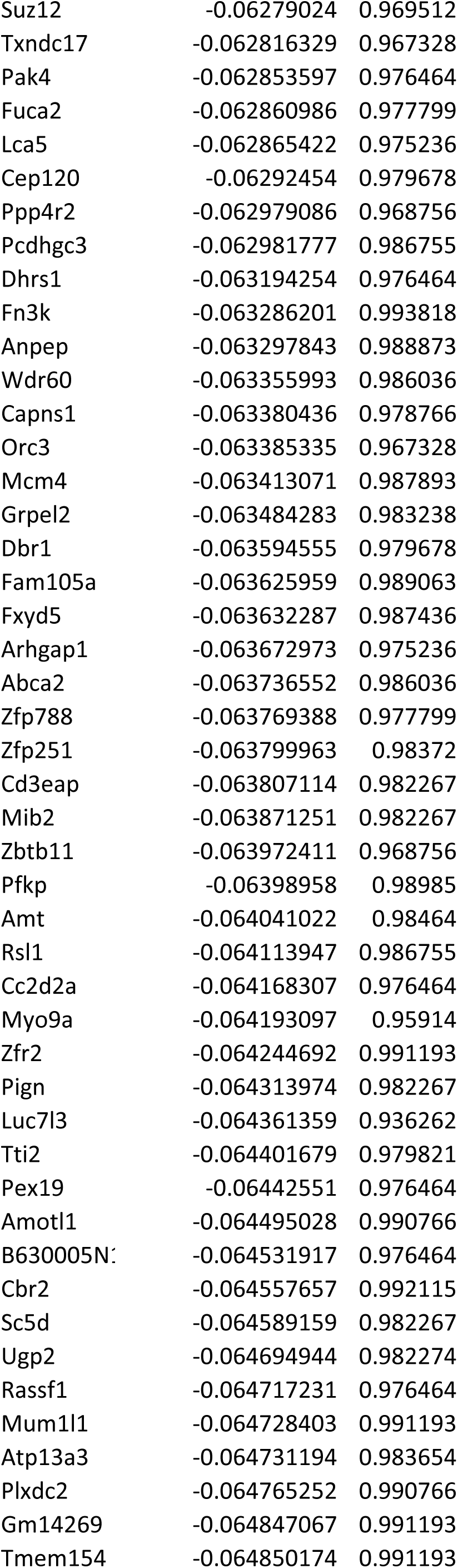
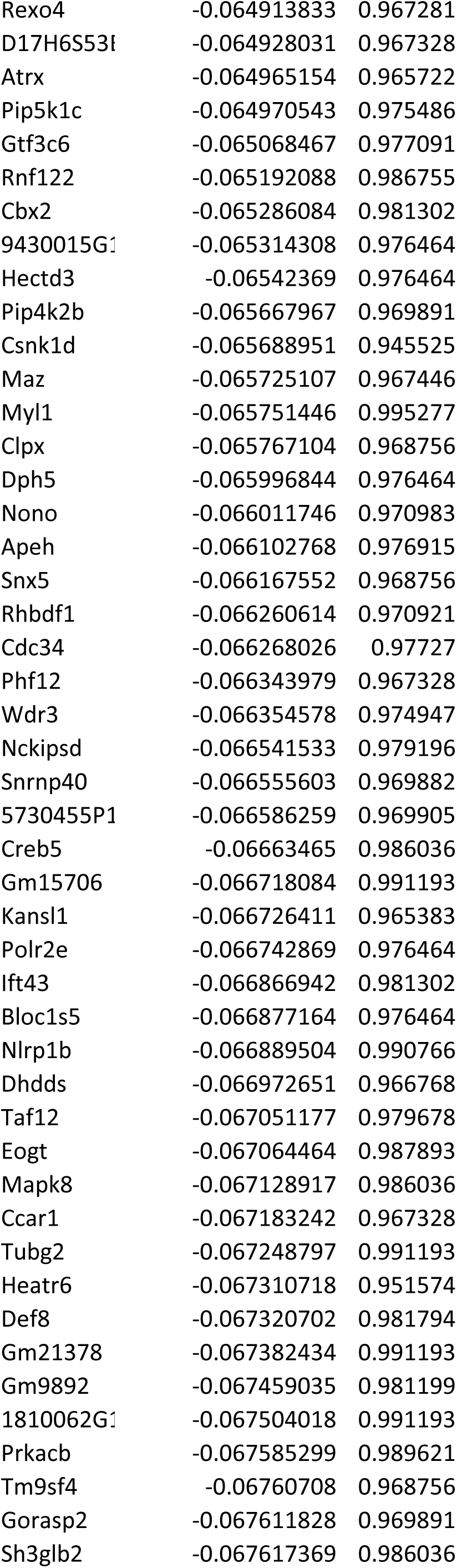
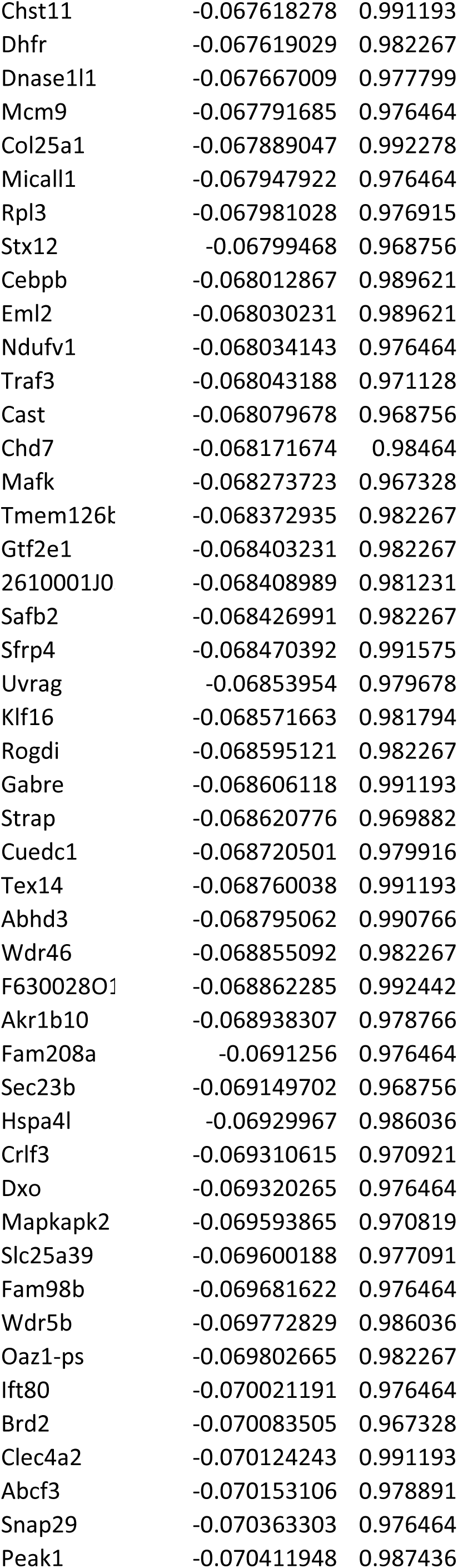
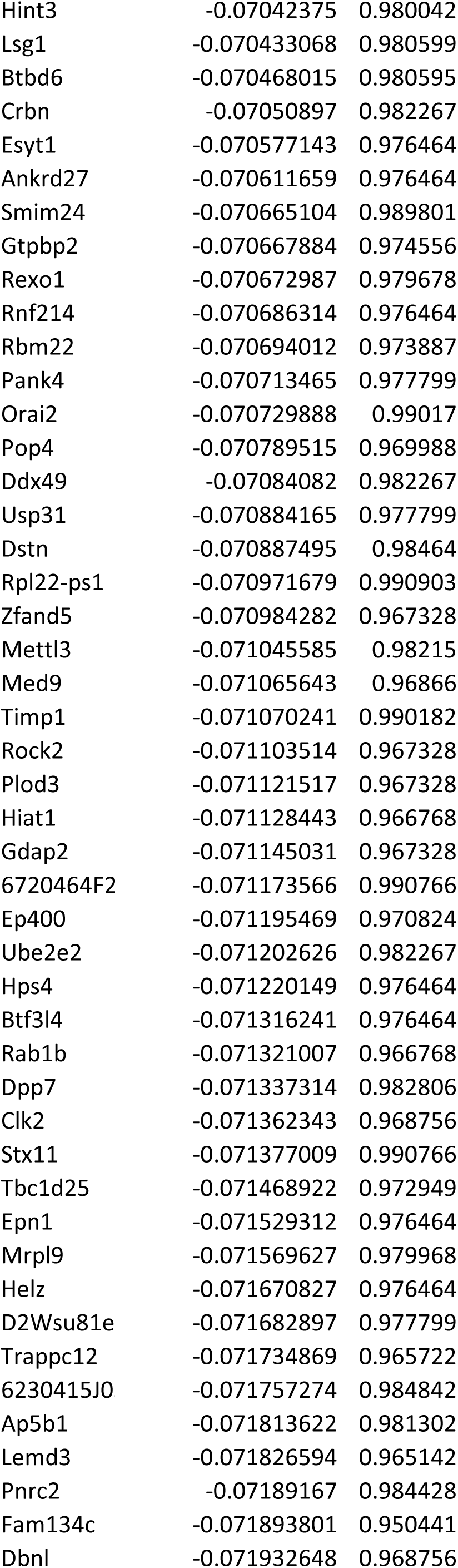
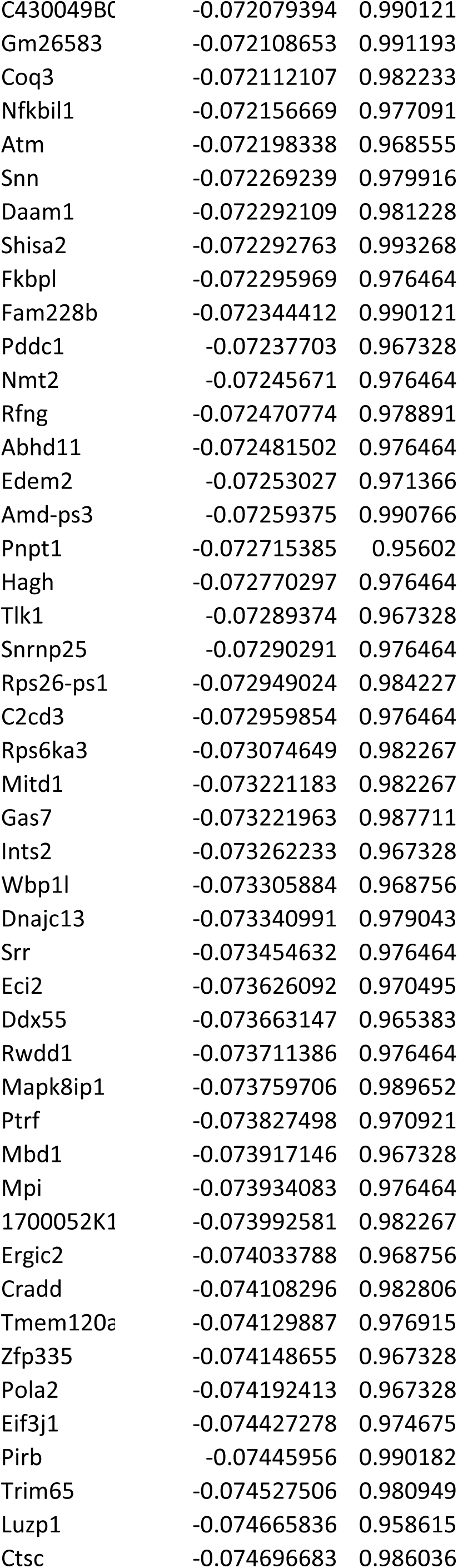
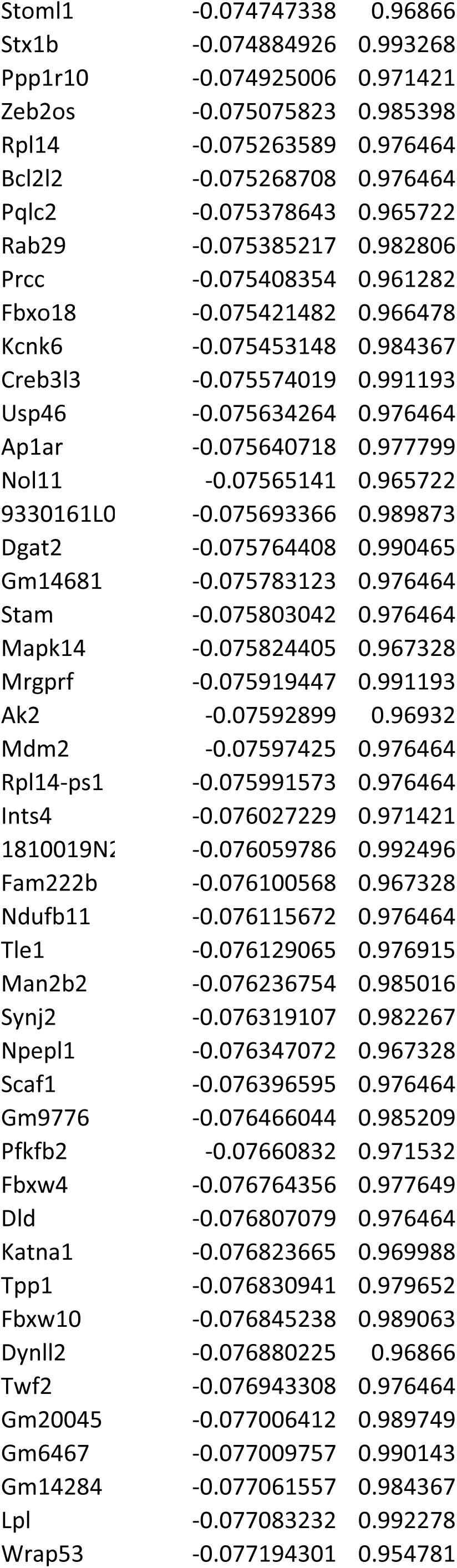
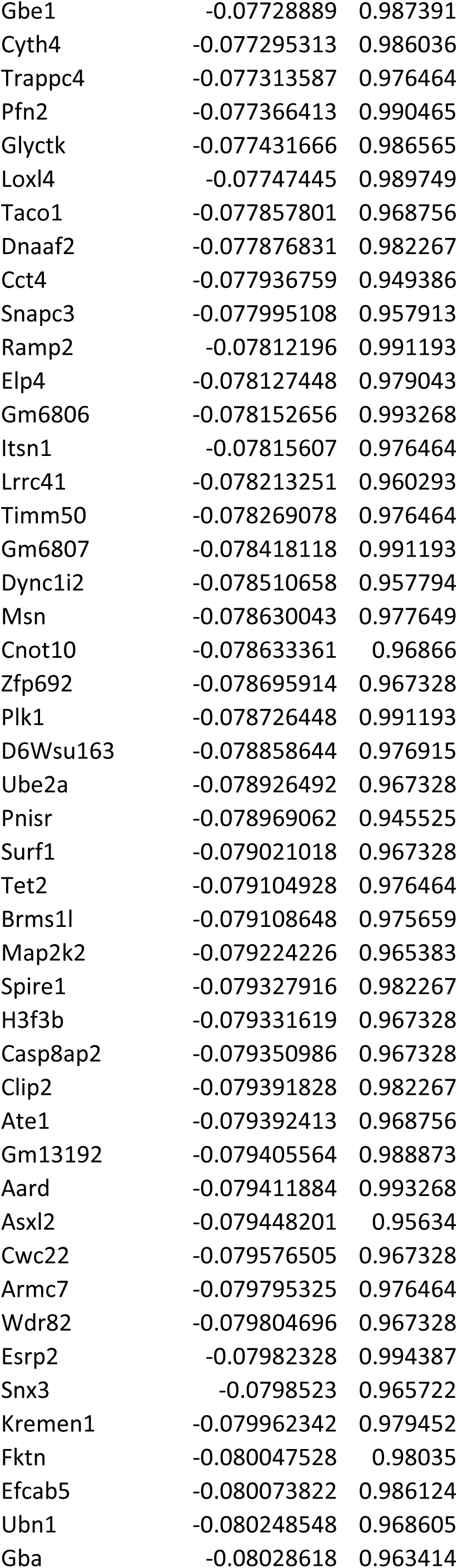
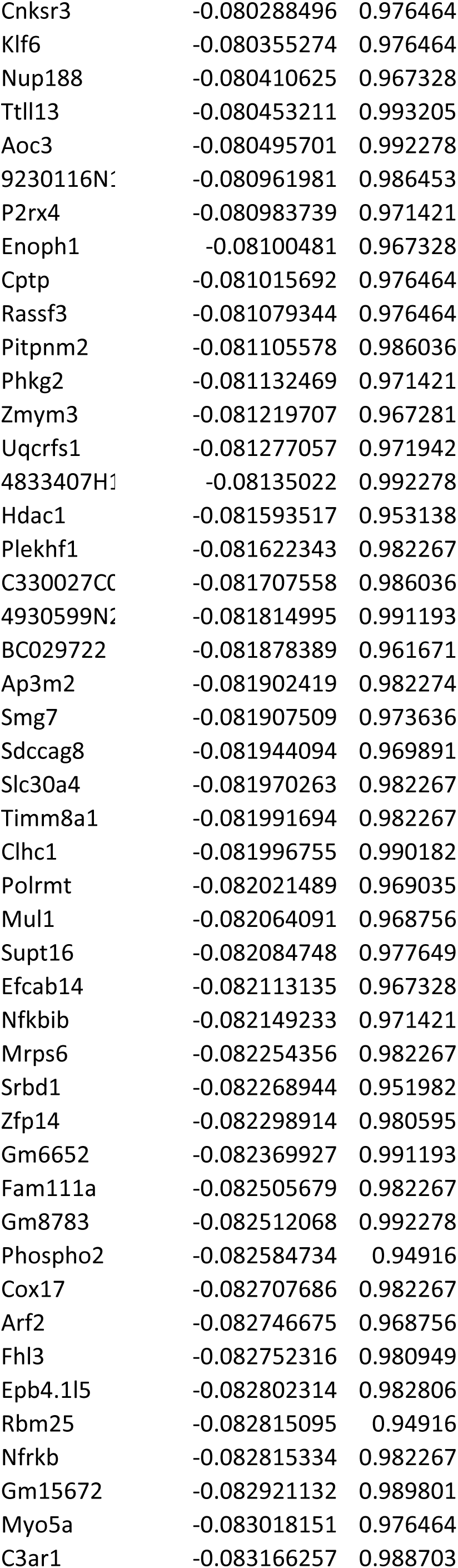
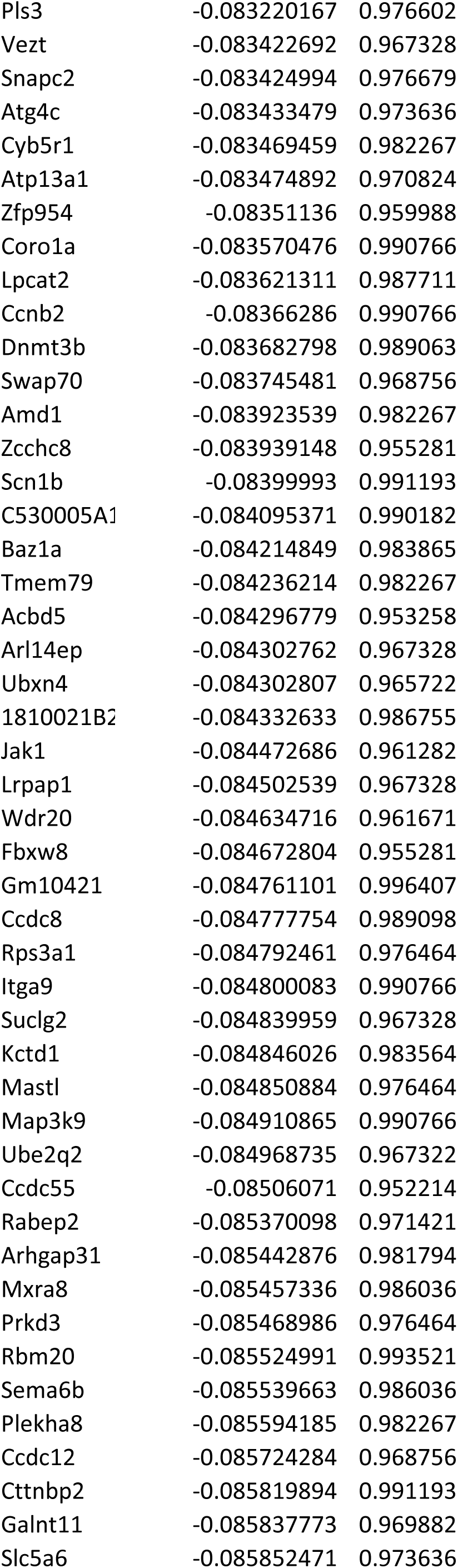
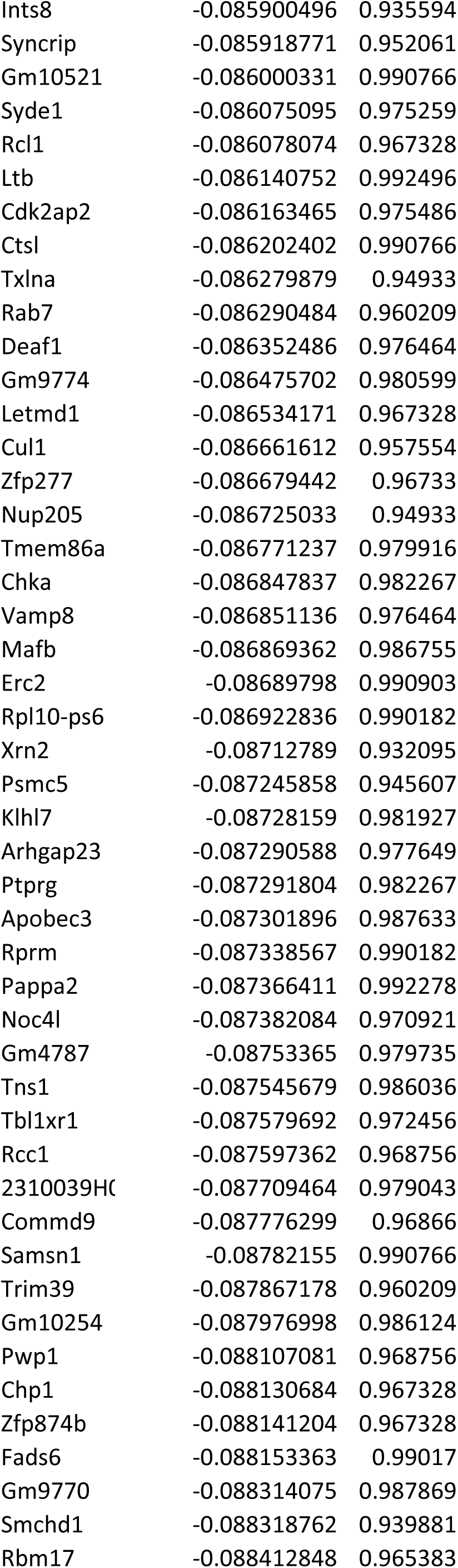
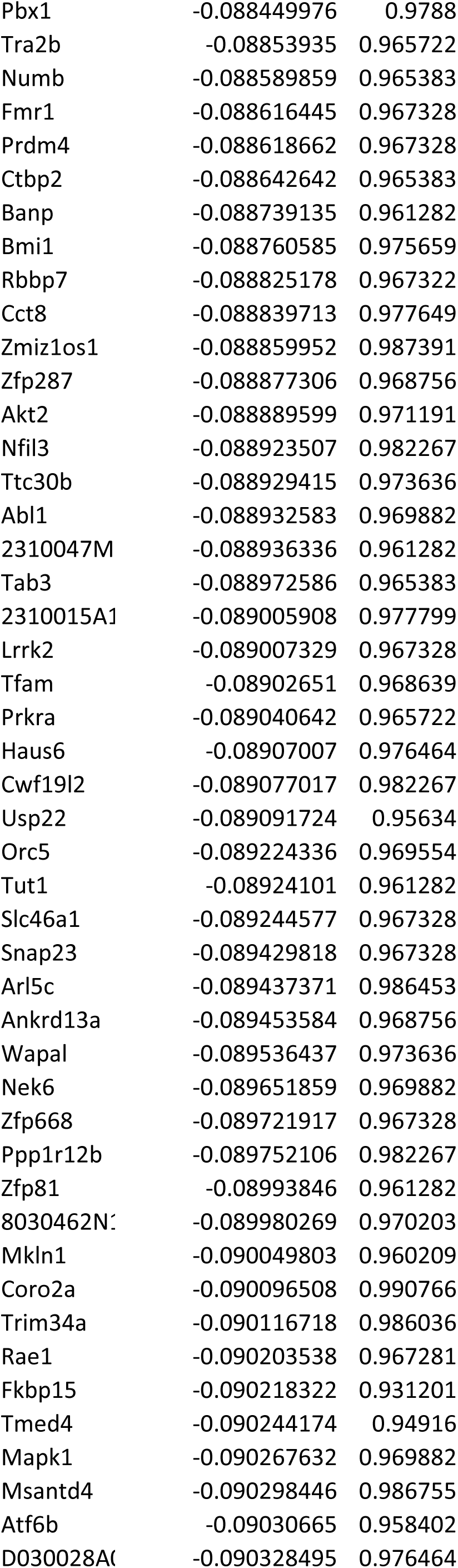
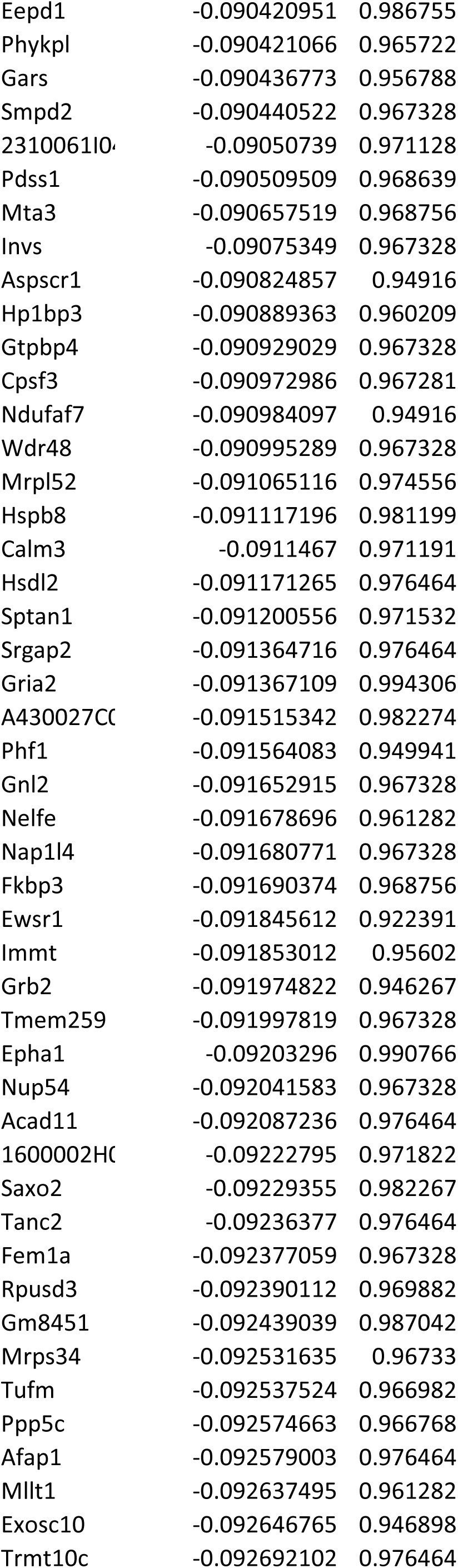
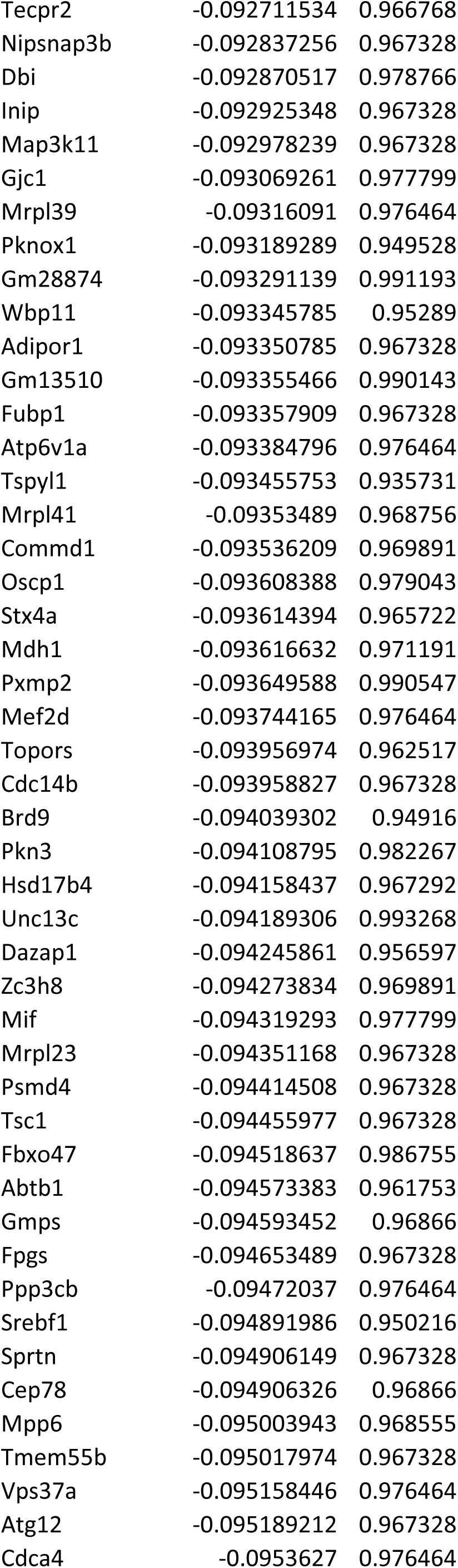
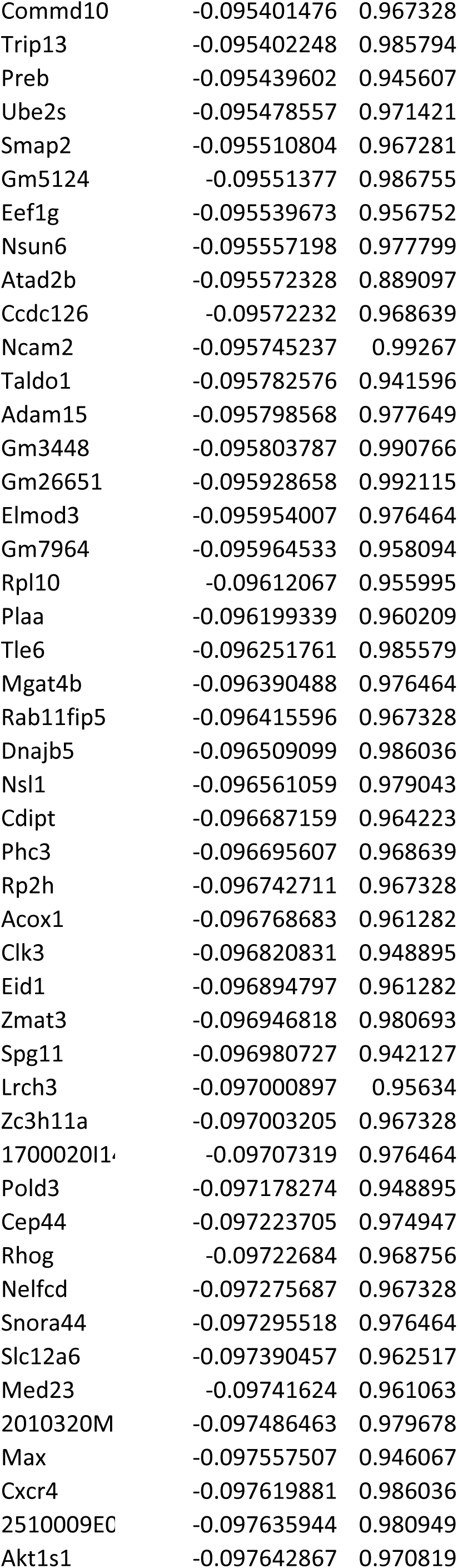
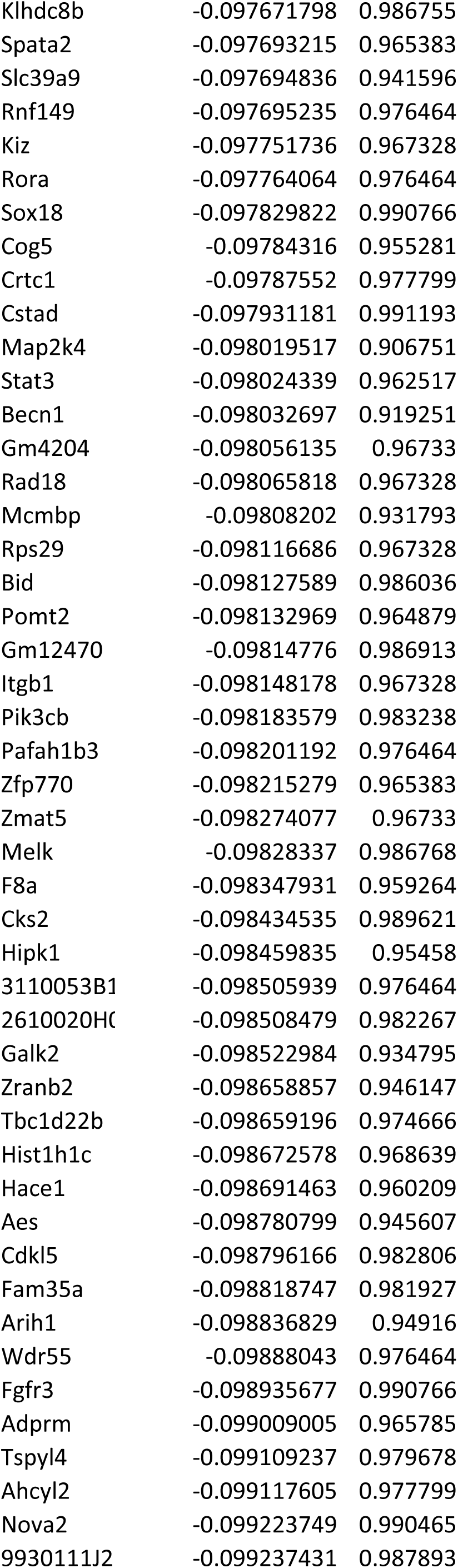
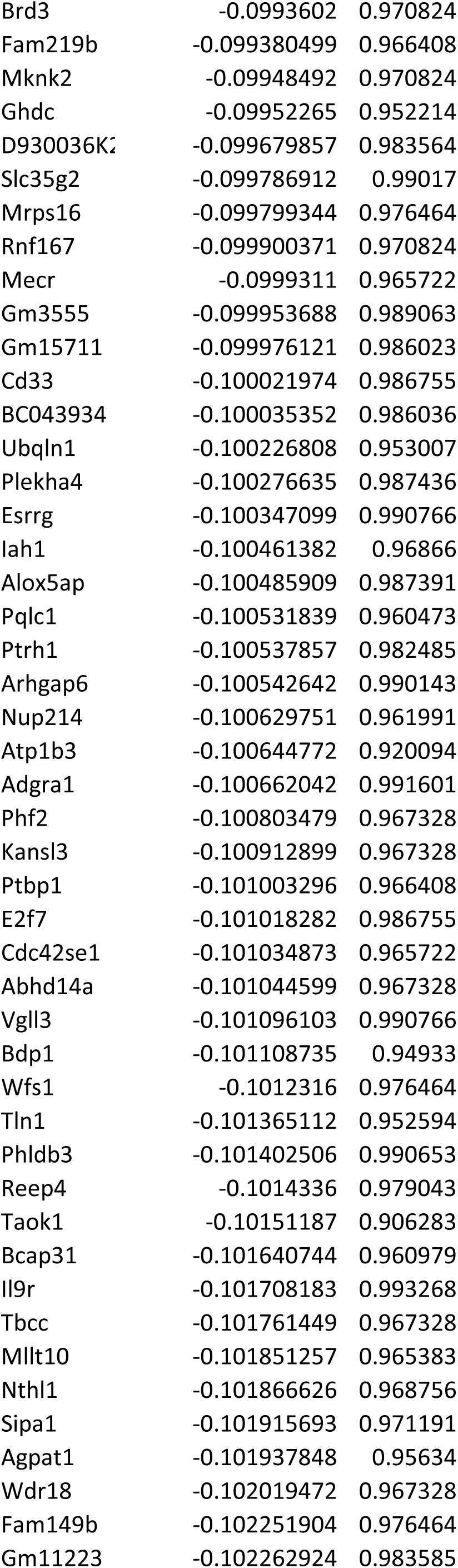
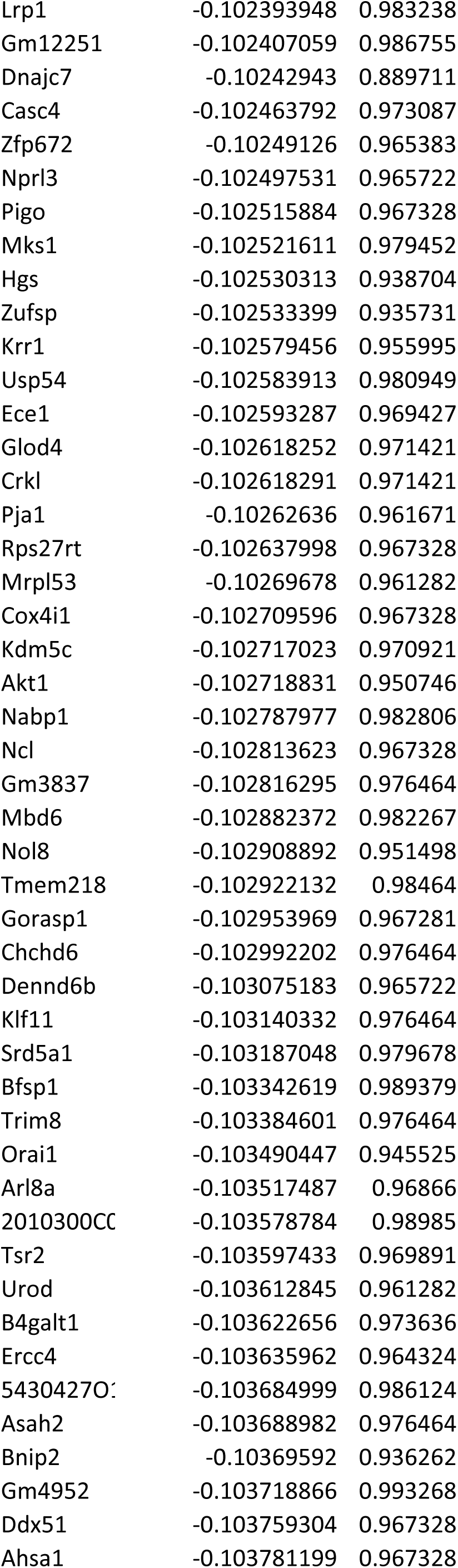
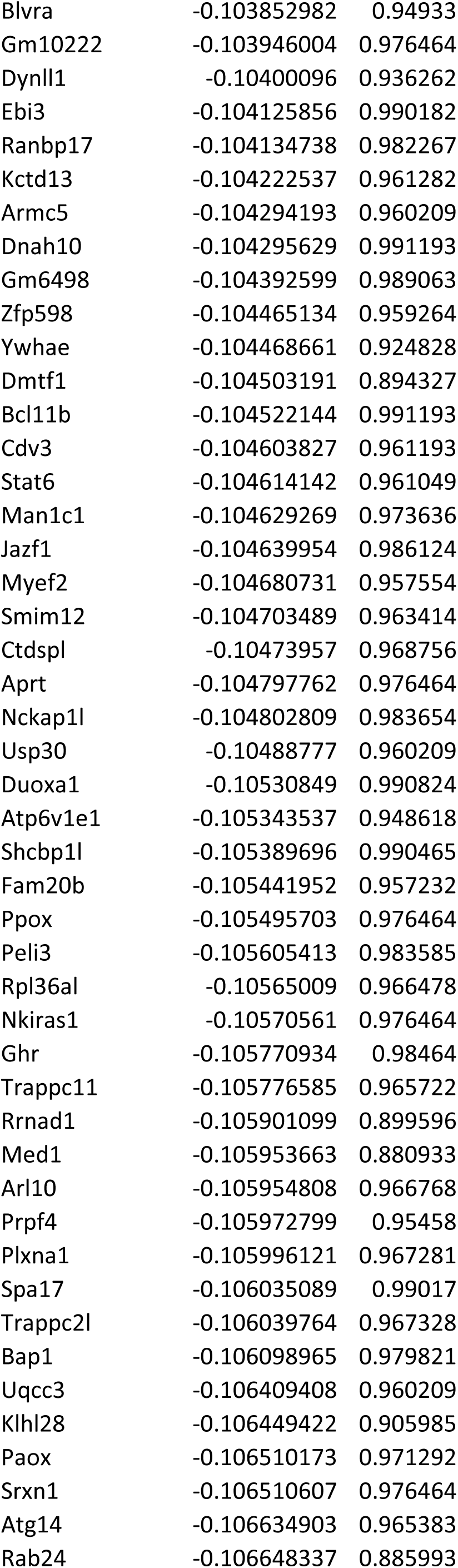
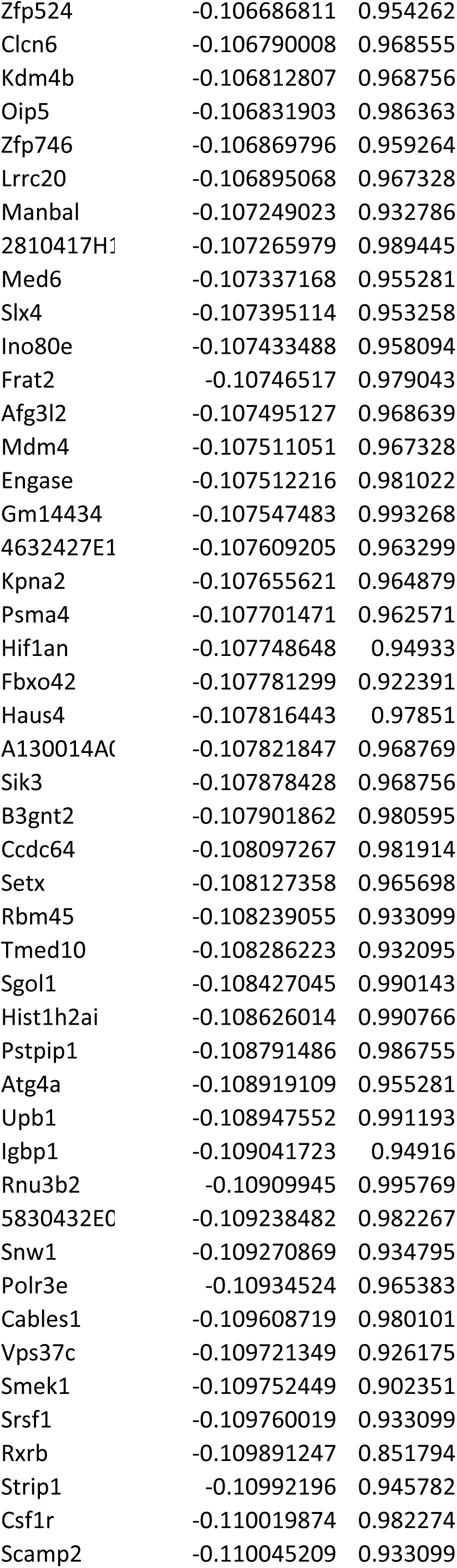
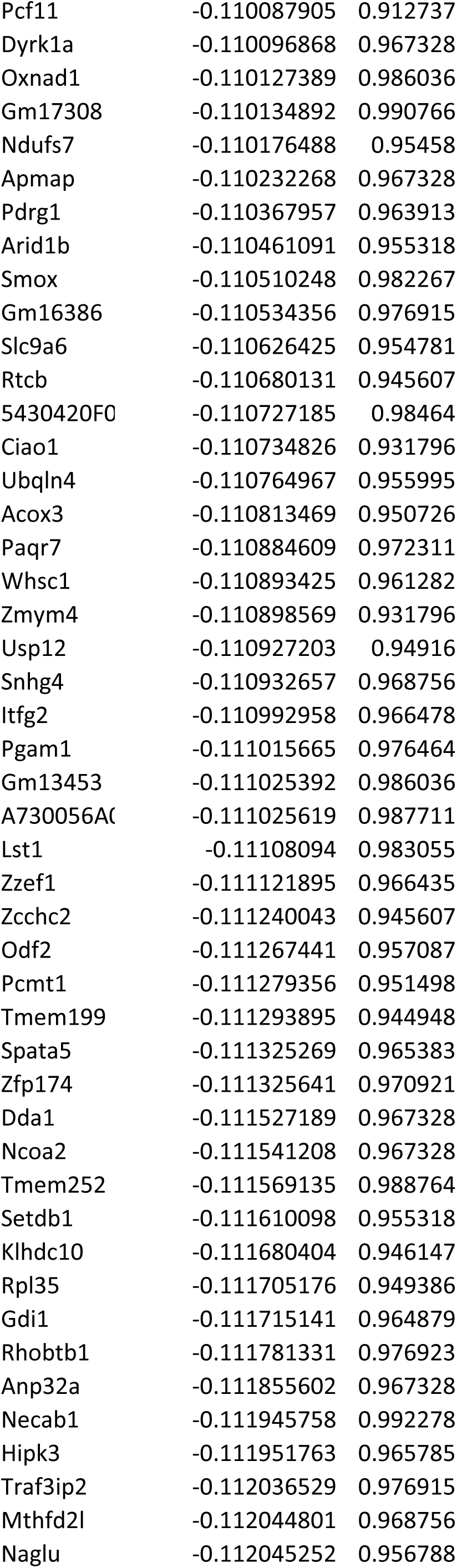
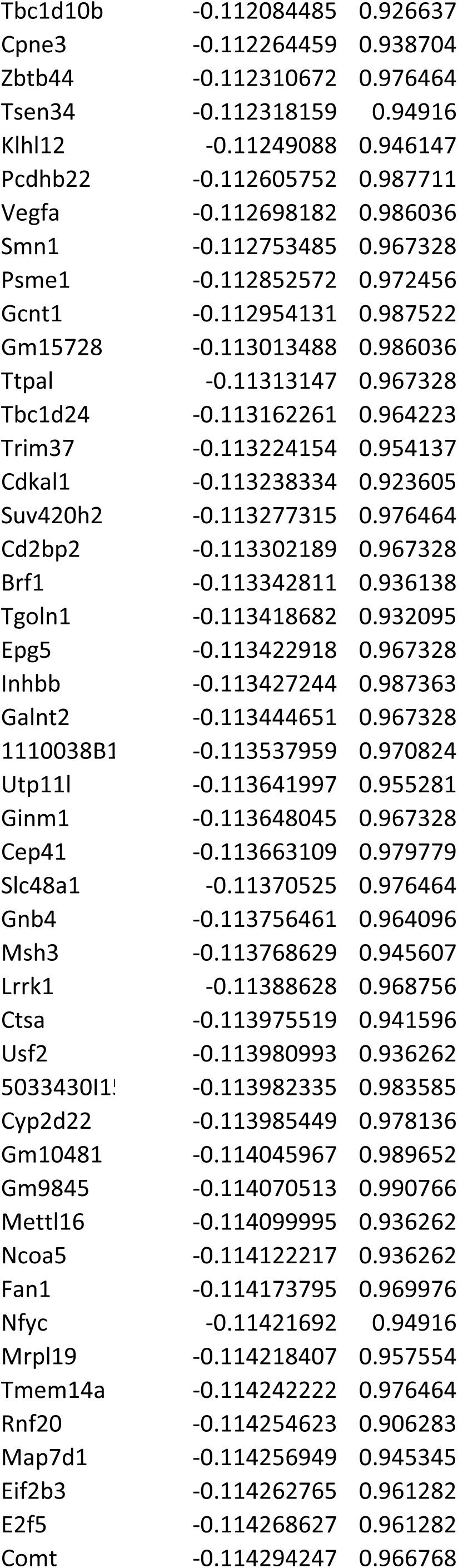
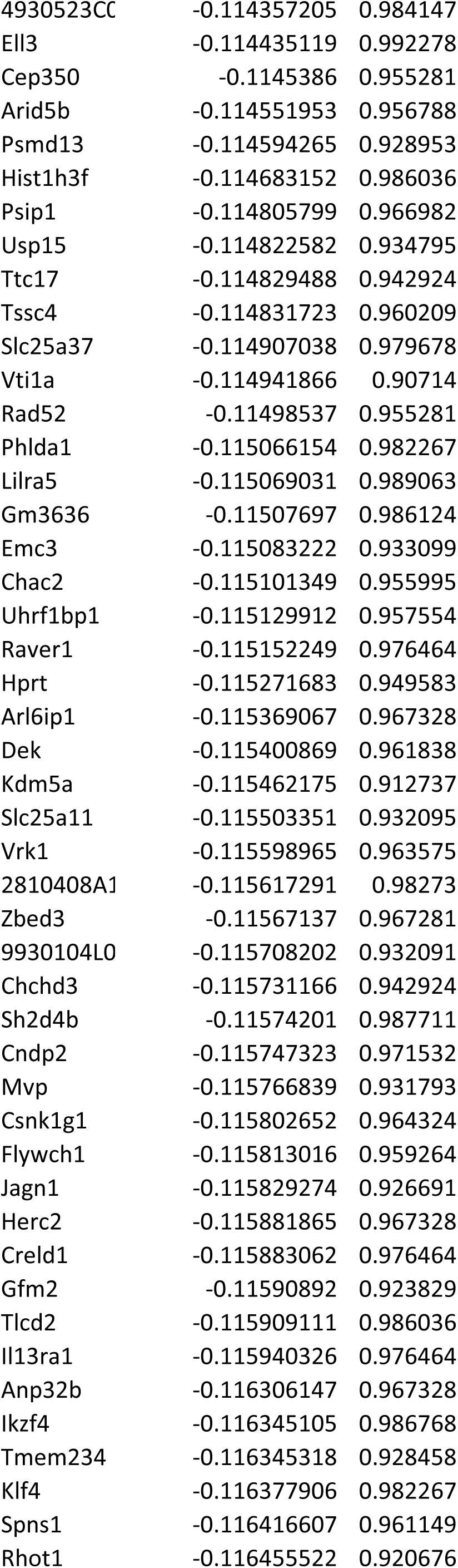
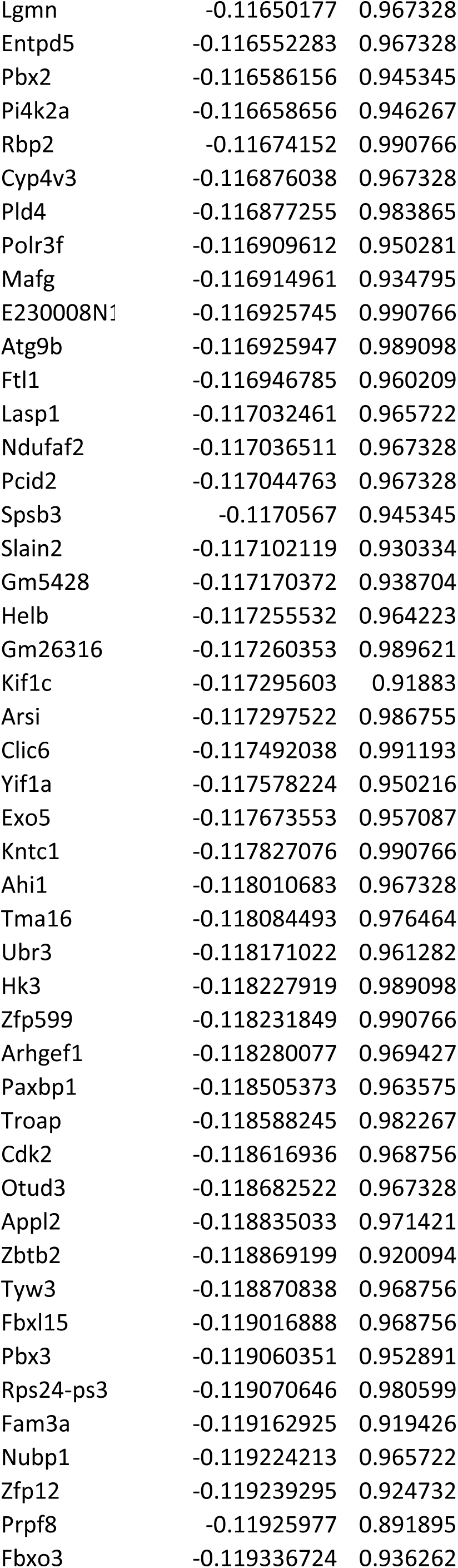
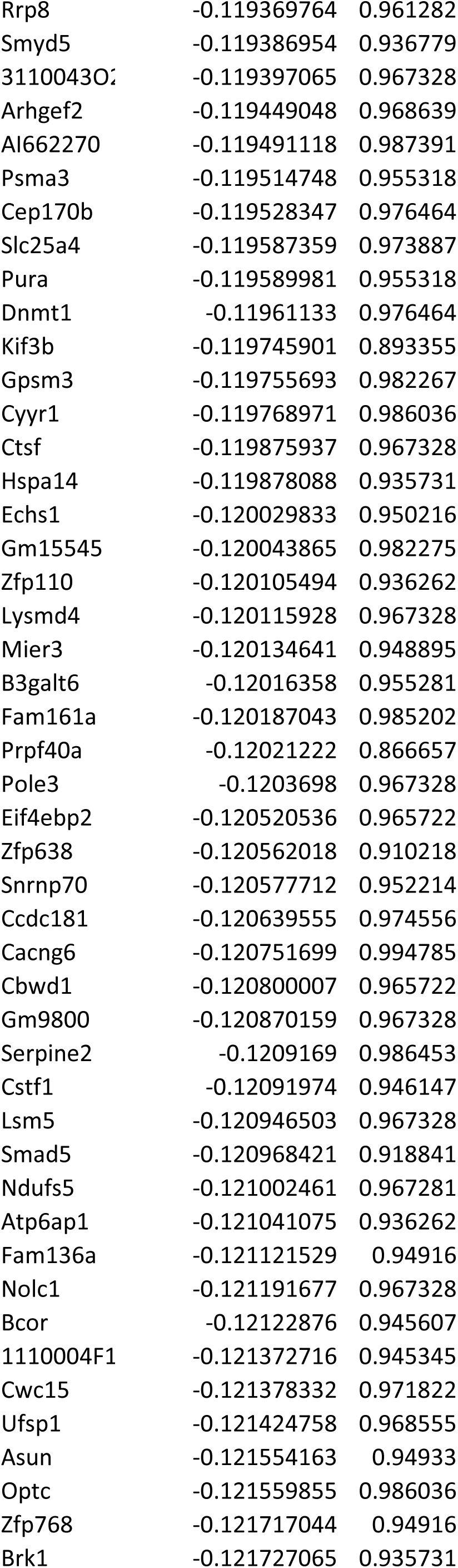
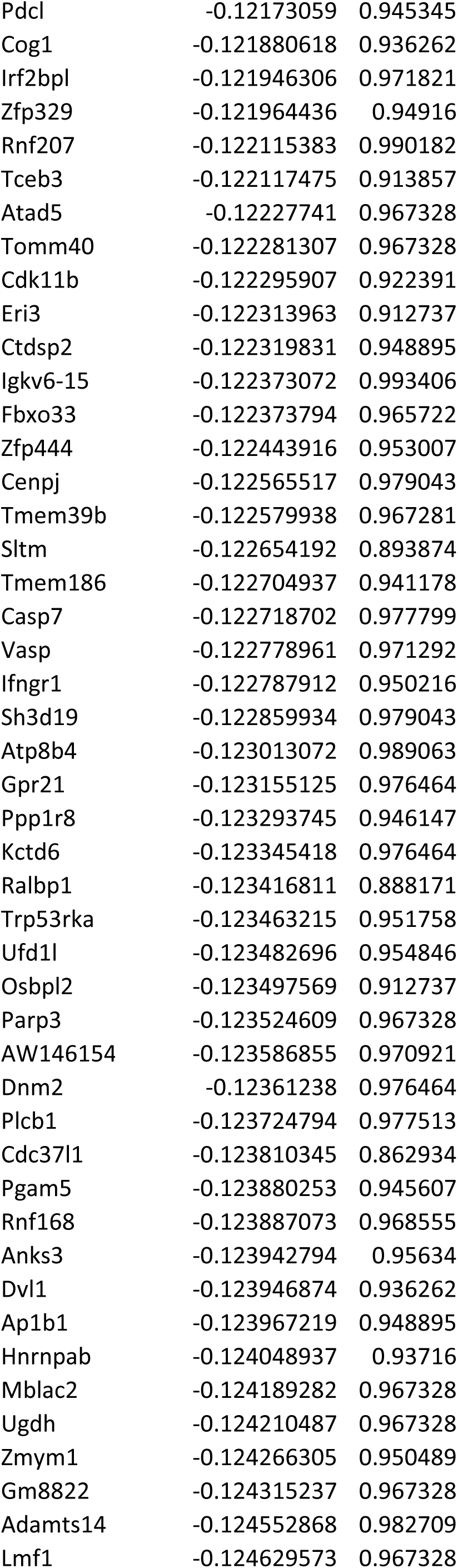
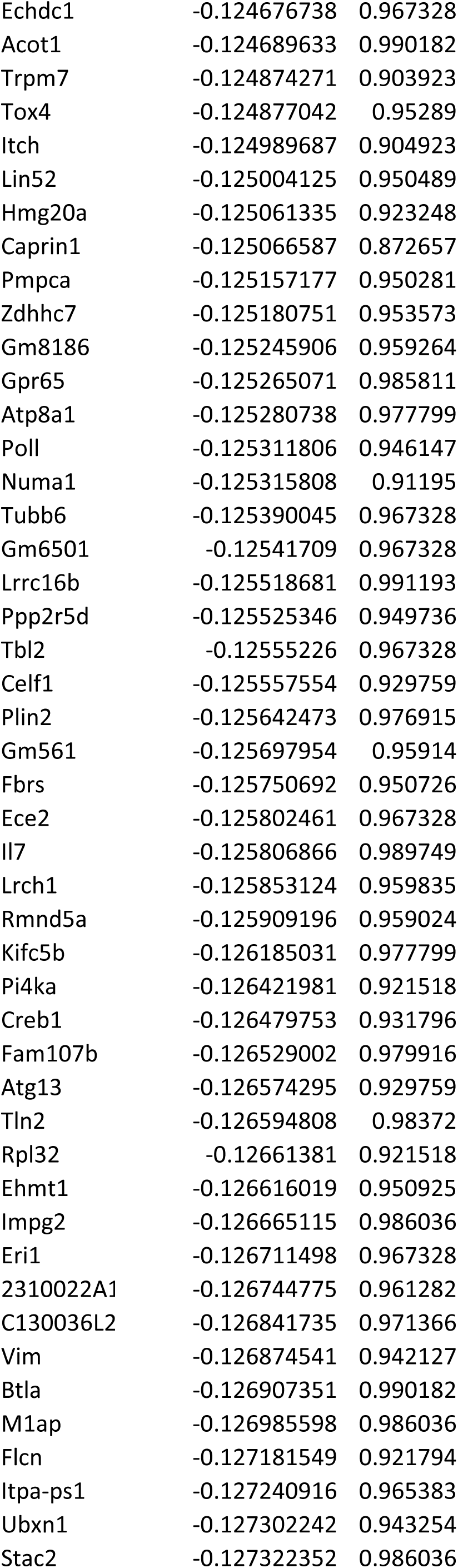
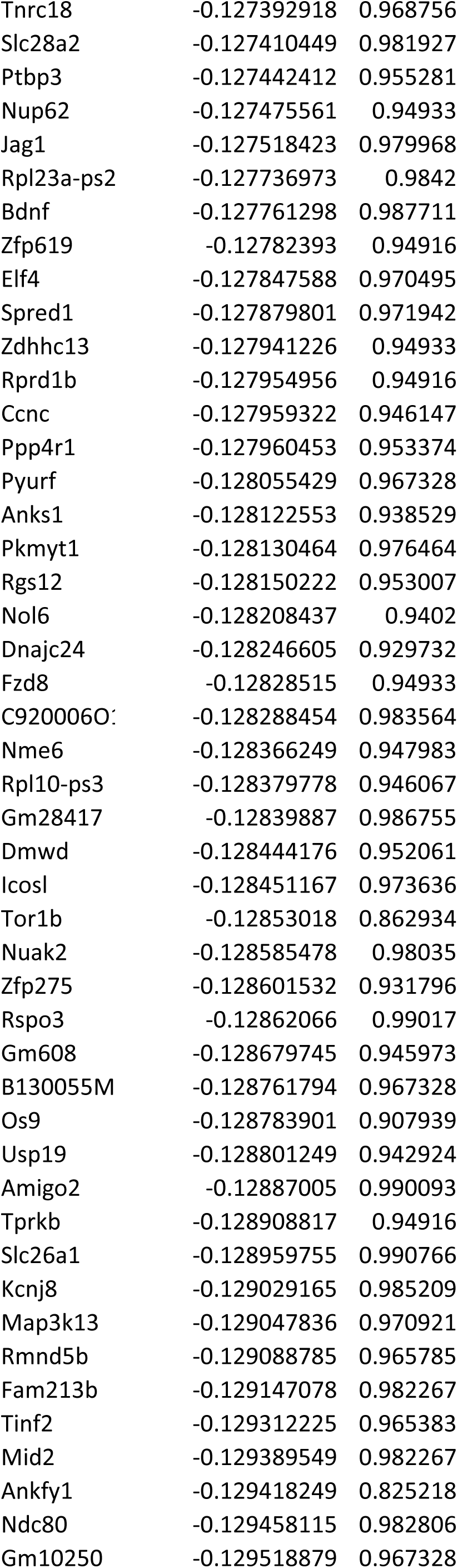
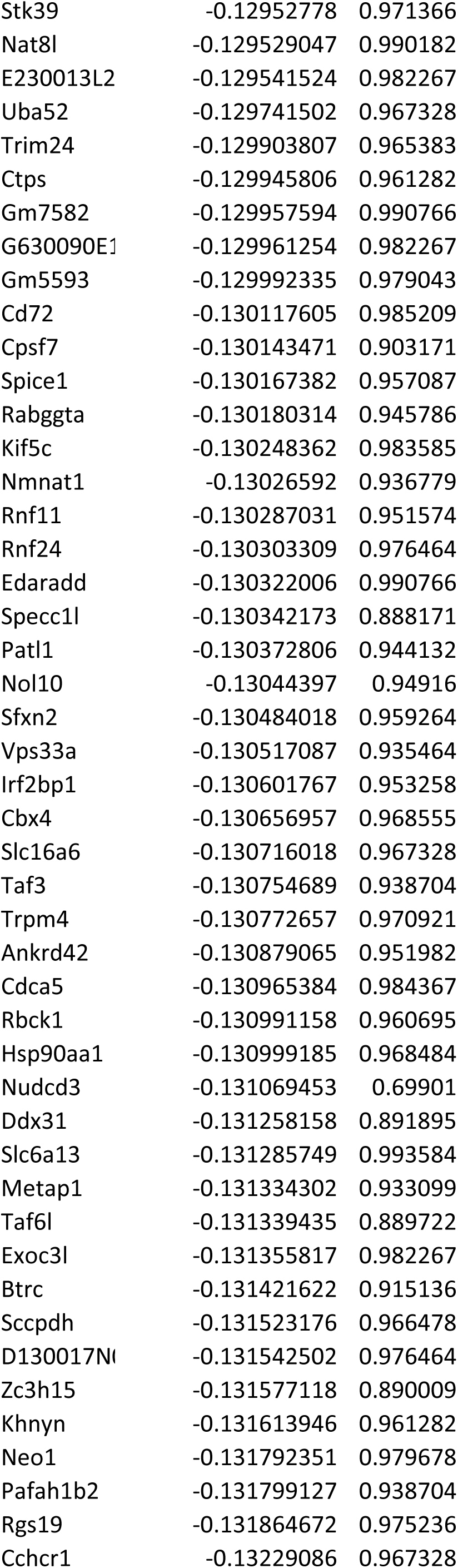
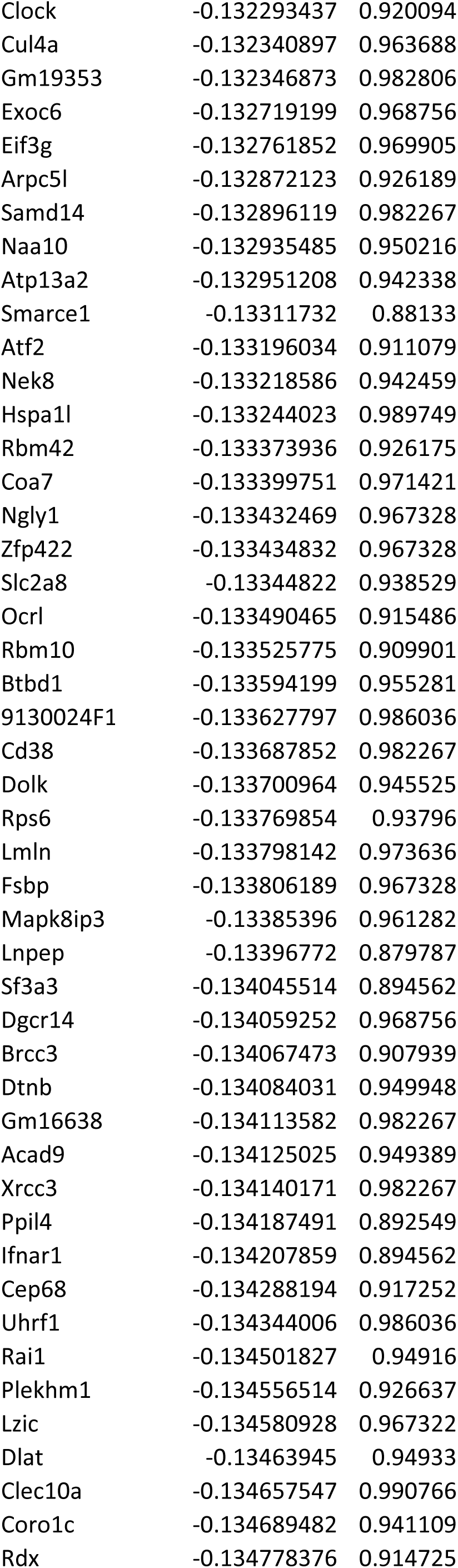
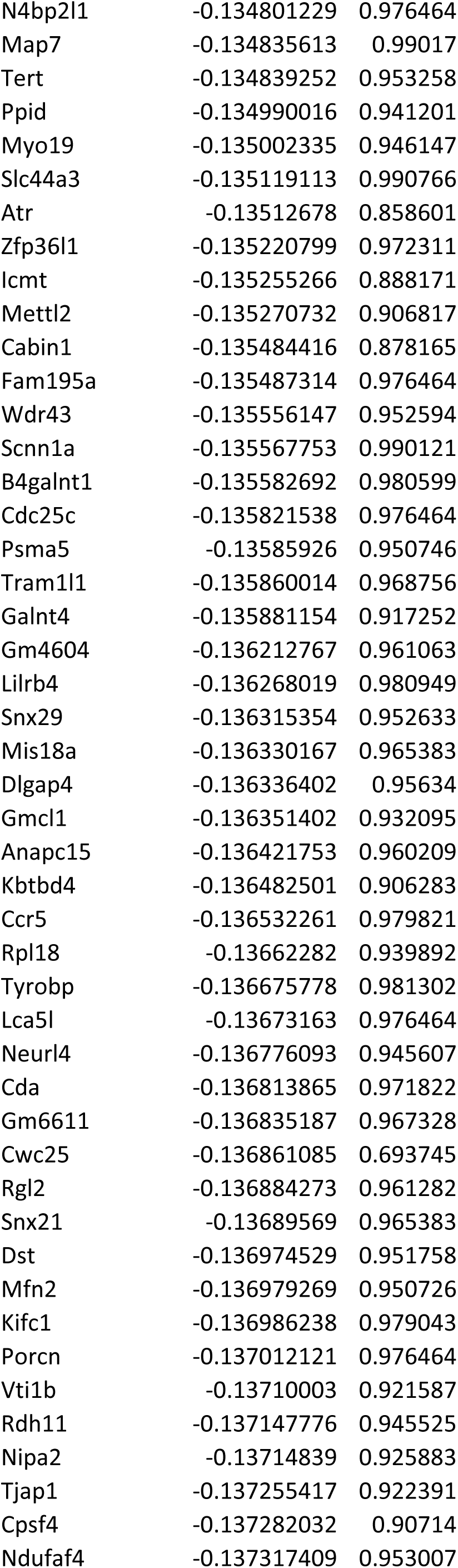
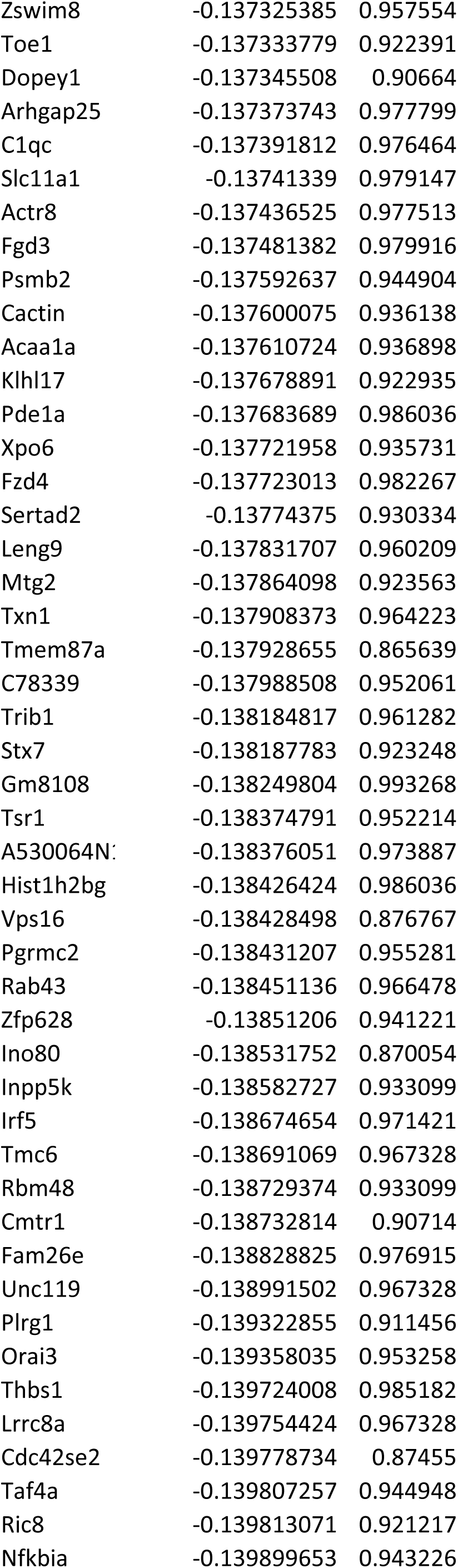
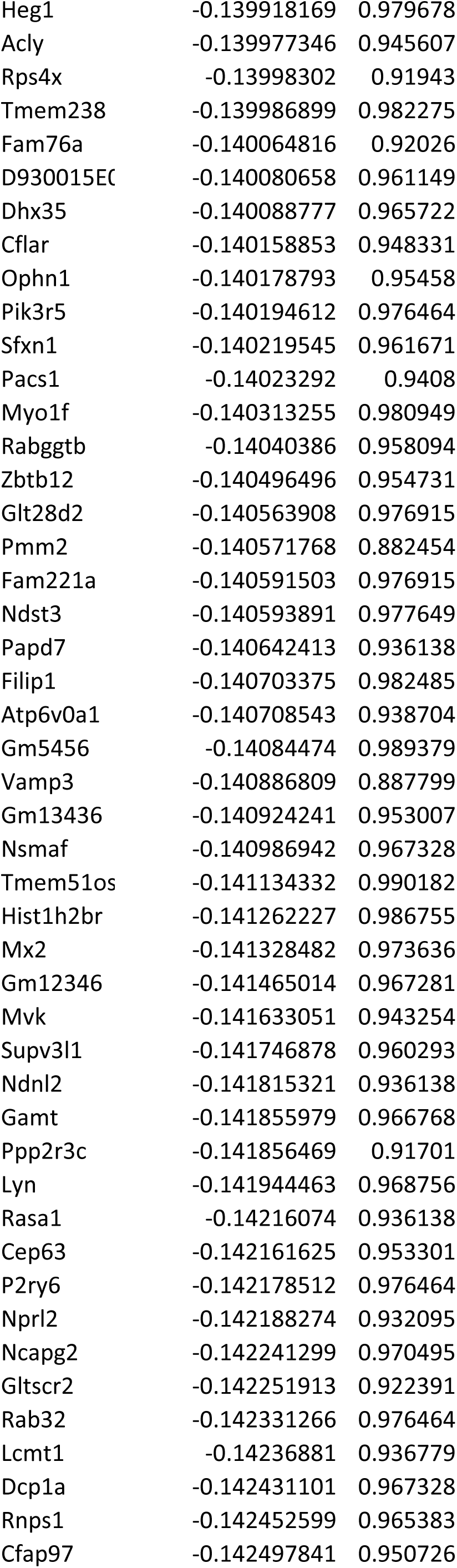
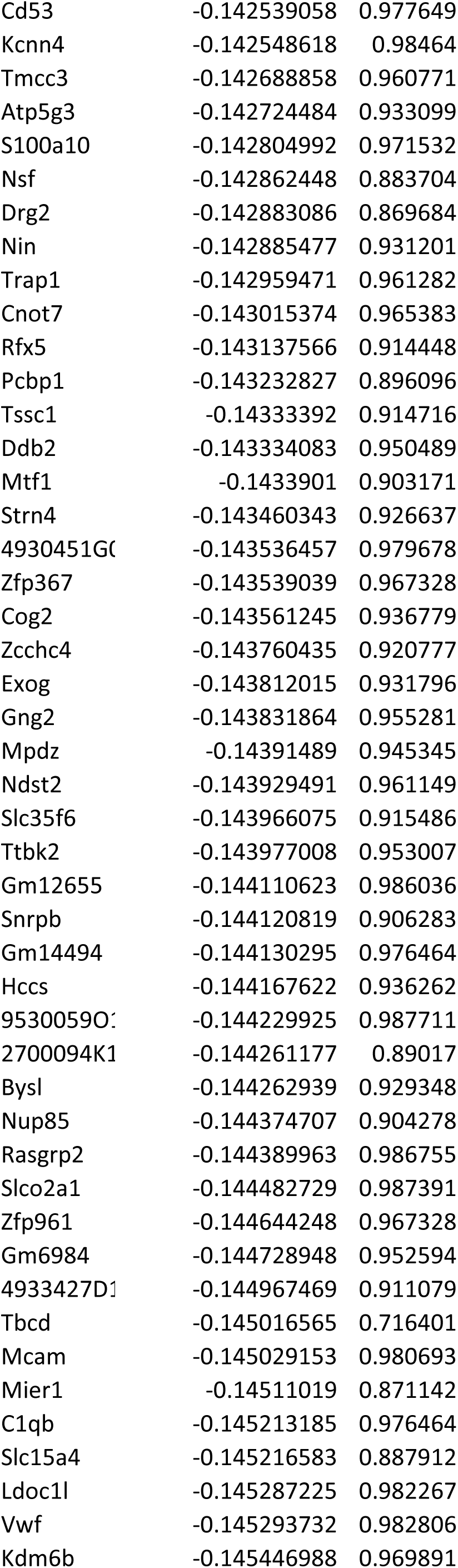
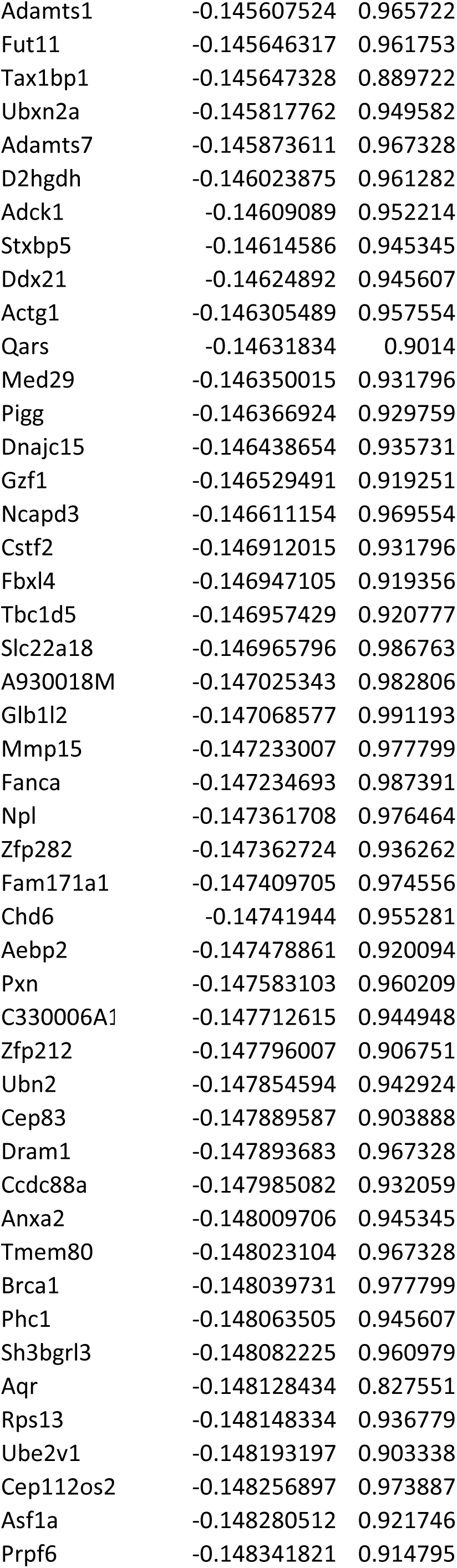
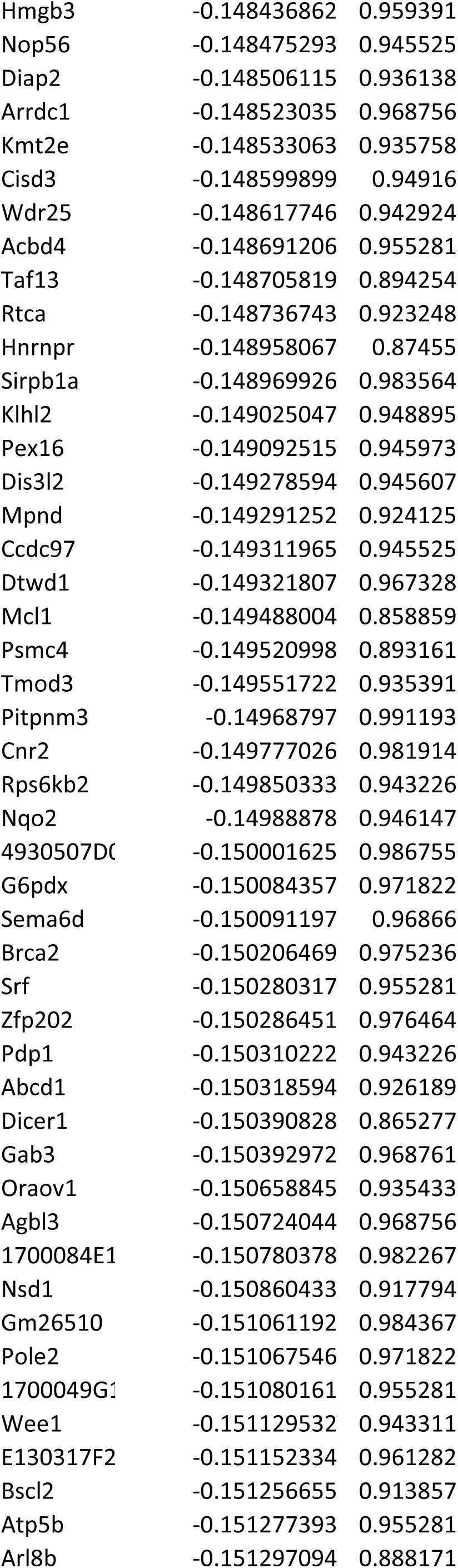
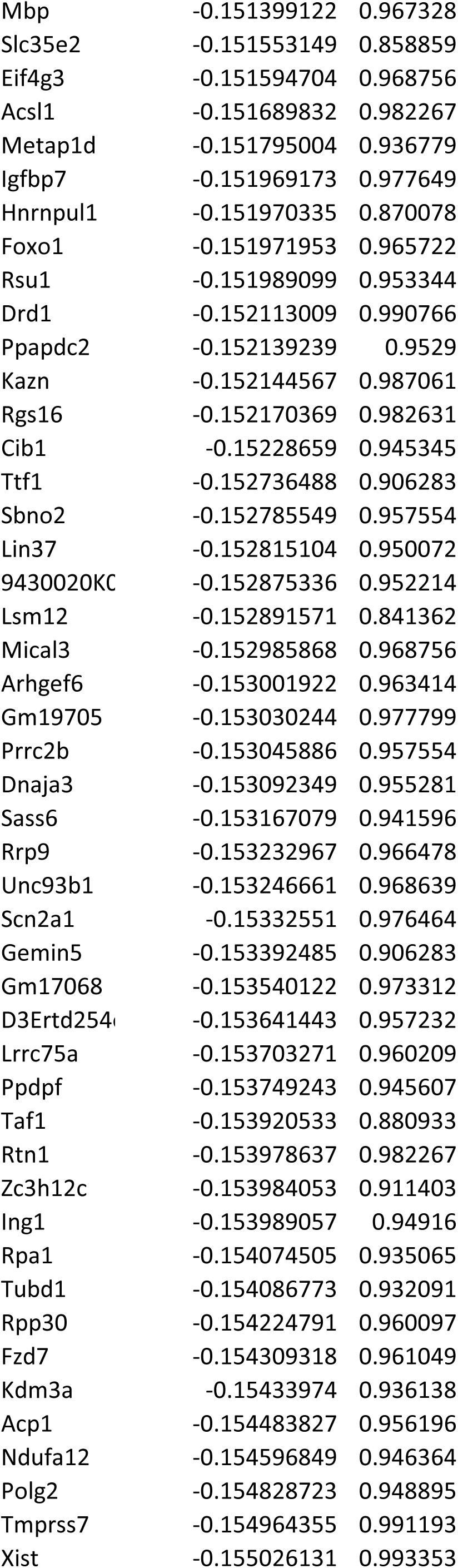
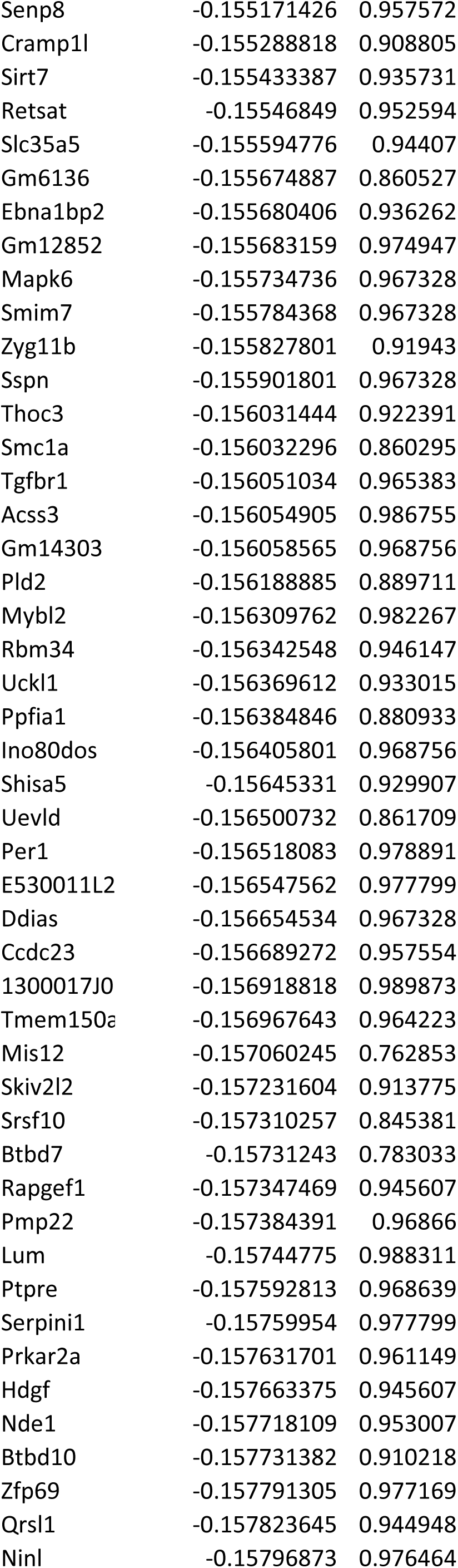
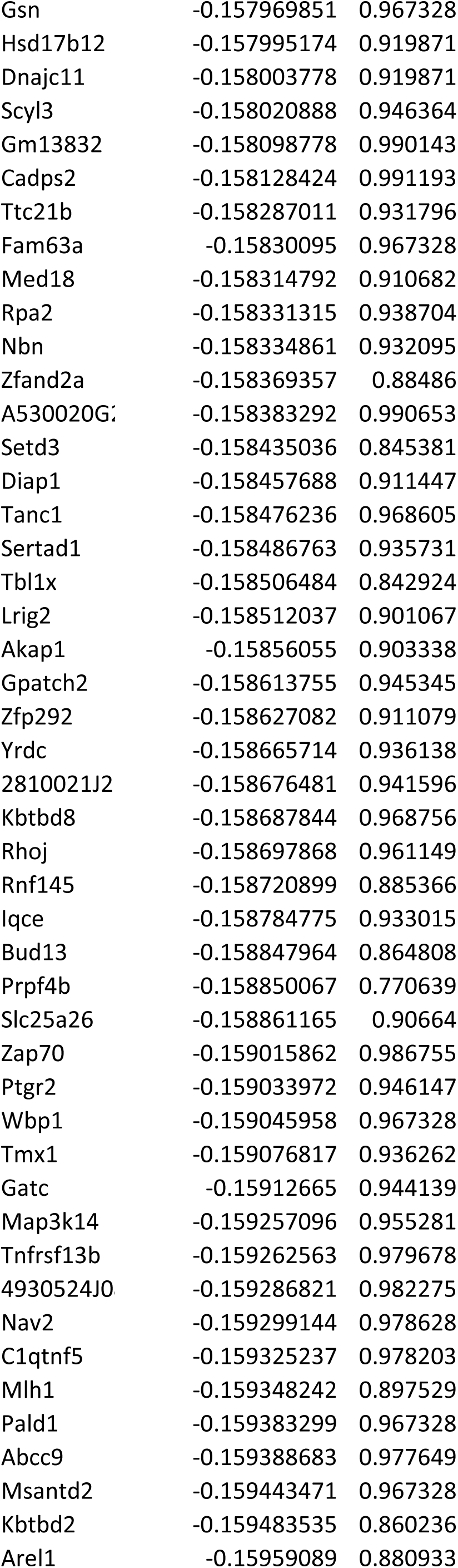
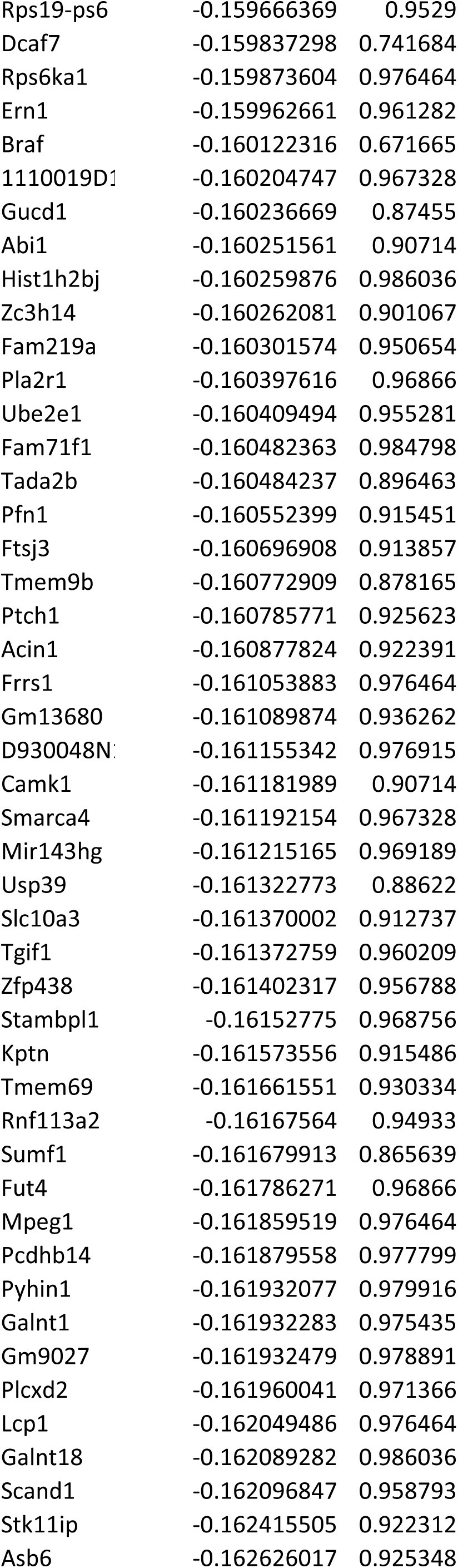
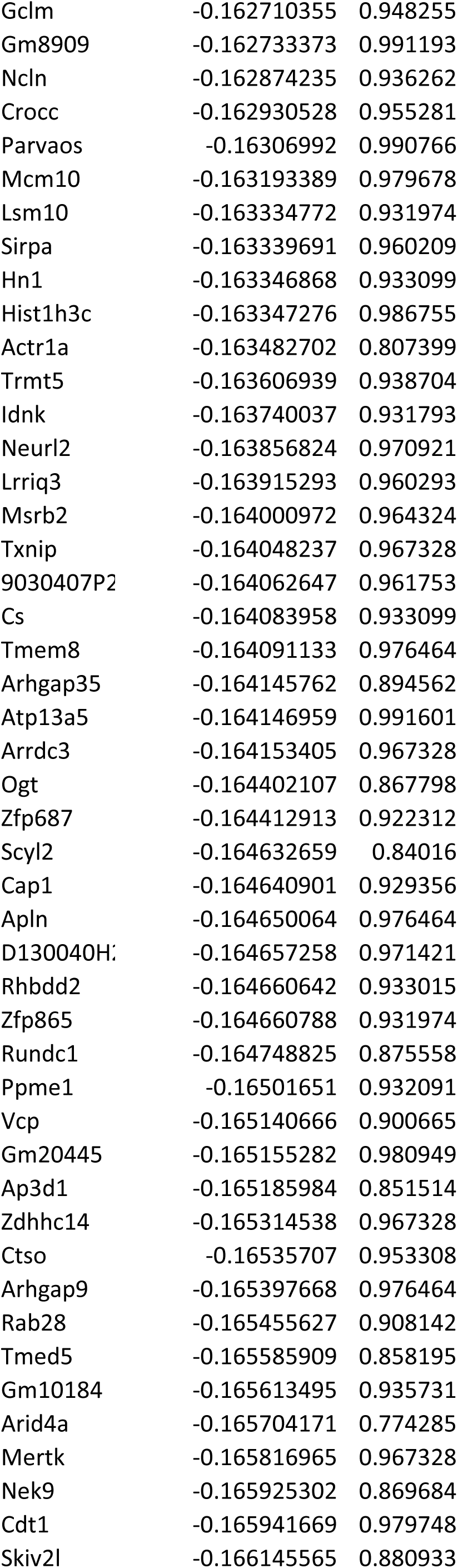
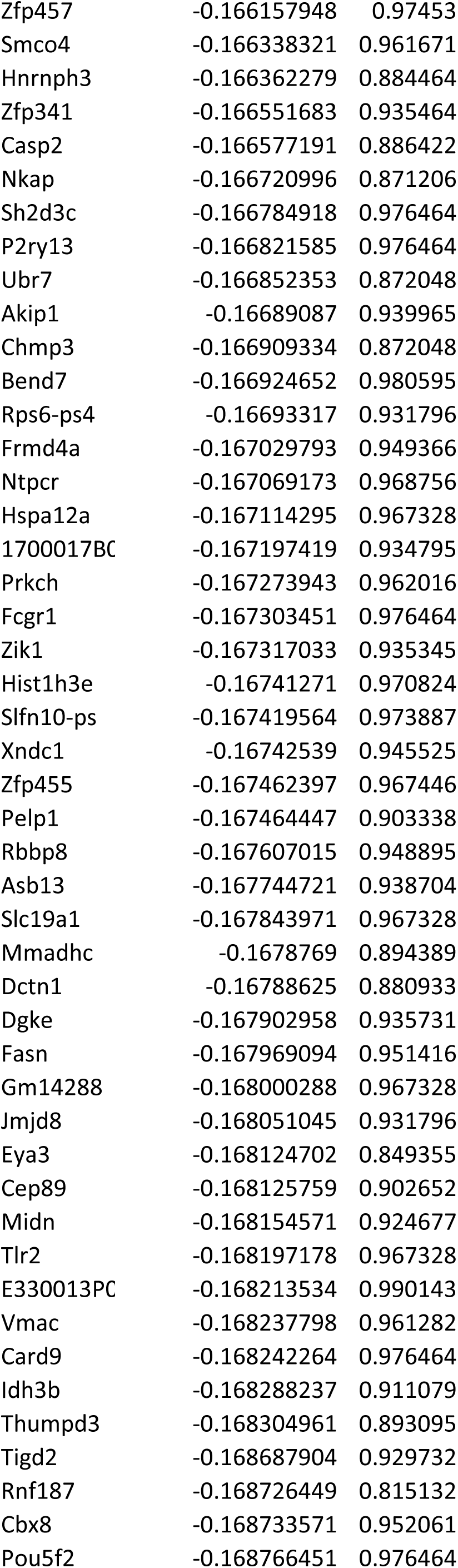
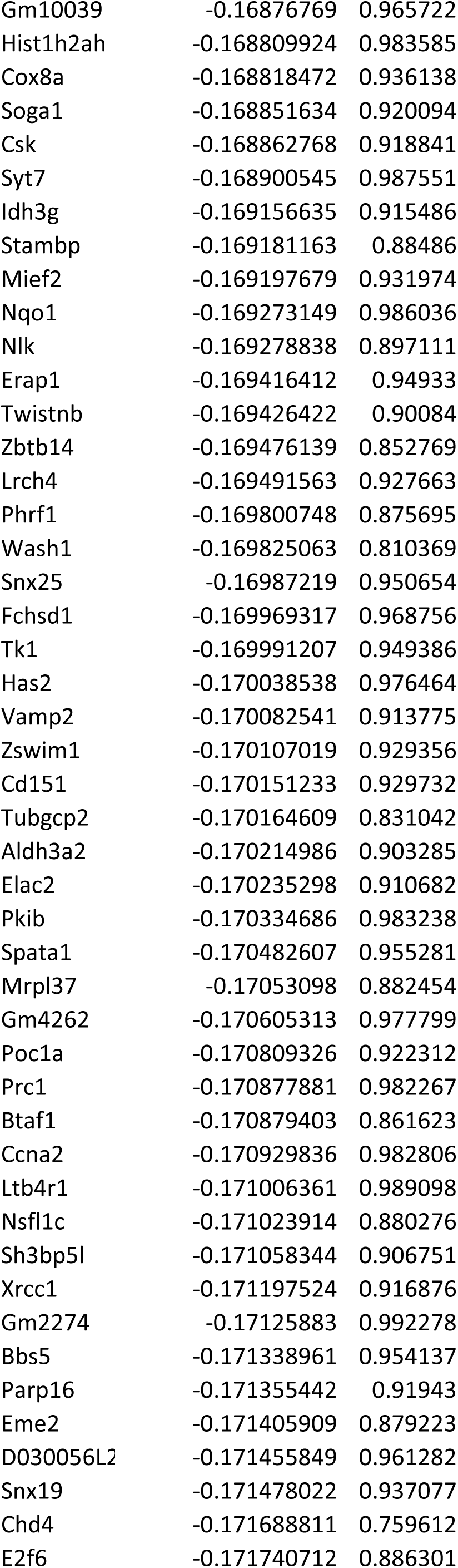
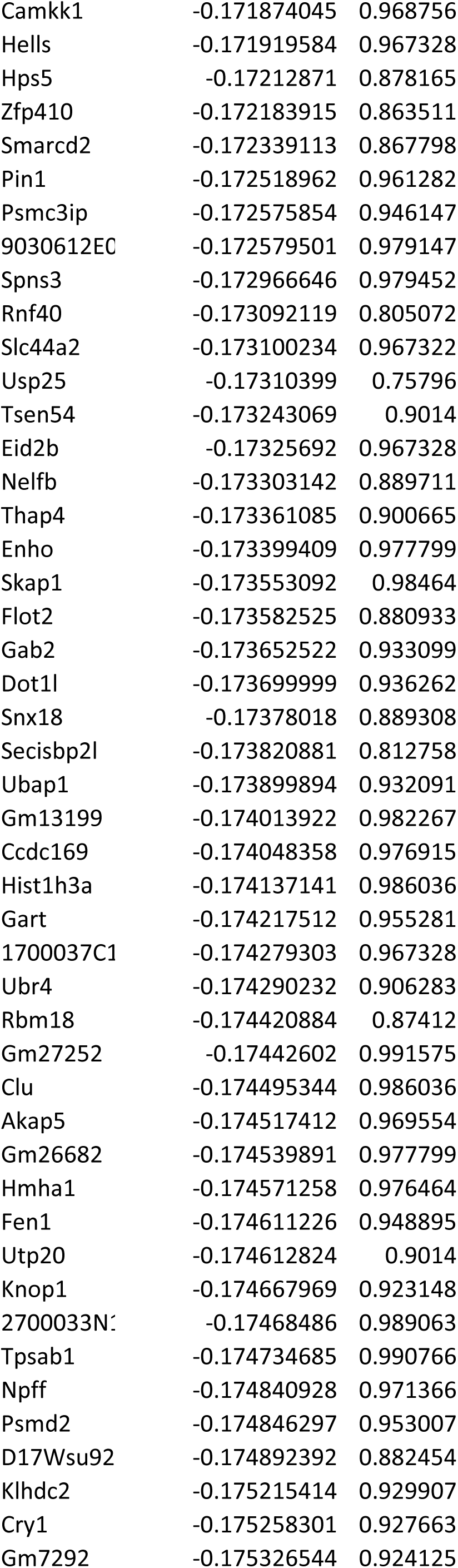
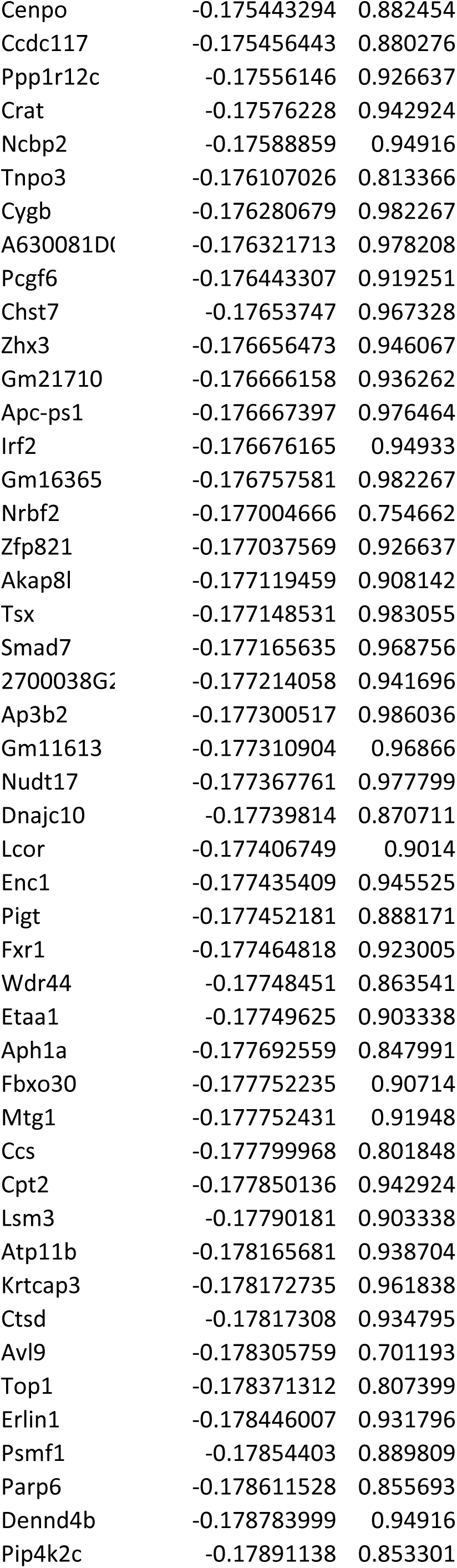
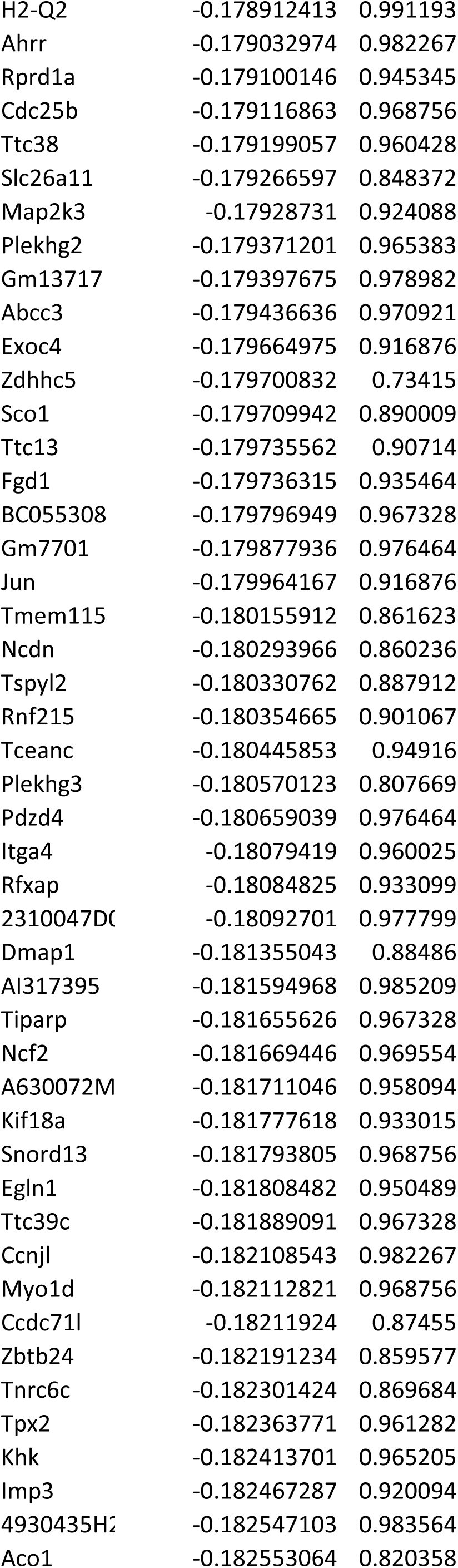
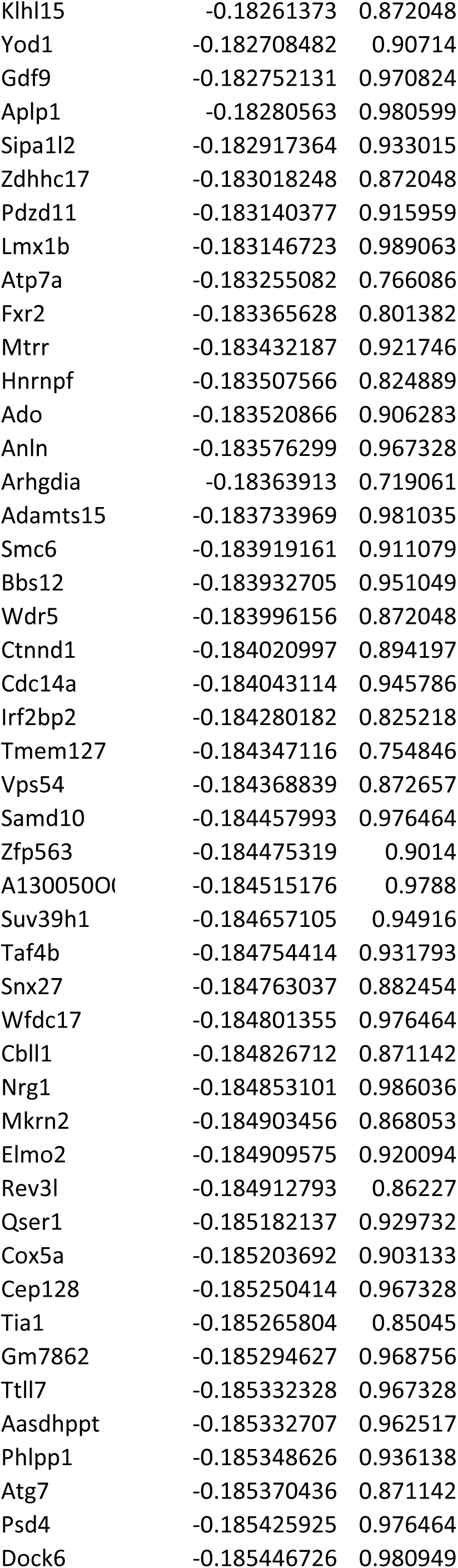
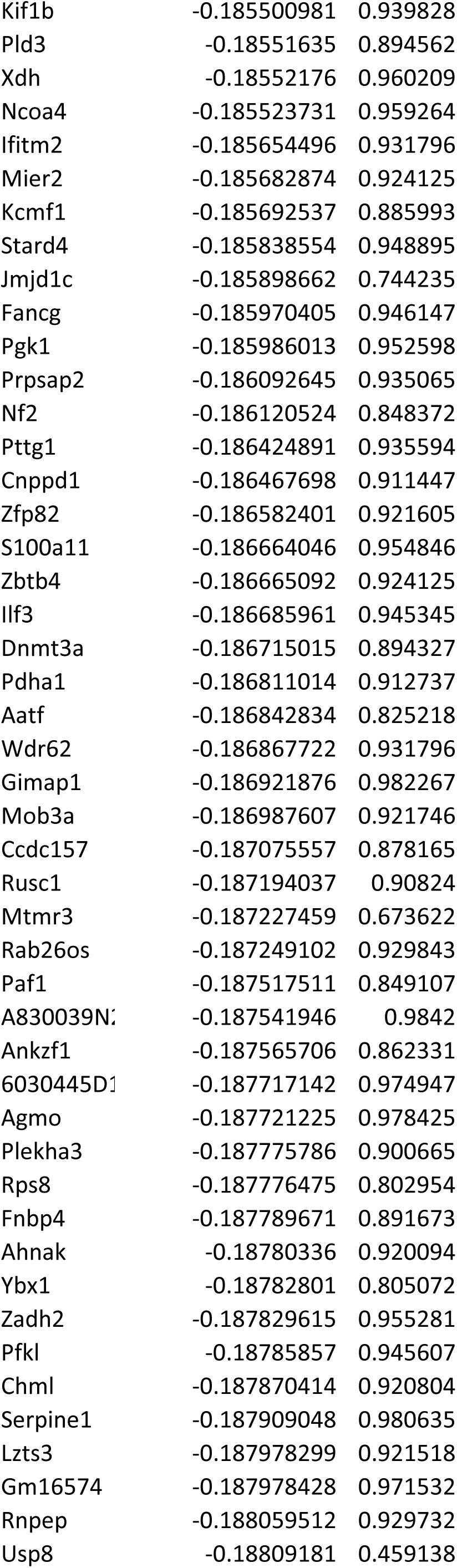
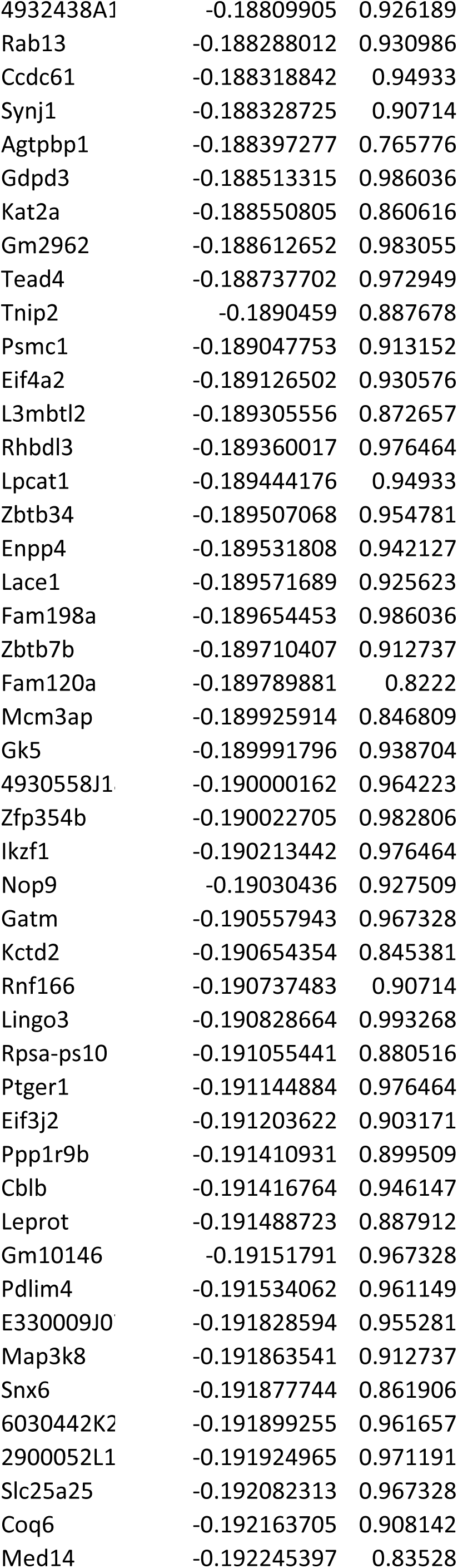
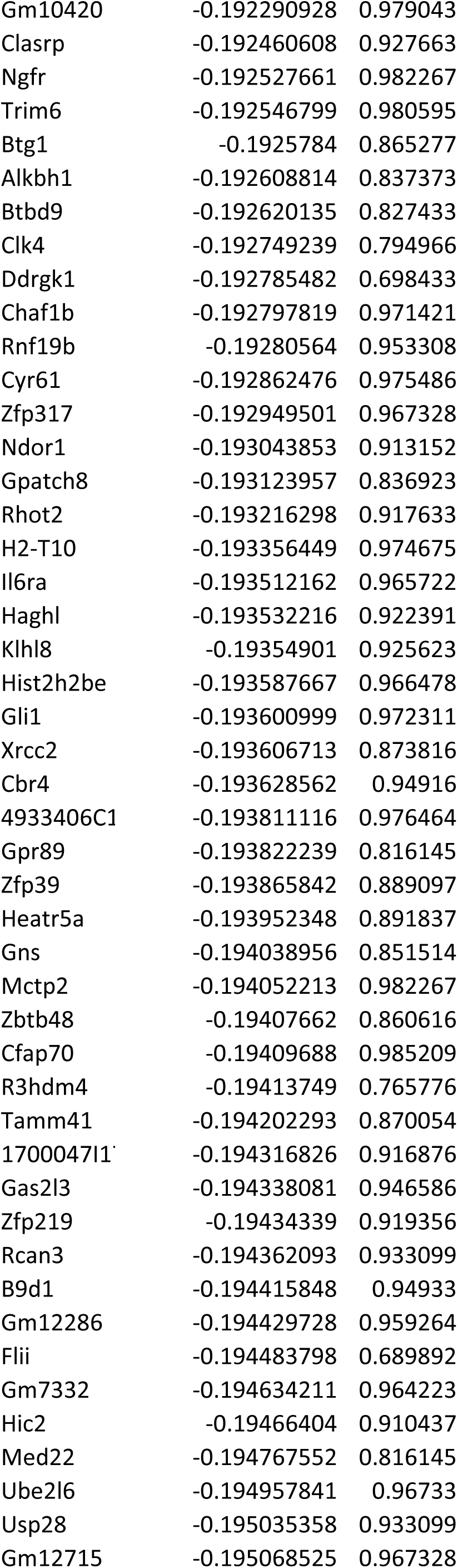
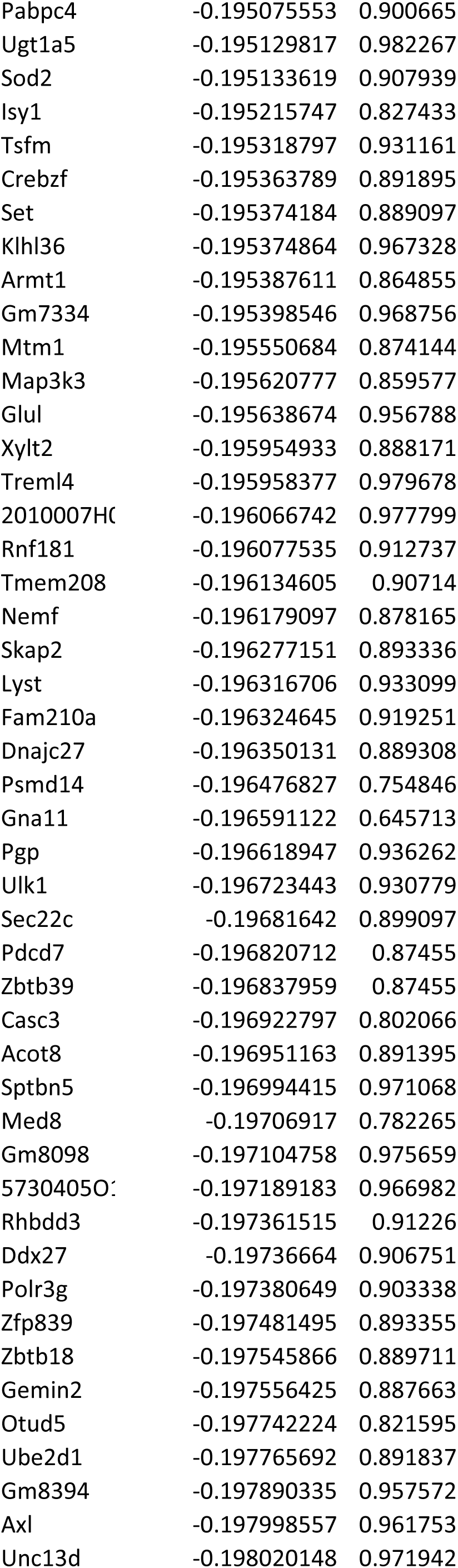
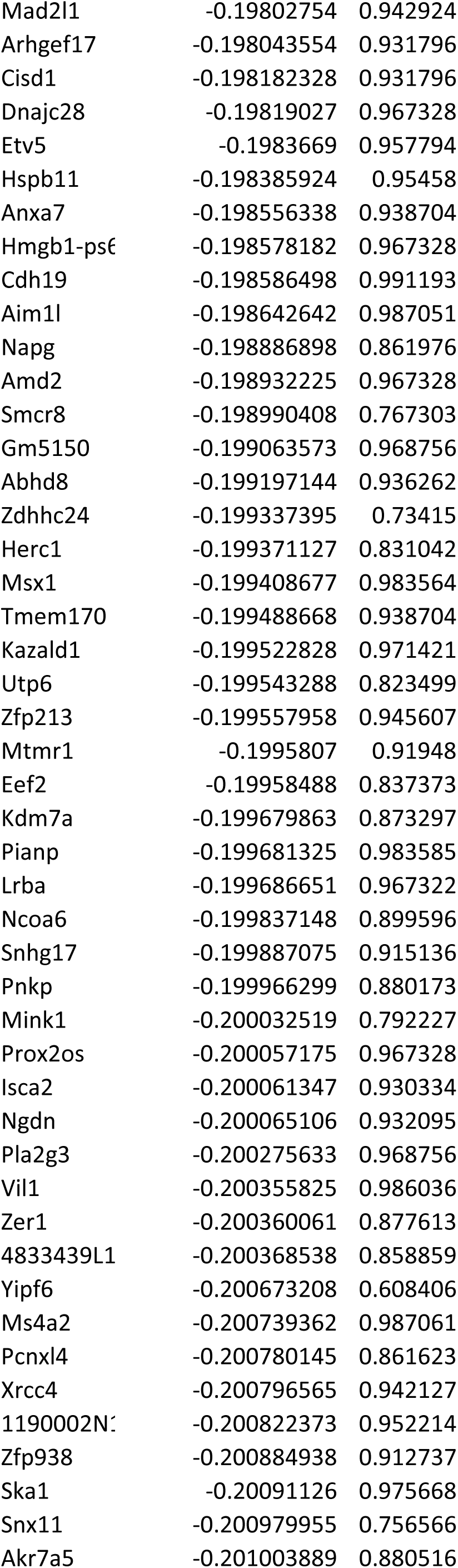
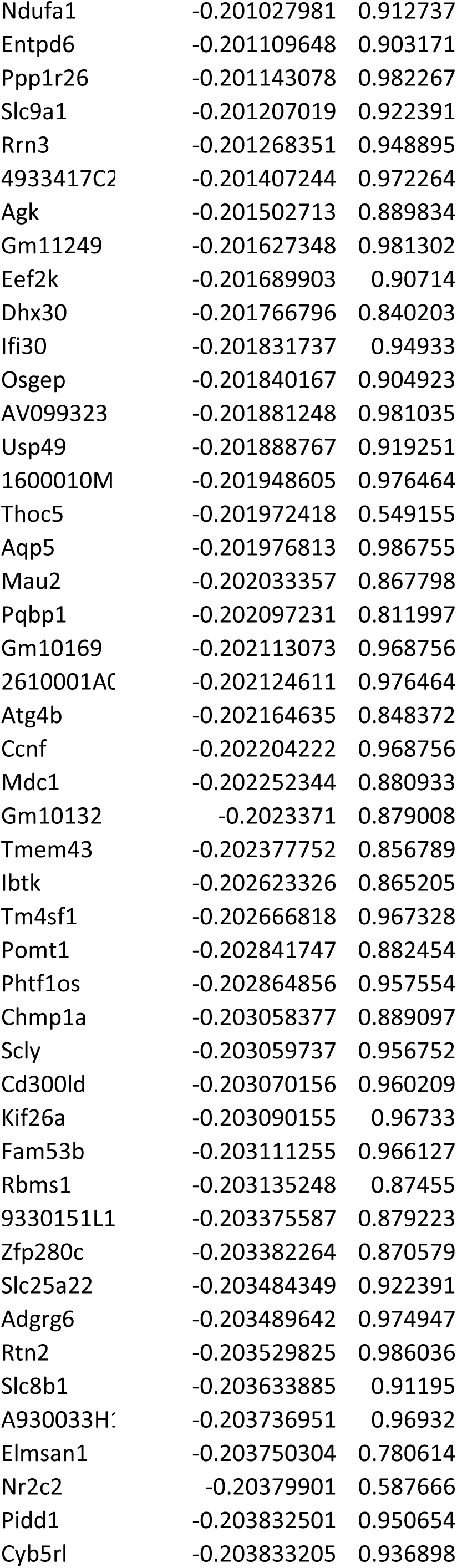
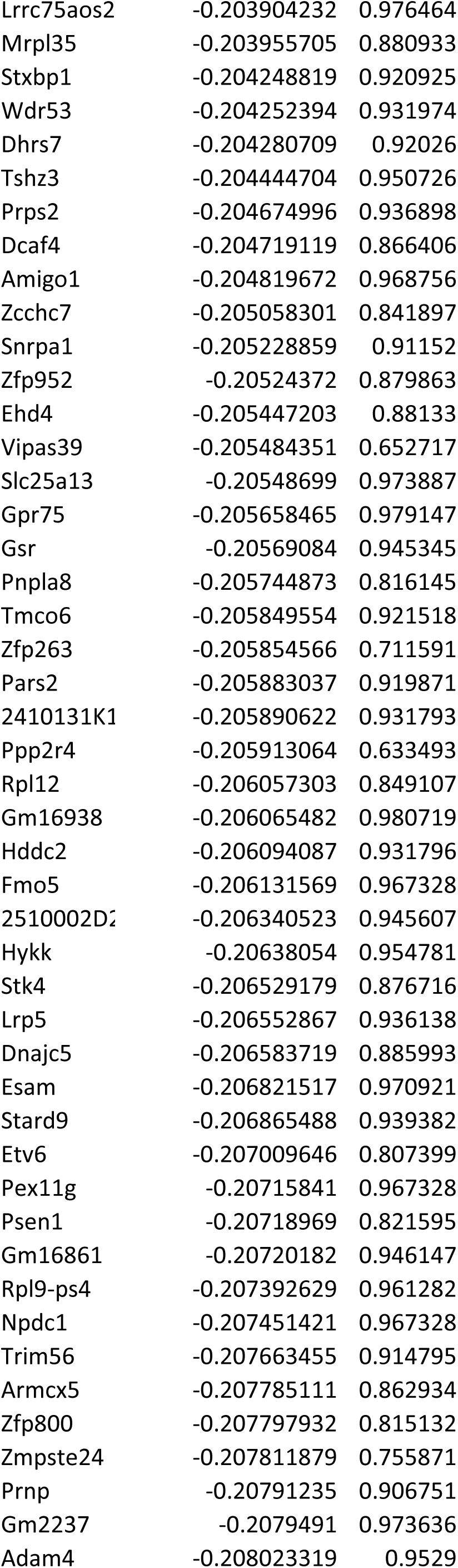
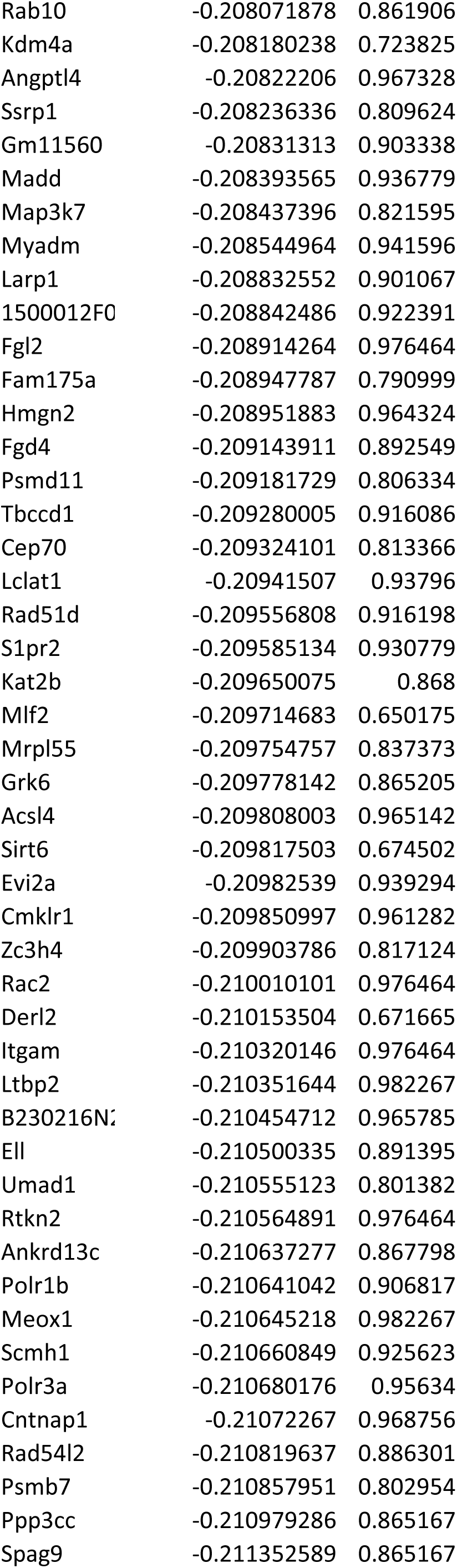
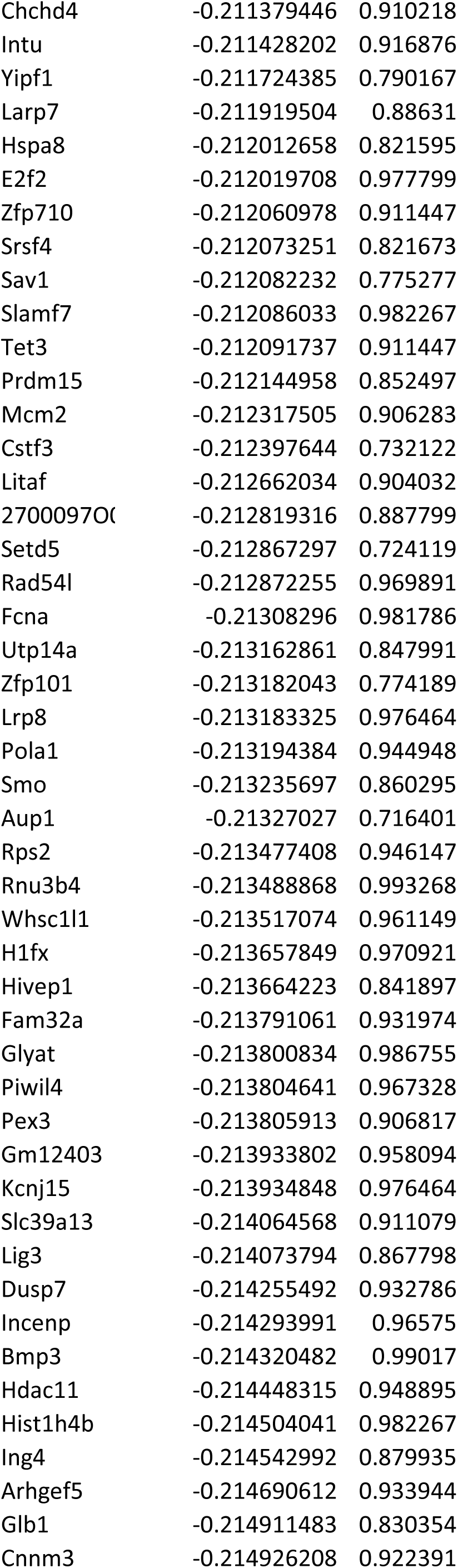
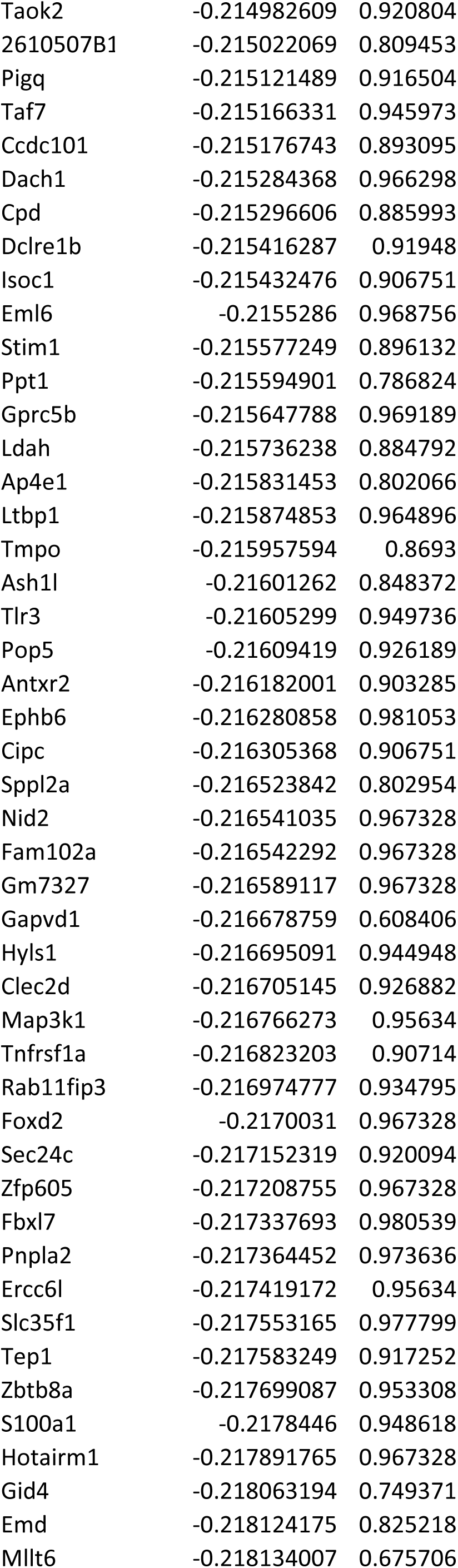
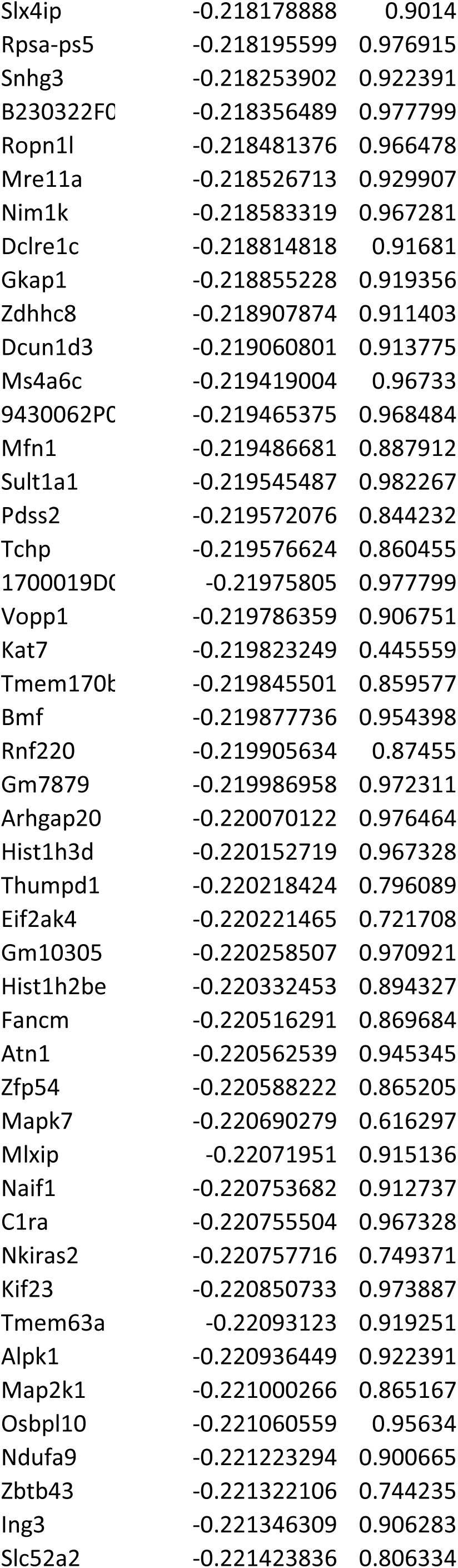
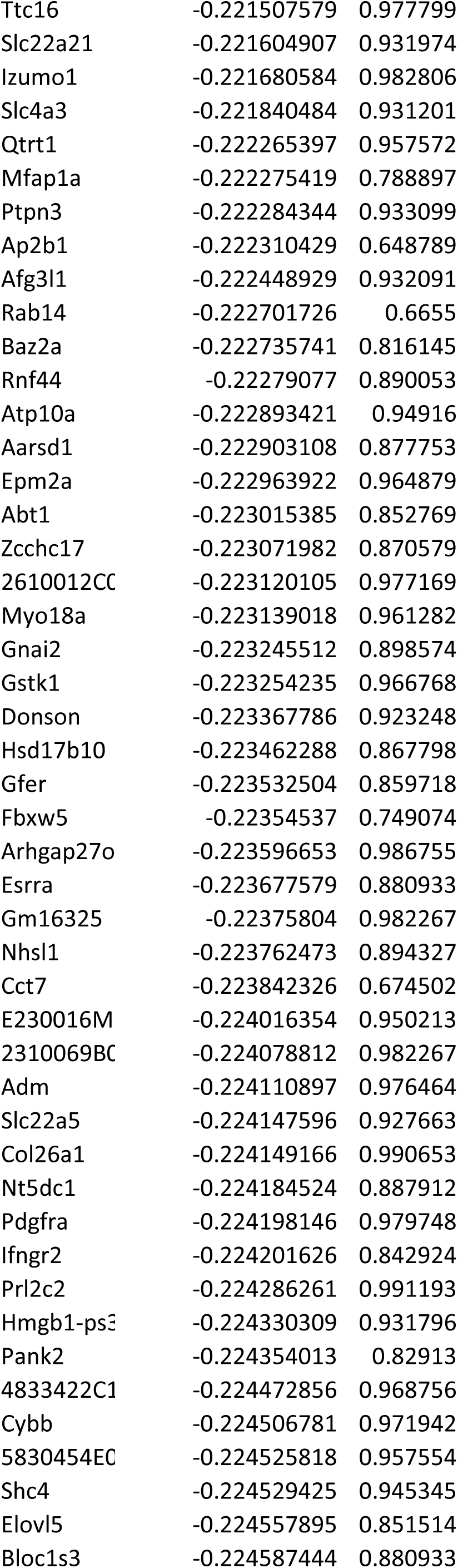
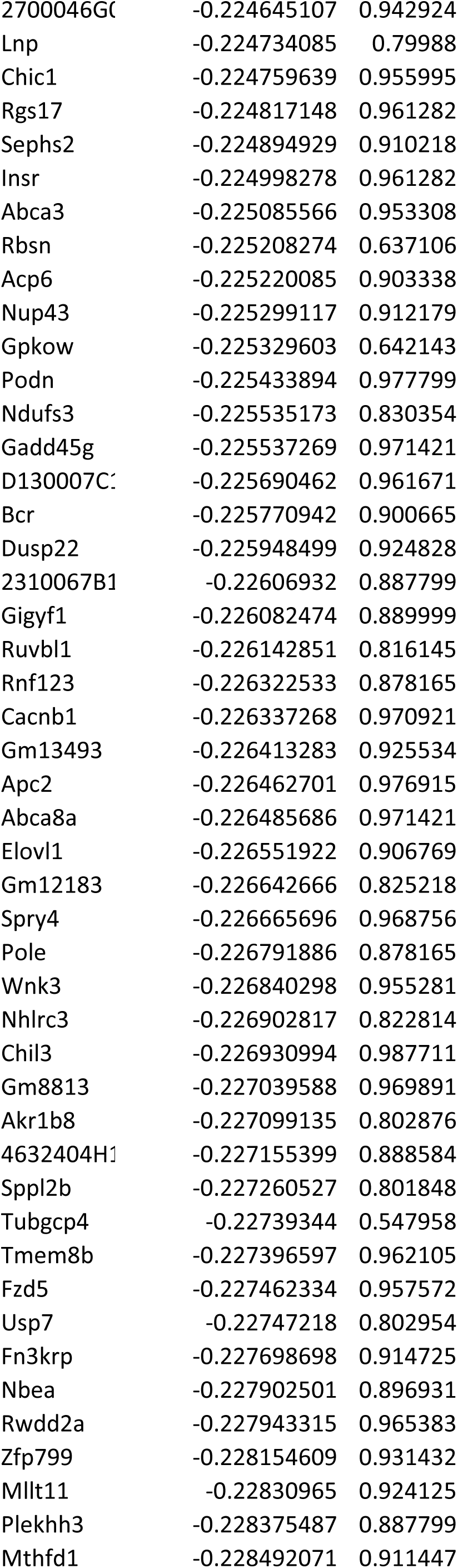
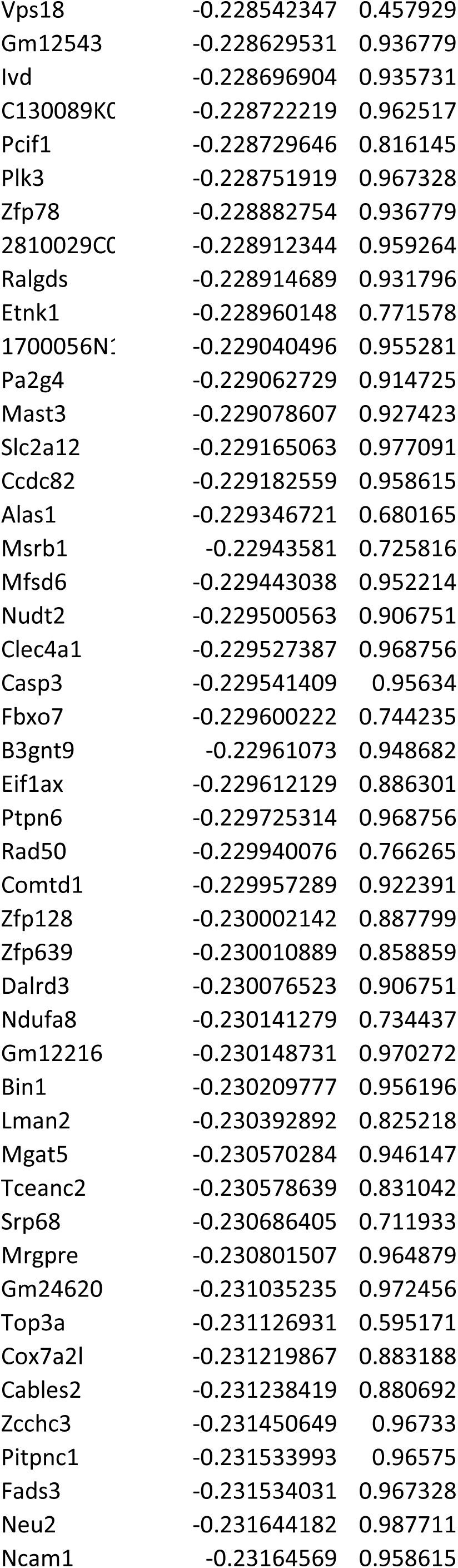
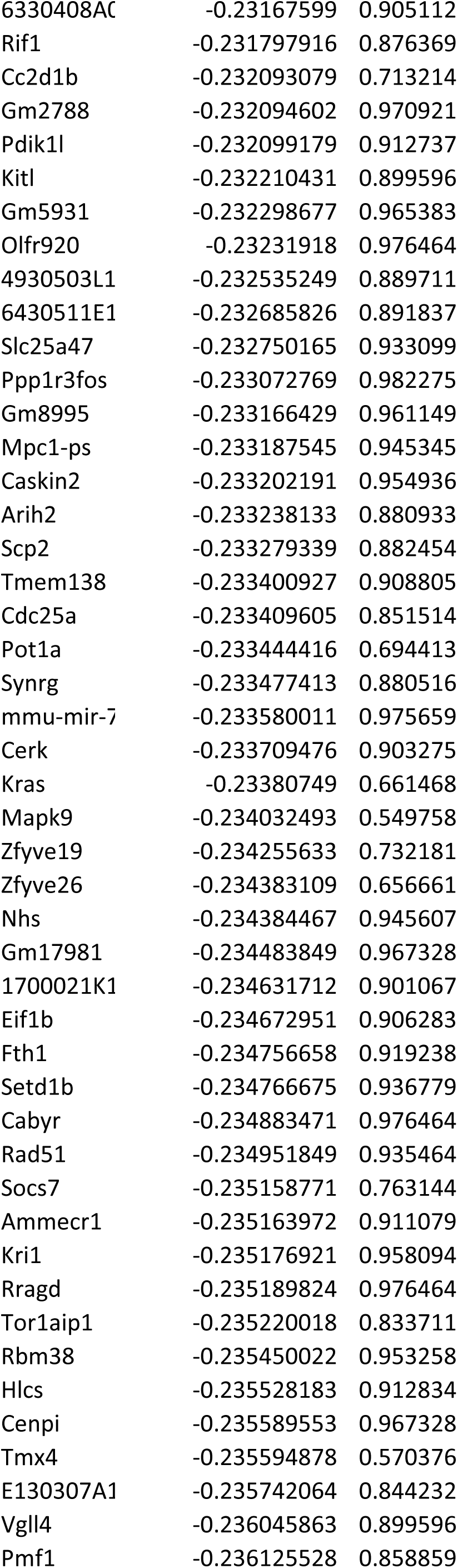
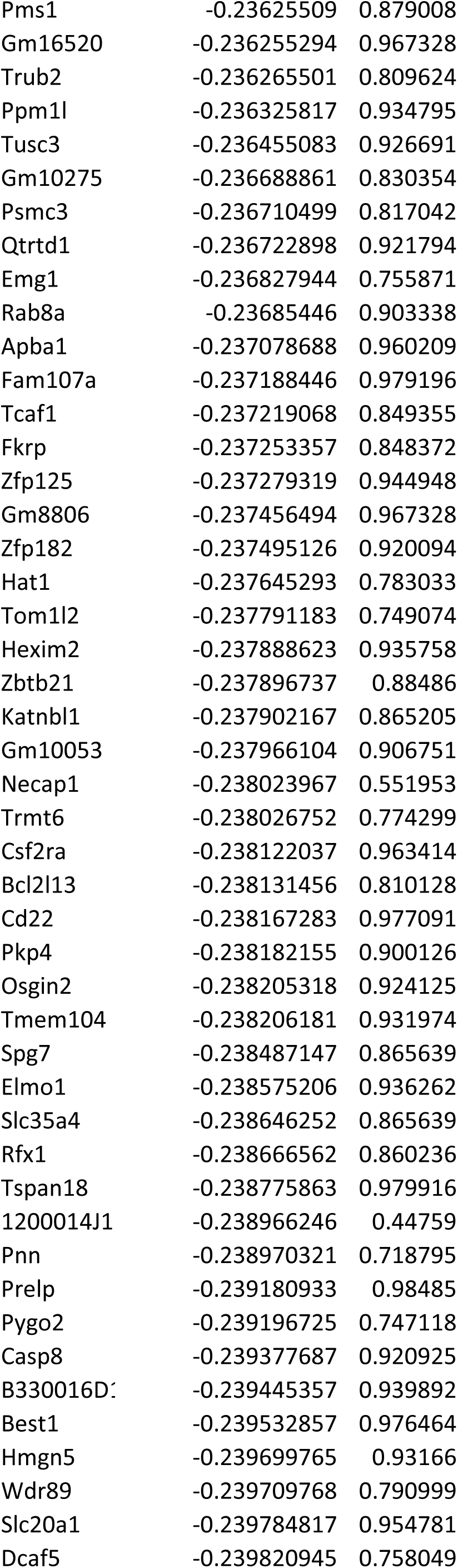
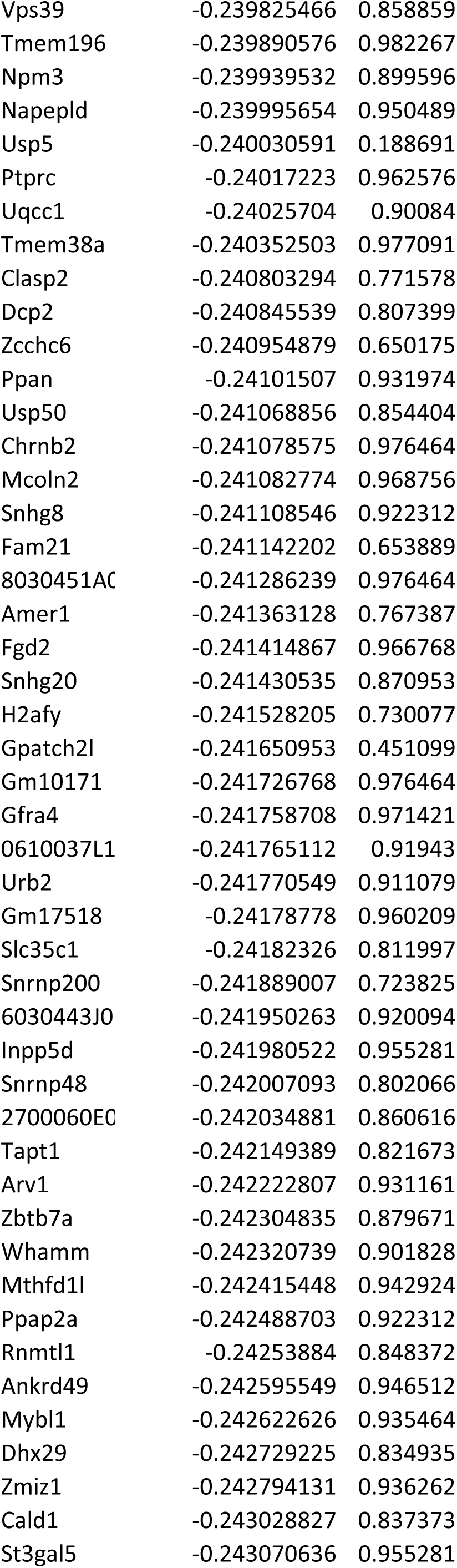
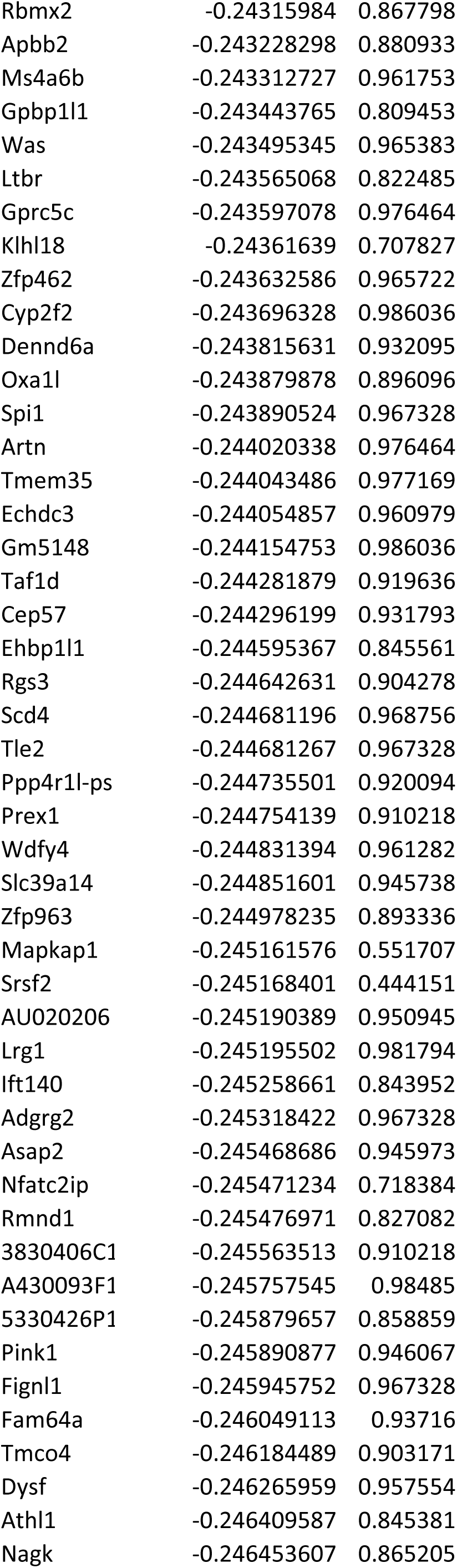
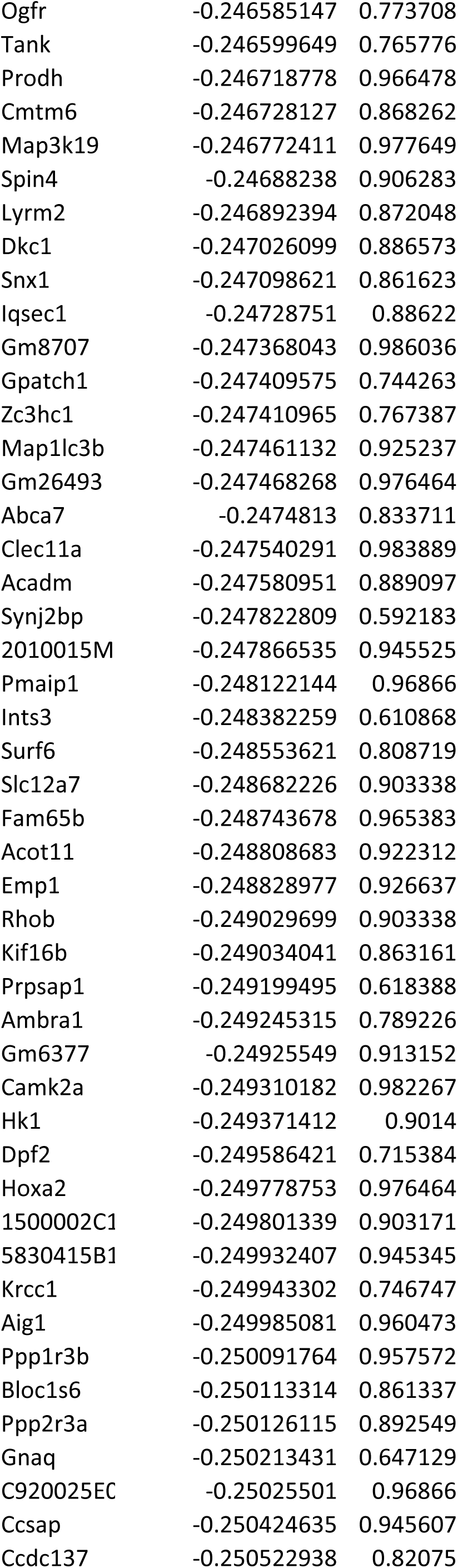
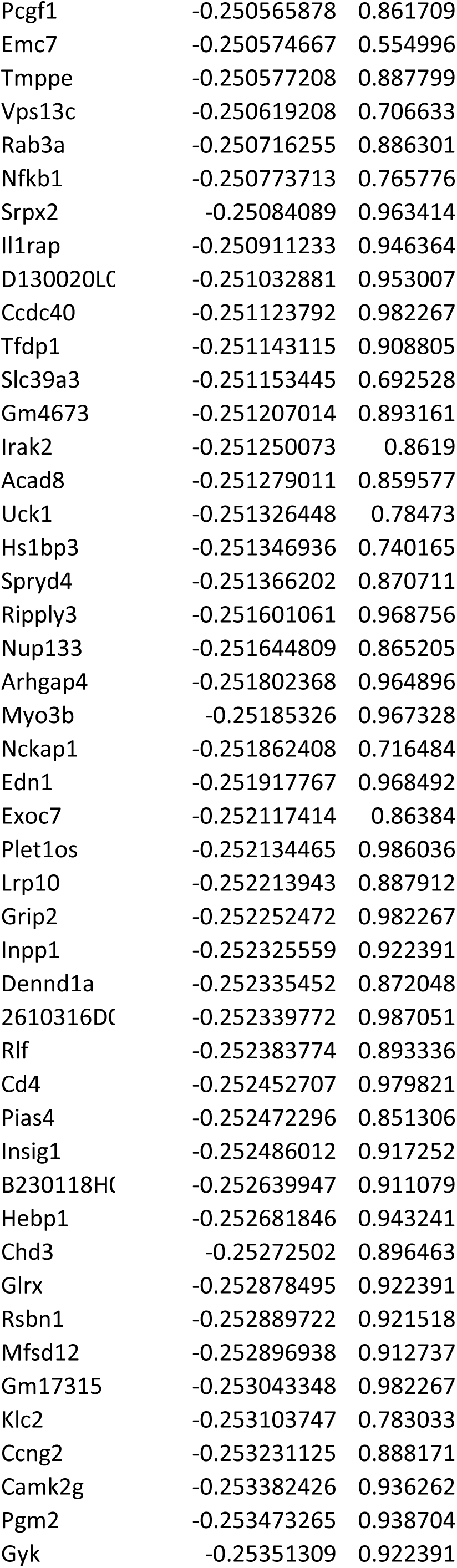
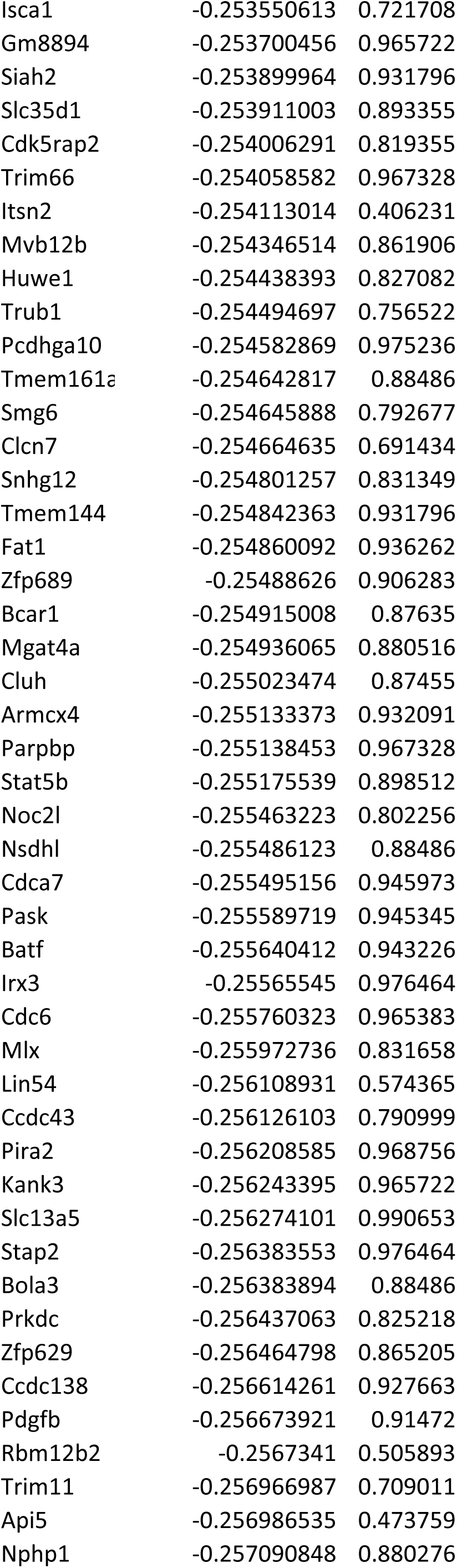
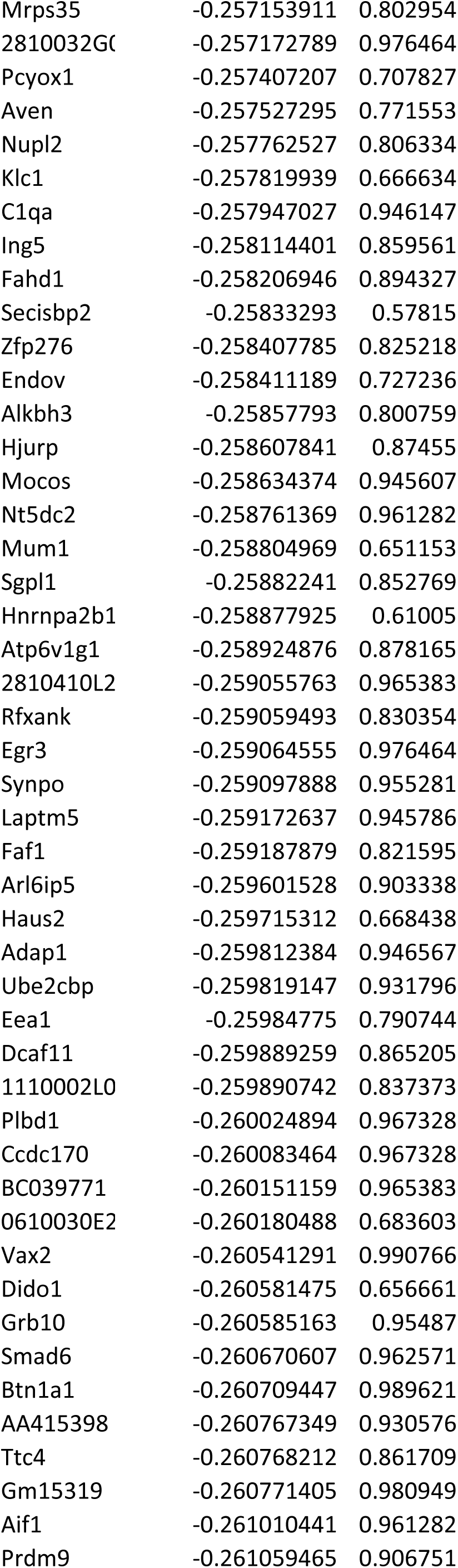
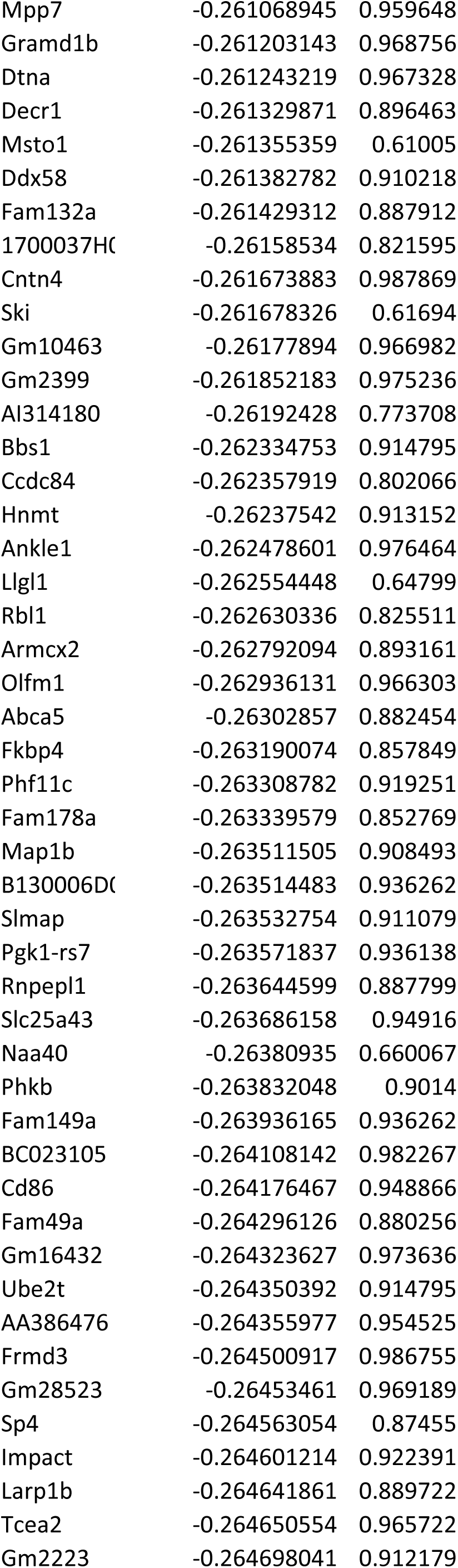
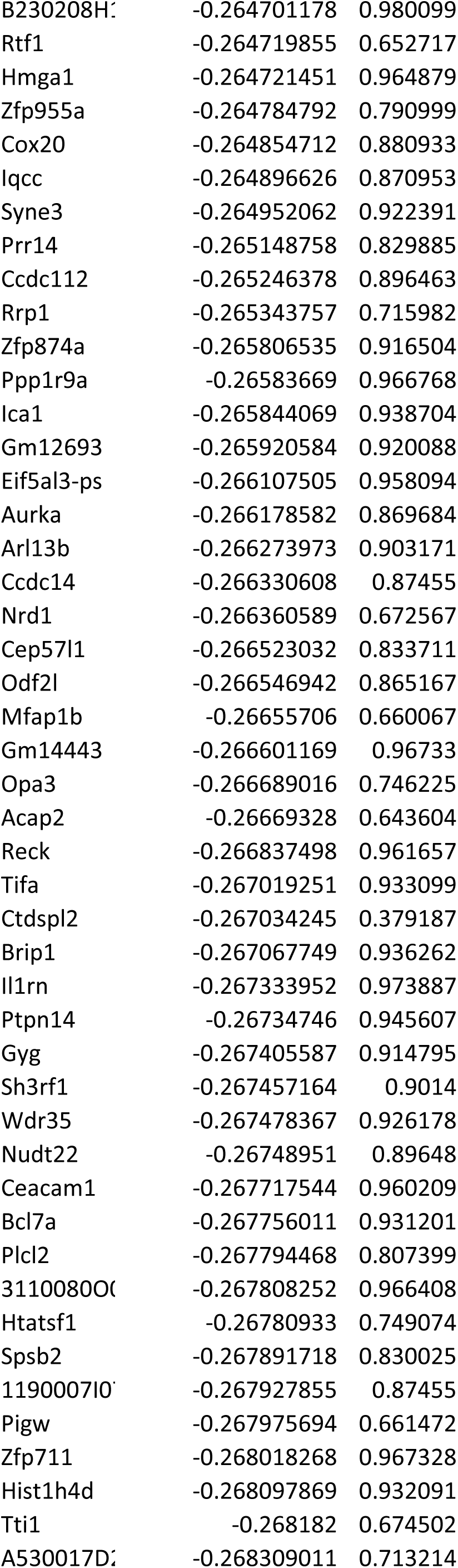
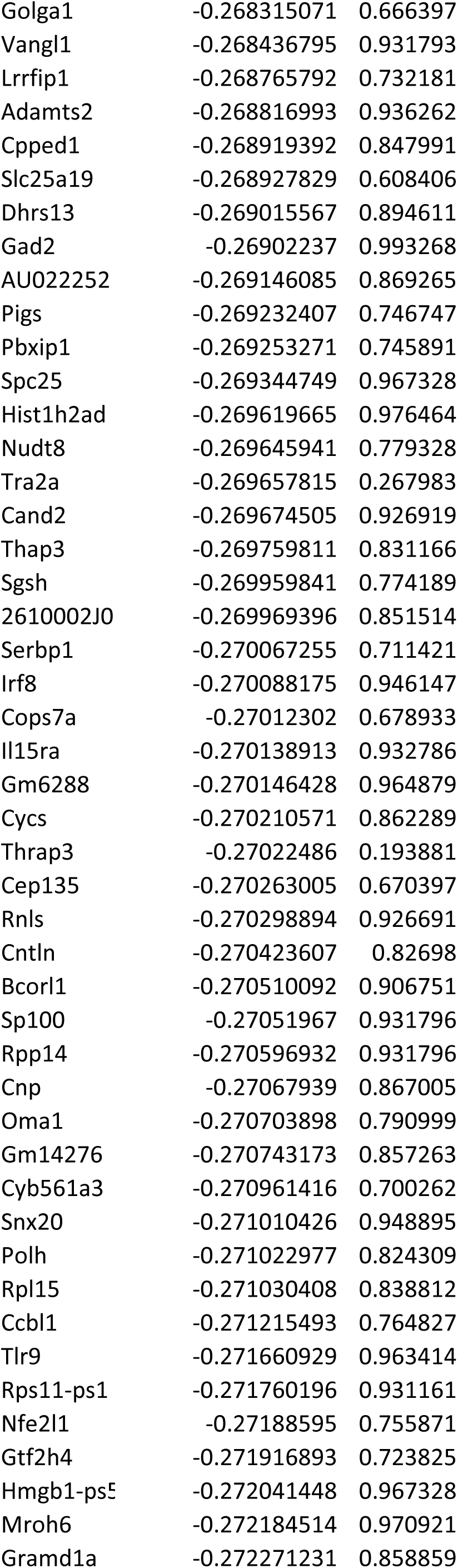
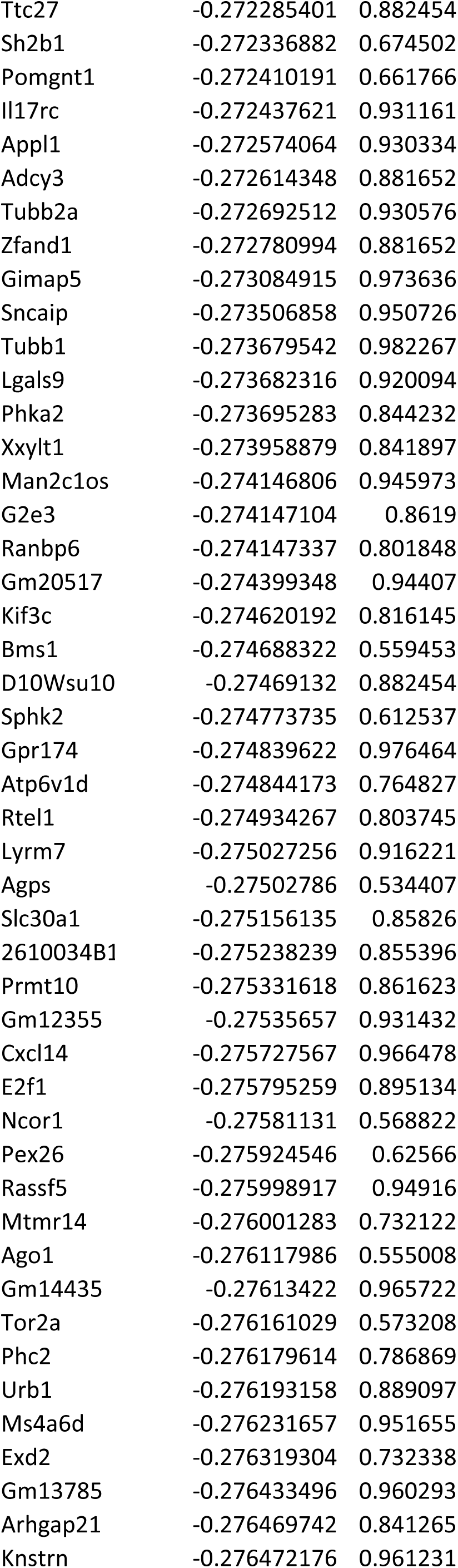
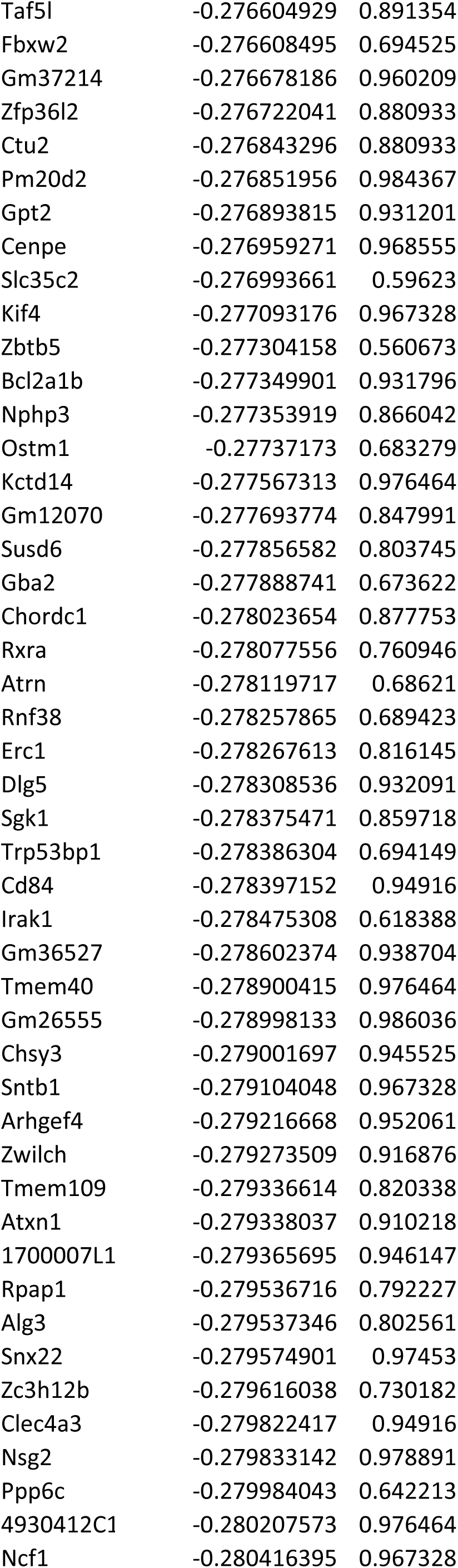
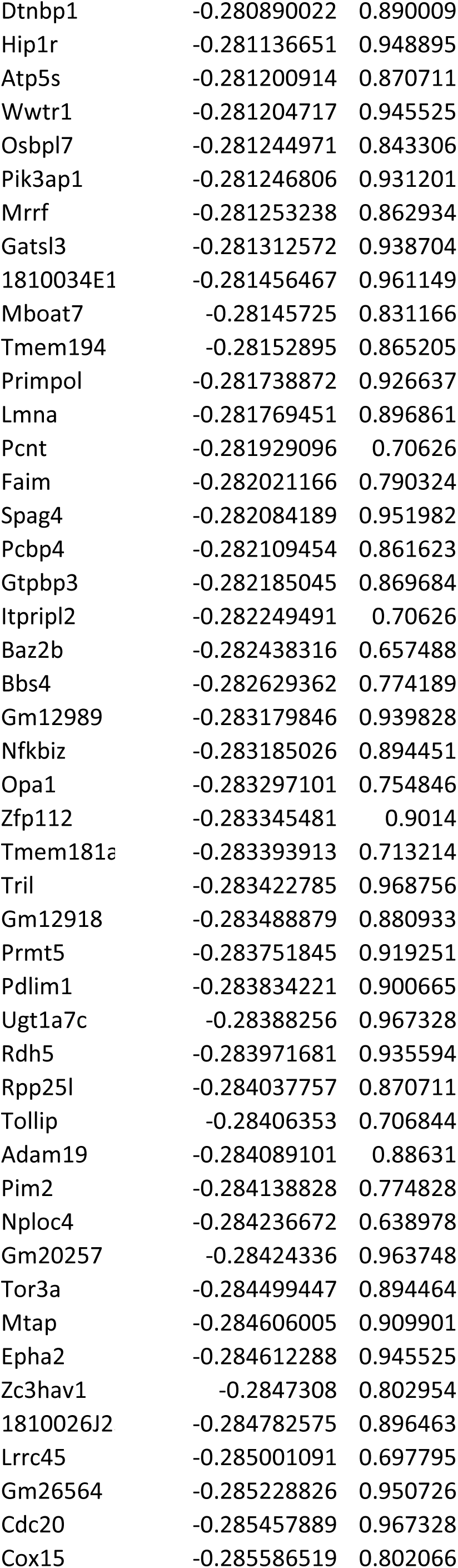
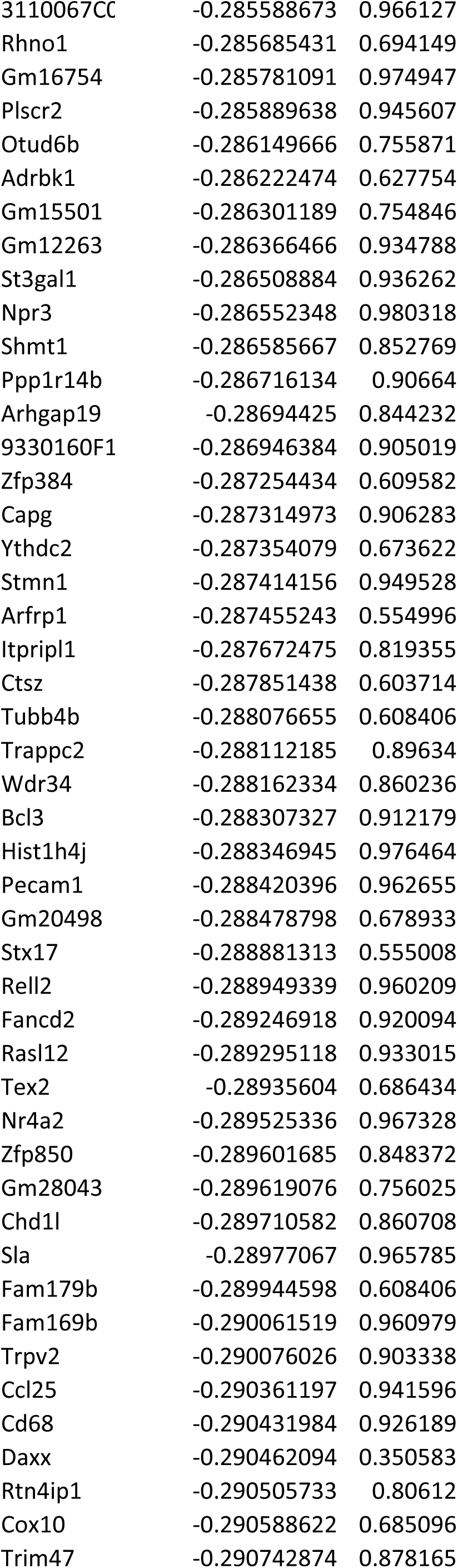
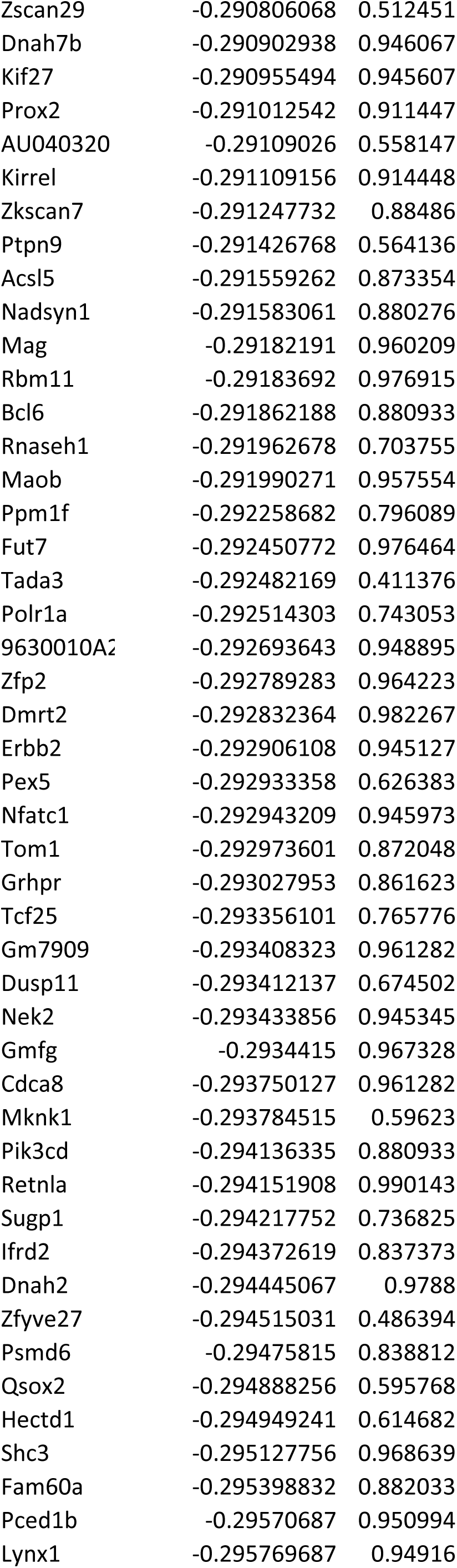
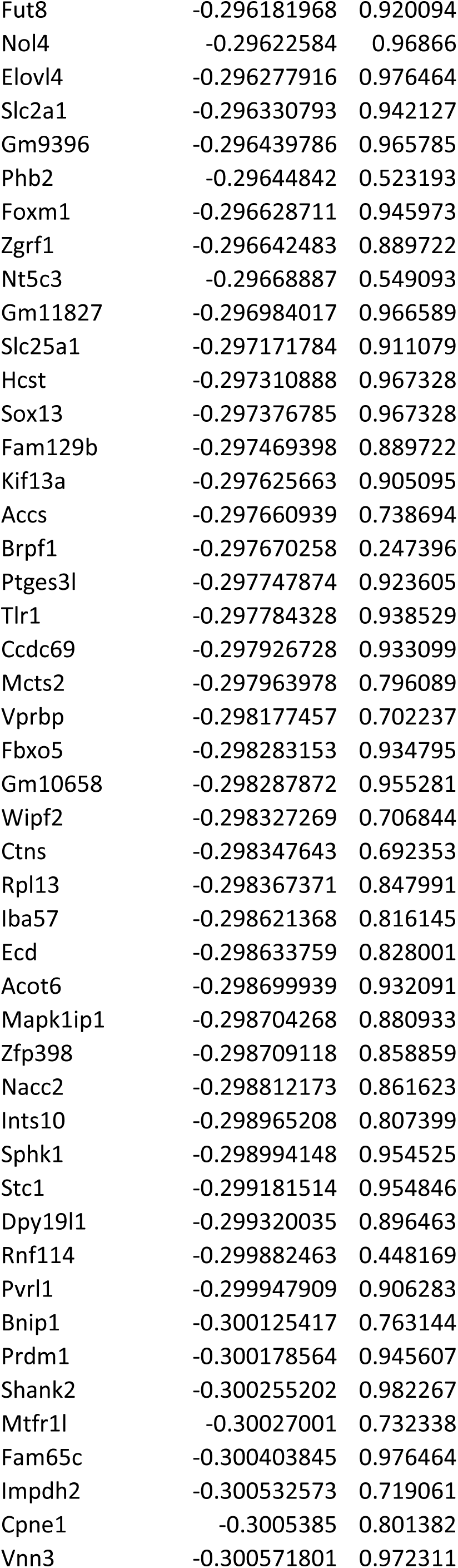
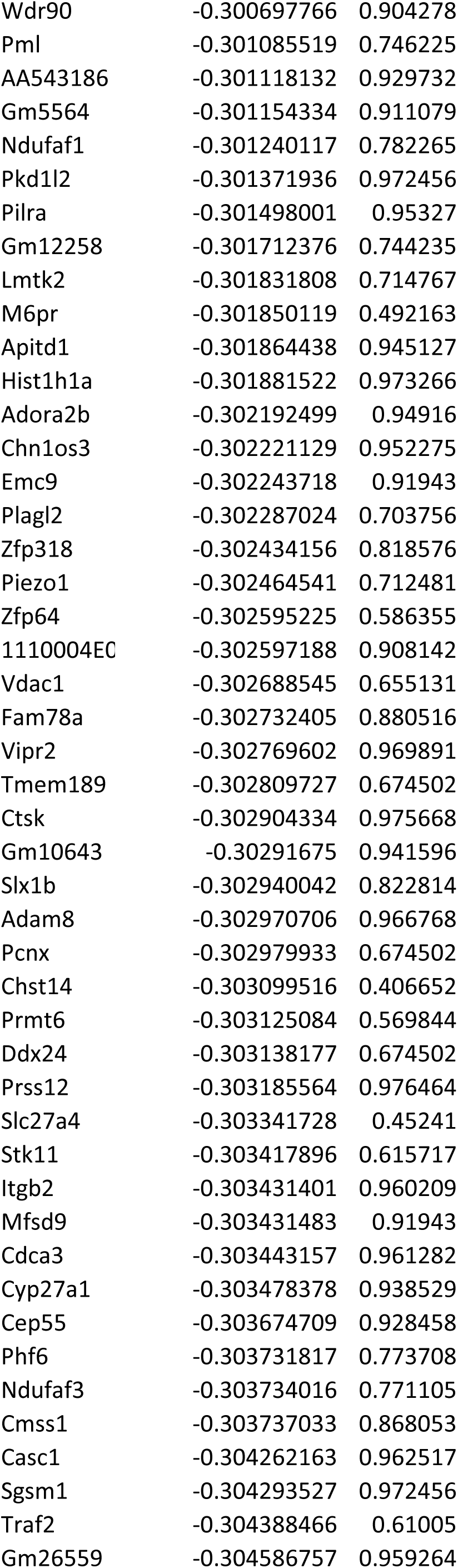
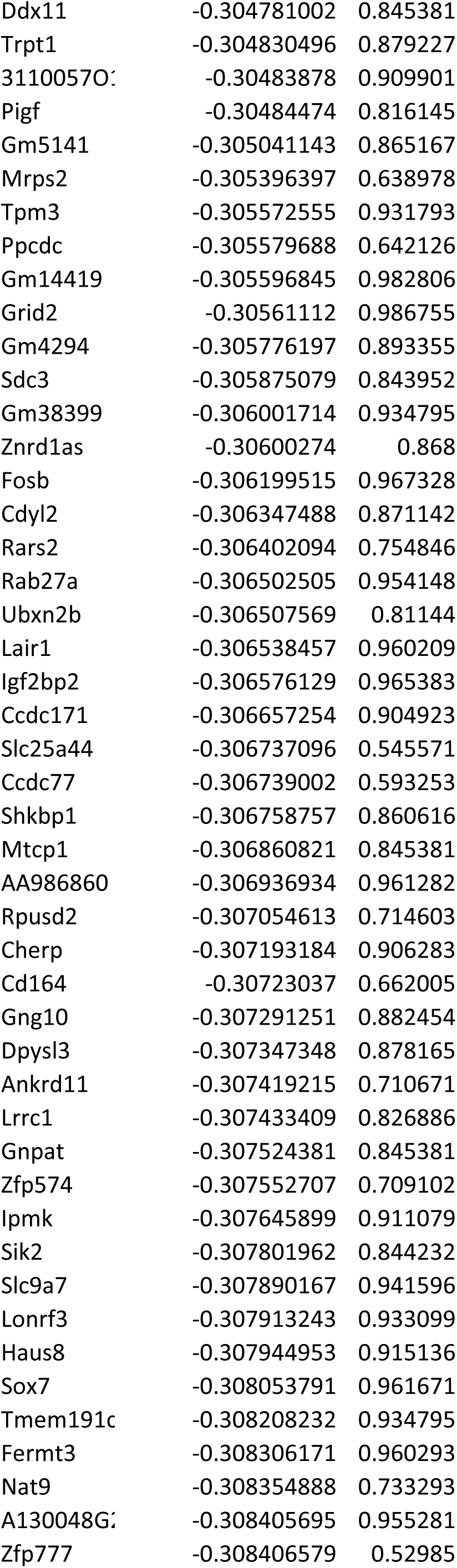
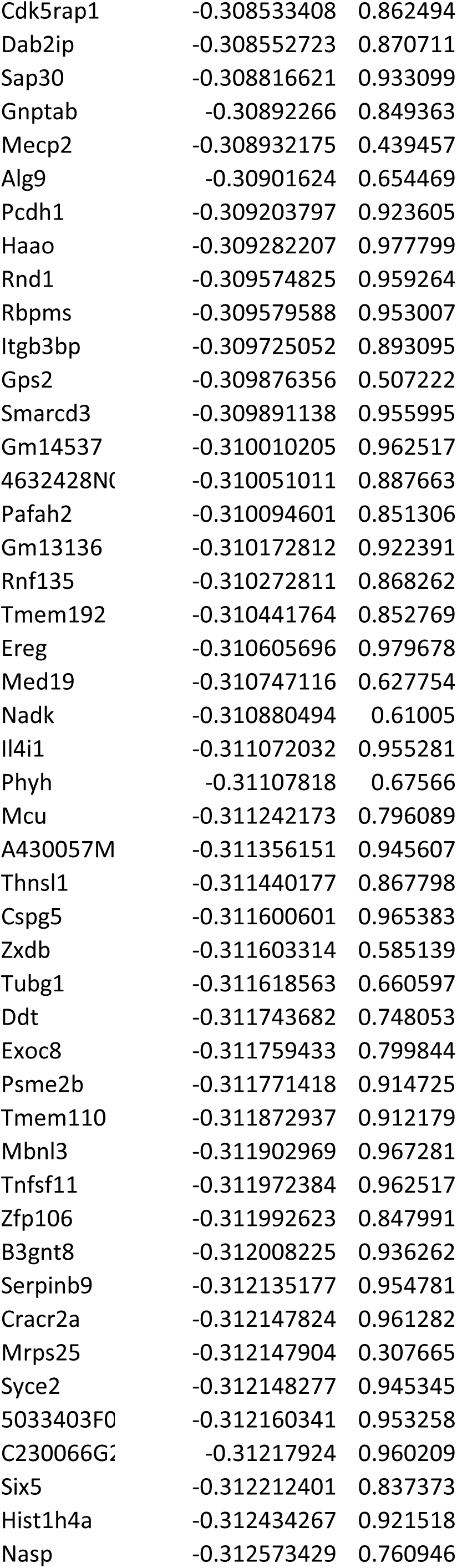
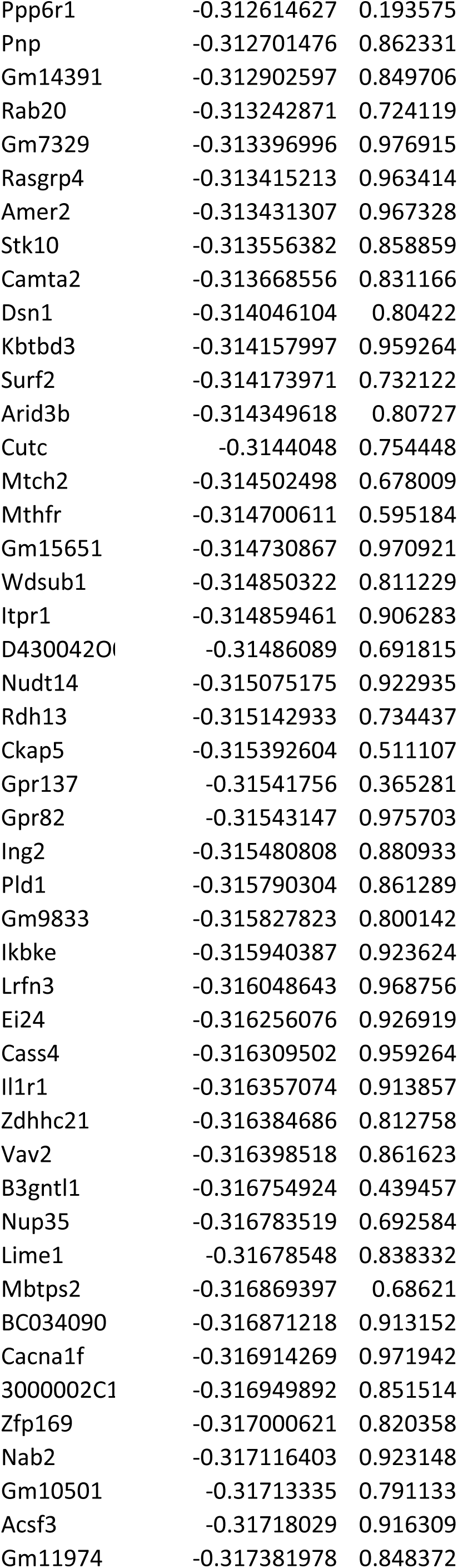
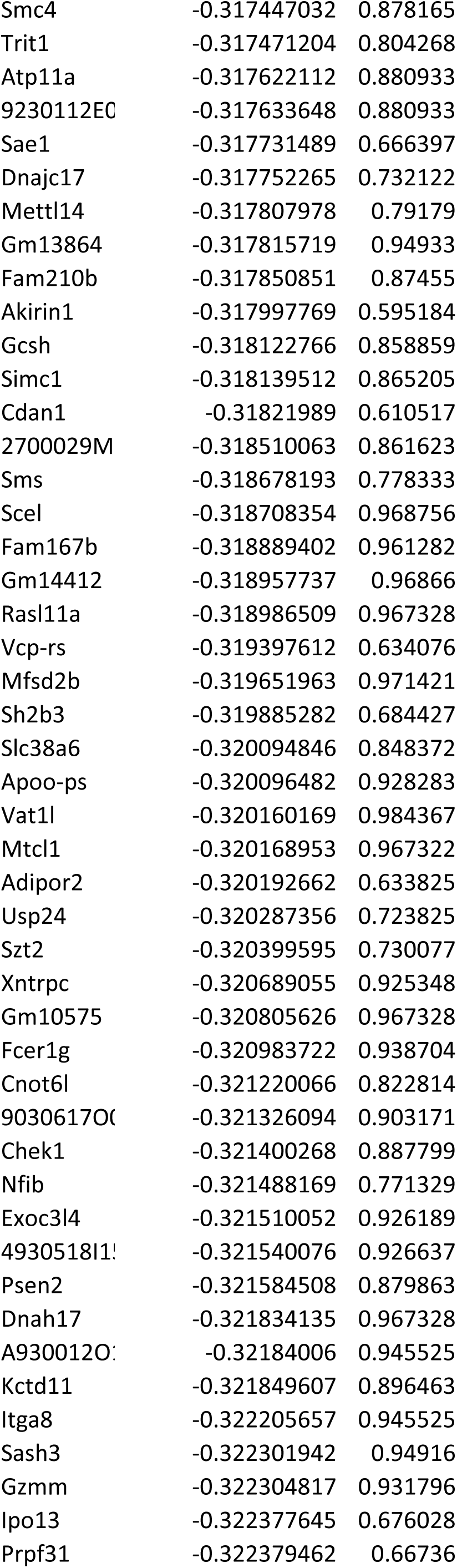
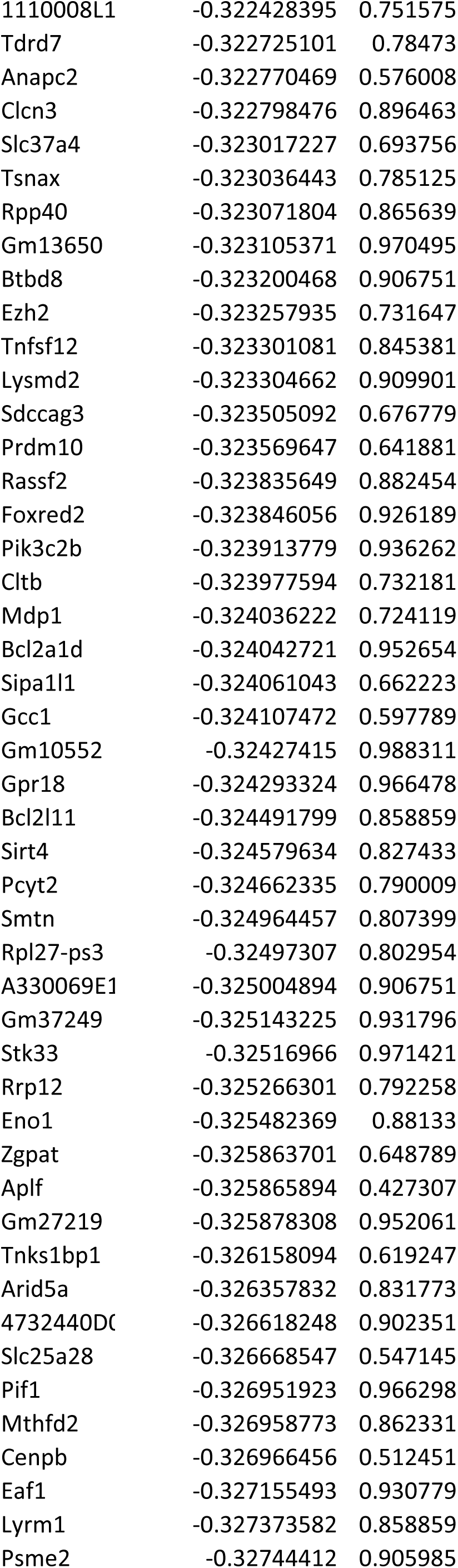
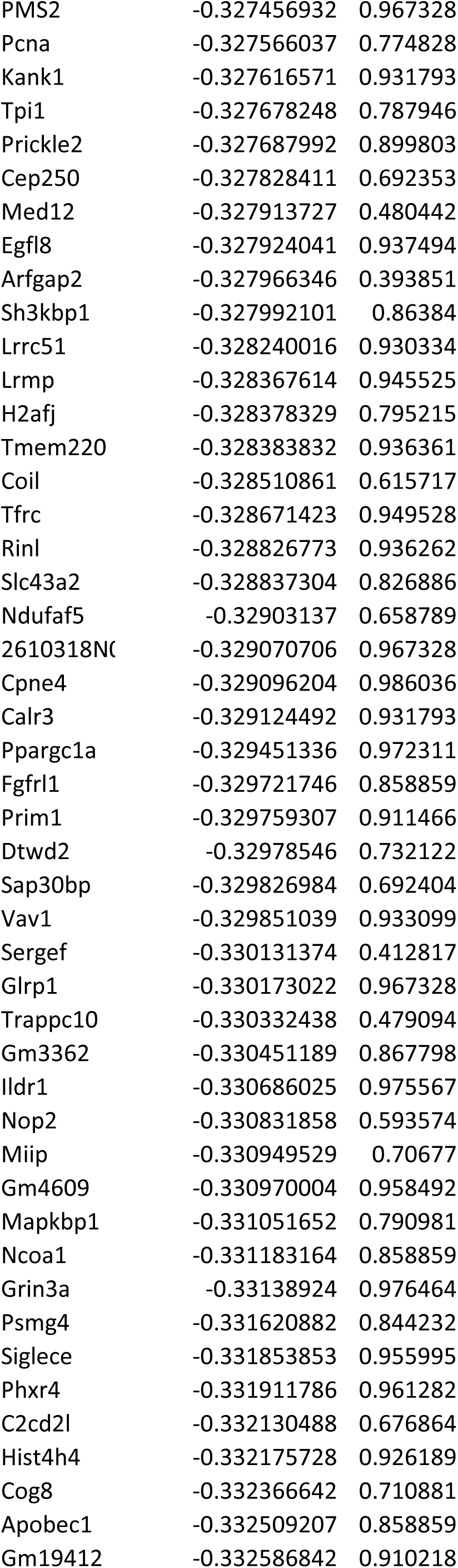
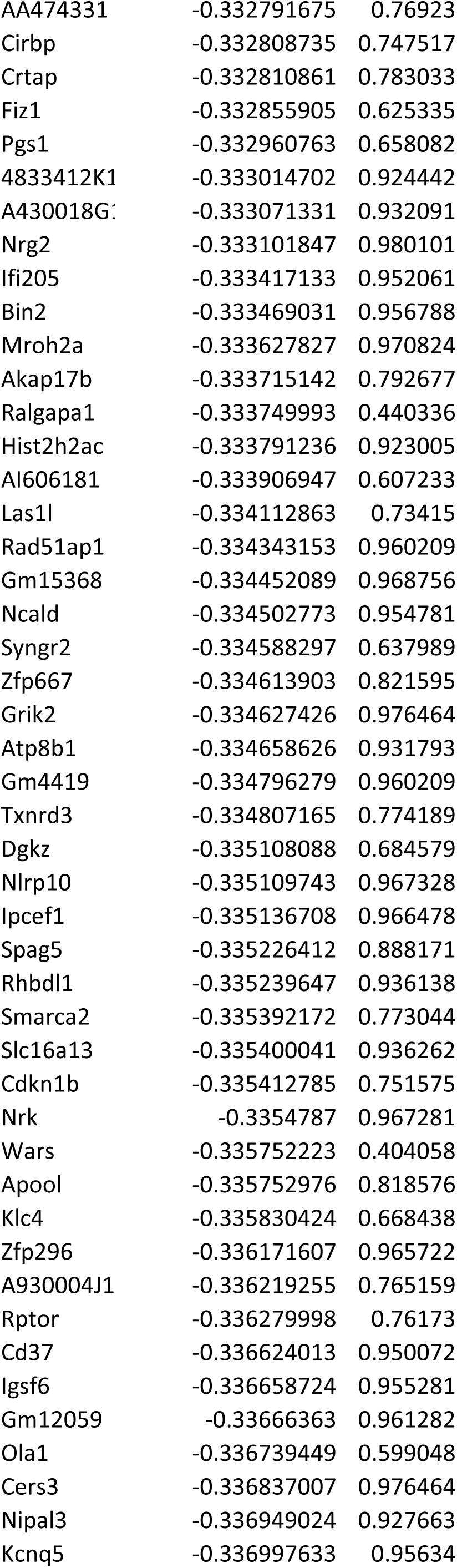
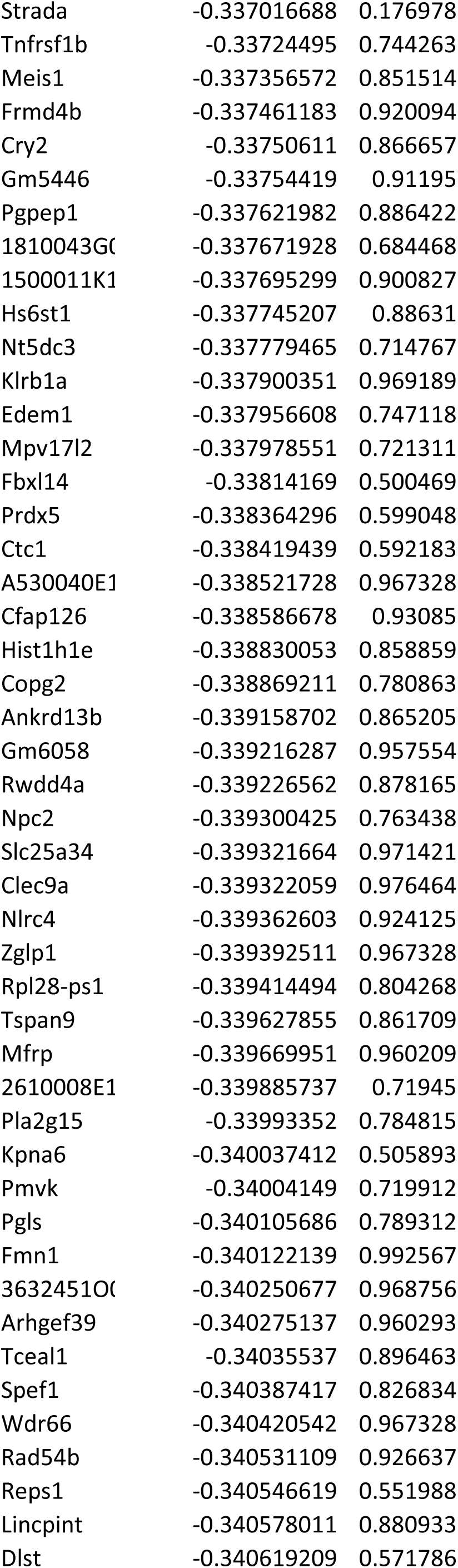
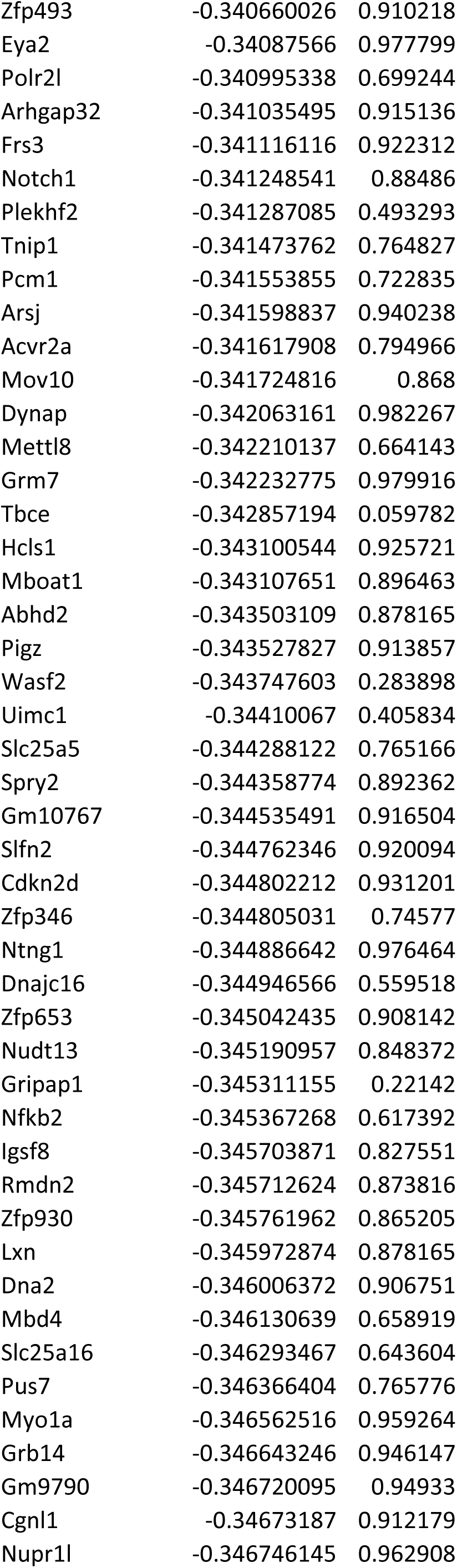
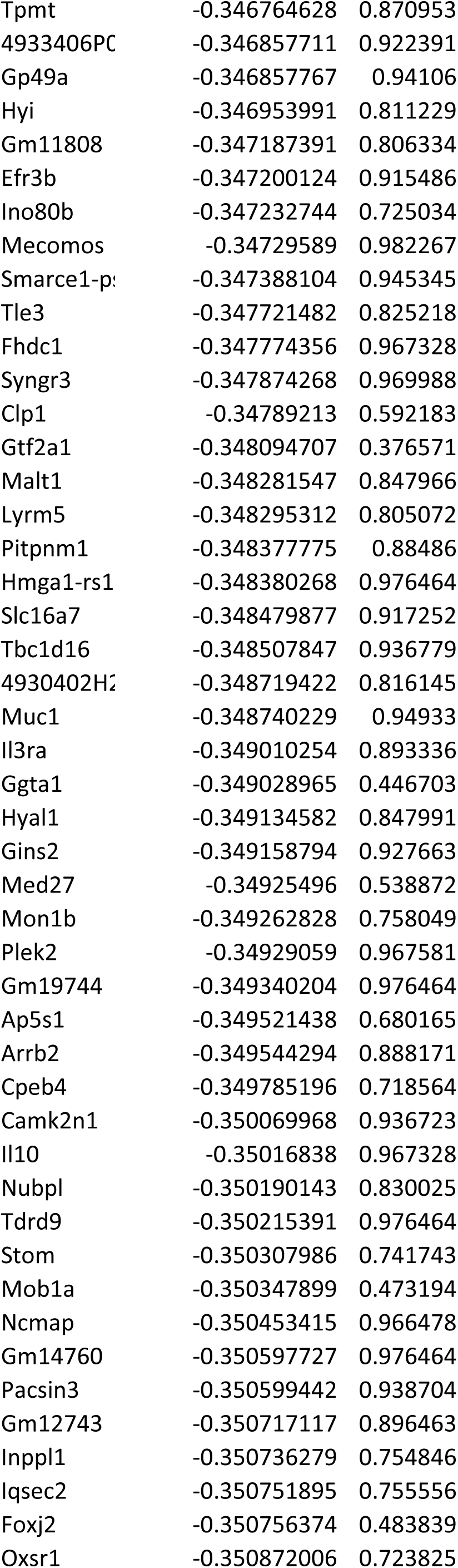
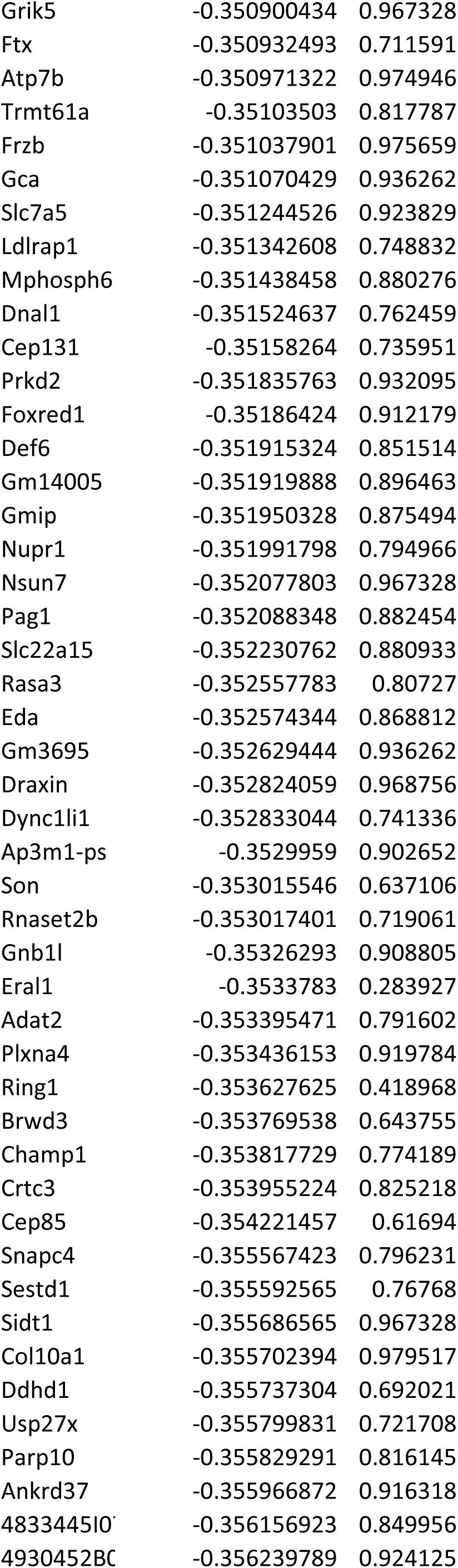
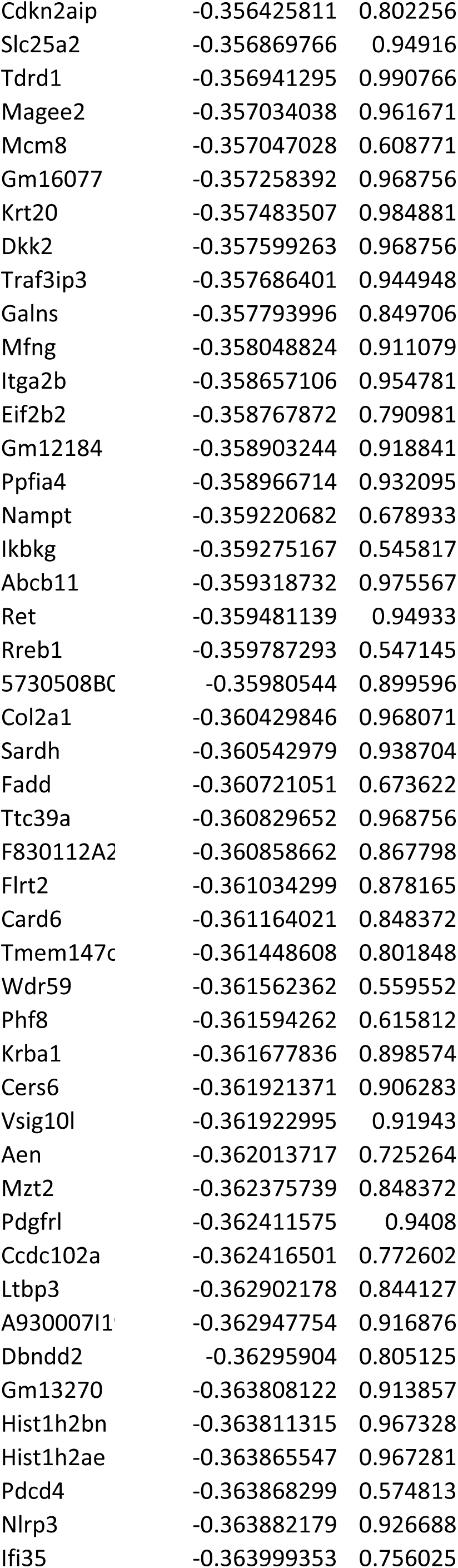
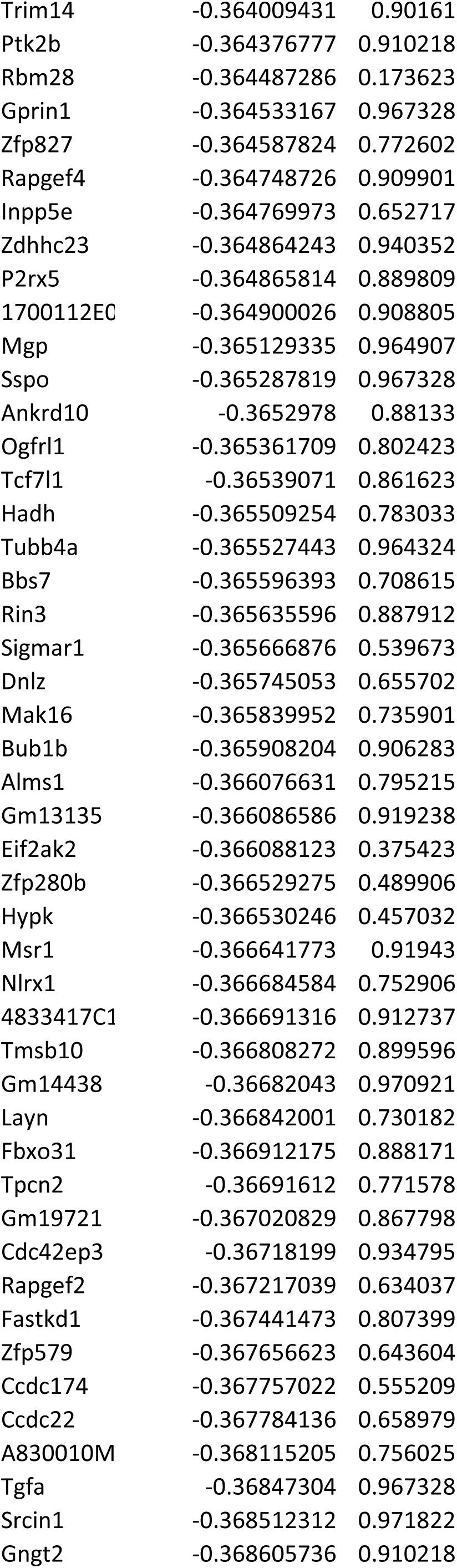
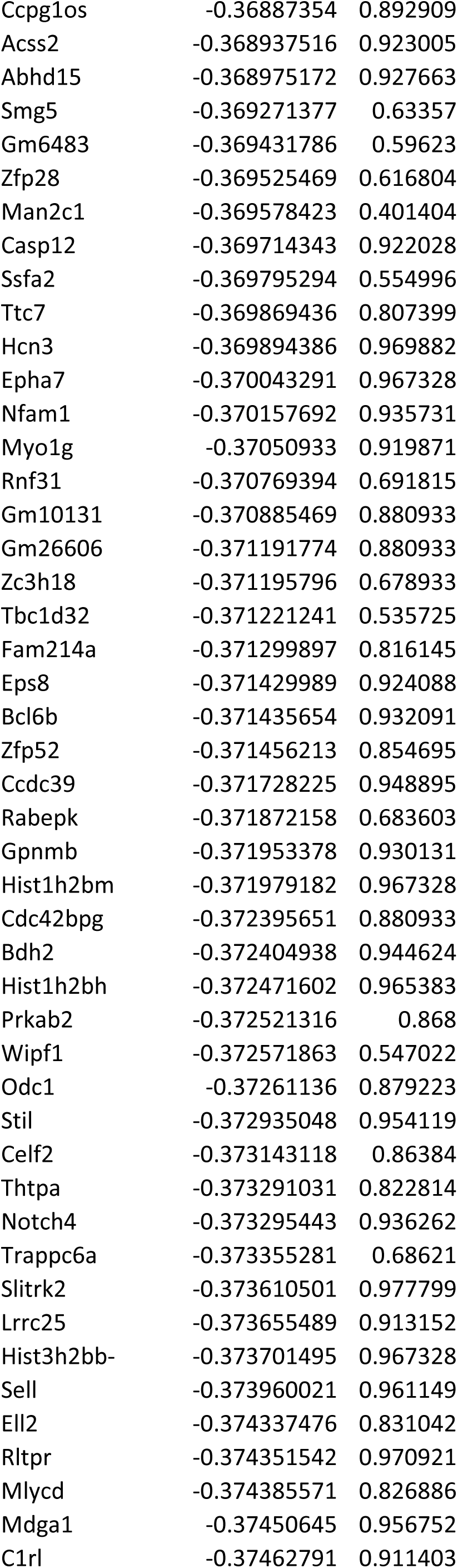
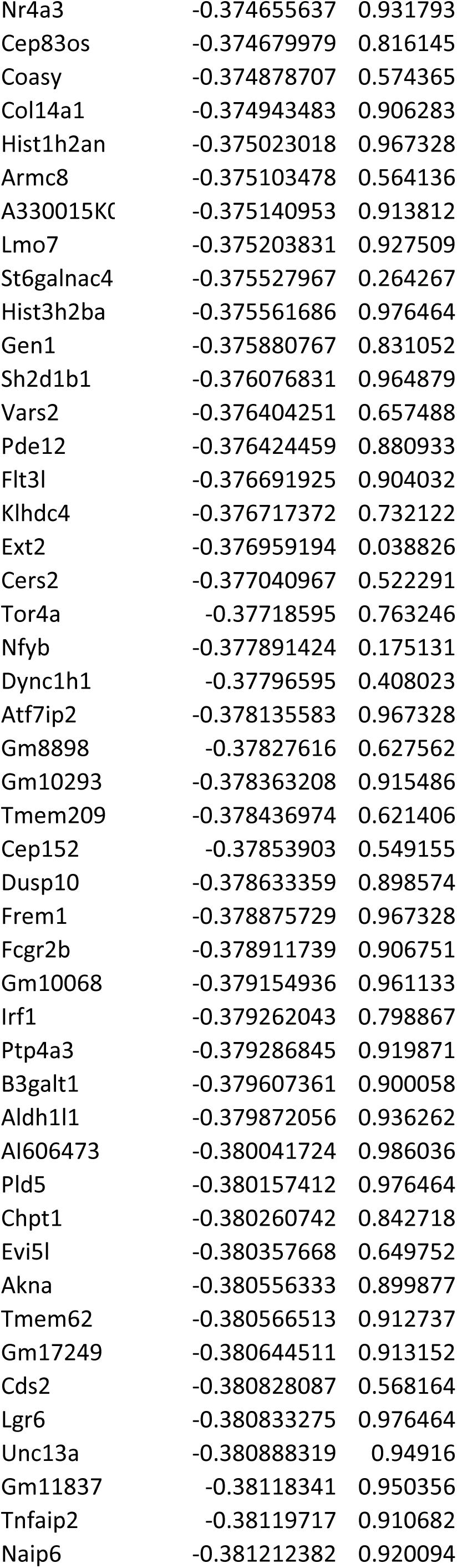
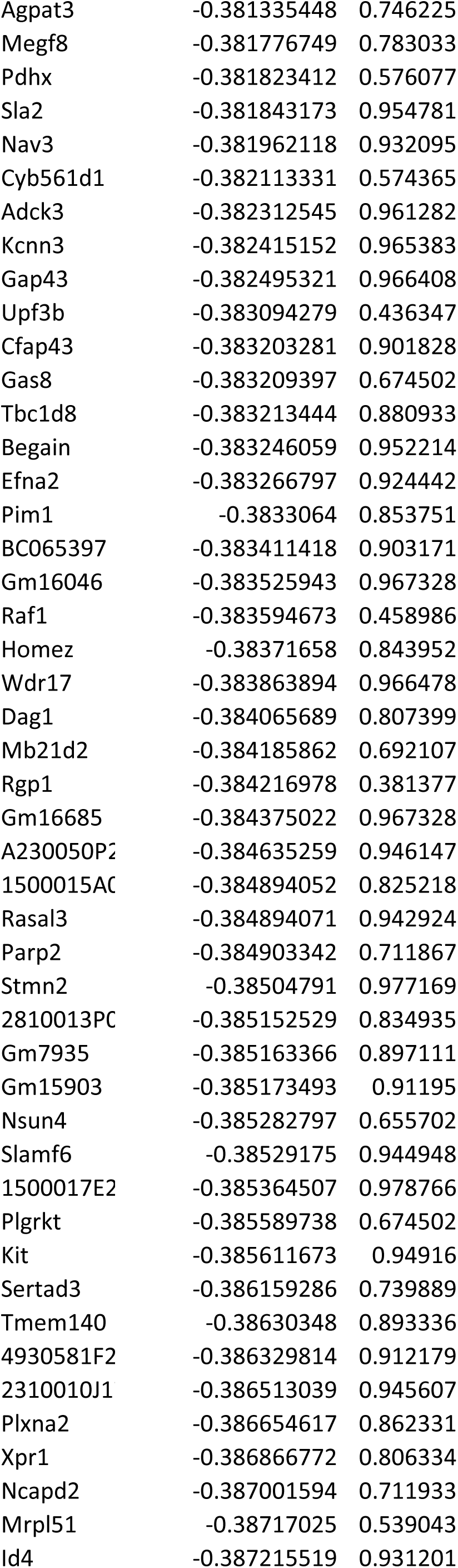
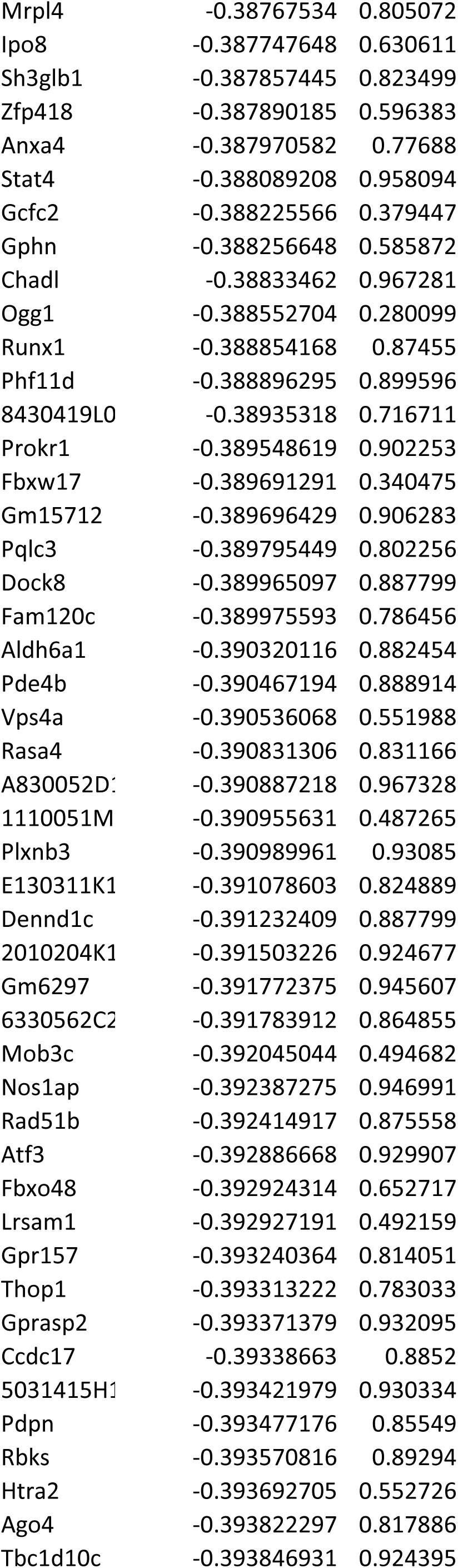
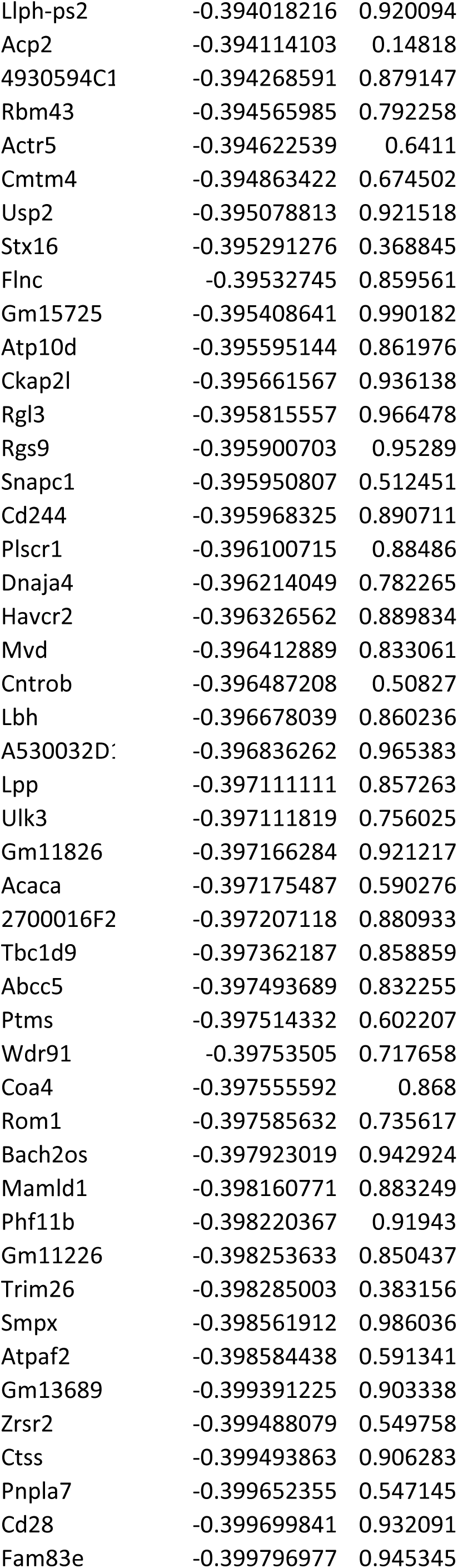
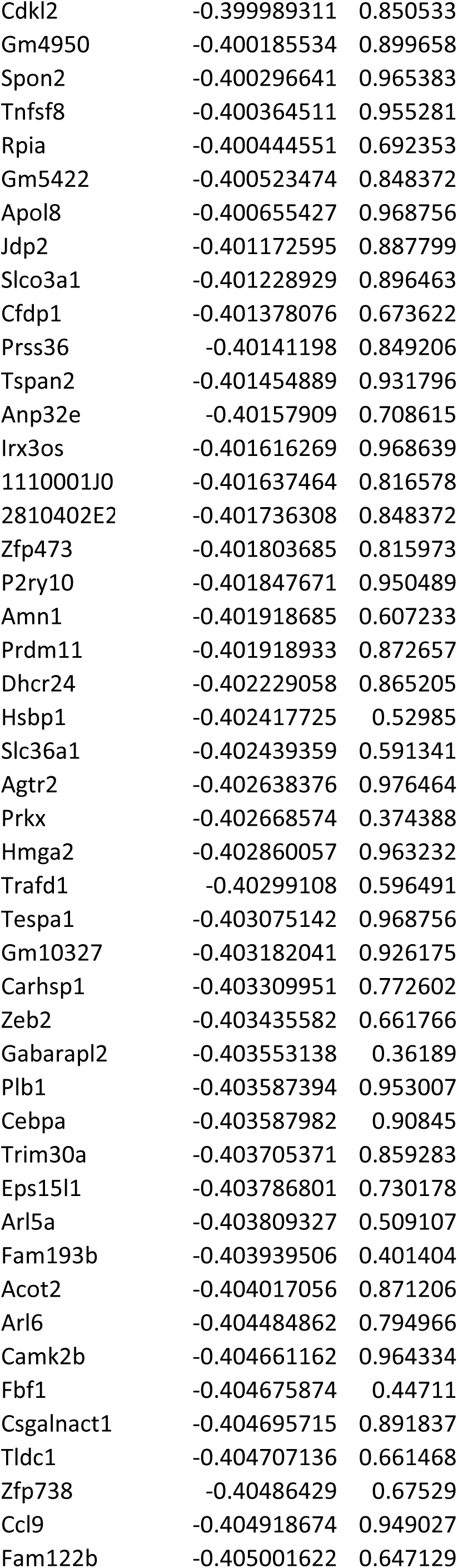
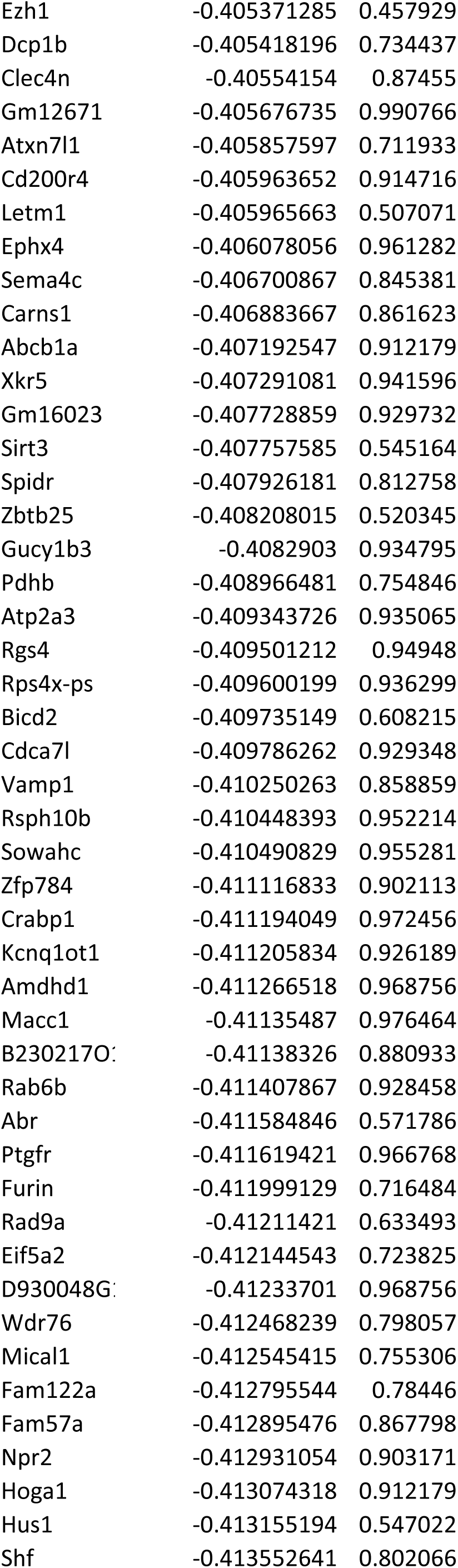
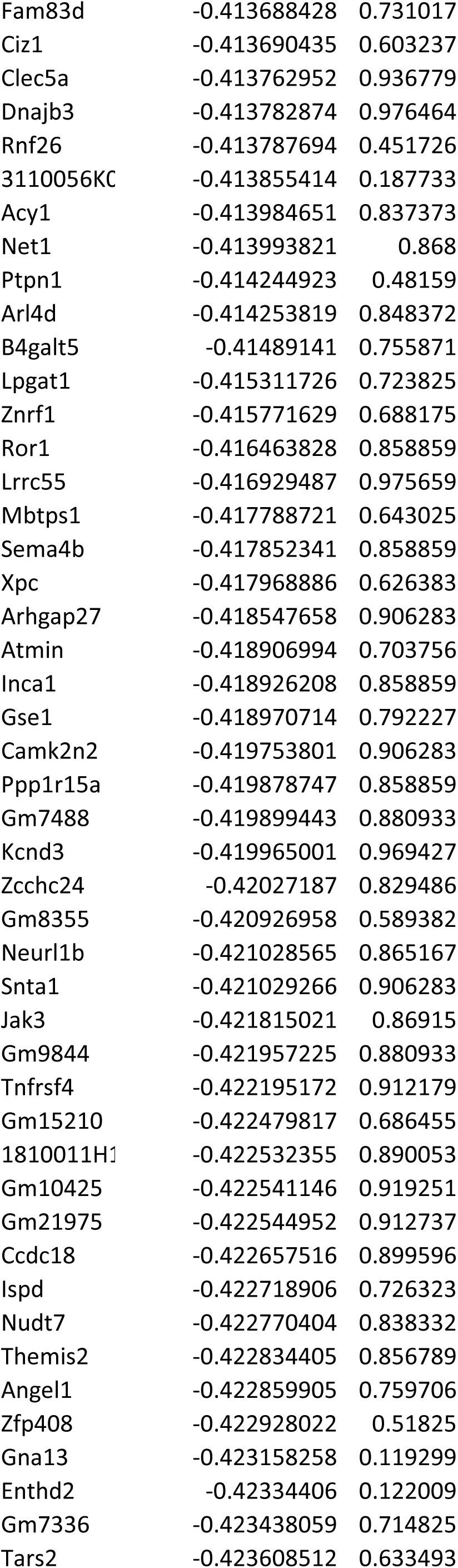
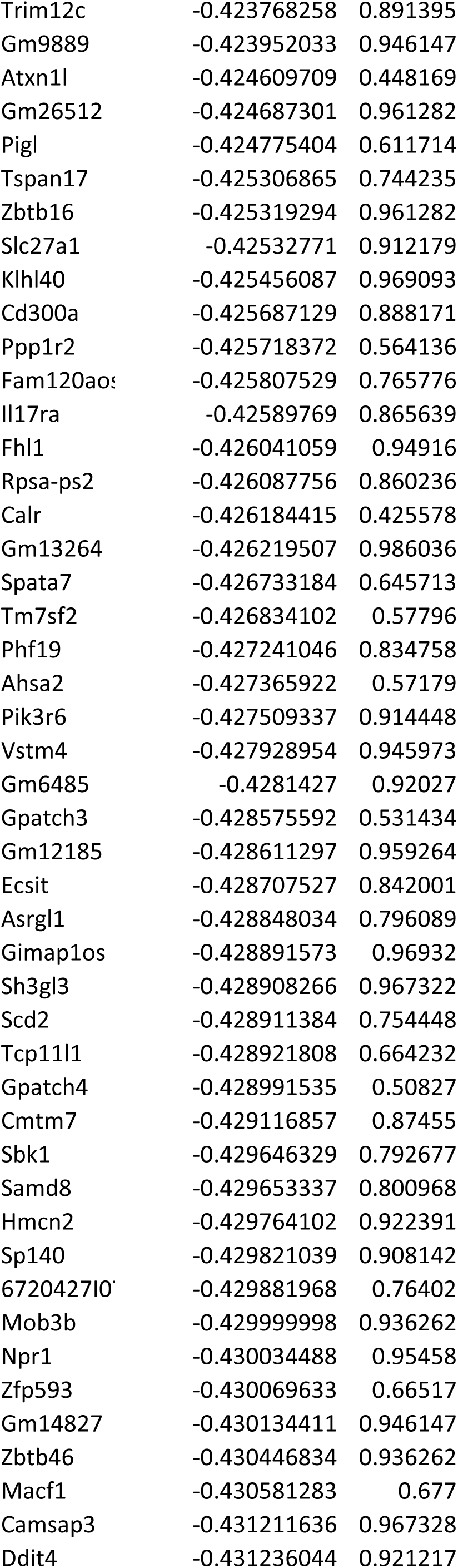
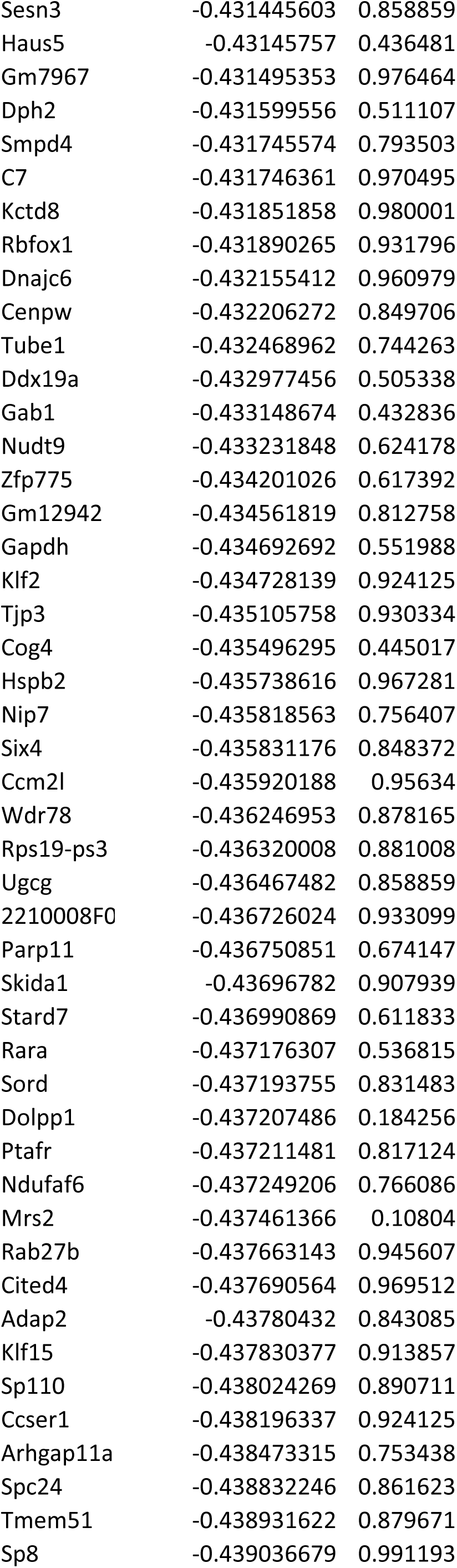
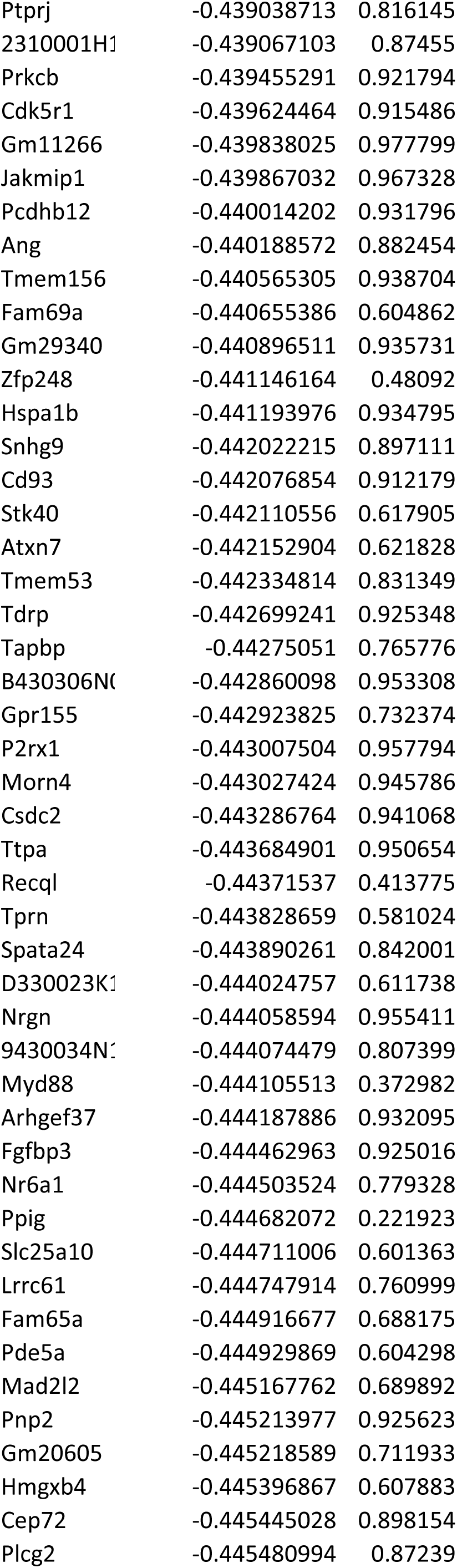
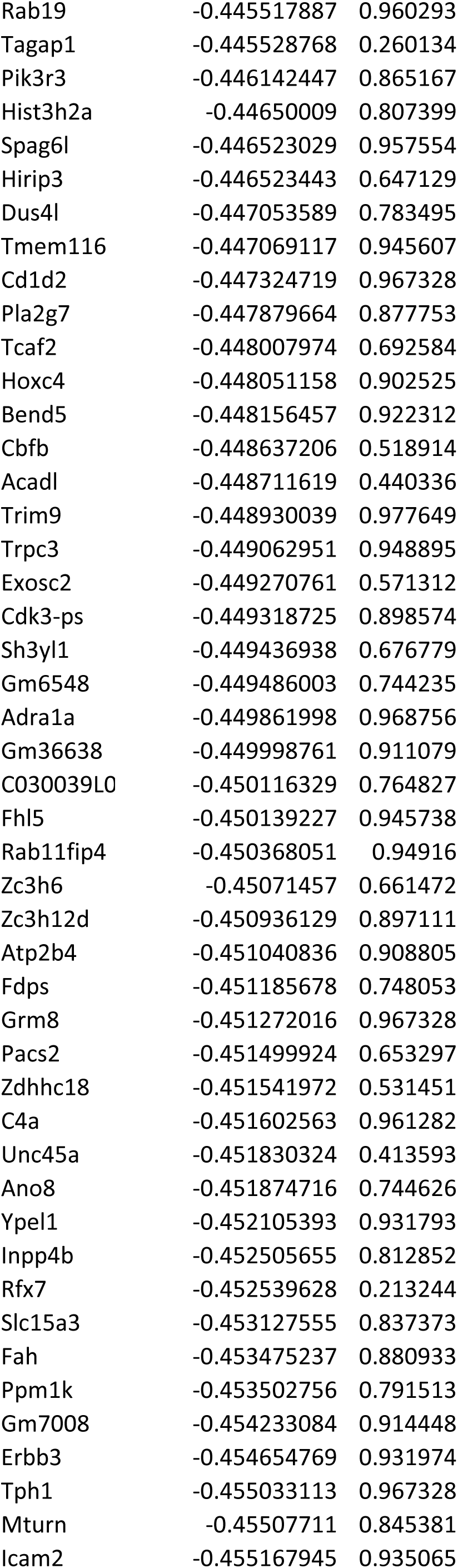
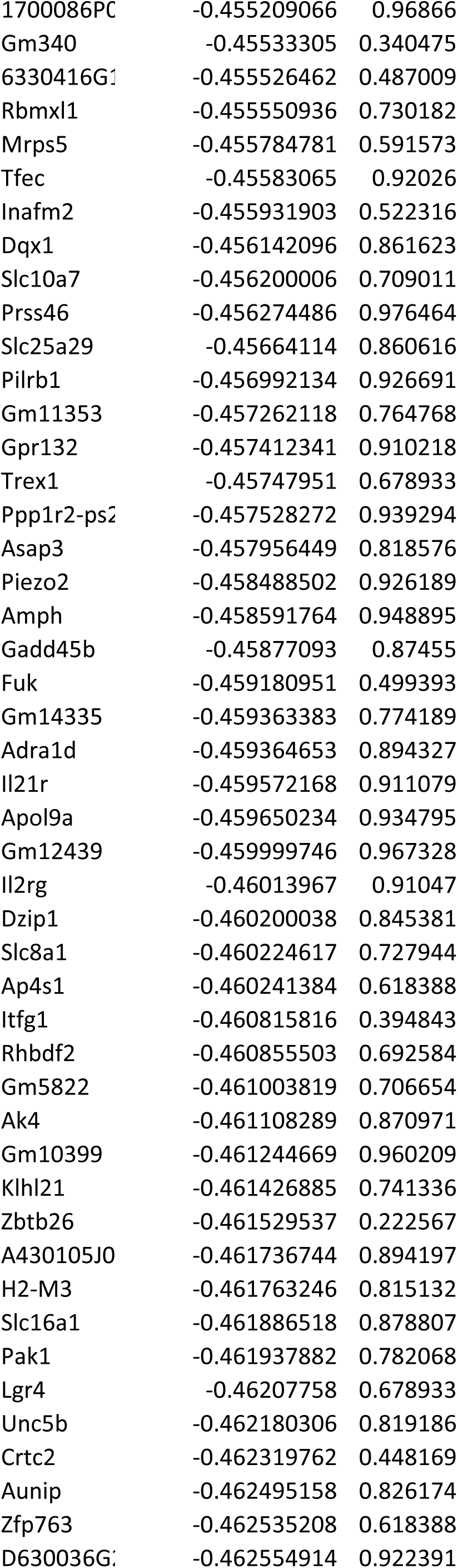
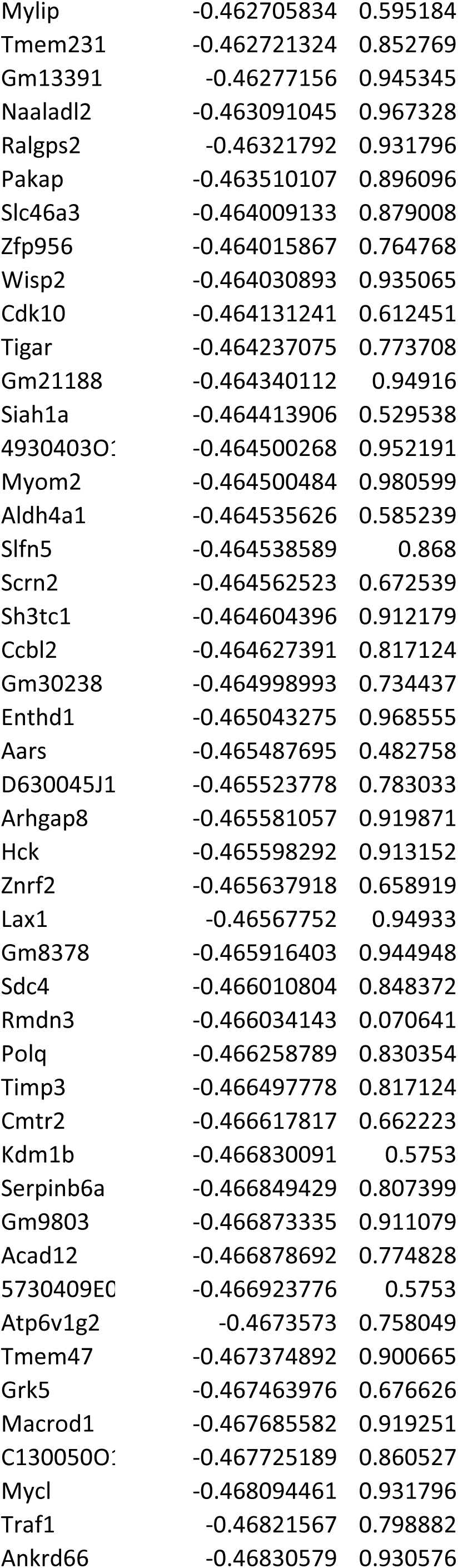
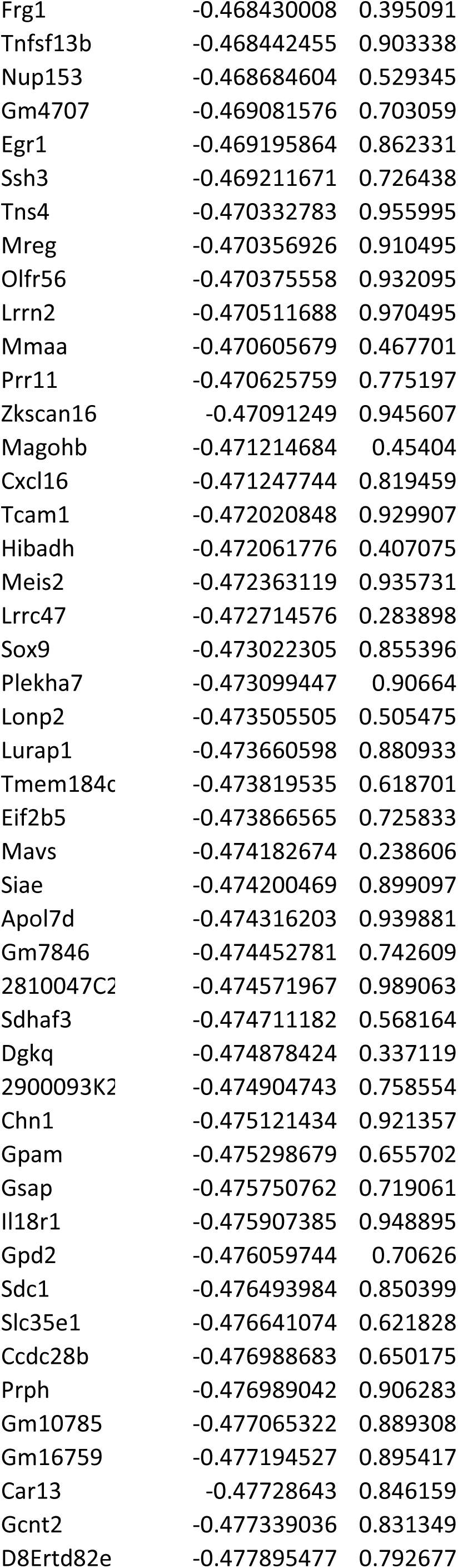
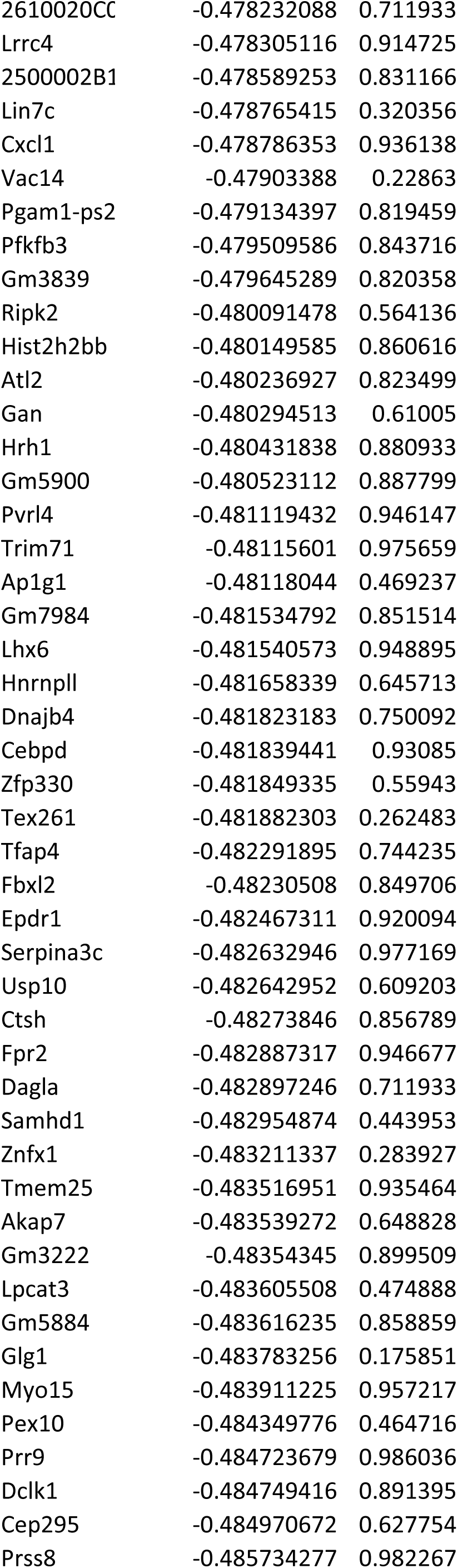
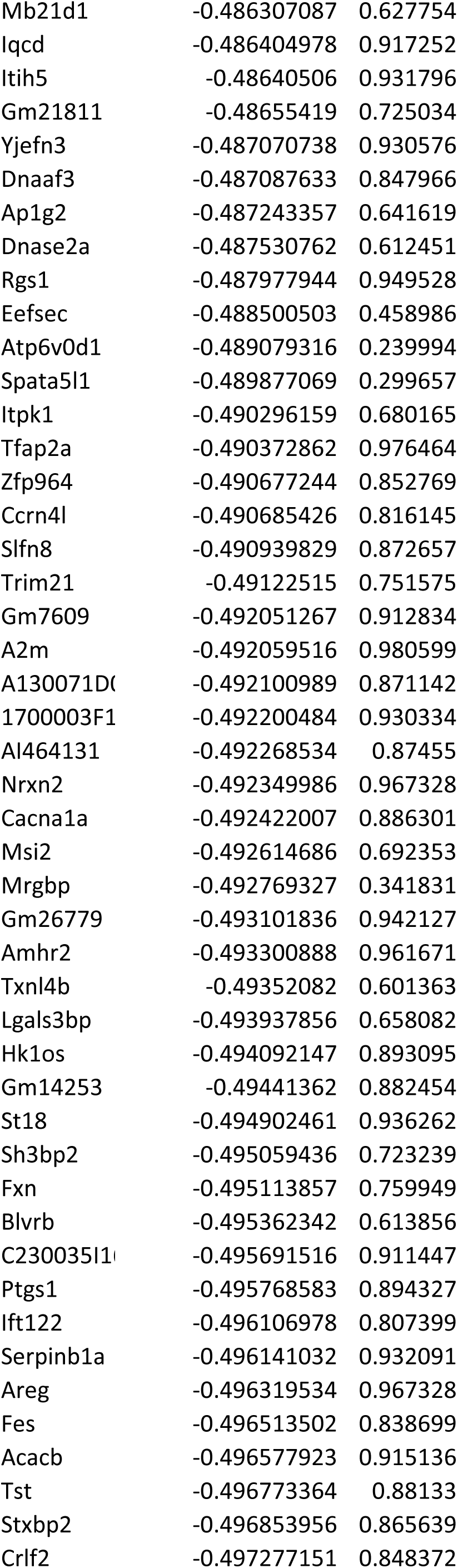
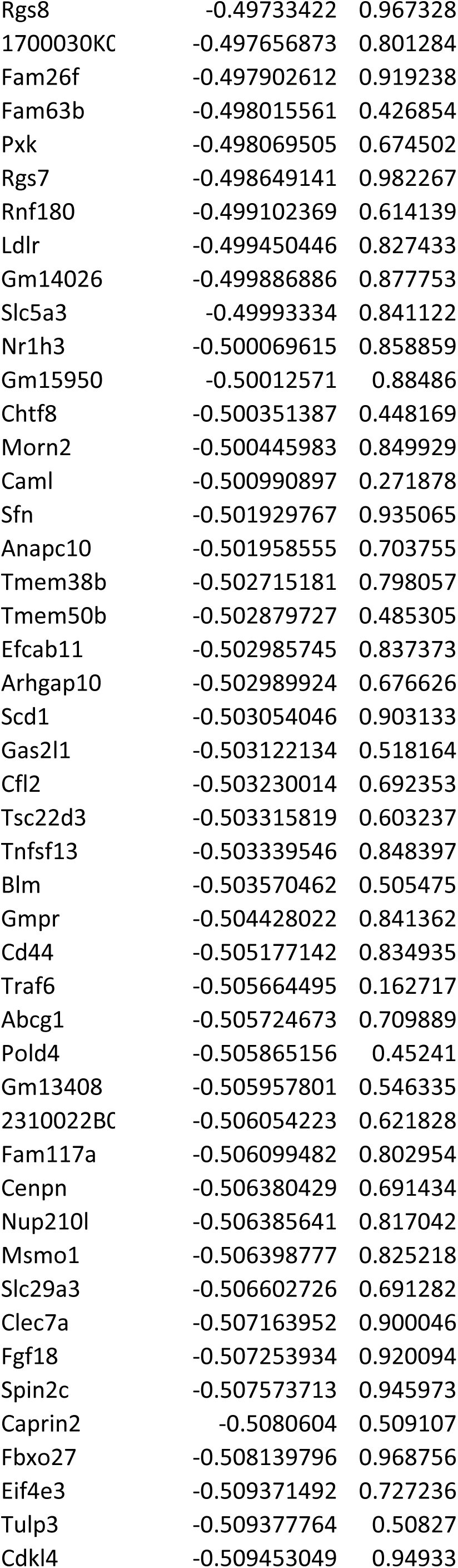
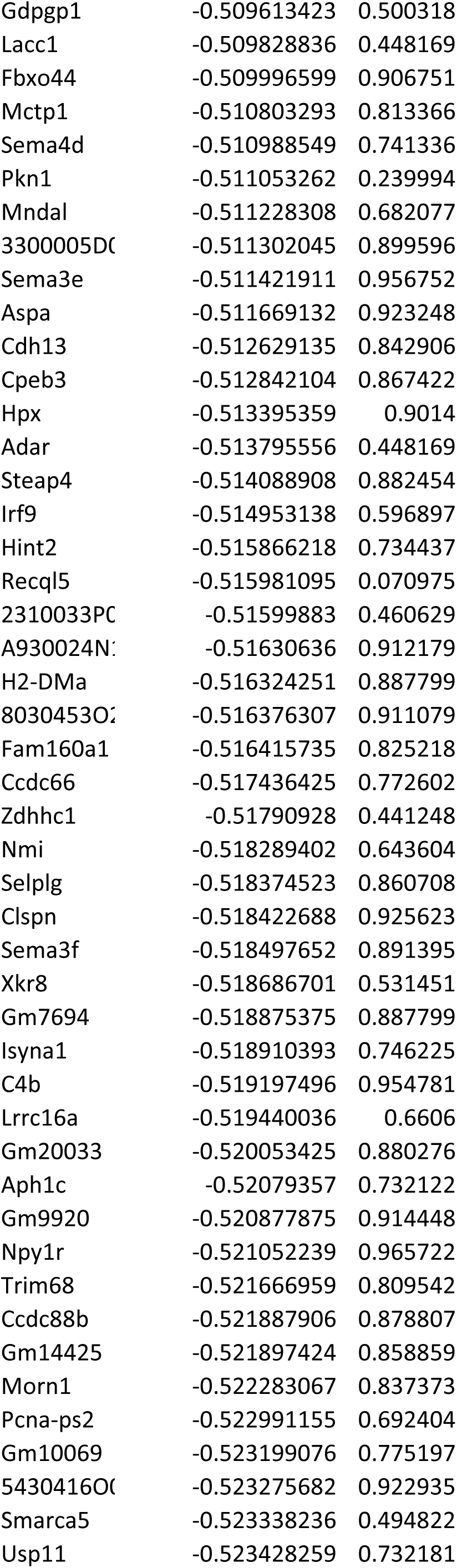
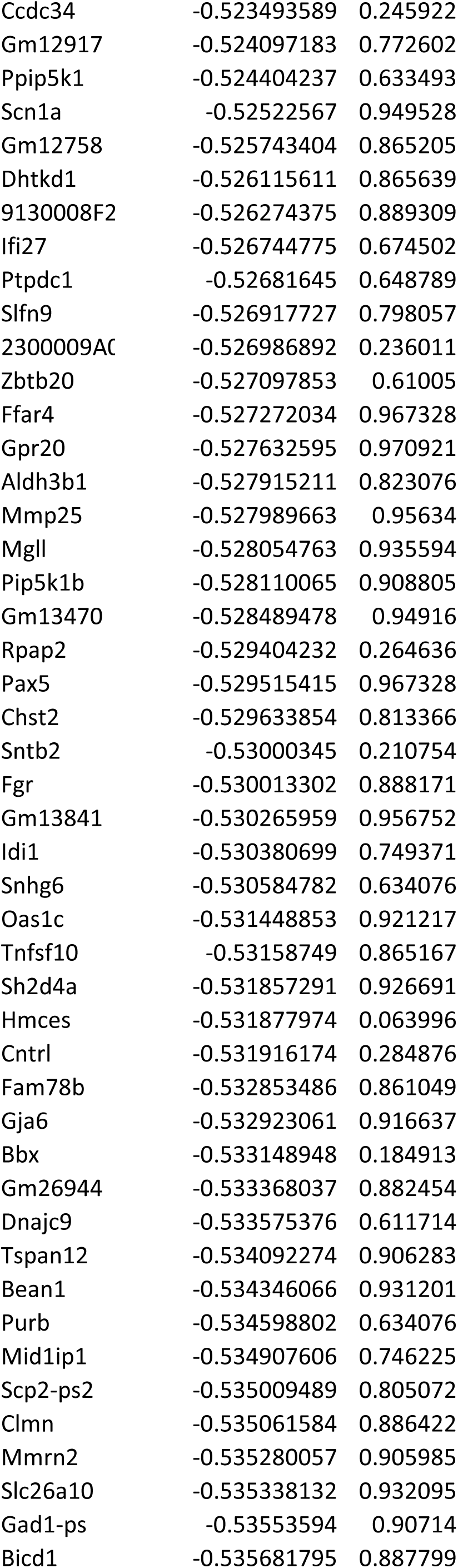
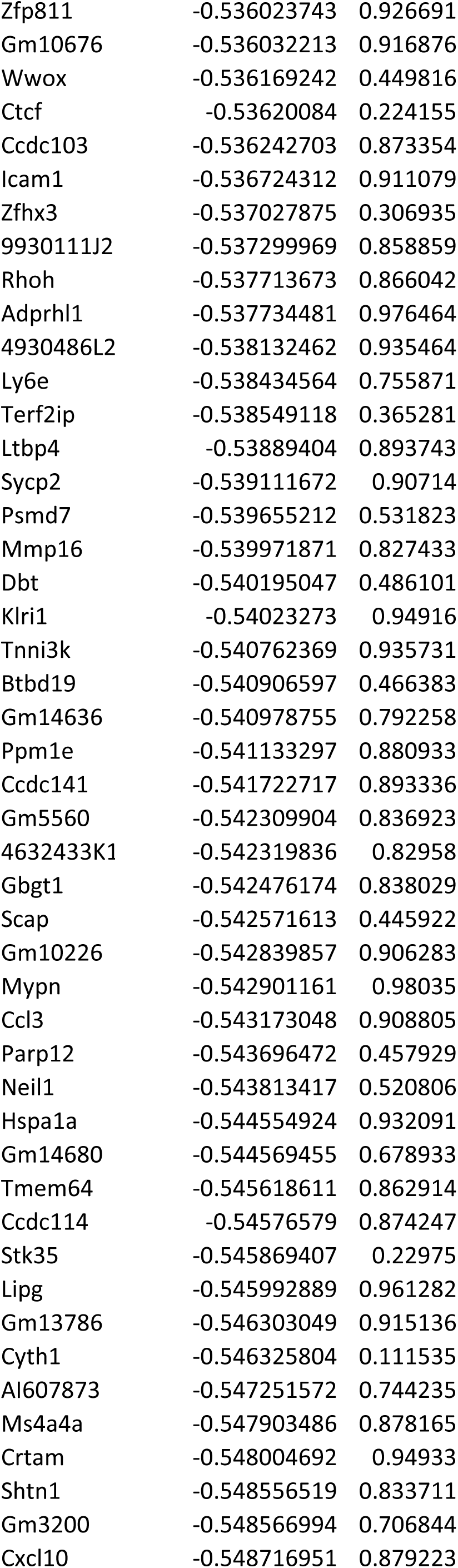
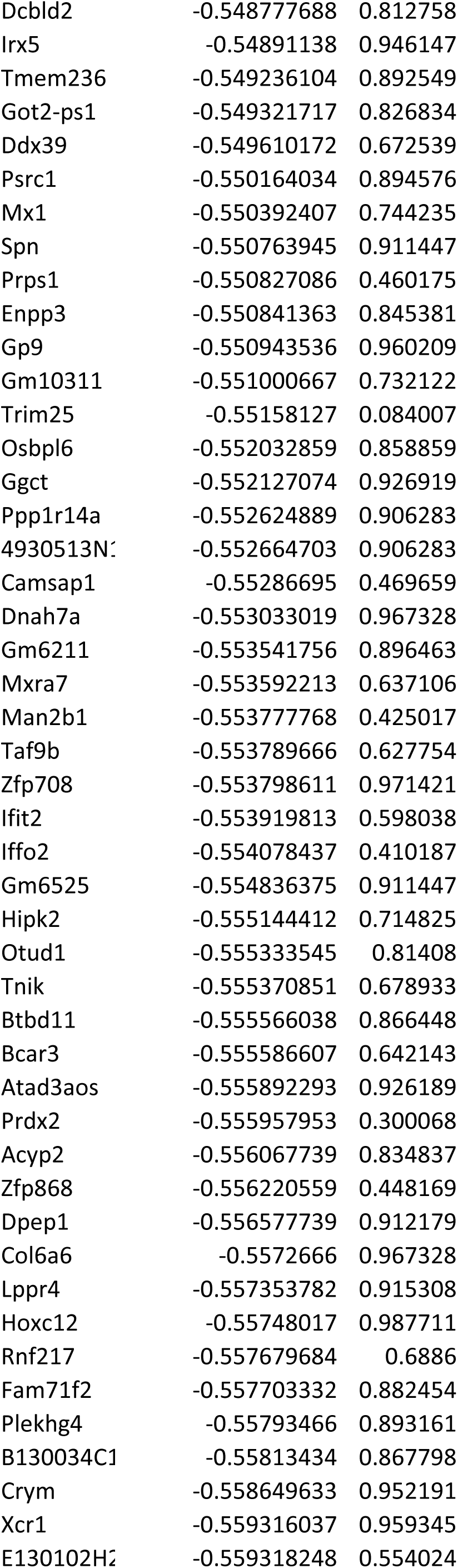
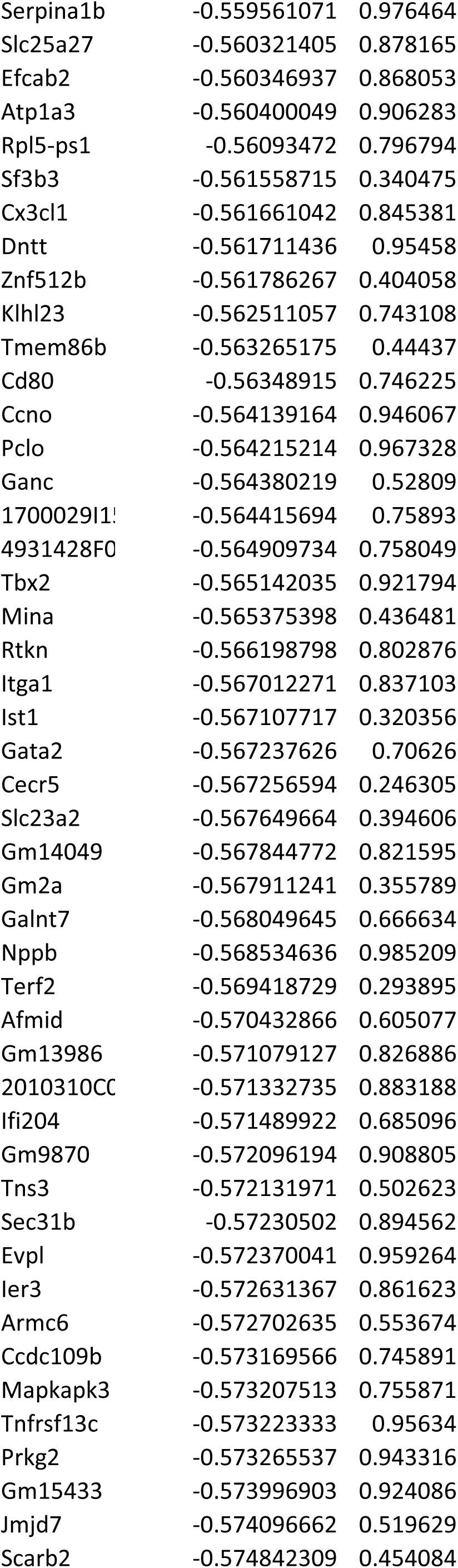
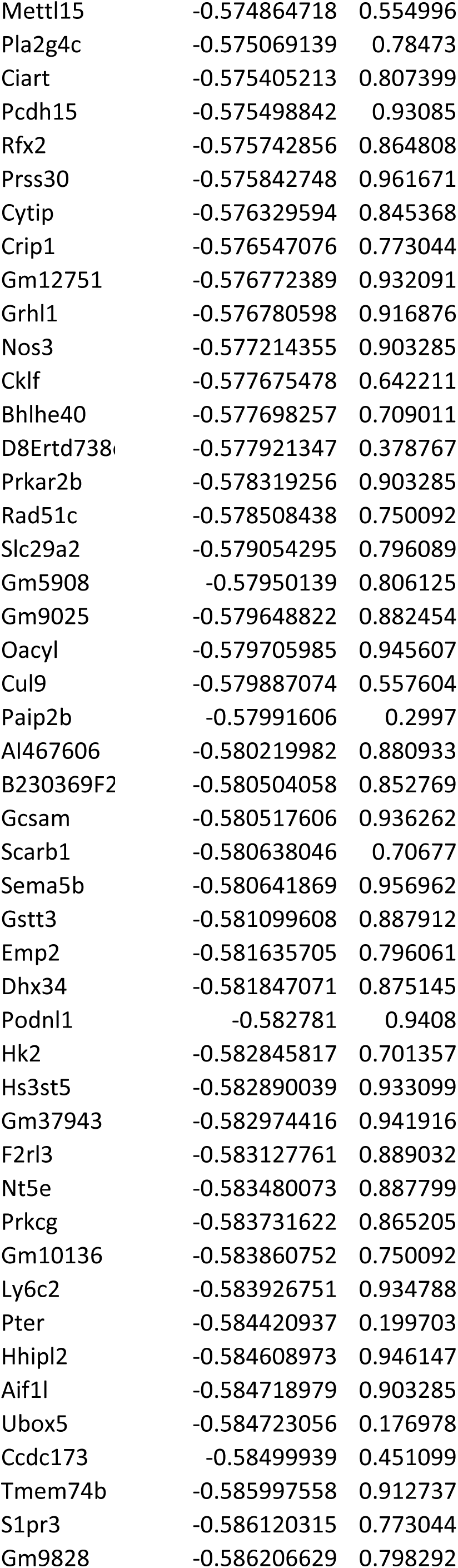
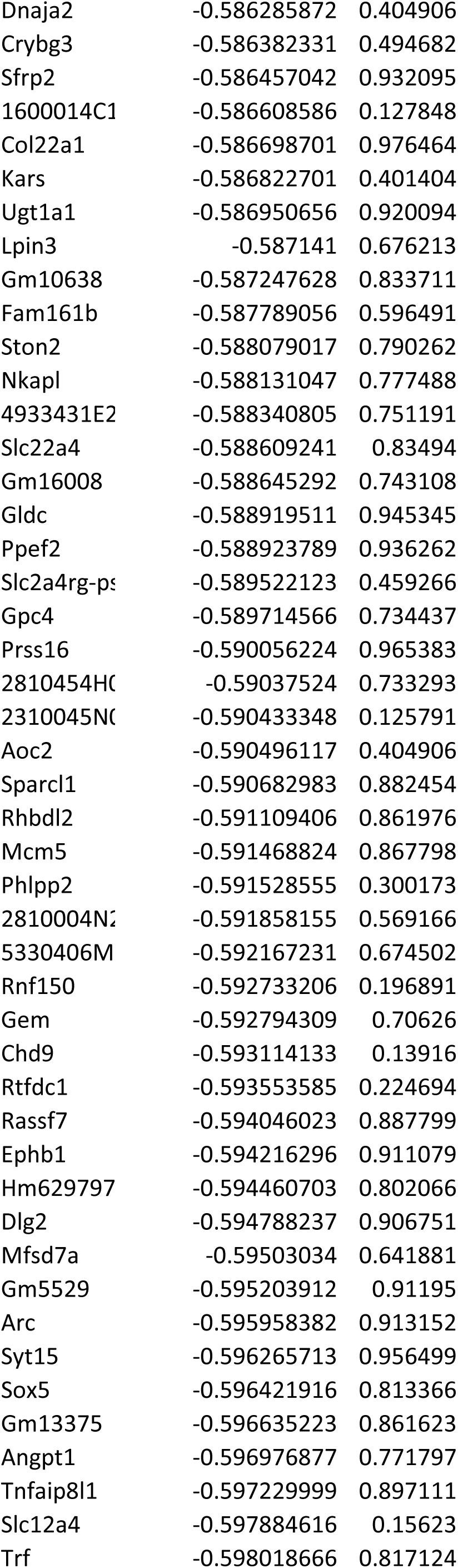
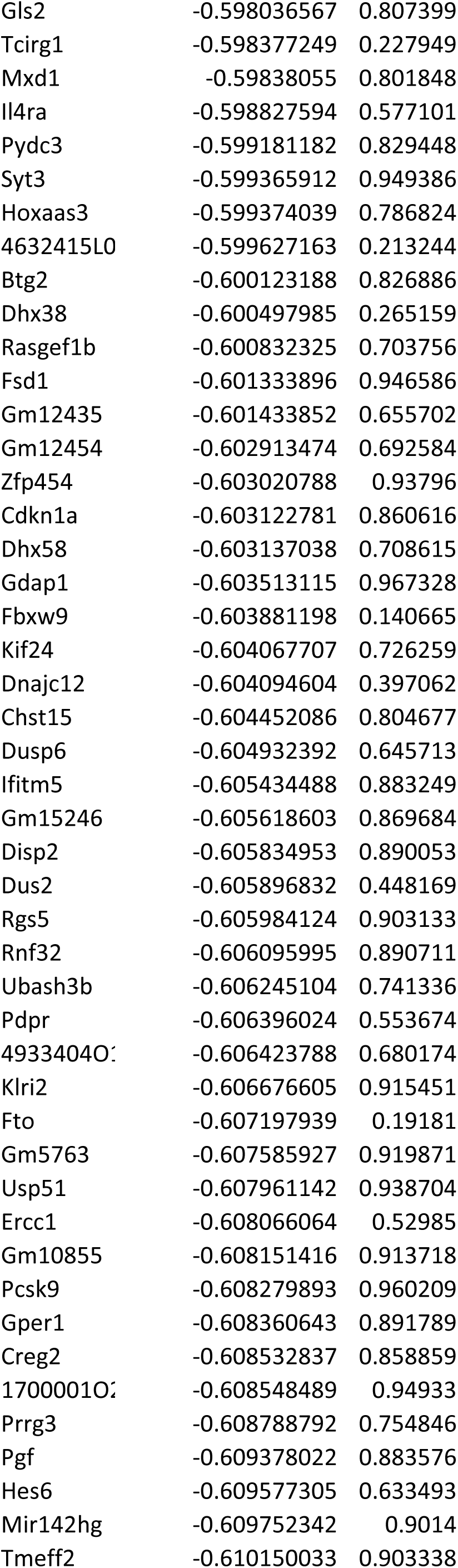
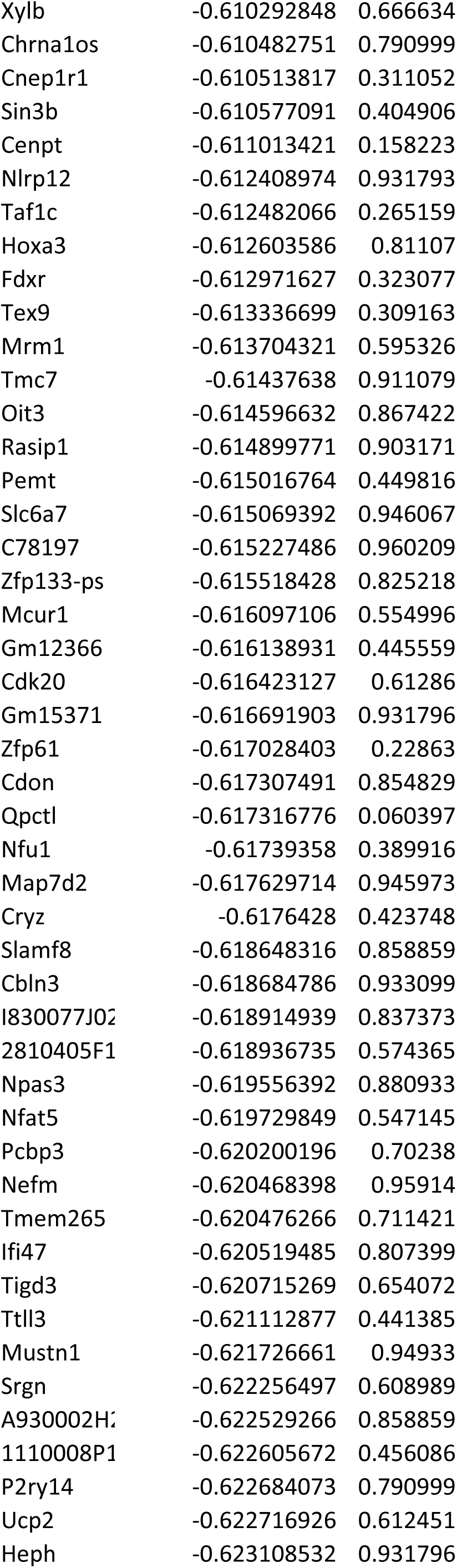
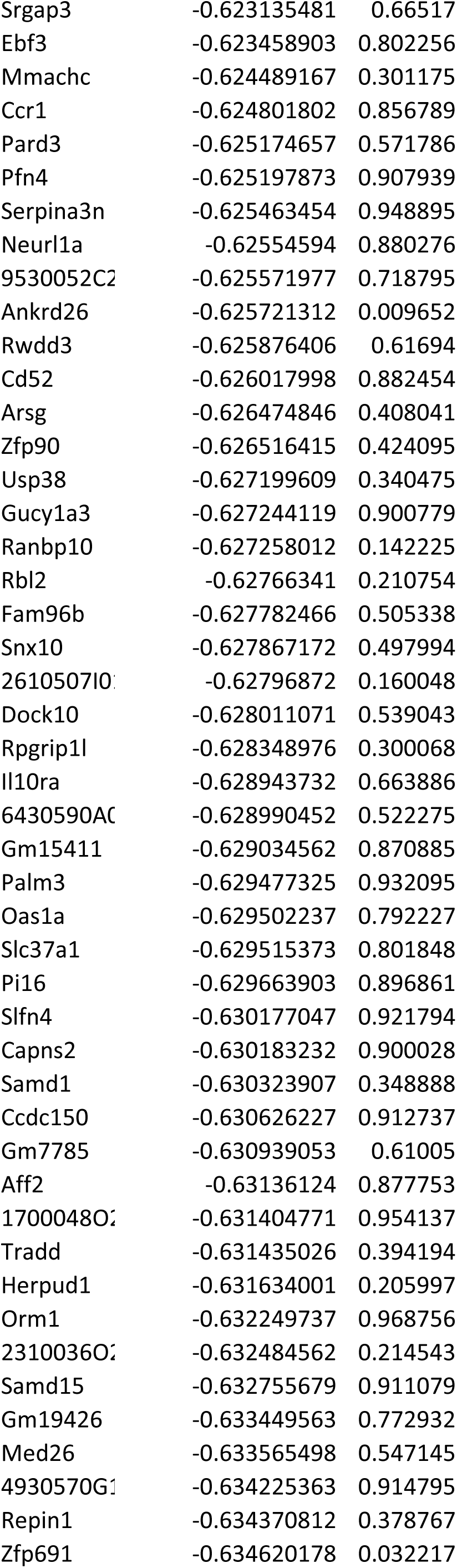
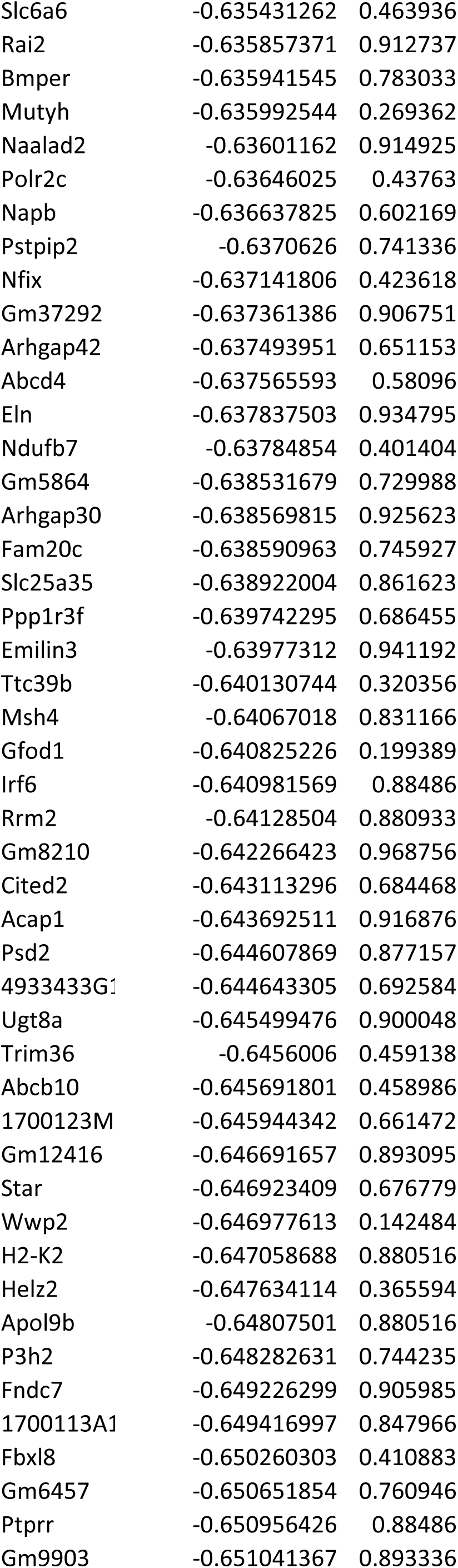
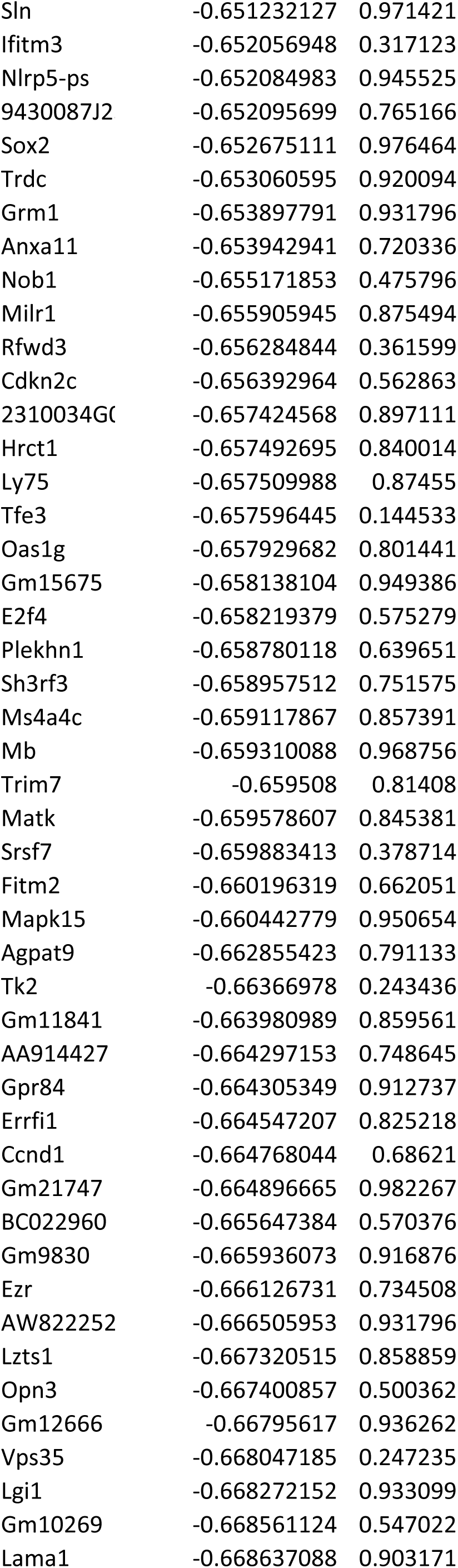
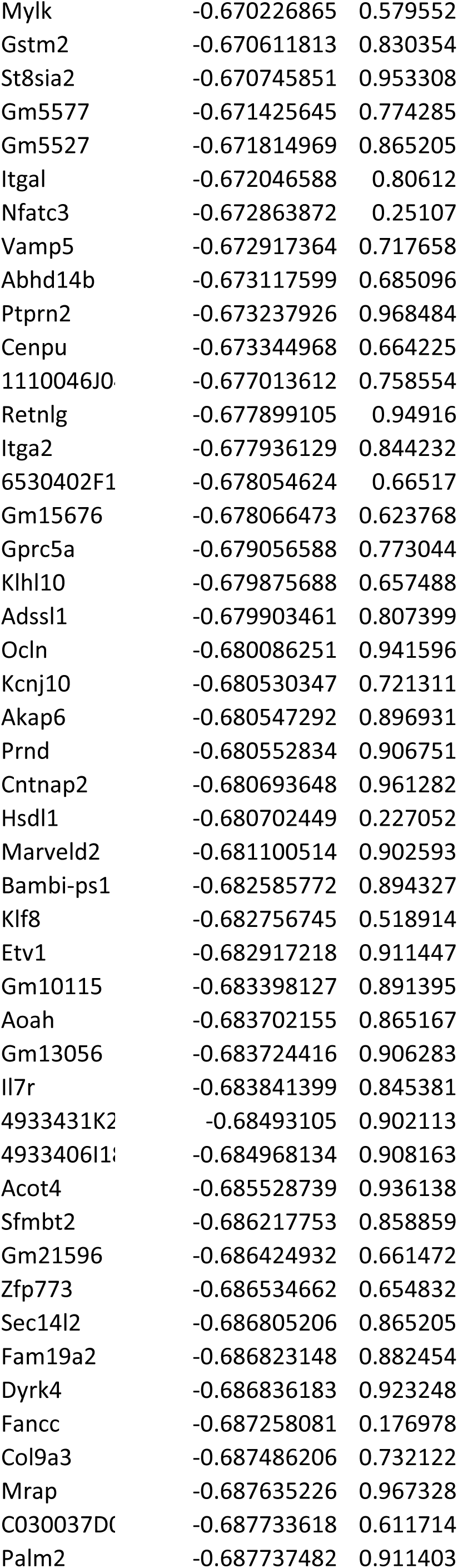
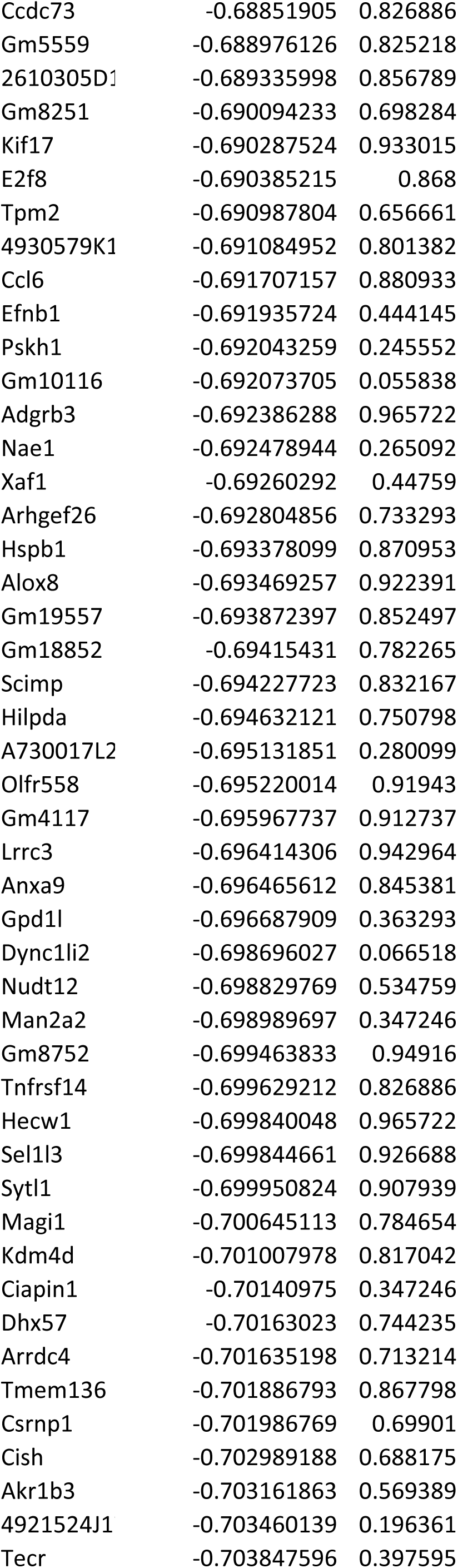
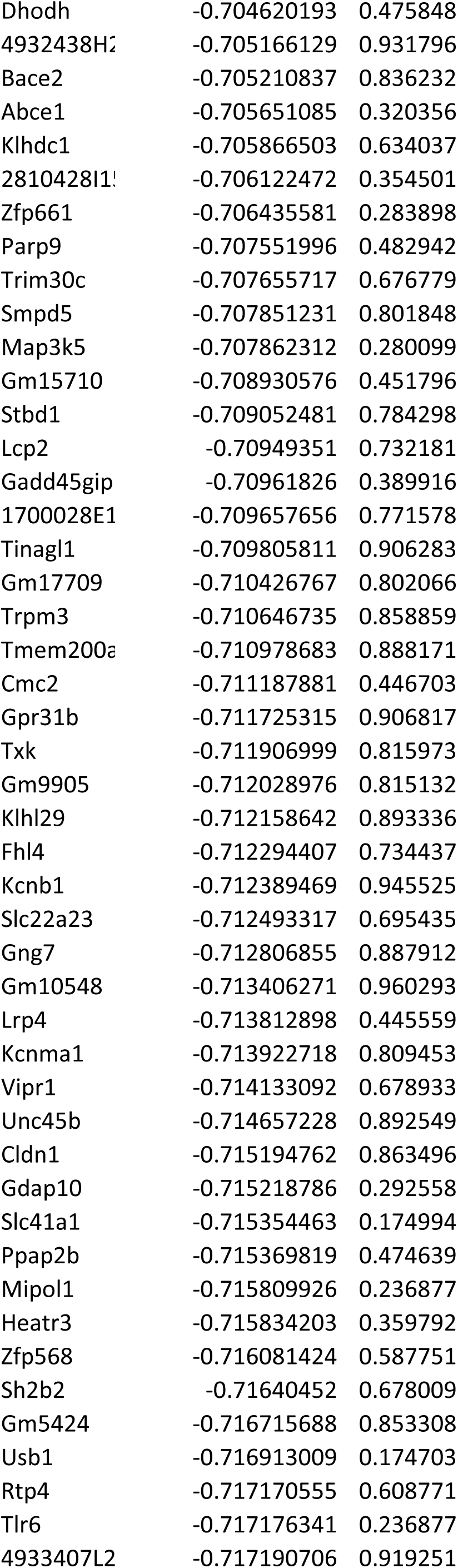
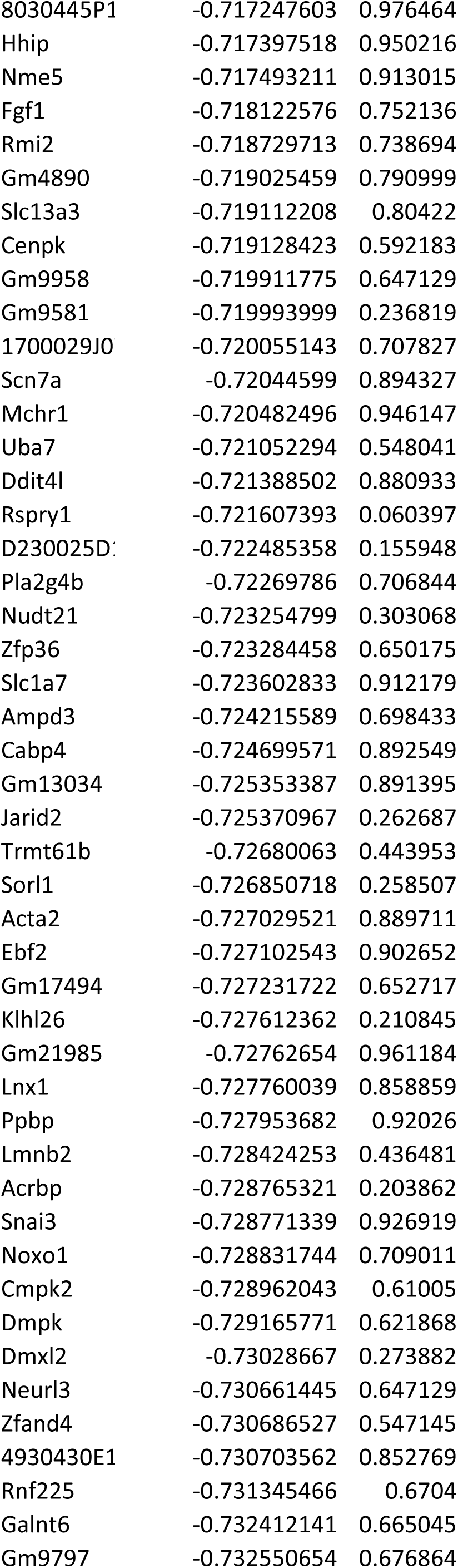
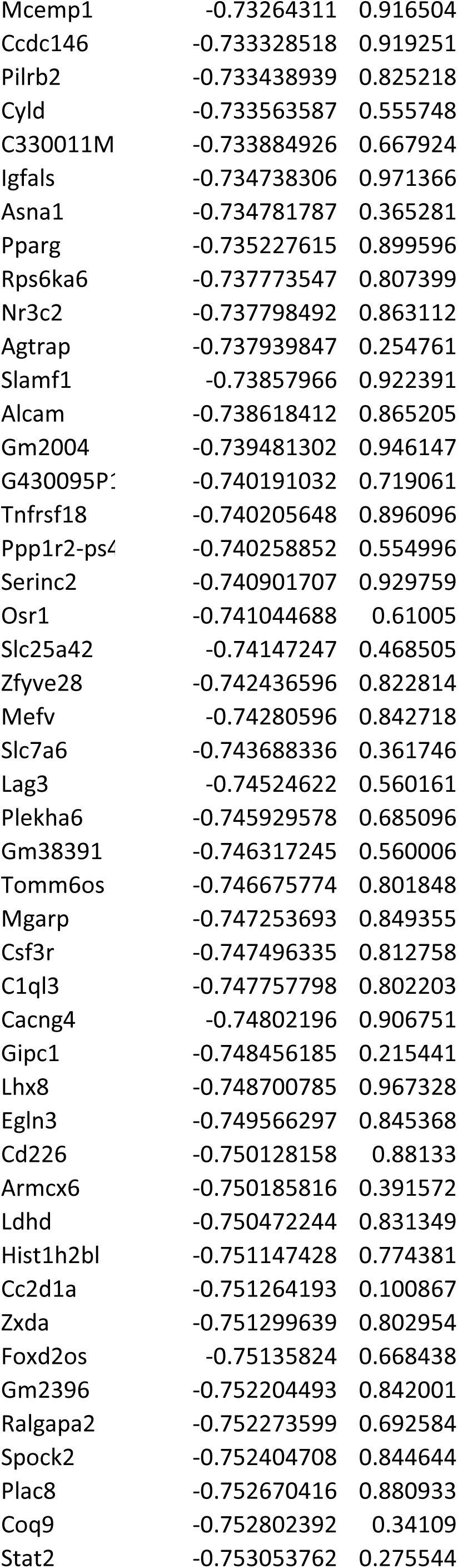
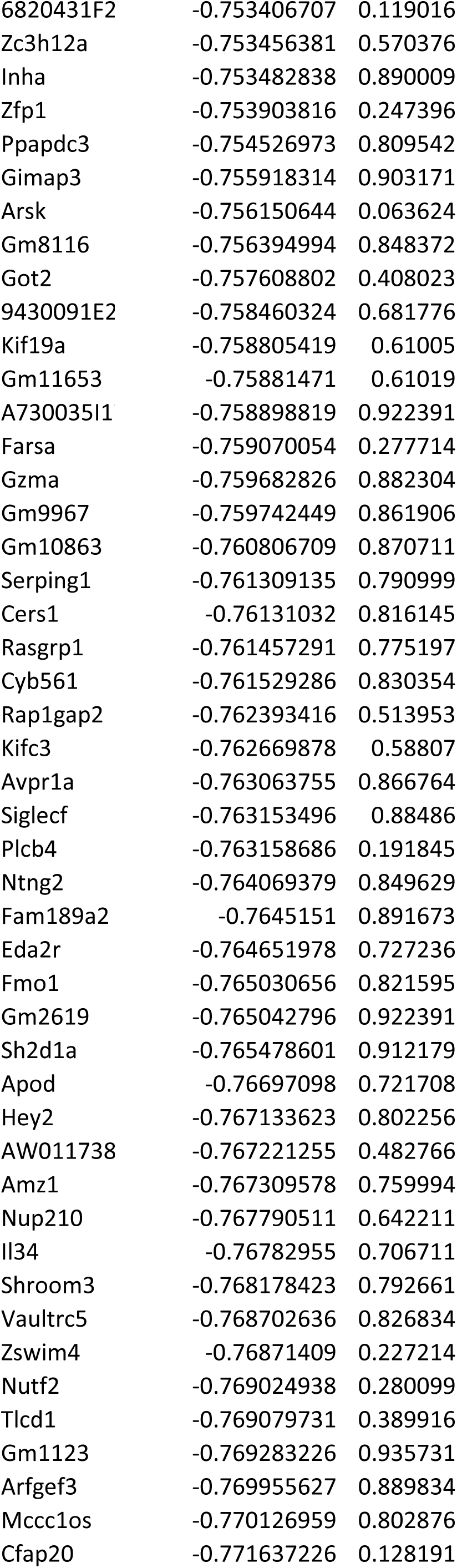
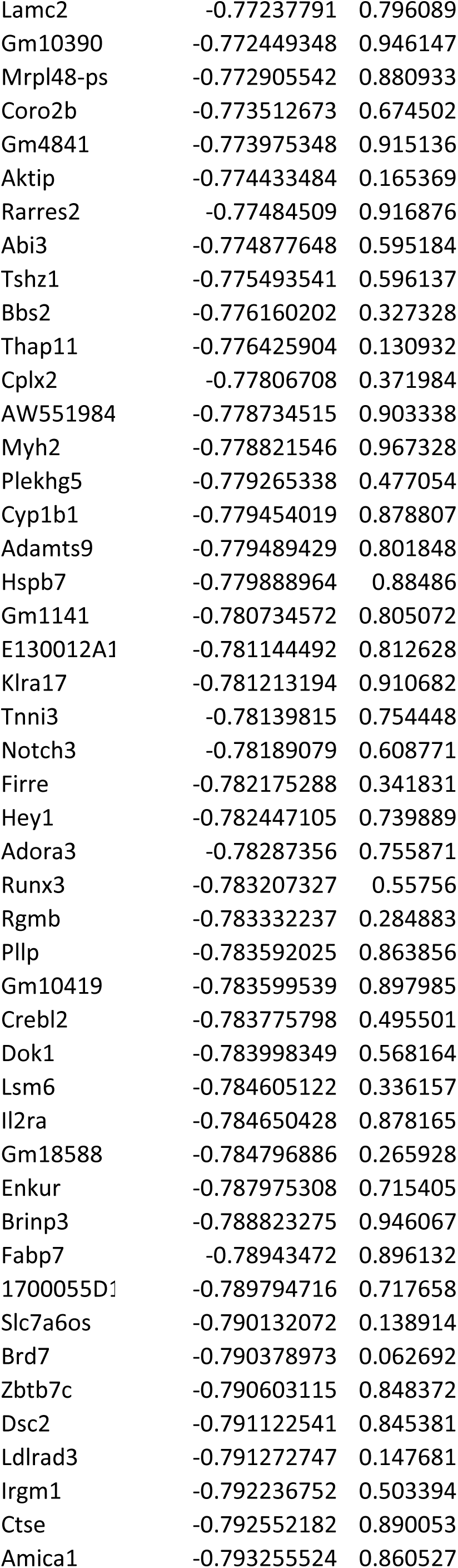
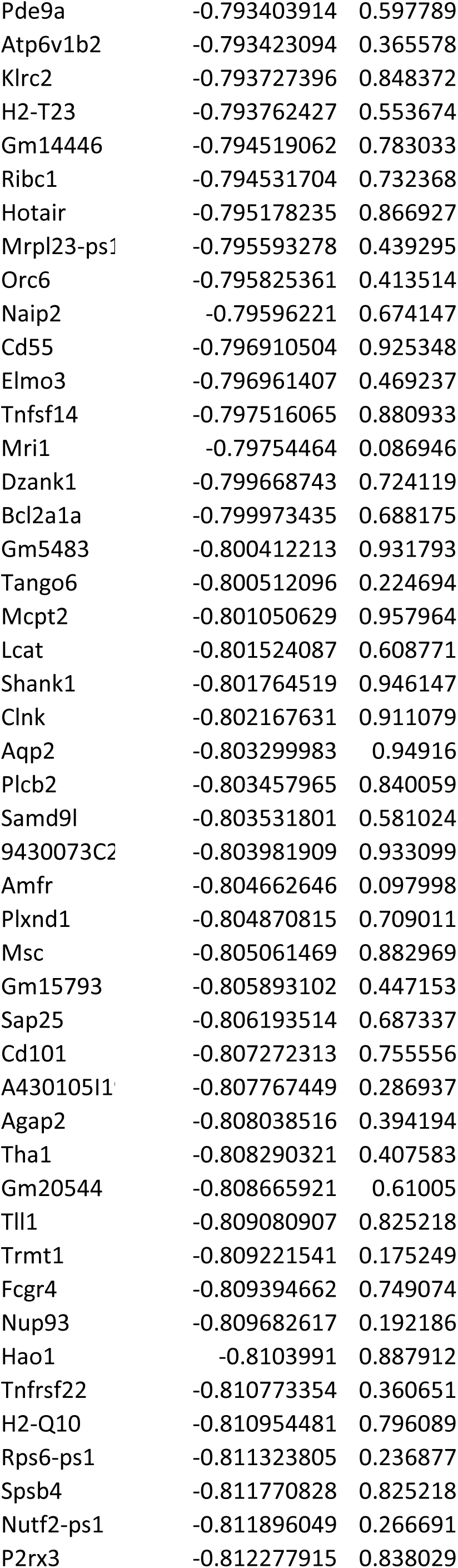
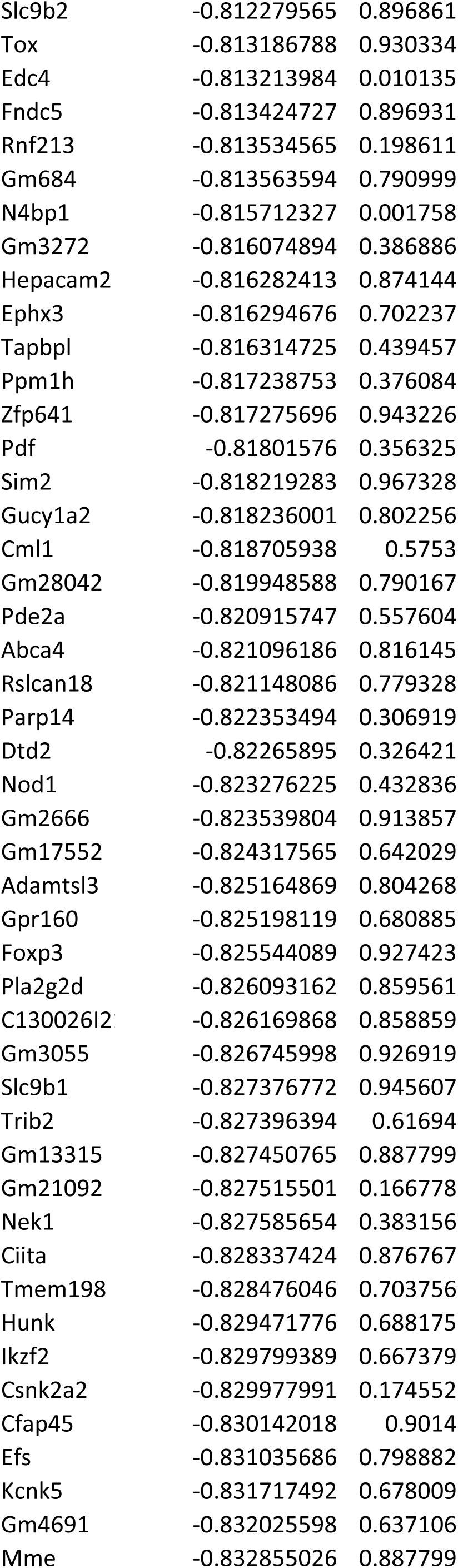
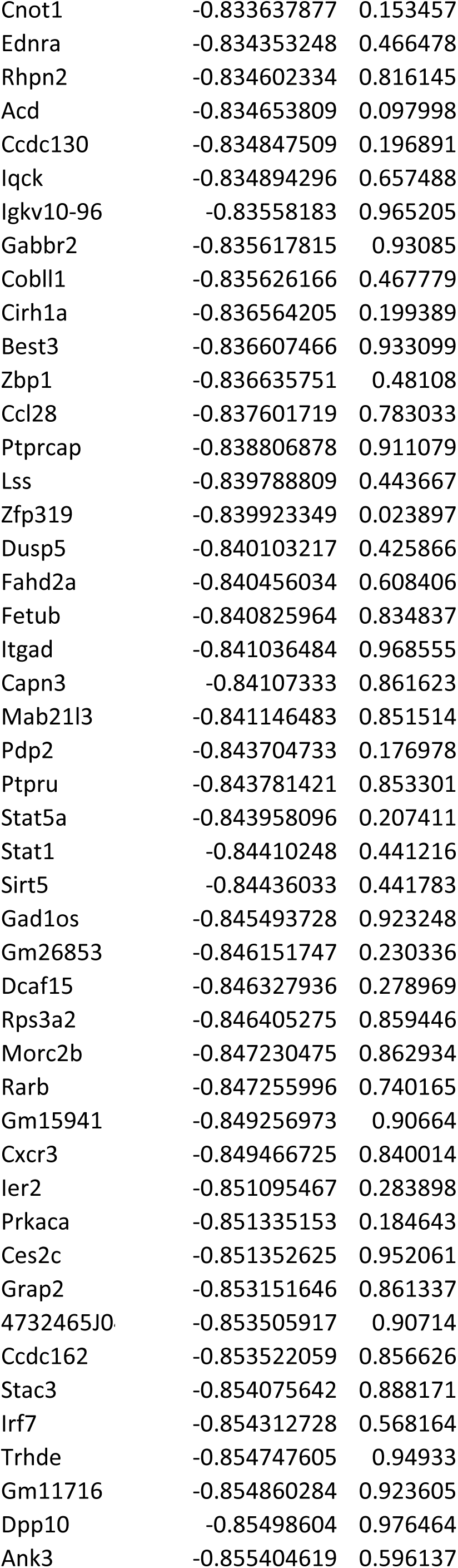
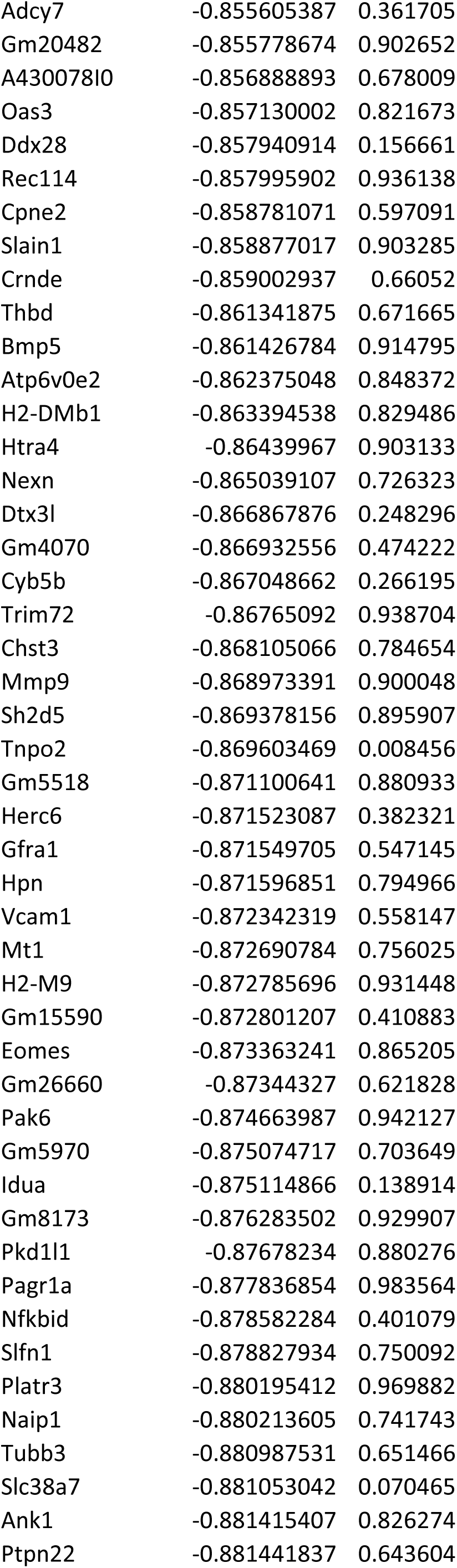
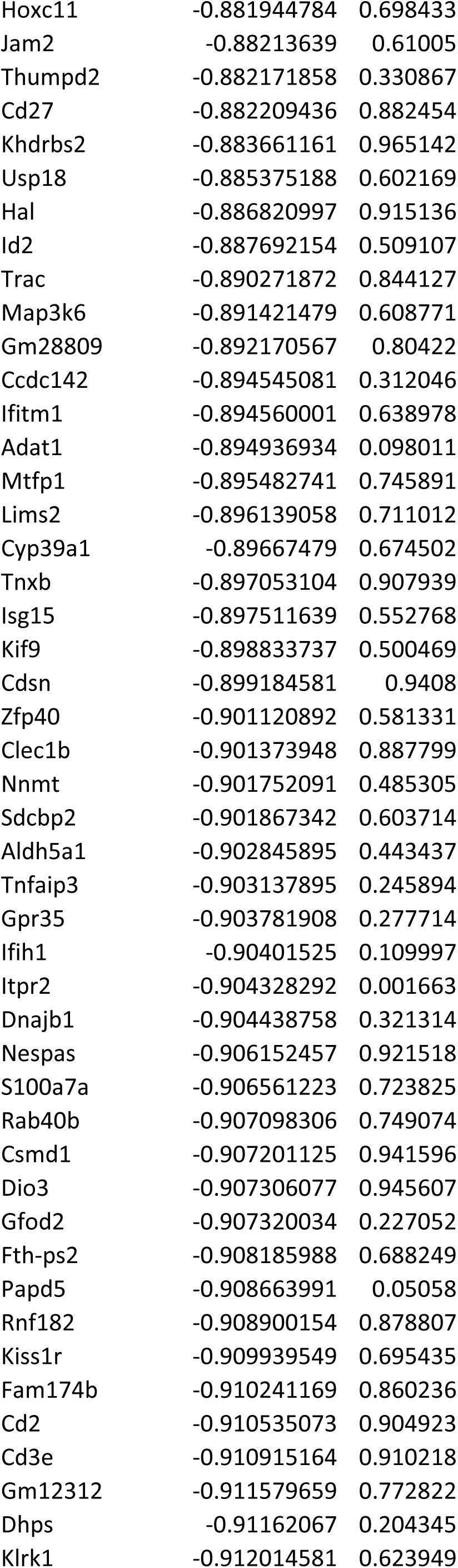
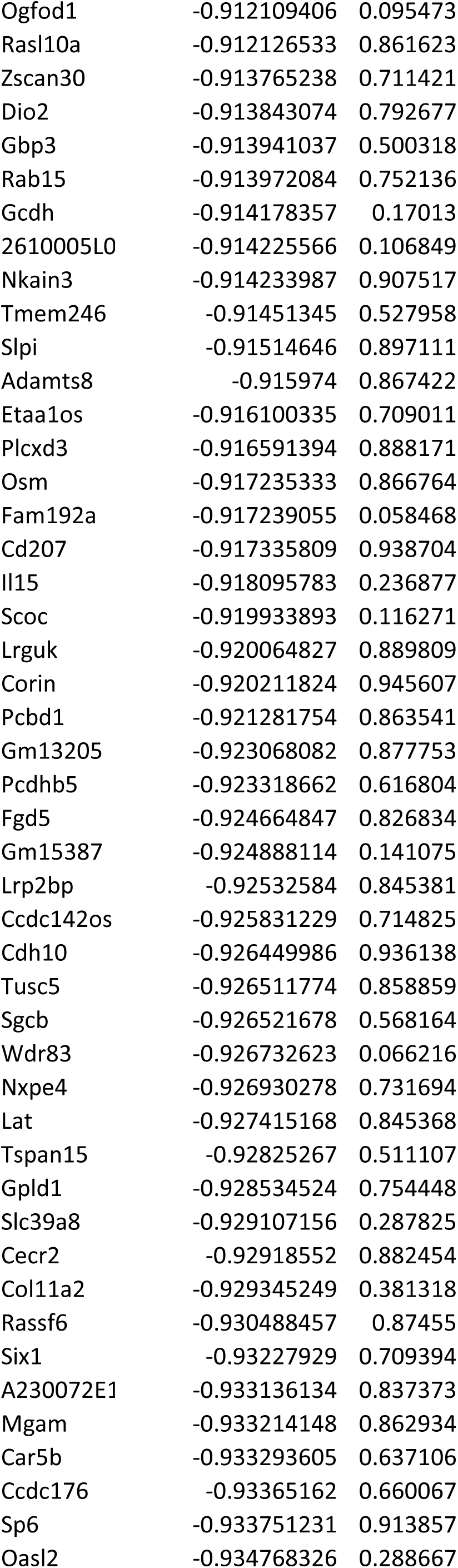
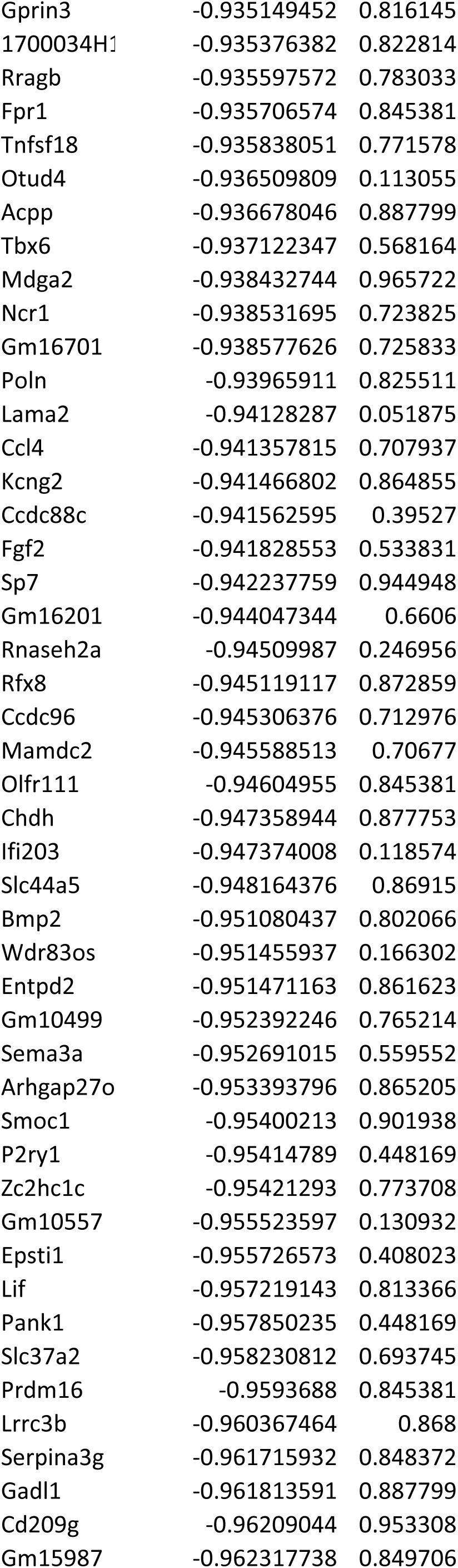
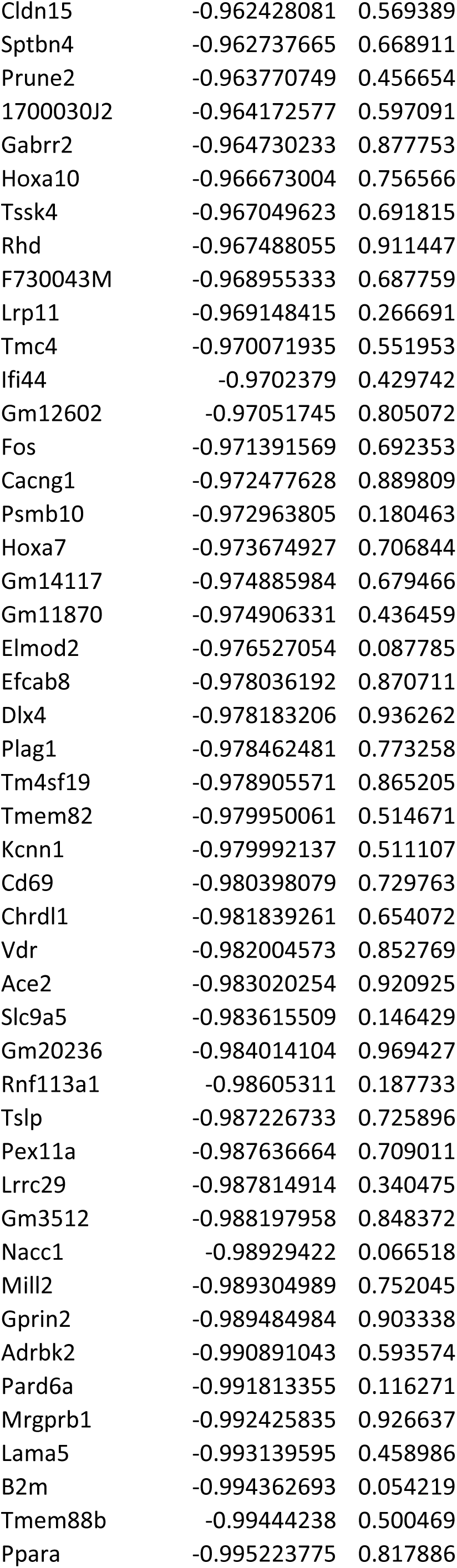
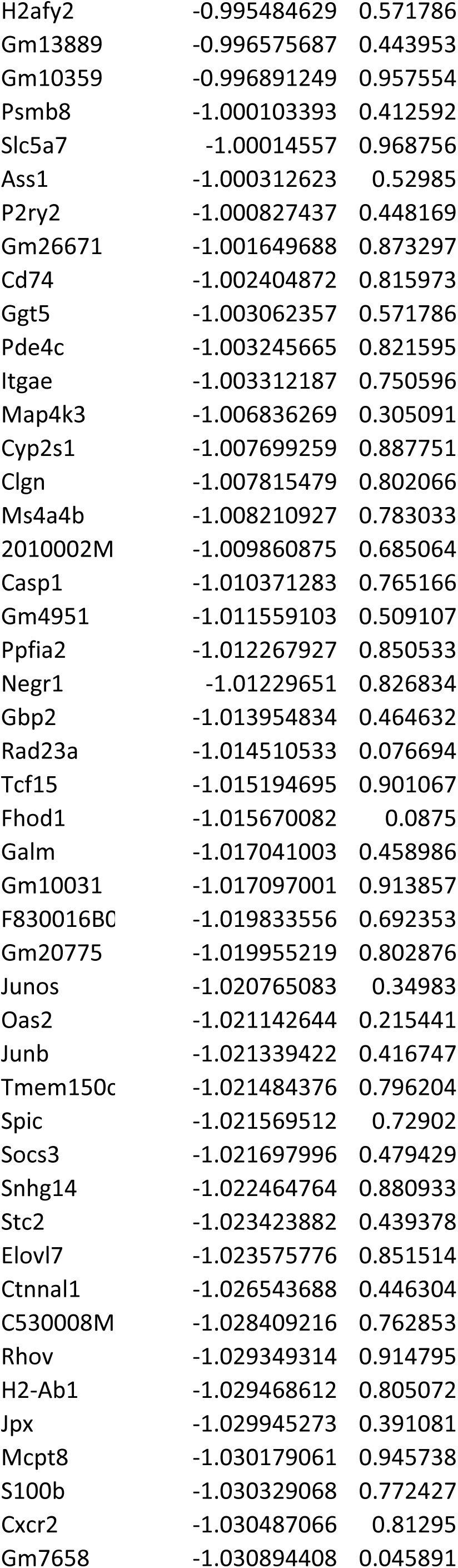
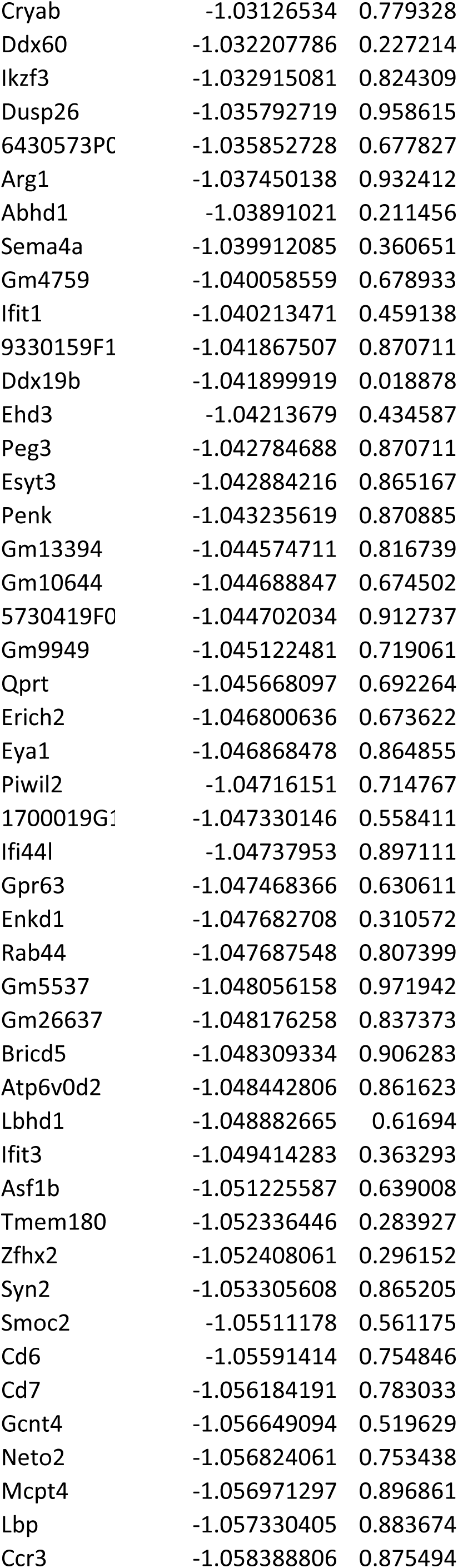
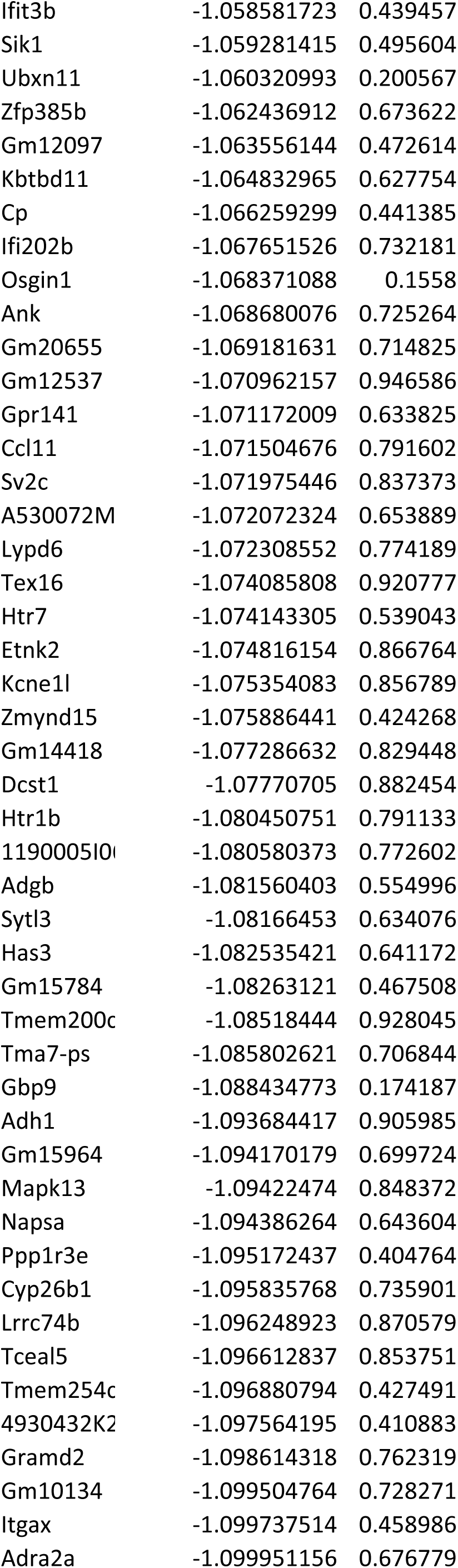
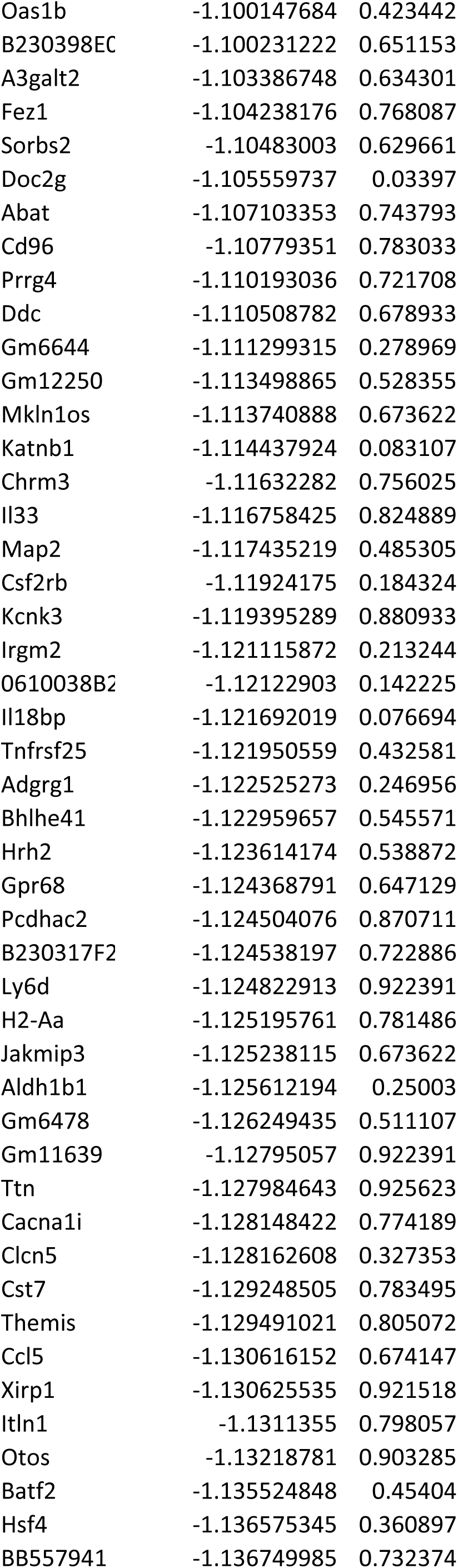
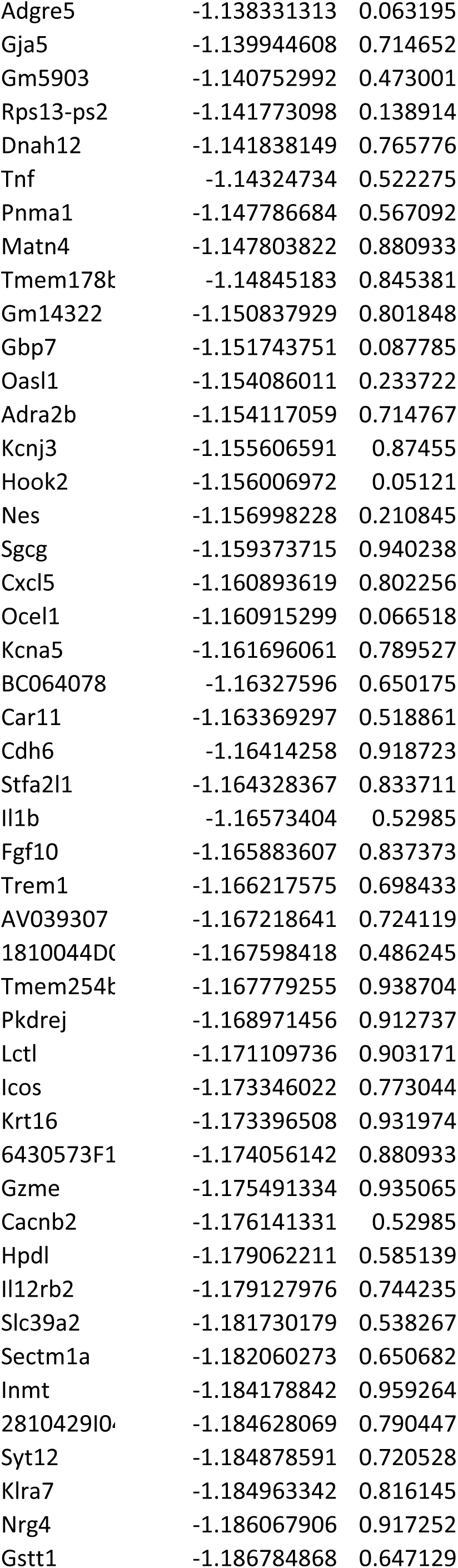
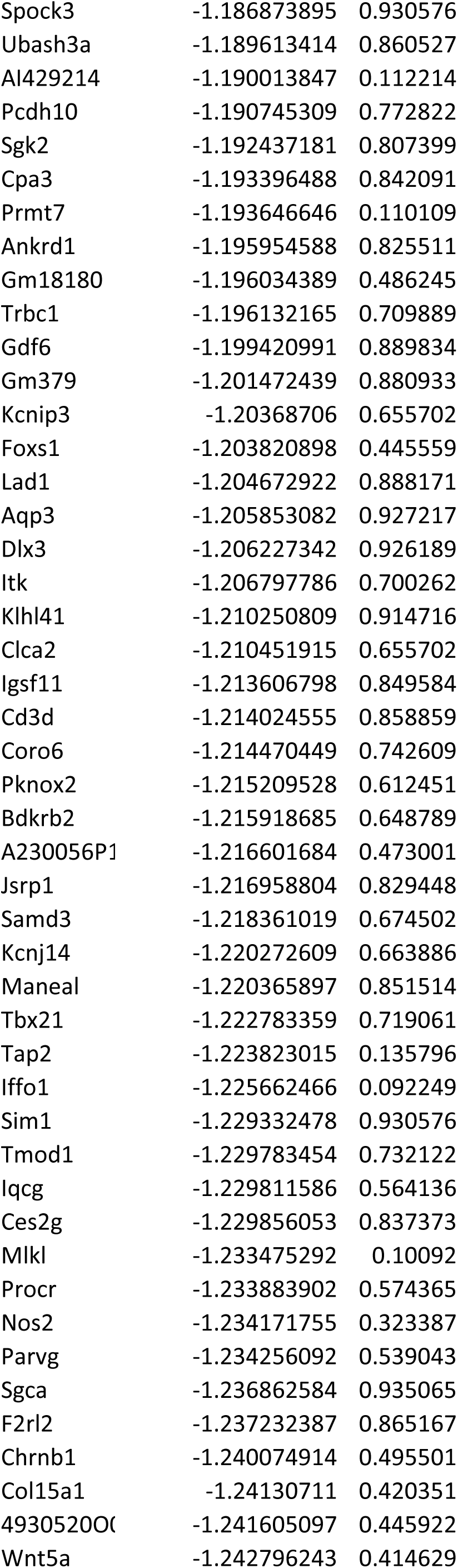
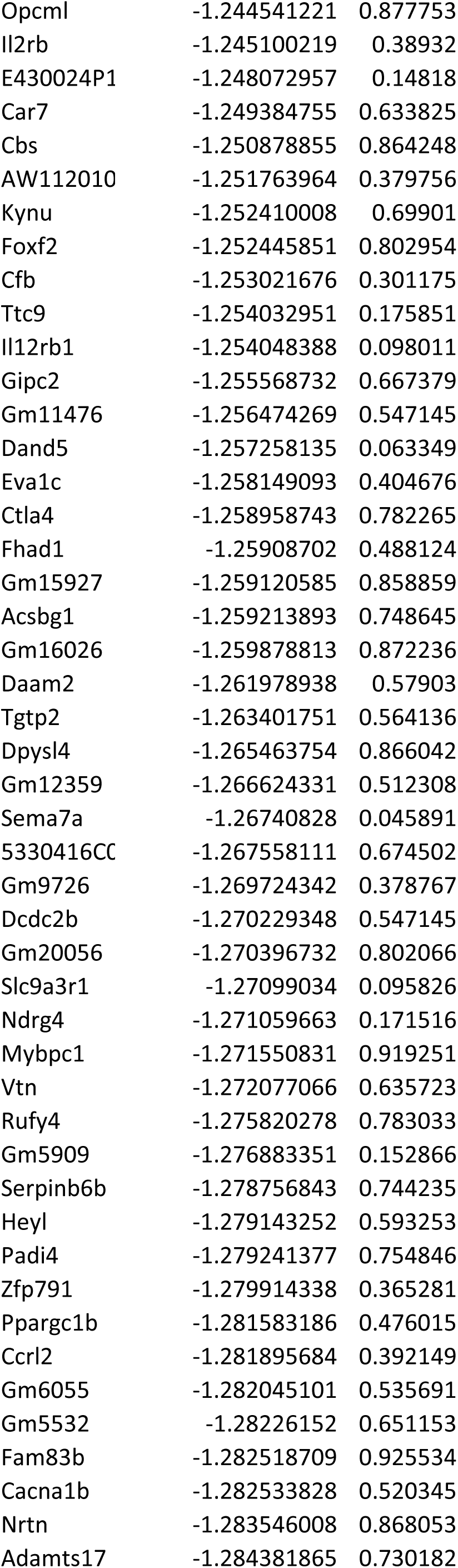
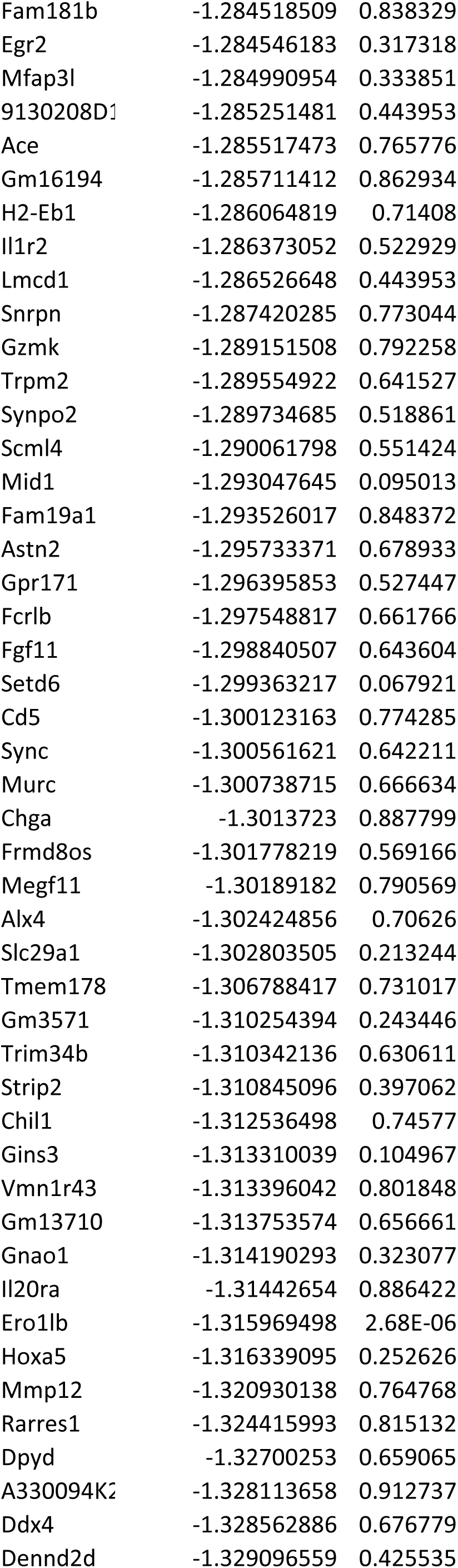
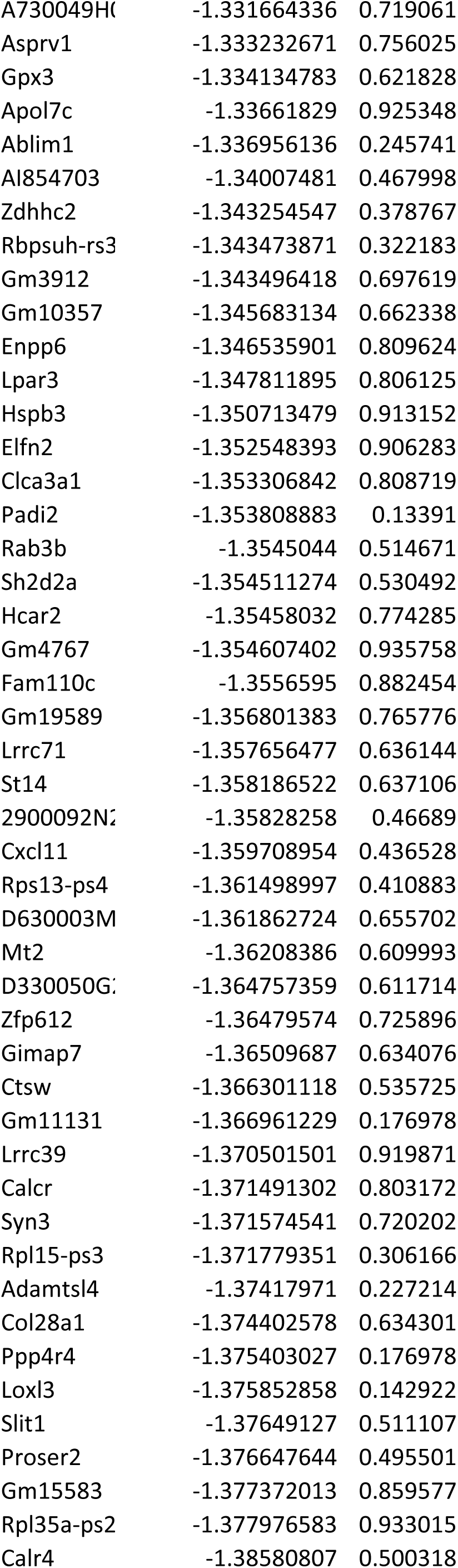
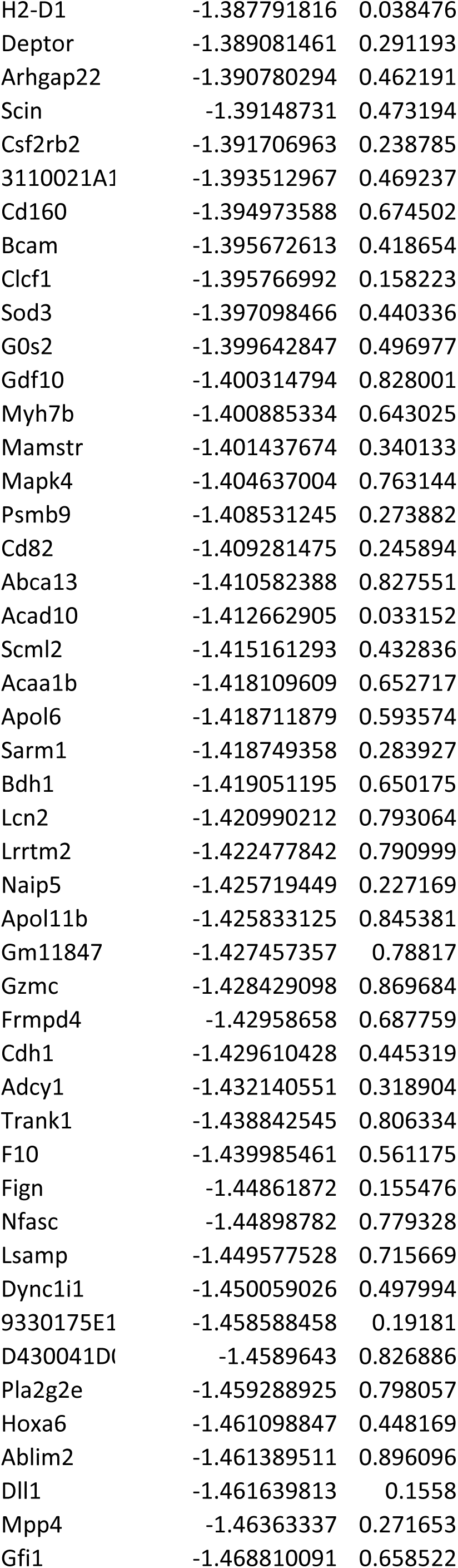
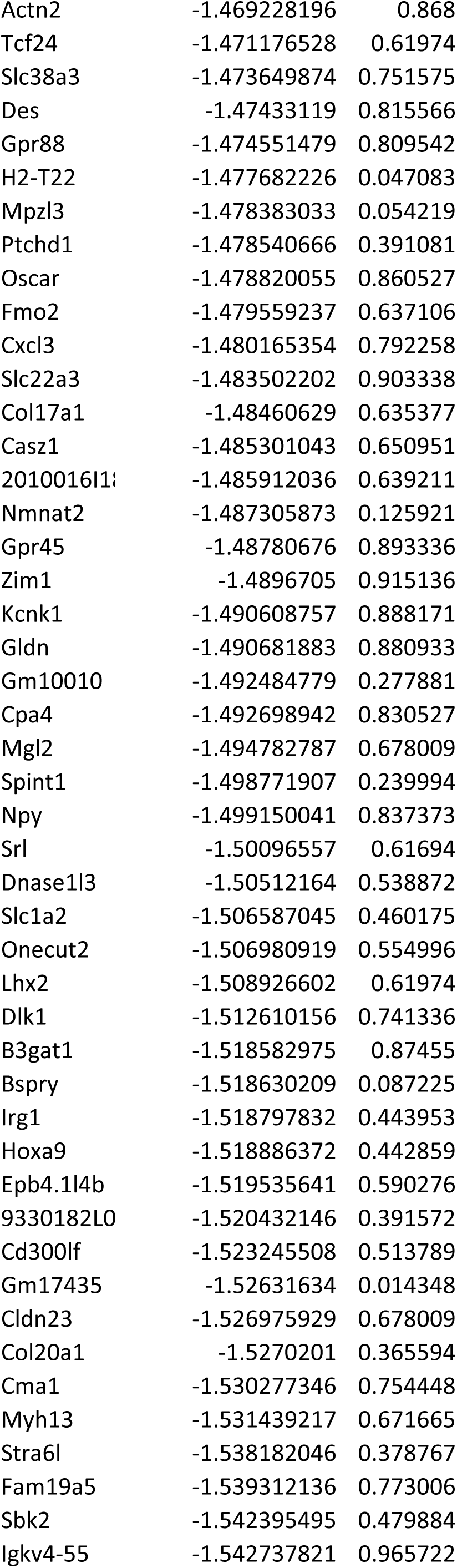
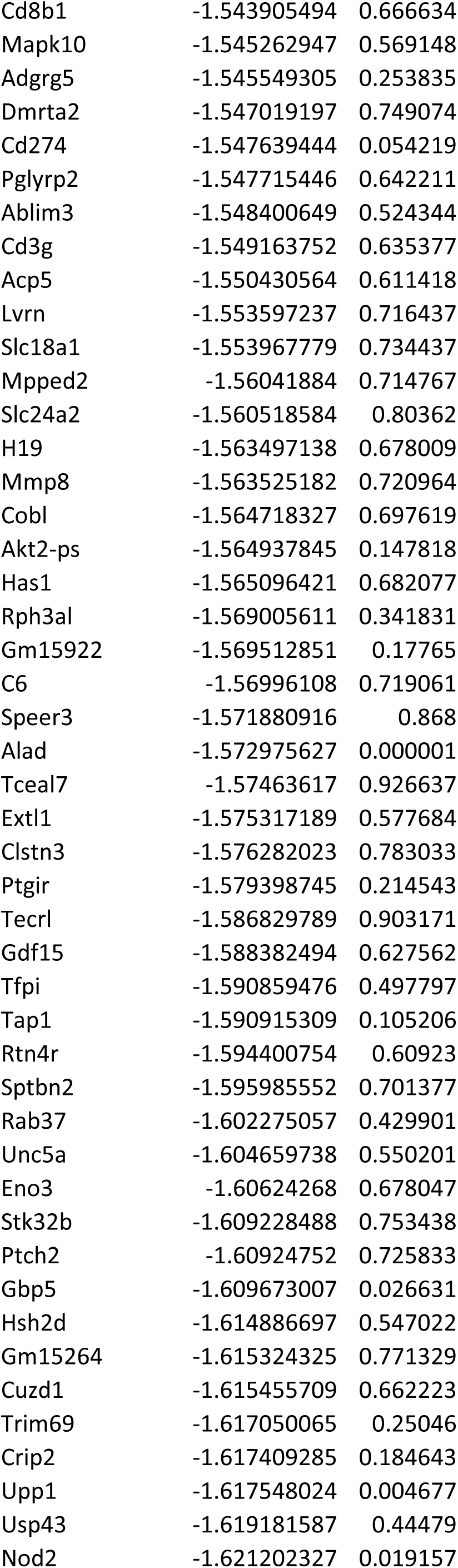
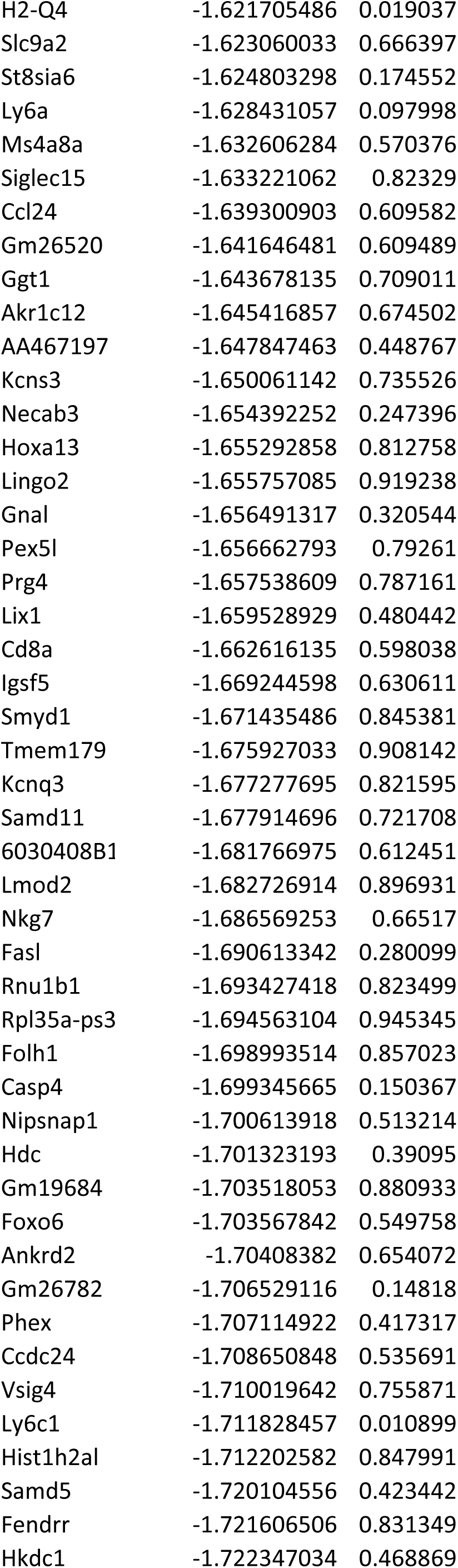
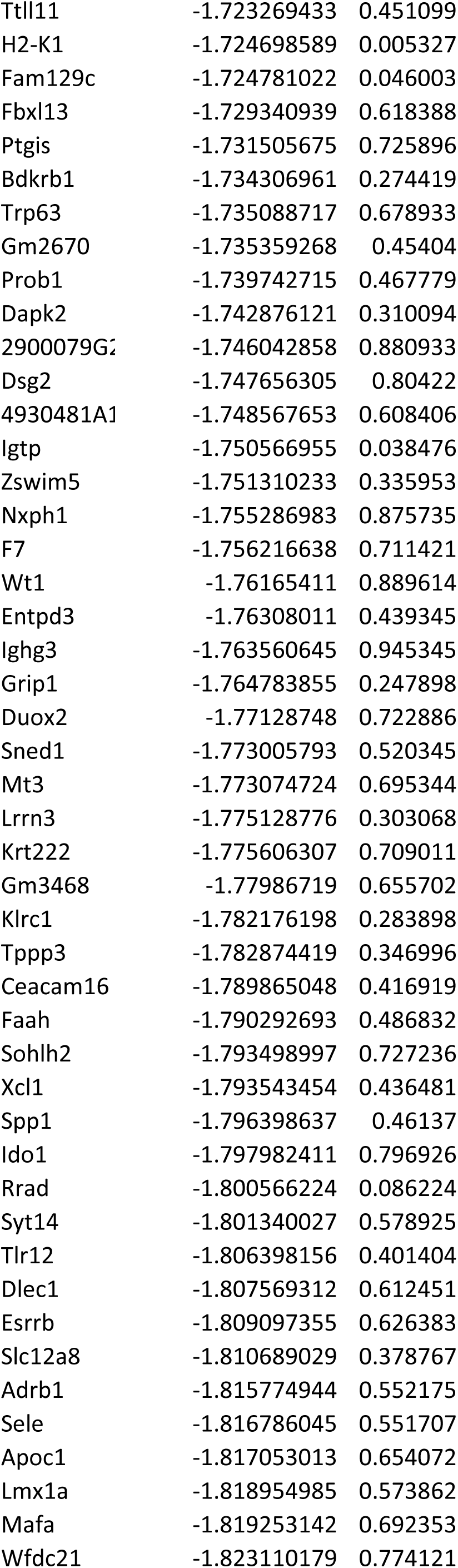
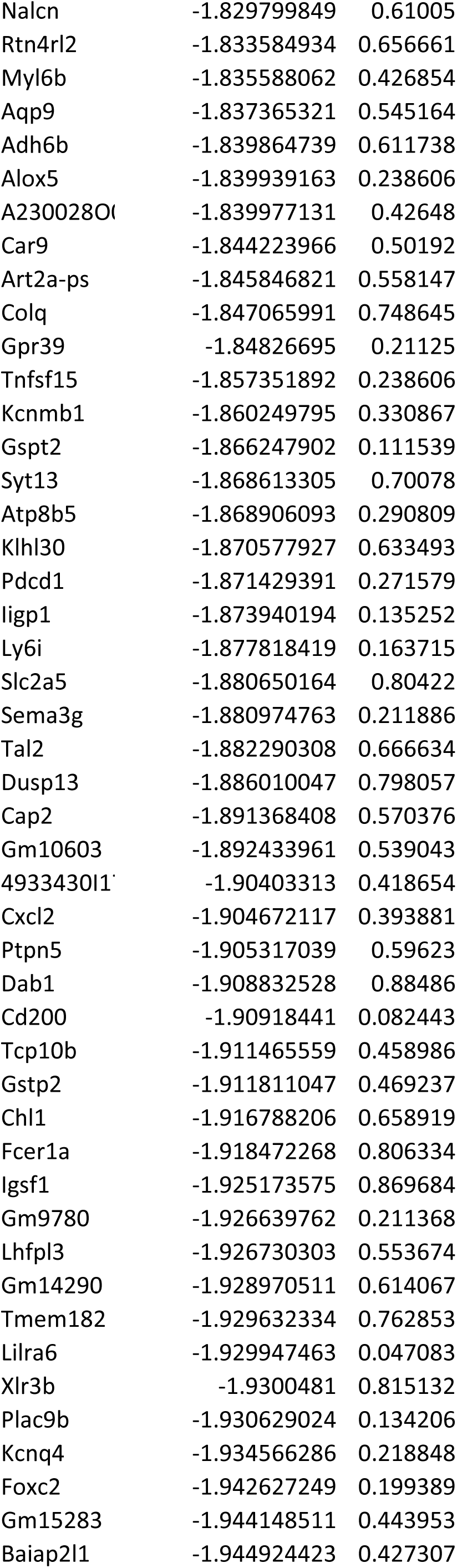
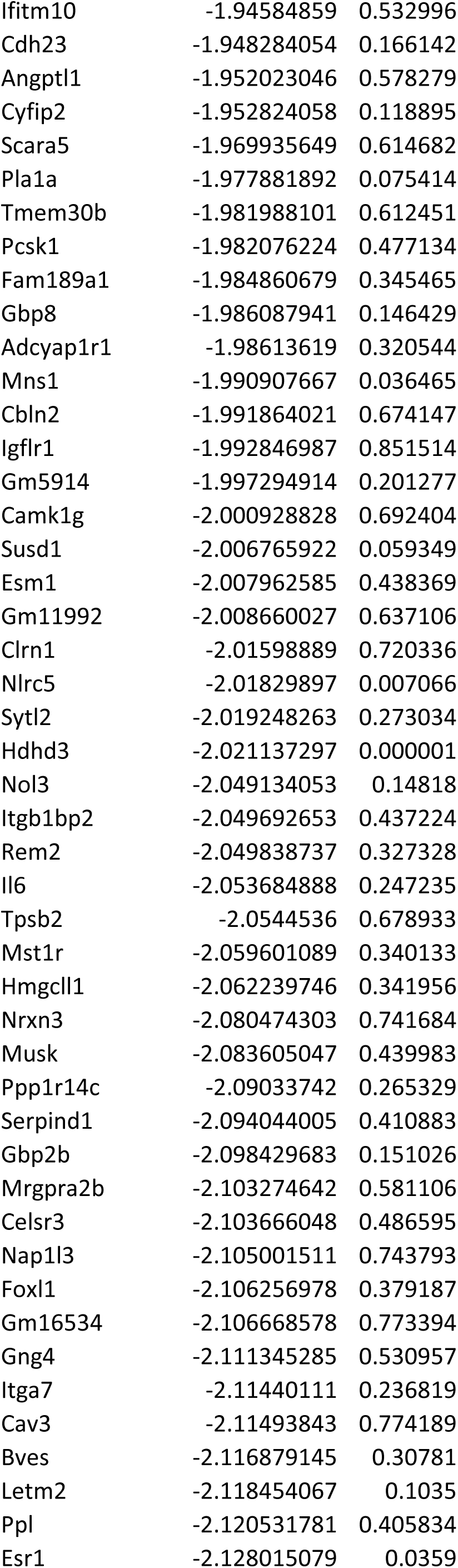
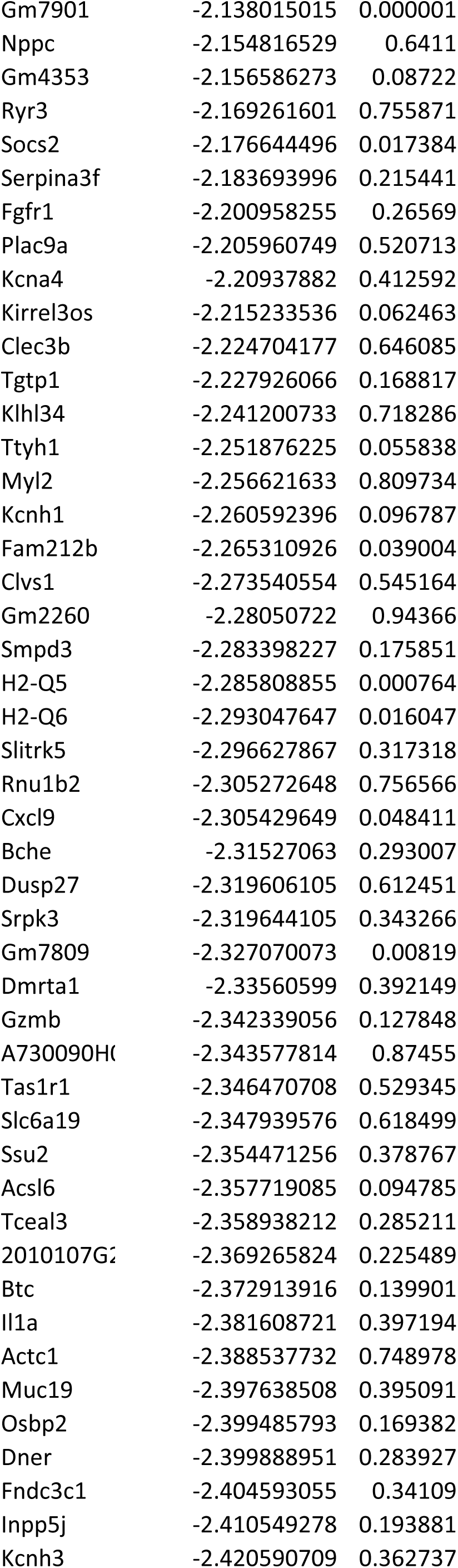
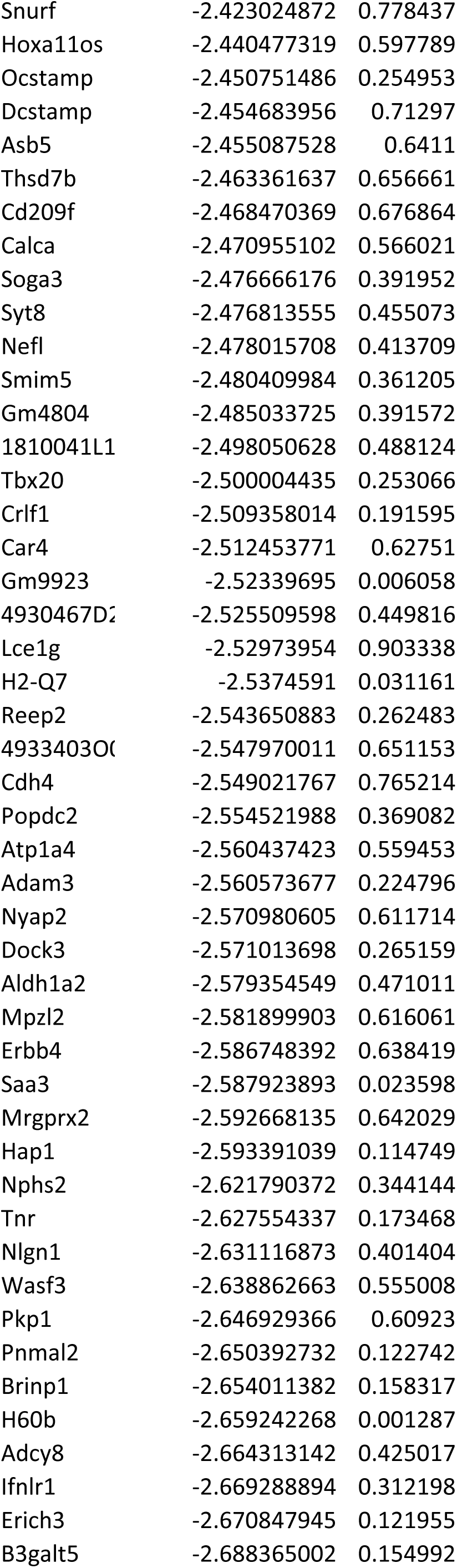
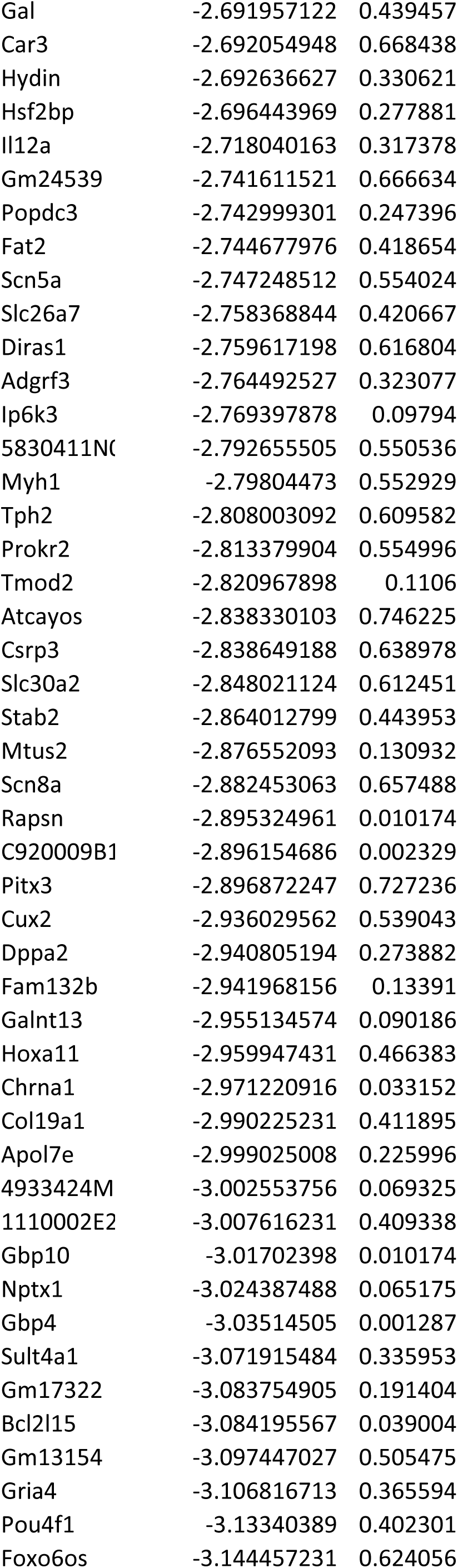
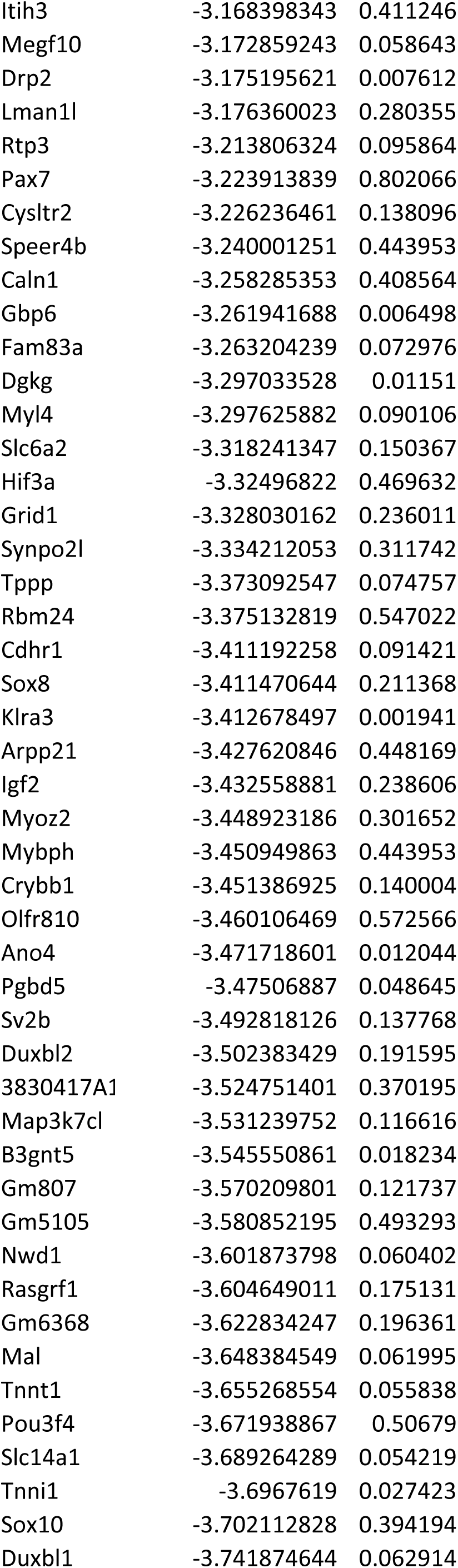
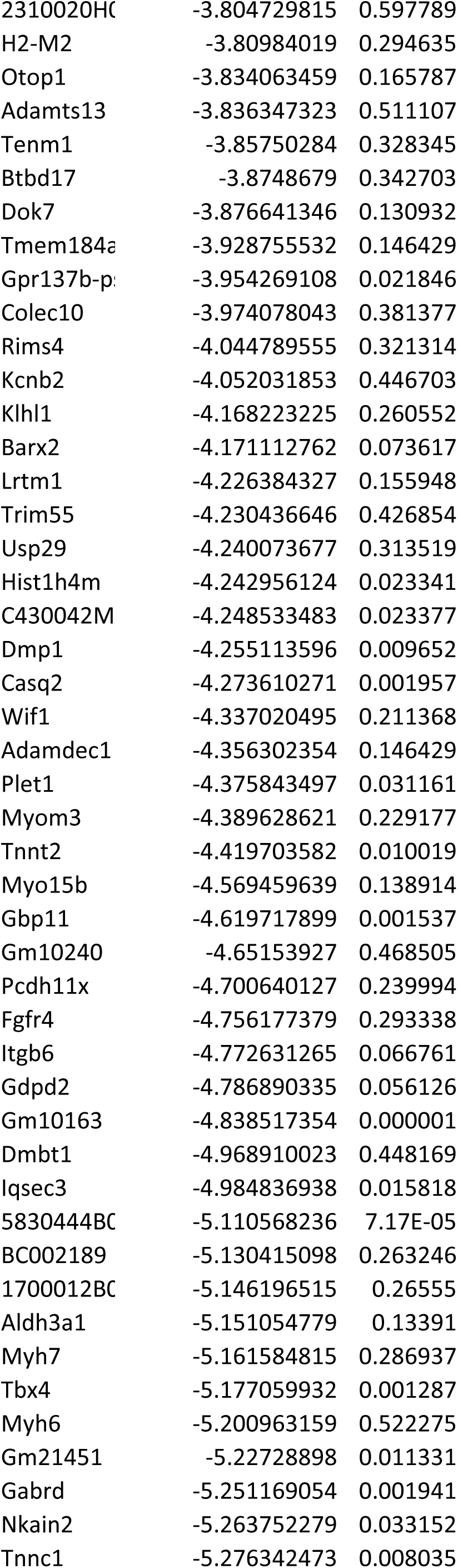
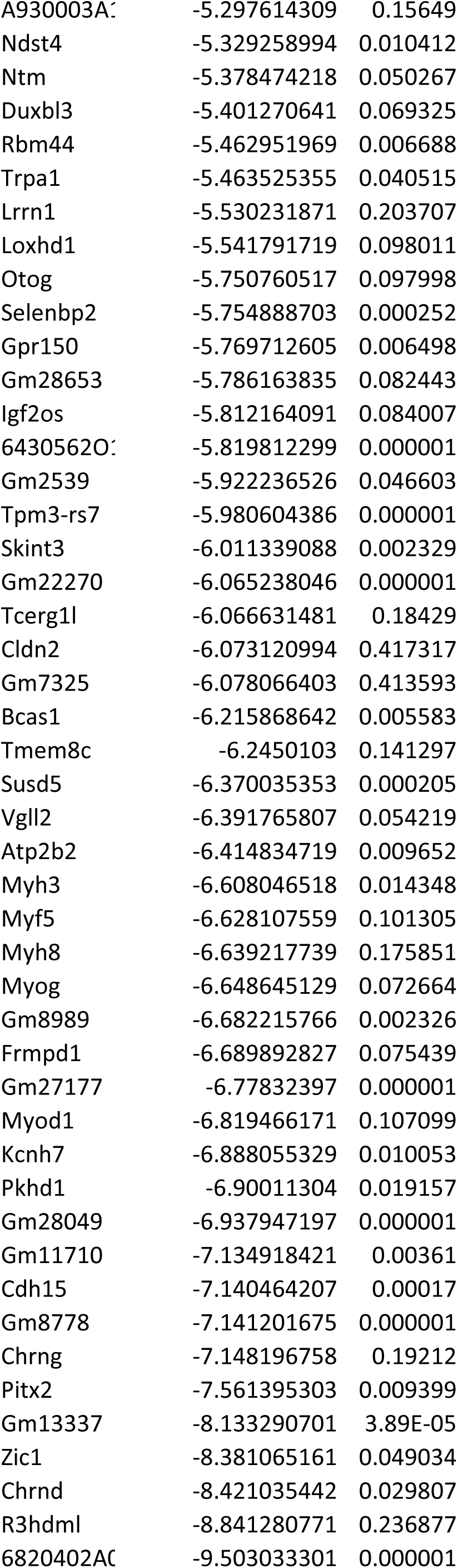
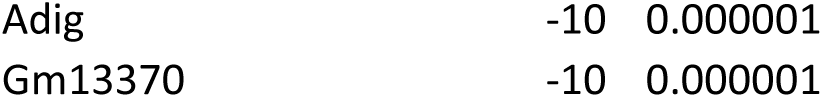
Differentially expressed genes from tumors arising in *2P*, *2PW*, *2PY*, and *2PWY* mice by total RNA-Seq.

